# Influence of Primary Coordination Sphere on Anion Rebound Selectivity in Nonheme Fe Enzyme-Catalyzed C(sp^3^)–H Functionalization: A Comparative Experimental and Computational Study of EgtB and ACCO

**DOI:** 10.64898/2026.07.10.737789

**Authors:** Liu-Peng Zhao, Rui Guo, Binh Khanh Mai, Heyu Chen, Peng Liu, Yang Yang

## Abstract

Developing enzymatic mechanisms for C–F bond formation remains a long-standing challenge. Here, we repurposed the biosynthetic nonheme Fe enzyme EgtB, which features a three-histidine facial triad, to catalyze C(sp^3^)–H fluorination reactions. Directed evolution of EgtB afforded two new-to-nature fluorine atom transferases with opposite enantiopreference, EgtB_CHF1_ and EgtB_CHF2_, with up to 28-fold improved total activity. In contrast to our previously evolved nonheme Fe fluorine atom transfer biocatalyst ACCO_CHF_, which contains a two-histidine-one-carboxylate facial triad, the evolved EgtB_CHF_ variants displayed unexpected hydroxylation activity. ^18^O-labeling experiments showed that the hydroxy group originated from water rather than residual O_2_. Computational studies suggested that the three-histidine-supported Fe(III) center exhibits enhanced Lewis acidity compared to the two-histidine-one-carboxylate system, allowing deprotonation of Fe(III)-bound water to form a Fe(III)–OH species to catalyze radical hydroxylation. Primary coordination-sphere mutagenesis in EgtB and ACCO further supported the critical role of Fe coordination chemistry in controlling radical rebound reactivity and selectivity. Computational studies revealed that Fe coordination chemistry strongly influences both fluorine atom abstraction and radical rebound, with the intrinsic C–X (X = F, OH, and N_3_) bond forming radical rebound preference following the order N_3_ > OH > F. Furthermore, multivariate linear regression analysis revealed that fluorine atom abstraction is primarily governed by the intrinsic Fe–F bond strength, whereas fluorine rebound is predominantly controlled by the electronic structure of the Fe(III) intermediate. Together, these findings provide mechanistic insights into nonheme Fe enzymology and reprogramming toward selective radical rebound reactions, including challenging C–H fluorination.

**Table of Contents (TOC):** 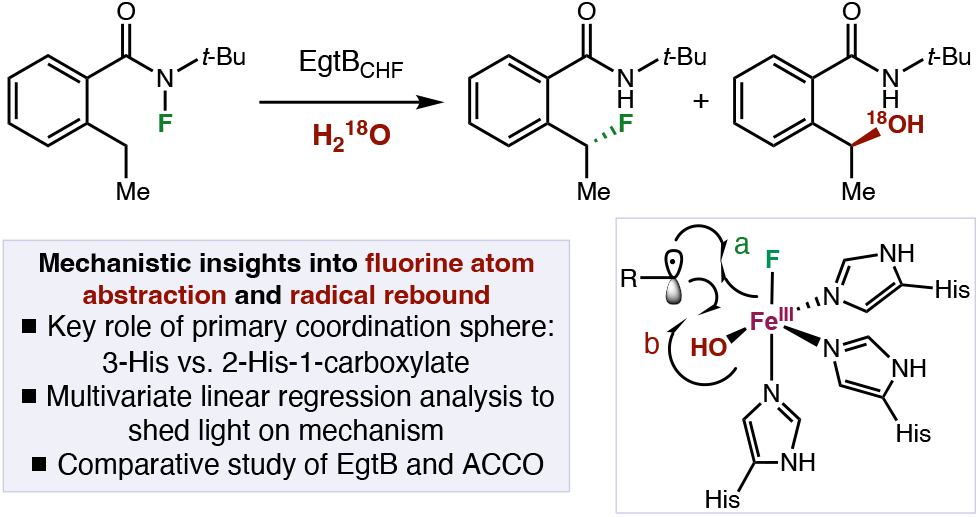

## Introduction

Organofluorine compounds exhibit unique physicochemical and biological properties, including enhanced metabolic stability, improved lipophilicity, and fine-tuned molecular conformation, making them indispensable structural elements in pharmaceuticals,^1^ agrochemicals,^2^ and functional materials.^3^ Due to their broad utility, the stereoselective incorporation of fluorine into organic molecules has been extensively studied by organic chemists over the past decade, leading to the development of an array of synthetically useful asymmetric fluorination methodologies.^4^ In contrast, enzymatic C– F bond formation is exceedingly rare in nature.^5^ To date, 5′-fluoro-5′-deoxyadenosine fluorinase remains the only known natural enzyme capable of catalyzing biological fluorination operating via an S_N_2 mechanism to convert *S*-adenosylmethionine (SAM) into 5′-fluoro-5′-deoxyadenosine (5’-FDA) and enabling the biosynthesis of downstream organofluorine natural products (Scheme 1(A)).^6^ Consequently, the scarcity of natural fluorination enzymes and mechanisms has rendered the design and development of unnatural enzymatic fluorination a long-standing challenge with significant implications for fluorine enzymology,^6^ biocatalysis^7^ and synthetic biology.^8^

In this context, as an important family of nonheme Fe enzymes,^9^ *α*-ketoglutarate (*α*KG)-dependent Fe halogenases^10^ have long captivated enzymologists and enzyme engineers due to their potential to enable C–F bond formation through mechanisms analogous to their native chlorination reactions, which proceed via a radical rebound pathway. Since the pioneering work of Walsh,^11^ a diverse array of *α*KG-dependent nonheme Fe halogenases has been discovered and characterized, establishing a powerful radical halogenation enzymology for C(sp^3^)–H chlorination^12, 13^ and bromination^12h, 12i, 12l, 13a, 13e, 13g^ mediated by nonheme Fe(IV)=O and Fe(III)–X species (Scheme 1(B)). Further biochemical studies demonstrated that these non-heme halogenases could be repurposed to catalyze non-native radical rebound reactions, including azidation^12f, 12l^ and nitration^12f^. More recently, through the use of non-native substrates^14^ and cooperative photobiocatalysis,^15^ our group, the Huang group and other researchers converted a broader range of nonheme Fe enzymes to catalyze fluorination,^14b, 14c^ azidation,^15–16^ thiocyanation,^15–16^ and isocyanation^13e, 15b^ via analogous radical rebound mechanisms. Despite these advances, reprogramming nonheme Fe halogenases as fluorinases has remained a daunting challenge.^17^ This difficulty has been attributed to the high hydration energy of fluoride,^18^ and the challenge of properly positioning the carbon-centered radical to favor fluorine rebound over competing hydroxylation^19^ pathways.

To address these limitations, inspired by our previous work on P450 bromine atom transferases,^20^ we envisioned that leveraging a similar fluorine atom transfer mechanism using non-native *N*-fluoroamide substrates could bypass the generation of Fe(IV)=O intermediates, thus enabling direct interrogation of the ability of nonheme Fe enzymes to catalyze C–F bond formation via a radical rebound mechanism. In 2024, our group^14c^ and the Huang group^14b^ contemporaneously reported biocatalytic enantioselective C(sp^3^)–H fluorination reactions using two functionally distinct nonheme Fe enzymes, including 1-amino-cyclopropane-1-carboxylic acid oxidase (ACCO)^21^ and (*S*)-2-hydroxypropylphosphonate epoxidase (HPPE),^22^ respectively. Mechanistically, these transformations proceed through a fluorine atom abstraction of the *N*-fluoroamide (**1**) with the ferrous nonheme Fe center, leading to a ferric fluoride (**II**) and a nitrogen-centered radical (**III**) (Scheme 1(C)). Subsequent intramolecular 1,5-hydrogen atom transfer (HAT) of III gives rise to a carbon-centered radical **IV**, which upon a fluorine atom rebound with the non-heme Fe(III)–F intermediate affords the fluorine atom transposition product (**2**) and regenerates the ferrous nonheme enzyme catalyst. Overall, these fluorine atom transfer enzymes provided a new biocatalytic strategy for stereocontrolled C–F bond formation via a C–H functionalization logic. Additionally, elegant studies from the Huang group showed that in the presence of exogenous azide ion, the same *N*-fluoroamide substrate could be selectively transformed into the corresponding C–H azidation product,^14a^ indicating divergent anion rebound^23^ pathways could be achieved in nonheme Fe enzyme-catalyzed radical C–H functionalization reactions.

**Scheme 1.**
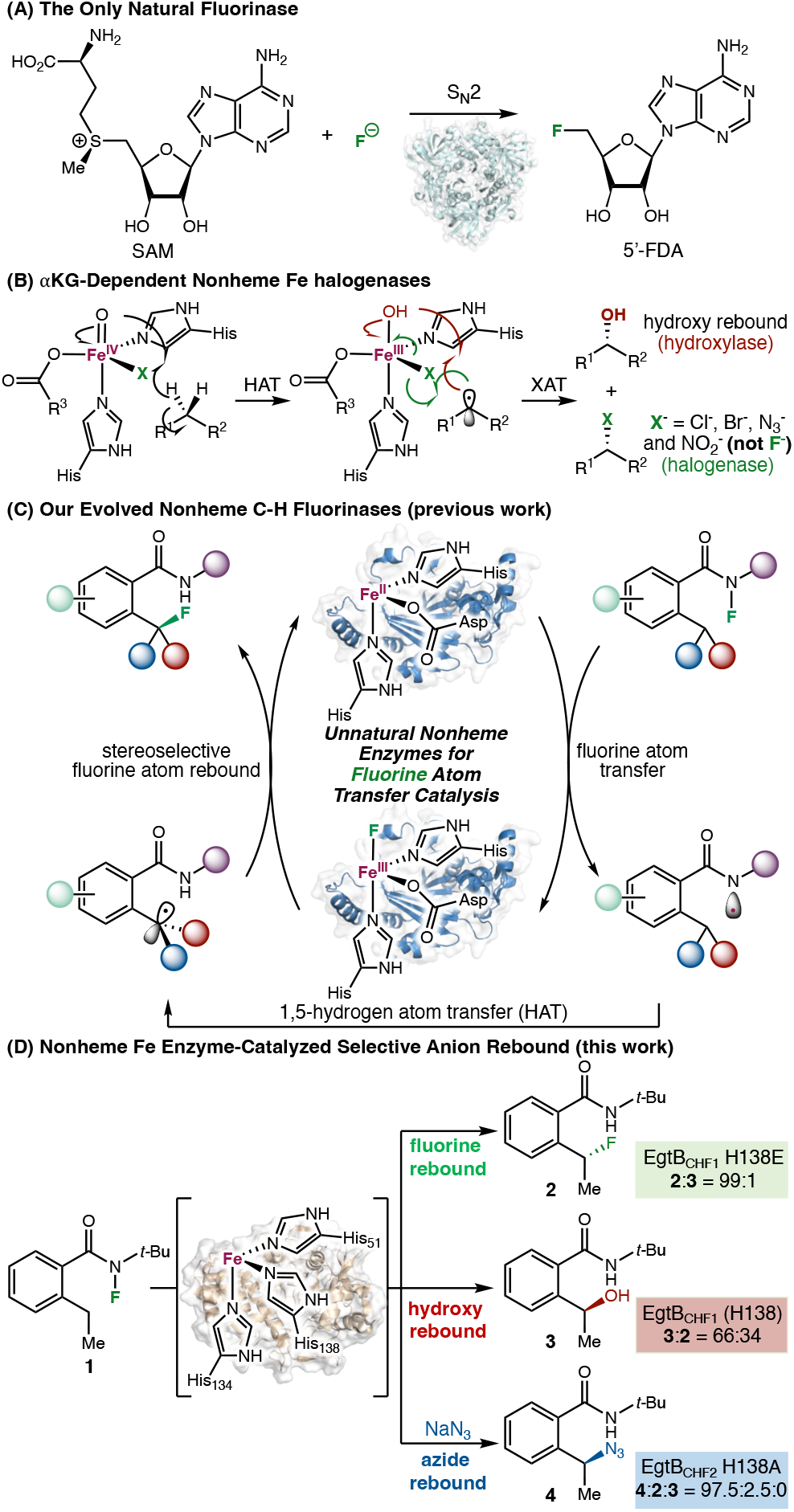
Nonheme Fe Enzymes for C–H Fluorination via a Fluorine Atom Transfer Mechanism. ^***a***^HAT: hydrogen atom transfer. XAT: halogen atom transfer. His: histidine (H); Asp: aspartate (D).

Despite these advances, the mechanism and origin of radical rebound anion selectivity in nonheme Fe-catalyzed C–H functionalization reactions remain to be elucidated. In particular, the lack of an in-depth understanding of nonheme Fe enzyme structure-activity relationships, including the impact of primary coordination sphere and active-site environment, has hampered the rational selection and rapid engineering of nonheme biocatalysts for radical-mediated C–H functionalization in a highly selective fashion. In this Article, combining experimental and computational approaches, we present a comparative investigation of two evolved nonheme Fe enzymes EgtB_CHF_ and ACCO_CHF_ featuring a three-histidine and a two-histidine-one-carboxylate facial triad, respectively, in diverse radical C– H functionalization via a rebound mechanism. First, upon the survey of an in-house collection of nonheme Fe enzymes with diverse coordination chemistry and further directed evolution, we identified a promiscuous enzyme EgtB from *Mycolicibacterium thermoresistibile* (*Mth*EgtB),^24^ a thermophilic sulfoxide synthase involved in ergothioneine biosynthesis with a three-histidine facial triad, that catalyzed the fluorine atom transfer reaction. Directed evolution of *Mth*EgtB led to a nonheme Fe biocatalysts EgtB_CHF1_ and EgtB_CHF2_ with substantial fluorination activity, but also unexpected hydroxylation activity. ^18^O isotope labeling experiments revealed that the hydroxy group was derived from the aqueous buffer (H_2_ ^18^O). To account for this unexpected finding, we proposed and computationally validated that the substantially higher Lewis acidity of the three-histidine Fe center in EgtB compared to the two-histidine-one-carboxylate Fe in ACCO allowed for the facile deprotonation of Fe(III)-bound H_2_O, giving rise to a Fe(III)–OH intermediate responsible for the promiscuous hydroxylation reaction. Systematic variation of Fe-binding residues in EgtB and ACCO confirmed the impact of Fe coordination chemistry on Lewis acidity and radical rebound anion selectivity, allowing the production of fluorination and azidation products with enhanced chemoselectivity (Scheme 1(D)). Furthermore, computational studies combining density functional theory (DFT), molecular dynamics (MD) and quantum mechanics/molecular mechanics (QM/MM) calculations were carried out to elucidate the mechanism of C–H fluorination and to evaluate how the primary coordination sphere and active-site environment facilitate this transformation. In addition, predictive models for fluorine atom abstraction and C–F bond forming radical rebound were developed using multivariate linear regression analysis, revealing how modulation of the primary coordination sphere governs reactivity and selectivity in nonheme Fe enzyme-catalyzed fluorine atom transfer.

## Results and discussion

### Discovery of enzymatic C–H fluorination and azidation activities

At the outset of this study, we evaluated our recently expanded collection of ca. 200 metalloproteins and their variants, including diverse nonheme Fe enzymes, for fluorine atom transfer using *N*-fluoroamide substrate **1** via high throughput experimentation using 24-well plates. All potential hits were then validated by nonheme Fe enzyme expression in 125 mL Erlenmeyer flasks, and representative results are summarized in Table 1 (entries 1–9). It was found that Fe- and *α*KG-dependent C–H halogenases, including SadA D157G (entry 1),^12i^ WelO5 (entry 2),^12e, 12h^ and BesD (entry 3)^12k, 12l, 12p^ were ineffective in mediating this C–H fluorination. Interestingly, all the nonheme Fe enzymes displaying the desired fluorine atom transfer activity were not natural C–H halogenases. Among these, quercetin 2,3-dioxygenase from *Bacillus subtilis* (*Bsu*QueD)^25^ with a three-histidine-one-carboxylate tetrad afforded **2** in 0.2% yield and 71:29 e.r. (entry 4). EvdO2 from *Micromonospora carbonacea*,^26^ an oxygenase involved in everninomicin biosynthesis, afforded **2** in marginal activity (0.2% yield, 36:64 e.r., entry 5). Furthermore, 1-aminocyclopropane-1-carboxylic acid oxidase from *Petunia hybrida* (*Phy*ACCO)^21^ exhibited excellent enantioselectivity in C–H fluorination (0.2% yield, 90:10 e.r., entry 6). Likewise, isopenicillin N synthase (IPNS)^27^ from *Emericella nidulans*, whose native function is to catalyze the oxidative cyclization forming isopenicillin N and recently engineered by our lab as a highly active biocatalyst for 1,3-nitrogen migration,^28^ provided the desired fluorination product 2 in 0.2% yield and 90:10 e.r. (entry 7). Similarly, metapyrocatechase from *Pseudomonas putida* (*Pp*MPC)^29^ (entry 8), an extradiol dioxygenase we recently engineered for photobiocatalytic azidation, thiocyanation and isocyanation,^15b^ was ineffective in this fluorine atom transfer reaction. In contrast, EgtB from *Mycolicibacterium thermoresistibile* (*Mth*EgtB), a thermophilic sulfoxide synthase involved in ergothioneine biosynthesis,^24^ furnished fluorinated **2** in 3.5% yield and 42:58 e.r. (entry 9). Among all the nonheme Fe enzymes exhibiting fluorine atom transferase activity, EgtB was the only one featuring a three-histidine facial triad (**Figure 1**). Collectively, these results demonstrated that a diverse range of Fe coordination chemistry, including a three-histidine and a two-histidine-one-carboxylate triad and a three-histidine-one-carboxylate tetrad, could support the long sought-after C–F bond formation activity.

**Table 1.**
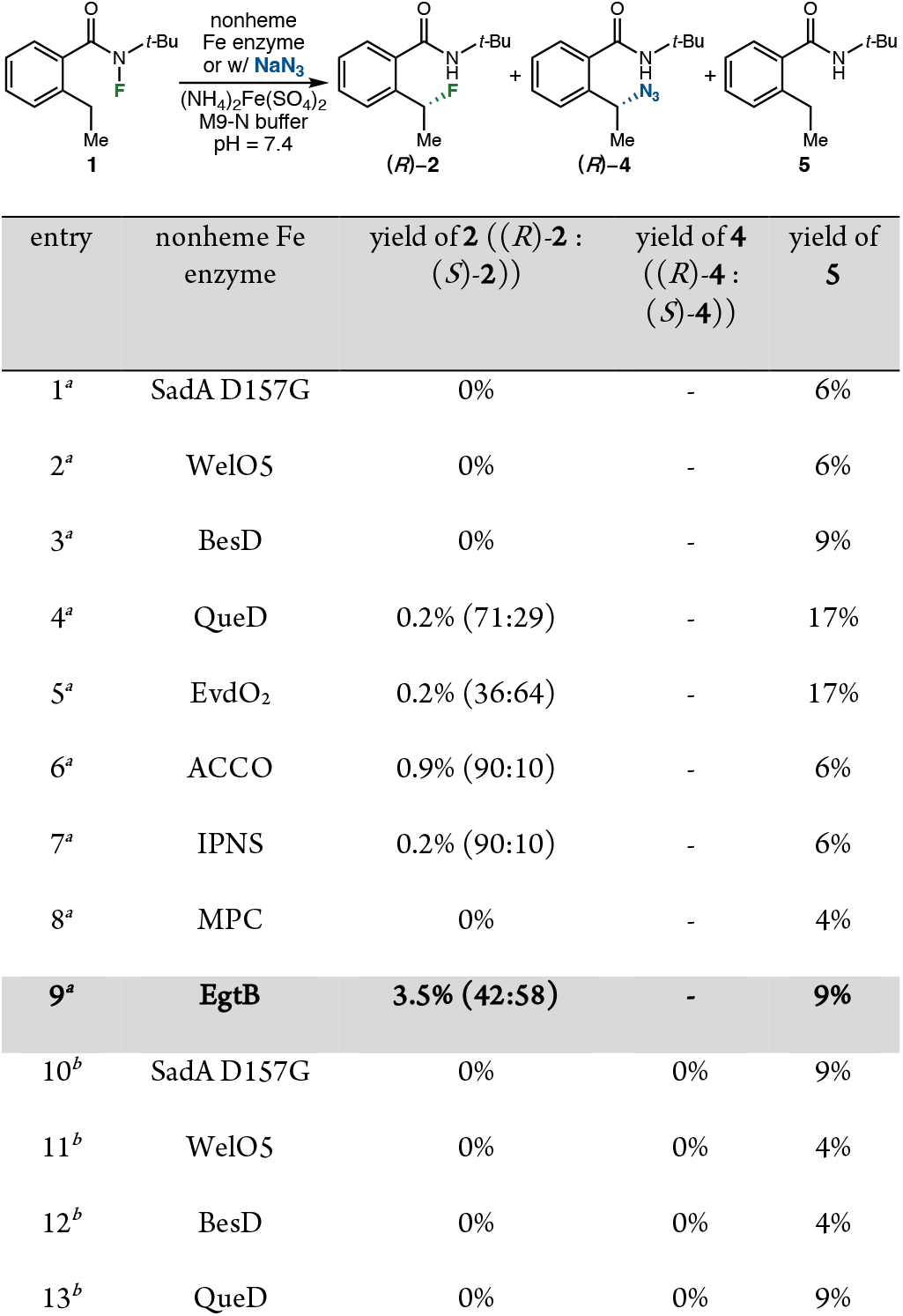
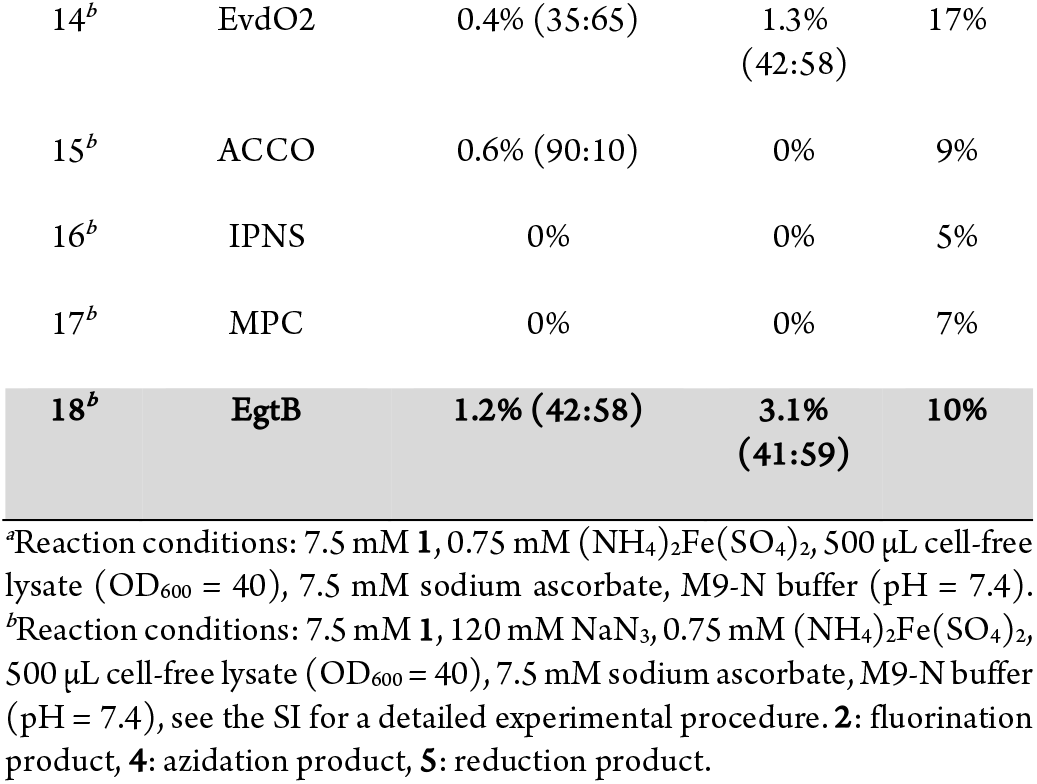
Evaluation of Nonheme Fe Enzymes for Promiscuous Radical Rebound Activities Including Fluorination and Azidation.

**Figure 1.**
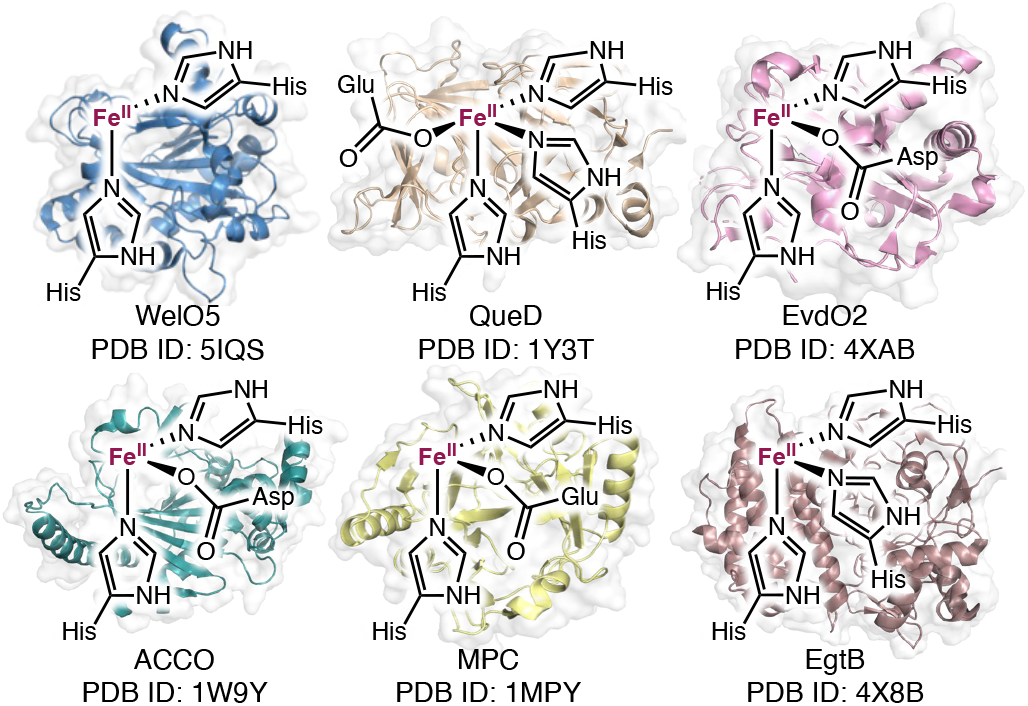
Representative nonheme Fe enzyme structures and PDB IDs screened in this reaction.

To better understand the radical rebound activity and anion selectivity of these nonheme Fe enzymes in the presence of exogenous anion, we re-evaluated their C–H functionalization activity in the presence of added NaN_3_ (Table 1, entries 10–18). It was found that *α*KG-dependent halogenases SadA D157G (entry 10), WelO5 (entry 11) and BesD (entry 12) did not promote either fluorination or azidation of **1** under these reaction conditions. Interestingly, many nonheme Fe enzymes showing activity in fluorine atom transfer, including QueD (entry 13), ACCO (entry 15) and IPNS (entry 16), did not promote azide rebound in the presence of exogenous NaN_3_. Among these, the addition of NaN_3_ shutdown the fluorine transfer activity of QueD (entry 13) and IPNS (entry 16), but preserved the fluorination activity of ACCO with identical enantioselectivity (90:10 e.r., entry 15). In contrast, EvdO2 afforded azidation product **4** in 1.3% yield (42:58 e.r.) and fluorination product **2** in 0.4% yield (35:65 e.r.) (entry 14). EgtB provided **4** and **2** in 3.1% yield (41:59 e.r.) and 1.2% yield (42:58 e.r.) (entry 18). Together, these results reveal the contrasting radical rebound activity and anion selectivity of nonheme Fe enzymes in C–H (pseudo)halogenation reactions.

### Directed evolution of fluorine atom transferase EgtB_CHF_

As EgtB was the only three-histidine nonheme Fe enzyme displaying C–F bond forming activity, further improving and understanding the radical rebound activity of EgtB would offer insights into non-heme Fe enzymology, particularly in the context of radical-mediated C–H fluorination and (pseudo)halogenation. Thus, we carried out directed evolution of *Mth*EgtB to improve its fluorination activity and enantioselectivity in this C(sp^3^)–H fluorination reaction (Table 2). Guided by the crystal structure of *Mth*EgtB (**Figure 2**, PDB ID: 4X8E)^24b^ and molecular docking studies, we performed site-saturation mutagenesis (SSM) and screening by targeting active site residues in proximity to the nonheme Fe center. For each round of engineering, SSM libraries of *Mth*EgtB were generated using the 22c-trick method,^30^ and 88 clones were evaluated in 24- or 96-well plates in the form of whole-cell biocatalysts. Enhancement of total activity and/or enantioselectivity was used as the selection criteria for directed evolution. Initially, beneficial mutations Y377W, W415R and R87D were identified to improve the enantioselectivity or fluorination activity, leading to an improved fluorinase variant *Mth*EgtB Y377W W415R R87D, furnishing the fluorinated product **2** with a total turnover number (TTN) of 26 and 34:66 e.r. Among these targeted active site residues, Y377 was previously established as an essential residue in the native function of EgtB for C–S bond formation and sulfoxidation.^24b, 24c^ Thus, the identification of Y377W as a beneficial mutation furnishing improved enantioselectivity and similar fluorination activity indicated a departure from the mechanism in native enzymatic oxidative C–S coupling. Additionally, as we improved the total activity of *Mth*EgtB toward C–H fluorination, an unexpected C–H hydroxylation product (**3**) was also discovered and further confirmed and characterized by HPLC and NMR analyses. Using *Mth*EgtB Y377W W415R R87D as the biocatalyst, the absolute stereochemistry of the major enantiomer of C–H hydroxylation product **3** was found to be the same as C–H fluorination product **2** (i.e., (*S*)-). This stereochemistry was established by comparing the enzymatic C–H hydroxylation product with enantioenriched sample independently synthesized using the Corey-Bakshi-Shibata reduction^31^ (see the SI for details).

**Table 2.**
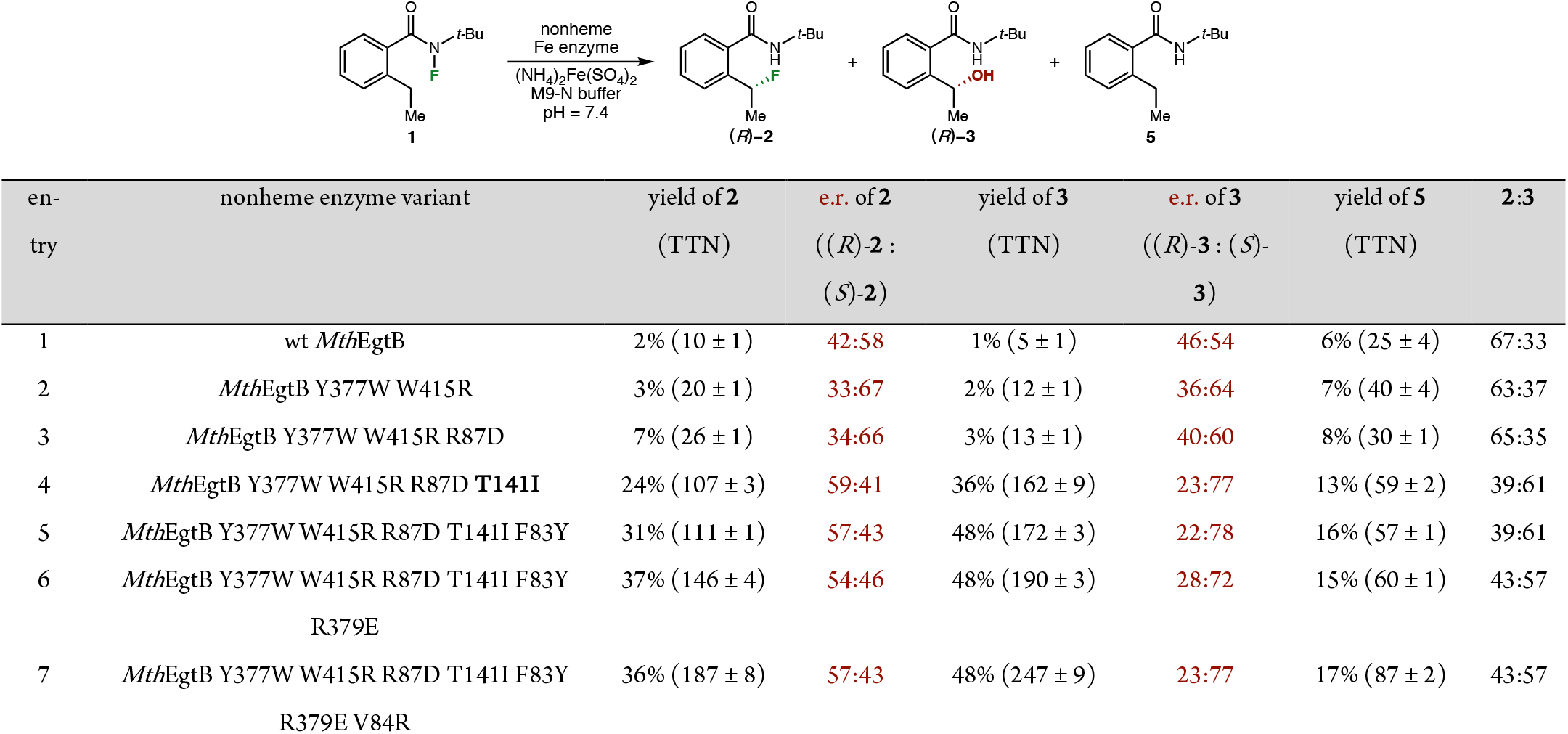
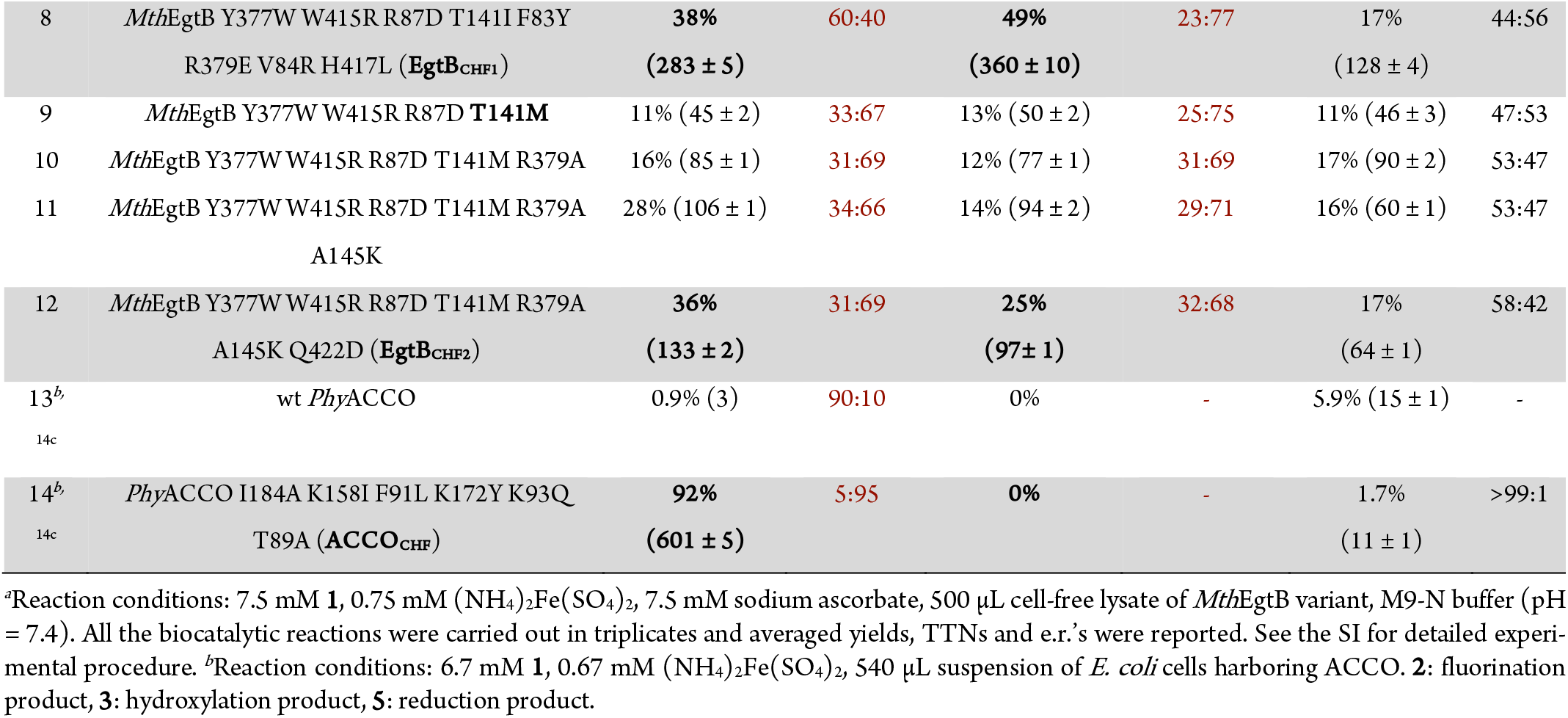
Directed Evolution of *Mth*EgtB as New-to-Nature Fluorine Atom Transferase^*a*^.

| en-try | nonheme enzyme variant | yield of **2**<br>(TTN) | e.r. of **2**<br>(*(R)*-**2** :<br>*(S)*-**2**) | yield of **3**<br>(TTN) | e.r. of **3**<br>(*(R)*-**3** : *(S)*-**3**) | yield of **5**<br>(TTN) | **2**:**3** |
| --- | --- | --- | --- | --- | --- | --- | --- |
| 1 | wt *MthEgtB* | 2% (10 ± 1) | 42:58 | 1% (5 ± 1) | 46:54 | 6% (25 ± 4) | 67:33 |
| 2 | *MthEgtB* Y377W W415R | 3% (20 ± 1) | 33:67 | 2% (12 ± 1) | 36:64 | 7% (40 ± 4) | 63:37 |
| 3 | *MthEgtB* Y377W W415R R87D | 7% (26 ± 1) | 34:66 | 3% (13 ± 1) | 40:60 | 8% (30 ± 1) | 65:35 |
| 4 | *MthEgtB* Y377W W415R R87D **T141I** | 24% (107 ± 3) | 59:41 | 36% (162 ± 9) | 23:77 | 13% (59 ± 2) | 39:61 |
| 5 | *MthEgtB* Y377W W415R R87D T141I F83Y | 31% (111 ± 1) | 57:43 | 48% (172 ± 3) | 22:78 | 16% (57 ± 1) | 39:61 |
| 6 | *MthEgtB* Y377W W415R R87D T141I F83Y<br>R379E | 37% (146 ± 4) | 54:46 | 48% (190 ± 3) | 28:72 | 15% (60 ± 1) | 43:57 |
| 7 | *MthEgtB* Y377W W415R R87D T141I F83Y<br>R379E V84R | 36% (187 ± 8) | 57:43 | 48% (247 ± 9) | 23:77 | 17% (87 ± 2) | 43:57 |
| 8 | *MthEgtB* Y377W W415R R87D T141I F83Y<br>R379E V84R H417L (**EgtB<sub>CHF1</sub>**) | 38%<br>(283 ± 5) | 60:40 | 49%<br>(360 ± 10) | 23:77 | 17%<br>(128 ± 4) | 44:56 |
| 9 | *MthEgtB* Y377W W415R R87D **T141M** | 11% (45 ± 2) | 33:67 | 13% (50 ± 2) | 25:75 | 11% (46 ± 3) | 47:53 |
| 10 | *MthEgtB* Y377W W415R R87D T141M R379A | 16% (85 ± 1) | 31:69 | 12% (77 ± 1) | 31:69 | 17% (90 ± 2) | 53:47 |
| 11 | *MthEgtB* Y377W W415R R87D T141M R379A<br>A145K | 28% (106 ± 1) | 34:66 | 14% (94 ± 2) | 29:71 | 16% (60 ± 1) | 53:47 |
| 12 | *MthEgtB* Y377W W415R R87D T141M R379A<br>A145K Q422D (**EgtB<sub>CHF2</sub>**) | 36%<br>(133 ± 2) | 31:69 | 25%<br>(97 ± 1) | 32:68 | 17%<br>(64 ± 1) | 58:42 |
| 13<sup>b</sup><br>14c | wt *PhyACCO* | 0.9% (3) | 90:10 | 0% | - | 5.9% (15 ± 1) | - |
| 14<sup>b</sup><br>14c | *PhyACCO* I184A K158I F91L K172Y K93Q<br>T89A (**ACCO<sub>CHF</sub>**) | 92%<br>(601 ± 5) | 5:95 | 0% | - | 1.7%<br>(11 ± 1) | >99:1 |

<sup>a</sup>Reaction conditions: 7.5 mM **1**, 0.75 mM (NH<sub>4</sub>)<sub>2</sub>Fe(SO<sub>4</sub>)<sub>2</sub>, 7.5 mM sodium ascorbate, 500 μL cell-free lysate of *MthEgtB* variant, M9-N buffer (pH = 7.4). All the biocatalytic reactions were carried out in triplicates and averaged yields, TTNs and e.r.'s were reported. See the SI for detailed experimental procedure. <sup>b</sup>Reaction conditions: 6.7 mM **1**, 0.67 mM (NH<sub>4</sub>)<sub>2</sub>Fe(SO<sub>4</sub>)<sub>2</sub>, 540 μL suspension of *E. coli* cells harboring ACCO. **2**: fluorination product, **3**: hydroxylation product, **5**: reduction product.

**Figure 2.**
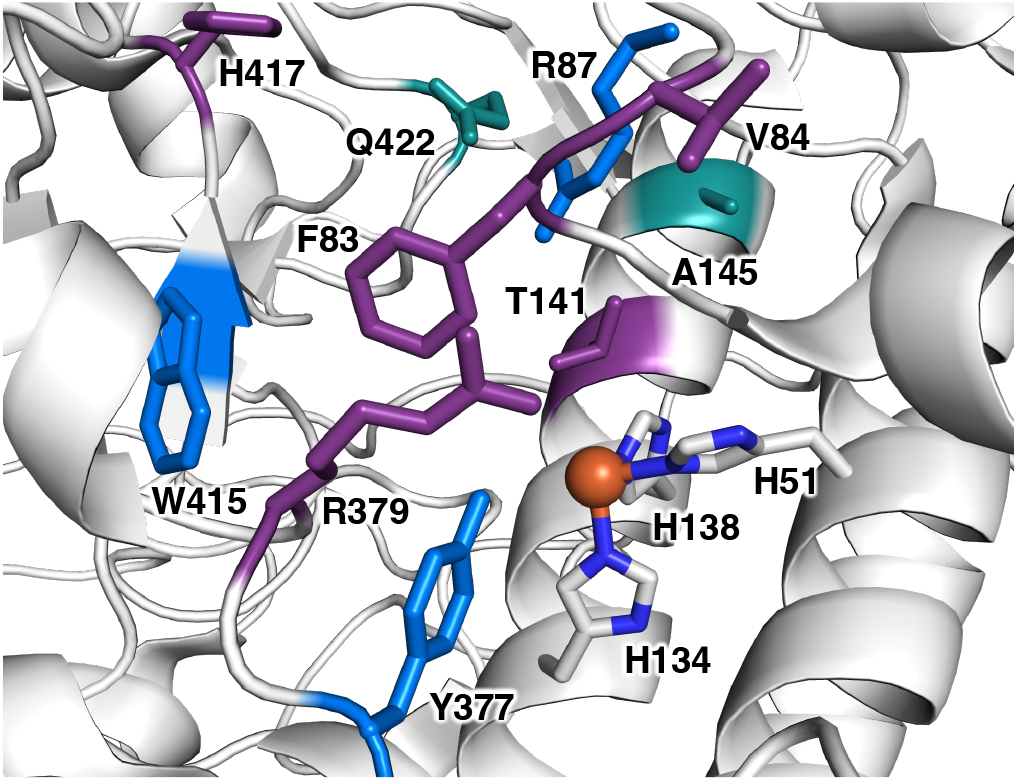
Active-site illustration of *Mth*EgtB (PDB ID: 4X8E) and beneficial mutations identified from directed evolution. Marine residues: beneficial mutations identified before the key T141I/T141M mutation. Magenta residues: beneficial mutations identified in the EgtB_CHF1_ lineage. Deep teal residues: beneficial mutations identified in the EgtB_CHF2_ lineage.

Next, SSM and screening revealed the critical role of T141 proximal to H134 and H138 residing in the same *α*-helix as these Fe-binding residues in modulating the enzyme activity and enantiopreference for C–H fluorination. In particular, the incorporation of the T141I mutation led to a 4.4-fold improvement in TTN for C–H fluorination, along with the reversal of absolute stereochemistry of **2** (59:41 e.r., entry 4). On the other hand, the introduction of T141M resulted in a 1.8-fold enhancement in fluorination activity with almost identical stereoselectivity of **2** (33:67 e.r., entry 9). Both T141I and T141M enhanced the C–H hydroxylation activity, with T141I and T141M providing a 2.0- and 1.4-fold higher TTN for hydroxylation product **3**, respectively.

With two variants EgtB Y377W W415R R87D T141I and EgtB Y377W W415R R87D T141M in hand, further SSM and screening afforded final variants of fluorine transfer enzymes with further improved activities, including EgtB Y377W W415R R87D T141I F83Y R379E V84R H417L (EgtB_CHF1_, CHF = C–H fluorinase, entries 4– 8) and EgtB Y377W W415R R87D T141M R379A A145K Q422D (EgtB_CHF2_, entries 9–12). Under optimized conditions, EgtB_CHF1_ provided C–H fluorination product **2** in (283 ± 5) TTN and 60:40 e.r. (entry 8), while EgtB_CHF2_ delivered **2** in (133 ± 2) TTN and 31:69 e.r. (entry 12). During the engineering of EgtB_CHF1_, F83Y (entry 5), R379E (entry 6) and H417L (entry 8) further improved the fluorination activity. V78R (entry 7) and H417L (entry 8) slightly enhanced fluorination enantioselectivity. Similarly, beneficial mutations R379A (entry 10), A145K (entry 11), and Q422D (entry 12) played an important role in further enhancing the fluorination activity of EgtB_CHF2_.

Throughout this directed evolution campaign, the C–H hydroxylation product **3** was observed along the evolutionary trajectory of both EgtB_CHF1_ and EgtB_CHF2_. As fluorination activity was progressively enhanced, the formation of the hydroxylated product also increased accordingly, with EgtB_CHF1_ affording **3** in 49% yield and 360 TTN, and EgtB_CHF2_ in 25% yield and 97 TTN. In contrast to C–H fluorination, the enantiopreference of C–H hydroxylation remained unchanged across all the evolved variants, with EgtB_CHF1_ and EgtB-_CHF2_ delivering **3** in 77:23 e.r. and 68:32 e.r., respectively, in favor of the (*S*)-enantiomer. These results suggest distinct enantioinduction mechanisms for C–H fluorination and hydroxylation catalyzed by the same enzyme variant. Finally, we note that this C–H hydroxylation activity was not observed with wt ACCO (entry 13) and the engineered ACCO_CHF_ (entry 14), both of which featured a two-histidine-one-carboxylate facial triad.^14c^

### Mechanism studies for EgtB-catalyzed C–H fluorination and hydroxylation

To gain further insight into the mechanism of this biocatalytic fluorination, a series of control experiments was conducted. Product analysis under the standard reaction conditions revealed the absence of the alkene and the lactam derived from carbocation intermediates (see the SI for details). These observations indicated that carbocationic intermediates are likely not involved. In addition, cyclopropyl radical clock experiments using substrate **6** with EgtB_CHF1_ or EgtB_CHF2_ resulted exclusively in the formation of the ring-opened product **8**, while the fluorinated products were not observed (**Figure 3A**, see the SI for details). These results further supported the intermediacy of a benzylic radical species under the biocatalytic conditions. Furthermore, in the presence of 2.0 equiv of TEMPO, the fluorination reaction was effectively suppressed, and the corresponding TEMPO-trapping adduct **9** was observed by LC–MS analysis (**Figure 3B**, see the SI for details). Collectively, these experiments supported the formation of radical intermediates in this biocatalytic transformation and disfavored the involvement of carbocationic intermediates.

**Figure 3.**
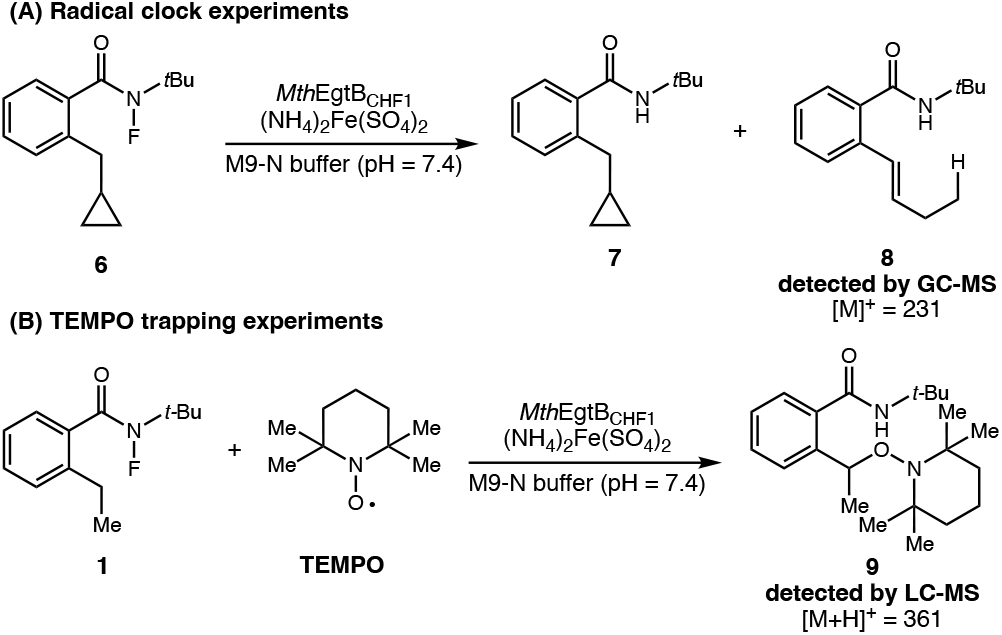
Mechanism studies for EgtB-catalyzed C–H functionalization. (A) Radical clock experiments. Reaction conditions: 7.5 mM **6**, 0.75 mM (NH_4_)_2_Fe(SO_4_)_2_, 7.5 mM sodium ascorbate, 500 μL cell-free lysate of *Mth*EgtB_CHF1_ variant, M9-N buffer (pH = 7.4) see the SI for detailed experimental procedures and GC-MS analysis results. (B) TEMPO trapping experiments. Reaction conditions: 7.5 mM **1**, 15.0 mM TEMPO, 0.75 mM (NH_4_)_2_Fe(SO_4_)_2_, 7.5 mM sodium ascorbate, 500 μL cell-free lysate of *Mth*EgtB_CHF1_ variant, M9-N buffer (pH = 7.4) see the SI for detailed experimental procedures and LC-MS analysis results.

To further understand the mechanism of this EgtB-catalyzed unexpected C–H hydroxylation reaction and the origin of the hydroxy group incorporated into the benzylic position of **3**, we performed isotope labeling experiments using ^18^O-labeled ^18^O_2_ and H_2_ ^18^O. To assess the possibility of C–H hydroxylation via O_2_ trapping of the nascent benzylic radical, we carried out this biotransformation in the presence of isotopically labeled dioxygen (^18^O_2_). When the biocatalytic C–H functionalization reaction was carried out under an atmosphere of ^18^O_2_, no ^18^O-incorporated hydroxylated product **3-**^**18**^**O** was observed by LC-MS analysis. Instead, non-^18^O-incorporated hydroxylation product **3-**^**16**^**O** was still observed under these conditions. This result indicated that the hydroxy group does not come from dioxygen. In contrast, when the biocatalytic reaction was performed in an aqueous buffer prepared from H_2_ ^18^O, substantial ^18^O incorporation into **3** (**3-**^**18**^**O**) was observed. Together, these results show that the hydroxy group in product **3** originates from H_2_O in the reaction buffer (**Figure 4**, see SI for detail).

**Figure 4.**
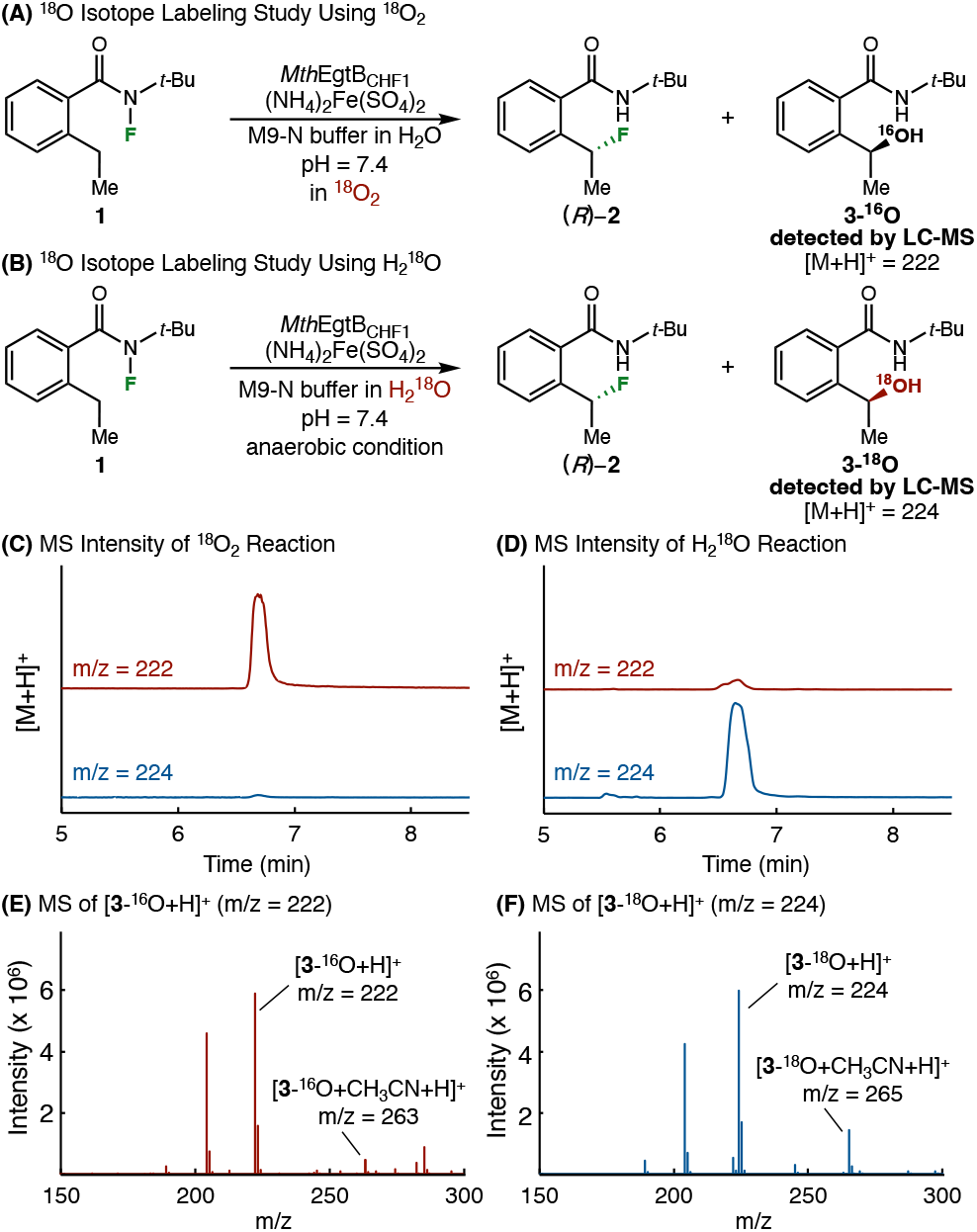
^18^O isotope labeling studies. (A) Isotope labeling experiment using ^18^O-labeled ^18^O_2_. (B) Isotope labeling experiment using ^18^O-labeled H_2_ ^18^O. Reaction conditions: 7.5 mM **1**, 0.75 mM (NH_4_)_2_Fe(SO_4_)_2_, 7.5 mM sodium ascorbate, 500 μL cell-free lysate of *Mth*EgtB_CHF1_ variant, M9-N buffer (pH = 7.4), see the SI for detailed experimental procedures and LC-MS analysis results. (C) Extracted ion chromatogram of the reaction in the presence of ^18^O_2_, monitoring *m*/*z* = 222 and 224. (D) Extracted ion chromatogram of the reaction using H_2_ ^18^O, monitoring *m*/*z* = 222 and 224. (E) Extracted mass spectrometry at peak retention time corresponding to **3-^16^O**. (F) Extracted mass spectrometry at peak retention time corresponding to **3-^18^O**.

To account for this unexpected hydroxylation activity, we propose that, unlike ACCO, nonheme Fe enzyme EgtB generates a Fe(III)–OH under the reaction conditions, thereby allowing the C– H hydroxylation product to form via a radical rebound mechanism involving this enzymatic Fe(III)–OH intermediate. Due to the higher Lewis acidity of EgtB’s Fe(III) center, supported by a three-histidine triad, compared to ACCO’s Fe(III) supported by a two-histidine-one-carboxylate triad, we hypothesized that the water molecule bound to EgtB’s Fe(III) is highly acidic and becomes readily deprotonated to furnish a Fe(III)–OH species.

To evaluate this proposal, we computed aqueous p*K*_a_ values of representative water-bound Fe(II) and Fe(III) complexes using both truncated and theozyme models with the scaled solvent-accessible surface (SMD_sSAS_) approach described by Smith *et al*.^32^ at the B3LYP-D3(BJ)/def2-TZVP/SMD_sSAS_(H_2_O)//B3LYP-D3(BJ)/6-31G(d)–SDD(Fe) level of theory (see SI for more details). For models representing EgtB with a three-histidine facial triad, Im_3_Fe(II)(H_2_O)_2_ and Im_3_Fe(III)(F)(H_2_O)_2_ have computed p*K*_a_ values of 13.6 and 8.7, respectively (**Scheme 2**). These results suggest that the Fe(II)-bound water remains protonated, whereas the Fe(III)-bound water in EgtB could be partially deprotonated under the experimental conditions (pH = 7.4), generating an Fe(III)–OH species (**12-A**) and giving rise to hydroxy rebound products (see **Figures S17-S18** for Fe-bound water deprotonation analysis). In contrast, in ferric ACCO, which contains a two-histidine–one-carboxylate facial triad, both Fe(II)- and Fe(III)-bound waters are computationally found to remain protonated (p*K*_a_ = 13.9 and 10.6, respectively, **Figure S15**). Thus, no Fe(III)–OH species forms with ACCO, precluding the formation of C–H hydroxylation products. Collectively, these results suggest that the distinct primary coordination environments of ACCO, featuring a two-His-one-carboxylate facial triad, and EgtB, featuring a three-Histidine facial triad, differentially modulate the formation of Fe(III)–OH species and the resulting radical rebound selectivity.

**Scheme 2.**
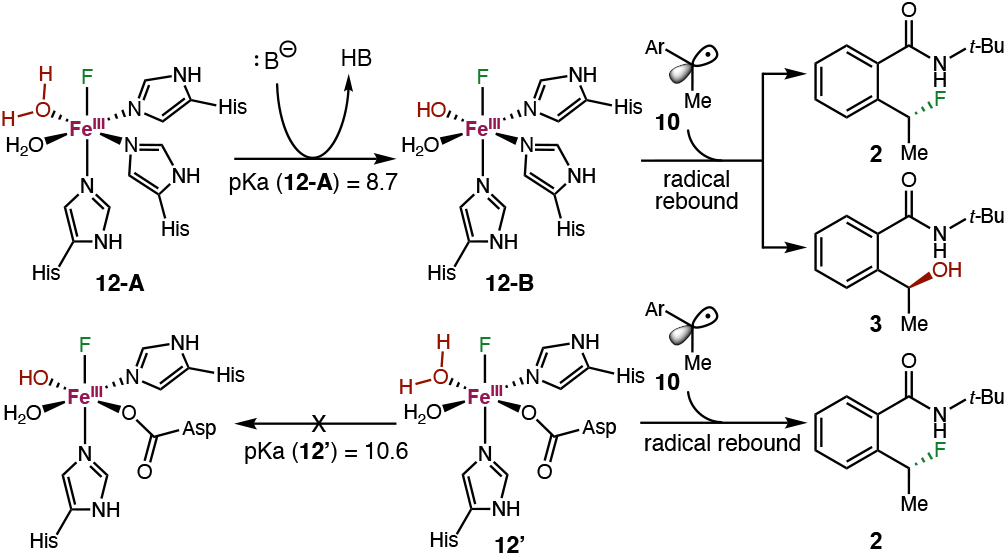
Influence of Nonheme Fe Coordination Chemistry on Fe(III) Lewis Acidity and Protonation State of Fe(III)-Bound Water^***a***^. ^***a***^p*K*_a_ values were calculated using density functional theory at the B3LYP-D3(BJ)/def2-TZVP/SMD_sSAS_(H_2_O)//B3LYP-D3(BJ)/6-31G(d)–SDD(Fe) level of theory.

To further probe the effects of Fe-bound-water deprotonation on hydroxylation activity, EgtB_CHF1_, EgtB_CHF2_ and ACCO_CHF_-catalyzed C–H functionalization reactions were performed over a pH range of 6.5–9.0. For EgtB_CHF1_, increasing the buffer pH led to an increase in hydroxylation activity, with the yield of hydroxylation product **3** increasing from 36% at pH 6.5 to 54% at pH 9.0, while fluorination activity remained largely unchanged (Table 3). These changes corresponded to an increase of the hydroxylation-to-fluorination ratio from nearly 1:1 to 2:1. A similar pH-dependent trend was also observed for MthEgtB_CHF2_, where higher pH led to increased hydroxylation activity and selectivity (see Table S14 for details). Notably, under the optimized condition (pH = 9.0), EgtB_CHF1_ afforded the hydroxylation product with 56% yield and 80:20 e.r.. In contrast to the engineered EgtB_CHF_ variants, no hydroxy rebound product was observed for ACCO_CHF_, even under basic conditions (Table S15). Additionally, similar pH-dependent trends were also observed under azidation conditions, with higher pH values consistently favoring hydroxylation over competing fluorinatation and azide rebound pathways (Tables S16–18). These observations are consistent with a mechanism in which higher pH facilitates deprotonation of Fe-bound water to generate a Fe(III)–OH species responsible for hydroxy rebound.

**Table 3.**
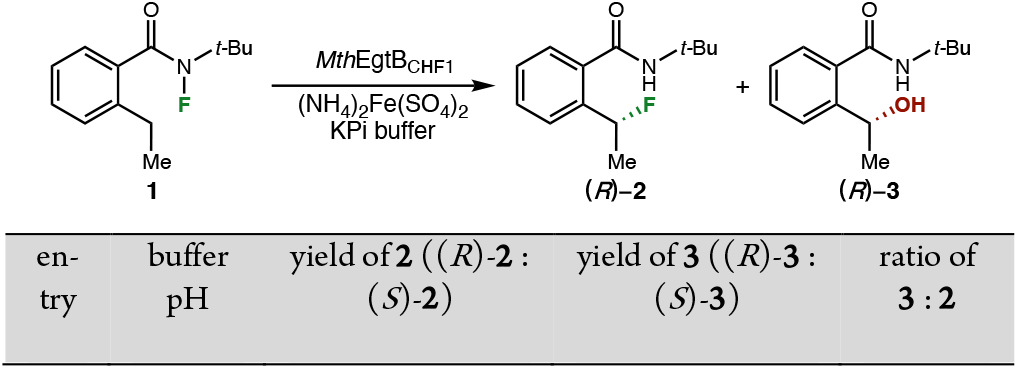
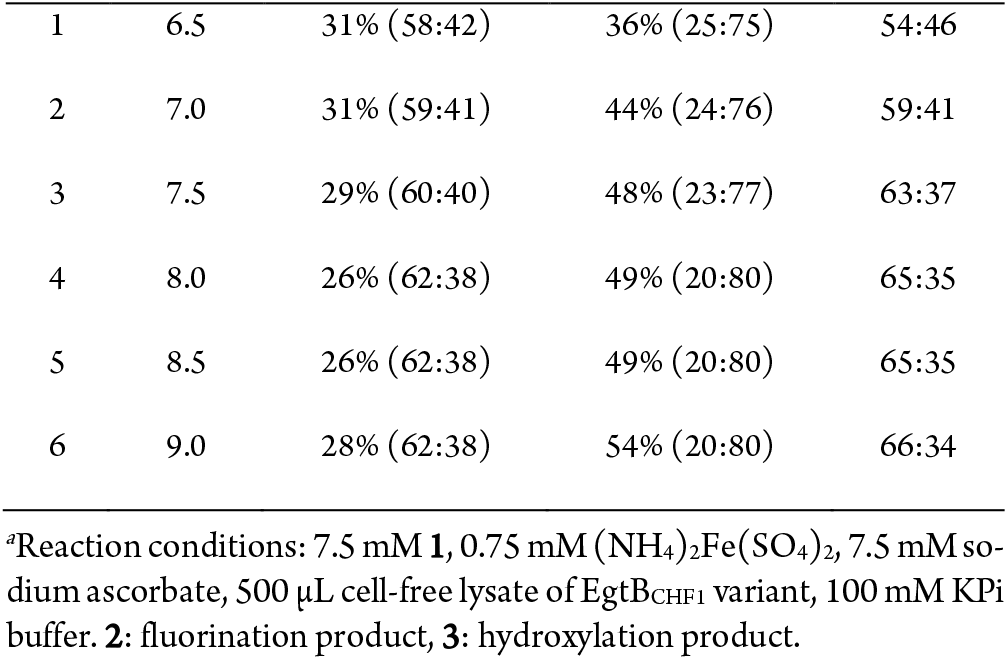
The Influence of Primary Coordination Sphere on Radical Re-bound Activity and Selectivity.

### Importance of Fe primary coordination sphere

To experimentally validate these findings, we conducted site-directed mutagenesis on Fe-binding residues of *Mth*EgtB_CHF_ and ACCO_CHF_ (**Figure 5**). We first mutated each of the three Fe-binding histidine residues of EgtB_CHF1_, including H51, H138 and H134, to an aspartate (D), a glutamate (E), and an alanine (A). Mutating H51 and H134 to D, E, or A completely abolished the EgtB’s fluorination and hydroxylation activity (Table 4, entries 2–7). Iron-content analysis showed that the H51 and H134 mutants exhibited reduced Fe incorporation (Table S4), suggesting that the loss of total activity may stem from reduced Fe binding affinity. Accordingly, absolute activity comparisons among these mutants should be interpreted with caution, whereas the observed changes in chemo- and enantioselectivity provide more direct insight into the role of the primary coordination sphere in controlling radical rebound selectivity. Importantly, it was found that the H138 mutants, including EgtB_CHF1_ H138D, H138E, and H138A, all retained its catalytic activity. Mutating H138 to either an aspartate (H138D, entry 8) or a glutamate (H138E, entry 9) abolished the hydroxylase activity while retaining their fluorinase activity, although this activity was reduced compared to the three-histidine enzyme EgtB_CHF1_ (entry 1). Notably, both H138D and H138E led to an inversion of enantiopreference for C– H fluorination, suggesting mutations to the primary coordination sphere residues not only perturb the coordination environment of the ferric center but also reshape the stereochemical course of the radical rebound event. Interestingly, EgtB_CHF1_ H138A, possessing the same two-histidine facial dyad found in naturally occurring *α*KG-dependent halogenases,^11c, 12e, 12h, 12l^ retained both fluorination and hydroxylation activities, affording a **2:3** ratio of 35:65. The enantioselectivities of products **2** and **3** remained unchanged relative to EgtB_CHF1_ (entry 10). Moreover, the observed preference for hydroxylation is consistent with the low p*K*_a_ value calculated for the (Im)_2_Fe(III)(F)(H_2_O)_3_ ^2+^ complex (p*K*_a_ = 5.3) that indicates the preferential formation of Fe(III)–OH species in EgtB_CHF1_ H138A (**Table 6**, entry 8, *vide infra*). Starting from the other evolved EgtB-_CHF2_, the introduction of H51/H134/H138 mutations led to a complete loss of catalytic activity (entries 12–14, see Table S6 for details). Because ACCO_CHF_ exhibited substantially higher activity and selectivity under whole-cell conditions, whereas the EgtB_CHF_ provided better fluorination results as cell-free lysates, the corresponding Fe-binding residue mutagenesis studies on ACCO_CHF_ were carried out under whole-cell reaction conditions to assess the catalytic activity and selectivity of these ACCO_CHF_ mutants. Additional detailed results for ACCO activity studies are available in Tables S7-S8. Finally, with our previously evolved ACCO_CHF_,^14c^ mutating its Fe-binding aspartate to a glutamate (entry 16), a histidine (entry 17), or an alanine (entry 18) fully abolished its activity. Collectively, these results further supported that nonheme Fe enzymes with a three-histidine and two-histidine coordination chemistry can support the promiscuous hydroxylation activity, highlighting the importance of coordination geometry, ligand identity, and electronic environment in controlling radical rebound selectivity in nonheme Fe enzyme-catalyzed reactions.

**Table 4.**
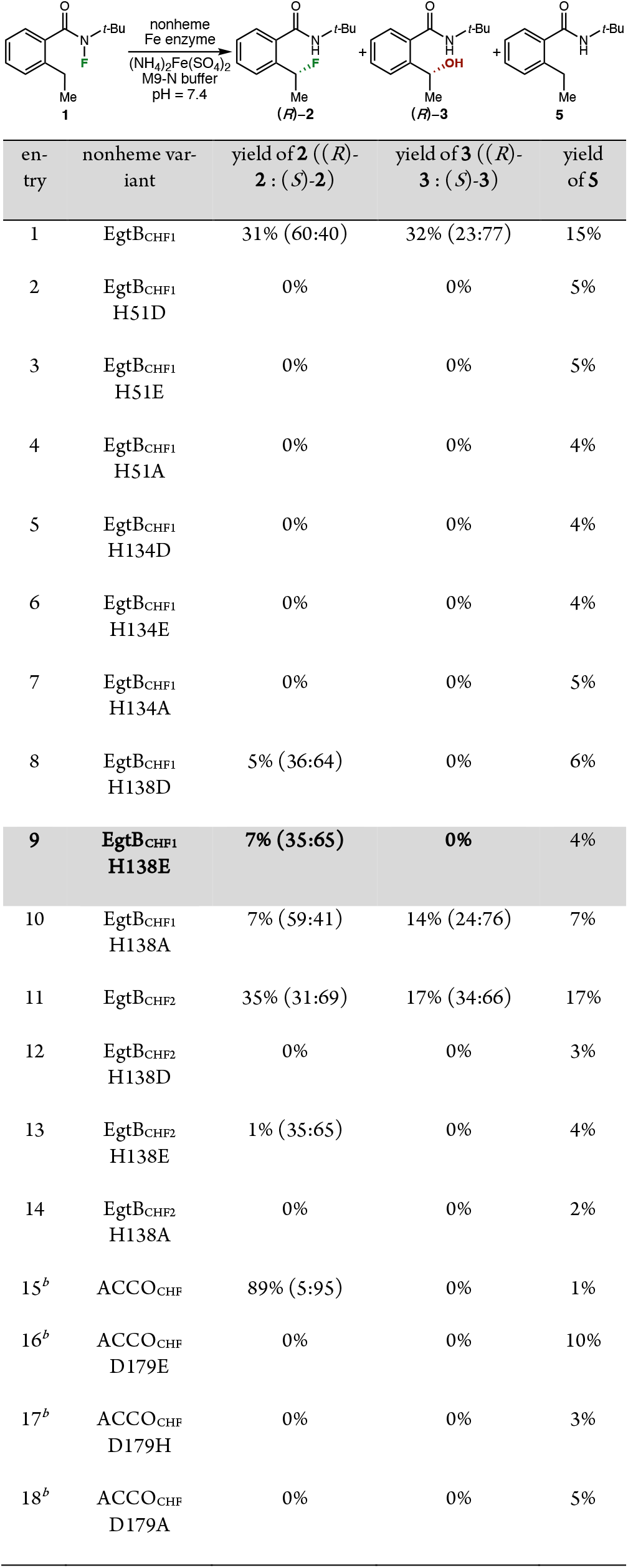
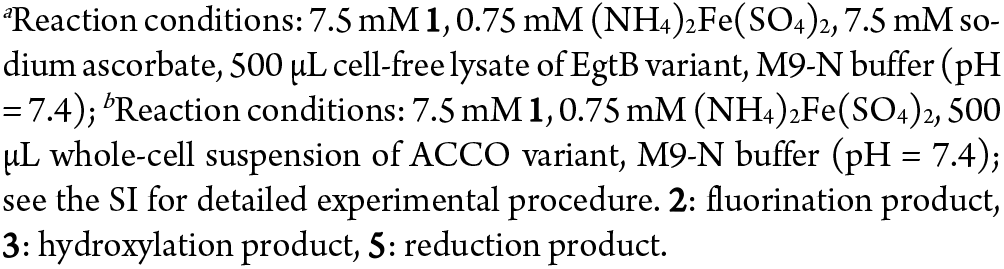
The Influence of Primary Coordination Sphere on Radical Rebound Activity Species and Anion Selectivity.

| entry | nonheme variant | yield of **2** ((*R*)-**2** : (*S*)-**2**) | yield of **3** ((*R*)-**3** : (*S*)-**3**) | yield of **5** |
| --- | --- | --- | --- | --- |
| 1 | EgtB<sub>CHF1</sub> | 31% (60:40) | 32% (23:77) | 15% |
| 2 | EgtB<sub>CHF1</sub> H51D | 0% | 0% | 5% |
| 3 | EgtB<sub>CHF1</sub> H51E | 0% | 0% | 5% |
| 4 | EgtB<sub>CHF1</sub> H51A | 0% | 0% | 4% |
| 5 | EgtB<sub>CHF1</sub> H134D | 0% | 0% | 4% |
| 6 | EgtB<sub>CHF1</sub> H134E | 0% | 0% | 4% |
| 7 | EgtB<sub>CHF1</sub> H134A | 0% | 0% | 5% |
| 8 | EgtB<sub>CHF1</sub> H138D | 5% (36:64) | 0% | 6% |
| **9** | **EgtB<sub>CHF1</sub> H138E** | **7% (35:65)** | **0%** | **4%** |
| 10 | EgtB<sub>CHF1</sub> H138A | 7% (59:41) | 14% (24:76) | 7% |
| 11 | EgtB<sub>CHF2</sub> | 35% (31:69) | 17% (34:66) | 17% |
| 12 | EgtB<sub>CHF2</sub> H138D | 0% | 0% | 3% |
| 13 | EgtB<sub>CHF2</sub> H138E | 1% (35:65) | 0% | 4% |
| 14 | EgtB<sub>CHF2</sub> H138A | 0% | 0% | 2% |
| 15<sup>b</sup> | ACCO<sub>CHF</sub> | 89% (5:95) | 0% | 1% |
| 16<sup>b</sup> | ACCO<sub>CHF</sub> D179E | 0% | 0% | 10% |
| 17<sup>b</sup> | ACCO<sub>CHF</sub> D179H | 0% | 0% | 3% |
| 18<sup>b</sup> | ACCO<sub>CHF</sub> D179A | 0% | 0% | 5% |

<sup>a</sup>Reaction conditions: 7.5 mM **1**, 0.75 mM (NH<sub>4</sub>)<sub>2</sub>Fe(SO<sub>4</sub>)<sub>2</sub>, 7.5 mM sodium ascorbate, 500 μL cell-free lysate of EgtB variant, M9-N buffer (pH = 7.4); <sup>b</sup>Reaction conditions: 7.5 mM **1**, 0.75 mM (NH<sub>4</sub>)<sub>2</sub>Fe(SO<sub>4</sub>)<sub>2</sub>, 500 μL whole-cell suspension of ACCO variant, M9-N buffer (pH = 7.4); see the SI for detailed experimental procedure. **2**: fluorination product, **3**: hydroxylation product, **5**: reduction product.

**Table 5.** Primary Coordination Sphere Control of Promiscuous Radical Rebound Activity in the Presence of Azide^*a*^.

| entry | nonheme variant | yield of **2** :<br>((*R*)-**2** :<br>(*S*)-**2**) | yield of **3** :<br>((*R*)-**3** :<br>(*S*)-**3**) | yield of **4** :<br>((*R*)-**4** :<br>(*S*)-**4**) | yield of **5** |
| --- | --- | --- | --- | --- | --- |
| **1** | **EgtB<sub>CHF1</sub>** | **15%**<br>**(60:40)** | **23%**<br>**(23:77)** | **50%**<br>**(44:56)** | 15% |
| 2 | EgtB<sub>CHF1</sub><br>H138D | 2%<br>(35:65) | 0% | 0% | 3% |
| 3 | EgtB<sub>CHF1</sub><br>H138E | 7%<br>(35:65) | 0% | 0% | 5% |
| 4 | EgtB<sub>CHF1</sub><br>H138A | 8%<br>(50:50) | 9%<br>(24:76) | 32%<br>(45:55) | 9% |
| 5 | EgtB<sub>CHF2</sub> | 15%<br>(31:69) | 10%<br>(33:67) | 40%<br>(39:61) | 12% |
| 6 | EgtB<sub>CHF2</sub><br>H138D | 0% | 0% | 0% | 3% |
| 7 | EgtB<sub>CHF2</sub><br>H138E | 1% | 0% | 1% | 5% |
| **8** | **EgtB<sub>CHF2</sub>**<br>**H138A** | **1%** | **0%** | **42%**<br>**(27:73)** | 8% |
| 9 | ACCO<sub>CHF</sub> | 77%<br>(6:94) | 0% | 12%<br>(49:51) | 5% |
| 10<sup>b</sup> | ACCO<sub>CHF</sub><br>D179E | 0% | 0% | 14%<br>(45:55) | 10% |
| 11<sup>b</sup> | ACCO<sub>CHF</sub><br>D179H | 0% | 0% | 0% | 3% |
| 12<sup>b</sup> | ACCO<sub>CHF</sub><br>D179A | 0% | 0% | 10%<br>(58:42) | 5% |

<sup>a</sup>Reaction conditions: 7.5 mM **1**, 120 mM NaN<sub>3</sub>, 0.75 mM (NH<sub>4</sub>)<sub>2</sub>Fe(SO<sub>4</sub>)<sub>2</sub>, 7.5 mM sodium ascorbate, 500 μL cell-free lysate of EgtB variant, M9-N buffer (pH = 7.4); <sup>b</sup>Reaction conditions: 7.5 mM **1**, 120 mM NaN<sub>3</sub>, 0.75 mM (NH<sub>4</sub>)<sub>2</sub>Fe(SO<sub>4</sub>)<sub>2</sub>, 500 μL whole-cell suspension of ACCO variant, M9-N buffer (pH = 7.4); see the SI for detailed experimental procedure. **2**: fluorination product, **3**: hydroxylation product, **4**: azidation product, **5**: reduction product.

**Table 6.** Computed activation Gibbs free energies (kcal/mol) for fluorine atom abstraction (**TS-1**) and radical rebound with fluorine (**TS-3**), hydroxy (**TS-3-OH**), and azide (**TS-3-N_3_**) intermediates; p*K*_a_ and LUMO energies (*E*_LUMO_, eV) of Fe(III)–F species; bond dissociation enthalpies (BDE, kcal/mol) of Fe(III)–F, Fe(III)–N_3_ and Fe(III)–OH bonds; and local force constants (*k*^a^, mDyne/Å) of Fe–F bonds across different primary coordination spheres. Fe(II) and Fe(III)–F intermediates were calculated at high-spin quintet and sextet states, respectively.^*a,b*^.

| entry | nonheme Fe<sup>II</sup> species (A) | resulting Fe<sup>III</sup>-F species (B) | Activation Gibbs free energy ($\Delta G^\ddagger$) | | | | Properties of Fe<sup>III</sup>-F species (B) | | |
| --- | --- | --- | --- | --- | --- | --- | --- | --- | --- |
|  |  |  | TS-1 | TS-3 (F)<br>(Fe-F BDE of B) | TS-3-OH<br>(Fe-OH BDE of B) | TS-3-N<sub>3</sub><br>(Fe-N<sub>3</sub> BDE of B) | p*K*<sub>a</sub> of Fe-bound H<sub>2</sub>O | *k*<sup>2</sup>(Fe-F) | *E*<sub>LUMO</sub> |
| 1 | (Im)<sub>3</sub>Fe<sup>II</sup>(H<sub>2</sub>O)<sub>2</sub> (**11**) | (Im)<sub>3</sub>Fe<sup>III</sup>F(H<sub>2</sub>O)<sub>2</sub> (**12-A**) | 13.3 | 0.0 (75.2) | - | - | 8.7 | 3.00 | -1.8 |
| 2 | (Im)<sub>3</sub>Fe<sup>II</sup>(N<sub>3</sub><sup>-</sup>)(H<sub>2</sub>O) | (Im)<sub>3</sub>Fe<sup>III</sup>F(N<sub>3</sub><sup>-</sup>)(H<sub>2</sub>O) | 12.1 | 6.6 (81.7) | - | 2.3 (29.2) | 13.2 | 2.58 | -0.9 |
| 3 | (Im)<sub>3</sub>Fe<sup>II</sup>(OH<sup>-</sup>)(H<sub>2</sub>O) | (Im)<sub>3</sub>Fe<sup>III</sup>F(OH<sup>-</sup>)(H<sub>2</sub>O) (**12-B/13**) | 10.2 | 9.3 (82.0) | 5.4 (45.3) | - | 15.2 | 2.21 | -0.8 |
| 4 | (Im)<sub>3</sub>Fe<sup>II</sup>(N<sub>3</sub><sup>-</sup>)(OH<sup>-</sup>) | (Im)<sub>3</sub>Fe<sup>III</sup>F(N<sub>3</sub><sup>-</sup>)(OH<sup>-</sup>) | 13.7 | 16.3 (91.5) | 11.3 (50.2) | 11.7 (43.5) | N/A | 2.40 | 0.1 |
| 5 | (Im)<sub>2</sub>(AcO)Fe<sup>II</sup>(H<sub>2</sub>O)<sub>2</sub> (**11'**) | (Im)<sub>2</sub>(AcO)Fe<sup>III</sup>F(H<sub>2</sub>O)<sub>2</sub> (**12'/13'**) | 11.8 | 3.4 (81.4) | - | - | 10.6 | 2.61 | -1.0 |
| 6 | (Im)<sub>2</sub>(AcO)Fe<sup>II</sup>(N<sub>3</sub><sup>-</sup>)(H<sub>2</sub>O) | (Im)<sub>2</sub>(AcO)Fe<sup>III</sup>F(N<sub>3</sub><sup>-</sup>)(H<sub>2</sub>O) | 16.1 | 10.1 (83.0) | - | 8.9 (33.1) | 16.7 | 2.90 | -0.1 |
| 7 | (Im)<sub>2</sub>(AcO)Fe<sup>II</sup>(OH<sup>-</sup>)(H<sub>2</sub>O) | (Im)<sub>2</sub>(AcO)Fe<sup>III</sup>F(OH<sup>-</sup>)(H<sub>2</sub>O) | 17.2 | 13.3 (87.3) | 13.2 (47.6) | - | 12.6 | 2.67 | 0.0 |
| 8 | (Im)<sub>2</sub>Fe<sup>II</sup>(H<sub>2</sub>O)<sub>3</sub> | (Im)<sub>2</sub>Fe<sup>III</sup>F(H<sub>2</sub>O)<sub>3</sub> | 13.0 | 0.0<sup>c</sup> (69.8) | - | - | 5.3 | 3.31 | -2.0 |
| 9 | (Im)<sub>2</sub>Fe<sup>II</sup>(N<sub>3</sub><sup>-</sup>)(H<sub>2</sub>O)<sub>2</sub> | (Im)<sub>2</sub>Fe<sup>III</sup>F(N<sub>3</sub><sup>-</sup>)(H<sub>2</sub>O)<sub>2</sub> | 7.4 | 6.2 (77.5) | - | 0.9 (24.8) | 11.8 | 2.40 | -1.1 |
| 10 | (Im)<sub>2</sub>Fe<sup>II</sup>(OH<sup>-</sup>)(H<sub>2</sub>O)<sub>2</sub> | (Im)<sub>2</sub>Fe<sup>III</sup>F(OH<sup>-</sup>)(H<sub>2</sub>O)<sub>2</sub> | 12.3 | 8.9 (83.1) | 4.7 (46.1) | - | 9.0 | 2.47 | -0.8 |
| 11 | (Im)<sub>2</sub>Fe<sup>II</sup>(N<sub>3</sub><sup>-</sup>)(OH<sup>-</sup>)(H<sub>2</sub>O) | (Im)<sub>2</sub>Fe<sup>III</sup>F(N<sub>3</sub><sup>-</sup>)(OH<sup>-</sup>)(H<sub>2</sub>O) | 11.0 | 13.3 (86.0) | 11.0 (44.9) | 10.3 (34.4) | 16.0 | 1.92 | -0.1 |

<sup>a</sup>Imidazole and acetate were used to model the Fe-bound histidine and aspartate residues, respectively. In alanine mutants (EgtB<sub>CHF</sub> H138A and ACCO<sub>CHF</sub> D179A), the mutated residue was replaced by a water ligand in the truncated computational models. <sup>b</sup>DFT calculations were performed at the B3LYP-D3(BJ)/def2-TZVP/SMD//B3LYP-D3(BJ)/6-31G(d)-SDD(Fe) level of theory. <sup>c</sup>This step is barrierless (no TS was located).

**Figure 5.**
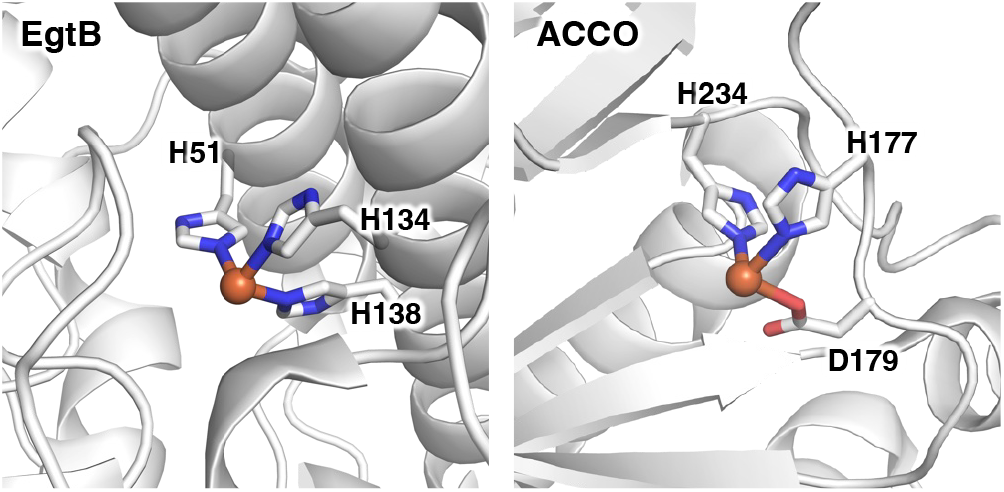
Active-site illustrations with the primary coordination sphere were made based on PDB ID 4X8E (*Mth*EgtB) and PDB ID 1W9Y (ACCO).

To further probe the impact of primary coordination sphere and active-site environment on other anion rebound activities with non-heme Fe enzymes, we investigated the same set of Fe-binding residue mutants of EgtB_CHF1_, EgtB_CHF2_ and ACCO_CHF_ in the presence of added NaN_3_. Both EgtB_CHF1_ and EgtB_CHF2_ displayed highly promiscuous radical rebound activities in the presence of exogenous azide anion, furnishing the corresponding C(sp^3^)–H azidation product **5** in 50% yield (56:44 e.r.) and 40% yield (61:39 e.r.), respectively (Table 5, entries 1 and 5). In addition, both the fluorinated and hydroxylated products were also observed. For EgtB_CHF1_, the fluorination product **2** was formed in 15% yield with 60:40 e.r., along with hydroxylation product **3** in 23% yield with 23:77 e.r.. On the other hand, EgtB_CHF2_ afforded **2** in 15% yield (31:69 e.r.) and **3** in 10% yield (33:67 e.r.). Next, we investigated the promiscuous activity of EgtB_CHF_ variants bearing different primary coordination sphere mutations. For both EgtB_CHF1_ and EgtB_CHF2_ variants, mutating H51 and H134 to D, E, or A completely abolished their catalytic activity (see Tables S9 and S10 for details), consistent with results in the absence of azide. Both EgtB_CHF1_ H138D (entry 2) and EgtB_CHF1_ H138E (entry 3) catalyzed C(sp^3^)–H fluorination in the presence of NaN_3_. Importantly, with EgtB_CHF1_ H138D (entry 2) and EgtB_CHF1_ H138E (entry 3), even upon addition of 16 equiv NaN_3_, neither the hydroxylation product nor the azidation product could be detected, highlighting the fluorination fidelity of these two-His-one-carboxylate mutants of EgtB. In contrast, the EgtB_CHF1_ H138A mutant displayed catalytic promiscuity, producing fluorination product **2**, hydroxylation product **3**, and azidation product **4** in 8% (50:50 e.r.), 9% (24:76 e.r.) and 32% (45:55 e.r.), respectively (entries 4). Starting from EgtB_CHF2_, the introduction of H138D and H138E nearly abolished their catalytic activity (entries 6 and 7). Interestingly, the EgtB_CHF2_ H138A mutant exhibited significantly enhanced azidation activity and selectivity, furnishing the azidation product **5** in 42% yield and 73:27 e.r. with an azidation: fluorination ratio of 97.5:2.5 (entry 8). The change in enantioselectivity between EgtB_CHF2_ (entry 5) and EgtB-_CHF2_ H138A (entry 10) further highlighted the role of the primary coordination sphere on controlling radical rebound reactivity and stereoselectivity. Next, we examined ACCO_CHF_ in the presence of 16 equiv NaN_3_. Under these conditions, ACCO_CHF_ still afforded the fluorination product **2** in 77% yield and 94:6 e.r., with only 12% yield of the azidation product in nearly a racemic fashion (51:49 e.r., entry 9). Taken together, these results illustrate the fine interplay of primary coordination sphere and active-site environment in nonheme Fe enzymes in controlling radical rebound activity and anion selectivity, providing a mechanistic basis for rationally modulating C(sp^3^)–H fluorination, hydroxylation, and azidation reactions.

### Computational studies

To gain further insights into the mechanism of nonheme Fe enzyme-catalyzed fluorine atom transfer and subsequent fluorine rebound, we performed DFT calculations of the reaction energy profile for this biocatalytic C–H fluorination using a truncated model of EgtB_CHF1_ (**Figure 6(A**), black pathway). The imidazole (Im) ligands were used to model the Fe-bound histidine residues^14c^ (see the Supporting Information for computational details). Non-heme Fe(II)–H_2_O and Fe(III)–OH species were used, as our p*K*_a_ calculations indicate that the water ligand bound to a ferrous center stays protonated while that bound to Fe(III) could be deprotonated in this ligand environment (see **Scheme 2**). These DFT calculations indicate that all the intermediates and transition states in the catalytic cycle feature high-spin quintet Fe(II) and sextet Fe(III)^14c,33^ (see **Figure S13** for the computed pathway at the less favorable intermediate-spin state). The computed reaction energy profile revealed a relatively low barrier of 13.3 kcal/mol for the initial fluorine atom abstraction (**TS-1, Figure 6(B)**) from the *N*-fluoroamide substrate **1** to the Fe(II) center. The highly exergonic nature of fluorine atom abstraction (*ΔG* = −10.2 kcal/mol) reflects conversion of the N–F bond in **1** (BDE (N–F) = 64.7 kcal/mol) to a stronger Fe(III)– F bond (BDE (Fe(III)–F of **12-A**) = 75.2 kcal/mol), providing a favorable thermodynamic driving force for substrate activation. Following fluorine atom abstraction, the Fe(III)-bound water undergoes kinetically feasible deprotonation by a basic residue via a water wire network (**Figures S17–S18**). Classical MD simulations indicate that D81 and D87 are connected to the Fe-bound water through a persistent hydrogen-bonding network, providing a plausible pathway for proton transfer and generation of the Fe(III)–OH species. Concurrently, the nitrogen-centered radical in **12-B** undergoes rapid 1,5-hydrogen atom transfer (1,5-HAT) via **TS-2** to generate the benzylic radical intermediate **13**. Subsequent radical rebound with the Fe(III)–F species to form the C–F bond via **TS-3** is also kinetically facile, with an activation barrier of 9.3 kcal/mol relative to intermediate **13**. The low barrier to C–F bond formation through an Fe(III)–F intermediate highlights the viability of radical rebound fluorination in nonheme Fe systems. These findings have broad implications for repurposing nonheme Fe enzymes as fluorinases to catalyze diverse C–F bond-forming reactions, an enzymatic function that has historically remained elusive to enzymologists. We also evaluated alternative pathways, including substrate reduction by the Fe(II) species and benzyl radical oxidation by the Fe(III)–F species (**Figures S23–S24**). Both pathways are highly endothermic, making them less likely under the reaction conditions.

**Figure 6.**
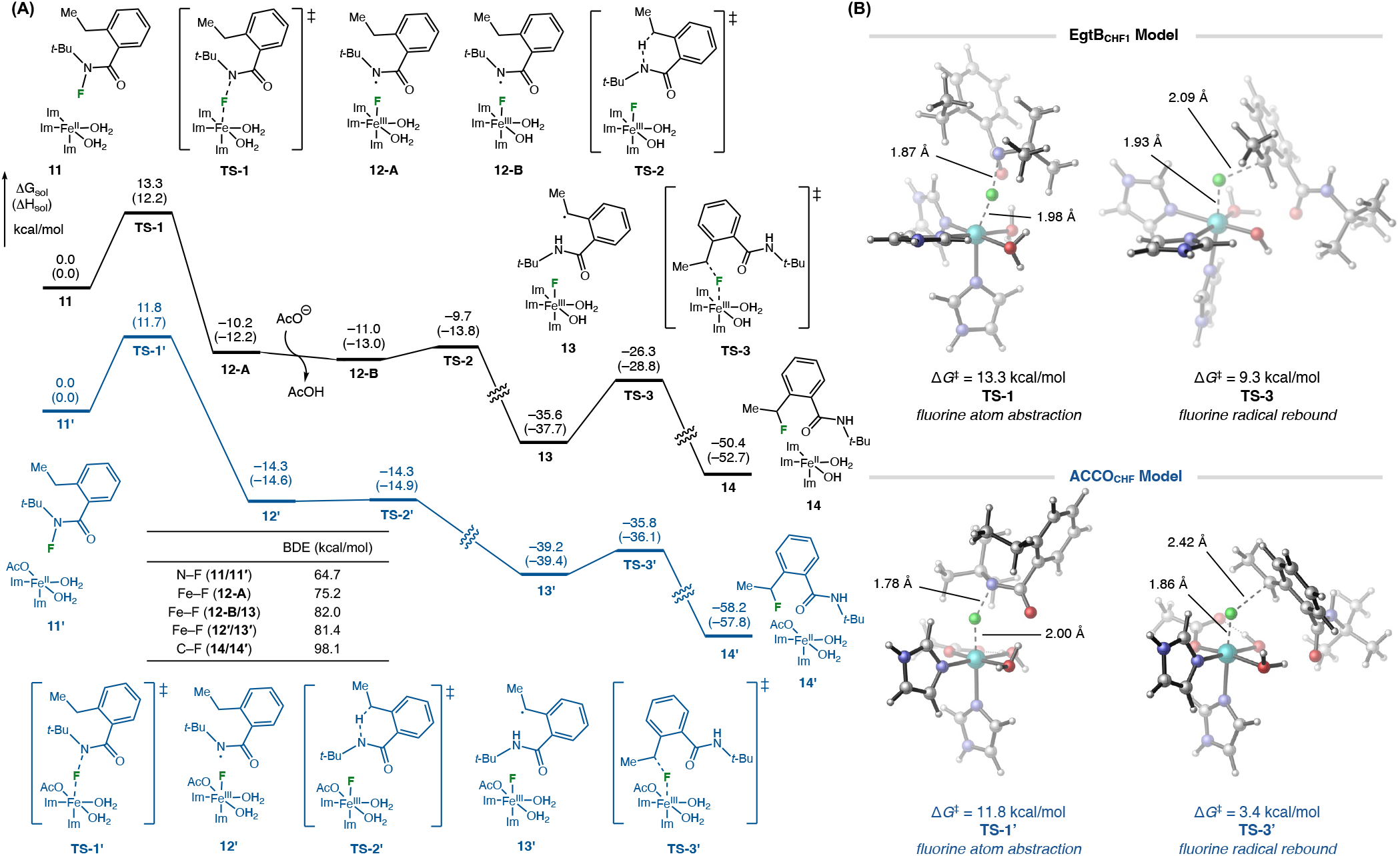
(A) DFT-computed reaction energy profile of EgtB_CHF1_ (black) and ACCO_CHF_-catalyzed (blue) fluorine atom transfer of *N*-fluoroamide 1 at quintet spin state using a truncated model at the B3LYP-D3(BJ)/def2-TZVP/SMD//B3LYP-D3(BJ)/6-31G(d)–SDD(Fe) level of theory. (B) Optimized geometries of the fluorine atom abstraction (**TS-1/TS-1′**) and fluorine radical rebound (**TS-3/TS-3′**) transition states with EgtB_CHF1_ and ACCO_CHF_ model systems. Imidazole (Im) and acetate (OAc) groups are models for histidine and aspartate residues, respectively. BDE, bond dissociation enthalpy.

We next compared the reaction energy profile catalyzed by ACCO_CHF_ featuring a two-histidine-one-carboxylate facial triad (**Figure 6(A)**, blue pathway) present in EgtB_CHF1_.^14c^ Although EgtB-_CHF1_ and ACCO_CHF_^14c^ follow the same overall reaction mechanism for the fluorine atom transfer, the computed reaction energy profiles reveal notable differences in their individual elementary steps. First, p*K*_a_ calculations suggest that the water molecules bound to both Fe(II) and Fe(III) complexes of ACCO_CHF_ remain as neutral aqua species (*vide supra*, see **Figures S15-S16**). Second, the activation barriers to fluorine atom abstraction and fluorine radical rebound steps are both lower in ACCO_CHF_ than those in EgtB_CHF1_. The lower barrier to the fluorine atom abstraction (**TS-1’**) could be attributed to the formation of a stronger Fe–F bond in **12’** (BDE = 81.4 kcal/mol compared with 75.2 kcal/mol for **12-A**), due to its more electron-rich Fe center supported by stronger *σ*-donor ligands. On the other hand, the lower barrier to fluorine radical rebound (**TS-3’**, *ΔG*^‡^ = 3.4 kcal/mol compared with 9.3 kcal/mol from **13** via **TS-3**) is not simply driven by thermodynamics, considering the comparable Fe–F BDEs in **13** and **13’** (82.0 and 81.4 kcal/mol, respectively). These results suggest that other factors beyond thermodynamic driving force contribute to the fluorine radical rebound reactivity. This prompted us to perform a more thorough quantitative analysis on factors determining fluorine atom abstraction and fluorine radical rebound reactivities.

To further elucidate how the primary coordination sphere of non-heme Fe enzymes controls fluorine atom abstraction and radical rebound activity and selectivity, we performed additional DFT calculations on truncated nonheme Fe models bearing a series of supporting ligands (**Table 6**). These models represent nonheme Fe centers coordinated by three-histidine (entries 1–4), two-histidine-one-carboxylate (entries 5–7), and two-histidine (entries 8–11) ligand sets, including those containing additional azide and/or hydroxy ligands. For the H138A and D179A models, the coordinating residue was replaced by a water ligand to preserve the octahedral coordination environment of the Fe center, consistent with solvent coordination upon loss of the coordinating side chain. Binding free energy calculations further indicate that azide anion binding to both the Fe(II) and Fe(III) species is thermodynamically favorable (**Figure S25**), supporting the feasibility of the corresponding Fe–N_3_ intermediates. Our truncated nonheme Fe models enabled direct comparison of the fluorine atom abstraction (**TS-1**) step and the competing radical rebound pathways involving fluorine (**TS-3**), hydroxy (**TS-3-OH**), and azide (**TS-3-N**_**3**_) across distinct coordination environments. Across all environments examined, the intrinsic rebound preference generally follows the order N_3_ > OH > F. Azide rebound exhibits the lowest activation barriers, likely facilitated by the weaker Fe–N_3_ bonds compared with Fe–OH and Fe–F species (see **Figure S20** for the correlation between activation barriers for radical rebound and BDEs). In contrast, the tightly bound and highly electronegative fluorine ligand increases the energetic penalty for Fe–F bond cleavage, rendering fluorine rebound intrinsically less favorable.

### Multivariate linear regression studies

Because the kinetic barriers to fluorine atom abstraction and radical rebounds are not governed solely by thermodynamics, we next sought to develop physics-informed, descriptor-based models for these elementary steps. Several chemically meaningful descriptors were evaluated to identify correlations with reactivity. These include the bond dissociation enthalpy, BDE(Fe–X), as a thermodynamic descriptor of Fe–X bond strength. In addition, we considered the local force constant, *k*^*a*^(Fe–F),^34^ and the vertical bond dissociation energy,^36^ BDE_vert_(Fe–X), to capture the intrinsic strength of the Fe–X bond at its equilibrium position. BDE_vert_(Fe–X) values were calculated as the energy required for homolytic Fe–X bond cleavage without geometric relaxation, thereby isolating the electronic contribution to bond cleavage. Lastly, we considered the LUMO energy of the Fe(III)–F complex, *E*_LUMO_(Fe^III^F), as an electronic descriptor reflecting the influence of the primary coordination sphere at the Fe center. Due to the limited size of the dataset (11 data points, **Table 6**), all data were included in the training set, and model robustness was assessed using leave-one-out cross-validation (*Q*^2^_LOOCV_).^35^

For fluorine atom abstraction, multivariate linear regression analysis revealed a two-parameter model based on BDE_vert_(Fe–F) and *k*^*a*^(Fe–F) (*R*^2^ = 0.85, *Q*^2^_LOOCV_ = 0.75) (**Figure 7(A)**), demonstrating that fluorine atom abstraction is predominantly governed by the intrinsic strength of the Fe–F bond. Although BDE_vert_(Fe–F) and local force constant *k*^*a*^(Fe–F) are both related to bond strength, the correlation between these two descriptors is only moderate (*R*^*2*^ = 0.68). The variance inflation factor^37^ value of 3.2 indicates that their multi-collinearity is well controlled and does not compromise the stability of this two-parameter model. Models based on the fully relaxed bond dissociation enthalpy (BDE) display lower descriptor–reactivity correlations (*R*^2^ = 0.82, *Q*^2^_LOOCV_ = 0.56, **Figure S21**), suggesting that inclusion of structural relaxation and thermodynamic contributions diminishes predictive power for the transition-state barrier.

**Figure 7.**
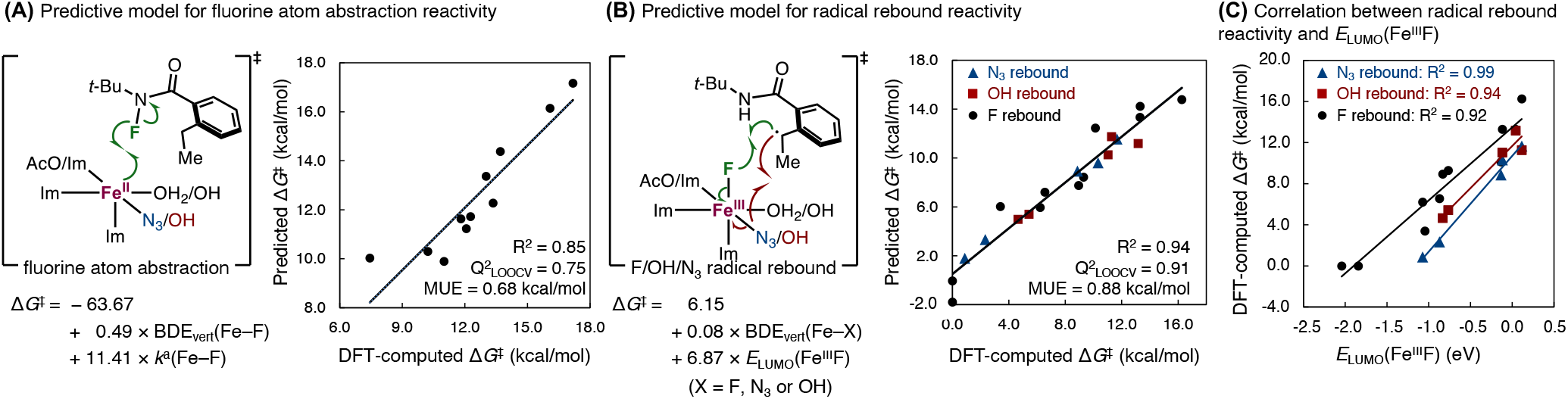
Predictive models for the **(A)** fluorine atom abstraction and **(B)** radical rebound reactivity. *Q*^2^_LOOCV_leave-one-out cross-validation coefficient of determination. (C) Correlations between radical rebound reactivity (Δ*G*^‡^ _DFT_) and LUMO energy of the Fe(III)–F species.

Next, we developed a multivariate linear regression model for the reactivity of all three competing radical rebound processes involving fluorine, azide, and hydroxy groups. The best performing two-parameter model uses BDE_vert_(Fe–F) and *E*_LUMO_(Fe^III^F) as descriptors (*R*^2^ = 0.94, *Q*^2^ _LOOCV_= 0.91, **Figure 7(B)**). The relatively small coefficient for BDE_vert_(Fe–F) (0.08) indicates that radical rebound activity is predominantly electronically controlled: more electron-deficient Fe(III) centers with lower LUMO energies exhibit higher rebound activity. An alternative model using the BDE of forming C–X bond and *E*_LUMO_(Fe^III^F) as descriptors gave comparable performance (*R*^2^ = 0.93, **Figure S22**). In fact, excellent correlations between rebound barriers and *E*_LUMO_(Fe^III^F) were observed for radical rebound reactions with the same functional group (*R*^2^ = 0.92, 0.94, and 0.99 for F, OH, and N_3_ rebound, respectively, **Figure 7(C)**), further reinforcing the importance of the electronic properties of the Fe(III) species in this step. This observation is consistent with previous study by Solomon and co-workers,^19a^ which highlighted the critical role of frontier molecular orbital energetics of Fe(III)–X intermediates in controlling radical rebound selectivity in nonheme Fe enzymes. Our results further suggest that modulation of the primary coordination sphere can alter these electronic properties and thereby influence radical rebound reactivity. Taken together, these descriptor-based regression analyses of fluorine atom abstraction and radical rebound provide mechanistic insights into how the primary coordination sphere modulates these individual elementary steps, thereby enabling control over reactivity and selectivity in non-heme Fe enzymes.

To gain insights into the preferred substrate binding mode and the roles of active site residues in EgtB-catalyzed C(sp^3^)–H fluorination, we performed classical molecular dynamics (MD) simulations for *N*-fluoroamide **1** bound to EgtB_CHF1_ (**Figure 8(A)**, left). To model the near-attack-conformations (NACs)^38^ prior to the fluorine atom abstraction step, the Fe–F distance between **1** and the Fe(II) center was restrained to 3.0–3.2 Å using a harmonic potential of 100 kcal·mol^−1^·Å^−2^. This restrained distance range was chosen based on DFT calculations, which indicate an Fe–F distance of 3.04 Å in a model Fe(II)–*N*-fluoroamide dative complex (**Figure S9**). Because the potential substrate binding site *trans* to H138 is occupied by Q55 and W377 residues, *N*-fluoroamide **1** may coordinate *trans* to either H51 or H134 at the Fe center. Thus, we examined two binding modes in our MD simulations: mode **A** (substrate *trans* to H51) and mode **B** (substrate *trans* to H134) (**Figure S10**). The simulations reveal that in binding mode A, *N*-fluoroamide 1 forms a persistent hydrogen bond with residue W415R (**Figure 8(A)**). The N–H···O distance remains below 2.5 Å in 89% of the simulation time (**Figure S11**). In contrast, in binding mode **B**, the substrate does not engage in any significant stabilizing interactions with active-site residues (**Figure S9(B)**). The hydrogen bond interactions with W415R observed in binding mode A may promote fluorine atom abstraction via two complementary effects. First, it anchors the *N*-fluoroamide substrate, in particular the N–F moiety, in proximity to the Fe(II) center. Second, it electronically activates the amide N–F bond, lowering its bond dissociation enthalpy (BDE) from 64.7 to 62.9 kcal/mol and facilitating electron transfer from the metal center to the fluoroamide substrate during fluorine atom abstraction (**Figure S14**).^39^ These computational results are consistent with experimental observations showing that the W415R mutation enhances fluorination activity (*vide supra*).

**Figure 8.**
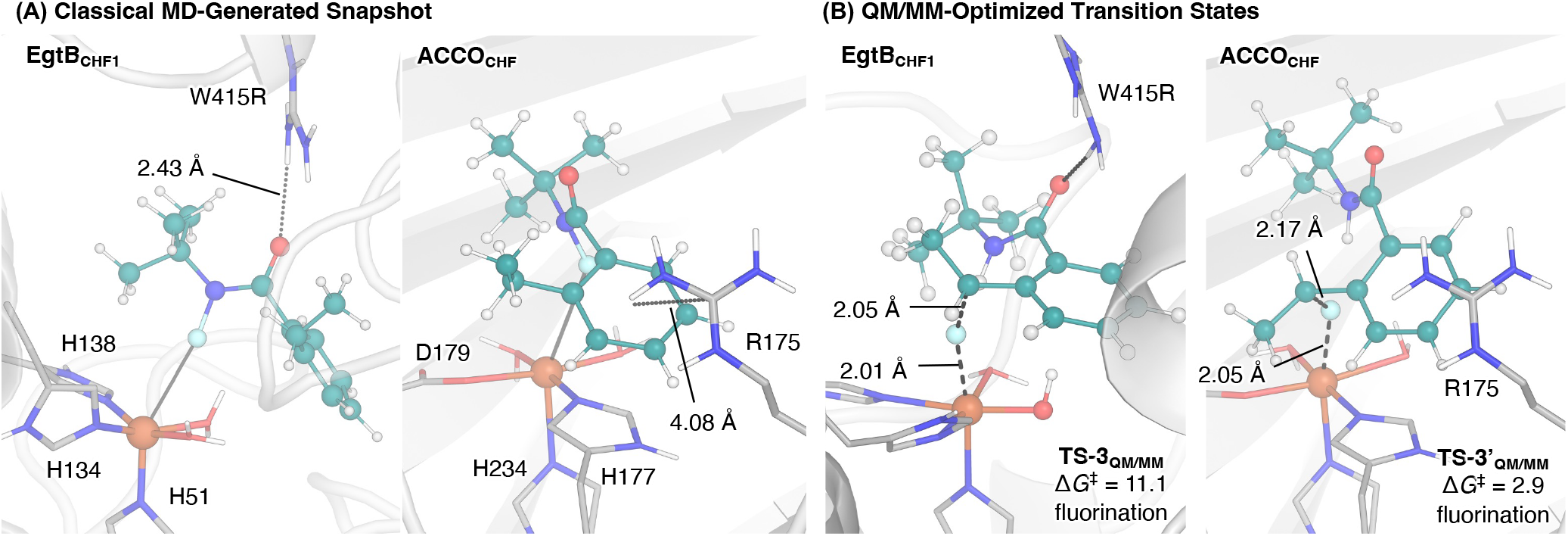
**(A)** The most populated structures from restrained classical MD simulations of *N*-fluoroamide **1** in the active sites of EgtB_CHF1_ and ACCO_CHF_. **(B)** QM/MM-optimized geometries of the radical rebound transition states for the C–H fluorination in EgtB_CHF1_ and ACCO_CHF_ at high-spin quintet state. Gibbs free energy values are in kcal/mol with respect to the Fe(III)–F intermediates. The *N*-fluoroamide substrate is shown in green.

In the NAC MD simulations with our previously engineered ACCO_CHF_,^14c^ a persistent arginine/*π* interaction^41^ between R175 and the phenyl group of the *N*-fluoroamide **1** substrate was observed (**Figure 8(A)**, right). This interaction may play a role in controlling radical rebound by restricting rotation of the phenyl ring and maintaining its proximity to the H177 ligand. Specifically, the arginine/*π* interaction likely limits the conformational flexibility of the benzyl radical intermediate; the resulting proximity of benzylic carbon to the Fe-bound fluoride may facilitate fluorine radical rebound while suppressing the completing azide rebound pathway in the presence of N_3_ ^-^.

To further understand the role of enzyme active site in affecting the C–H fluorination and hydroxylation, we performed QM/MM calculations using the ONIOM algorithm to locate the fluorine radical rebound transition states with EgtB_CHF1_ and ACCO_CHF_.^40^ The radical rebound with the Fe(III)–F species in EgtB_CHF1_ to form a C–F bond via **TS-3**_**QM/MM**_ is kinetically facile, requiring an activation barrier of 11.1 kcal/mol with respect to the Fe(III)–F intermediate **13**_**QM/MM**_ (**Figure S26**). In this structure, the hydrogen bond between the W415R side chain and the amide carbonyl of the substrate anchors the benzylic radical in proximity to the Fe(III)–F moiety, facilitating the radical rebound step. Additional QM/MM calculations of the competing hydroxy rebound pathway in EgtB_CHF1_ indicate that fluorine and hydroxy rebound are energetically comparable (Figure S26), supporting the experimentally observed competition between fluorination and hydroxylation. In the QM/MM-computed fluorine radical rebound transition state with ACCO_CHF_, a significantly low activation barrier of 2.9 kcal/mol relative to **13’**_**QM/MM**_ was observed. These results indicate that ACCO_CHF_ is a more efficient biocatalyst for fluorine radical rebound, consistent with the reactivity trend observed in the DFT small model calculations (*vide supra*).

## Conclusion

In summary, we repurposed and evolved a biosynthetic nonheme Fe enzyme EgtB which had not been previously investigated for un-natural enzymatic activity to enable C(sp^3^)–H fluorination, hydroxylation and azidation. Directed evolution of EgtB furnished two fluorine atom transferases with opposite enantiopreference, while comparative studies with our previously evolved nonheme Fe biocatalyst ACCO_CHF_ highlighted how subtle differences in primary coordination sphere profoundly influence reactivity and selectivity. Combined experimental and computational investigations delineate how the nonheme Fe ligand environment governs anion rebound selectivity in biocatalytic radical C(sp^*3*^)–H functionalization. Isotope-labeling experiments, systematic Fe-binding residue mutagenesis, DFT calculations, multivariate linear regression analysis, MD simulations and QM/MM studies collectively reveal that the enhanced Lewis acidity of a three-histidine-coordinated Fe(III) center enables facile deprotonation of Fe-bound water to generate a Fe(III)–OH species, thereby leading to promiscuous hydroxy rebound. In contrast, a two-histidine-one-carboxylate Fe(III) center in ACCO does not support the deprotonation of Fe-bound water due to its low levels of Lewis acidity, thereby enabling fluorine rebound with excellent anion fidelity. Quantitative descriptor analysis further demonstrates that the activation barriers for fluorine atom abstraction and radical rebound are controlled by distinct electronic parameters, revealing a previously unrecognized trend in how the identity of the Fe coordination chemistry modulates individual elementary steps. In particular, fluorine atom abstraction is governed by the intrinsic strength of the Fe(III)–F bond. Stronger *σ*-donor ligands at the nonheme Fe center, such as anionic hydroxy and carboxylate ligands, facilitates fluorine atom abstraction with the Fe(II) enzyme, driven by the formation of a more stable Fe(III)–F bond. In contrast, the reactivity of radical rebound does not strongly correlate with thermodynamic driving force and is instead primarily controlled by the electrophilicity of the Fe(III) intermediate: more electron-deficient Fe(III) centers are more reactive toward radical rebound. Additionally, across all the ferric enzyme model systems examined, the intrinsic C–X (X = F, OH, and N_3_) bond forming radical rebound reactivity follows the trend N_3_ > OH > F, with azidation displaying the lowest activation energy and fluorination the highest. Together, these results establish mechanistic principles for tuning reactivity and chemoselectivity in nonheme Fe-catalyzed radical rebound chemistry. More broadly, this combined experimental and computational study highlights primary coordination-sphere engineering as a powerful yet previously underexplored strategy for reprogramming nonheme enzymatic activity and selectivity, further expanding the toolbox for biocatalytic, enantioselective C(sp^*3*^)–X bond formation via C–H functionalization.

## ASSOCIATED CONTENT

### Supporting Information

The Supporting Information is available free of charge on the ACS Publications website.

Additional experimental and computational results, characterization data, Cartesian coordinates, and energies of DFT-computed structures (PDF)

## AUTHOR INFORMATION

### Author Contributions

#L.Z., R.G. and B.K.M. contributed equally.

### Notes

The authors declare no competing financial interest.

## ACKNOWLEDGMENT

The experimental work is supported by the National Institutes of Health (R35GM147387 to Y.Y.). Computational study is supported by the National Science Foundation (CHE-2247505 to P.L.). Y.Y. is an Alfred P. Sloan Research Fellow (FG-2024-22244), a Camille Dreyfus Teacher-Scholar Awardee (TC-25-084), a David & Lucile Packard Fellow (2023-76169) and a Howard Hughes Medical Institute Freeman Hrabowski Scholar. Computations were carried out at the University of Pittsburgh Center for Research Computing and Data and the Advanced Cyber-infrastructure Coordination Ecosystem: Services & Support (ACCESS) Program, supported by NSF awards OAC-2117681, OAC-1928147, OAC-1928224 and CHE-260005. We thank Dr. Qi Zhou, Dr. Wenzhen Fu, Dr. Shuyun Ju and Chongtao Li in our laboratory (University of California Santa Barbara) for their preliminary studies on fluorine atom transferases.

