## Supporting Information for "Influence of Primary Coordination Sphere on Anion Rebound Selectivity in Nonheme Fe Enzyme-Catalyzed C(sp^3^)–H Functionalization: A Comparative Experimental and Computational Study of EgtB and ACCO"

#### I. General methods

**General.** Unless otherwise noted, all chemicals and reagents were obtained from commercial suppliers (Sigma-Aldrich, VWR, Alfa Aesar, Combi-Blocks, AmBeed, Oakwood, and Enamine) and used without further purification. Silica gel chromatography was carried out using AMD Silica Gel 60, 230-400 mesh. $^1$H and $^{13}$C NMR spectra were recorded on Varian 500, Varian 600, Bruker 400, or Bruker 500 MHz instruments in $CDCl_3$ and are referenced to residual solvent signals. $^{19}$F NMR spectra (where applicable) were recorded on a Bruker 400 MHz. Data for $^1$H NMR are reported as follows: chemical shift (δ: ppm), multiplicity (s = singlet, d = doublet, t = triplet, q = quartet, p = pentet, sext = sextet, m = multiplet, dd = doublet of doublets, dt = doublet of triplets, dq = doublet of quartets, ddd = doublet of doublet of doublets, ddt = doublet of doublet of triplets, brs = broad singlet), coupling constant (Hz), integration. Sonication on a small scale was performed using a BioLogics ultrasonic homogenizer (model 150VT) equipped with a stepped microtip or a Qsonica Q500 sonicator with a 24-tip horn. All IR spectra were recorded on a Thermo Scientific Nicolet iS5 spectrometer (iD5 ATR, diamond). High-resolution mass spectrometry data were obtained at the Mass Spectral Facilities at the University of California Santa Barbara and the University of California Irvine. High-resolution accurate mass (HRAM) ESI data was analyzed on a Waters LCT Premier mass spectrometer with a LEAP PAL autosampler with isocratic MeOH flow (no column; direct injection). Molecular formulas (MF) were validated by lock mass calibration to sodiated polyethylene glycol polymer or monomethyl ether polyethylene glycol polymer standards. High-resolution accurate mass (HRAM) EI data was acquired using a Waters LCT Premier time-of-flight (TOF) mass spectrometer. Masses of positively charged ions were calibrated using a methanol solution of polyethylene glycol or polyethylene glycol monomethyl ether as an internal standard. All samples were dissolved in $CH_2Cl_2$ and were directly infused unless otherwise noted. Synthetic reactions were monitored by thin-layer chromatography (TLC, Silicycle TLG-R10014BK-323 gel plates) using a UV lamp or an appropriate TLC stain for visualization.

Cell cultures were grown in Luria-Bertani medium (LB) and Terrific Broth medium (TB) or Hyper Broth medium (AthenaES) (HB). M9-N minimal medium (M9-N buffer, pH = 7.4) was used as a buffering system for whole-cell and lysate biotransformation. The M9-N buffer contains 47.7 mM $Na_2HPO_4$, 22.0 mM $KH_2PO_4$, 8.6 mM NaCl, 2.0 mM $MgSO_4$, and 0.1 mM $CaCl_2$. The extraction solution was prepared by adding 0.5 mmol mesitylene to 500 mL diisopropyl ether (*i*-$Pr_2O$). Diisopropyl ether was used as the extraction solvent, as conventional extraction solvent such as EtOAc would co-elude with the *N*-fluoroamide substrate in our HPLC analysis. Our studies showed that the substrates and products are highly soluble in *i*-$Pr_2O$.

**Cloning and site-saturation mutagenesis.** pET-28b(+) was used as the cloning and expression vector for *Mycobacterium thermoresistible* EgtB (*Mth*EgtB) with an *N*-terminal 6×His-tag. The gene encoding *Mth*EgtB was codon-optimized for protein production using *E. coli* as the host organism and purchased as gBlocks from GeneralBiol.[1] The gene was cloned into pET-28b(+) between restriction sites Nde I and Hind III. Nonheme Fe enzymes with a C-terminal 6×His-tag were cloned into pET-22b(+) between Nde I and Xho I. Site-saturation mutagenesis was performed using the "22c-trick" method described by Kille *et al.*[2] PCR primers were ordered from IDT or Azenta (formerly Genewiz). The PCR products were gel purified and ligated using a Gibson mix prepared from 5X isothermal (ISO) reaction buffer (25% PEG-8000, 500 mM Tris-HCl pH 7.5, 50 mM $MgCl_2$, 50 mM DTT, 1 mM each of the dNTPs, and 5 mM NAD), T5 exonuclease, Phusion DNA polymerase, and Taq DNA ligase.[3] The ligation mixture was used directly to transform electrocompetent *E. coli* strain *E. cloni* BL21(DE3) cells (Lucigen).

**Procedure for the directed evolution of EgtB$_{CHF}$ variants**

**General note.** Initially, due to the relatively low total activity of the new-to-nature fluorinase, directed evolution was carried out using 24-well deep-well plates to ensure excellent reproducibility. When the total activity of an intermediate fluorinase variant is sufficiently high, directed evolution was carried out using 96-well deep-well plates to have higher throughput.

**Expression of *Mth*EgtB variants in 24-well plates.** For $EgtB_{CHF1}$, from wt *Mth*EgtB to *Mth*EgtB Y377W W415R R87D T141I, directed evolution was performed in 24-well plates. For $EgtB_{CHF2}$, from wt *Mth*EgtB to *Mth*EgtB Y377W W415R R87D T141M R379A, directed evolution of $EgtB_{CHF2}$ was performed in 24-well plates. Single colonies from $LB_{kan}$ agar plates were picked using sterile toothpicks and cultured in deep-well 96-well plates containing $LB_{kan}$ (400 µL/well) at 37 °C, 250 rpm shaking overnight. $TB_{kan}$ (4.0 mL/well) in four 24-well plates was then inoculated with an aliquot (200 µL/well) of these overnight cultures and allowed to shake for 2.5 h at 37 °C and 230 rpm. The plates were cooled on ice for 20 min before induction with 0.5 mM isopropyl β-D-1-thiogalactopyranoside (IPTG, final concentration). Expression was conducted at 22 °C, 150 rpm for 22 h.

**Whole-cell reaction screening in 24-well plates.** *E. coli* cells in deep-well 24-well plates were then pelleted by centrifugation (4000 rpm, 5 min, 4 °C) using an Eppendorf 5910R tabletop centrifuge. After centrifugation, the supernatant was removed, and the cell pellet was resuspended in M9-N buffer (600 µL) by gentle shaking using a Fisher Scientific microplate shaker (800 rpm, 3 min). The 24-well plate was then transferred into a Coy anaerobic chamber. In the Coy chamber, the *N*-fluoroamide substrate **1** (10 µL, 0.4 M in EtOH) and ferrous ammonium sulfate (10 µL, 40 mM in $ddH_2O$) were added to each well in succession using an Eppendorf Xplorer multichannel pipette. The plates were sealed with an aluminum foil and shaken on a Corning microplate shaker at room temperature and 680 rpm. After 6–18 h, the 24-well plate was taken out of the anaerobic chamber, the seal was removed, and the analytical scale reactions were worked up following the appropriate method below.

**Chiral HPLC analysis.** The reaction mixtures were quenched by the addition of 600 µL extraction solution (0.5 mM mesitylene in diisopropyl ether, a non-UV active polar solvent) using an Eppendorf Xplorer plus, 12-channel, 50–1000 µL electronic pipette. The plate was tightly sealed with a reusable silicone mat, shaken vigorously for 30 times, and centrifuged (4500 rpm, 20 min) to separate the organic and the aqueous layers. The organic layers (350 μL/well) were transferred to 500 μL vial inserts using an Eppendorf Xplorer, 12-channel, 15–300 μL electronic pipette. Chiral HPLC analysis was conducted using a Shimadzu i-series (66 MPa) HPLC with IPA/hexanes as the mobile phase. The Chiralpak IB-N5 (4.6 mm × 25 cm, 5 micron), Chiralpak IC (4.6 mm × 25 cm, 5 micron) and Chiralpak IG (4.6 mm × 25 cm, 5 micron) were used for this normal phase HPLC analysis (*vide infra*).

**Expression of *Mth*EgtB variants in 96-well plates.** For $EgtB_{CHF1}$, from *Mth*EgtB Y377W W415R R87D T141I to the final variant, directed evolution of was performed in 96-well plates. For $EgtB_{CHF2}$, from *Mth*EgtB Y377W W415R R87D T141M R379A to the final variant, directed evolution of was performed in 96-well plates. Single colonies from $LB_{kan}$ agar plates were picked using sterile toothpicks and cultured in deep-well 96-well plates containing $LB_{kan}$ (400 μL/well) at 37 °C, 250 rpm shaking overnight. $TB_{kan}$ (1000 μL/well) in a deep-well 96-well plate was then inoculated with an aliquot (50 μL/well) of these overnight cultures and allowed to shake for 3 h at 37 °C and 250 rpm. The plates were cooled on ice for 20 min before induction with 0.5 mM IPTG (final concentration). Expression was conducted at 22 °C, 220 rpm for 22 h.

**Whole-cell reaction screening in 96-well plates.** *E. coli* cells were then pelleted by centrifugation (4000 rpm, 5 min, 4 °C) using an Eppendorf 5910R tabletop centrifuge. After centrifugation, the supernatant was removed, and the cell pellet was resuspended in M9-N buffer (400 μL) by gently shaking using a Fisher Scientific microplate shaker (800 rpm, 3 min). The 24-well plate was then transferred into a Coy anaerobic chamber. In the Coy chamber, the *N*-fluoroamide substrate **1** (10 μL, 0.4 M in EtOH) and ferrous ammonium sulfate (10 μL, 40 mM in $ddH_2O$) were added in succession using an Eppendorf Xplorer multichannel pipette. The plates were sealed with an aluminum foil and shaken on a Corning microplate shaker at room temperature and 680 rpm. After 6–18 h, the 24-well plate was taken out of the anaerobic chamber, the seal was removed, and the analytical scale reactions were worked up following the method described above.

**Procedure for analytical scale biocatalytic C–H functionalization reactions**

**Expression of *Mth*EgtB variants and other nonheme Fe enzymes in 125 mL Erlenmeyer flasks.** *E. coli* (*E. cloni* BL21(DE3)) cells carrying the *Mth*EgtB plasmid in a pET-28b(+) vector were grown overnight in 4 mL $LB_{kan}$. $TB_{kan}$ (30 mL in a 125 mL Erlenmeyer flask) was then inoculated with 1.5 mL preculture and incubated at 37 °C, 230 rpm for 2.5 h until $OD_{600}$ reached ca. 2.0. The cultures were cooled on ice for 20 min before induction with 0.5 mM IPTG (final concentration). Expression was conducted at 22 °C, 150 rpm for 22 h. Nonheme enzyme cloned into pET-22b(+) were expressed in $TB_{amp}$ in an analogous manner.

**Biocatalytic C–H fluorination using cell-free lysates.** *E. coli* cells were pelleted by centrifugation (3000 rpm, 5 min, 4 °C) using an Eppendorf 5910R tabletop centrifuge. Supernatant was removed and the resulting cell pellet was resuspended in M9-N buffer to $OD_{600}$ = 15–60 (typically 40). *E. coli* cells were lysed by sonication using a BioLogics ultrasonic homogenizer (model 150 V/T) equipped with a stepped microtip (6 min in total, 1 sec on, 1 sec off, 45% amplitude, 2 cycles), samples were carefully submerged in wet ice to avoid overheating. (Note: overheating in the sonication step will lead to a significant loss of enzyme activity.) The resulting lysed solutions were kept on ice until further use.

The cell-free lysate of the nonheme enzyme (500 µL) was added to a 2 mL vial. In a Coy anaerobic chamber, ferrous ammonium sulfate (10 µL, 40 mM in $ddH_2O$), sodium ascorbate (10 µL, 400 mM in M9-N buffer) and the *N*-fluoroamide substrate **1** (10 µL, 0.4 M in EtOH) were added to 500 µL cell-free lysate in a 2 mL vial in succession. The vials were sealed and shaken on a Corning microplate shaker at room temperature and 680 rpm for 20 h. All reactions were run in triplicates and averaged total turnover numbers along with standard deviation were reported.

**Biocatalytic C–H azidation using cell-free lysates.** Cell-free lysates were prepared using the procedure described above. In an anaerobic chamber, ferrous ammonium sulfate (10 µL, 40 mM in $ddH_2O$), sodium ascorbate (10 µL, 400 mM in M9-N buffer), $NaN_3$ (20 µL, 3.2 M in $ddH_2O$)

and the *N*-fluoroamide substrate **1** (10 μL, 0.4 M in EtOH) were added to 500 μL cell-free lysate in a 2 mL vial in succession. The vials were sealed and shaken on a Corning microplate shaker at room temperature and 680 rpm for 20 h. All reactions were run in triplicates.

**Chiral HPLC analysis.** The reaction mixtures were quenched by the addition of 600 μL extraction solution (0.5 mM mesitylene in diisopropyl ether). The vials were vortexed vigorously to ensure good mixing and extraction. The mixture in each vial was transferred to a 1.5 mL microcentrifuge tube, and the layers were separated by centrifugation (15000 rpm, 15 min) using an Eppendorf tabletop centrifuge 5425. The organic layer (350 μL) was transferred to a 500 μL vial insert placed in a 2 mL vial. Chiral HPLC analysis was conducted using a Shimadzu i-series (66 MPa) HPLC with IPA/hexanes as the mobile phase. Chiralpak IB-N5 (4.6 mm × 25 cm, 5 micron), Chiralpak IC (4.6 mm × 25 cm, 5 micron), Chiralpak IG (4.6 mm × 25 cm, 5 micron), and Chiralpak IH (4.6 mm × 25 cm, 5 micron) columns were used for analysis.

**HPLC calibration curve.** Stock solutions of authentic compounds (40 mM in EtOAc) were prepared. Aliquots (120 μL, 100 μL, 80 μL, 40 μL, 20 μL, 10 μL) were transferred to individual vials, and EtOAc was removed under vacuum. Note: In our HPLC analysis, EtOAc co-eluded with the *N*-fluoroamide substrate **1** and complicated the analysis. As a result, EtOAc was removed from samples used for calibration curve. To each vial were added 500 μL M9-N buffer and 600 μL extraction solution (0.5 mM mesitylene in diisopropyl ether). The mixture was vortexed and transferred to a microcentrifuge tube. After centrifugation (15000 rpm, 15 min) using an Eppendorf tabletop centrifuge 5425, the organic layer (350 μL) was transferred to a 500 μL vial insert placed in a 2 mL vial. Chiral HPLC analysis was conducted using a Shimadzu i-series (66 MPa) HPLC with IPA/hexanes as the mobile phase. Chiralpak IB-N5 (4.6 mm × 25 cm, 5 micron), Chiralpak IC (4.6 mm × 25 cm, 5 micron) and Chiralpak IG (4.6 mm × 25 cm, 5 micron) columns were used for analysis.

**Determination of nonheme enzyme concentration by SDS-PAGE analysis.** SDS-PAGE analysis was performed using a Bio-Rad Mini-PROTEAN Tetra cell powered by a Bio-Rad Power Pac Basic power supply. The lysate was diluted by 40 times in M9-N buffer and centrifuged (15,000 rpm, 10 min, 4 °C) using an Eppendorf tabletop centrifuge 5424R. The resulting supernatant was used for SDS-PAGE analysis. A stock solution of bovine serum albumin (BSA, 1.0 mg/mL in M9-N buffer) was freshly prepared. A series of aliquots (10 –100 µL) of the BSA stock solution was diluted to 1000 µL in M9-N buffer as the standard solution for SDS-PAGE analysis. The nonheme enzyme samples or BSA standards (30 µL) mixed with 10 µL Bio-Rad 4× Laemmli sample buffer were incubated at 90 °C for 10 min in a Bio-Rad T100 Thermal Cycler and then cooled to room temperature. (Note: incubation in a thermalcycler with a heated lid was found to be essential to ensure excellent reproducibility. Incubation in a thermal mixer without a heated lid will lead to water condensation on PCR tube caps, resulting in a change of protein concentration.)

10 µL denatured protein sample was loaded into the wells of a Bio-Rad Mini-PROTEAN TGX Stain-Free gel. 5 µL Bio-Rad Precision Plus Protein All Blue Standards was used as the protein marker in electrophoresis. Upon the completion, the SDS-PAGE gel was stained by Coomassie blue by microwaving the gel in Bio-Rad Coomassie Brilliant Blue R-250 Staining Solution for 30 s. The gel was then destained in a destaining solution (AcOH/MeOH/dd$H_2O$ = 1:2:7) on a rocker overnight. The gel image was acquired using a Bio-Rad Gel Doc EZ Imager, and the protein bands were analyzed by the Bio-Rad Gel Doc™ EZ System. A calibration curve was generated from the BSA standards on the same gel to determine the concentrations of *Mth*EgtB and other nonheme Fe enzymes.

#### II. Evaluation of Fe-dependent proteins

**Table S1.** Detailed information of Fe-dependent proteins descripted in the manuscript.

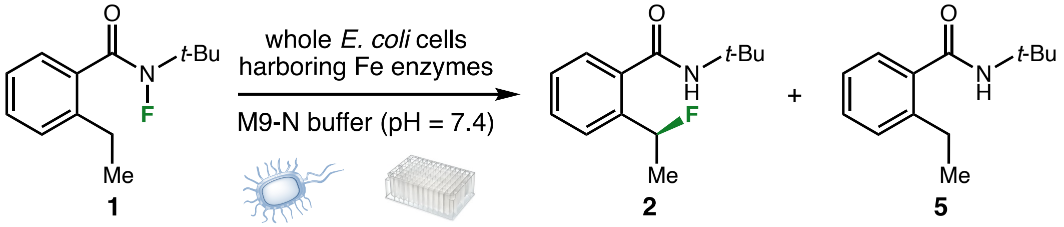

| metallo protein | UniProt ID | organism | annotation | mutations |
|---|---|---|---|---|
| **BesD** | G8XHD5 | *Streptantibioticus cattleyicolor* | 2OG-dependent halogenase | none |
| **EvdO2** | A0A0M3KL01 | *Micromonospora carbonacea* | everninomicin dioxygenase | none |
| **SadA D157G** | Q0B2N4 | *Burkholderia ambifaria* | 2OG-dependent halogenase | D157G |
| **WelO5** | A0A067YX61 | *Hapalosiphon welwitschii* | 2OG-dependent halogenase | none |
| **EgtB** | G7CFI3 | *Mycobacterium thermoresistibile* | Ergothioneine-biosynthetic sulfoxide synthase | none |
| **ACCO** | Q08506 | *Petunia hybrida* | 1-aminocyclopropane-1-carboxylate oxidase | |
| **IPNS** | P05326 | *Emericella nidulans* | isopenicillin N synthase | |
| **MPC** | P06622 | *Pseudomonas putida* | metapyrocatechase | none |

#### III. Protein and DNA sequences

##### Protein sequences

##### *Mth*EgtB

MGSSHHHHHHSSGLVPRGSHMTGVAVPHRAELARQLIDARNRTLRLVDFDDAELRRQYDPLMSP
LVWDLAHIGQQEELWLLRGGDPRRPGLLEPAVEQLYDAFVHPRASRVHLPLLSPAQARRFCATV
RSAVLDALDRLPEDADTFAFGMVVSHEHQHDETMLQALNLRSGEPLLGSGTALPPGRPGVAGTS
VLVPGGPFVLGVDLADEPYALDNERPAHVVDVPAFRIGRVPVTNAEWRAFIDDGGYRQRRWWSD
AGWAYRCEAGLTAPQFWNPDGTRTRFGHVEDIPPDEPVQHVTYFEAEAYAAWAGARLPTEIEWE
KACAWDPATGRRRRYPWGDAAPTAALANLGGDALRPAPVGAYPAGASACGAEQMLGDVWEWTSS
PLRPWPGFTPMIYQRYSQPFFEGAGSGDYRVLRGGSWAVAADILRPSFRNWDHPIRRQIFAGVR
LAWDVDRQTARPGPVGGC*

##### *Mth*EgtB Y377W W415R R87D T141I F83Y R379E V84R H417L (EgtB$_{\mathrm{CHF1}}$)

MGSSHHHHHHSSGLVPRGSHMTGVAVPHRAELARQLIDARNRTLRLVDFDDAELRRQYDPLMSP
LVWDLAHIGQQEELWLLRGGDPRRPGLLEPAVEQLYDA**YR**HP**D**ASRVHLPLLSPAQARRFCATV
RSAVLDALDRLPEDADTFAFGMVVSHEHQHDE**I**MLQALNLRSGEPLLGSGTALPPGRPGVAGTS
VLVPGGPFVLGVDLADEPYALDNERPAHVVDVPAFRIGRVPVTNAEWRAFIDDGGYRQRRWWSD
AGWAYRCEAGLTAPQFWNPDGTRTRFGHVEDIPPDEPVQHVTYFEAEAYAAWAGARLPTEIEWE
KACAWDPATGRRRRYPWGDAAPTAALANLGGDALRPAPVGAYPAGASACGAEQMLGDVWEWTSS
PLRPWPGFTPMI**W**Q**E**YSQPFFEGAGSGDYRVLRGGSWAVAADILRPSFRN**R**D**L**PIRRQIFAGVR
LAWDVDRQTARPGPVGGC*

##### *Mth*EgtB Y377W W415R R87D T141I F83Y R379E V84R H417L H138A (EgtB$_{\mathrm{CHF1}}$ H138A)

MGSSHHHHHHSSGLVPRGSHMTGVAVPHRAELARQLIDARNRTLRLVDFDDAELRRQYDPLMSP
LVWDLAHIGQQEELWLLRGGDPRRPGLLEPAVEQLYDA**YR**HP**D**ASRVHLPLLSPAQARRFCATV
RSAVLDALDRLPEDADTFAFGMVVSHEHQ**A**DE**I**MLQALNLRSGEPLLGSGTALPPGRPGVAGTS

VLVPGGPFVLGVDLADEPYALDNERPAHVVDVPAFRIGRVPVTNAEWRAFIDDGGYRQRRWWSD
AGWAYRCEAGLTAPQFWNPDGTRTRFGHVEDIPPDEPVQHVTYFEAEAYAAWAGARLPTEIEWE
KACAWDPATGRRRRYPWGDAAPTAALANLGGDALRPAPVGAYPAGASACGAEQMLGDVWEWTSS
PLRPWPGFTPMI**W**Q**E**YSQPFFEGAGSGDYRVLRGGSWAVAADILRPSFRN**R**D**L**PIRRQIFAGVR
LAWDVDRQTARPGPVGGC*

***Mth*EgtB Y377W W415R R87D T141M R379A A145K Q422D (EgtB$_{CHF2}$)**

MGSSHHHHHHSSGLVPRGSHMTGVAVPHRAELARQLIDARNRTLRLVDFDDAELRRQYDPLMSP
LVWDLAHIGQQEELWLLRGGDPRRPGLLEPAVEQLYDAFVHP**D**ASRVHLPLLSPAQARRFCATV
RSAVLDALDRLPEDADTFAFGMVVSHEHQHDE**M**MLQ**K**LNLRSGEPLLGSGTALPPGRPGVAGTS
VLVPGGPFVLGVDLADEPYALDNERPAHVVDVPAFRIGRVPVTNAEWRAFIDDGGYRQRRWWSD
AGWAYRCEAGLTAPQFWNPDGTRTRFGHVEDIPPDEPVQHVTYFEAEAYAAWAGARLPTEIEWE
KACAWDPATGRRRRYPWGDAAPTAALANLGGDALRPAPVGAYPAGASACGAEQMLGDVWEWTSS
PLRPWPGFTPMI**W**Q**A**YSQPFFEGAGSGDYRVLRGGSWAVAADILRPSFRN**R**DHPIRR**D**IFAGVR
LAWDVDRQTARPGPVGGC*

***Mth*EgtB Y377W W415R R87D T141M R379A A145K Q422D H138A (EgtB$_{CHF2}$ H138A)**

MGSSHHHHHHSSGLVPRGSHMTGVAVPHRAELARQLIDARNRTLRLVDFDDAELRRQYDPLMSP
LVWDLAHIGQQEELWLLRGGDPRRPGLLEPAVEQLYDAFVHP**D**ASRVHLPLLSPAQARRFCATV
RSAVLDALDRLPEDADTFAFGMVVSHEHQ**A**DE**M**MLQ**K**LNLRSGEPLLGSGTALPPGRPGVAGTS
VLVPGGPFVLGVDLADEPYALDNERPAHVVDVPAFRIGRVPVTNAEWRAFIDDGGYRQRRWWSD
AGWAYRCEAGLTAPQFWNPDGTRTRFGHVEDIPPDEPVQHVTYFEAEAYAAWAGARLPTEIEWE
KACAWDPATGRRRRYPWGDAAPTAALANLGGDALRPAPVGAYPAGASACGAEQMLGDVWEWTSS
PLRPWPGFTPMI**W**Q**A**YSQPFFEGAGSGDYRVLRGGSWAVAADILRPSFRN**R**DHPIRR**D**IFAGVR
LAWDVDRQTARPGPVGGC*

**ACCO I184A K158I F91L K172Y K93Q T89A (ACCO$_{CHF}$):**

MGSSHHHHHHSSGLVPRGSHMENFPIISLDKVNGVERAATMEMIKDACENWGFFELVNHGIPRE
VMDTVEKMTKGHYKKCMEQRFKELVASKALEGVQAEVTDMDWES**A**F**L**L**Q**HLPISNISEVPDLDE
EYREVMRDFAKRLEKLAEELLDLLCENLGLEKGYLKNAFYGSKGPNFGT**I**VSNYPPCPKPDLI**Y**
GLRAHTDAGGI**A**LLFQDDKVSGLQLLKDGQWIDVPPMRHSIVVNLGDQLEVITNGKYKSVMHRV
IAQKDGARMSLASFYNPGSDAVIYPAPALVEKEAEENKQVYPKFVFDDYMKLYAGLKFQAKEPR
FEAMKAMETDVKMDPIATV*

**DNA sequences**

***Mth*EgtB**

ATGGGCAGCAGCCATCATCATCATCATCACAGCAGCGGCCTGGTGCCGCGCGGCAGCCATATGA
CTGGAGTAGCAGTGCCCCACCGTGCAGAATTGGCCCGTCAGTTAATTGACGCTCGCAATCGTAC
CCTGCGTCTTGTTGATTTCGATGACGCCGAGCTGCGCCGCCAGTATGACCCTCTTATGAGCCCC
TTGGTATGGGACTTGGCTCATATTGGACAGCAGGAGGAGTTATGGTTGCTTCGTGGAGGCGACC
CCCGTCGCCCAGGGCTGTTAGAGCCAGCAGTTGAGCAGTTATACGACGCATTCGTACATCCACG
TGCCTCCCGCGTCCACCTTCCGTTGTTATCTCCCGCTCAAGCCCGCCGCTTCTGTGCTACCGTG
CGTAGCGCGGTTCTGGACGCTTTAGACCGTCTTCCTGAAGATGCCGATACGTTCGCCTTCGGGA
TGGTCGTCTCGCACGAACACCAGCACGATGAGACGATGTTGCAAGCGCTGAATTTACGCTCAGG
TGAGCCCTTGCTTGGGTCTGGGACGGCGTTACCTCCGGGTCGCCCTGGCGTCGCGGGCACAAGC
GTTTTAGTACCTGGAGGTCCGTTTGTGCTGGGTGTCGATTTGGCTGACGAGCCATACGCATTAG
ATAACGAACGTCCGGCCCACGTAGTAGATGTGCCAGCGTTCCGTATCGGTCGCGTGCCAGTGAC
AAACGCTGAGTGGCGTGCATTTATCGACGATGGTGGATATCGCCAACGTCGTTGGTGGTCGGAT
GCTGGTTGGGCTTATCGCTGTGAAGCCGGGCTTACCGCCCCGCAGTTCTGGAATCCCGATGGCA
CTCGTACCCGCTTCGGACATGTAGAGGATATTCCCCCCGATGAGCCAGTTCAACACGTCACTTA
TTTTGAAGCAGAGGCTTATGCAGCATGGGCAGGGGCGCGCTTACCCACGGAAATTGAGTGGGAG
AAGGCTTGCGCCTGGGACCCTGCCACAGGTCGTCGCCGCCGCTACCCCTGGGGAGACGCGGCTC
CCACTGCGGCACTTGCGAATCTTGGTGGCGACGCGTTGCGTCCAGCGCCTGTAGGAGCCTATCC
TGCGGGGGCAAGCGCGTGTGGAGCTGAGCAGATGTTGGGGGATGTTTGGGAATGGACTTCAAGT

CCGCTGCGCCCATGGCCGGGGTTTACACCGATGATCTATCAGCGCTACTCTCAACCATTTTTTG

AGGGAGCAGGCAGCGGAGACTACCGTGTATTGCGTGGGGGCAGTTGGGCAGTCGCTGCTGACAT

CCTTCGTCCGTCATTCCGCAATTGGGACCACCCCATTCGTCGTCAAATTTTCGCAGGGGTCCGT

CTGGCGTGGGACGTTGACCGTCAGACGGCTCGTCCCGGACCAGTGGGCGGATGTTGA

**_Mth_EgtB Y377W W415R R87D T141I F83Y R379E V84R H417L (EgtB$_{CHF1}$)**

ATGGGCAGCAGCCATCATCATCATCATCACAGCAGCGGCCTGGTGCCGCGCGGCAGCCATATGA

CTGGAGTAGCAGTGCCCCACCGTGCAGAATTGGCCCGTCAGTTAATTGACGCTCGCAATCGTAC

CCTGCGTCTTGTTGATTTCGATGACGCCGAGCTGCGCCGCCAGTATGACCCTCTTATGAGCCCC

TTGGTATGGGACTTGGCTCATATTGGACAGCAGGAGGAGTTATGGTTGCTTCGTGGAGGCGACC

CCCGTCGCCCAGGGCTGTTAGAGCCAGCAGTTGAGCAGTTATACGACGCA**TATCGT**CATCCA**GA**

**T**GCCTCCCGCGTCCACCTTCCGTTGTTATCTCCCGCTCAAGCCCGCCGCTTCTGTGCTACCGTG

CGTAGCGCGGTTCTGGACGCTTTAGACCGTCTTCCTGAAGATGCCGATACGTTCGCCTTCGGGA

TGGTCGTCTCGCACGAACACCAGCACGATGAG**ATT**ATGTTGCAAGCGCTGAATTTACGCTCAGG

TGAGCCCTTGCTTGGGTCTGGGACGGCGTTACCTCCGGGTCGCCCTGGCGTCGCGGGCACAAGC

GTTTTAGTACCTGGAGGTCCGTTTGTGCTGGGTGTCGATTTGGCTGACGAGCCATACGCATTAG

ATAACGAACGTCCGGCCCACGTAGTAGATGTGCCAGCGTTCCGTATCGGTCGCGTGCCAGTGAC

AAACGCTGAGTGGCGTGCATTTATCGACGATGGTGGATATCGCCAACGTCGTTGGTGGTCGGAT

GCTGGTTGGGCTTATCGCTGTGAAGCCGGGCTTACCGCCCCGCAGTTCTGGAATCCCGATGGCA

CTCGTACCCGCTTCGGACATGTAGAGGATATTCCCCCCGATGAGCCAGTTCAACACGTCACTTA

TTTTGAAGCAGAGGCTTATGCAGCATGGGCAGGGGCGCGCTTACCCACGGAAATTGAGTGGGAG

AAGGCTTGCGCCTGGGACCCTGCCACAGGTCGTCGCCGCCGCTACCCCTGGGGAGACGCGGCTC

CCACTGCGGCACTTGCGAATCTTGGTGGCGACGCGTTGCGTCCAGCGCCTGTAGGAGCCTATCC

TGCGGGGGCAAGCGCGTGTGGAGCTGAGCAGATGTTGGGGGATGTTTGGGAATGGACTTCAAGT

CCGCTGCGCCCATGGCCGGGGTTTACACCGATGATC**TGG**CAG**GAG**TACTCTCAACCATTTTTTG

AGGGAGCAGGCAGCGGAGACTACCGTGTATTGCGTGGGGGCAGTTGGGCAGTCGCTGCTGACAT

CCTTCGTCCGTCATTCCGCAAT**CGT**GAC**CTG**CCCATTCGTCGTCAAATTTTCGCAGGGGTCCGT
CTGGCGTGGGACGTTGACCGTCAGACGGCTCGTCCCGGACCAGTGGGCGGATGTTGA

***Mth*EgtB Y377W W415R R87D T141M R379A A145K Q422D (EgtB$_{CHF2}$)**

ATGGGCAGCAGCCATCATCATCATCATCACAGCAGCGGCCTGGTGCCGCGCGGCAGCCATATGA
CTGGAGTAGCAGTGCCCCACCGTGCAGAATTGGCCCGTCAGTTAATTGACGCTCGCAATCGTAC
CCTGCGTCTTGTTGATTTCGATGACGCCGAGCTGCGCCGCCAGTATGACCCTCTTATGAGCCCC
TTGGTATGGGACTTGGCTCATATTGGACAGCAGGAGGAGTTATGGTTGCTTCGTGGAGGCGACC
CCCGTCGCCCAGGGCTGTTAGAGCCAGCAGTTGAGCAGTTATACGACGCATTCGTACATCCA**GA**
**T**GCCTCCCGCGTCCACCTTCCGTTGTTATCTCCCGCTCAAGCCCGCCGCTTCTGTGCTACCGTG
CGTAGCGCGGTTCTGGACGCTTTAGACCGTCTTCCTGAAGATGCCGATACGTTCGCCTTCGGGA
TGGTCGTCTCGCACGAACACCAGCACGATGAG**ATG**ATGTTGCAA**AAG**CTGAATTTACGCTCAGG
TGAGCCCTTGCTTGGGTCTGGGACGGCGTTACCTCCGGGTCGCCCTGGCGTCGCGGGCACAAGC
GTTTTAGTACCTGGAGGTCCGTTTGTGCTGGGTGTCGATTTGGCTGACGAGCCATACGCATTAG
ATAACGAACGTCCGGCCCACGTAGTAGATGTGCCAGCGTTCCGTATCGGTCGCGTGCCAGTGAC
AAACGCTGAGTGGCGTGCATTTATCGACGATGGTGGATATCGCCAACGTCGTTGGTGGTCGGAT
GCTGGTTGGGCTTATCGCTGTGAAGCCGGGCTTACCGCCCCGCAGTTCTGGAATCCCGATGGCA
CTCGTACCCGCTTCGGACATGTAGAGGATATTCCCCCCGATGAGCCAGTTCAACACGTCACTTA
TTTTGAAGCAGAGGCTTATGCAGCATGGGCAGGGGCGCGCTTACCCACGGAAATTGAGTGGGAG
AAGGCTTGCGCCTGGGACCCTGCCACAGGTCGTCGCCGCCGCTACCCCTGGGGAGACGCGGCTC
CCACTGCGGCACTTGCGAATCTTGGTGGCGACGCGTTGCGTCCAGCGCCTGTAGGAGCCTATCC
TGCGGGGGCAAGCGCGTGTGGAGCTGAGCAGATGTTGGGGGATGTTTGGGAATGGACTTCAAGT
CCGCTGCGCCCATGGCCGGGGTTTACACCGATGATC**TGG**CAG**GCG**TACTCTCAACCATTTTTTG
AGGGAGCAGGCAGCGGAGACTACCGTGTATTGCGTGGGGGCAGTTGGGCAGTCGCTGCTGACAT
CCTTCGTCCGTCATTCCGCAAT**CGT**GACCACCCCATTCGTCGT**GAT**ATTTTCGCAGGGGTCCGT
CTGGCGTGGGACGTTGACCGTCAGACGGCTCGTCCCGGACCAGTGGGCGGATGTTGA

**_Mth_EgtB Y377W W415R R87D T141I F83Y R379E V84R H417L H138A (EgtB$_{CHF1}$ H138A)**

ATGGGCAGCAGCCATCATCATCATCATCACAGCAGCGGCCTGGTGCCGCGCGGCAGCCATATGA
CTGGAGTAGCAGTGCCCCACCGTGCAGAATTGGCCCGTCAGTTAATTGACGCTCGCAATCGTAC
CCTGCGTCTTGTTGATTTCGATGACGCCGAGCTGCGCCGCCAGTATGACCCTCTTATGAGCCCC
TTGGTATGGGACTTGGCTCATATTGGACAGCAGGAGGAGTTATGGTTGCTTCGTGGAGGCGACC
CCCGTCGCCCAGGGCTGTTAGAGCCAGCAGTTGAGCAGTTATACGACGCA**TATCGT**CATCCA**GA**
**T**GCCTCCCGCGTCCACCTTCCGTTGTTATCTCCCGCTCAAGCCCGCCGCTTCTGTGCTACCGTG
CGTAGCGCGGTTCTGGACGCTTTAGACCGTCTTCCTGAAGATGCCGATACGTTCGCCTTCGGGA
TGGTCGTCTCGCACGAACACCAG**GCG**GATGAG**ATT**ATGTTGCAAGCGCTGAATTTACGCTCAGG
TGAGCCCTTGCTTGGGTCTGGGACGGCGTTACCTCCGGGTCGCCCTGGCGTCGCGGGCACAAGC
GTTTTAGTACCTGGAGGTCCGTTTGTGCTGGGTGTCGATTTGGCTGACGAGCCATACGCATTAG
ATAACGAACGTCCGGCCCACGTAGTAGATGTGCCAGCGTTCCGTATCGGTCGCGTGCCAGTGAC
AAACGCTGAGTGGCGTGCATTTATCGACGATGGTGGATATCGCCAACGTCGTTGGTGGTCGGAT
GCTGGTTGGGCTTATCGCTGTGAAGCCGGGCTTACCGCCCCGCAGTTCTGGAATCCCGATGGCA
CTCGTACCCGCTTCGGACATGTAGAGGATATTCCCCCCGATGAGCCAGTTCAACACGTCACTTA
TTTTGAAGCAGAGGCTTATGCAGCATGGGCAGGGGCGCGCTTACCCACGGAAATTGAGTGGGAG
AAGGCTTGCGCCTGGGACCCTGCCACAGGTCGTCGCCGCCGCTACCCCTGGGGAGACGCGGCTC
CCACTGCGGCACTTGCGAATCTTGGTGGCGACGCGTTGCGTCCAGCGCCTGTAGGAGCCTATCC
TGCGGGGGCAAGCGCGTGTGGAGCTGAGCAGATGTTGGGGGATGTTTGGGAATGGACTTCAAGT
CCGCTGCGCCCATGGCCGGGGTTTACACCGATGATC**TGG**CAG**GAG**TACTCTCAACCATTTTTTG
AGGGAGCAGGCAGCGGAGACTACCGTGTATTGCGTGGGGGCAGTTGGGCAGTCGCTGCTGACAT
CCTTCGTCCGTCATTCCGCAAT**CGT**GAC**CTG**CCCATTCGTCGTCAAATTTTCGCAGGGGTCCGT
CTGGCGTGGGACGTTGACCGTCAGACGGCTCGTCCCGGACCAGTGGGCGGATGTTGA

**_Mth_EgtB Y377W W415R R87D T141M R379A A145K Q422D H138A (EgtB$_{CHF2}$ H138A)**

ATGGGCAGCAGCCATCATCATCATCATCACAGCAGCGGCCTGGTGCCGCGCGGCAGCCATATGA
CTGGAGTAGCAGTGCCCCACCGTGCAGAATTGGCCCGTCAGTTAATTGACGCTCGCAATCGTAC
CCTGCGTCTTGTTGATTTCGATGACGCCGAGCTGCGCCGCCAGTATGACCCTCTTATGAGCCCC
TTGGTATGGGACTTGGCTCATATTGGACAGCAGGAGGAGTTATGGTTGCTTCGTGGAGGCGACC
CCCGTCGCCCAGGGCTGTTAGAGCCAGCAGTTGAGCAGTTATACGACGCATTCGTACATCCA**GA**
**T**GCCTCCCGCGTCCACCTTCCGTTGTTATCTCCCGCTCAAGCCCGCCGCTTCTGTGCTACCGTG
CGTAGCGCGGTTCTGGACGCTTTAGACCGTCTTCCTGAAGATGCCGATACGTTCGCCTTCGGGA
TGGTCGTCTCGCACGAACACCAG**GCG**GATGAG**ATG**ATGTTGCAA**AAG**CTGAATTTACGCTCAGG
TGAGCCCTTGCTTGGGTCTGGGACGGCGTTACCTCCGGGTCGCCCTGGCGTCGCGGGCACAAGC
GTTTTAGTACCTGGAGGTCCGTTTGTGCTGGGTGTCGATTTGGCTGACGAGCCATACGCATTAG
ATAACGAACGTCCGGCCCACGTAGTAGATGTGCCAGCGTTCCGTATCGGTCGCGTGCCAGTGAC
AAACGCTGAGTGGCGTGCATTTATCGACGATGGTGGATATCGCCAACGTCGTTGGTGGTCGGAT
GCTGGTTGGGCTTATCGCTGTGAAGCCGGGCTTACCGCCCCGCAGTTCTGGAATCCCGATGGCA
CTCGTACCCGCTTCGGACATGTAGAGGATATTCCCCCCGATGAGCCAGTTCAACACGTCACTTA
TTTTGAAGCAGAGGCTTATGCAGCATGGGCAGGGGCGCGCTTACCCACGGAAATTGAGTGGGAG
AAGGCTTGCGCCTGGGACCCTGCCACAGGTCGTCGCCGCCGCTACCCCTGGGGAGACGCGGCTC
CCACTGCGGCACTTGCGAATCTTGGTGGCGACGCGTTGCGTCCAGCGCCTGTAGGAGCCTATCC
TGCGGGGGCAAGCGCGTGTGGAGCTGAGCAGATGTTGGGGGATGTTTGGGAATGGACTTCAAGT
CCGCTGCGCCCATGGCCGGGGTTTACACCGATGATC**TGG**CAG**GCG**TACTCTCAACCATTTTTTG
AGGGAGCAGGCAGCGGAGACTACCGTGTATTGCGTGGGGGCAGTTGGGCAGTCGCTGCTGACAT
CCTTCGTCCGTCATTCCGCAAT**CGT**GACCACCCCATTCGTCGT**GAT**ATTTTCGCAGGGGTCCGT
CTGGCGTGGGACGTTGACCGTCAGACGGCTCGTCCCGGACCAGTGGGCGGATGTTGA

**ACCO I184A K158I F91L K172Y K93Q T89A (ACCO$_{CHF}$)**

ATGGGCAGCAGCCATCATCATCATCATCACAGCAGCGGCCTGGTGCCGCGCGGCAGCCATATGG
AAAACTTCCCGATTATTAGTCTGGATAAAGTTAATGGTGTGGAACGCGCCGCCACAATGGAAAT
GATTAAGGATGCCTGTGAAAATTGGGGCTTTTTCGAACTGGTTAATCATGGCATTCCGCGTGAA

GTGATGGATACCGTGGAAAAAATGACCAAAGGCCATTATAAAAAGTGCATGGAACAGCGTTT**CT**

**G**AGAACTGGTTGCAAGTAAAGCCCTGGAAGGCGTGCAGGCAGAAGTTACCGATATGGATTGGGA

AAGC**GCG**TTTTTCCTG**CAG**CATCTGCCGATTAGTAATATTAGCGAAGTGCCGGATCTGGATGAA

GAATATCGTGAAGTGATGCGCGATTTTGCAAAACGTCTGGAAAAACTGGCCGAAGAACTGCTGG

ATCTGCTGTGTGAAAATCTGGGCCTGGAAAAAGGTTATCTGAAAAATGCATTTTACGGTAGTAA

GGGTCCGAATTTTGGTACA**ATT**GTTAGCAATTATCCGCCGTGTCCGAAACCGGATCTGATT**TAT**

GGTCTGCGTGCACATACCGATGCAGGTGGTATT**GCG**CTGCTGTTTCAGGATGATAAAGTGAGCG

GCCTGCAGCTGCTGAAAGATGGTCAGTGGATTGATGTTCCGCCGATGCGCCATAGTATTGTTGT

TAATCTGGGTGACCAGCTGGAAGTTATTACCAATGGCAAATATAAAAGCGTTATGCATCGTGTG

ATTGCCCAGAAAGATGGCGCACGCATGAGTCTGGCCAGTTTTTATAATCCGGGCAGCGATGCCG

TTATCTATCCGGCCCCGGCACTGGTTGAAAAAGAAGCCGAAGAAAATAAGCAGGTTTATCCGAA

ATTTGTGTTTGATGATTACATGAAACTGTACGCCGGCCTGAAATTTCAGGCAAAAGAACCGCGC

TTTGAAGCAATGAAAGCAATGGAAACCGATGTTAAAATGGACCCTATTGCAACCGTGTAA

**Table S2.** Primers used for the directed evolution of $EgtB_{CHF}$ and $ACCO_{CHF}$ via site-saturation mutagenesis.

| DNA Template | name | 5'-3' primer sequences |
|---|---|---|
| pET-28b(+) | HindIII_fwd | GAAAGCTTGCGGCCGCACTCGAG |
| | HindIII_rev | CTCGAGTGCGGCCGCAAGCTTTCAACATCCGCCCACTGGTCC |
| | NdeI_fwd | GTGCCGCGCGGCAGCCATATGACTGGAGTAGCAGTGCCCCAC |
| | NdeI_rev | CATATGGCTGCCGCGCGGCACCAG |
| *Mth*EgtB | Y377X_NDT | CATGGCCGGGGTTTACACCGATGATC**NDT**CAGCGCTACTC |
| | Y377X_VHG | CATGGCCGGGGTTTACACCGATGATC**VHG**CAGCGCTACTC |
| | Y377X_TGG | CATGGCCGGGGTTTACACCGATGATC**TGG**CAGCGCTACTC |
| | Y377X_rev | GATCATCGGTGTAAACCCCGGCCATG |
| *Mth*EgtB Y377W | W415X_NDT | CTTCGTCCGTCATTCCGCAAT**NDT**GACCACCCCATTC |
| | W415X_VHG | CTTCGTCCGTCATTCCGCAAT**VHG**GACCACCCCATTC |
| | W415X_rev | TTGCGGAATGACGGACGAAGGATGTCAG |
| *Mth*EgtB Y377W W415R | R87X_NDT | ATACGACGCATTCGTACATCCA**NDT**GCCTCCCGCGTC |
| | R87X_VHG | ATACGACGCATTCGTACATCCA**VHG**GCCTCCCGCGTC |
| | R87X_TGG | ATACGACGCATTCGTACATCCA**TGG**GCCTCCCGCGTC |
| | R87X_rev | GATGTACGAATGCGTCGTATAACTGCTCAAC |
| *Mth*EgtB Y377W W415R R87D | T141X_NDT | CACGAACACCAGCACGATGAG**NDT**ATGTTGCAAGC |
| | T141X_VHG | CACGAACACCAGCACGATGAG**VHG**ATGTTGCAAGC |
| | T141X_TGG | CACGAACACCAGCACGATGAG**TGG**ATGTTGCAAGC |
| | T141X_rev | CTCATCGTGCTGGTGTTCGTGCGAGACG |
| *Mth*EgtB Y377W W415R R87D T141I | F83X_NDT | CAGTTGAGCAGTTATACGACGCA**NDT**GTACATCCAC |
| | F83X_VHG | CAGTTGAGCAGTTATACGACGCA**VHG**GTACATCCAC |
| | F83X_TGG | CAGTTGAGCAGTTATACGACGCA**TGG**GTACATCCAC |
| | F83X_rev | GTCGTATAACTGCTCAACTGCTGGCTC |

| | | |
|---|---|---|
| *Mth*EgtB Y377W W415R R87D T141I F83Y | R379X_NDT | GTTTACACCGATGATCTGGCAG**NDT**TACTCTCAAC |
| | R379X_VHG | GTTTACACCGATGATCTGGCAG**VHG**TACTCTCAAC |
| | R379X_TGG | GTTTACACCGATGATCTGGCAG**TGG**TACTCTCAAC |
| | R379X_rev | CAGATCATCGGTGTAAACCCCGGCCATG |
| *Mth*EgtB Y377W W415R R87D T141I F83Y R379E | V84X_NDT | CAGTTGAGCAGTTATACGACGCATAT**NDT**CATCCAGATG |
| | V84X_VHG | CAGTTGAGCAGTTATACGACGCATAT**VHG**CATCCAGATG |
| | V84X_TGG | CAGTTGAGCAGTTATACGACGCATAT**TGG**CATCCAGATG |
| | V84X_rev | CGTCGTATAACTGCTCAACTGCTG |
| *Mth*EgtB Y377W W415R R87D T141I F83Y R379E V84R | H417X_NDT | CGTCCGTCATTCCGCAATCGTGAC**NDT**CCCATTCG |
| | H417X_VHG | CGTCCGTCATTCCGCAATCGTGAC**VHG**CCCATTCG |
| | H417X_TGG | CGTCCGTCATTCCGCAATCGTGAC**TGG**CCCATTCG |
| | H417X_rev | GATTGCGGAATGACGGACGAAGGATGTC |
| *Mth*EgtB Y377W W415R R87D T141M | R379X_NDT | GTTTACACCGATGATCTGGCAG**NDT**TACTCTCAAC |
| | R379X_VHG | GTTTACACCGATGATCTGGCAG**VHG**TACTCTCAAC |
| | R379X_TGG | GTTTACACCGATGATCTGGCAG**TGG**TACTCTCAAC |
| | R379X_rev | CAGATCATCGGTGTAAACCCCGGCCATG |
| *Mth*EgtB Y377W W415R R87D T141M R379A | A145X_NDT | CGATGAGATGATGTTGCAA**NDT**CTGAATTTAC |
| | A145X_VHG | CGATGAGATGATGTTGCAA**VHG**CTGAATTTAC |
| | A145X_TGG | CGATGAGATGATGTTGCAA**TGG**CTGAATTTAC |
| | A145X_rev | TTGCAACATCATCTCATCGTGCTGGTG |
| *Mth*EgtB Y377W W415R R87D T141M R379A A145K | Q422X_NDT | CGTGACCACCCCATTCGTCGT**NDT**ATTTTCGCAG |
| | Q422X_VHG | CGTGACCACCCCATTCGTCGT**VHG**ATTTTCGCAG |
| | Q422X_TGG | CGTGACCACCCCATTCGTCGT**TGG**ATTTTCGCAG |
| | Q422X_rev | ACGACGAATGGGGTGGTCACGATTG |

| | | |
|---|---|---|
| *Mth*EgtB Y377W W415R R87D T141I F83Y R379E V84R H417L (*Mth*EgtB$_{CHF1}$) | H51X_rev | AGCCAAGTCCCATACCAAGG |
| | H51D_fwd | CCTTGGTATGGGACTTGGCT**GAT**ATTGGACAGC |
| | H51E_fwd | CCTTGGTATGGGACTTGGCT**GAA**ATTGGACAGC |
| | H51A_fwd | CCTTGGTATGGGACTTGGCT**GCG**ATTGGACAGC |
| | H134X_rev | CGAGACGACCATCCCGAAGG |
| | H134D_fwd | CCTTCGGGATGGTCGTCTCG**GAT**GAACACCAGC |
| | H134E_fwd | CCTTCGGGATGGTCGTCTCG**GAA**GAACACCAGC |
| | H134A_fwd | CCTTCGGGATGGTCGTCTCG**GCG**GAACACCAGC |
| | H138X_rev | CTGGTGTTCGTGCGAGACGAC |
| | H138A_fwd | GTCGTCTCGCACGAACACCAG**GCG**GATGAGATTATG |
| | H138D_fwd | GTCGTCTCGCACGAACACCAG**GAT**GATGAGATTATG |
| | H138E_fwd | GTCGTCTCGCACGAACACCAG**GAA**GATGAGATTATG |
| *Mth*EgtB Y377W W415R R87D T141M R379A A145K H417L (*Mth*EgtB$_{CHF2}$) | H51X_rev | AGCCAAGTCCCATACCAAGG |
| | H51D_fwd | CCTTGGTATGGGACTTGGCT**GAT**ATTGGACAGC |
| | H51E_fwd | CCTTGGTATGGGACTTGGCT**GAA**ATTGGACAGC |
| | H51A_fwd | CCTTGGTATGGGACTTGGCT**GCG**ATTGGACAGC |
| | H134X_rev | CGAGACGACCATCCCGAAGG |
| | H134D_fwd | CCTTCGGGATGGTCGTCTCG**GAT**GAACACCAGC |
| | H134E_fwd | CCTTCGGGATGGTCGTCTCG**GAA**GAACACCAGC |
| | H134E_fwd | CCTTCGGGATGGTCGTCTCG**GCG**GAACACCAGC |
| | H138X_rev | CTGGTGTTCGTGCGAGACGAC |
| | H138A_fwd | GTCGTCTCGCACGAACACCAG**GCG**GATGAGATGATG |
| | H138D_fwd | GTCGTCTCGCACGAACACCAG**GAT**GATGAGATGATG |
| | H138E_fwd | GTCGTCTCGCACGAACACCAG**GAA**GATGAGATGATG |
| ACCO I184A K158I | D179X_rev | GGTATGTGCACGCAGACCATAAATCAGATCC |
| | D179E_fwd | ATGGTCTGCGTGCACATACC**GAA**GCAGGTGGTATTG |

| | | |
|---|---|---|
| K172V K93Q T89A (ACCO$_{CHF}$) | D179A_fwd | ATGGTCTGCGTGCACATACC**GCG**GCAGGTGGTATTG |
| | D179H_fwd | ATGGTCTGCGTGCACATACC**CAT**GCAGGTGGTATTG |

#### IV. Determination of nonheme enzyme concentrations

##### 1. Determination of enzyme concentrations by SDS-PAGE analysis.

A representative example using SDS-PAGE analysis to determine the concentration of nonheme enzymes in the cell-free lysate is provided below. Typical concentrations of the entire $EgtB_{CHF}$ lineage are provided in Table **S3**.

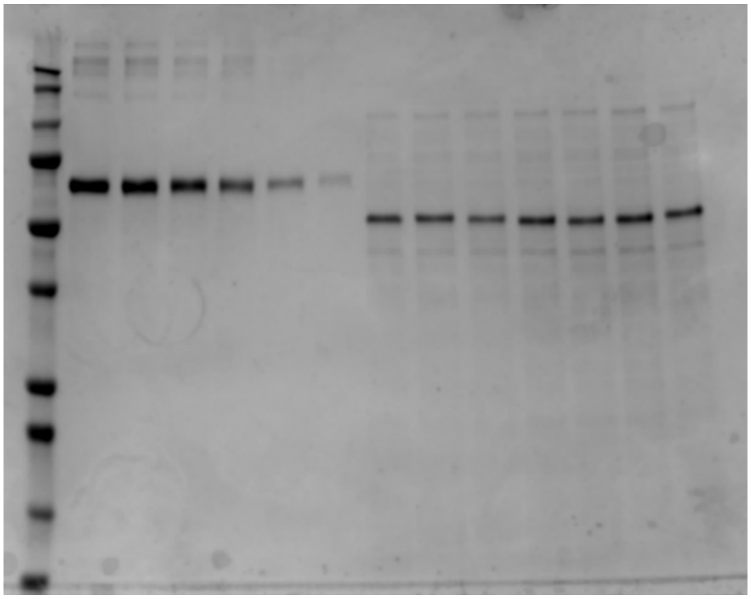

**Figure S1.** SDS-PAGE analysis for the determination of *Mth*EgtB concentration in cell-free lysates (Lane 1: protein ladder, Lanes 2-7: BSA standards, Lane 8-14: $EgtB_{CHF}$ variants).

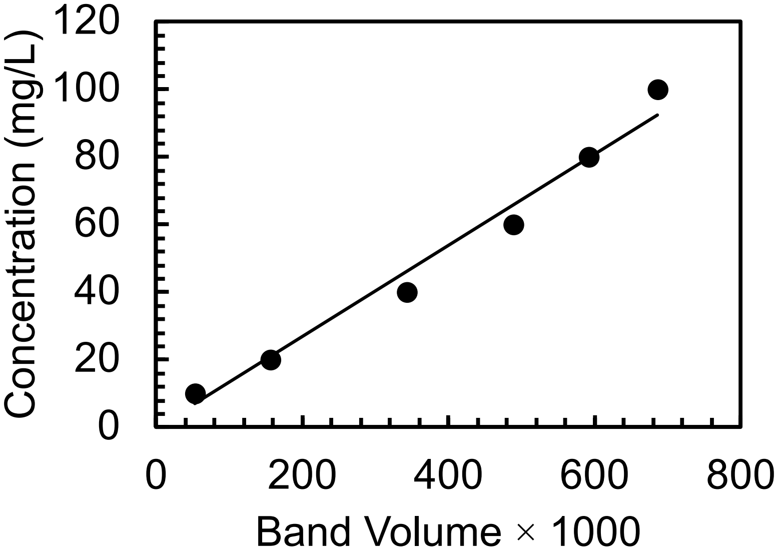

**Figure S2**. Calibration curve for nonheme enzyme concentration determination.

**Table S3.** Typical enzyme concentrations of EgtB$_{CHF}$ variants in cell-free lysate (CFL) determined using this method.

| entry | EgtB$_{CHF}$ variant | Enzyme concentration (□M) in CFL |
|---|---|---|
| 1 | wt *Mth*EgtB | 18.5 |
| 2 | *Mth*EgtB Y377W W415R | 13.9 |
| 3 | *Mth*EgtB Y377W W415R R87D | 21.4 |
| 4 | *Mth*EgtB Y377W W415R R87D T141I | 17.8 |
| 5 | *Mth*EgtB Y377W W415R R87D T141I F83Y | 22.3 |
| 6 | *Mth*EgtB Y377W W415R R87D T141I F83Y R379E | 20.3 |
| 7 | *Mth*EgtB Y377W W415R R87D T141I F83Y R379E V84R | 15.4 |
| 8 | *Mth*EgtB Y377W W415R R87D T141I F83Y R379E V84R H417L (**EgtB$_{CHF1}$**) | 10.7 |
| 9 | *Mth*EgtB Y377W W415R R87D T141I F83Y R379E V84R H417L H138A (**EgtB$_{CHF1}$ H138A**) | 15.6 |
| 10 | *Mth*EgtB Y377W W415R R87D T141M | 19.7 |
| 11 | *Mth*EgtB Y377W W415R R87D T141M R379A | 15.0 |
| 12 | *Mth*EgtB Y377W W415R R87D T141M R379A A145K | 21.2 |
| 13 | *Mth*EgtB Y377W W415R R87D T141M R379A A145K Q422D (**EgtB$_{CHF2}$**) | 21.6 |
| 14 | *Mth*EgtB Y377W W415R R87D T141M R379A A145K Q422D H138A (**EgtB$_{CHF2}$ H138A**) | 16.3 |

**2. Determination of iron concentrations in purified nonheme Fe enzymes.**

Iron in purified nonheme Fe enzyme was quantified using a modified ferrozine-based colorimetric method.[4] 500 μL purified nonheme Fe enzyme (100 μM) in KPi buffer (100 mM, pH = 7.4) was mixed with 250 μL of freshly prepared Reagent A (iron-releasing reagent; 1:1 mixture of 1.2 M HCl and 4.5% w/v $KMnO_4$) and incubated at 60 °C for 2 h in a sealed microcentrifuge tube to release enzyme bound iron. After cooling to room temperature, 50 μL of Reagent B (reducing/chelating reagent; 6.5 mM ferrozine, 13.1 mM neocuproine, 2.0 M ascorbic acid, and 5.0 M ammonium acetate in water) was added. The mixture was incubated at room temperature for 30 min to allow complete formation of the Fe(II)-ferrozine complex. Absorbance was measured at 562 nm using a UV–vis spectrophotometer. Iron concentrations were determined using a calibration curve developed using ferrous ethylenediammonium sulfate with 4 μg/mL, 2.4 μg/mL, 1.6 μg/mL, 0.8 μg/mL, 0.4 μg/mL, 0.2 μg/mL, and 0.1μg/mL.

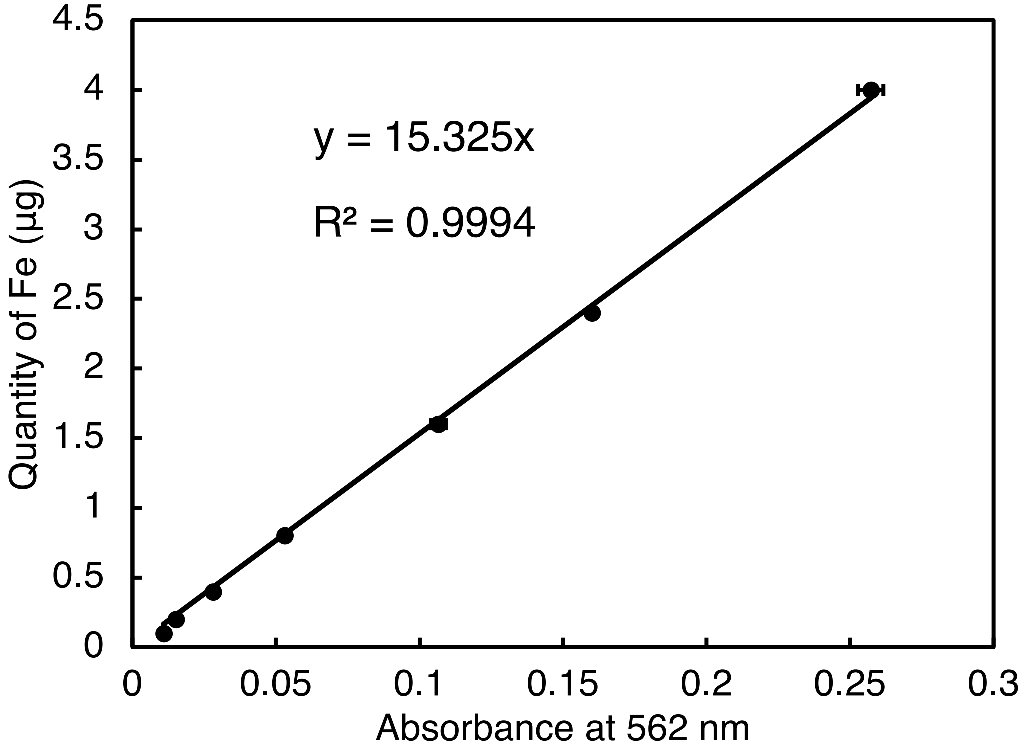

**Figure S3.** Ferrozine assay calibration curve for the determination of Fe concentration in purified nonheme Fe enzyme.

**Table S4**. Summary of Fe quantity in selected purified EgtB and ACCO variants determined by the ferrozine assay.

| enzyme variant | absorbance at 562 nm[a] | Fe concentration (μg/mL) | Fe occupancy |
|---|---|---|---|
| $EgtB_{CHF1}$ | 0.153 ± 0.006 | 2.34 ± 0.09 | 42.5% |
| $EgtB_{CHF1}$ H51D | 0.038 ± 0.001 | 0.58 ± 0.01 | 10.5% |
| $EgtB_{CHF1}$ H134E | 0.019 ± 0.003 | 0.30 ± 0.05 | 5.4% |
| $EgtB_{CHF1}$ H138A | 0.053 ± 0.002 | 0.81 ± 0.04 | 14.8% |
| $EgtB_{CHF1}$ H138D | 0.097 ± 0.006 | 1.50 ± 0.09 | 27.2% |
| $EgtB_{CHF1}$ H138E | 0.091 ± 0.005 | 1.41 ± 0.08 | 25.6% |
| $EgtB_{CHF2}$ | 0.180 ± 0.003 | 2.77 ± 0.05 | 50.4% |
| $EgtB_{CHF2}$ H138A | 0.078 ± 0.007 | 1.20 ± 0.10 | 21.8% |
| $ACCO_{CHF}$ | 0.125 ± 0.004 | 1.92 ± 0.06 | 34.9% |
| $ACCO_{CHF}$ D179A | 0.086 ± 0.004 | 1.32 ± 0.05 | 24.1% |
| $ACCO_{CHF}$ D179E | 0.103 ± 0.007 | 1.58 ± 0.10 | 28.8% |
| $ACCO_{CHF}$ D179H | 0.045 ± 0.005 | 0.69 ± 0.08 | 12.5% |

[a]All the experiments were performed in technical triplicates, and the average absorbance values and standard deviations are reported.

Iron incorporation analysis revealed that primary sphere mutations altered Fe incorporation to varying degrees. Although excess Mohr's salt was supplemented during the biotransformations, the reduced iron occupancy observed for several mutants suggests that perturbation of the primary coordination sphere can compromise productive metal binding, thereby lowering the overall catalytic activity. Nevertheless, the changes in chemoselectivity and stereoselectivity observed for active mutants indicate that these mutations also modulate the intrinsic rebound selectivity, rather than merely affecting metal loading.

#### V. Mechanism studies.

##### 1. Product analysis

Analysis of the reaction products obtained under the standard conditions revealed that no alkene side product, which would be expected from carbocation elimination pathways, were observed. In addition, no intramolecularly trapped lactam products arising from carbocation intermediates were detected. These observations argue against the involvement of a long-lived benzylic carbocation intermediate under the reaction conditions, consistent with previous experimental findings reported for Fe-catalyzed, fluoroamide-directed C–H fluorination reactions.[5]

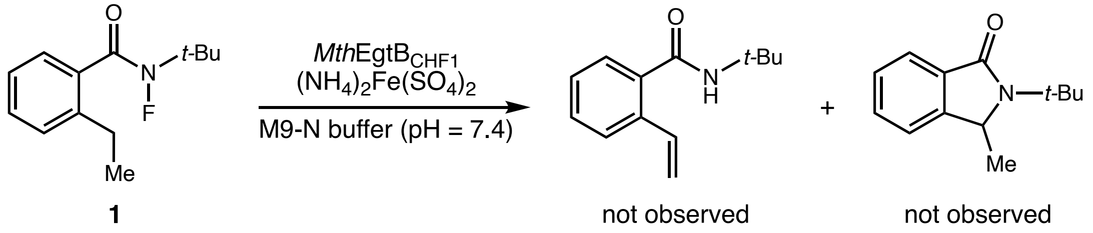

##### 2. Radical clock experiments

Following the standard conditions, the cyclopropyl substrate **6** was subjected to reactions with cell-free lysate containing EgtB$_{CHF1}$ or EgtB$_{CHF2}$. GC-MS analysis revealed that neither the directed fluorinated product **S1** nor the ring-opened fluorinated product **S2** was detected. Instead, the ring-opened byproduct, (*E*)-2-(but-1-en-1-yl)-N-(tert-butyl)benzamide (**8**), was observed.

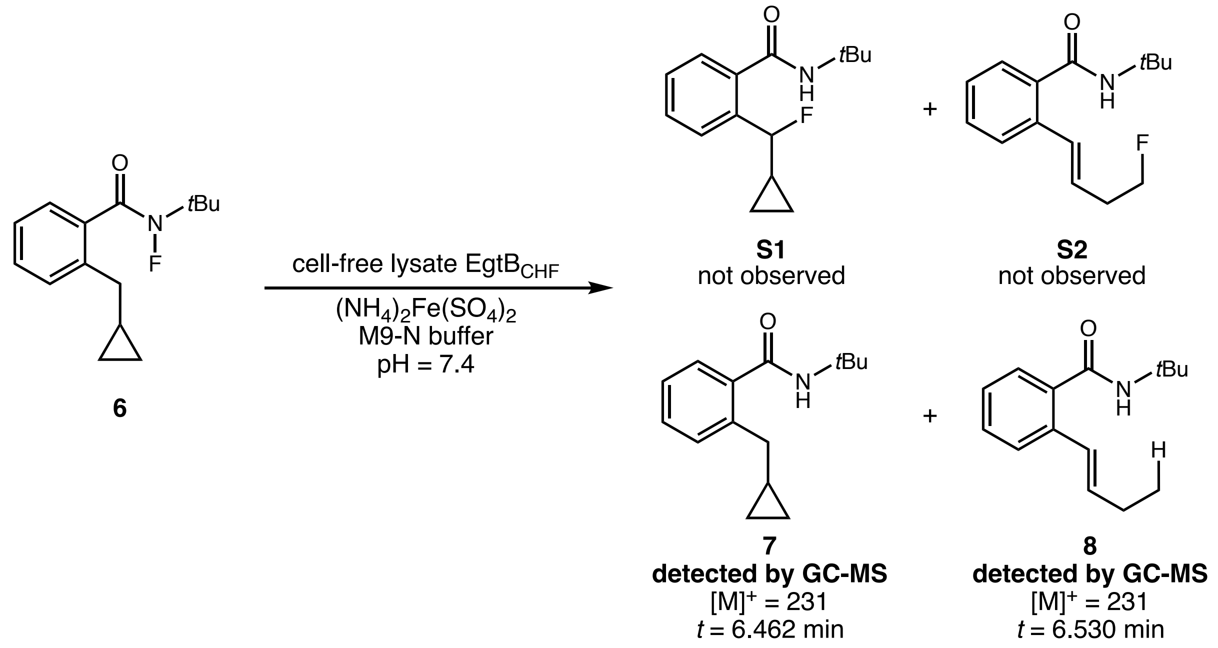

These results support the formation of intermediate **C** from intermediate **B** through a radical-mediated ring-opening process.

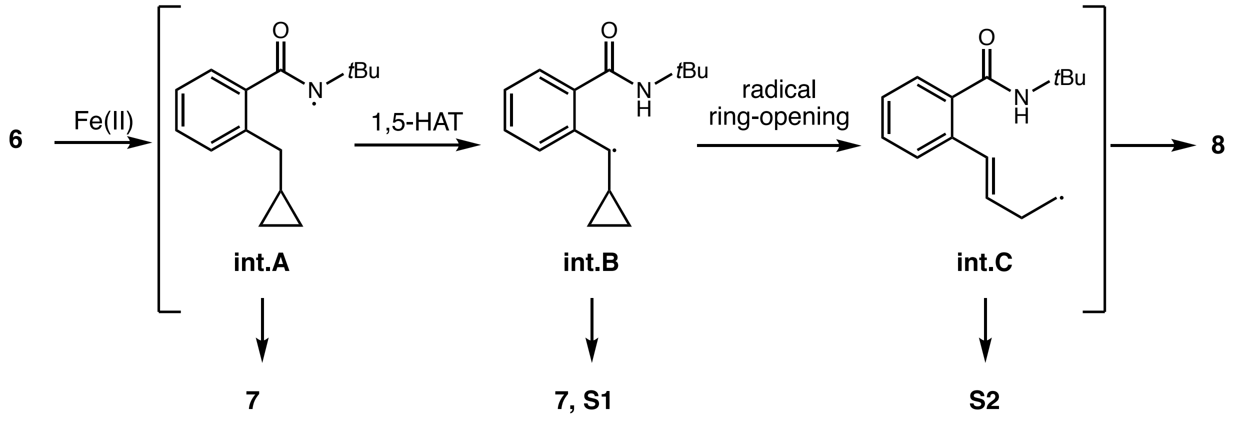

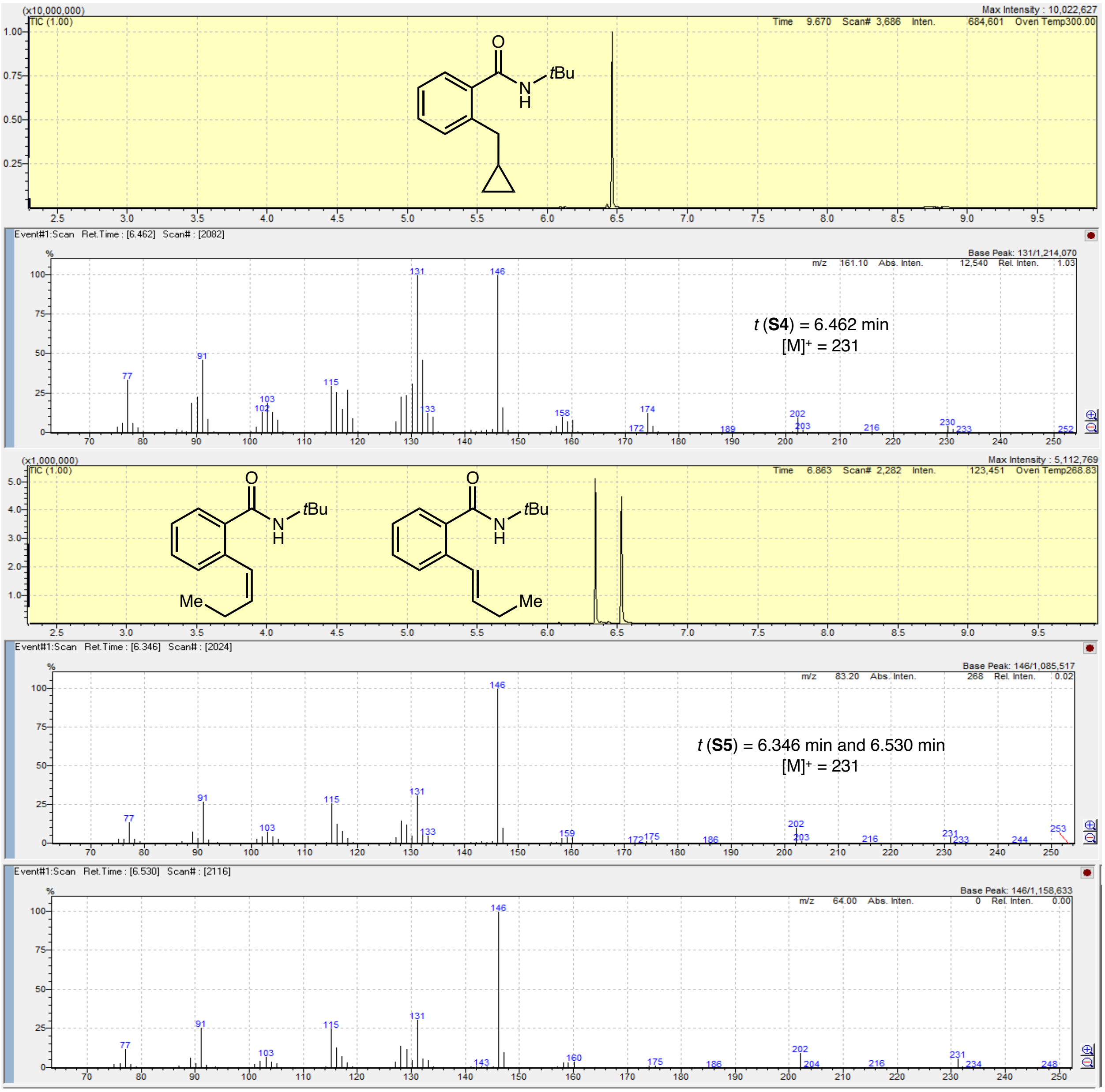

**Figure S4**. GC-MS analysis of authentic product **7** and (*Z*/*E*)-**8**.

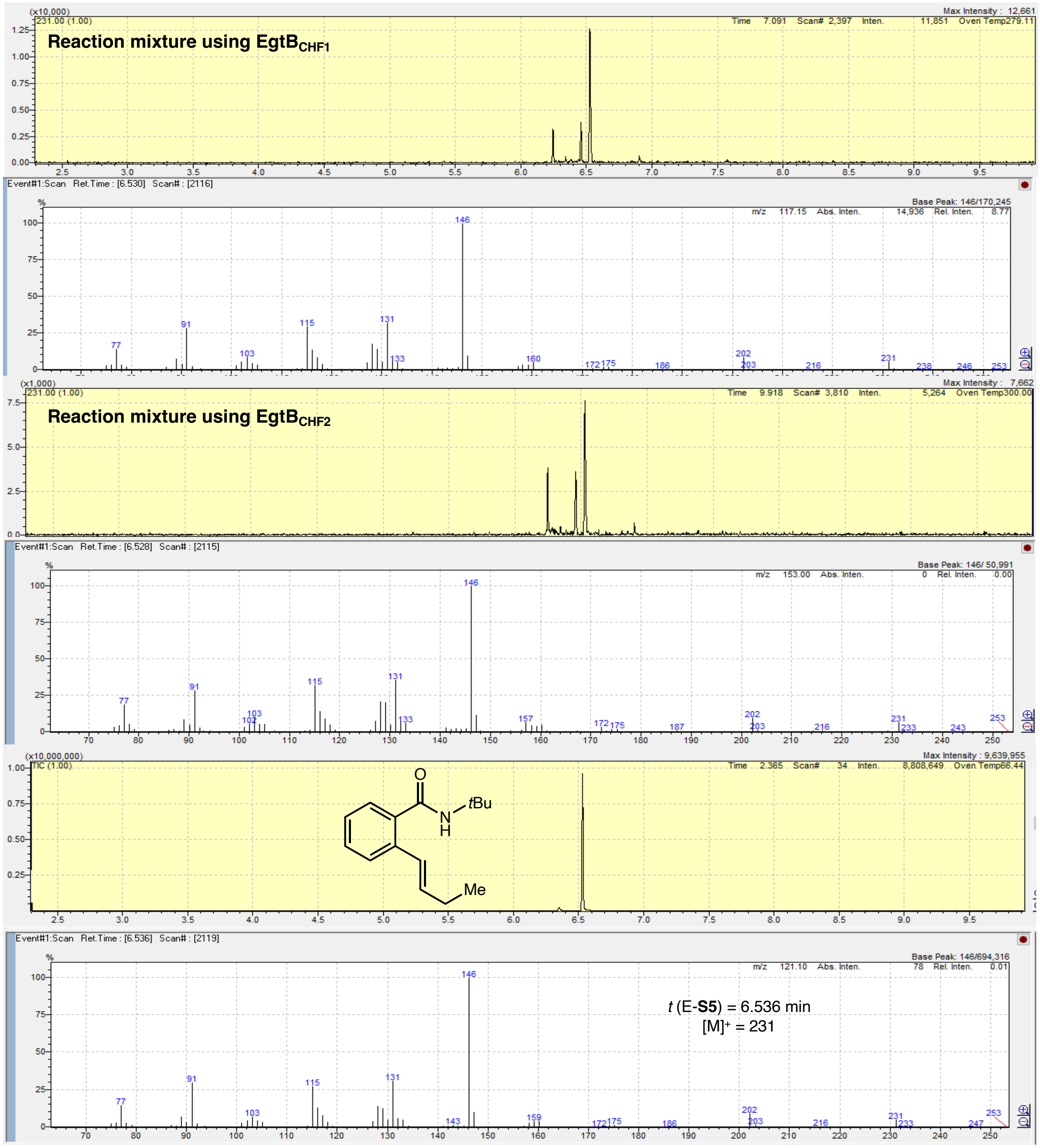

**Figure S5**. GC-MS analysis of reaction mixture.

#### 3. TEMPO trapping experiments

Following the standard conditions, TEMPO trapping experiments was performed with EgtB$_{CHF1}$, EgtB$_{CHF2}$ and ACCO$_{CHF}$. Upon addition of 2.0 equiv of TEMPO to the reaction mixture, no desired fluorinated product was observed in any of the three enzymatic systems. Instead, the radical trapping adduct **9** was detected by LC-MS analysis. These results support the radical-mediated nature of this biocatalytic transformation.

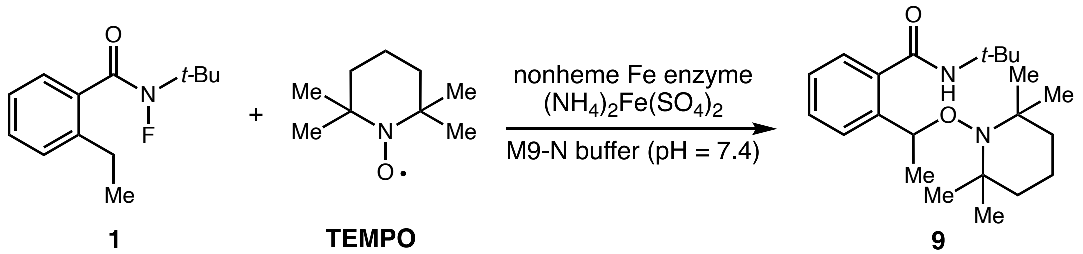

**1** **TEMPO** **9**

**detected by LC-MS**

$[M+H]^+ = 361$

Together, the cyclopropyl radical clock and TEMPO trapping experiments provide strong evidence for the involvement of radical intermediates in this biocatalytic transformation.

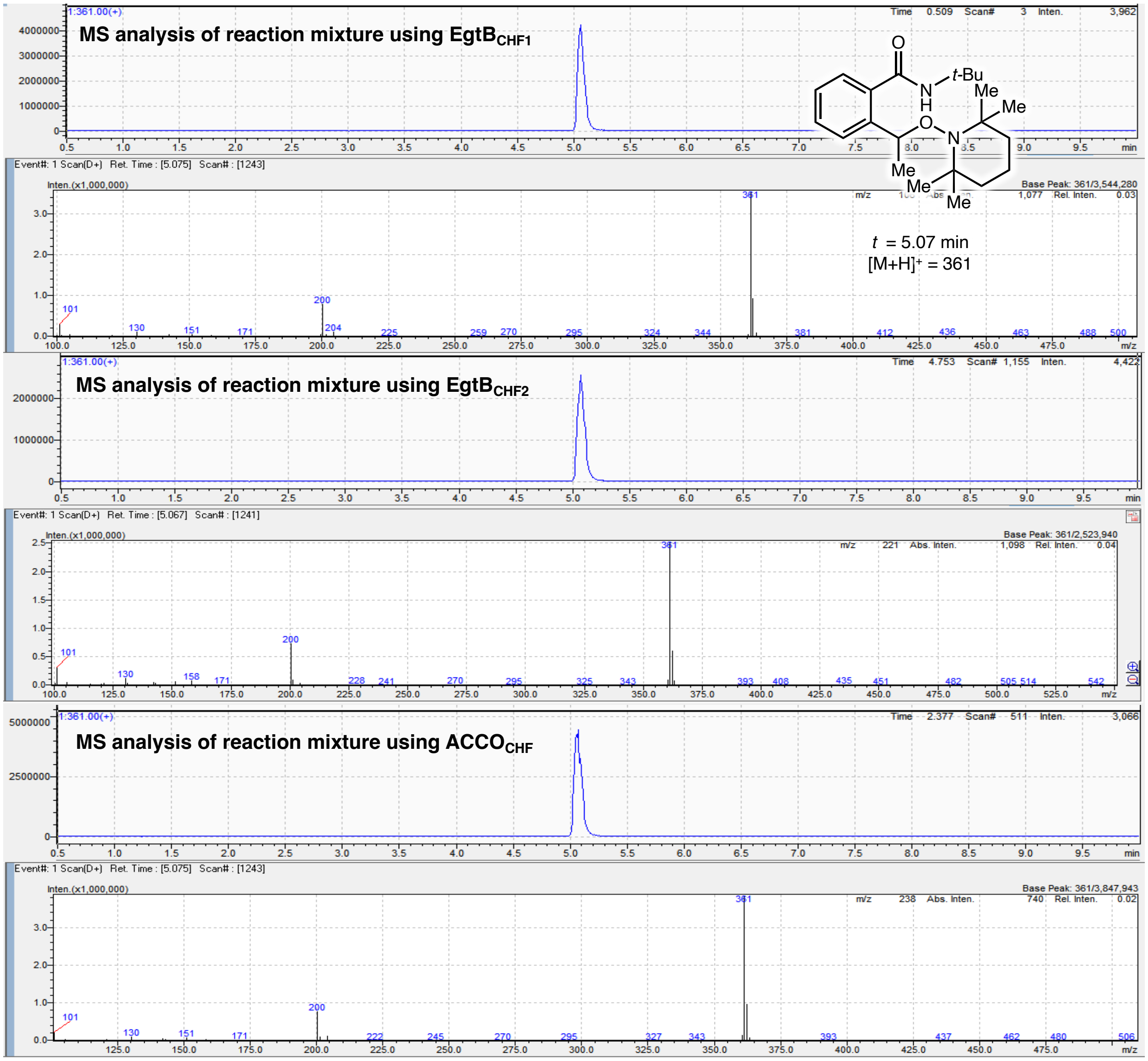

**Figure S6**. LC-MS analysis of reaction mixture.

#### 4. $^{18}$O-labeling experiments

$^{18}$O-labeling experiments was conducted using either $^{18}O_2$ or $H_2{}^{18}O$. When the reaction was performed under an atmosphere of $^{18}O_2$, no $^{18}$O-incorporated hydroxylated product was detected, whereas clear $^{18}$O incorporation was observed when the reaction was carried out in $H_2{}^{18}O$ containing buffer.

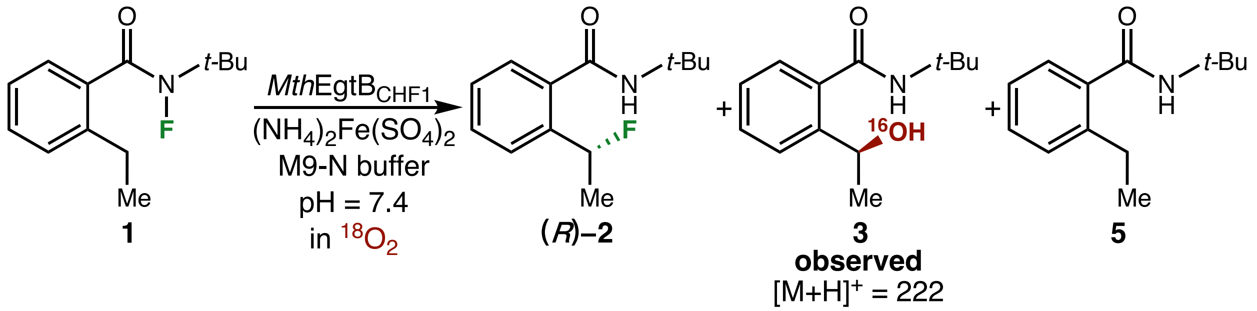

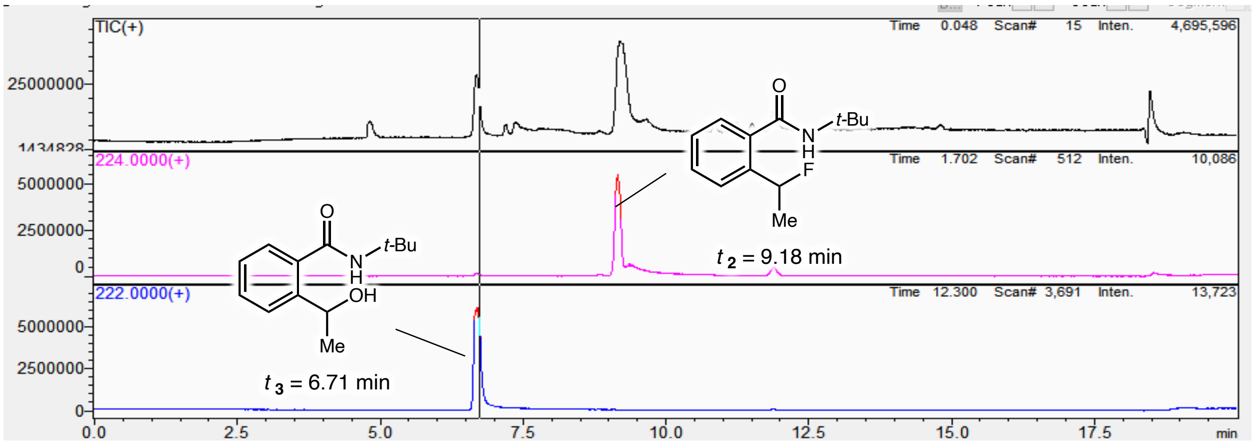

| retention time (min) | extracted ion peak area ($\mathbf{3}_{m/z=222}$) | extracted ion peak area ($\mathbf{3}_{m/z=224}$) | area ratio ($\mathbf{3}_{m/z=222}/\mathbf{3}_{m/z=224}$) |
|---|---|---|---|
| 6.71 | $5.4*10^7$ | $7.2*10^5$ | 99:1 |

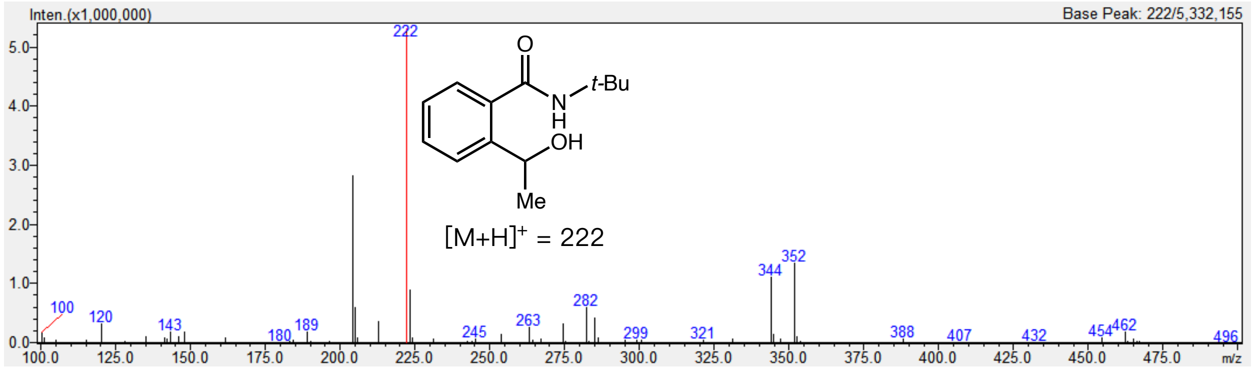

**Figure S7**. LC-MS analysis of reaction mixture.

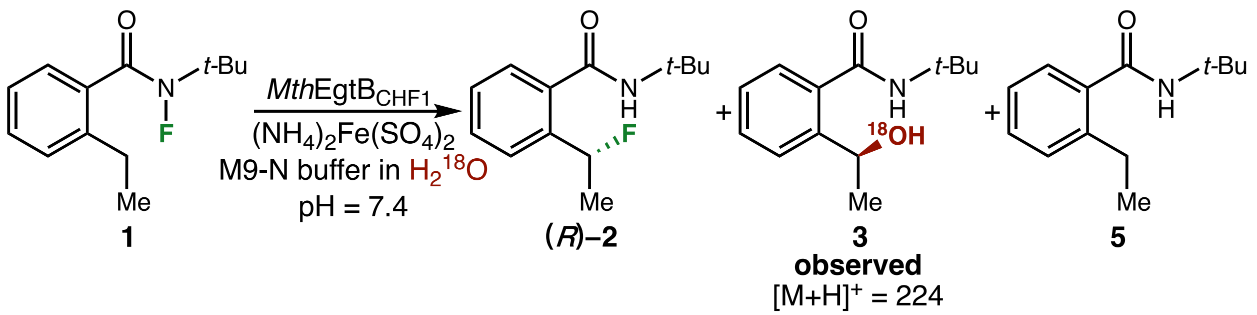

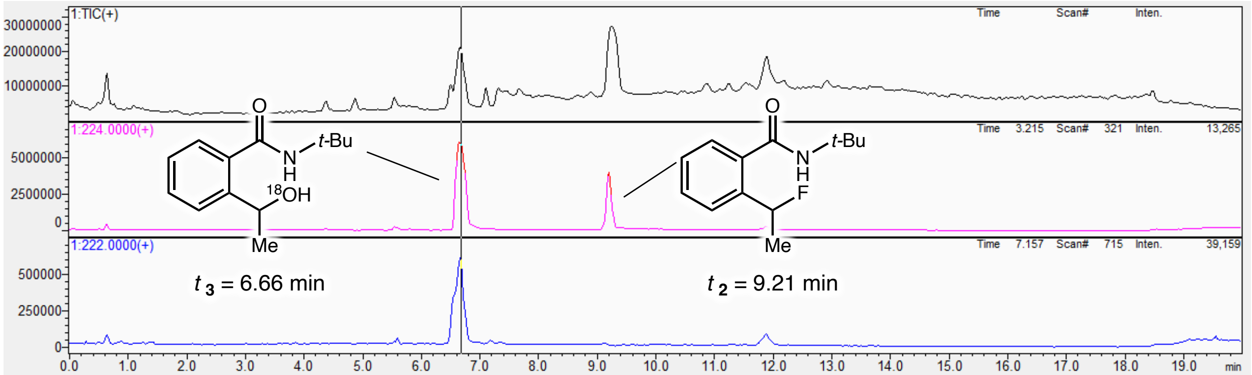

| retention time (min) | extracted ion peak area ($\mathbf{3}_{m/z=222}$) | extracted ion peak area ($\mathbf{3}_{m/z=224}$) | area ratio ($\mathbf{3}_{m/z=224}$/$\mathbf{3}_{m/z=222}$) |
|---|---|---|---|
| 6.66 | $6.4*10^6$ | $6.8*10^7$ | 91:9 |

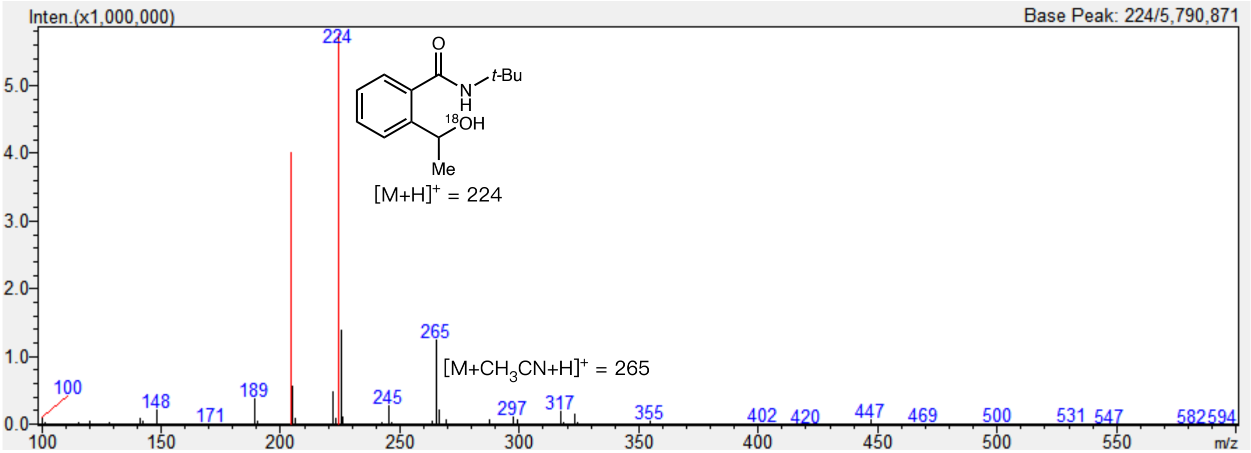

**Figure S8**. LC-MS analysis of reaction mixture.

#### VI. Primary coordination sphere effects on radical rebound activity and selectivity

To study the primary coordination sphere effect of the nonheme Fe enzyme on radical rebound activity and selectivity, we performed site-directed mutagenesis on evolved *Mth*EgtB to mutate the Fe-binding histidine (H) residues to aspartate (D), glutamate (E), and alanine (A) in the Fe center. A similar mutagenesis study was also carried out with our previously evolved C-H fluorinase $ACCO_{CHF}$ to mutate the Fe-binding aspartate (D) to E, H and A. A detailed discussion is provided in the main text.

**Table S5.** Primary coordination sphere effects on C–H fluorination with $EgtB_{CHF1}$

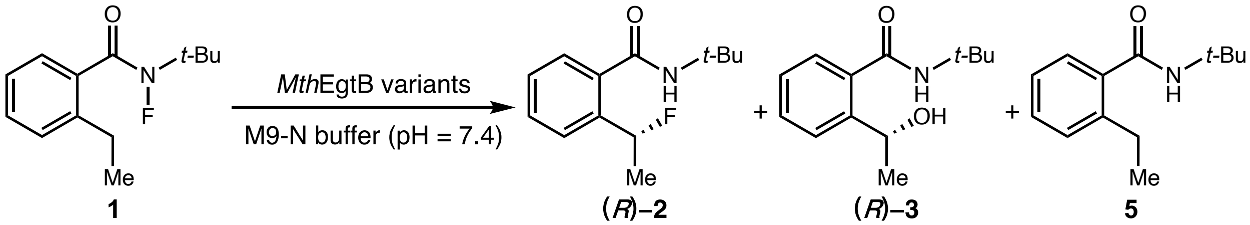

| entry | variants | 2 | | 3 | | 5 |
|---|---|---|---|---|---|---|
| | | yield (%) | e.r. ((*R*)-**2** : (*S*)-**2**) | yield (%) | e.r. ((*R*)-**3** : (*S*)-**3**) | yield (%) |
| 1 | $EgtB_{CHF1}$ | 30.7 ± 0.7 | 60:40 | 32 ± 1 | 23:77 | 14.7 ± 0.3 |
| 2 | $EgtB_{CHF1}$ H51D | 0 | - | 0 | - | 5 |
| 3 | $EgtB_{CHF1}$ H51E | 0 | - | 0 | - | 5 |
| 4 | $EgtB_{CHF1}$ H51A | 0 | - | 0 | - | 4 |
| 5 | $EgtB_{CHF1}$ H134D | 0 | - | 0 | - | 4 |
| 6 | $EgtB_{CHF1}$ H134E | 0 | - | 0 | - | 4 |
| 7 | $EgtB_{CHF1}$ H134A | 0 | - | 0 | - | 5 |
| 8 | $EgtB_{CHF1}$ H138D | 5.0 ± 0.1 | 36:64 | 0 | - | 6.3 ± 0.2 |
| 9 | $EgtB_{CHF1}$ H138E | 6.6 ± 0.1 | 35:65 | 0 | - | 4.4 ± 0.1 |

| | | | | | | |
|---|---|---|---|---|---|---|
| 10 | EgtB$_{CHF1}$ H138A | 7.3 ± 0.4 | 59:41 | 13.5 ± 0.1 | 24:76 | 7.0 ± 0.2 |

Reaction conditions: 7.5 mM **1**, 0.75 mM $(NH_4)_2Fe(SO_4)_2$, 7.5 mM sodium ascorbate, 500 μL cell-free lysate of EgtB variant, M9-N buffer (pH = 7.4). **2**: fluorination product, **3**: hydroxylation product, **5**: reduction product.

**Table S6.** Primary coordination sphere effects on C–H fluorination with EgtB$_{CHF2}$

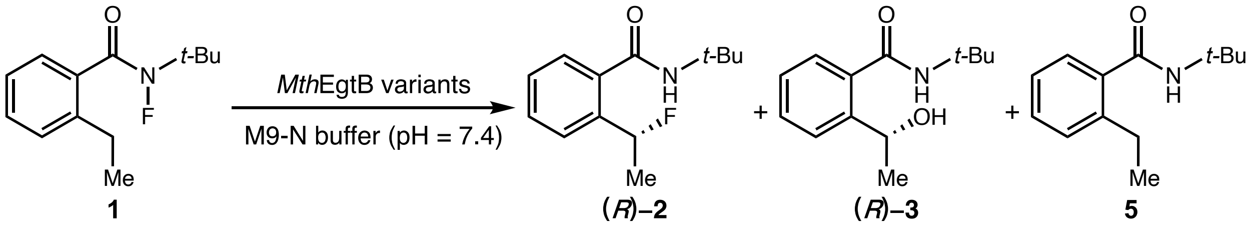

| entry | variants | 2 | | 3 | | 5 |
|---|---|---|---|---|---|---|
| | | yield (%) | e.r. ((*R*)-**2** : (*S*)-**2**) | yield (%) | e.r. ((*R*)-**3** : (*S*)-**3**) | yield (%) |
| 1 | EgtB$_{CHF2}$ | 35.5 ± 0.1 | 31:69 | 17.3 ± 0.5 | 34:66 | 16.9 ± 0.6 |
| 2 | EgtB$_{CHF2}$ H51D | 0 | - | 0 | - | 5 |
| 3 | EgtB$_{CHF2}$ H51E | 0 | - | 0 | - | 5 |
| 4 | EgtB$_{CHF2}$ H51A | 0 | - | 0 | - | 5 |
| 5 | EgtB$_{CHF2}$ H134D | 0 | - | 0 | - | 5 |
| 6 | EgtB$_{CHF2}$ H134E | 0 | - | 0 | - | 5 |
| 7 | EgtB$_{CHF2}$ H134A | 0 | - | 0 | - | 5 |
| 8 | EgtB$_{CHF2}$ H138D | 0 | - | 0 | - | 2.6 ± 0.2 |
| 9 | EgtB$_{CHF2}$ H138E | 1.3 ± 0.1 | 35:65 | 0 | - | 4.4 ± 0.1 |
| 10 | EgtB$_{CHF2}$ H138A | 0 | - | 0 | - | 2.1 ± 0.2 |

Reaction conditions: 7.5 mM **1**, 0.75 mM $(NH_4)_2Fe(SO_4)_2$, 7.5 mM sodium ascorbate, 500 μL cell-free lysate of EgtB variant, M9-N buffer (pH = 7.4). **2**: fluorination product, **3**: hydroxylation product, **5**: reduction product.

**Table S7.** Primary coordination sphere effects on C–H fluorination with $ACCO_{CHF}$ under whole-cell reaction conditions

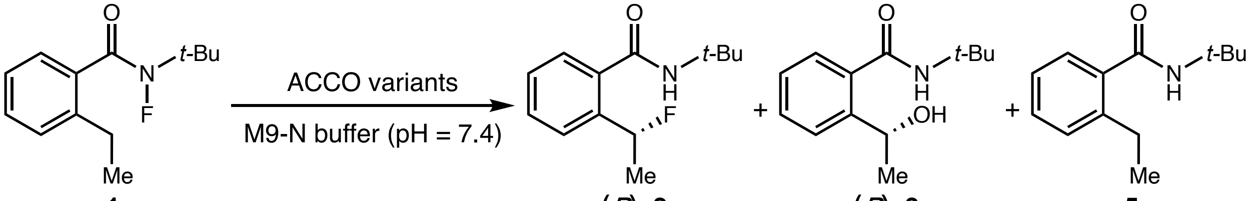

| entry | variants | 2 | | 3 | | 5 |
|---|---|---|---|---|---|---|
| | | yield (%) | e.r. ((*R*)-**2** : (*S*)-**2**) | yield (%) | e.r. ((*R*)-**3** : (*S*)-**3**) | yield (%) |
| 1 | $ACCO_{CHF}$ | 89 ± 3 | 5:95 | 0 | - | 1.5 ± 0.3 |
| 2 | $ACCO_{CHF}$ D179E | 0 | - | 0 | - | 9.7 ± 0.2 |
| 3 | $ACCO_{CHF}$ D179H | 0 | - | 0 | - | 2.6 ± 0.2 |
| 4 | $ACCO_{CHF}$ D179A | 0 | - | 0 | - | 4.9 ± 0.4 |

Reaction conditions: 7.5 mM **1**, 0.75 mM $(NH_4)_2Fe(SO_4)_2$, 500 μL whole-cell suspension of ACCO variant, M9-N buffer (pH = 7.4). **2**: fluorination product, **3**: hydroxylation product, **5**: reduction product.

**Table S8.** Primary coordination sphere effects on C–H fluorination with $ACCO_{CHF}$ under cell-free lysate reaction conditions

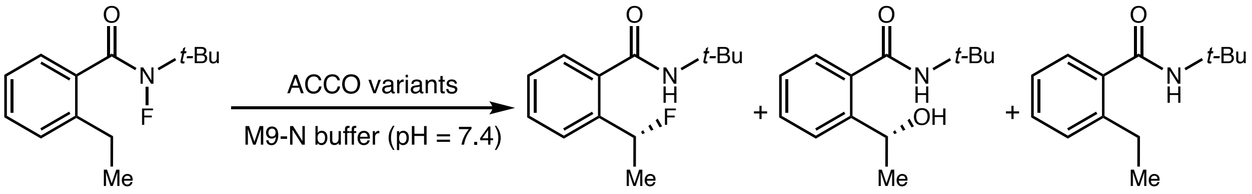

| entry | variants | **2** | | **3** | | **5** |
|---|---|---|---|---|---|---|
| | | yield (%) | e.r. ((*R*)-**2** : (*S*)-**2**) | yield (%) | e.r. ((*R*)-**3** : (*S*)-**3**) | yield (%) |
| 1 | $ACCO_{CHF}$ | 43 ± 2 | 11:89 | 0 | - | 3.0 ± 0.3 |
| 2 | $ACCO_{CHF}$ D179E | 0 | - | 0 | - | 2.7 ± 0.2 |
| 3 | $ACCO_{CHF}$ D179H | 0 | - | 0 | - | 1.2 ± 0.1 |
| 4 | $ACCO_{CHF}$ D179A | 0 | - | 0 | - | 2.9 ± 0.3 |

Reaction conditions: 7.5 mM **1**, 0.75 mM $(NH_4)_2Fe(SO_4)_2$, 7.5 mM sodium ascorbate, 500 μL cell-free lysate of ACCO variant, M9-N buffer (pH = 7.4). **2**: fluorination product, **3**: hydroxylation product, **5**: reduction product.

**Primary coordination sphere effects on C–H azidation**

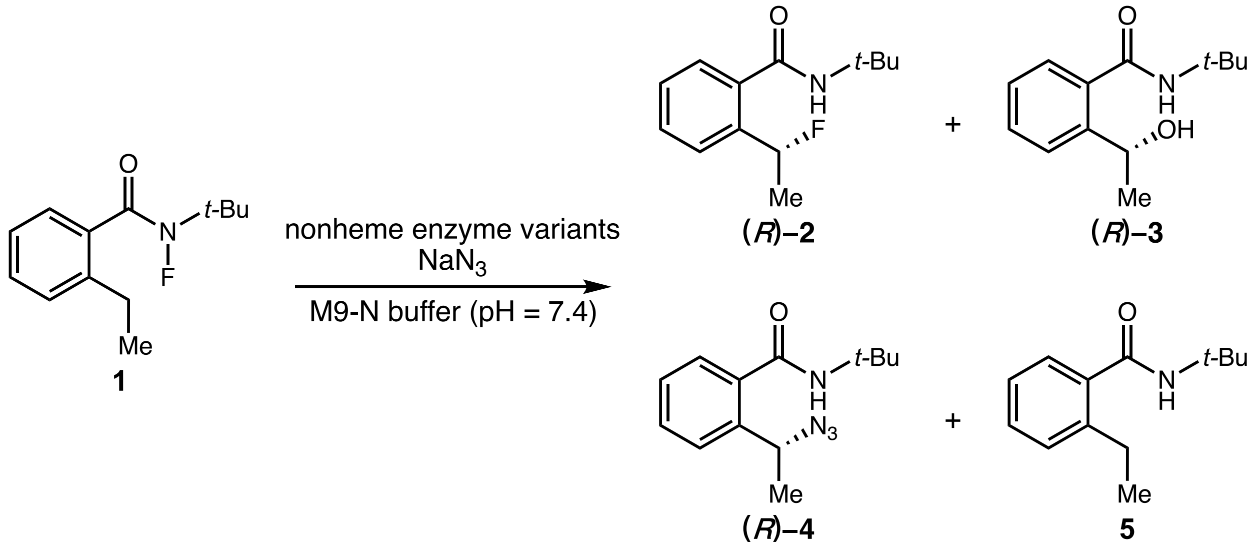

**Table S9.** Primary coordination sphere effects on C–H azidation with $EgtB_{CHF1}$

| entry | variants | **2** | | **3** | | **4** | | **5** |
|---|---|---|---|---|---|---|---|---|
| | | yield (%) | e.r. ((*R*)-**2** : (*S*)-**2**) | yield (%) | e.r. ((*R*)-**3** : (*S*)-**3**) | yield (%) | e.r. ((*R*)-**4** : (*S*)-**4**) | yield (%) |
| 1 | $EgtB_{CHF1}$ | 15 ± 2 | 60:40 | 23 ± 1 | 23:77 | 50 ± 2 | 44:56 | 15.4 ± 0.9 |
| 2 | $EgtB_{CHF1}$ H51D | 0 | - | 0 | - | 0 | - | 5 |
| 3 | $EgtB_{CHF1}$ H51E | 0 | - | 0 | - | 0 | - | 5 |
| 4 | $EgtB_{CHF1}$ H51A | 0 | - | 0 | - | 0 | - | 5 |
| 5 | $EgtB_{CHF1}$ H134D | 0 | - | 0 | - | 0 | - | 5 |
| 6 | $EgtB_{CHF1}$ H134E | 0 | - | 0 | - | 0 | - | 5 |
| 7 | $EgtB_{CHF1}$ H134A | 0 | - | 0 | - | 0 | - | 5 |
| 8 | $EgtB_{CHF1}$ H138D | 2.3 ± 0.2 | 35:65 | 0 | - | 0 | - | 3.3 ± 0.1 |
| 9 | $EgtB_{CHF1}$ H138E | 6.7 ± 0.2 | 35:65 | 0 | - | <1 | - | 5.1 ± 0.2 |
| 10 | $EgtB_{CHF1}$ H138A | 7.5 ± 0.3 | 50:50 | 9.0 ± 0.9 | 24:76 | 31.2 ± 0.8 | 45:55 | 9.1 ± 0.3 |

Reaction conditions: 7.5 mM **1**, 120 mM $NaN_3$, 0.75 mM $(NH_4)_2Fe(SO_4)_2$, 7.5 mM sodium ascorbate, 500 μL cell-free lysate of EgtB variant, M9-N buffer (pH = 7.4). **2**: fluorination product, **3**: hydroxylation product, **4**: azidation product, **5**: reduction product.

**Table S10.** Primary coordination sphere effects on C–H azidation with $EgtB_{CHF2}$

| entry | variants | **2** | | **3** | | **4** | | **5** |
|---|---|---|---|---|---|---|---|---|
| | | yield (%) | e.r. ((*R*)-**2** : (*S*)-**2**) | yield (%) | e.r. ((*R*)-**3** : (*S*)-**3**) | yield (%) | e.r. ((*R*)-**4** : (*S*)-**4**) | yield (%) |
| 1 | $EgtB_{CHF2}$ | 14.9 ± 0.5 | 31:69 | 9.7 ± 0.4 | 33:67 | 40 ± 2 | 39:61 | 12.1 ± 0.5 |
| 2 | $EgtB_{CHF2}$ H51D | 0 | - | 0 | - | <1 | - | 5 |
| 3 | $EgtB_{CHF2}$ H51E | 0 | - | 0 | - | <1 | - | 5 |
| 4 | $EgtB_{CHF2}$ H51A | 0 | - | 0 | - | <1 | - | 5 |
| 5 | $EgtB_{CHF2}$ H134D | 0 | - | 0 | - | <1 | - | 5 |
| 6 | $EgtB_{CHF2}$ H134E | 0 | - | 0 | - | <1 | - | 5 |
| 7 | $EgtB_{CHF2}$ H134A | 0 | - | 0 | - | <1 | - | 5 |
| 8 | $EgtB_{CHF2}$ H138D | 0 | - | 0 | - | <1 | - | 3 |
| 9 | $EgtB_{CHF2}$ H138E | 1.4 ± 0.1 | - | 0 | - | 1.0 ± 0.1 | - | 5.3 ± 0.2 |
| 10 | $EgtB_{CHF2}$ H138A | 1.0 ± 0.1 | - | 0 | - | 42 ± 2 | 27:73 | 7.6 ± 0.1 |

Reaction conditions: 7.5 mM **1**, 120 mM $NaN_3$, 0.75 mM $(NH_4)_2Fe(SO_4)_2$, 7.5 mM sodium ascorbate, 500 µL cell-free lysate of EgtB variant, M9-N buffer (pH = 7.4). **2**: fluorination product, **3**: hydroxylation product, **4**: azidation product, **5**: reduction product.

**Table S11.** Primary coordination sphere effects on C–H azidation with $ACCO_{CHF}$ under whole-cell reaction conditions

| entry | variants | **2** | | **3** | | **4** | | **5** |
|---|---|---|---|---|---|---|---|---|
| | | yield (%) | e.r. ((*R*)-**2** : (*S*)-**2**) | yield (%) | e.r. ((*R*)-**3** : (*S*)-**3**) | yield (%) | e.r. ((*R*)-**4** : (*S*)-**4**) | yield (%) |
| 1 | $ACCO_{CHF}$ | 77 ± 3 | 6:94 | 0 | - | 12 ± 1 | 49:51 | 5 ± 1 |
| 2 | $ACCO_{CHF}$ D179E | 0 | - | 0 | - | 14 ± 2 | 45:55 | 9.7 ± 0.2 |
| 3 | $ACCO_{CHF}$ D179H | 0 | - | 0 | - | 0 | - | 2.6 ± 0.2 |
| 4 | $ACCO_{CHF}$ D179A | 0 | - | 0 | - | 10 ± 1 | 42:58 | 4.9 ± 0.4 |

Reaction conditions: 7.5 mM **1**, 120 mM $NaN_3$, 0.75 mM $(NH_4)_2Fe(SO_4)_2$, 500 µL whole-cell suspension of ACCO variant, M9-N buffer (pH = 7.4). **2**: fluorination product, **3**: hydroxylation product, **4**: azidation product, **5**: reduction product.

**Table S12.** Primary coordination sphere effects on C–H azidation with $ACCO_{CHF}$ under cell-free lysate reaction conditions

| entry | variants | **2** | | **3** | | **4** | | **5** |
|---|---|---|---|---|---|---|---|---|
| | | yield (%) | e.r. ((*R*)-**2** : (*S*)-**2**) | yield (%) | e.r. ((*R*)-**3** : (*S*)-**3**) | yield (%) | e.r. ((*R*)-**4** : (*S*)-**4**) | yield (%) |
| 1 | $ACCO_{CHF}$ | 51 ± 2 | 18:82 | 0 | - | 22 ± 1 | 42:58 | 8 ± 1 |
| 2 | $ACCO_{CHF}$ D179E | 0 | - | 0 | - | 35 ± 2 | 45:55 | 15 ± 1 |
| 3 | $ACCO_{CHF}$ D179H | 0 | - | 0 | - | 0 | - | 2.1 ± 0.2 |
| 4 | $ACCO_{CHF}$ D179A | 0 | - | 0 | - | 35 ± 2 | 43:57 | 15 ± 1 |

Reaction conditions: 7.5 mM **1**, 120 mM $NaN_3$, 0.75 mM $(NH_4)_2Fe(SO_4)_2$, 7.5 mM sodium ascorbate, 500 µL cell-free lysate of ACCO variant, M9-N buffer (pH = 7.4). **2**: fluorination product, **3**: hydroxylation product, **4**: azidation product, **5**: reduction product.

#### VII. pH effects on radical rebound activity and selectivity

To further investigate the effect of pH on hydroxyl rebound activity and selectivity in nonheme Fe enzymes, we performed fluorination and azidation reactions using the evolved variants *Mth*EgtB$_{CHF1}$, *Mth*EgtB$_{CHF2}$ and ACCO$_{CHF}$ over a range of buffer pH values (6.5–9.0). The results revealed that higher pH conditions generally promoted both the activity and selectivity of hydroxylation. In particular, *Mth*EgtB$_{CHF1}$ exhibited the highest hydroxylation performance at pH 9.0, affording the hydroxylation product in 56% yield with a hydroxylation-to-fluorination ratio of 2:1.

**Table S13.** pH effects on C–H hydroxylation with EgtB$_{CHF1}$

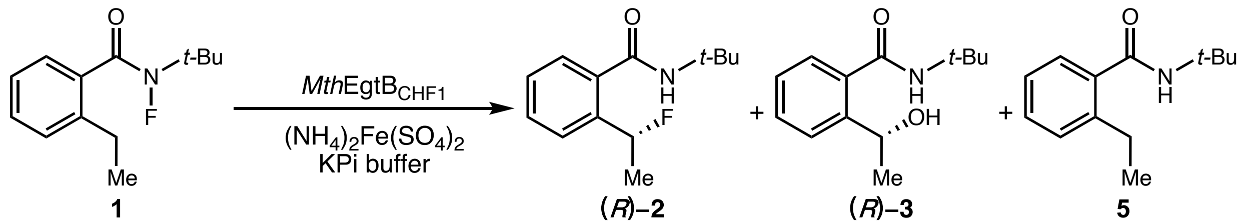

| entry | buffer pH | 2 | | 3 | | 5 |
|---|---|---|---|---|---|---|
| | | yield (%) | e.r. ((*R*)-**2** : (*S*)-**2**) | yield (%) | e.r. ((*R*)-**3** : (*S*)-**3**) | yield (%) |
| 1 | 6.5 | 31 ± 2 | 58:42 | 36 ± 2 | 25:75 | 13 ± 1 |
| 2 | 7.0 | 31 ± 1 | 59:41 | 44 ± 1 | 24:76 | 14 ± 1 |
| 3 | 7.5 | 29 ± 1 | 60:40 | 48 ± 1 | 23:77 | 13 ± 1 |
| 4 | 8.0 | 26 ± 1 | 62:38 | 49 ± 1 | 20:80 | 13 ± 1 |
| 5 | 8.5 | 26 ± 1 | 62:38 | 49 ± 1 | 20:80 | 14 ± 1 |
| 6 | 9.0 | 28 ± 1 | 62:38 | 54 ± 1 | 20:80 | 15 ± 1 |

Reaction conditions: 7.5 mM **1**, 0.75 mM $(NH_4)_2Fe(SO_4)_2$, 7.5 mM sodium ascorbate, 500 µL cell-free lysate of EgtB$_{CHF1}$ variant, 100 mM KPi buffer. **2**: fluorination product, **3**: hydroxylation product, **5**: reduction product.

**Table S14.** pH effects on C–H hydroxylation with $EgtB_{CHF2}$

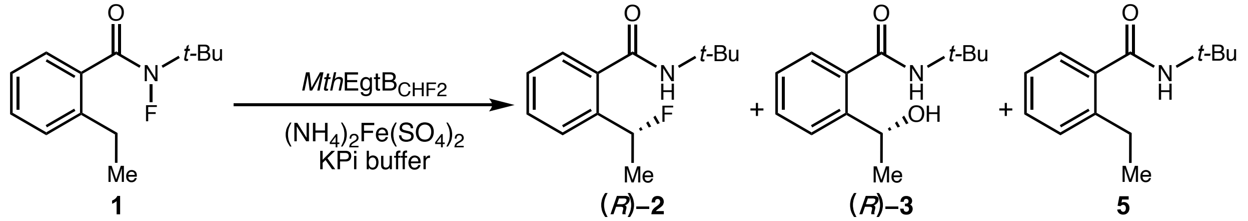

| entry | buffer pH | 2 | | 3 | | 5 |
|---|---|---|---|---|---|---|
| | | yield (%) | e.r. ((*R*)-**2** : (*S*)-**2**) | yield (%) | e.r. ((*R*)-**3** : (*S*)-**3**) | yield (%) |
| 1 | 6.5 | 36 ± 2 | 30:70 | 17 ± 1 | 32:68 | 15 ± 1 |
| 2 | 7.0 | 32 ± 1 | 32:68 | 17 ± 1 | 30:70 | 14 ± 1 |
| 3 | 7.5 | 40 ± 1 | 31:69 | 18 ± 1 | 34:66 | 17 ± 1 |
| 4 | 8.0 | 38 ± 4 | 31:69 | 22 ± 1 | 29:71 | 17 ± 2 |
| 5 | 8.5 | 46 ± 2 | 31:69 | 25 ± 2 | 32:68 | 21 ± 1 |
| 6 | 9.0 | 43 ± 1 | 31:69 | 28 ± 2 | 31:69 | 21 ± 1 |

Reaction conditions: 7.5 mM **1**, 0.75 mM $(NH_4)_2Fe(SO_4)_2$, 7.5 mM sodium ascorbate, 500 µL cell-free lysate of $EgtB_{CHF2}$ variant, 100 mM KPi buffer. **2**: fluorination product, **3**: hydroxylation product, **5**: reduction product.

**Table S15.** pH effects on C–H hydroxylation with $ACCO_{CHF}$

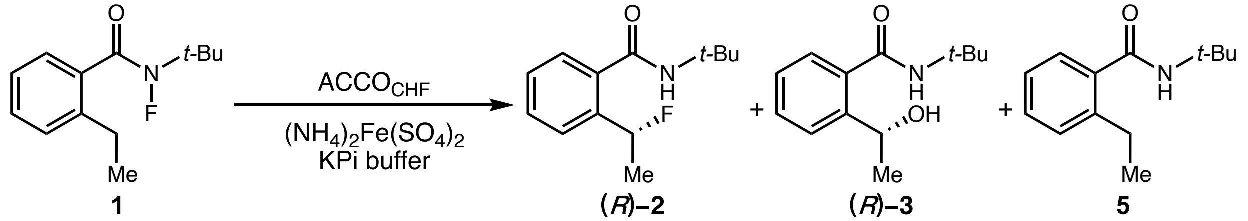

| entry | buffer pH | **2** | | **3** | | **5** |
|---|---|---|---|---|---|---|
| | | yield (%) | e.r. ((*R*)-**2** : (*S*)-**2**) | yield (%) | e.r. ((*R*)-**3** : (*S*)-**3**) | yield (%) |
| 1 | 6.5 | 73 ± 3 | 6:94 | 0 | - | 3.9 ± 0.6 |
| 2 | 7.0 | 71 ± 1 | 6:94 | 0 | - | 4.1 ± 0.2 |
| 3 | 7.5 | 83 ± 5 | 5:95 | 0 | - | 5.1 ± 0.6 |
| 4 | 8.0 | 77 ± 4 | 5:95 | 0 | - | 5.7 ± 0.3 |
| 5 | 8.5 | 79 ± 4 | 6:94 | 0 | - | 6.8 ± 0.5 |
| 6 | 9.0 | 80 ± 6 | 6:94 | 0 | - | 8.1 ± 0.6 |

Reaction conditions: 7.5 mM **1**, 0.75 mM $(NH_4)_2Fe(SO_4)_2$, 500 μL whole-cell suspension of $ACCO_{CHF}$ variant, 100 mM KPi buffer. **2**: fluorination product, **3**: hydroxylation product, **5**: reduction product.

**pH effects on C–H azidation**

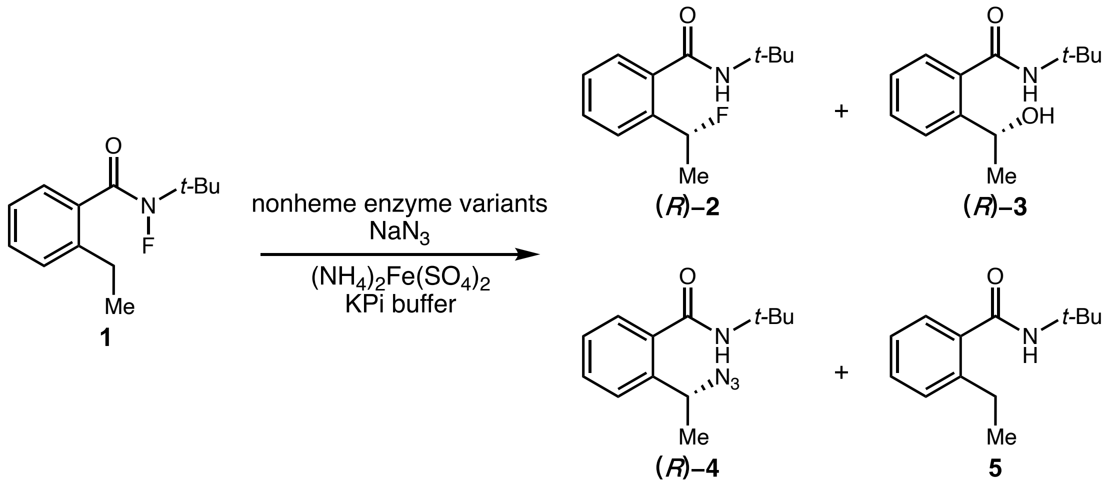

**Table S16.** pH effects on C–H azidation with $EgtB_{CHF1}$

| entry | buffer pH | **2** | | **3** | | **4** | | **5** |
|---|---|---|---|---|---|---|---|---|
| | | yield (%) | e.r. ((*R*)-**2** : (*S*)-**2**) | yield (%) | e.r. ((*R*)-**3** : (*S*)-**3**) | yield (%) | e.r. ((*R*)-**4** : (*S*)-**4**) | yield (%) |
| 1 | 6.5 | 13 ± 1 | 58:42 | 23 ± 2 | 24:76 | 52 ± 3 | 44:56 | 12 ± 1 |
| 2 | 7.0 | 14 ± 1 | 59:41 | 27 ± 1 | 23:77 | 55 ± 2 | 44:56 | 13 ± 1 |
| 3 | 7.5 | 14 ± 1 | 61:39 | 28 ± 1 | 21:79 | 53 ± 1 | 44:56 | 13 ± 1 |
| 4 | 8.0 | 14 ± 1 | 62:38 | 31 ± 1 | 19:81 | 47 ± 1 | 43:57 | 13 ± 1 |
| 5 | 8.5 | 14 ± 1 | 63:37 | 31 ± 1 | 18:82 | 49 ± 2 | 43:57 | 13 ± 1 |
| 6 | 9.0 | 14 ± 1 | 63:37 | 34 ± 1 | 19:81 | 50 ± 1 | 42:57 | 14 ± 1 |

Reaction conditions: 7.5 mM **1**, 120 mM $NaN_3$, 0.75 mM $(NH_4)_2Fe(SO_4)_2$, 7.5 mM sodium ascorbate, 500 μL cell-free lysate of $EgtB_{CHF1}$ variant, 100 mM KPi buffer. **2**: fluorination product, **3**: hydroxylation product, **4**: azidation product, **5**: reduction product.

**Table S17.** pH effects on C–H azidation with $EgtB_{CHF2}$

| entry | buffer pH | **2** | | **3** | | **4** | | **5** |
|---|---|---|---|---|---|---|---|---|
| | | yield (%) | e.r. ((*R*)-**2** : (*S*)-**2**) | yield (%) | e.r. ((*R*)-**3** : (*S*)-**3**) | yield (%) | e.r. ((*R*)-**4** : (*S*)-**4**) | yield (%) |
| 1 | 6.5 | 14 ± 1 | 25:75 | 10 ± 1 | 31:69 | 46 ± 3 | 38:62 | 18 ± 1 |
| 2 | 7.0 | 17 ± 1 | 28:72 | 15 ± 1 | 28:72 | 44 ± 1 | 38:62 | 13 ± 1 |
| 3 | 7.5 | 22 ± 4 | 25:75 | 11 ± 2 | 28:72 | 45 ± 3 | 38:62 | 17 ± 3 |
| 4 | 8.0 | 20 ± 1 | 29:71 | 16 ± 1 | 29:71 | 47 ± 1 | 37:63 | 15 ± 1 |
| 5 | 8.5 | 26 ± 3 | 26:74 | 14 ± 1 | 32:68 | 45 ± 1 | 38:62 | 18 ± 1 |
| 6 | 9.0 | 25 ± 2 | 28:72 | 17 ± 1 | 30:70 | 44 ± 1 | 38:62 | 19 ± 1 |

Reaction conditions: 7.5 mM **1**, 120 mM $NaN_3$, 0.75 mM $(NH_4)_2Fe(SO_4)_2$, 7.5 mM sodium ascorbate, 500 µL cell-free lysate of $EgtB_{CHF2}$ variant, 100 mM KPi buffer. **2**: fluorination product, **3**: hydroxylation product, **4**: azidation product, **5**: reduction product.

**Table S18.** pH effects on C–H azidation with $ACCO_{CHF}$

| entry | buffer pH | **2** yield (%) | **2** e.r. ((*R*)-**2** : (*S*)-**2**) | **3** yield (%) | **3** e.r. ((*R*)-**3** : (*S*)-**3**) | **4** yield (%) | **4** e.r. ((*R*)-**4** : (*S*)-**4**) | **5** yield (%) |
|---|---|---|---|---|---|---|---|---|
| 1 | 6.5 | 61 ± 4 | 10:90 | 0 | - | 15 ± 2 | 43:57 | 12 ± 2 |
| 2 | 7.0 | 63 ± 5 | 9:91 | 0 | - | 15 ± 2 | 42:58 | 13 ± 2 |
| 3 | 7.5 | 66 ± 1 | 7:93 | 0 | - | 15 ± 1 | 42:58 | 13 ± 3 |
| 4 | 8.0 | 74 ± 4 | 7:93 | 0 | - | 16 ± 2 | 42:58 | 14 ± 2 |
| 5 | 8.5 | 75 ± 3 | 9:91 | 0 | - | 16 ± 2 | 41:59 | 14 ± 2 |
| 6 | 9.0 | 79 ± 2 | 8:92 | 0 | - | 17 ± 1 | 41:59 | 15 ± 1 |

Reaction conditions: 7.5 mM **1**, 120 mM $NaN_3$, 0.75 mM $(NH_4)_2Fe(SO_4)_2$, 500 µL whole-cell suspension of $ACCO_{CHF}$ variant, 100 mM KPi buffer. **2**: fluorination product, **3**: hydroxylation product, **4**: azidation product, **5**: reduction product.

**VIII. Synthesis and characterization of substrates and products.**

**Synthetic procedures to prepare substrates and racemic products.**

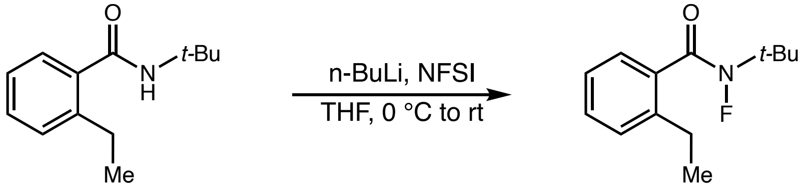

This procedure is modified from a previously described procedure.[5] A round bottom flask was dried in an oven (110 °C) and charged with a stir bar. This flask was evacuated and backfilled with $N_2$ and this procedure was repeated for a total of three times. To this flask were added the amide (17.8 g, 86.7 mmol, 1.0 equiv) and anhydrous THF (300 mL, 0.29 M). The mixture was cooled to 0 °C in an ice bath, and *n*-BuLi (38.2 mL, 2.5 M in hexanes, 1.1 equiv) was added dropwise. The reaction mixture was allowed to stir at 0 °C for 0.5 h before the addition of a solution of NFSI (41.0 g, 130 mmol, 1.5 equiv) in THF (180 mL). The reaction mixture was allowed to warm to room temperature slowly and stirred overnight. Saturated aq. $NaHCO_3$ (200 mL) was then added, and the reaction mixture was extracted with EtOAc (3 × 300 mL). The combined organic phases were washed with brine (200 mL), dried over anhydrous $Na_2SO_4$, and concentrated under vacuum with the aid of a rotary evaporator. The residue was purified by Biotage (100 g SNAP cartridge, 0-4% EtOAc/hexanes for 8 CV, then 4% EtOAc/hexanes for 10 CV) to afford the product as a pale-yellow oil (7.9 g, 45% yield).

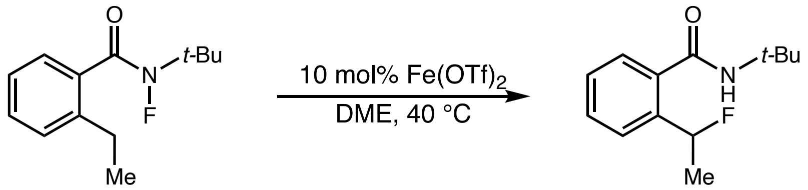

A round bottom flask was dried in an oven (110 °C) and charged with a stir bar. This flask was evacuated and backfilled with $N_2$ and this procedure was repeated for a total of three times. To this flask were added anhydrous $Fe(OTf)_2$ (318 mg, 0.898 mmol, 0.2 equiv) and a solution of fluoroamide (1.00 g, 4.49 mmol, 1.0 equiv) in anhydrous DME (22 mL, 0.2 M). The mixture was heated to 40 °C for 4 h. Then the mixture was diluted with 22 mL EtOAc and filtered through a silica plug and eluded with EtOAc (3 × 20 mL). The filtrate was concentrated under vacuum with the aid of a rotary evaporator. The residue was purified by Biotage (50 g SNAP cartridge, 0-10% EtOAc/hexanes for 8 CV, then 10% EtOAc/hexanes for 5 CV) to afford the fluoride product as a white powder (831 mg, 83% yield).

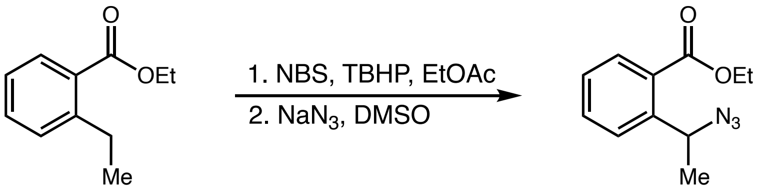

To a solution of methyl 2-ethylbenzoate (2.67 g, 15.0 mmol, 1.0 equiv) in ethyl acetate (30 mL, 0.2 M) was added *N*-bromosuccinimide (NBS, 3.20 g, 18.0 mmol, 1.2 equiv) and benzoyl peroxide (BPO, 0.440 g, 1.80 mmol, 0.12 eq). The mixture was purged with $N_2$ for 10 min, refluxed for 2 h, and then cooled to room temperature. 50 mL water was added, the organic layer was separated, and the reaction mixture was extracted with EtOAc (3 × 50 mL). The combined organic layers were dried over anhydrous $Na_2SO_4$ and concentrated under vacuum with the aid of a rotary evaporator. The crude product was dissolved in DMSO (30 mL, 0.2 M). $NaN_3$ (1.07 g, 16.5 mmol, 1.1 equiv) was then added. The mixture was allowed to stir at 22 °C for 12 h before saturated aq. $NaHCO_3$ (50 mL) was added, and the reaction mixture was extracted with EtOAc (50 mL) for a total of three times. The combined organic phases were washed with brine (50 mL), dried over anhydrous $Na_2SO_4$, and concentrated under vacuum with the aid of a rotary evaporator. The residue was purified by Biotage (50 g SNAP cartridge, 0-5% EtOAc/hexanes for 8 CV, then 5% EtOAc/hexanes for 10 CV) to afford the azide product as a colorless oil (2.46 g, 75% two steps). $^1$H NMR (400 MHz, $CDCl_3$) δ: 7.91 (dd, $J$ = 7.9, 1.2 Hz, 1H), 7.61 (dd, $J$ = 7.9, 1.4 Hz, 1H), 7.55 (td, $J$ = 7.7, 1.4 Hz, 1H), 7.35 (td, $J$ = 7.8, 1.4 Hz, 1H), 5.68 (q, $J$ = 6.7 Hz, 1H), 4.37 (q, $J$ = 7.1 Hz, 2H), 1.53 (d, $J$ = 6.7 Hz, 3H), 1.40 (t, $J$ = 7.1 Hz, 3H) ppm. $^{13}$C NMR (101 MHz, $CDCl_3$) δ: 167.1, 143.2, 132.7, 130.6, 128.9, 127.6, 126.9, 61.3, 57.4, 22.4, 14.3 ppm.

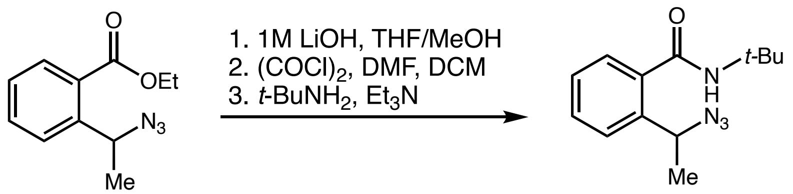

To a stirred solution of azide (500 mg, 2.28 mmol, 1.0 equiv) in THF/MeOH (20 mL, 1:1) was added aq. LiOH (12 mL, 1.0 M) slowly at 0 °C. The mixture was allowed to warm to room temperature for 2 h before aq. HCl (15 mL, 1.0 M) was added. The mixture was extracted with EtOAc (3 × 20 mL). The combined organic phases were washed with brine (20 mL), dried over anhydrous $Na_2SO_4$, and concentrated under vacuum with the aid of a rotary evaporator. The crude product was used directly for the next step without further purification.

To a stirred solution of the benzoic acid (330 mg, 1.72 mmol, 1.0 equiv) in $CH_2Cl_2$ (10 mL, 0.17 M) was added one drop *N,N*-dimethylformamide and oxalyl chloride (0.17 mL, 2.06 mmol, 1.50 equiv). The reaction was stirred at room temperature for 1 h. To another flask with a stir bar were added *tert*-butylamine (0.26 mL, 2.58mmol, 1.5 equiv), triethylamine (0.62 mL, 4.3 mmol, 2.5 equiv) and $CH_2Cl_2$ (5 mL, 0.3 M) at 0 ºC. The freshly prepared acid chloride was slowly added into this flask. The reaction mixture was allowed to warm to room temperature and stirred for another 1 h before aq. HCl (5 mL, 1.0 M) was added. The resultant mixture was extracted with EtOAc (3 × 10 mL). The combined organic phases were washed with brine (20 mL), dried over anhydrous $Na_2SO_4$, and concentrated under vacuum with the aid of a rotary evaporator. The residue was purified by Biotage (10 g SNAP cartridge, 0-10% EtOAc/hexanes for 8 CV, then 10% EtOAc/hexanes for 10 CV) to afford the amide product as a white solid (130 mg, 28% over two steps).

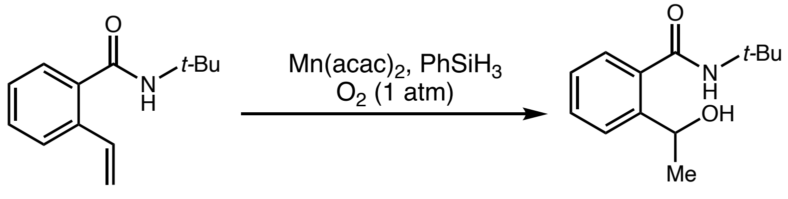

To a stirred solution of the styrene (1.11 g, 5.46 mmol, 1.0 equiv) and $PhSiH_3$ (3.4 mL, 27.3 mmol, 5 equiv) in EtOH (25 mL, 0.2 M) was added Mn(acac)$_2$ (138 mg, 0.546 mmol, 10 mol%)

at 22 °C. Oxygen was then bubbled into the mixture for 10 min. The mixture was then allowed to stir under an oxygen atmosphere at 22 °C for 5 h. The volatiles were removed under vacuum, and the residue was purified by Biotage (50 g SNAP cartridge, 0-30% EtOAc/hexanes for 8 CV, then 30% EtOAc/hexanes for 10 CV) to afford the alcohol as a colorless oil (800 mg, 66%).

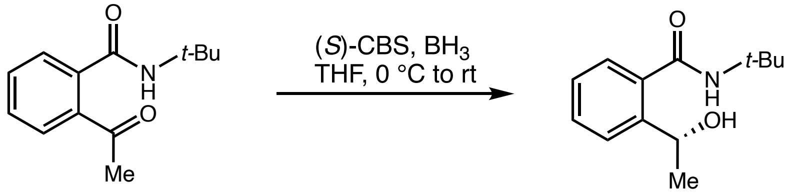

This procedure is modified from a previously described procedure, and the (*R*)-product was obtained in accordance with the Corey-Bakshi-Shibata reduction and its stereochemistry outcome.[6] To a stirred solution of (*S*)-1-Methyl-3,3-diphenylhexahydropyrrolo[1,2-c][1,3,2]oxazaborole (219 mg, 1.0 mmol, 1.0 equiv) and $BH_3$ in THF (1 M in THF, 1.5 mL, 1.5 mmol, 1.5 equiv) in THF (5 mL, 0.2 M) was added the ketone (219 mg, 1.0 mmol, 1.0 equiv) at 0 °C. The reaction mixture was allowed to warm to room temperature and stirred for another 6 h before 3 mL aq. HCl (1.0 M) was added. The resulting mixture was extracted with EtOAc (3 × 10 mL). The combined organic phases were washed with brine (20 mL), dried over anhydrous $Na_2SO_4$, and concentrated under vacuum with the aid of a rotary evaporator. The residue was purified by Biotage (10 g SNAP cartridge, 0-20% EtOAc/hexanes for 10 CV, then 20% EtOAc/hexanes for 5 CV) to afford the amide product as a white solid (88 mg, 40% yield, 60:40 e.r.). The absolute stereochemistry was established based on assignments in literature.[6]

**Characterization data of substrates and products**

***N*-(*Tert*-butyl)-2-ethyl-*N*-fluorobenzamide (1)**

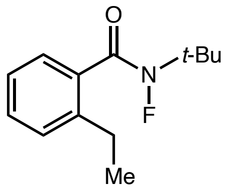

$^{1}$H NMR (400 MHz, $CDCl_3$) δ: 7.35 (ddd, $J$ = 7.6, 7.6, 1.5 Hz, 1H), 7.33 – 7.25 (m, 1H), 7.26 (s, 1H), 7.21 (ddd, $J$ = 7.4, 7.4, 1.3 Hz, 1H), 2.74 (q, $J$ = 7.6 Hz, 1H), 1.56 (d, $J$ = 2.0 Hz, 9H), 1.24 (t, $J$ = 7.6 Hz, 3H) ppm. $^{13}$C NMR (101 MHz, $CDCl_3$) δ: 175.2 (d, $J$ = 11.0 Hz), 141.7 (d, $J$ = 2.2 Hz), 134.8, 130.2 (d, $J$ = 1.4 Hz), 129.0, 127.2 (d, $J$ = 4.4 Hz), 125.5, 64.4 (d, $J$ = 10.5 Hz), 27.3 (d, $J$ = 5.7 Hz), 26.2, 15.8 ppm.$^{19}$F NMR (376 MHz, $CDCl_3$) δ: -63.4 (s) ppm. Spectral data match those previously reported.[5]

***N*-(*Tert*-butyl)-2-(1-fluoroethyl)benzamide (2)**

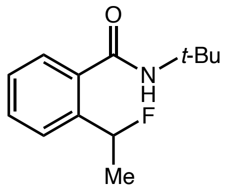

$^{1}$H NMR (400 MHz, $CDCl_3$) δ: 7.57 (d, $J$ = 7.7 Hz, 1H), 7.45 (ddd, $J$ = 7.5, 7.5, 1.6 Hz, 1H), 7.39 – 7.28 (m, *2*H), 6.03 (dq, $J$ = 47.7, 6.3 Hz, 1H), 5.77 (s, 1H), 1.68 (dd, $J$ = 24.1, 6.3 Hz, 3H), 1.45 (s, 9H) ppm. $^{13}$C NMR (101 MHz, $CDCl_3$) δ: 168.7, 140.0 (d, $J$ = 19.4 Hz), 135.7 (d, $J$ = 5.1 Hz), 130.3, 128.1 (d, $J$ = 1.9 Hz), 126.6, 125.9 (d, $J$ = 8.9 Hz), 88.8 (d, $J$ = 165.1 Hz), 52.1, 28.9, 23.2 (d, $J$ = 25.3 Hz) ppm.$^{19}$F NMR (376 MHz, $CDCl_3$) δ: -166.58 (dq, $J$ = 48.4, 24.1 Hz) ppm. Spectral data match those previously reported.[5]

For analytical scale biotransformation of substrate **1**, the HPLC analysis conditions are as follows: IC column, 97:3 hexanes: *i*-PrOH, 1.2 mL/min, 40 °C.

For biocatalytic reaction with $EgtB_{CHF1}$, HPLC analysis indicated 31% yield, 60:40 e.r.: $t_R$ = 9.74 (*R*, major), 10.38 (*S*, minor) min.

For biocatalytic reaction with $EgtB_{CHF2}$, HPLC analysis indicated 36% yield, 31:69 e.r.: $t_R$ = 9.75 (*R*, minor), 10.37 (*S*, major) min.

For biocatalytic reaction with $ACCO_{CHF}$, HPLC analysis indicated 89% yield, 5:95 e.r.: $t_R$ = 9.56 (*R*, minor), 10.12 (*S*, major) min.

**N-(tert-butyl)-2-(1-hydroxyethyl)benzamide (3)**

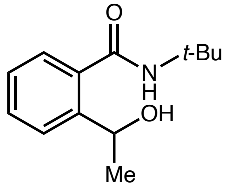

$^1$H NMR (400 MHz, $CDCl_3$) δ: 7.47 – 7.33 (m, 3H), 7.29 – 7.23 (m, 1H), 6.06 (s, 1H), 4.91 (q, $J$ = 6.6 Hz, 1H), 1.53 (d, $J$ = 6.6 Hz, 3H), 1.46 (s, 9H). $^{13}$C NMR (126 MHz, $CDCl_3$) δ: 170.7, 143.4, 136.6, 130.4, 127.4, 127.4, 126.9, 68.1, 52.1, 28.7, 22.0 ppm. IR: ν: 3280, 2970, 1633, 1539, 1452, 1393, 1364, 1313, 1221, 1076, 761, 665 $cm^{-1}$. HRMS (*m/z*): $[M + Na]^+$ calcd for $C_{13}H_{19}N_1O_2Na^+$ 244.1313, found 244.1314.

For analytical scale biotransformation of substrate **1**, the HPLC analysis conditions are as follows: IB-N5 column, 98:2 hexanes: *i*-PrOH, 1.2 mL/min, 40 °C.

For biocatalytic reaction with $EgtB_{CHF1}$, HPLC analysis indicated 32% yield, 75:25 e.r.: $t_R$ = 10.41 (major, *S*), 11.78 (minor, *R*) min.

For biocatalytic reaction with $EgtB_{CHF2}$, HPLC analysis indicated 17% yield, 66:34 e.r.: $t_R$ = 10.28 (major, *S*), 11.57 (minor, *R*) min.

For CBS reduction, HPLC analysis indicated 40:60 e.r. and allowed the establishment of absolute configuration based on literature report[6]: $t_R$ = 10.56 (minor, *S*), 11.588 (major, *R*) min.

**2-(1-Azidoethyl)-N-(tert-butyl)benzamide (4)**

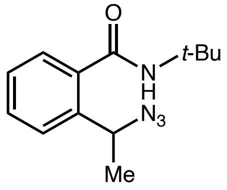

$^1$H NMR (400 MHz, $CDCl_3$) δ: 7.50 (dd, $J$ = 7.8, 1.4 Hz, 1H), 7.44 (ddd, $J$ = 7.9, 7.5, 1.6 Hz, 1H), 7.37 (dd, $J$ = 7.6, 1.6 Hz, 1H), 7.31 (td, $J$ = 7.4, 1.4 Hz, 1H), 5.76 (s, 1H), 5.15 (q, $J$ = 6.8 Hz, 1H), 1.58 (d, $J$ = 5.9 Hz, 3H), 1.47 (s, 9H) ppm. $^{13}$C NMR (101 MHz, $CDCl_3$) δ: 168.9, 139.1, 136.9, 130.4, 128.0, 127.0, 126.7, 57.5, 52.2, 28.9, 21.5 ppm. Spectral data match those previously reported.[7]

For analytical scale biotransformation of substrate **1**, the HPLC analysis condition are as follows: IC column, 97:3 hexanes: *i*-PrOH, 1.2 mL/min, 40 °C.

For biocatalytic reaction with $EgtB_{CHF1}$, HPLC analysis indicated 50% yield, 56:44 e.r.: $t_R$ = 7.05 (*S*, major), 7.62 (*R*, minor) min.

For biocatalytic reaction with $EgtB_{CHF2}$, HPLC analysis indicated 40% yield, 61:39 e.r.: $t_R$ = 7.16 (*S*, major), 7.74 (*R*, minor) min.

For biocatalytic reaction with $EgtB_{CHF2}$ H138A, HPLC analysis indicated 42% yield, 73:27 e.r.: $t_R$ = 7.17 (*S*, major), 7.76 (*R*, minor) min.

**IX. HPLC calibration curves**

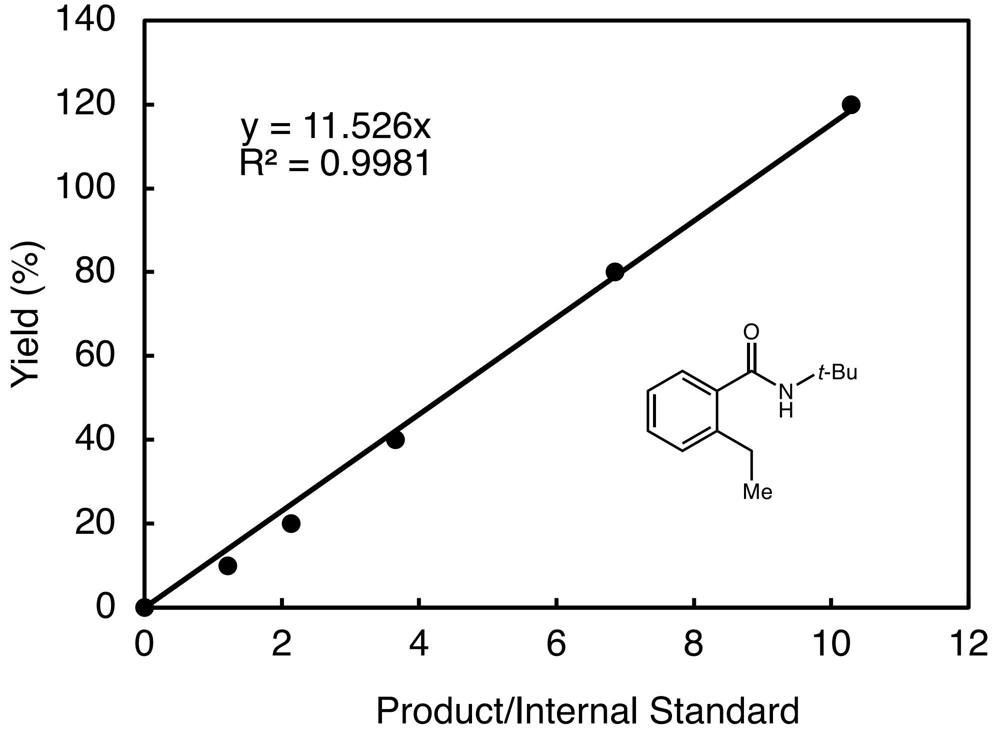

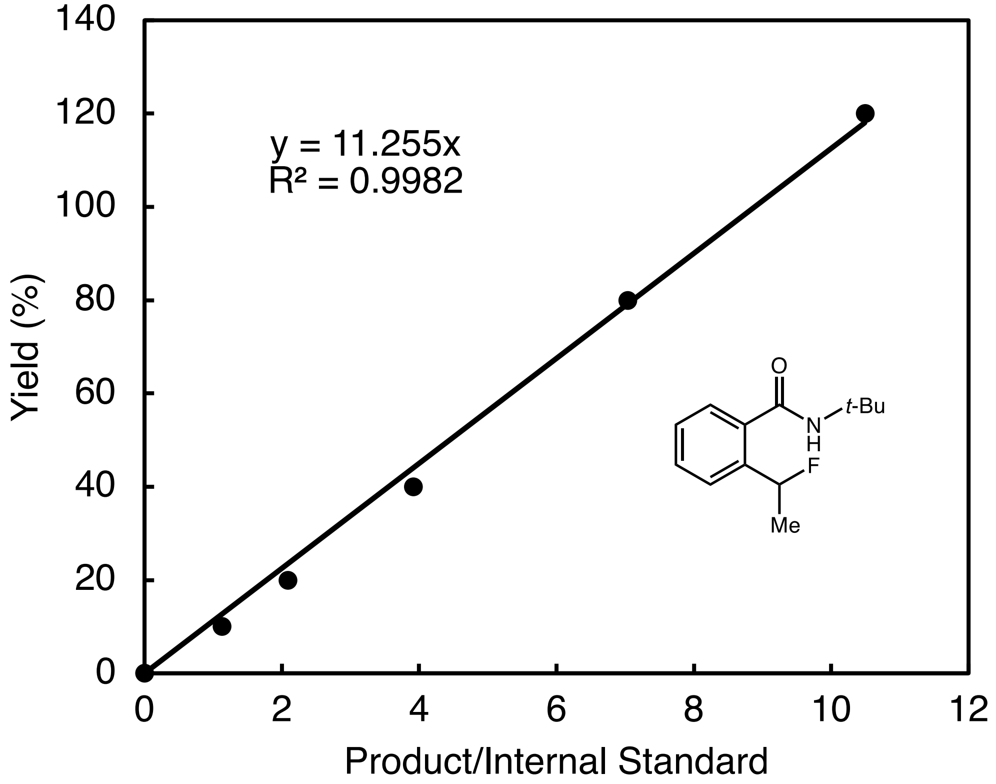

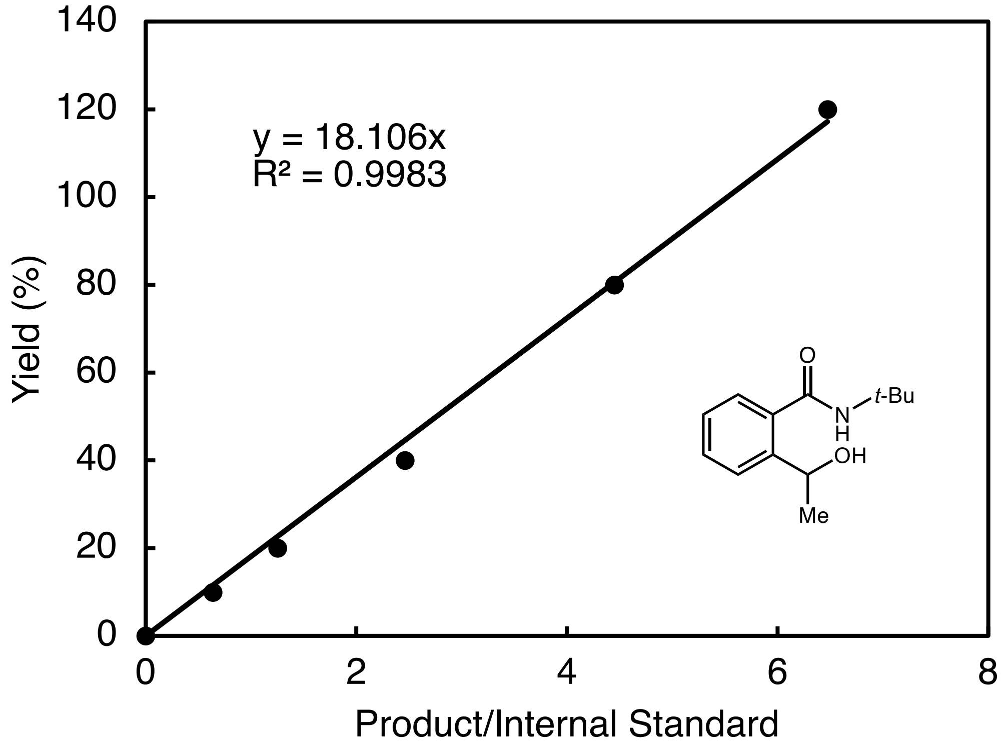

y = 18.106x
R² = 0.9983
Yield (%)
Product/Internal Standard
O
N
H
t-Bu
OH
Me

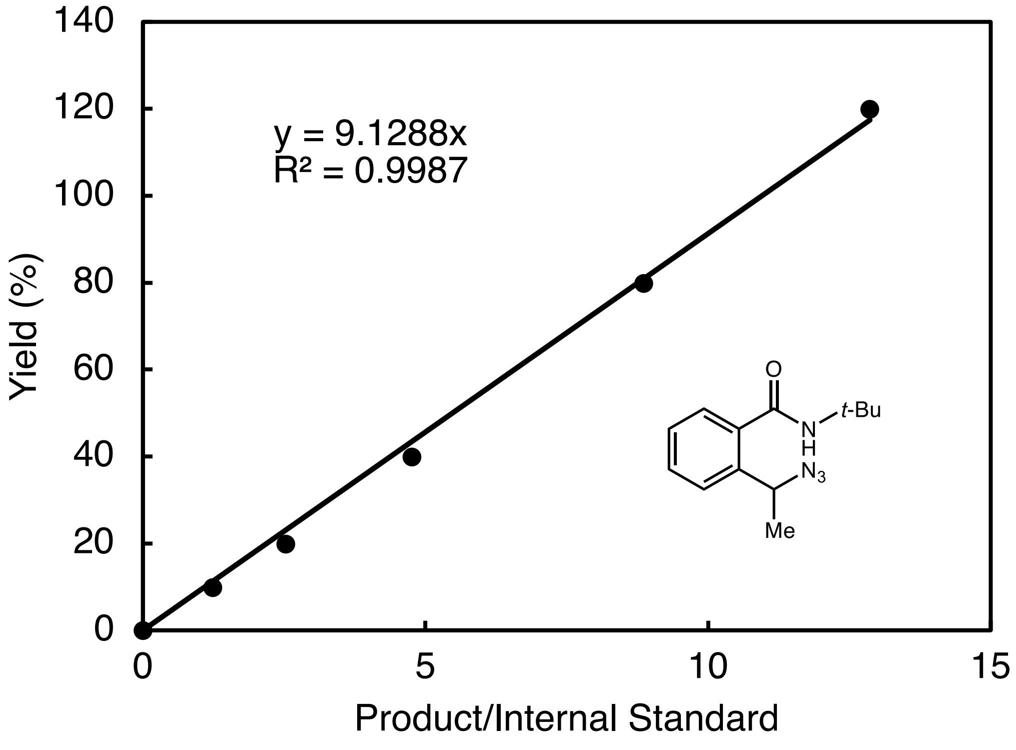

y = 9.1288x
R² = 0.9987
Yield (%)
Product/Internal Standard
O
N
H
t-Bu
N3
Me

#### X. Chiral HPLC traces

##### *N*-(*tert*-butyl)-2-(1-fluoroethyl)benzamide (2)

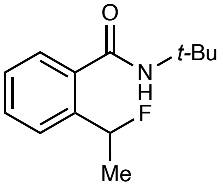

Racemic **2**:

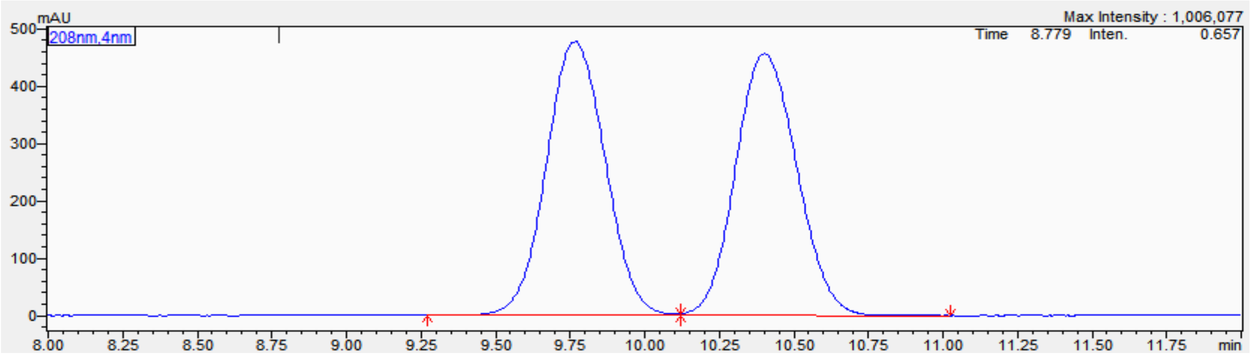

| Peak# | Ret. Time | Height | Area | Area% |
|---|---|---|---|---|
| 1 | 9.768 | 477341 | 6585286 | 50.151 |
| 2 | 10.403 | 456823 | 6545684 | 49.849 |
| Total | | 934164 | 13130970 | 100.000 |

Enantioenriched **2** obtained using EgtB$_{CHF1}$ (HPLC analysis): 60:40 e.r.

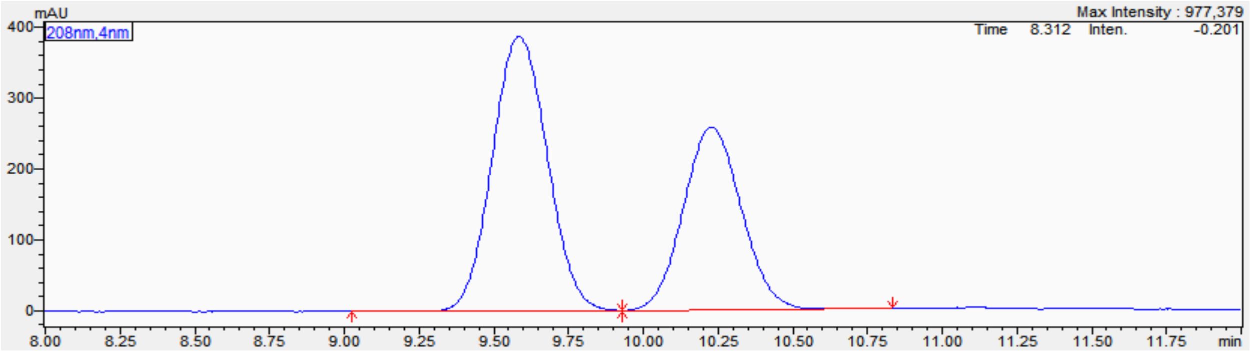

| Peak# | Ret. Time | Height | Area | Area% |
|---|---|---|---|---|
| 1 | 9.738 | 396542 | 5096452 | 59.984 |
| 2 | 10.384 | 255957 | 3399952 | 40.016 |
| Total | | 652499 | 8496404 | 100.000 |

Enantioenriched **2** obtained using $EgtB_{CHF2}$ (HPLC analysis): 31:69 e.r.

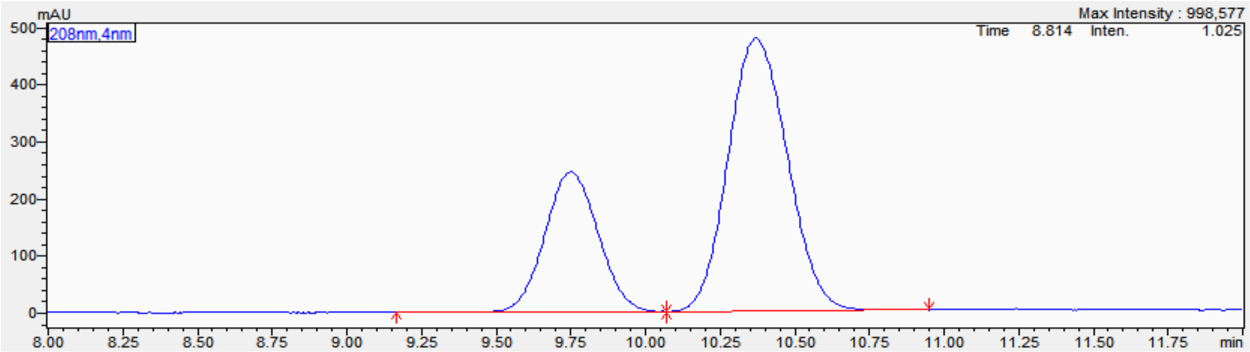

| Peak# | Ret. Time | Height | Area | Area% |
|---|---|---|---|---|
| 1 | 9.751 | 245367 | 3026199 | 31.425 |
| 2 | 10.370 | 477051 | 6603864 | 68.575 |
| Total | | 722419 | 9630064 | 100.000 |

Enantioenriched **2** obtained using $ACCO_{CHF}$ (HPLC analysis): 5.5:94.5 e.r.

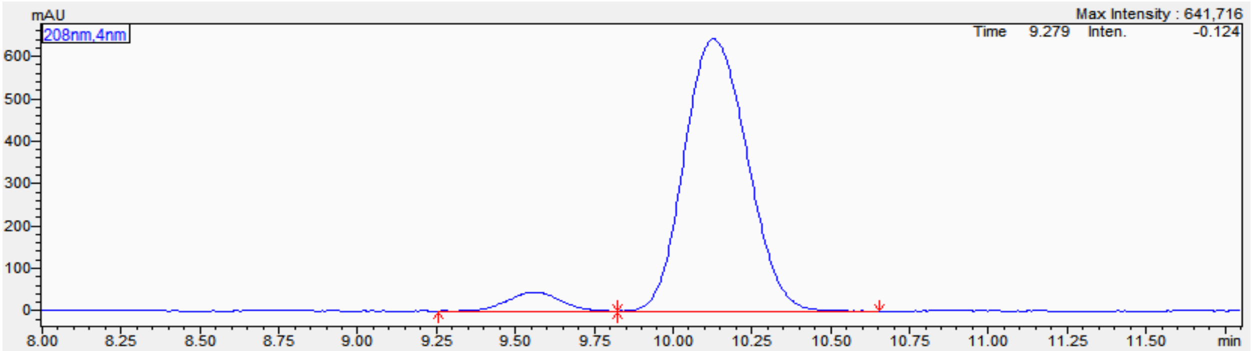

| Peak# | Ret. Time | Height | Area | Area% |
|---|---|---|---|---|
| 1 | 9.561 | 44496 | 523733 | 5.558 |
| 2 | 10.127 | 642184 | 8899936 | 94.442 |
| Total | | 686681 | 9423669 | 100.000 |

***N*-(*tert*-butyl)-2-(1-hydroxyethyl)benzamide (3)**

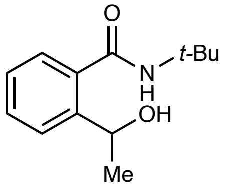

Racemic **3:**

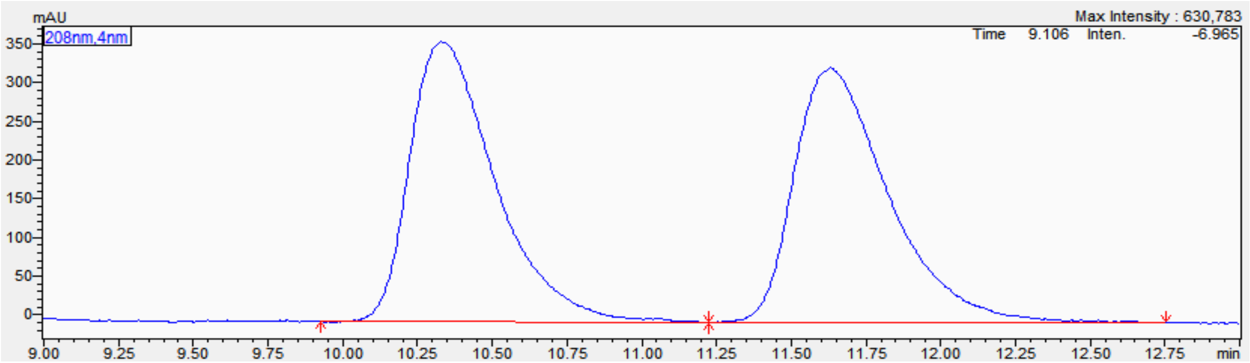

| Peak# | Ret. Time | Height | Area | Area% |
|---|---|---|---|---|
| 1 | 10.330 | 361986 | 7249855 | 49.904 |
| 2 | 11.631 | 329963 | 7277751 | 50.096 |
| Total | | 691949 | 14527606 | 100.000 |

Enantioenriched **3** obtained using $EgtB_{CHF1}$ (HPLC analysis):75:25 e.r.

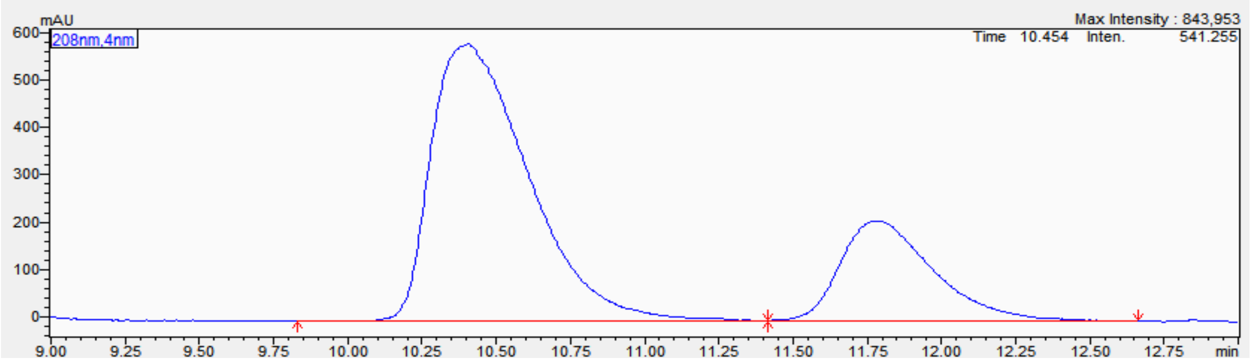

| Peak# | Ret. Time | Height | Area | Area% |
|---|---|---|---|---|
| 1 | 10.405 | 584949 | 13314664 | 74.564 |
| 2 | 11.780 | 210188 | 4542003 | 25.436 |
| Total | | 795138 | 17856667 | 100.000 |

Enantioenriched **3** obtained using $EgtB_{CHF2}$ (HPLC analysis): 66:34 e.r.

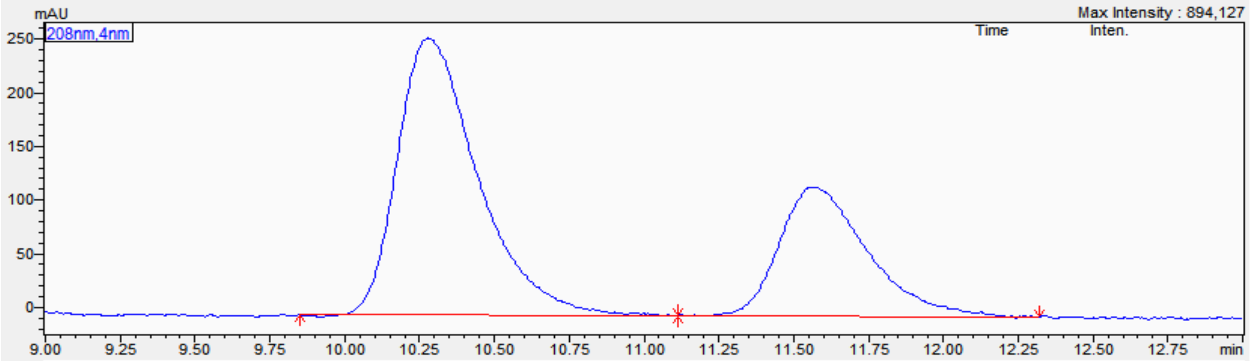

| Peak# | Ret. Time | Height | Area | Area% |
|---|---|---|---|---|
| 1 | 10.279 | 258023 | 4814053 | 66.358 |
| 2 | 11.573 | 120106 | 2440602 | 33.642 |
| Total | | 378129 | 7254654 | 100.000 |

Enantioenriched **3** obtained via CBS reduction (HPLC analysis): 40:60 e.r.

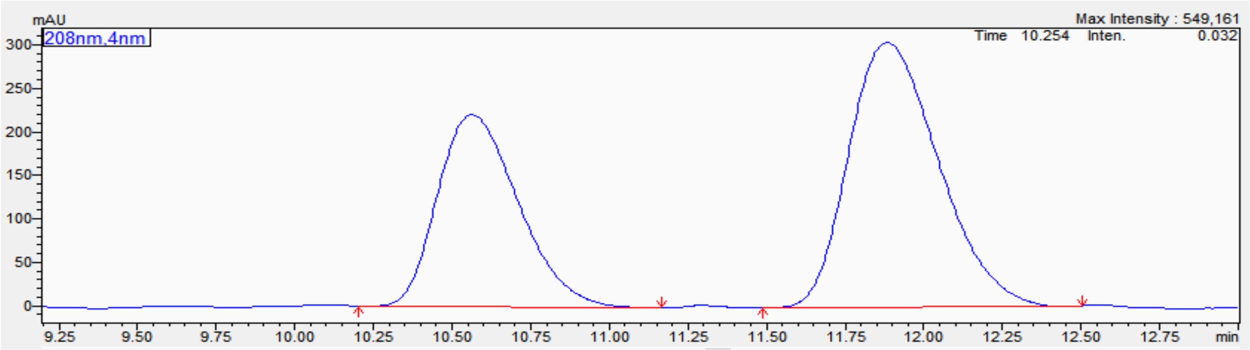

| Peak# | Ret. Time | Height | Area | Area% |
|---|---|---|---|---|
| 1 | 10.563 | 220395 | 3929951 | 39.411 |
| 2 | 11.884 | 303573 | 6041720 | 60.589 |
| Total | | 523968 | 9971671 | 100.000 |

**2-(1-azidoethyl)-*N*-(*tert*-butyl)benzamide (4)**

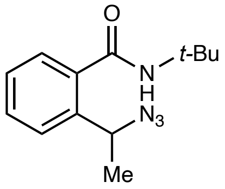

Racemic **4**:

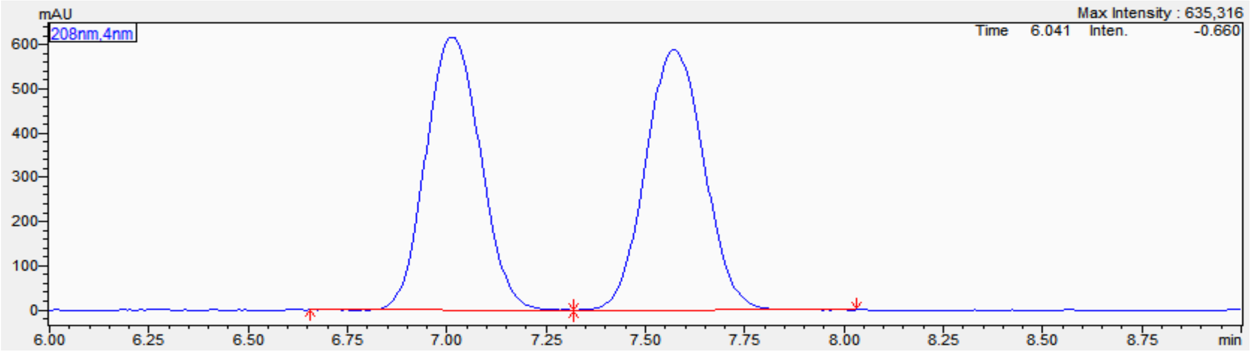

| Peak# | Ret. Time | Height | Area | Area% |
|---|---|---|---|---|
| 1 | 7.014 | 615700 | 5903252 | 49.427 |
| 2 | 7.572 | 589108 | 6040034 | 50.573 |
| Total | | 1204808 | 11943286 | 100.000 |

Enantioenriched **4** obtained using EgtB$_{CHF1}$ (HPLC analysis): 56:44 e.r.

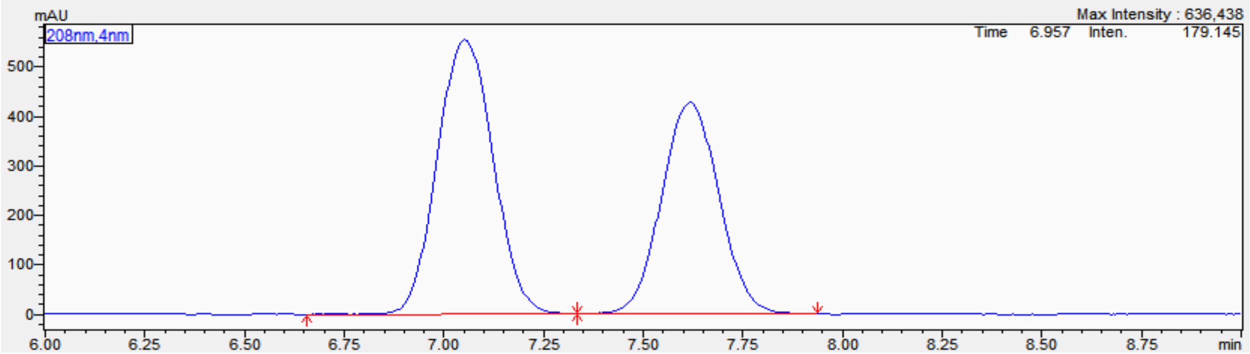

| Peak# | Ret. Time | Height | Area | Area% |
|---|---|---|---|---|
| 1 | 7.052 | 554920 | 5447961 | 55.841 |
| 2 | 7.617 | 427933 | 4308165 | 44.159 |
| Total | | 982853 | 9756126 | 100.000 |

Enantioenriched **4** obtained using $EgtB_{CHF2}$ (HPLC analysis): 61:39 e.r.

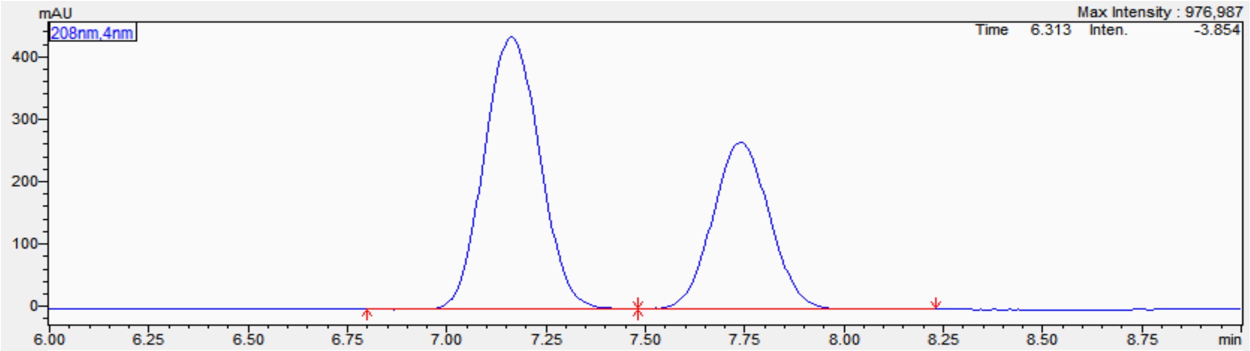

| Peak# | Ret. Time | Height | Area | Area% |
|---|---|---|---|---|
| 1 | 7.163 | 435750 | 4252681 | 61.494 |
| 2 | 7.740 | 267127 | 2662953 | 38.506 |
| Total | | 702877 | 6915634 | 100.000 |

Enantioenriched **4** obtained using $EgtB_{CHF2}$ H138A (HPLC analysis): 73:27 e.r.

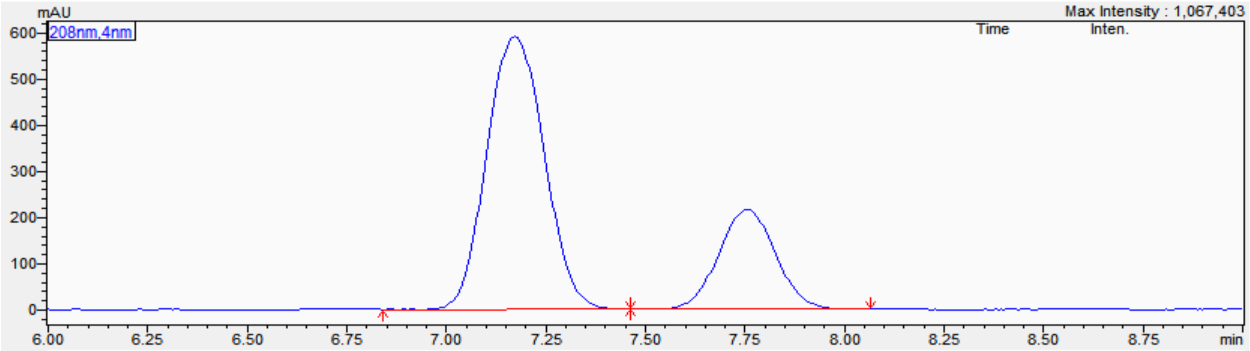

| Peak# | Ret. Time | Height | Area | Area% |
|---|---|---|---|---|
| 1 | 7.172 | 591130 | 5939553 | 73.479 |
| 2 | 7.755 | 216100 | 2143791 | 26.521 |
| Total | | 807230 | 8083344 | 100.000 |

#### XI. Computational Details

##### 1. Classical Molecular Dynamics (MD) Simulations

Classical MD simulations were performed to study the binding modes of *N*-fluoramide substrate (**1**) and active site residue–substrate interactions that promote the fluorine atom transfer from the substrate to the Fe(II) center. The starting structure of *Mth*EgtB was obtained from Protein Data Bank (PDB ID: 4X8B). To generate C–H fluorinase variants $EgtB_{CHF1}$ and $EgtB_{CHF2}$, appropriate residues were mutated using the Mutagenesis tool in PyMOL.[8] For variant $EgtB_{CHF1}$, mutated residues include Y377W W415R R87D, T141I, F83Y, R379E, V84R, H417L, and A145T. For variant $EgtB_{CHF2}$, mutated residues include Y377W, W415R, R87D, T141M, R379A, A145K, and Q422D (**Table 2** of the manuscript). The geometry of substrate *N*-fluoroamide **1** was optimized using the B3LYP functional[9] and 6-31G(d,p) basis set in *Gaussian 16*.[10] The optimized geometry of the substrate was then docked into $EgtB_{CHF1}$ and $EgtB_{CHF2}$ variants using AutoDock[11] with the Lamarckian genetic algorithm. A grid box with dimensions of 40 Å, 40 Å, and 40 Å was used, which center was set to be close to the Fe center. Docking parameters were set as follows: genetic algorithm run of 30, population size of 150, and 25 million energy evaluations. Because *N*-fluoroamide **1** may approach the Fe center at the coordination site *trans* to either residue H51 or H134, in our MD simulations, we considered these two binding modes **A** and **B**, where *N*-fluoroamide **1** is *trans* to H51 and H134, respectively (**Figure S10**). The best scored geometry for each binding mode from the docking calculations were then used to construct the initial input geometry for classical MD simulations. Because the potential substrate binding site *trans* to H138 is occupied by Q55 and W377 residues and our docking calculations did not locate docked substrate located proximal to that site, this binding mode was not considered in subsequent classical MD simulations.

Classical MD simulations were carried out using the pmemd module[12] of the GPU-accelerated Amber 16 software.[13] The Amber ff14SB force field[14] was used in all classical MD simulations. Force field parameters for the $Fe(II)(His)_3(H_2O)_2$(**1**) complexes in two binding modes were generated using the MCPB.py module[15] with the general Amber force field (gaff).[16] Using the

Merz-Singh-Kollman scheme,[17] RESP charge fitting[18] on electrostatic potential generated at the B3LYP/6-31G(d) level of theory was performed to generate partial charges at the high-spin quintet state, which was calculated to be the ground state of non-heme Fe(II) complexes.[19] Protonation states of enzyme residues were determined using the H++ server.[20] The enzyme was then put into a solvated cuboid box with the periodic boundary condition using the TIP3P water model.[21] The minimum distance between the enzyme surface and the edge of the water box was set to 10 Å. Water molecules were treated with the SHAKE algorithm.[22] The system was neutralized by adding $Na^+$ counterions. Long-range electrostatic was calculated using the particle-mesh-Ewald method.[23] Lennard-Jones and electrostatic interaction cut-offs were set to 12 Å.

We first performed energy minimization with positional restraints for the protein by applying a force constant of 500 $kcal\cdot mol^{-1}\cdot Å^{-2}$ in 30,000 steps. Next, the system was gradually heated from 0 K to 300 K in 200 ps, which is followed by an equilibration using the isothermal–isobaric ensemble (NPT) in the next 1.5 ns. Finally, production MD simulations were run in 500 ns using the same conditions as the equilibration with a time step of 2 fs. In our MD simulations, to simulate the substrate near attack conformation in the fluorine atom transfer process and prevent undesired substrate dissociation events, the Fe–F distance was restrained in a range of 3.0 – 3.2 Å with a harmonic potential of 100 $kcal\cdot mol^{-1}\cdot Å^{-2}$. The restrained Fe–F distance was determined from DFT calculation with a small model, *i.e.*, an $Fe(II)(Im)_3(H_2O)_2$(**1**) complex, where the fluorine atom of substrate **1** form a weak dative bond with the Fe center with an Fe–F distance of 3.18 Å (**Figure S9**). After the MD simulations, clustering analysis was carried out using the cpptraj module[24] to identify the most populated structure in the last 400 ns of each MD simulation. The RMSD value was used as the distance metric for clustering analysis.

Our MD simulations demonstrate that in binding mode **A**, *N*-fluoroamide **1** forms hydrogen bonds with residue W415R in the simulations with both $EgtB_{CHF1}$ and $EgtB_{CHF2}$ variants. The N–H···O distance between the side chain of residue W415R and the amide carbonyl oxygen of **1** is less than 2.5 Å in most of the MD simulation times (61.2% and 81.9% for $EgtB_{CHF1}$ and $EgtB_{CHF2}$ variants, respectively; see **Figure S11**). On the other hand, in binding mode **B**, *N*-fluoroamide **1** does not have any noticeable stabilizing interaction with residues in the active site. This result suggests that binding mode **B** is less favorable than binding mode **A**.

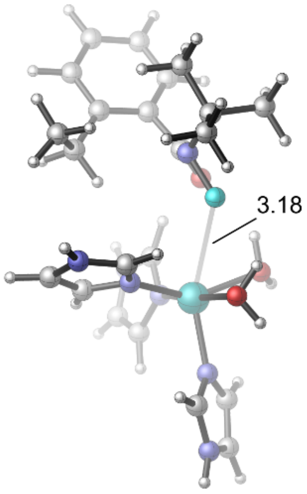

**Figure S9**. Optimized structure of the complex of *N*-fluoroamide (**1**) and a small model Fe(II)(Im)$_3$($H_2O$)$_2$ where histidine residues are replaced by imidazole molecules at high-spin quintet state. The geometry optimization was carried out at the B3LYP-D3(BJ)/SDD(Fe)–6-31G(d) level of theory.

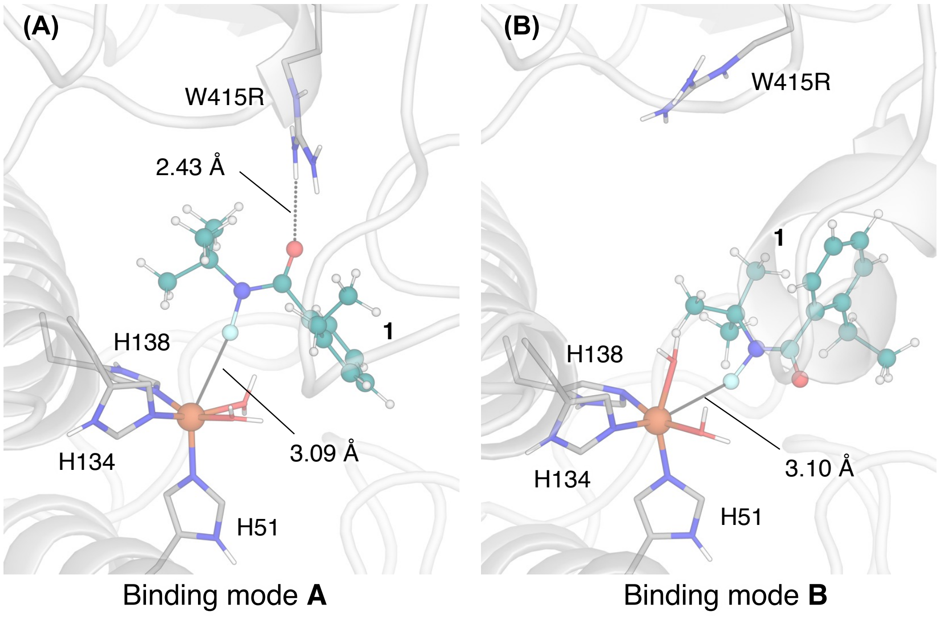

**Figure S10**. The most populated structure of the EgtB$_{CHF1}$ variant in complex with *N*-fluoroamide **1** with in (**A**) binding mode **A** (**1** is *trans* to H51) and (**B**) binding mode **B** (**1** is *trans* to H134).

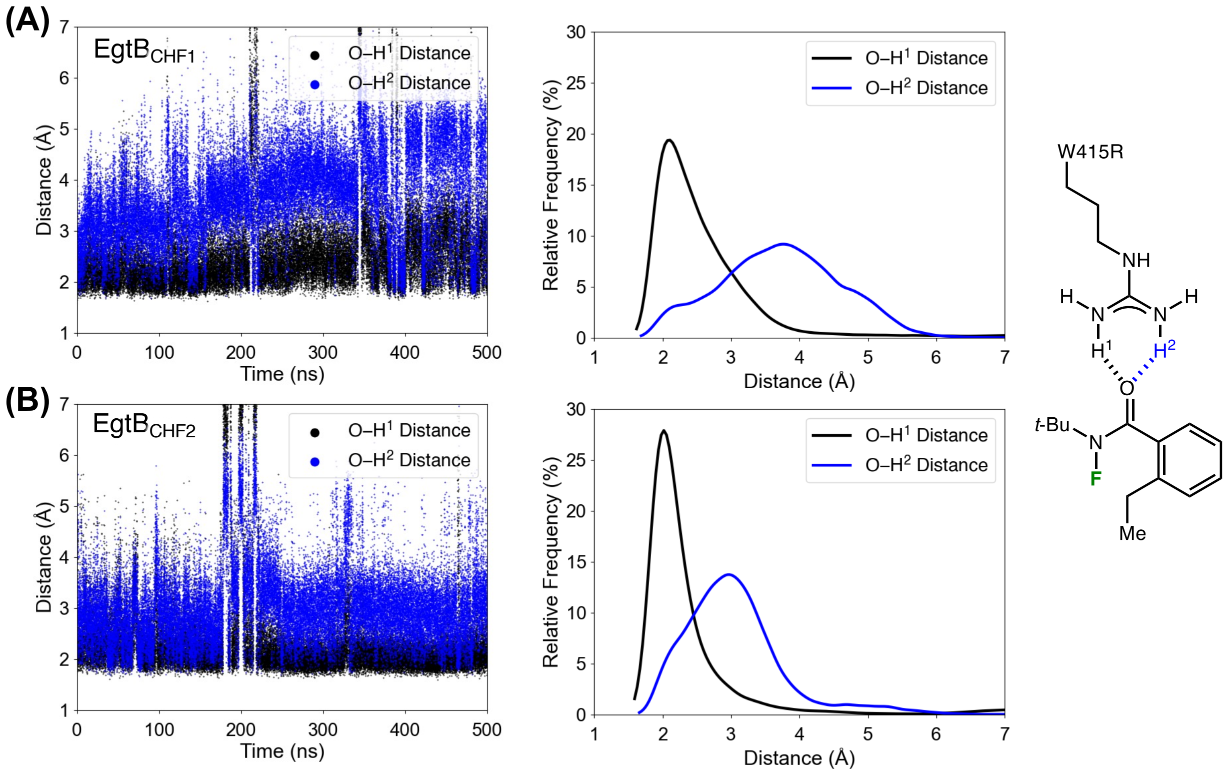

**Figure S11**. Time evolution and distribution of N–H···O distances between the side chain of residue W415R and the amide carbonyl oxygen of **1** in complex with (**A**) EgtB$_{CHF1}$ and (**B**) EgtB$_{CHF2}$ variants in binding mode **A** (**1** is *trans* to H51).

#### 2. Arginine/π interaction between R175 and the phenyl group of the *N*-fluoroamide 1 in ACCO$_{CHF}$

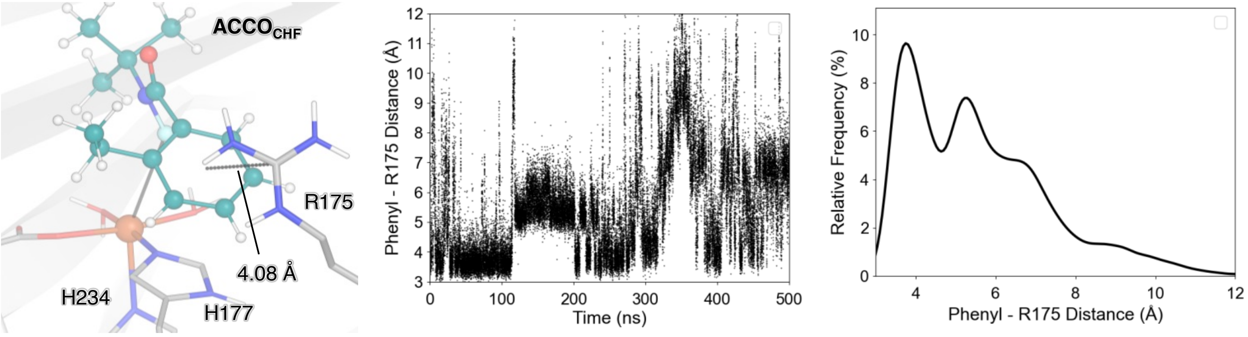

**Figure S12**. Time evolution and distribution of arginine/π interaction between R175 and the phenyl group of the *N*-fluoroamide **1** in ACCO$_{CHF}$.

#### 3. Density Functional Theory Calculations

DFT calculations were performed using the Gaussian 16 software.[10] Geometries of intermediates and transition states were optimized using the dispersion-corrected[25] B3LYP-D3(BJ) functional[17c, 26] with the SDD basis set for Fe atom and 6-31G(d) basis set for other atoms. Vibrational frequency calculations were performed at the same level of theory as their respective geometry optimizations to confirm that the obtained transition states connect their respective minima. Quasi-harmonic approximation was performed with the Cramer-Truhlar approach[27] using the GoodVibes package[28], in which all vibrational frequencies below 100 $cm^{-1}$ were shifted to 100 $cm^{-1}$ in entropy calculations. Single-point energy calculations were carried out using the B3LYP-D3(BJ) functional with the def2-TZVP basis set.[29] Solvation energy corrections were calculated in the single-point energy calculations using the SMD solvation model.[30] Diethyl ether solvent ($\varepsilon$ = 4.3) was used to mimic the actual dielectric constant within enzyme active site.[31] High and intermediate spin states for all species were calculated (*i.e.*, quintet and triplet states for Fe(II) species and sextet and quartet states for Fe(III) species).

Local force constants of Fe–F bond in all $Fe^{III}F$ species were calculated using the local modes method[32] at B3LYP-D3(BJ)/6-31G(d)–SDD(Fe) level of theory (i.e., optimization level).

#### 4. QM/MM Calculations

The ONIOM algorithm[33] implemented in Gaussian 16 was used in QM/MM (hybrid quantum mechanics/classical mechanics) geometry optimizations of intermediates and transition states. The entire enzyme, substrate, water molecules, and counter ions within 5 Å from the enzyme were included in the QM/MM calculations. The QM region includes the Fe atom, H51, H134, H138, W415R residues, and the substrate **1**, which has 107 atoms and a total charge of +1. Only high-spin state was calculated because our DFT calculations (see later) indicated this state is the ground states of intermediates and TSs.

For the QM region, the B3LYP-D3(BJ)/6-31G(d)–SDD(Fe) level of theory was used in geometry optimization and vibrational frequency calculations, and the B3LYP-D3(BJ)/def2-TZVP level of theory was used in single-point energy calculations. For the MM region, the same force field parameters from the classical MD simulations were used. Residues, water molecules, and $Na^+$ counterions greater than 10 Å away from the QM region were fixed during geometry optimization. The quadratic coupled algorithm[34] and the mechanical embedding scheme were used in geometry optimization. Single-point energy calculations were performed with the electronic embedding scheme, which better describes electrostatic interactions between QM and MM regions.[35]

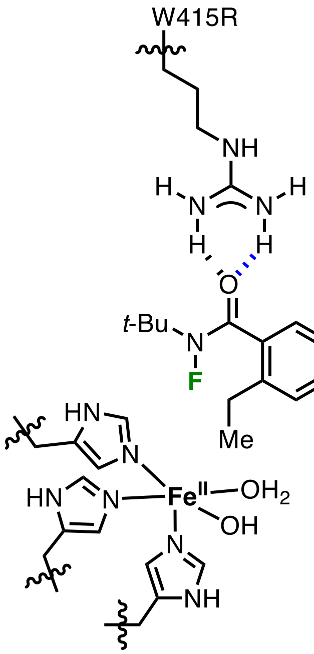

**Scheme S1**. QM region used in QM/MM Calculations. Wavy lines represent QM-MM boundary.

#### 5. Bond Dissociation Enthalpy Calculations and Effect of Hydrogen Bond on N–F Bond

Bond dissociation enthalpies (BDEs) of N–F, Fe–F, and C–F bonds were calculated at the B3LYP-D3(BJ)/def2-TZVP/SMD($Et_2O$)//B3LYP-D3(BJ)/6-31G(d)–SDD(Fe) level of theory (**Figure S13**).

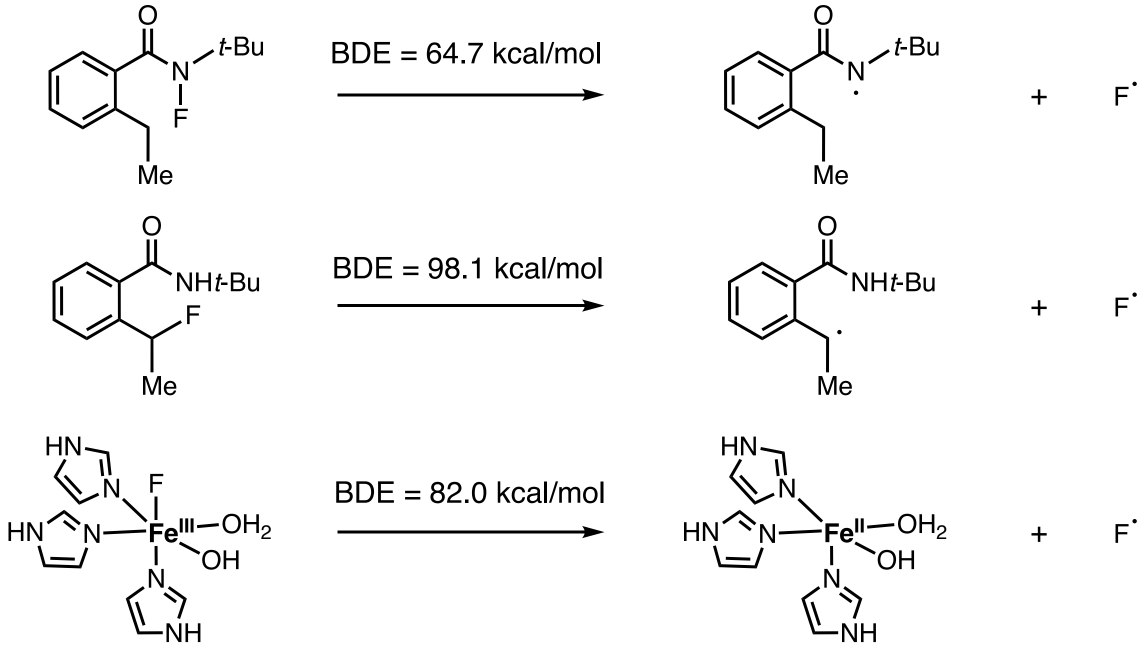

**Figure S13**. Bond dissociation enthalpy (BDE) calculations. Fe(II) and Fe(III)–F intermediates were calculated at high-spin quintet and sextet states, respectively.

Our calculations show that the N–F BDE decreases to 62.9 kcal/mol in the hydrogen bond complex with guanidium, compared to the N–F BDE of 64.7 kcal/mol in *N*-fluoroamide **1** itself (**Figure S14**). On the other hand, the N–F BDE in a hydrogen bond complex with a water molecule is 64.1 kcal/mol, which is only 0.6 kcal/mol weaker than the N–F BDE of **1**. These results indicate that the activation effect of hydrogen bond with the guanidium group in the W415R side chain is stronger than the hydrogen bond with water.

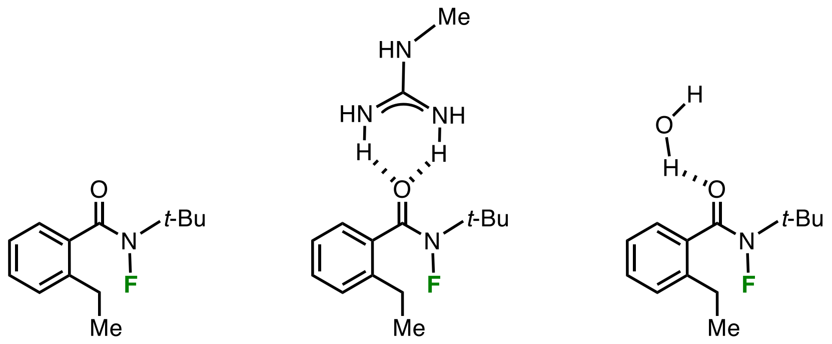

BDE (N–F) 64.7 kcal/mol | BDE (N–F) 62.9 kcal/mol | BDE (N–F) 64.1 kcal/mol

**Figure S14**. Effect of hydrogen bonds on N–F BDEs.

#### 6. Protonation State of Water Molecules Bound to the Fe Center

Because the protonation states of water molecules bound to the Fe center are unknown, we calculated the *pK*<sub></sub>$pK_a$ values of several water-coordinated $(Im)_3$Fe(II) and $(Im)_3$Fe(III)–F species (Im = imidazole). We used the $SMD_{sSAS}$ approach described by Smith *et al.*[36] to calculate the *p*Ka values in aqueous solution (**Scheme S2**). We used the same scaling factor (α = 0.485) proposed by Smith to generate the solute cavity. In our own benchmark studies[37] using hexaaqua Fe(III) complex as a test compound, a better agreement with experimental *p*Ka value was obtained with the $SMD_{sSAS}$ approach compared with the calculations using default SMD parameters.

We computed Gibbs free energies of the protonated form (*i.e.*, two water molecule bound) of the Fe(II) and Fe(III)–F species ($G^*_{\mathrm{aq}}(\mathrm{AH}^+)$) and the corresponding conjugate bases ($G^*_{\mathrm{aq}}(\mathrm{A})$) using DFT at the B3LYP-D3(BJ)/def2-TZVP/SMD$_{\mathrm{sSAS}}$($H_2O$)//B3LYP-D3(BJ)/6-31G(d)–SDD(Fe) level of theory. The Gibbs free energy of a proton in aqueous solution ($G^*_{\mathrm{aq}}(\mathrm{H}^+)$) was calculated from the gas-phase free energy of proton ($G^{\mathrm{o}}_{\mathrm{g}}(\mathrm{H}^+) = -6.28\ \mathrm{kcal/mol}$)[38] and its hydration free energy ( $\Delta G_{\mathrm{aq,solv}}(\mathrm{H}^+) = -265.9\ \mathrm{kcal/mol}$ ) from literature.[39] $\Delta G^{\mathrm{o}\rightarrow *} = RT\ln(24.47) = 1.89\ \mathrm{kcal/mol}$ is the free energy change to convert from the standard state of ideal gas (1 atm) to the standard state in solution (1 M).[30]

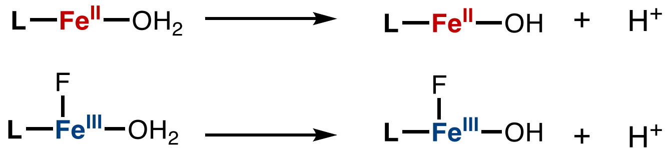

$$\mathrm{AH}^+(\mathrm{aq}) \xrightarrow{\Delta G^*_{\mathrm{aq}}} \mathrm{A}(\mathrm{aq}) + \mathrm{H}^+(\mathrm{aq}) \quad \textbf{(S1)}$$

$$\mathrm{p}K_{\mathrm{a}} = \frac{\Delta G^*_{\mathrm{aq}}}{2.303RT} \quad \textbf{(S2)}$$

$$\Delta G^*_{\mathrm{aq}} = G^*_{\mathrm{aq}}(\mathrm{A}) + G^*_{\mathrm{aq}}(\mathrm{H}^+) - G^*_{\mathrm{aq}}(\mathrm{AH}^+) \quad \textbf{(S3)}$$

$$G^*_{\mathrm{aq}}(\mathrm{H}^+) = G^{\mathrm{o}}_{\mathrm{g}}(\mathrm{H}^+) + \Delta G_{\mathrm{aq,solv}}(\mathrm{H}^+) + \Delta G^{\mathrm{o}\rightarrow *} \quad \textbf{(S4)}$$

**Scheme S2**. Acid dissociation constant (*p*Ka) calculations.

The computed aqueous *p*Ka values indicate that the *p*Ka of water significantly decreases when bound to either an Fe(II) or Fe(III) center. $H_2O$ bound to Fe(III) shows a lower *p*Ka relative to that bound to Fe(II). It should be noted that the enzyme active-sit environment further modulates the *p*Ka of Fe-bound water molecules. Therefore, these computational results alone cannot conclusively identify the protonation state of the Fe(II) species although Fe(III)–F compounds in EgtB$_{\mathrm{CHF}}$ (three-histidine facial triad and two-histidine facial dyad) are likely deprotonated under the experimental conditions considering their low computed *p*Ka values.

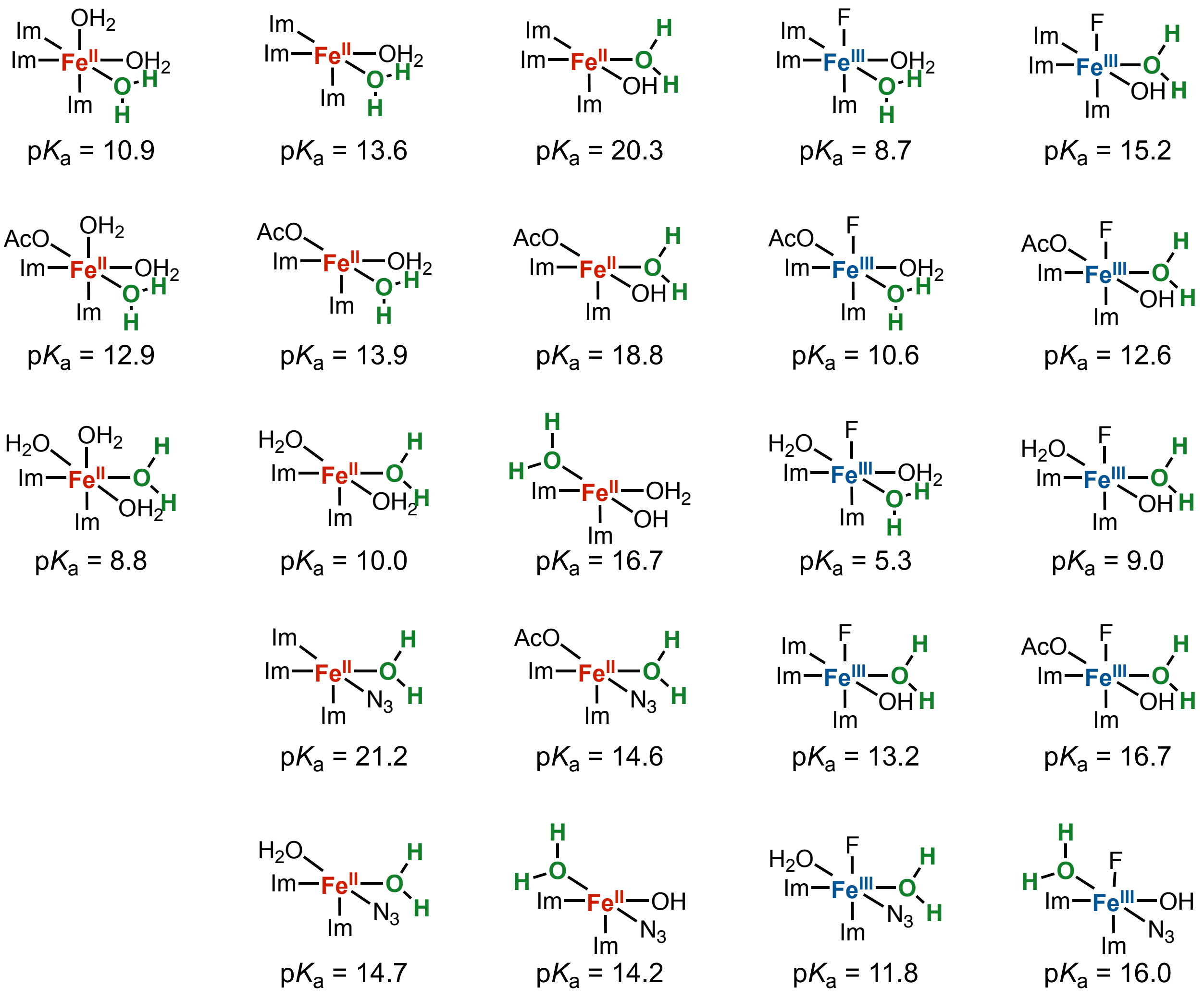

**Figure S15**. DFT-computed *p*Ka values of water-coordinated Fe(II) and Fe(III)–F compounds (the specific $H_2O$ studied is highlighted in green). Fe(II) and Fe(III)–F intermediates were calculated at high-spin quintet and sextet states, respectively.

Considering that the enzyme environment can substantially modulate *p*Ka values through hydrogen-bonding interactions, active-site water organization, and substrate-induced effects, *p*Ka calculations were performed using theozyme models. The initial geometries of all theozyme models were extracted from the most populated structures obtained from classical MD simulations. Our theozyme model for $EgtB_{CHF1}$ enzyme consists of *N*-fluoroamide **1**, the Fe atom, two water molecules bound to the Fe atom, and five active site residues represented by their side chains and

Cα atoms: H51, H134, and H138, which coordinate the Fe center; W415R, which forms a hydrogen-bonding interaction with substrate **1**; and Q137, which is in close contact with the substrate. In addition, two second-shell water molecules that form hydrogen bond to the Fe-bound water ligands were included.

In addition, the theozyme model for $ACCO_{CHF}$ enzyme consists of *N*-fluoroamide **1**, the Fe atom, two water molecules bound to the Fe atom, and five active site residues represented by their side chains and Cα atoms: D179, H177, and H234, which coordinate the Fe center; R175, which forms a cation–π interaction with substrate **1**; N216, which hydrogen bonds with D179 and the Fe-bound water molecules; and T178, which is located between D179 and H177 in the active site.. Three additional second-shell water molecules that participate in hydrogen-bonding interactions with substrate **1**, the Fe-bound water ligands, and D179 were also included. Main chain carbonyl and amino groups of these residues were replaced with C–H bonds. To maintain the locations of amino acid residues as in the most populated structure of the enzyme, the positions of Cα and added H atoms of all residues included in the theozyme were fixed during geometry optimizations. DFT calculations were performed at the B3LYP-D3(BJ)/def2-TZVP/$SMD_{sSAS}$($H_2O$)//B3LYP-D3(BJ)/6-31G(d)–SDD(Fe) level of theory. Similar active site cluster/theozyme approaches have previously been applied to investigate protonation equilibria and pKa shifts in enzyme active sites, including carbonic anhydrase[40] and papain.[41]

Interestingly, the *p*Ka values obtained from the theozyme calculations give a consistent trend with those derived from the truncated models. In both models, the water molecule bound to Fe(III) is predicted to be substantially more acidic than that bound to Fe(II). Notably, the enzyme active-site environment further modulates the acidity of the Fe-bound water ligands through local electrostatic and hydrogen-bonding interactions. Taken together, these results suggest that the Fe(III)–F intermediates in $EgtB_{CHF}$ (three-histidine facial triad and two-histidine facial dyad) are predominantly present in their deprotonated forms under the experimental conditions.

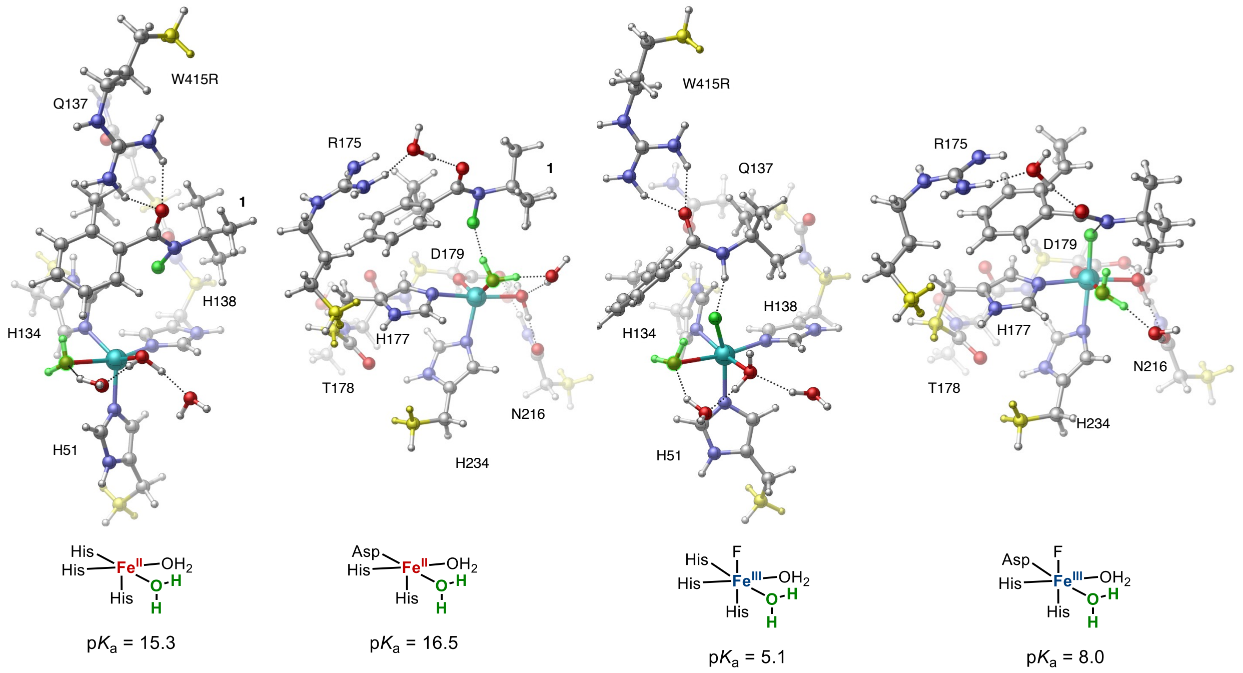

**Figure S16**. *p*Ka calculation using theozyme models. The specific $H_2O$ studied is highlighted in green. Positions of the Cα and added H atoms (colored in yellow) of active site residues were constrained during geometry optimizations. DFT calculations were performed at high-spin quintet state.

#### 7. Protonation Abstraction from Fe(III)–F Species

The proton abstraction step from Fe(III)–F species by acetate as a base was compute at the B3LYP-D3(BJ)/def2-TZVP/$SMD_{sSAS}$($H_2O$)//B3LYP-D3(BJ)/6-31G(d)–SDD(Fe) level of theory.

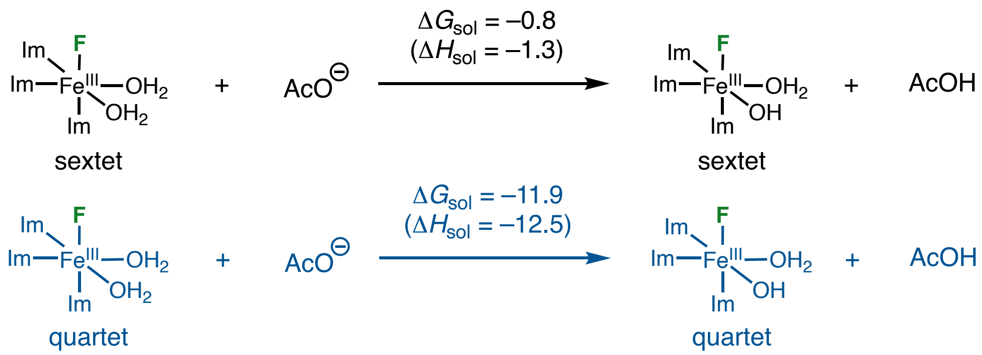

**Figure S17**. Proton abstraction from Fe(III)–F species. Fe(II) and Fe(III)–F intermediates were calculated at high-spin quintet and sextet states, respectively.

To identify potential proton acceptors involved in deprotonation of the Fe-bound water molecule, we further analyzed the classical MD simulation trajectories. In the most populated conformations of both $EgtB_{CHF1}$ and $EgtB_{CHF2}$ enzymes, we found that the D81 and D87 residues are located proximity to the non-heme iron center. The carboxylate groups of these residues are within ~7 Å of the Fe center and are connected to the active site through a network of water molecules. Such an arrangement could enable proton transfer from the Fe-bound water molecule through a water-wire network, implicating D81 and D87 as potential proton acceptors. (**Figure S18A**).

To further evaluate the feasibility of this process, we performed DFT calculations using a theozyme model that explicitly includes the Fe-bound water molecule, the bridging water molecules, and the D81 and D87 proton acceptors. The calculated activation enthalpies (**Figure S18B**) indicates that proton transfer through the proposed water-wire pathway is kinetically facile, with a sufficiently low barrier to compete with the subsequent 1,5-hydrogen atom transfer step. These results support the feasibility of Fe(III)–OH species formation prior to radical rebound, providing a mechanistic rationale for the experimentally observed hydroxylation pathway.

Taken together, the observed water organization provides a structural basis for proton transfer from the Fe-bound water molecule to D81/D87 through the hydrogen-bonded water molecules. Combined with the DFT calculations showing kinetically feasible proton transfer, these results support formation of the Fe(III)–OH species under the experimental conditions, consistent with the observed pH dependence of hydroxylation.

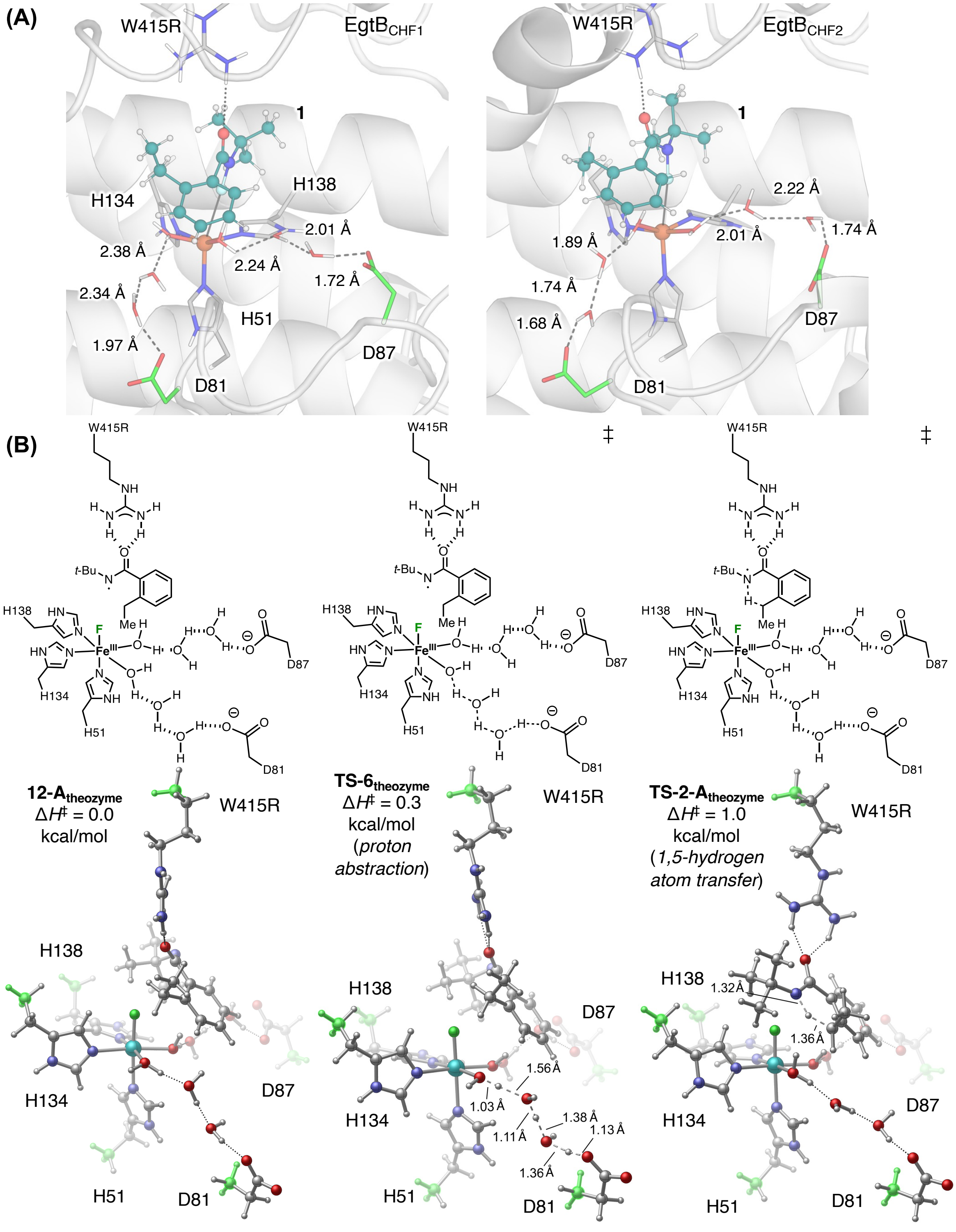

**Figure S18.** Computational studies for the protonation abstraction from Fe(III)–F species. (**A**) The most populated structure of the EgtB$_{CHF1}$ and EgtB$_{CHF2}$ variants obtained from classical MD simulations. Residues D81 and D87 are proximal to the non-heme iron center and connected to the active site through water molecules, suggesting D81 and D87 as potential proton acceptors for the water deprotonation. (**B**) DFT calculations using theozyme model. Positions of the Cα and added H atoms (highlighted in green) of active site residues were constrained during geometry optimizations. DFT calculations were performed at high-spin quintet state. Enthalpies are relative to **12-A**$_{\mathbf{theozyme}}$.

#### 8. Reaction Energy Profile of Nonheme Fe Enzyme-Catalyzed C–H Fluorination

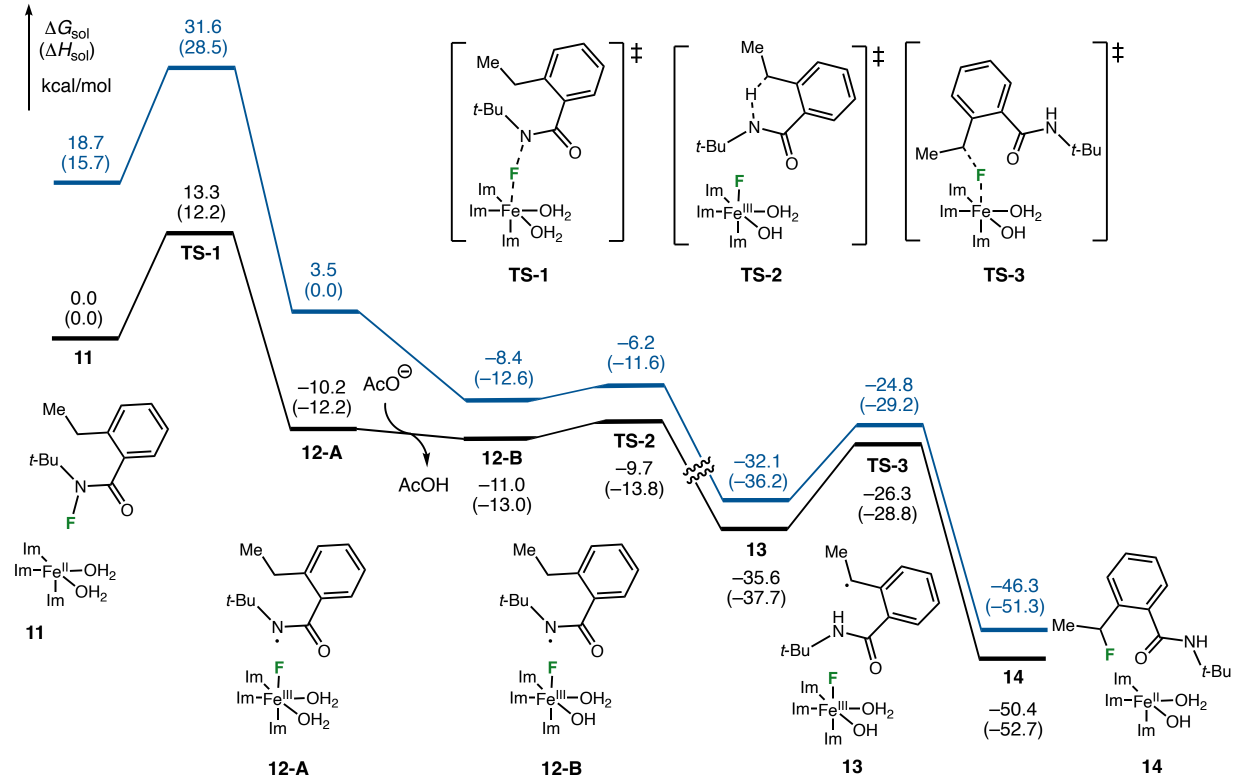

**Figure S19**. Reaction energy profile of the C($sp^3$)–H fluorination at quintet (black) and triplet (dark blue) spin states computed using the truncated model at the B3LYP-D3(BJ)/def2-TZVP/SMD($Et_2O$)//B3LYP-D3(BJ)/6-31G(d)–SDD(Fe) level of theory.

#### 9. Correlation Between Activation Barrier and Other Parameters for Radical Rebound

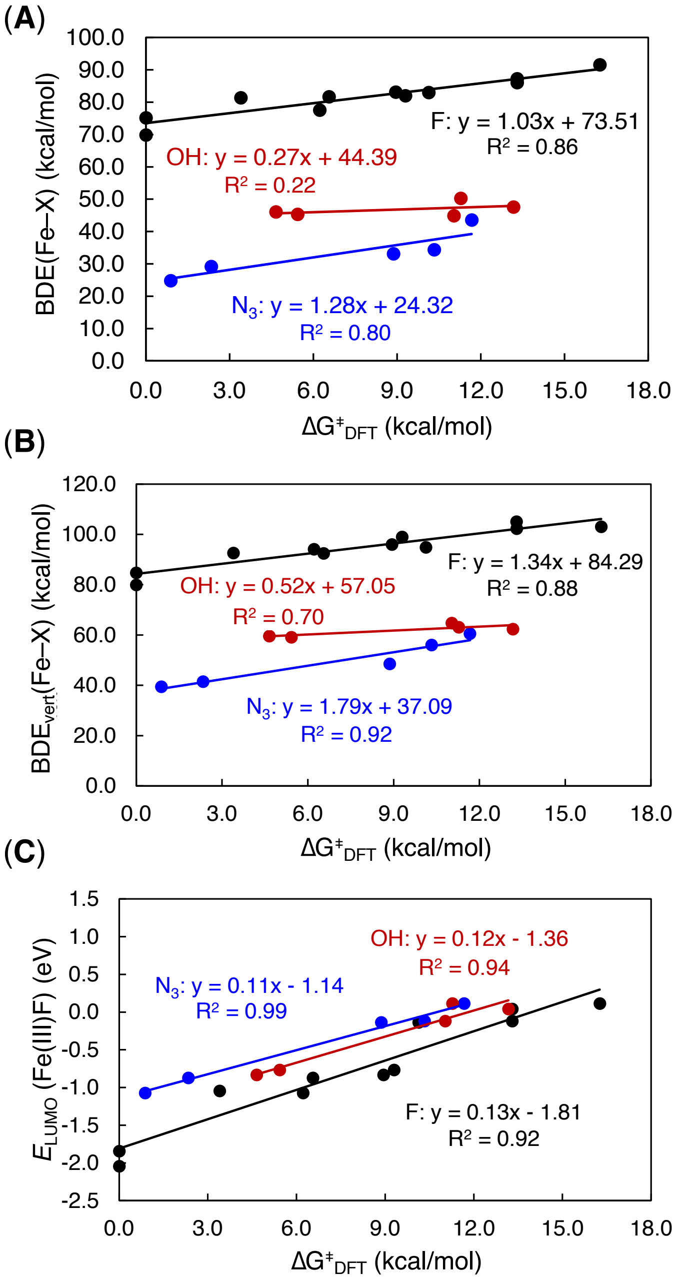

**Figure S20**. Correlation between activation barriers ($\Delta G^{\ddagger}$) for the radical rebound step with (**A**) bond dissociation enthalpies (BDE(Fe–X), X = F, $N_3$ or OH) (**B**) vertical bond dissociation energies ($BDE_{vert}$(Fe–X)), and (**C**) LUMO energies of $Fe^{III}F$ species ($E_{LUMO}(Fe^{III}F)$). The black, blue, and red colors represent the fluorine, azide, and hydroxyl radical rebound reactions, respectively.

#### 10. Alternative Regression Model for Fluorine Atom Abstraction

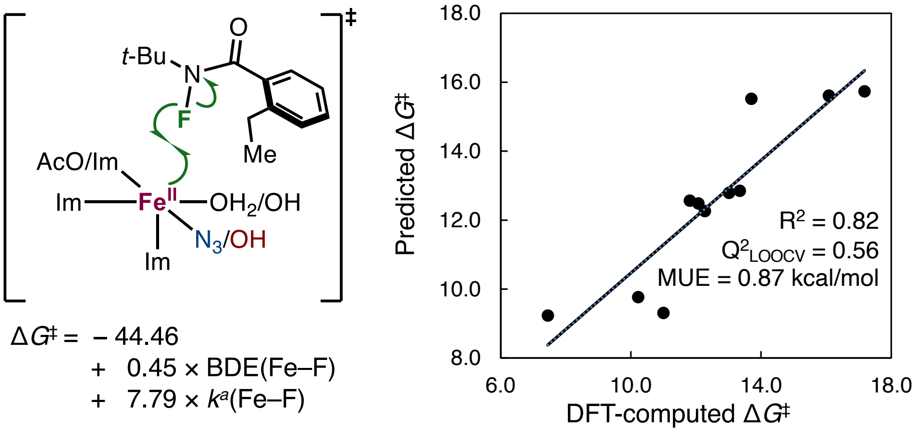

**Figure S21**. Alternative regression model for the fluorine atom abstraction step. BDE(Fe–F) is the bond dissociation enthalpy of Fe–F bond.

#### 11. Alternative Regression Model for Radical Rebound

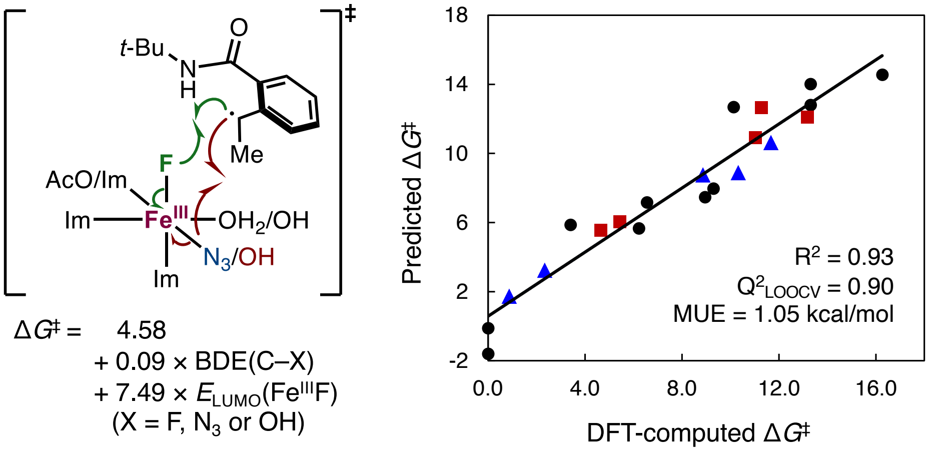

**Figure S22**.. Predictive models for the radical rebound step using the bond dissociation enthalpy of C–X bond, BDE(C–X), and LUMO energy of $Fe^{III}F$ species, $E_{LUMO}(Fe^{III}F)$, as descriptors.

#### 12. Substrate Reduction by Fe(II) Species

The oxidation potentials of *N*-fluoroamide substrate **1** and Fe(II) intermediates were calculated from the reaction Gibbs free energies of the half-reactions at the B3LYP-D3(BJ)/def2-

TZVP/SMD($Et_2O$)//B3LYP-D3(BJ)/6-31G(d)–SDD(Fe) level of theory. The Gibbs free energy of electron of −0.867 kcal/mol.[42]

Since

$$\Delta G_{rxn} = -n \times F \times E_{red}^{abs}$$

where *n* is the number of electron transferred (one), *F* is the Faraday constant. The absolute standard reduction potential can be calculated as

$$E_{red}^{abs} = \frac{\Delta G_{rxn}}{-n \times F}$$

Based on our DFT calculation, the electron transfers from Fe(II) to *N*-fluoroamide substrate **1** are highly endergonic (101.8 and 71.5 kcal/mol for $(Im)_3Fe^{II}(H_2O)_2$ (**11**) and $(Im)_2(AcO)Fe^{II}(H_2O)_2$ (**11'**), respectively). The highly unfavorable thermodynamics thus rule out the single electron transfer (SET) pathway.

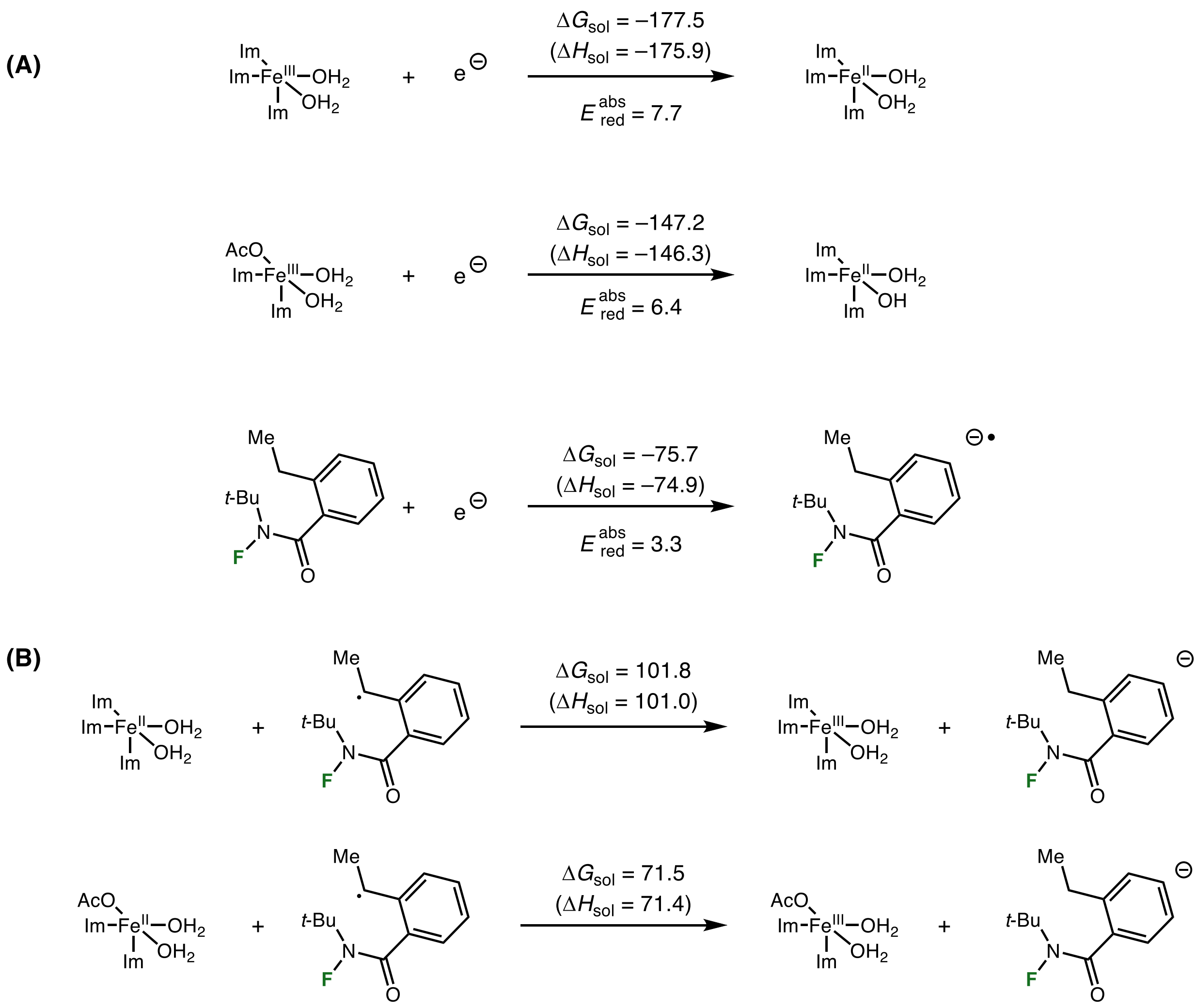

**Figure S23**. (**A**) Absolute standard reduction potential for half-reaction. (**B**) Electron transfer processes from Fe(II) to *N*-fluoroamide substrate **1**. Fe(II) and Fe(III)–F intermediates were calculated at high-spin quintet and sextet states, respectively.

#### 13. Benzyl Radical Oxidation Pathway by Fe(III)–F Species

We further evaluated the oxidation potentials of the benzyl radical and Fe(III)–F intermediates at the B3LYP-D3(BJ)/def2-TZVP/SMD($Et_2O$)//B3LYP-D3(BJ)/6-31G(d)–SDD(Fe) level of theory. Electron transfer from the benzyl radical to the Fe(III)–F intermediate is highly unfavorable, being uphill by 23.1 and 33.9 kcal/mol for $(Im)_3Fe^{III}F(H_2O)_2$ (**13**) and

$(Im)_2(AcO)Fe^{III}F(H_2O)_2$ (**13'**), respectively. Therefore, the formation of a carbocation intermediate via a single electron transfer mechanism can be ruled out.

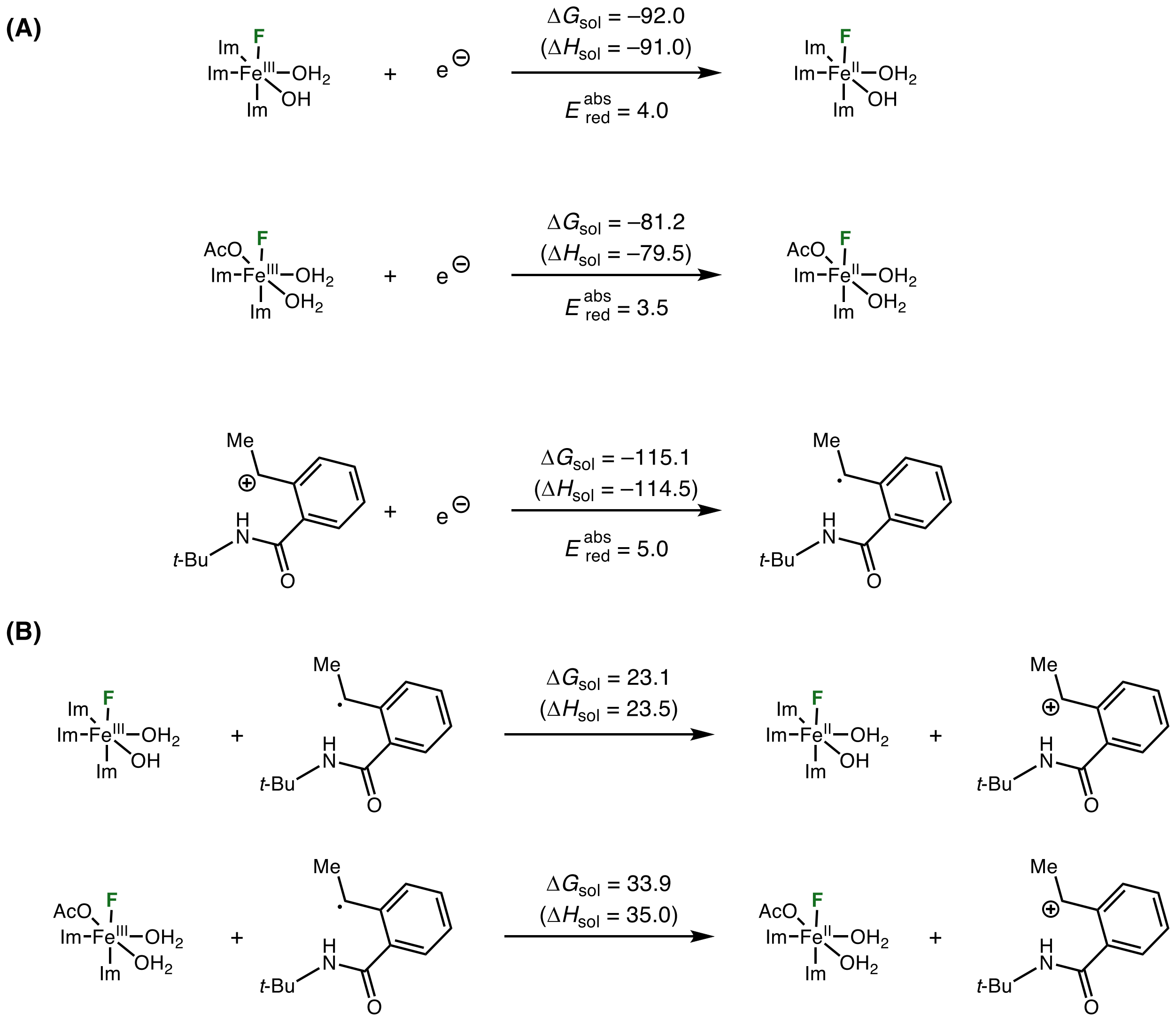

**Figure S24**. (**A**) Absolute standard reduction potential for half-reaction. (**B**) The formation of a carbocation intermediate via a single electron transfer mechanism. Fe(II) and Fe(III)–F intermediates were calculated at high-spin quintet and sextet states, respectively.

#### 14. Binding Free Energies of Azide Anion to Fe(II) and Fe(III)–F Species

The free energies associated with azide binding to Fe(II) and Fe(III)–F species were calculated at the B3LYP-D3(BJ)/def2-TZVP/SMD($Et_2O$)//B3LYP-D3(BJ)/6-31G(d)–SDD(Fe) level of theory. The calculated binding processes are uniformly exergonic ($\Delta G = -13.2$ to $-62.6$ kcal/mol), indicating a strong thermodynamic preference for azide coordination to the iron center.

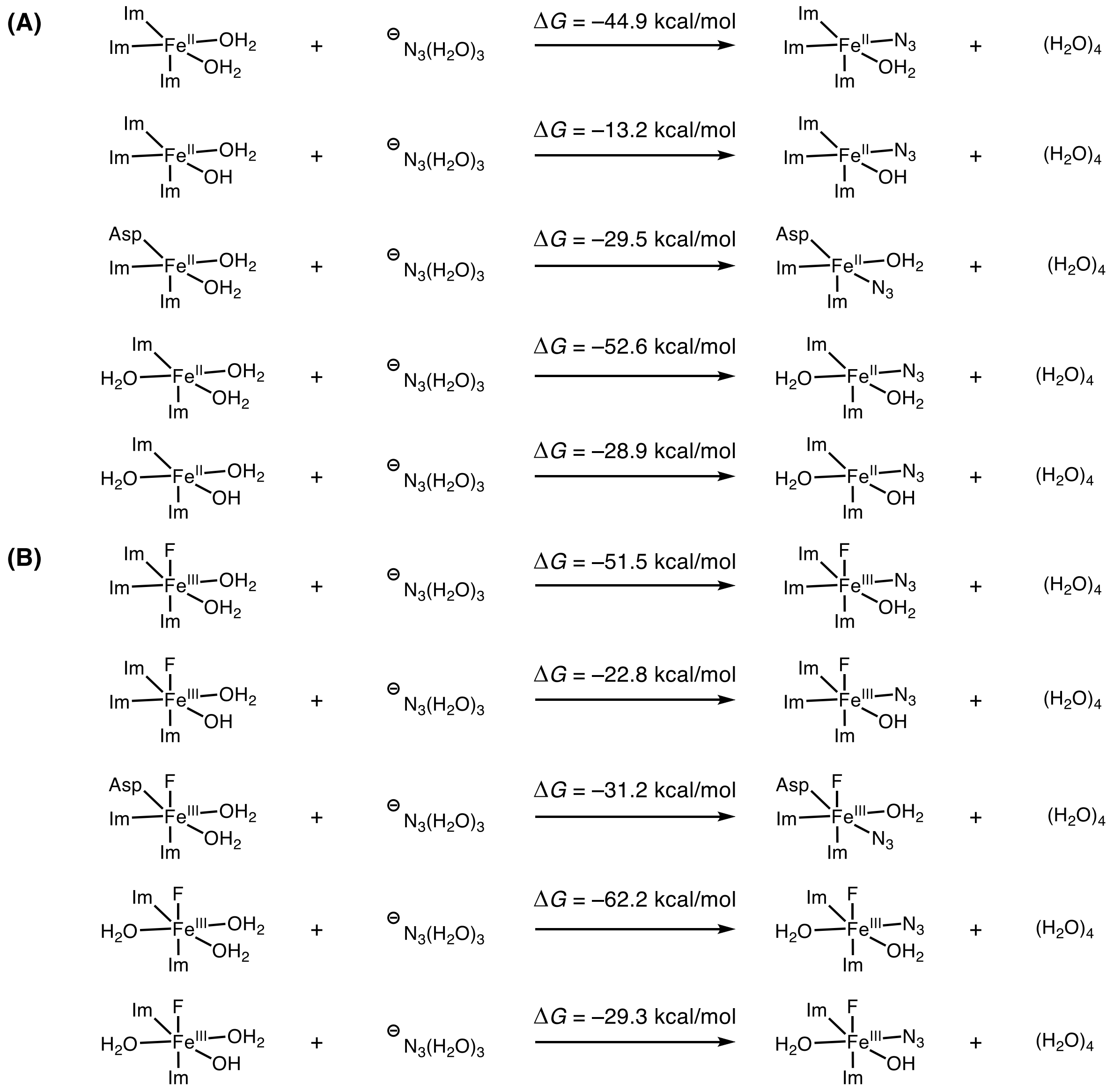

**Figure S25**. Calculated free energies for azide binding to Fe(II) and Fe(III)–F intermediates. Fe(II) and Fe(III)–F intermediates were calculated at high-spin quintet and sextet states, respectively.

#### 15. Radical Rebound Transition States from QM/MM Calculations

Transition states of the fluorine and hydroxy rebound pathways in $EgtB_{CHF1}$ were investigated using QM/MM ONIOM calculations. The fluorine rebound transition states leading to (*R*)-**2** and (*S*)-**2** are nearly identical in energy, in agreement with the modest enantioselectivity observed experimentally. Interestingly, because of the steric repulsion between phenyl group on the benzyl radical and H134, the hydroxy rebound **TS-3-OH-S$_{QM/MM}$** is calculated to be 1.0 kcal/mol higher in energy than the fluorine radical rebound **TS-3$_{QM/MM}$**. Given the relatively small activation free energy difference, these calculations suggest that fluorine and hydroxy radical rebound are energetically competitive.

It is worth noting that the truncated model and the QM/MM calculations predict different trends in the relative fluorine and hydroxy rebound barriers. In the truncated model, hydroxy rebound from the $Fe^{III}(F)(OH)$ intermediates are calculated to be more favorable than the corresponding fluorine radical rebounds (**Table 6**). In contrast, the QM/MM calculations predict fluorine rebound to be slightly lower in energy. This difference likely reflects the explicit protein environment included in the QM/MM calculations, which captures steric and noncovalent interactions absent from the truncated model. Nevertheless, both computational approaches consistently indicate that fluorine and hydroxy rebound pathways have similar activation barriers.

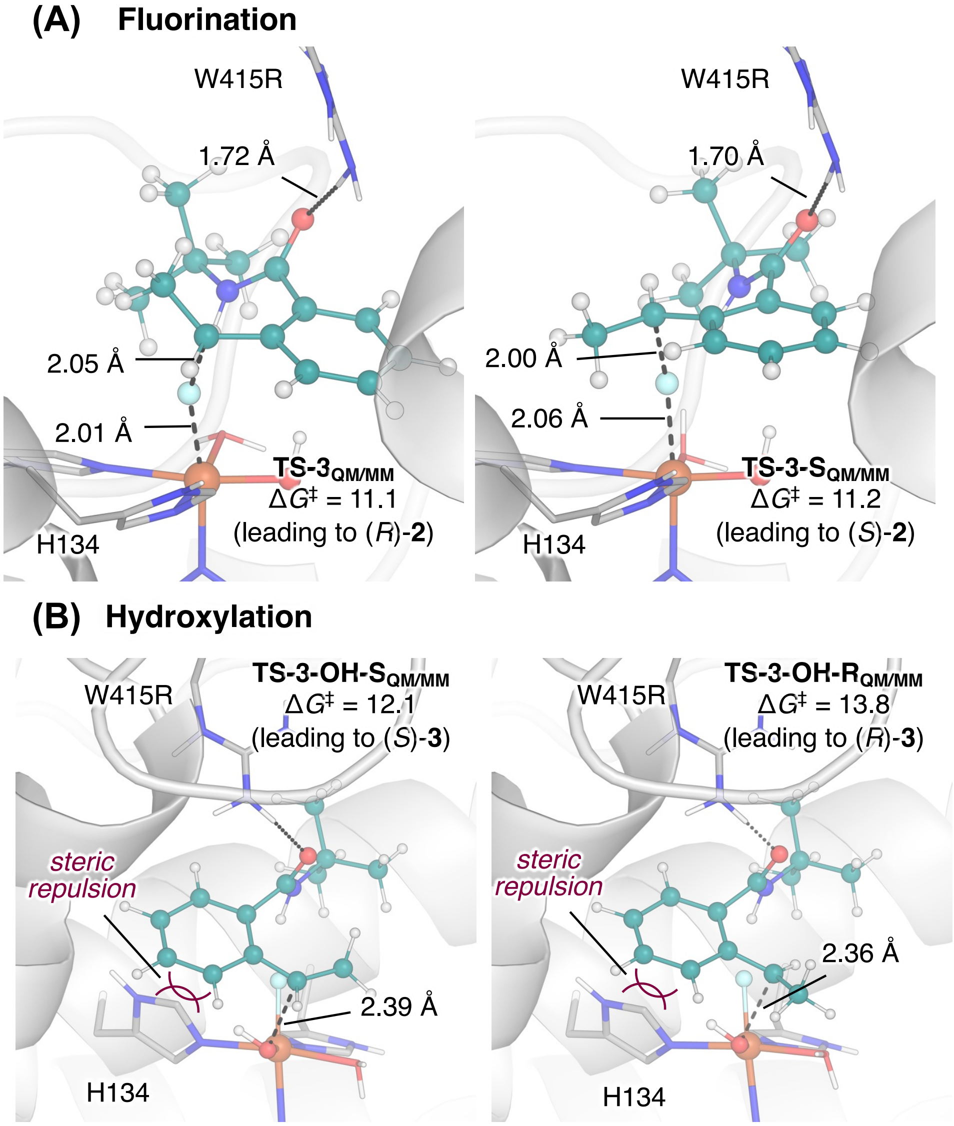

**Figure S26**. QM/MM-optimized geometries of the radical rebound transition states for the C–H fluorination and hydroxylation at high-spin quintet state. Gibbs free energy values are in kcal/mol with respect to the Fe(III)–F intermediates.

Furthermore, we have also considered the hydroxylation pathway involved in the equatorial-to-axial isomerization (**Figure S27**). However, the isomerization transition state **TS-4** is calculated to be 3.0 kcal/mol higher in energy than **TS-3-OH-S**$_{\mathbf{QM/MM}}$, which disfavors this scenario.

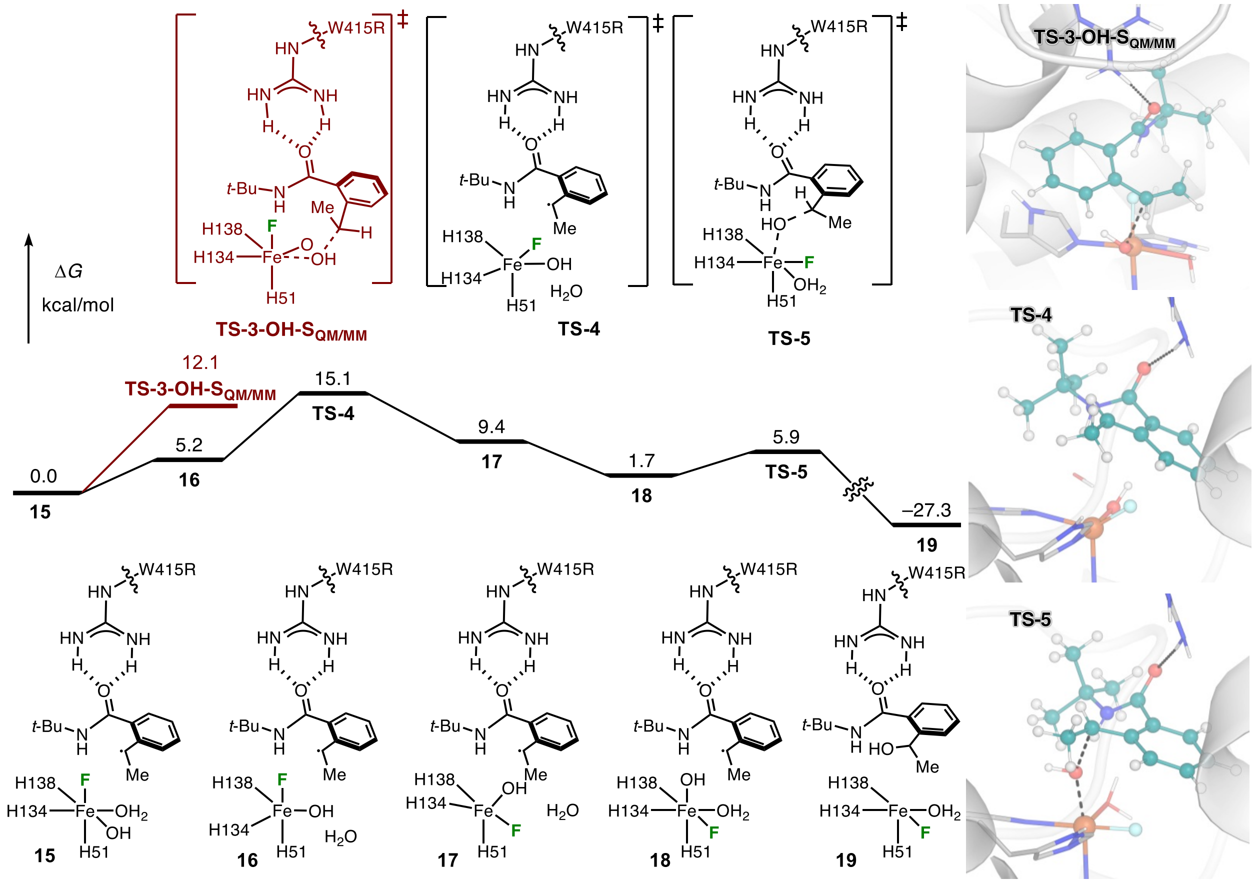

**Figure S27**. QM/MM-calculated free energy profile for the hydroxyl rebound pathway involving equatorial-to-axial isomerization of the hydroxyl ligand at high-spin quintet state. Gibbs free energy values are in kcal/mol with respect to the Fe(III)–F intermediates **15**.

#### 16. Classical MD Simulations for EgtB$_{CHF1}$ H138D Variant

To gain qualitative insight into the origin of the enantioselectivity in EgtB$_{CHF1}$ and EgtB$_{CHF1}$ H138D variants, we analyzed the N···$H^R$/$H^S$ distances in the near-attack-conformation (NACs). Interestingly, in EgtB$_{CHF1}$ enzyme, the $H^R$ atom is closer to nitrogen atom in *N*-fluoroamide **1**, favoring the activation of the C–$H^R$ bond. On the other hand, in EgtB$_{CHF1}$ H138D, the H138D mutation repositions the substrate toward the 375–385 loop and induces rotation of the ethyl group to relieve steric interactions. As a result, the $H^S$ atom becomes preferentially oriented toward the nitrogen atom nitrogen atom in *N*-fluoroamide **1**, favoring the activation of C–$H^S$ bond. These MD simulations provide a qualitative structural rationale for the altered enantioselectivity observed in the H138D variant.

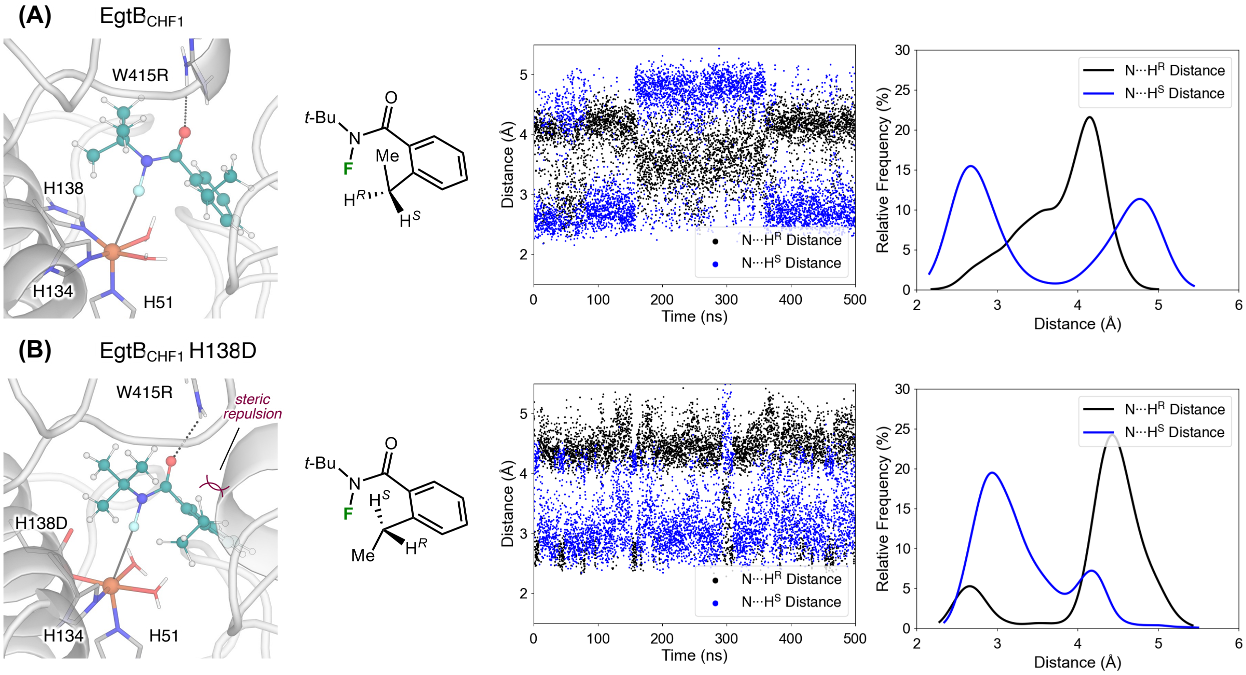

**Figure S28**. The most populated structures of $EgtB_{CHF1}$ and $EgtB_{CHF1}$ H138D; Time evolution and distribution of N···$H^R$/$H^S$ distances in *N*-fluoroamide **1** obtained from classical MD simulation.

(10) Frisch, M. J.; Trucks, G. W.; Schlegel, H. B.; Scuseria, G. E.; Robb, M. A.; Cheeseman, J. R.; Scalmani, G.; Barone, V.; Petersson, G. A.; Nakatsuji, H.; Li, X.; Caricato, M.; Marenich, A. V.; Bloino, J.; Janesko, B. G.; Gomperts, R.; Mennucci, B.; Hratchian, H. P.; Ortiz, J. V.; Izmaylov, A. F.; Sonnenberg, J. L.; Williams; Ding, F.; Lipparini, F.; Egidi, F.; Goings, J.; Peng, B.; Petrone, A.; Henderson, T.; Ranasinghe, D.; Zakrzewski, V. G.; Gao, J.; Rega, N.; Zheng, G.; Liang, W.; Hada, M.; Ehara, M.; Toyota, K.; Fukuda, R.; Hasegawa, J.; Ishida, M.; Nakajima, T.; Honda, Y.; Kitao, O.; Nakai, H.; Vreven, T.; Throssell, K.; Montgomery Jr., J. A.; Peralta, J. E.; Ogliaro, F.; Bearpark, M. J.; Heyd, J. J.; Brothers, E. N.; Kudin, K. N.; Staroverov, V. N.; Keith, T. A.; Kobayashi, R.; Normand, J.; Raghavachari, K.; Rendell, A. P.; Burant, J. C.; Iyengar, S. S.; Tomasi, J.; Cossi, M.; Millam, J. M.; Klene, M.; Adamo, C.; Cammi, R.; Ochterski, J. W.; Martin,

R. L.; Morokuma, K.; Farkas, O.; Foresman, J. B.; Fox, D. J. Gaussian 16 Rev. C.01, Wallingford, CT, **2016**.

(11) Morris, G. M.; Huey, R.; Lindstrom, W.; Sanner, M. F.; Belew, R. K.; Goodsell, D. S.; Olson, A. J. AutoDock4 and AutoDockTools4: Automated docking with selective receptor flexibility. *J. Comput. Chem.* **2009**, *30* (16), 2785-2791.

(12) Salomon-Ferrer, R.; Gotz, A. W.; Poole, D.; Le Grand, S.; Walker, R. C. Routine microsecond molecular dynamics simulations with AMBER on GPUs. 2. Explicit solvent particle mesh Ewald. *J. Chem. Theory Comput.* **2013**, *9* (9), 3878-3888.

(13) Case, D. A.; Aktulga, H. M.; Belfon, K.; Ben-Shalom, I. Y.; Berryman, J. T.; Brozell, S. R.; Cerutti, D. S.; T.E. Cheatham, I.; Cisneros, G. A.; Cruzeiro, V. W. D.; Darden, T. A.; Duke, R. E.; Giambasu, G.; Gilson, M. K.; Gohlke, H.; Goetz, A. W.; Harris, R.; Izadi, S.; Izmailov, S. A.; Kasavajhala, K.; Kaymak, M. C.; King, E.; Kovalenko, A.; Kurtzman, T.; Lee, T. S.; LeGrand, S.; Li, P.; Lin, C.; Liu, J.; Luchko, T.; Luo, R.; Machado, M.; Man, V.; Manathunga, M.; Merz, K. M.; Miao, Y.; Mikhailovskii, O.; Monard, G.; Nguyen, H.; O'Hearn, K. A.; Onufriev, A.; F. Pan, S. P.; Qi, R.; Rahnamoun, A.; Roe, D. R.; Roitberg, A.; Sagui, C.; Schott-Verdugo, S.; Shajan, A.; Shen, J.; Simmerling, C. L.; Skrynnikov, N. R.; Smith, J.; Swails, J.; Walker, R. C.; Wang, J.; Wang, J.; Wei, H.; Wolf, R. M.; Wu, X.; Xiong, Y.; Xue, Y.; York, D. M.; Zhao, S.; Kollman, P. A. Amber 20, University of California, San Francisco, CA, **2016**.

(14) Maier, J. A.; Martinez, C.; Kasavajhala, K.; Wickstrom, L.; Hauser, K. E.; Simmerling, C. ff14SB: improving the accuracy of protein side chain and backbone parameters from ff99SB. *J. Chem. Theory Comput.* **2015**, *11* (8), 3696-3713.

(15) Li, P.; Merz Jr, K. M. MCPB. py: A python based metal center parameter builder. *J. Chem. Inf. Model.* **2016**, *56*, 599-604.

(16) Wang, J.; Wolf, R. M.; Caldwell, J. W.; Kollman, P. A.; Case, D. A. Development and testing of a general amber force field. *J. Comput. Chem.* **2004**, *25* (9), 1157-1174.

(17) (a) Singh, U. C.; Kollman, P. A. An approach to computing electrostatic charges for molecules. *J. Comp. Chem.* **1984**, *5* (2), 129-145.; (b) Besler, B. H.; Merz Jr, K. M.; Kollman, P.

A. Atomic charges derived from semiempirical methods. *J. Comp. Chem.* **1990**, *11* (4), 431-439.; (c) Becke, A. D. Density-functional thermochemistry. III. The role of exact exchange. *J. Chem. Phys.* **1993**, *98* (7), 5648-5652.

#### XIII. Cartesian Coordinates and Energies

**1**

Charge: 0
Multiplicity: 1
B3LYP-D3(BJ)/6-31G(d)-SDD SCF energy (au): -736.044094994
B3LYP-D3(BJ)/6-31G(d)-SDD enthalpy (au): -735.737527994
B3LYP-D3(BJ)/6-31G(d)-SDD free energy (au): -735.798616994
B3LYP-D3(BJ)/def2-TZVP/SMD SCF energy (au): -736.354925435
B3LYP-D3(BJ)/def2-TZVP/SMD enthalpy (au): -736.048358435
B3LYP-D3(BJ)/def2-TZVP/SMD free energy (au): -736.109447435
B3LYP-D3(BJ)/def2-TZVP/SMD free energy (quasi-harmonic) (au): -736.107411441

Cartesian coordinates

| ATOM | X | Y | Z |
|---|---|---|---|
| C | -1.0265550000 | -0.3727870000 | -0.1814150000 |
| C | -1.2114450000 | -1.7075490000 | -0.5763740000 |
| C | -2.4573880000 | -2.3142090000 | -0.4723780000 |
| C | -3.5295060000 | -1.5916820000 | 0.0548480000 |
| C | -3.3442360000 | -0.2728520000 | 0.4609660000 |
| C | -2.1045140000 | 0.3697300000 | 0.3433570000 |
| H | -0.3686280000 | -2.2612130000 | -0.9745870000 |
| H | -2.5901710000 | -3.3428730000 | -0.7938510000 |
| H | -4.5067760000 | -2.0562220000 | 0.1529220000 |
| H | -4.1791180000 | 0.2831760000 | 0.8805380000 |
| C | -1.9995290000 | 1.8170780000 | 0.7741850000 |
| H | -0.9805570000 | 2.0554870000 | 1.0885850000 |
| H | -2.6493150000 | 1.9640330000 | 1.6460830000 |
| C | -2.4187110000 | 2.7867910000 | -0.3431180000 |
| H | -2.3715360000 | 3.8235390000 | 0.0089900000 |
| H | -3.4442480000 | 2.5845510000 | -0.6728330000 |
| H | -1.7487040000 | 2.6793070000 | -1.1991670000 |
| C | 0.2999870000 | 0.2778010000 | -0.3985520000 |
| O | 0.4288820000 | 1.4113690000 | -0.8444520000 |
| N | 1.4283510000 | -0.5229330000 | -0.1756700000 |
| C | 2.7973820000 | 0.0295320000 | 0.0265410000 |
| C | 3.7286180000 | -1.1618490000 | 0.3000080000 |
| C | 3.2321710000 | 0.7043560000 | -1.2851520000 |
| C | 2.8086710000 | 1.0135120000 | 1.2055670000 |
| H | 3.4960550000 | -1.6490090000 | 1.2485120000 |
| H | 3.6524860000 | -1.9037220000 | -0.5013390000 |
| H | 4.7607210000 | -0.8001930000 | 0.3398810000 |
| H | 2.6327340000 | 1.5870920000 | -1.5006850000 |
| H | 4.2838110000 | 0.9961430000 | -1.1923350000 |
| H | 3.1435440000 | 0.0036870000 | -2.1218300000 |
| H | 3.8292560000 | 1.3618530000 | 1.3957390000 |
| H | 2.1805530000 | 1.8814520000 | 0.9901880000 |
| H | 2.4392490000 | 0.5211370000 | 2.1109130000 |
| F | 1.1594720000 | -1.4509850000 | 0.8735580000 |

**1_Arg**

Charge: 1
Multiplicity: 1

B3LYP-D3(BJ)/6-31G(d)-SDD SCF energy (au): -981.175059921
B3LYP-D3(BJ)/6-31G(d)-SDD enthalpy (au): -980.740712921
B3LYP-D3(BJ)/6-31G(d)-SDD free energy (au): -980.822855921
B3LYP-D3(BJ)/def2-TZVP/SMD SCF energy (au): -981.653755640
B3LYP-D3(BJ)/def2-TZVP/SMD enthalpy (au): -981.219408640
B3LYP-D3(BJ)/def2-TZVP/SMD free energy (au): -981.301551640
B3LYP-D3(BJ)/def2-TZVP/SMD free energy (quasi-harmonic) (au): -981.295012719

Cartesian coordinates

| ATOM | X | Y | Z |
|---|---|---|---|
| H | 33.5365590000 | 40.6350500000 | 30.5518230000 |
| C | 34.5394700000 | 40.2054550000 | 30.4413190000 |
| N | 34.4948450000 | 38.7591470000 | 30.2435280000 |
| C | 34.1806730000 | 37.8774770000 | 31.2034570000 |
| N | 33.8266860000 | 38.2956400000 | 32.4244650000 |
| N | 34.2187650000 | 36.5619120000 | 30.9617780000 |
| H | 35.0128240000 | 40.6555860000 | 29.5685320000 |
| H | 35.1454120000 | 40.4455690000 | 31.3198370000 |
| H | 34.5695340000 | 38.4047800000 | 29.3003440000 |
| H | 33.5223080000 | 39.2436480000 | 32.5847590000 |
| H | 33.6606940000 | 37.5881400000 | 33.1400200000 |
| H | 34.7887590000 | 36.1884600000 | 30.2159660000 |
| H | 34.0087440000 | 35.9450150000 | 31.7511550000 |
| N | 34.7058890000 | 34.4889160000 | 35.2554620000 |
| O | 33.7461420000 | 35.7418380000 | 33.6438900000 |
| C | 35.3452010000 | 34.1303340000 | 32.9150280000 |
| C | 34.6113240000 | 33.3413410000 | 32.0194690000 |
| C | 35.2415060000 | 32.7480780000 | 30.9270140000 |
| C | 36.6053240000 | 32.9635460000 | 30.7231560000 |
| C | 37.3277680000 | 33.7618020000 | 31.6110100000 |
| C | 36.7213920000 | 34.3578640000 | 32.7229320000 |
| C | 34.5637140000 | 34.8512700000 | 33.9691190000 |
| C | 34.2575870000 | 35.1841480000 | 36.5055070000 |
| C | 34.9292110000 | 36.5643020000 | 36.5618970000 |
| C | 32.7252360000 | 35.2961880000 | 36.4609130000 |
| C | 34.6767740000 | 34.3265980000 | 37.7059780000 |
| C | 37.5194760000 | 35.2753060000 | 33.6232480000 |
| C | 37.2791280000 | 36.7628900000 | 33.3124540000 |
| F | 35.7103220000 | 33.5458850000 | 35.4589050000 |
| H | 33.5494840000 | 33.1873110000 | 32.1897910000 |
| H | 34.6742630000 | 32.1193680000 | 30.2480610000 |
| H | 37.1091140000 | 32.5054350000 | 29.8773950000 |
| H | 38.3890060000 | 33.9257620000 | 31.4442990000 |
| H | 37.2829610000 | 35.0721660000 | 34.6717610000 |
| H | 38.5841910000 | 35.0511730000 | 33.5007570000 |
| H | 37.8861030000 | 37.3981970000 | 33.9654320000 |
| H | 37.5464870000 | 36.9901040000 | 32.2741850000 |
| H | 36.2292460000 | 37.0371580000 | 33.4571740000 |
| H | 34.6231360000 | 37.0841300000 | 37.4747270000 |
| H | 36.0194950000 | 36.4669220000 | 36.5735830000 |
| H | 34.6384810000 | 37.1783570000 | 35.7052310000 |
| H | 32.3888870000 | 35.8888410000 | 35.6101380000 |
| H | 32.2698530000 | 34.3032720000 | 36.4009920000 |

| | | | |
|---|---|---|---|
| H | 32.3835520000 | 35.7755820000 | 37.3833500000 |
| H | 35.7622860000 | 34.2547140000 | 37.7982810000 |
| H | 34.2857400000 | 34.8015270000 | 38.6098510000 |
| H | 34.2608450000 | 33.3177890000 | 37.6407250000 |

**1_water**
Charge: 0
Multiplicity: 1
B3LYP-D3(BJ)/6-31G(d)-SDD SCF energy (au): -812.469508689
B3LYP-D3(BJ)/6-31G(d)-SDD enthalpy (au): -812.135146689
B3LYP-D3(BJ)/6-31G(d)-SDD free energy (au): -812.203476689
B3LYP-D3(BJ)/def2-TZVP/SMD SCF energy (au): -812.833279017
B3LYP-D3(BJ)/def2-TZVP/SMD enthalpy (au): -812.498917017
B3LYP-D3(BJ)/def2-TZVP/SMD free energy (au): -812.567247017
B3LYP-D3(BJ)/def2-TZVP/SMD free energy (quasi-harmonic) (au): -812.564434328

Cartesian coordinates

| ATOM | X | Y | Z |
|---|---|---|---|
| C | -5.5517070000 | -2.4789390000 | -4.4590300000 |
| C | -4.9199410000 | -1.2405870000 | -4.4499820000 |
| C | -3.6473320000 | -1.0835590000 | -5.0185640000 |
| C | -2.9770620000 | -2.1835020000 | -5.5896320000 |
| C | -3.6254050000 | -3.4253590000 | -5.5643500000 |
| C | -4.8969090000 | -3.5783880000 | -5.0173250000 |
| H | -6.5425680000 | -2.5872130000 | -4.0283620000 |
| H | -5.4123290000 | -0.3809000000 | -4.0082440000 |
| H | -3.1131350000 | -4.2873950000 | -5.9846890000 |
| H | -5.3736770000 | -4.5545830000 | -5.0207430000 |
| C | -1.6058220000 | -2.0915890000 | -6.2253200000 |
| C | -1.6824450000 | -1.9033030000 | -7.7498500000 |
| H | -1.0610460000 | -3.0174380000 | -6.0029530000 |
| H | -1.0215920000 | -1.2720280000 | -5.7999720000 |
| H | -0.6763350000 | -1.8754350000 | -8.1862380000 |
| H | -2.2292660000 | -2.7265360000 | -8.2229140000 |
| H | -2.1972990000 | -0.9690040000 | -7.9879860000 |
| C | -3.0381600000 | 0.2786480000 | -5.0953740000 |
| O | -2.6050960000 | 0.7454920000 | -6.1503200000 |
| N | -3.0779960000 | 1.0504360000 | -3.9506300000 |
| C | -2.2103300000 | 2.2160610000 | -3.6100390000 |
| C | -2.2626660000 | 3.2383990000 | -4.7575000000 |
| C | -0.7766650000 | 1.7316700000 | -3.3471950000 |
| H | -1.8153650000 | 4.1697640000 | -4.3945060000 |
| H | -1.7102030000 | 2.9060850000 | -5.6337340000 |
| H | -3.2966740000 | 3.4459160000 | -5.0496610000 |
| H | -0.7722980000 | 0.9759220000 | -2.5558130000 |
| H | -0.3243370000 | 1.3114150000 | -4.2497240000 |
| H | -0.1588940000 | 2.5743030000 | -3.0189840000 |
| C | -2.8163950000 | 2.8573910000 | -2.3504470000 |
| H | -3.8536930000 | 3.1580260000 | -2.5304990000 |
| H | -2.7929510000 | 2.1753090000 | -1.4988030000 |
| H | -2.2365790000 | 3.7493070000 | -2.0950270000 |
| O | 0.1573460000 | 0.8986120000 | -6.5151620000 |
| H | -0.8178580000 | 0.9313660000 | -6.5188880000 |

```
H      0.3781170000    0.3485070000   -7.2810040000
F     -3.2162990000    0.2416240000   -2.7953080000
```

**2**

Charge: 0
Multiplicity: 1
B3LYP-D3(BJ)/6-31G(d)-SDD SCF energy (au):               -736.115155606
B3LYP-D3(BJ)/6-31G(d)-SDD enthalpy (au):                 -735.807546606
B3LYP-D3(BJ)/6-31G(d)-SDD free energy (au):              -735.868788606
B3LYP-D3(BJ)/def2-TZVP/SMD SCF energy (au):              -736.438269050
B3LYP-D3(BJ)/def2-TZVP/SMD enthalpy (au):                -736.130660050
B3LYP-D3(BJ)/def2-TZVP/SMD free energy (au):             -736.191902050
B3LYP-D3(BJ)/def2-TZVP/SMD free energy (quasi-harmonic) (au): -736.189814444

Cartesian coordinates

```
ATOM    X               Y               Z
C      3.0339090000   -0.0772010000   -0.9129220000
O      2.4149420000   -0.5693690000   -1.8504540000
N      3.9812500000   -0.7334360000   -0.1880280000
H      4.4345370000   -0.2100210000    0.5460560000
C      4.2484810000   -2.1859690000   -0.2439180000
C      5.3115810000   -2.4705610000    0.8243040000
C      2.9603390000   -2.9635460000    0.0732370000
C      4.7839220000   -2.5608520000   -1.6351110000
H      6.2280180000   -1.8995130000    0.6316740000
H      4.9431610000   -2.2155920000    1.8258820000
H      5.5701250000   -3.5337190000    0.8234570000
H      2.1962540000   -2.7504270000   -0.6769160000
H      3.1627240000   -4.0405150000    0.0778150000
H      2.5699090000   -2.6705550000    1.0521850000
H      4.9961020000   -3.6351160000   -1.6806360000
H      4.0463010000   -2.3115370000   -2.4000220000
H      5.7100540000   -2.0156780000   -1.8489950000
C      2.8634270000    1.3792800000   -0.5373490000
C      3.2503230000    2.3179010000   -1.5028660000
C      2.2981970000    1.8228520000    0.6700220000
C      3.1330350000    3.6840310000   -1.2655280000
H      3.6457520000    1.9555860000   -2.4469150000
C      2.1676180000    3.2006160000    0.8892720000
C      2.5932680000    4.1284480000   -0.0571310000
H      3.4494380000    4.3970880000   -2.0211080000
H      1.7101970000    3.5481510000    1.8132510000
H      2.4843630000    5.1912650000    0.1384930000
C      1.7948680000    0.8975070000    1.7616730000
C      2.7975910000    0.6638100000    2.8862340000
H      3.0670820000    1.6126080000    3.3605600000
H      2.3654850000    0.0005220000    3.6420450000
H      3.7103030000    0.1993330000    2.5026450000
F      1.4311920000   -0.3441570000    1.2326300000
H      0.8791850000    1.3313290000    2.1828270000
```

**3**

Charge: 0

Multiplicity: 1
B3LYP-D3(BJ)/6-31G(d)-SDD SCF energy (au): -712.103839381
B3LYP-D3(BJ)/6-31G(d)-SDD enthalpy (au): -711.782950381
B3LYP-D3(BJ)/6-31G(d)-SDD free energy (au): -711.843650381
B3LYP-D3(BJ)/def2-TZVP/SMD SCF energy (au): -712.415375784
B3LYP-D3(BJ)/def2-TZVP/SMD enthalpy (au): -712.094486784
B3LYP-D3(BJ)/def2-TZVP/SMD free energy (au): -712.155186784
B3LYP-D3(BJ)/def2-TZVP/SMD free energy (quasi-harmonic) (au): -712.153135403

Cartesian coordinates

| ATOM | X | Y | Z |
|---|---|---|---|
| C | -0.2702010000 | 0.3631770000 | -3.6333980000 |
| O | -0.7789540000 | 0.0901640000 | -2.5363520000 |
| N | -0.5874100000 | -0.2979630000 | -4.7754110000 |
| H | -0.2298230000 | 0.0917840000 | -5.6361090000 |
| C | -1.5790390000 | -1.3899070000 | -4.8968930000 |
| C | -1.5837340000 | -1.7989340000 | -6.3753850000 |
| C | -1.1472800000 | -2.5785700000 | -4.0239130000 |
| C | -2.9722500000 | -0.8865580000 | -4.4838510000 |
| H | -1.8860320000 | -0.9617720000 | -7.0167360000 |
| H | -0.5919300000 | -2.1428260000 | -6.6921320000 |
| H | -2.2928030000 | -2.6163240000 | -6.5359830000 |
| H | -1.1005360000 | -2.2838420000 | -2.9741910000 |
| H | -1.8656250000 | -3.3989130000 | -4.1300830000 |
| H | -0.1605490000 | -2.9413340000 | -4.3320290000 |
| H | -3.7144680000 | -1.6800410000 | -4.6251990000 |
| H | -2.9739860000 | -0.5887350000 | -3.4339160000 |
| H | -3.2662270000 | -0.0245600000 | -5.0923030000 |
| C | 0.7766110000 | 1.4304510000 | -3.8011530000 |
| C | 1.8971390000 | 1.1341240000 | -4.5945900000 |
| C | 0.6836620000 | 2.6854460000 | -3.1647990000 |
| C | 2.9273340000 | 2.0542570000 | -4.7668950000 |
| H | 1.9729990000 | 0.1521020000 | -5.0527520000 |
| C | 1.7232730000 | 3.6013270000 | -3.3660520000 |
| C | 2.8358670000 | 3.3004180000 | -4.1488010000 |
| H | 3.7907930000 | 1.7987230000 | -5.3738700000 |
| H | 1.6502490000 | 4.5795610000 | -2.8962840000 |
| H | 3.6239140000 | 4.0370090000 | -4.2777110000 |
| C | -0.4612050000 | 3.1231380000 | -2.2481640000 |
| C | -0.1365660000 | 2.7819820000 | -0.7856600000 |
| H | 0.8076440000 | 3.2400940000 | -0.4711570000 |
| H | -0.9405020000 | 3.1488640000 | -0.1389300000 |
| H | -0.0518430000 | 1.6973940000 | -0.6712550000 |
| H | -0.5340090000 | 4.2132260000 | -2.3425290000 |
| O | -1.7369820000 | 2.6435460000 | -2.6288990000 |
| H | -1.7535790000 | 1.7007920000 | -2.3800440000 |

**4**
Charge: 0
Multiplicity: 1
B3LYP-D3(BJ)/6-31G(d)-SDD SCF energy (au): -800.474281268
B3LYP-D3(BJ)/6-31G(d)-SDD enthalpy (au): -800.153471268
B3LYP-D3(BJ)/6-31G(d)-SDD free energy (au): -800.218264268

B3LYP-D3(BJ)/def2-TZVP/SMD SCF energy (au): -800.810997620
B3LYP-D3(BJ)/def2-TZVP/SMD enthalpy (au): -800.490187620
B3LYP-D3(BJ)/def2-TZVP/SMD free energy (au): -800.554980620
B3LYP-D3(BJ)/def2-TZVP/SMD free energy (quasi-harmonic) (au): -800.552547352

Cartesian coordinates

| ATOM | X | Y | Z |
|---|---|---|---|
| C | -0.1638520000 | 0.3334340000 | -3.6034810000 |
| O | -0.5963860000 | 0.0340990000 | -2.4903130000 |
| N | -0.5322340000 | -0.2922260000 | -4.7546340000 |
| H | -0.2288500000 | 0.1428600000 | -5.6148030000 |
| C | -1.5592650000 | -1.3506150000 | -4.8669030000 |
| C | -1.6301630000 | -1.7223570000 | -6.3534700000 |
| C | -1.1304980000 | -2.5724010000 | -4.0393920000 |
| C | -2.9207580000 | -0.8203560000 | -4.3879550000 |
| H | -1.9330060000 | -0.8612460000 | -6.9623130000 |
| H | -0.6605910000 | -2.0837900000 | -6.7160460000 |
| H | -2.3675250000 | -2.5158680000 | -6.5070190000 |
| H | -1.0392580000 | -2.3042860000 | -2.9854990000 |
| H | -1.8745030000 | -3.3704190000 | -4.1395380000 |
| H | -0.1652040000 | -2.9532370000 | -4.3909930000 |
| H | -3.6912790000 | -1.5884560000 | -4.5187020000 |
| H | -2.8766200000 | -0.5455490000 | -3.3331160000 |
| H | -3.2128470000 | 0.0628540000 | -4.9667840000 |
| C | 0.8969750000 | 1.3886750000 | -3.8038700000 |
| C | 2.0803900000 | 1.0162100000 | -4.4544900000 |
| C | 0.7515600000 | 2.7041750000 | -3.3257130000 |
| C | 3.1281120000 | 1.9212230000 | -4.6195480000 |
| H | 2.1816770000 | -0.0038650000 | -4.8136920000 |
| C | 1.8036260000 | 3.6049670000 | -3.5146830000 |
| C | 2.9879780000 | 3.2237010000 | -4.1451290000 |
| H | 4.0427010000 | 1.6102110000 | -5.1157930000 |
| H | 1.6870070000 | 4.6284660000 | -3.1663590000 |
| H | 3.7897870000 | 3.9450580000 | -4.2733870000 |
| C | -0.4909780000 | 3.2002550000 | -2.6042490000 |
| C | -0.4657110000 | 2.8546800000 | -1.1101660000 |
| H | 0.4371340000 | 3.2803080000 | -0.6614850000 |
| H | -1.3345810000 | 3.2795840000 | -0.5951580000 |
| H | -0.4517660000 | 1.7711880000 | -0.9780810000 |
| H | -0.5049700000 | 4.2925850000 | -2.7084870000 |
| N | -1.7025520000 | 2.7221530000 | -3.3300830000 |
| N | -2.6196560000 | 2.2580010000 | -2.6511550000 |
| N | -3.5390110000 | 1.8103100000 | -2.1420590000 |

**$^{5}$11’**

Charge: 1
Multiplicity: 5
B3LYP-D3(BJ)/6-31G(d)-SDD SCF energy (au): -1693.80364091
B3LYP-D3(BJ)/6-31G(d)-SDD enthalpy (au): -1693.22260791
B3LYP-D3(BJ)/6-31G(d)-SDD free energy (au): -1693.33763091
B3LYP-D3(BJ)/def2-TZVP/SMD SCF energy (au): -2834.36043901
B3LYP-D3(BJ)/def2-TZVP/SMD enthalpy (au): -2833.77940601
B3LYP-D3(BJ)/def2-TZVP/SMD free energy (au): -2833.89442901

B3LYP-D3(BJ)/def2-TZVP/SMD free energy (quasi-harmonic) (au): -2833.88358210

Cartesian coordinates

| ATOM | X | Y | Z |
|---|---|---|---|
| Fe | -0.0347160000 | -0.8276320000 | -1.1589560000 |
| C | -0.5051100000 | -1.3740230000 | 3.1461640000 |
| N | 0.7100660000 | -0.7207370000 | 3.0584800000 |
| H | 1.2905520000 | -0.4404400000 | 3.8366910000 |
| C | 0.9888260000 | -0.5129900000 | 1.7479060000 |
| H | 1.8663040000 | -0.0107320000 | 1.3705120000 |
| N | 0.0147880000 | -1.0023750000 | 0.9941390000 |
| C | -0.9262280000 | -1.5405800000 | 1.8534400000 |
| H | -1.8330310000 | -1.9856330000 | 1.4744750000 |
| H | -0.9449600000 | -1.6462560000 | 4.0926150000 |
| C | 1.4609840000 | -4.5345150000 | -2.7561190000 |
| N | 1.5247000000 | -4.7947280000 | -1.3986490000 |
| H | 1.8807620000 | -5.6395860000 | -0.9733070000 |
| C | 1.0335090000 | -3.7219820000 | -0.7365260000 |
| H | 0.9633850000 | -3.6476640000 | 0.3379800000 |
| N | 0.6563390000 | -2.7886000000 | -1.6000550000 |
| C | 0.9190180000 | -3.2833340000 | -2.8670470000 |
| H | 0.6986510000 | -2.6919210000 | -3.7418650000 |
| H | 1.8012820000 | -5.2448870000 | -3.4928100000 |
| O | -2.1297290000 | -1.3998930000 | -1.1387150000 |
| H | -2.5666910000 | -0.6033580000 | -1.5464830000 |
| H | -2.2797780000 | -2.1329050000 | -1.7581540000 |
| C | -1.2400670000 | 4.2256280000 | -5.2158520000 |
| C | -1.2399630000 | 3.3772770000 | -4.1110590000 |
| C | -2.4527640000 | 2.8756290000 | -3.6132430000 |
| C | -3.6843340000 | 3.1857920000 | -4.2296970000 |
| C | -3.6517850000 | 4.0609410000 | -5.3188120000 |
| C | -2.4512180000 | 4.5792150000 | -5.8091060000 |
| H | -0.3012280000 | 4.6023810000 | -5.6089080000 |
| H | -0.3037210000 | 3.0729000000 | -3.6524790000 |
| H | -4.5878930000 | 4.3320270000 | -5.7987700000 |
| H | -2.4648430000 | 5.2480770000 | -6.6644380000 |
| C | -5.0064090000 | 2.6344320000 | -3.7394210000 |
| C | -5.4312570000 | 3.1739030000 | -2.3644850000 |
| H | -4.9533750000 | 1.5422550000 | -3.6973400000 |
| H | -5.7773240000 | 2.8867480000 | -4.4747520000 |
| H | -6.4201810000 | 2.7926330000 | -2.0910300000 |
| H | -4.7337660000 | 2.8636550000 | -1.5784760000 |
| H | -5.4770360000 | 4.2683720000 | -2.3653460000 |
| C | -2.3841720000 | 1.9057830000 | -2.4925480000 |
| O | -2.8293950000 | 0.7479350000 | -2.5979820000 |
| N | -1.6414370000 | 2.2931700000 | -1.4106360000 |
| C | -1.5443370000 | 3.5215020000 | -0.5524860000 |
| C | -1.9671850000 | 4.7757460000 | -1.3270220000 |
| C | -0.0768960000 | 3.6399560000 | -0.1203960000 |
| H | -1.9042760000 | 5.6123620000 | -0.6247220000 |
| H | -1.3101430000 | 4.9873070000 | -2.1701080000 |
| H | -2.9948620000 | 4.7140550000 | -1.6891380000 |
| H | 0.2556970000 | 2.7388630000 | 0.4007800000 |

| | | | |
|---|---|---|---|
| H | 0.5627470000 | 3.7949980000 | -0.9934520000 |
| H | 0.0371300000 | 4.4950790000 | 0.5524120000 |
| F | -1.4290170000 | 1.1723660000 | -0.5553930000 |
| C | -2.4773250000 | 3.3338880000 | 0.6578640000 |
| H | -3.5143380000 | 3.2002420000 | 0.3349970000 |
| H | -2.1844660000 | 2.4767440000 | 1.2665360000 |
| H | -2.4285890000 | 4.2309720000 | 1.2816510000 |
| C | 1.8683210000 | 1.3393320000 | -1.9124600000 |
| O | 1.3042760000 | 1.5508690000 | -3.0059370000 |
| O | 1.5084350000 | 0.4290650000 | -1.0653980000 |
| C | 3.0685200000 | 2.1690860000 | -1.5062410000 |
| H | 2.9278310000 | 2.5694250000 | -0.4973850000 |
| H | 3.9561920000 | 1.5271270000 | -1.4811410000 |
| H | 3.2342310000 | 2.9823290000 | -2.2140690000 |
| O | -0.3290320000 | -0.4651380000 | -3.3028720000 |
| H | 0.2925140000 | 0.3308930000 | -3.3646100000 |
| H | -1.2222140000 | -0.1106290000 | -3.4687760000 |

**$^{3}$11'**

Charge: 1
Multiplicity: 3
B3LYP-D3(BJ)/6-31G(d)-SDD SCF energy (au): -1693.77431645
B3LYP-D3(BJ)/6-31G(d)-SDD enthalpy (au): -1693.19253245
B3LYP-D3(BJ)/6-31G(d)-SDD free energy (au): -1693.30447145
B3LYP-D3(BJ)/def2-TZVP/SMD SCF energy (au): -2834.33412287
B3LYP-D3(BJ)/def2-TZVP/SMD enthalpy (au): -2833.75233887
B3LYP-D3(BJ)/def2-TZVP/SMD free energy (au): -2833.86427787
B3LYP-D3(BJ)/def2-TZVP/SMD free energy (quasi-harmonic) (au): -2833.85420442

Cartesian coordinates

| ATOM | X | Y | Z |
|---|---|---|---|
| Fe | 0.0133060000 | -0.7957500000 | -1.0716660000 |
| C | -0.2725360000 | -1.6762200000 | 2.9992700000 |
| N | 0.9375620000 | -1.0111420000 | 2.9302900000 |
| H | 1.5530910000 | -0.8156440000 | 3.7073000000 |
| C | 1.1572120000 | -0.6493020000 | 1.6441870000 |
| H | 2.0124360000 | -0.1027620000 | 1.2808740000 |
| N | 0.1458570000 | -1.0546550000 | 0.8863420000 |
| C | -0.7565960000 | -1.6966480000 | 1.7192000000 |
| H | -1.6819760000 | -2.0912040000 | 1.3319550000 |
| H | -0.6706380000 | -2.0536430000 | 3.9277990000 |
| C | 1.2799490000 | -4.5403640000 | -2.7674370000 |
| N | 1.2632910000 | -4.8637840000 | -1.4227040000 |
| H | 1.5311990000 | -5.7525820000 | -1.0231840000 |
| C | 0.8305590000 | -3.7828130000 | -0.7318320000 |
| H | 0.7188290000 | -3.7533720000 | 0.3412600000 |
| N | 0.5669800000 | -2.7845860000 | -1.5644760000 |
| C | 0.8454100000 | -3.2450840000 | -2.8406340000 |
| H | 0.7110730000 | -2.6019380000 | -3.6955550000 |
| H | 1.5938290000 | -5.2432950000 | -3.5228280000 |
| O | -2.0609270000 | -1.3385170000 | -1.0117090000 |
| H | -2.5001770000 | -0.5479440000 | -1.4270660000 |
| H | -2.1791520000 | -2.0637910000 | -1.6474400000 |

| | | | |
|---|---|---|---|
| C | -1.2268210000 | 4.1857320000 | -5.2504930000 |
| C | -1.2236980000 | 3.3748300000 | -4.1176940000 |
| C | -2.4316120000 | 2.8628570000 | -3.6208680000 |
| C | -3.6600990000 | 3.1229720000 | -4.2639070000 |
| C | -3.6323390000 | 3.9616390000 | -5.3817190000 |
| C | -2.4369170000 | 4.4910750000 | -5.8723010000 |
| H | -0.2913110000 | 4.5719140000 | -5.6424050000 |
| H | -0.2880980000 | 3.1093300000 | -3.6341160000 |
| H | -4.5674010000 | 4.1943230000 | -5.8834470000 |
| H | -2.4530020000 | 5.1307960000 | -6.7495920000 |
| C | -4.9740600000 | 2.5537630000 | -3.7723030000 |
| C | -5.4353550000 | 3.1313270000 | -2.4246440000 |
| H | -4.8908230000 | 1.4656700000 | -3.6884840000 |
| H | -5.7405230000 | 2.7559200000 | -4.5275430000 |
| H | -6.4166870000 | 2.7309910000 | -2.1510110000 |
| H | -4.7411870000 | 2.8729470000 | -1.6171080000 |
| H | -5.5138030000 | 4.2230890000 | -2.4681210000 |
| C | -2.3593820000 | 1.9301870000 | -2.4681410000 |
| O | -2.7367130000 | 0.7442050000 | -2.5651170000 |
| N | -1.7124410000 | 2.3796300000 | -1.3587890000 |
| C | -1.6052240000 | 3.6554480000 | -0.5774140000 |
| C | -2.0535230000 | 4.8638180000 | -1.4085360000 |
| C | -0.1322570000 | 3.8063380000 | -0.1746710000 |
| H | -2.0068440000 | 5.7327860000 | -0.7453520000 |
| H | -1.4026190000 | 5.0507120000 | -2.2617780000 |
| H | -3.0804210000 | 4.7645640000 | -1.7645600000 |
| H | 0.2123760000 | 2.9370320000 | 0.3909440000 |
| H | 0.4941890000 | 3.9157250000 | -1.0640140000 |
| H | -0.0136390000 | 4.6977110000 | 0.4484240000 |
| F | -1.4878720000 | 1.3063650000 | -0.4595790000 |
| C | -2.5186390000 | 3.5264690000 | 0.6557240000 |
| H | -3.5594330000 | 3.3713640000 | 0.3551540000 |
| H | -2.2122020000 | 2.7035280000 | 1.3033770000 |
| H | -2.4648120000 | 4.4549020000 | 1.2311800000 |
| C | 1.8519490000 | 1.4131220000 | -1.7583990000 |
| O | 1.3058900000 | 1.6151410000 | -2.8667580000 |
| O | 1.4890840000 | 0.5081470000 | -0.9130410000 |
| C | 3.0336700000 | 2.2615600000 | -1.3373290000 |
| H | 2.8754850000 | 2.6591310000 | -0.3301680000 |
| H | 3.9318740000 | 1.6346760000 | -1.3049860000 |
| H | 3.1931710000 | 3.0777100000 | -2.0432180000 |
| O | -0.2128690000 | -0.4269190000 | -3.0460750000 |
| H | 0.3735510000 | 0.4057130000 | -3.1457600000 |
| H | -1.1272250000 | -0.1227810000 | -3.2153380000 |

**$^{5}$11**

Charge: 2
Multiplicity: 5
B3LYP-D3(BJ)/6-31G(d)-SDD SCF energy (au): -1691.25400454
B3LYP-D3(BJ)/6-31G(d)-SDD enthalpy (au): -1690.65150554
B3LYP-D3(BJ)/6-31G(d)-SDD free energy (au): -1690.76988654
B3LYP-D3(BJ)/def2-TZVP/SMD SCF energy (au): -2831.91639985
B3LYP-D3(BJ)/def2-TZVP/SMD enthalpy (au): -2831.31390085

B3LYP-D3(BJ)/def2-TZVP/SMD free energy (au): -2831.43228185
B3LYP-D3(BJ)/def2-TZVP/SMD free energy (quasi-harmonic) (au): -2831.42071131

Cartesian coordinates

| ATOM | X | Y | Z |
|---|---|---|---|
| Fe | 0.0398830000 | -1.1210000000 | -0.7156970000 |
| C | 1.3301560000 | 2.1531860000 | 1.8256930000 |
| N | 1.7837640000 | 2.5930540000 | 0.5971800000 |
| H | 2.2921800000 | 3.4511590000 | 0.4269640000 |
| C | 1.4448800000 | 1.6761920000 | -0.3383750000 |
| H | 1.6655910000 | 1.7749350000 | -1.3891240000 |
| N | 0.7975950000 | 0.6635190000 | 0.2286010000 |
| C | 0.7204330000 | 0.9522560000 | 1.5819740000 |
| H | 0.2269750000 | 0.2867180000 | 2.2727170000 |
| C | -3.0064880000 | -1.7330810000 | 2.2780750000 |
| N | -3.7572060000 | -1.1914580000 | 1.2529330000 |
| H | -4.7522480000 | -1.0090620000 | 1.2797450000 |
| C | -2.9436700000 | -0.9529320000 | 0.2006640000 |
| H | -3.2669540000 | -0.5142540000 | -0.7299270000 |
| N | -1.6995010000 | -1.3151480000 | 0.4990890000 |
| C | -1.7276920000 | -1.8055550000 | 1.7973490000 |
| H | -0.8376350000 | -2.1801770000 | 2.2783960000 |
| O | 1.6087730000 | -0.9950130000 | -2.2360680000 |
| H | 1.7222580000 | -1.8732760000 | -2.6396220000 |
| H | 1.2152380000 | -0.4273410000 | -2.9249350000 |
| H | -3.4400950000 | -2.0130240000 | 3.2254290000 |
| H | 1.4843760000 | 2.7169270000 | 2.7325060000 |
| C | 2.1797410000 | -4.2227730000 | 1.4548350000 |
| N | 3.1118680000 | -3.2059040000 | 1.3958760000 |
| H | 4.0533810000 | -3.2450870000 | 1.7656370000 |
| C | 2.5584390000 | -2.1520250000 | 0.7560940000 |
| H | 3.0669460000 | -1.2190960000 | 0.5675970000 |
| N | 1.3118040000 | -2.4391070000 | 0.3951440000 |
| C | 1.0663450000 | -3.7337070000 | 0.8274800000 |
| H | 0.1111220000 | -4.2127880000 | 0.6750630000 |
| H | 2.3894650000 | -5.1714330000 | 1.9238250000 |
| O | -0.5977120000 | -2.7852400000 | -2.0850330000 |
| H | -1.3205190000 | -2.4326420000 | -2.6319510000 |
| H | -0.9030610000 | -3.6413780000 | -1.7430170000 |
| C | -5.2150830000 | 1.5195840000 | -3.1873420000 |
| C | -3.8789190000 | 1.1835380000 | -3.3888330000 |
| C | -2.9170890000 | 1.5283170000 | -2.4291280000 |
| C | -3.2746250000 | 2.1692030000 | -1.2278010000 |
| C | -4.6249880000 | 2.5008590000 | -1.0552850000 |
| C | -5.5836290000 | 2.1939690000 | -2.0206450000 |
| H | -5.9590480000 | 1.2629940000 | -3.9342200000 |
| H | -3.5770190000 | 0.6688580000 | -4.2970200000 |
| H | -4.9257180000 | 3.0075040000 | -0.1420160000 |
| H | -6.6205000000 | 2.4712290000 | -1.8585460000 |
| C | -2.2723610000 | 2.4788280000 | -0.1349880000 |
| C | -1.5895850000 | 3.8501640000 | -0.2624700000 |
| H | -1.5060960000 | 1.6999050000 | -0.0973880000 |
| H | -2.7954880000 | 2.4365190000 | 0.8271510000 |

| | | | |
|---|---|---|---|
| H | -0.9686830000 | 4.0430850000 | 0.6180040000 |
| H | -0.9441810000 | 3.8910290000 | -1.1450710000 |
| H | -2.3254760000 | 4.6564960000 | -0.3405320000 |
| C | -1.5108250000 | 1.0860040000 | -2.6604220000 |
| O | -1.1214310000 | -0.0351490000 | -2.2671160000 |
| N | -0.6697130000 | 1.9526040000 | -3.2676940000 |
| C | -0.7819820000 | 3.0783390000 | -4.2790130000 |
| C | -2.0471010000 | 3.9131090000 | -4.0383630000 |
| C | -0.8020200000 | 2.4451610000 | -5.6765100000 |
| H | -1.9742380000 | 4.7813130000 | -4.6996400000 |
| H | -2.9623300000 | 3.3797710000 | -4.2889470000 |
| H | -2.1153910000 | 4.2781390000 | -3.0127390000 |
| H | 0.0926880000 | 1.8443460000 | -5.8572330000 |
| H | -1.6885360000 | 1.8163250000 | -5.8071400000 |
| H | -0.8359520000 | 3.2375010000 | -6.4295440000 |
| F | 0.6088850000 | 1.3523630000 | -3.4399620000 |
| C | 0.4537050000 | 3.9692240000 | -4.0753390000 |
| H | 0.4789960000 | 4.3750220000 | -3.0588430000 |
| H | 1.3841650000 | 3.4347240000 | -4.2733480000 |
| H | 0.3939050000 | 4.8090500000 | -4.7720020000 |

**$^{3}$11**

Charge: 2
Multiplicity: 3
B3LYP-D3(BJ)/6-31G(d)-SDD SCF energy (au): -1691.21920689
B3LYP-D3(BJ)/6-31G(d)-SDD enthalpy (au): -1690.61588889
B3LYP-D3(BJ)/6-31G(d)-SDD free energy (au): -1690.72999089
B3LYP-D3(BJ)/def2-TZVP/SMD SCF energy (au): -2831.89220848
B3LYP-D3(BJ)/def2-TZVP/SMD enthalpy (au): -2831.28889048
B3LYP-D3(BJ)/def2-TZVP/SMD free energy (au): -2831.40299248
B3LYP-D3(BJ)/def2-TZVP/SMD free energy (quasi-harmonic) (au): -2831.39095059

Cartesian coordinates

| ATOM | X | Y | Z |
|---|---|---|---|
| Fe | 0.6459010000 | -1.0644500000 | -0.1404990000 |
| C | -0.4712530000 | 2.2903930000 | 2.0963950000 |
| N | 0.3580980000 | 2.8640600000 | 1.1499070000 |
| H | 0.6158330000 | 3.8414830000 | 1.1036690000 |
| C | 0.7629010000 | 1.9101410000 | 0.2897300000 |
| H | 1.4134690000 | 2.0824430000 | -0.5503620000 |
| N | 0.2280470000 | 0.7380310000 | 0.6335610000 |
| C | -0.5493350000 | 0.9682470000 | 1.7636900000 |
| H | -1.0892890000 | 0.1727980000 | 2.2509030000 |
| C | -2.5924020000 | -2.8926500000 | 1.7013590000 |
| N | -3.2046820000 | -2.1534460000 | 0.7085850000 |
| H | -4.2004800000 | -2.1099090000 | 0.5325190000 |
| C | -2.2588260000 | -1.5122770000 | -0.0104910000 |
| H | -2.4538030000 | -0.8536690000 | -0.8473190000 |
| N | -1.0550280000 | -1.8118880000 | 0.4801540000 |
| C | -1.2508180000 | -2.6746680000 | 1.5505080000 |
| H | -0.4246130000 | -3.0714800000 | 2.1186180000 |
| O | 2.3652090000 | -0.2374630000 | -0.9524160000 |
| H | 3.1891370000 | -0.7420890000 | -0.8439870000 |

| | | | |
|---|---|---|---|
| H | 2.0951690000 | -0.3559800000 | -1.8897750000 |
| H | -3.1497710000 | -3.4944940000 | 2.4019500000 |
| H | -0.9126840000 | 2.8607950000 | 2.8982050000 |
| C | 2.0076560000 | -5.0506640000 | -0.0525780000 |
| N | 3.0108160000 | -4.4168900000 | 0.6495720000 |
| H | 3.8159670000 | -4.8666690000 | 1.0682790000 |
| C | 2.7344930000 | -3.0957510000 | 0.7001260000 |
| H | 3.3474520000 | -2.3722520000 | 1.2177350000 |
| N | 1.5925140000 | -2.8470460000 | 0.0593160000 |
| C | 1.1276900000 | -4.0663630000 | -0.4115400000 |
| H | 0.1991130000 | -4.1479140000 | -0.9542440000 |
| H | 2.0100340000 | -6.1150360000 | -0.2289750000 |
| O | 0.3166570000 | -1.0252580000 | -2.3267380000 |
| H | -0.4427380000 | -0.3903090000 | -2.5103950000 |
| H | 0.1092030000 | -1.8514230000 | -2.7921100000 |
| C | -5.5916060000 | 2.3978030000 | -3.6178110000 |
| C | -4.3071290000 | 1.8657800000 | -3.6061350000 |
| C | -3.4242470000 | 2.1898810000 | -2.5647430000 |
| C | -3.8334910000 | 2.9839390000 | -1.4784800000 |
| C | -5.1365750000 | 3.4973340000 | -1.5101390000 |
| C | -6.0001020000 | 3.2286070000 | -2.5700510000 |
| H | -6.2697130000 | 2.1662690000 | -4.4325100000 |
| H | -3.9720140000 | 1.2201910000 | -4.4126800000 |
| H | -5.4818140000 | 4.1069140000 | -0.6792540000 |
| H | -7.0008430000 | 3.6489430000 | -2.5695630000 |
| C | -2.9621210000 | 3.2314020000 | -0.2623180000 |
| C | -2.3716600000 | 4.6466610000 | -0.1768620000 |
| H | -2.1535570000 | 2.4946400000 | -0.2196100000 |
| H | -3.5782190000 | 3.0559160000 | 0.6280270000 |
| H | -1.9074860000 | 4.8131020000 | 0.8015930000 |
| H | -1.6120970000 | 4.8058470000 | -0.9479910000 |
| H | -3.1458980000 | 5.4094360000 | -0.3043330000 |
| C | -2.0583260000 | 1.6099530000 | -2.6215010000 |
| O | -1.8648800000 | 0.3949820000 | -2.4498830000 |
| N | -1.0223180000 | 2.5013980000 | -2.7779130000 |
| C | -0.8227000000 | 3.5407530000 | -3.8704470000 |
| C | -2.0405620000 | 4.4684610000 | -3.9732440000 |
| C | -0.5959980000 | 2.8021490000 | -5.1965030000 |
| H | -1.7700400000 | 5.2661060000 | -4.6713170000 |
| H | -2.9234310000 | 3.9663170000 | -4.3663150000 |
| H | -2.2932260000 | 4.9289330000 | -3.0173910000 |
| H | 0.2560580000 | 2.1210420000 | -5.1335620000 |
| H | -1.4873710000 | 2.2334470000 | -5.4822730000 |
| H | -0.3934050000 | 3.5289090000 | -5.9882590000 |
| F | 0.2038950000 | 1.7717150000 | -2.6895500000 |
| C | 0.4064240000 | 4.3640110000 | -3.4615160000 |
| H | 0.2459410000 | 4.8462410000 | -2.4917400000 |
| H | 1.3095110000 | 3.7529060000 | -3.4123280000 |
| H | 0.5696110000 | 5.1481700000 | -4.2051950000 |

**$^{5}$12'**

Charge: 1

Multiplicity: 5

B3LYP-D3(BJ)/6-31G(d)-SDD SCF energy (au): -1693.82554952
B3LYP-D3(BJ)/6-31G(d)-SDD enthalpy (au): -1693.24635252
B3LYP-D3(BJ)/6-31G(d)-SDD free energy (au): -1693.36048152
B3LYP-D3(BJ)/def2-TZVP/SMD SCF energy (au): -2834.38194382
B3LYP-D3(BJ)/def2-TZVP/SMD enthalpy (au): -2833.80274682
B3LYP-D3(BJ)/def2-TZVP/SMD free energy (au): -2833.91687582
B3LYP-D3(BJ)/def2-TZVP/SMD free energy (quasi-harmonic) (au): -2833.90619230

Cartesian coordinates

| ATOM | X | Y | Z |
|---|---|---|---|
| Fe | -0.1333820000 | 0.1252450000 | -0.4657430000 |
| C | -3.7528320000 | -2.0747990000 | 0.5303000000 |
| N | -4.1529380000 | -0.7502900000 | 0.5665800000 |
| H | -5.0812470000 | -0.4163510000 | 0.7865270000 |
| C | -3.0930510000 | 0.0297750000 | 0.2654700000 |
| H | -3.1056750000 | 1.1060500000 | 0.2044410000 |
| N | -2.0297110000 | -0.7340260000 | 0.0341700000 |
| C | -2.4272540000 | -2.0496200000 | 0.1957340000 |
| H | -1.7363880000 | -2.8651510000 | 0.0563370000 |
| H | -4.4299230000 | -2.8869150000 | 0.7426200000 |
| C | 1.3209460000 | -3.7875320000 | -1.6711370000 |
| N | 1.8173520000 | -3.6713620000 | -0.3845620000 |
| H | 2.4150610000 | -4.3386700000 | 0.0837920000 |
| C | 1.3942790000 | -2.4973950000 | 0.1332250000 |
| H | 1.6326640000 | -2.1314740000 | 1.1191190000 |
| N | 0.6445700000 | -1.8582360000 | -0.7565560000 |
| C | 0.5928700000 | -2.6508320000 | -1.8901910000 |
| H | 0.0557580000 | -2.3274250000 | -2.7665720000 |
| H | 1.5346020000 | -4.6408910000 | -2.2949780000 |
| O | -0.8268660000 | -0.0527370000 | -2.5243920000 |
| H | -0.5442620000 | 0.8085600000 | -2.9429110000 |
| H | -1.7975950000 | 0.0118580000 | -2.4975800000 |
| F | -0.9312190000 | 1.7718530000 | -0.3103940000 |
| C | -0.9328470000 | 3.2850440000 | -2.7666480000 |
| O | -0.0698420000 | 2.4477870000 | -3.1461210000 |
| N | -0.6127350000 | 4.5796430000 | -2.6114160000 |
| H | -3.3490090000 | 4.1818620000 | 0.2911210000 |
| C | 0.5661710000 | 5.1867290000 | -2.0172910000 |
| C | 0.2320700000 | 6.6649520000 | -1.7458660000 |
| C | 1.0447840000 | 4.4932600000 | -0.7358630000 |
| C | 1.6595030000 | 5.1149410000 | -3.1262670000 |
| H | -0.1363390000 | 7.1531550000 | -2.6521540000 |
| H | -0.5295690000 | 6.7550540000 | -0.9664570000 |
| H | 1.1335980000 | 7.1835410000 | -1.4066850000 |
| H | 1.2804960000 | 3.4444010000 | -0.9023890000 |
| H | 1.9489360000 | 4.9874090000 | -0.3679690000 |
| H | 0.2867980000 | 4.5439370000 | 0.0507420000 |
| H | 2.5433360000 | 5.6470760000 | -2.7593340000 |
| H | 1.9219240000 | 4.0790150000 | -3.3458860000 |
| H | 1.3102220000 | 5.5958470000 | -4.0433170000 |
| C | -2.3745640000 | 2.8982830000 | -2.6820980000 |
| C | -2.8280440000 | 1.9668400000 | -3.6317310000 |
| C | -3.2445550000 | 3.3654630000 | -1.6777200000 |

| | | | |
|---|---|---|---|
| C | -4.1263800000 | 1.4671520000 | -3.5832360000 |
| H | -2.1545810000 | 1.6556190000 | -4.4244160000 |
| C | -4.5441350000 | 2.8389760000 | -1.6410270000 |
| C | -4.9862450000 | 1.8997840000 | -2.5704820000 |
| H | -4.4680790000 | 0.7633500000 | -4.3358280000 |
| H | -5.2223800000 | 3.1890860000 | -0.8663500000 |
| H | -6.0034840000 | 1.5222900000 | -2.5197490000 |
| C | -2.8620830000 | 4.4166630000 | -0.6631610000 |
| C | -3.2871640000 | 5.8241490000 | -1.1162350000 |
| H | -4.3720670000 | 5.8774650000 | -1.2548080000 |
| H | -3.0039070000 | 6.5728840000 | -0.3691280000 |
| H | -2.8121500000 | 6.0870360000 | -2.0660190000 |
| H | -1.7878630000 | 4.3838370000 | -0.4697000000 |
| C | 1.6606070000 | 1.0395550000 | 1.6508080000 |
| O | 2.3723460000 | 1.6988290000 | 0.8811390000 |
| O | 0.7824320000 | 0.1508820000 | 1.2444030000 |
| C | 1.7505760000 | 1.2059080000 | 3.1474380000 |
| H | 0.8165460000 | 1.6478490000 | 3.5124690000 |
| H | 1.8571860000 | 0.2301370000 | 3.6310700000 |
| H | 2.5887840000 | 1.8523820000 | 3.4092590000 |
| O | 1.5400670000 | 0.9196640000 | -1.4448600000 |
| H | 2.0225710000 | 1.3165560000 | -0.6435790000 |
| H | 1.2576800000 | 1.6557150000 | -2.0300300000 |

**$^{3}$12'**

Charge: 1
Multiplicity: 3
B3LYP-D3(BJ)/6-31G(d)-SDD SCF energy (au): -1693.79805708
B3LYP-D3(BJ)/6-31G(d)-SDD enthalpy (au): -1693.21993008
B3LYP-D3(BJ)/6-31G(d)-SDD free energy (au): -1693.33406008
B3LYP-D3(BJ)/def2-TZVP/SMD SCF energy (au): -2834.36237751
B3LYP-D3(BJ)/def2-TZVP/SMD enthalpy (au): -2833.78425051
B3LYP-D3(BJ)/def2-TZVP/SMD free energy (au): -2833.89838051
B3LYP-D3(BJ)/def2-TZVP/SMD free energy (quasi-harmonic) (au): -2833.88684843

Cartesian coordinates

| ATOM | X | Y | Z |
|---|---|---|---|
| Fe | -0.2734510000 | -0.1775650000 | -0.1454740000 |
| C | -3.0219730000 | -2.9070000000 | 1.3598440000 |
| N | -2.5284660000 | -2.2971190000 | 2.5001310000 |
| H | -2.8227640000 | -2.4864870000 | 3.4486140000 |
| C | -1.5874050000 | -1.3990950000 | 2.1415220000 |
| H | -1.0125600000 | -0.7770570000 | 2.8085440000 |
| N | -1.4607200000 | -1.4089870000 | 0.8212210000 |
| C | -2.3458030000 | -2.3425290000 | 0.3144800000 |
| H | -2.4139200000 | -2.5390950000 | -0.7435610000 |
| H | -3.7858170000 | -3.6671850000 | 1.3971630000 |
| C | 1.4705170000 | -3.4067930000 | -2.2008040000 |
| N | 2.2125790000 | -3.3547580000 | -1.0343390000 |
| H | 3.0039240000 | -3.9415500000 | -0.8078770000 |
| C | 1.7190110000 | -2.3726510000 | -0.2503360000 |
| H | 2.0962540000 | -2.0839300000 | 0.7179210000 |
| N | 0.6919530000 | -1.8003210000 | -0.8640850000 |

| | | | |
|---|---|---|---|
| C | 0.5222830000 | -2.4285210000 | -2.0838300000 |
| H | -0.2523340000 | -2.1187270000 | -2.7661780000 |
| H | 1.6833530000 | -4.1127550000 | -2.9876490000 |
| O | -1.6289460000 | -0.2352270000 | -1.9857820000 |
| H | -1.3416930000 | 0.6037660000 | -2.4335580000 |
| H | -2.4884900000 | 0.0066050000 | -1.6005640000 |
| F | -1.2637180000 | 1.2024780000 | 0.3508650000 |
| C | -1.3155390000 | 3.2429070000 | -2.8297550000 |
| O | -0.6600390000 | 2.1647840000 | -2.7764400000 |
| N | -0.6740790000 | 4.4227990000 | -2.6955250000 |
| H | -2.7350910000 | 5.2455540000 | -1.2265040000 |
| C | 0.2838280000 | 4.8161240000 | -1.6690380000 |
| C | 0.3302240000 | 6.3553990000 | -1.6536260000 |
| C | -0.0635010000 | 4.2679000000 | -0.2766770000 |
| C | 1.6676830000 | 4.2984350000 | -2.1589200000 |
| H | 0.5552160000 | 6.7483120000 | -2.6490700000 |
| H | -0.6241570000 | 6.7723290000 | -1.3197240000 |
| H | 1.1084450000 | 6.6863340000 | -0.9594110000 |
| H | -0.2649710000 | 3.1967000000 | -0.2862350000 |
| H | 0.7662730000 | 4.4468790000 | 0.4133690000 |
| H | -0.9504500000 | 4.7725230000 | 0.1211920000 |
| H | 2.4307230000 | 4.6728440000 | -1.4692730000 |
| H | 1.7117610000 | 3.2098500000 | -2.1729480000 |
| H | 1.8815540000 | 4.6701180000 | -3.1641790000 |
| C | -2.7514530000 | 3.2342390000 | -3.2084380000 |
| C | -3.2103350000 | 2.0908250000 | -3.8851990000 |
| C | -3.6445310000 | 4.2923530000 | -2.9086030000 |
| C | -4.5411260000 | 1.9709380000 | -4.2659750000 |
| H | -2.5010530000 | 1.3133030000 | -4.1443270000 |
| C | -4.9810990000 | 4.1409740000 | -3.2930680000 |
| C | -5.4299440000 | 3.0013000000 | -3.9601660000 |
| H | -4.8774790000 | 1.0918880000 | -4.8066660000 |
| H | -5.6901310000 | 4.9290990000 | -3.0700530000 |
| H | -6.4744040000 | 2.9251180000 | -4.2476660000 |
| C | -3.1783870000 | 5.5402310000 | -2.1864310000 |
| C | -4.2363040000 | 6.6153810000 | -1.9280010000 |
| H | -5.0479220000 | 6.2419850000 | -1.2943540000 |
| H | -3.7791790000 | 7.4662020000 | -1.4137730000 |
| H | -4.6715650000 | 6.9872020000 | -2.8612850000 |
| H | -2.3593510000 | 5.9816140000 | -2.7709190000 |
| C | 1.7471770000 | 0.8752090000 | 1.7973040000 |
| O | 2.0838840000 | 1.7245930000 | 0.9307850000 |
| O | 0.9812780000 | -0.1329160000 | 1.5693880000 |
| C | 2.2454860000 | 1.0486580000 | 3.2150180000 |
| H | 1.4497840000 | 1.5111120000 | 3.8118300000 |
| H | 2.4762930000 | 0.0787880000 | 3.6634820000 |
| H | 3.1200830000 | 1.7004020000 | 3.2397630000 |
| O | 0.9887160000 | 0.8926190000 | -1.1041810000 |
| H | 1.5146480000 | 1.3733880000 | -0.2920300000 |
| H | 0.5222950000 | 1.5410820000 | -1.6880850000 |

**$^{5}$12-A**

Charge: 2

Multiplicity: 5
B3LYP-D3(BJ)/6-31G(d)-SDD SCF energy (au): -1691.26926371
B3LYP-D3(BJ)/6-31G(d)-SDD enthalpy (au): -1690.66828371
B3LYP-D3(BJ)/6-31G(d)-SDD free energy (au): -1690.78412971
B3LYP-D3(BJ)/def2-TZVP/SMD SCF energy (au): -2831.93432570
B3LYP-D3(BJ)/def2-TZVP/SMD enthalpy (au): -2831.33334570
B3LYP-D3(BJ)/def2-TZVP/SMD free energy (au): -2831.44919170
B3LYP-D3(BJ)/def2-TZVP/SMD free energy (quasi-harmonic) (au): -2831.43688999

Cartesian coordinates

| ATOM | X | Y | Z |
|---|---|---|---|
| Fe | -0.1814540000 | -0.0992020000 | 0.2184810000 |
| C | 2.6499270000 | -1.1884790000 | 3.2656830000 |
| N | 3.0896060000 | 0.0628880000 | 2.8760470000 |
| H | 3.8522230000 | 0.5793900000 | 3.2974490000 |
| C | 2.3276030000 | 0.4918890000 | 1.8508060000 |
| H | 2.4387530000 | 1.4419670000 | 1.3516950000 |
| N | 1.4158040000 | -0.4325540000 | 1.5559670000 |
| C | 1.6030150000 | -1.4870430000 | 2.4385530000 |
| H | 0.9731060000 | -2.3626860000 | 2.4159670000 |
| C | -2.6011560000 | -3.1778910000 | 1.9981840000 |
| N | -2.8453210000 | -2.0693830000 | 2.7891780000 |
| H | -3.4505370000 | -2.0438540000 | 3.6012630000 |
| C | -2.1557300000 | -1.0230020000 | 2.2985250000 |
| H | -2.1531810000 | -0.0263170000 | 2.7129510000 |
| N | -1.4714000000 | -1.4092910000 | 1.2191200000 |
| C | -1.7421540000 | -2.7559660000 | 1.0222290000 |
| H | -1.3160980000 | -3.3037250000 | 0.1970600000 |
| O | 1.0659310000 | 1.1956590000 | -0.9138350000 |
| H | 1.5213590000 | 0.7529890000 | -1.6504830000 |
| H | 0.4924340000 | 1.9157630000 | -1.3248360000 |
| H | -3.0549750000 | -4.1368410000 | 2.1945020000 |
| H | 3.1081440000 | -1.7339680000 | 4.0759370000 |
| C | 0.6380680000 | -3.2118550000 | -2.6936610000 |
| N | 1.7393990000 | -3.3340210000 | -1.8681850000 |
| H | 2.4903580000 | -4.0046370000 | -1.9792230000 |
| C | 1.6554690000 | -2.4054540000 | -0.8941290000 |
| H | 2.3830830000 | -2.2826680000 | -0.1075950000 |
| N | 0.5448960000 | -1.6842870000 | -1.0516950000 |
| C | -0.0995390000 | -2.1822290000 | -2.1781720000 |
| H | -1.0269410000 | -1.7543330000 | -2.5215200000 |
| H | 0.4871470000 | -3.8489730000 | -3.5512580000 |
| O | -1.6778240000 | 0.2026640000 | -1.2848250000 |
| H | -1.5206170000 | 1.1201840000 | -1.6824260000 |
| H | -2.5543050000 | 0.2382250000 | -0.8639380000 |
| F | -0.8116500000 | 1.3310480000 | 1.1238820000 |
| C | -1.4318430000 | 3.5997350000 | -2.3699120000 |
| O | -0.8221690000 | 2.5204940000 | -2.0597280000 |
| N | -0.7993670000 | 4.7677940000 | -2.2028030000 |
| H | -4.3683760000 | 6.3536510000 | -2.0948510000 |
| C | -0.1003040000 | 5.2967470000 | -1.0371140000 |
| C | -0.1475500000 | 6.8362280000 | -1.1280370000 |
| C | -0.6943480000 | 4.7990370000 | 0.2875580000 |

| | | | |
|---|---|---|---|
| C | 1.3818110000 | 4.8555780000 | -1.1955930000 |
| H | 0.2290930000 | 7.1814250000 | -2.0940000000 |
| H | -1.1654570000 | 7.2097120000 | -0.9955100000 |
| H | 0.4764090000 | 7.2572680000 | -0.3346490000 |
| H | -0.6665200000 | 3.7086700000 | 0.3655390000 |
| H | -0.1266210000 | 5.2173590000 | 1.1244510000 |
| H | -1.7329340000 | 5.1298400000 | 0.3935120000 |
| H | 1.9680650000 | 5.3542000000 | -0.4171660000 |
| H | 1.4917740000 | 3.7759930000 | -1.0760700000 |
| H | 1.7723530000 | 5.1501340000 | -2.1725750000 |
| C | -2.7394550000 | 3.5337720000 | -3.0524160000 |
| C | -3.0461770000 | 2.3156890000 | -3.6957990000 |
| C | -3.6666560000 | 4.6020610000 | -3.0827250000 |
| C | -4.2505270000 | 2.1417670000 | -4.3603030000 |
| H | -2.3066310000 | 1.5231460000 | -3.7057780000 |
| C | -4.8818250000 | 4.3920840000 | -3.7468800000 |
| C | -5.1777530000 | 3.1879650000 | -4.3794680000 |
| H | -4.4636310000 | 1.2079740000 | -4.8704450000 |
| H | -5.6083590000 | 5.1993020000 | -3.7661330000 |
| H | -6.1272890000 | 3.0665270000 | -4.8914980000 |
| C | -3.4183350000 | 5.9607580000 | -2.4741490000 |
| C | -2.8311020000 | 6.9512160000 | -3.4959930000 |
| H | -3.5178070000 | 7.0866280000 | -4.3370930000 |
| H | -2.6667860000 | 7.9289000000 | -3.0324660000 |
| H | -1.8766010000 | 6.5910320000 | -3.8907910000 |
| H | -2.7589890000 | 5.8794420000 | -1.6043730000 |

**$^{3}$12-A**

Charge: 2
Multiplicity: 3
B3LYP-D3(BJ)/6-31G(d)-SDD SCF energy (au): -1691.24777634
B3LYP-D3(BJ)/6-31G(d)-SDD enthalpy (au): -1690.64601234
B3LYP-D3(BJ)/6-31G(d)-SDD free energy (au): -1690.75890734
B3LYP-D3(BJ)/def2-TZVP/SMD SCF energy (au): -2831.91574111
B3LYP-D3(BJ)/def2-TZVP/SMD enthalpy (au): -2831.31397711
B3LYP-D3(BJ)/def2-TZVP/SMD free energy (au): -2831.42687211
B3LYP-D3(BJ)/def2-TZVP/SMD free energy (quasi-harmonic) (au): -2831.41509377

Cartesian coordinates

| ATOM | X | Y | Z |
|---|---|---|---|
| Fe | -0.2956380000 | -0.2590180000 | -0.6342280000 |
| C | 1.7281990000 | 1.3621860000 | 2.8167810000 |
| N | 2.4207780000 | 1.9466940000 | 1.7740430000 |
| H | 3.1504370000 | 2.6427830000 | 1.8659690000 |
| C | 1.9582070000 | 1.4430120000 | 0.6103690000 |
| H | 2.3076300000 | 1.7327540000 | -0.3682980000 |
| N | 1.0020660000 | 0.5526330000 | 0.8568010000 |
| C | 0.8428760000 | 0.4978880000 | 2.2330110000 |
| H | 0.1072390000 | -0.1432810000 | 2.6935300000 |
| C | -3.3117790000 | -1.4932070000 | 2.0026680000 |
| N | -3.7784980000 | -0.2875530000 | 1.5092720000 |
| H | -4.6541850000 | 0.1545560000 | 1.7612240000 |
| C | -2.8817840000 | 0.2117350000 | 0.6405810000 |

| | | | |
|---|---|---|---|
| H | -2.9825170000 | 1.1363200000 | 0.0939710000 |
| N | -1.8485310000 | -0.6308410000 | 0.5494120000 |
| C | -2.1061350000 | -1.7016640000 | 1.3951210000 |
| H | -1.4236470000 | -2.5298320000 | 1.4903900000 |
| O | 1.0729580000 | 0.1219290000 | -2.0980650000 |
| H | 1.5173640000 | -0.6667520000 | -2.4557920000 |
| H | 0.5742150000 | 0.5528820000 | -2.8572620000 |
| H | -3.8669790000 | -2.0775710000 | 2.7198180000 |
| H | 1.9219890000 | 1.6103100000 | 3.8487190000 |
| C | 0.7981390000 | -4.2984040000 | -0.5076020000 |
| N | 1.7257800000 | -3.7422330000 | 0.3502930000 |
| H | 2.4566450000 | -4.2453360000 | 0.8391360000 |
| C | 1.5141160000 | -2.4132700000 | 0.4231330000 |
| H | 2.0896050000 | -1.7308780000 | 1.0262590000 |
| N | 0.4799190000 | -2.0890020000 | -0.3549490000 |
| C | 0.0217000000 | -3.2614620000 | -0.9446680000 |
| H | -0.8015490000 | -3.2524490000 | -1.6387190000 |
| H | 0.7765220000 | -5.3537350000 | -0.7311510000 |
| O | -1.4474780000 | -1.1067770000 | -2.2954980000 |
| H | -1.2690220000 | -0.4689450000 | -3.0495290000 |
| H | -2.4000060000 | -0.9950920000 | -2.1244460000 |
| C | -4.7995770000 | 0.3404960000 | -3.1790490000 |
| C | -3.5348120000 | 0.6052270000 | -3.6959810000 |
| C | -2.8247700000 | 1.7569320000 | -3.3100310000 |
| C | -3.3984770000 | 2.6819910000 | -2.4157620000 |
| C | -4.6668780000 | 2.3832780000 | -1.8938720000 |
| C | -5.3618400000 | 1.2322090000 | -2.2598440000 |
| H | -5.3513370000 | -0.5342810000 | -3.5085420000 |
| H | -3.0990080000 | -0.0630570000 | -4.4330840000 |
| H | -5.1221020000 | 3.0887860000 | -1.2031150000 |
| H | -6.3517660000 | 1.0443750000 | -1.8546680000 |
| C | -2.7428360000 | 3.9854790000 | -2.0261840000 |
| C | -3.2361010000 | 5.1492340000 | -2.9048000000 |
| H | -1.6543230000 | 3.8985770000 | -2.0867000000 |
| H | -2.9745830000 | 4.2023440000 | -0.9768110000 |
| H | -2.7499770000 | 6.0859650000 | -2.6143740000 |
| H | -3.0232900000 | 4.9610290000 | -3.9616040000 |
| H | -4.3171740000 | 5.2838760000 | -2.8006290000 |
| C | -1.4322340000 | 1.8849640000 | -3.8269510000 |
| O | -0.6889510000 | 0.8503400000 | -3.9190400000 |
| N | -1.0300490000 | 3.0407830000 | -4.3276760000 |
| C | 0.2359270000 | 3.7375610000 | -4.4078710000 |
| C | -0.0651300000 | 5.2152840000 | -4.7339160000 |
| C | 0.9788610000 | 3.0960970000 | -5.6176060000 |
| H | 0.8774590000 | 5.7358560000 | -4.9240300000 |
| H | -0.6965460000 | 5.2960480000 | -5.6220880000 |
| H | -0.5697610000 | 5.7047500000 | -3.8971300000 |
| H | 1.1850900000 | 2.0404220000 | -5.4301510000 |
| H | 0.3823750000 | 3.1866750000 | -6.5279930000 |
| H | 1.9218780000 | 3.6348820000 | -5.7557920000 |
| F | -0.9501190000 | 1.3619400000 | -0.9465900000 |
| C | 1.0662920000 | 3.6215690000 | -3.1213300000 |
| H | 0.5221820000 | 4.0268320000 | -2.2629220000 |

H 1.3331320000 2.5835550000 -2.9083630000
H 1.9924270000 4.1924590000 -3.2384080000

**[5]12-B**

Charge: 1
Multiplicity: 5
B3LYP-D3(BJ)/6-31G(d)-SDD SCF energy (au): -1690.94778462
B3LYP-D3(BJ)/6-31G(d)-SDD enthalpy (au): -1690.35964562
B3LYP-D3(BJ)/6-31G(d)-SDD free energy (au): -1690.47692462
B3LYP-D3(BJ)/def2-TZVP/SMD SCF energy (au): -2831.50199626
B3LYP-D3(BJ)/def2-TZVP/SMD enthalpy (au): -2830.91385726
B3LYP-D3(BJ)/def2-TZVP/SMD free energy (au): -2831.03113626
B3LYP-D3(BJ)/def2-TZVP/SMD free energy (quasi-harmonic) (au): -2831.01777864

Cartesian coordinates

| ATOM | X | Y | Z |
|---|---|---|---|
| Fe | -0.0769010000 | 0.0505360000 | 0.0890980000 |
| C | 2.6005710000 | -1.0796200000 | 3.3471840000 |
| N | 3.2390520000 | -0.0210880000 | 2.7244280000 |
| H | 4.1052050000 | 0.4071230000 | 3.0206570000 |
| C | 2.4979230000 | 0.3596340000 | 1.6590630000 |
| H | 2.7348110000 | 1.1587370000 | 0.9732860000 |
| N | 1.4181130000 | -0.4060400000 | 1.5686270000 |
| C | 1.4668720000 | -1.3068030000 | 2.6159250000 |
| H | 0.6917500000 | -2.0428240000 | 2.7594340000 |
| C | -2.4486480000 | -3.0781440000 | 2.2475980000 |
| N | -2.8462860000 | -1.8844430000 | 2.8218140000 |
| H | -3.4939210000 | -1.7814650000 | 3.5908120000 |
| C | -2.2255490000 | -0.8713690000 | 2.1717400000 |
| H | -2.3259700000 | 0.1765420000 | 2.4069020000 |
| N | -1.4512870000 | -1.3536810000 | 1.2088080000 |
| C | -1.5810580000 | -2.7305630000 | 1.2466400000 |
| H | -1.0495930000 | -3.3634000000 | 0.5539920000 |
| O | 1.1821470000 | 1.0291140000 | -0.8464580000 |
| H | 0.7793330000 | 1.7484100000 | -1.3687490000 |
| H | -2.8153320000 | -4.0327360000 | 2.5908650000 |
| H | 3.0024010000 | -1.5562450000 | 4.2272820000 |
| C | 0.4577810000 | -3.3719550000 | -2.5984700000 |
| N | 1.7429450000 | -3.1445110000 | -2.1399920000 |
| H | 2.5786280000 | -3.6155590000 | -2.4572100000 |
| C | 1.6939780000 | -2.1574320000 | -1.2141710000 |
| H | 2.5557850000 | -1.7538050000 | -0.7068570000 |
| N | 0.4464150000 | -1.7498090000 | -1.0505510000 |
| C | -0.3372220000 | -2.4934510000 | -1.9129560000 |
| H | -1.3987900000 | -2.3209410000 | -1.9866340000 |
| H | 0.2387460000 | -4.1121740000 | -3.3515850000 |
| O | -1.6749300000 | 0.2344920000 | -1.4065930000 |
| H | -1.3978600000 | 1.0646240000 | -1.8839380000 |
| H | -2.4192250000 | 0.5178990000 | -0.8484740000 |
| F | -1.0206080000 | 1.3575390000 | 1.0075800000 |
| C | -1.3640780000 | 3.6174910000 | -2.4837600000 |
| O | -0.7067990000 | 2.5677810000 | -2.2788180000 |
| N | -0.7916980000 | 4.8179710000 | -2.2146580000 |

| | | | |
|---|---|---|---|
| H | -4.4073530000 | 6.2863770000 | -2.0839050000 |
| C | -0.1699370000 | 5.2112530000 | -0.9523930000 |
| C | -0.1577820000 | 6.7509250000 | -0.9222750000 |
| C | -0.8820820000 | 4.6354080000 | 0.2807080000 |
| C | 1.3004200000 | 4.7100690000 | -1.0131940000 |
| H | 0.3277290000 | 7.1517770000 | -1.8163730000 |
| H | -1.1727610000 | 7.1545760000 | -0.8698910000 |
| H | 0.3902230000 | 7.0929650000 | -0.0390640000 |
| H | -0.8774460000 | 3.5424990000 | 0.2891160000 |
| H | -0.3743690000 | 4.9792730000 | 1.1880750000 |
| H | -1.9187300000 | 4.9878510000 | 0.3295630000 |
| H | 1.8447280000 | 5.1630010000 | -0.1773750000 |
| H | 1.3497680000 | 3.6255470000 | -0.9173550000 |
| H | 1.7766540000 | 5.0142920000 | -1.9489560000 |
| C | -2.6824470000 | 3.5648180000 | -3.1714360000 |
| C | -2.9569880000 | 2.3768010000 | -3.8757280000 |
| C | -3.6426430000 | 4.6016190000 | -3.1391290000 |
| C | -4.1607000000 | 2.1993650000 | -4.5435130000 |
| H | -2.1968950000 | 1.6052050000 | -3.9129160000 |
| C | -4.8542640000 | 4.3919520000 | -3.8106600000 |
| C | -5.1184350000 | 3.2150520000 | -4.5058960000 |
| H | -4.3491520000 | 1.2824270000 | -5.0929700000 |
| H | -5.6046680000 | 5.1774740000 | -3.7826300000 |
| H | -6.0680650000 | 3.0915400000 | -5.0181120000 |
| C | -3.4381800000 | 5.9342740000 | -2.4564370000 |
| C | -2.8468030000 | 6.9887240000 | -3.4079140000 |
| H | -3.5087780000 | 7.1475210000 | -4.2654390000 |
| H | -2.7218350000 | 7.9464810000 | -2.8918160000 |
| H | -1.8702530000 | 6.6690650000 | -3.7815270000 |
| H | -2.7974810000 | 5.8251840000 | -1.5775000000 |

**$^{3}$12-B**

Charge: 1
Multiplicity: 3
B3LYP-D3(BJ)/6-31G(d)-SDD SCF energy (au): -1690.93631724
B3LYP-D3(BJ)/6-31G(d)-SDD enthalpy (au): -1690.34751024
B3LYP-D3(BJ)/6-31G(d)-SDD free energy (au): -1690.46082024
B3LYP-D3(BJ)/def2-TZVP/SMD SCF energy (au): -2831.47842479
B3LYP-D3(BJ)/def2-TZVP/SMD enthalpy (au): -2830.88961779
B3LYP-D3(BJ)/def2-TZVP/SMD free energy (au): -2831.00292779
B3LYP-D3(BJ)/def2-TZVP/SMD free energy (quasi-harmonic) (au): -2830.99173955

Cartesian coordinates

| ATOM | X | Y | Z |
|---|---|---|---|
| Fe | -0.2510690000 | -0.1707470000 | -0.7686630000 |
| C | 1.7436040000 | 1.5526530000 | 2.7532960000 |
| N | 2.5226050000 | 1.9594470000 | 1.6848900000 |
| H | 3.3113950000 | 2.5892190000 | 1.7318140000 |
| C | 2.0345100000 | 1.3849210000 | 0.5584400000 |
| H | 2.4249880000 | 1.5237750000 | -0.4379970000 |
| N | 0.9887250000 | 0.6281720000 | 0.8535720000 |
| C | 0.7925390000 | 0.7261180000 | 2.2169980000 |
| H | -0.0146970000 | 0.2041160000 | 2.7075810000 |

| | | | |
|---|---|---|---|
| C | -3.2533350000 | -1.2852890000 | 2.0346120000 |
| N | -3.7883530000 | -0.1824840000 | 1.3943910000 |
| H | -4.6895060000 | 0.2350300000 | 1.5808440000 |
| C | -2.9025710000 | 0.2557500000 | 0.4721140000 |
| H | -3.0397000000 | 1.1086710000 | -0.1737550000 |
| N | -1.8249700000 | -0.5205040000 | 0.4885850000 |
| C | -2.0279500000 | -1.4856080000 | 1.4590460000 |
| H | -1.2911430000 | -2.2467810000 | 1.6571620000 |
| O | 1.1232930000 | 0.0670760000 | -1.9073560000 |
| H | 0.7414620000 | 0.3627160000 | -2.7560580000 |
| H | -3.7790350000 | -1.8109080000 | 2.8159290000 |
| H | 1.9346340000 | 1.8815690000 | 3.7626460000 |
| C | 0.7699490000 | -4.3112380000 | -0.4573120000 |
| N | 1.9148090000 | -3.6837000000 | 0.0002300000 |
| H | 2.7806310000 | -4.1388120000 | 0.2535630000 |
| C | 1.6906990000 | -2.3500010000 | 0.0230050000 |
| H | 2.4168600000 | -1.6084010000 | 0.3130910000 |
| N | 0.4582430000 | -2.0955980000 | -0.3867650000 |
| C | -0.1284670000 | -3.3082340000 | -0.6984940000 |
| H | -1.1283940000 | -3.3569690000 | -1.0962370000 |
| H | 0.7069890000 | -5.3815160000 | -0.5730340000 |
| O | -1.4836240000 | -1.1497410000 | -2.3556420000 |
| H | -1.2308170000 | -0.5950200000 | -3.1378180000 |
| H | -2.4021870000 | -0.8796910000 | -2.1871840000 |
| C | -4.8778310000 | 0.3018590000 | -3.2629140000 |
| C | -3.6011950000 | 0.5674310000 | -3.7517300000 |
| C | -2.8980880000 | 1.7175870000 | -3.3579080000 |
| C | -3.4937540000 | 2.6410990000 | -2.4761480000 |
| C | -4.7747020000 | 2.3474790000 | -1.9860100000 |
| C | -5.4632460000 | 1.1956200000 | -2.3633490000 |
| H | -5.4140250000 | -0.5830000000 | -3.5917430000 |
| H | -3.1387370000 | -0.1091040000 | -4.4634700000 |
| H | -5.2400380000 | 3.0513470000 | -1.2997730000 |
| H | -6.4595870000 | 1.0059690000 | -1.9741710000 |
| C | -2.8386050000 | 3.9340370000 | -2.0517860000 |
| C | -3.2814390000 | 5.1135120000 | -2.9342760000 |
| H | -1.7522820000 | 3.8282900000 | -2.0667780000 |
| H | -3.1085740000 | 4.1404150000 | -1.0088450000 |
| H | -2.7990450000 | 6.0418770000 | -2.6102640000 |
| H | -3.0175830000 | 4.9345790000 | -3.9805720000 |
| H | -4.3656300000 | 5.2575220000 | -2.8773820000 |
| C | -1.4974620000 | 1.8490440000 | -3.8708890000 |
| O | -0.7939540000 | 0.8337990000 | -4.0868780000 |
| N | -1.0919840000 | 3.0704450000 | -4.2810780000 |
| C | 0.2340800000 | 3.6604600000 | -4.2048890000 |
| C | 0.0707640000 | 5.1681250000 | -4.4696090000 |
| C | 1.0354350000 | 3.0267700000 | -5.3809960000 |
| H | 1.0569900000 | 5.6378560000 | -4.5343600000 |
| H | -0.4644080000 | 5.3413110000 | -5.4073170000 |
| H | -0.4854850000 | 5.6476380000 | -3.6589650000 |
| H | 1.1766890000 | 1.9576930000 | -5.2179950000 |
| H | 0.5110100000 | 3.1768100000 | -6.3282690000 |
| H | 2.0095020000 | 3.5249240000 | -5.4328080000 |

| | | | |
|---|---|---|---|
| F | -1.0102640000 | 1.4504810000 | -1.1136430000 |
| C | 0.9563830000 | 3.4225230000 | -2.8703670000 |
| H | 0.3996160000 | 3.8642080000 | -2.0387850000 |
| H | 1.0903220000 | 2.3624570000 | -2.6566640000 |
| H | 1.9422640000 | 3.8985550000 | -2.9116840000 |

**$^5$13'**
Charge: 1
Multiplicity: 5
B3LYP-D3(BJ)/6-31G(d)-SDD SCF energy (au): -1693.85868757
B3LYP-D3(BJ)/6-31G(d)-SDD enthalpy (au): -1693.27916557
B3LYP-D3(BJ)/6-31G(d)-SDD free energy (au): -1693.39509557
B3LYP-D3(BJ)/def2-TZVP/SMD SCF energy (au): -2834.42177427
B3LYP-D3(BJ)/def2-TZVP/SMD enthalpy (au): -2833.84225227
B3LYP-D3(BJ)/def2-TZVP/SMD free energy (au): -2833.95818227
B3LYP-D3(BJ)/def2-TZVP/SMD free energy (quasi-harmonic) (au): -2833.94602970

Cartesian coordinates

| ATOM | X | Y | Z |
|---|---|---|---|
| Fe | -0.0366110000 | 0.0610980000 | -0.5321600000 |
| C | -3.6348650000 | -2.1100640000 | 0.6179660000 |
| N | -3.9494630000 | -0.7885340000 | 0.8833270000 |
| H | -4.8185600000 | -0.4541260000 | 1.2762120000 |
| C | -2.8951920000 | -0.0166910000 | 0.5405490000 |
| H | -2.8458800000 | 1.0572240000 | 0.6291220000 |
| N | -1.9185010000 | -0.7826260000 | 0.0655440000 |
| C | -2.3661030000 | -2.0907470000 | 0.1080950000 |
| H | -1.7476390000 | -2.9056190000 | -0.2318420000 |
| H | -4.3242940000 | -2.9161020000 | 0.8130620000 |
| C | 1.1596150000 | -3.7584740000 | -2.2221640000 |
| N | 1.7577560000 | -3.7863670000 | -0.9744130000 |
| H | 2.3633000000 | -4.5151480000 | -0.6223470000 |
| C | 1.4219950000 | -2.6566950000 | -0.3134350000 |
| H | 1.7495940000 | -2.3971530000 | 0.6806030000 |
| N | 0.6317820000 | -1.9094680000 | -1.0740850000 |
| C | 0.4621200000 | -2.5830740000 | -2.2711920000 |
| H | -0.1269180000 | -2.1558710000 | -3.0660650000 |
| H | 1.2912570000 | -4.5519220000 | -2.9405960000 |
| O | -0.9008090000 | 0.1110770000 | -2.5290050000 |
| H | -0.6028200000 | 1.0079950000 | -2.8732840000 |
| H | -1.8629410000 | 0.2047380000 | -2.4277330000 |
| F | -0.7586510000 | 1.7049770000 | -0.1490660000 |
| C | -0.9315450000 | 3.3833780000 | -2.5402040000 |
| O | -0.0589360000 | 2.5789140000 | -2.9600530000 |
| N | -0.6466880000 | 4.6225610000 | -2.1277410000 |
| H | -1.8523700000 | 3.9097570000 | -0.0516430000 |
| C | 0.6905290000 | 5.2590390000 | -1.9697590000 |
| C | 0.4084260000 | 6.7138160000 | -1.5707570000 |
| C | 1.4681720000 | 4.5528840000 | -0.8495390000 |
| C | 1.4571110000 | 5.2185060000 | -3.2996730000 |
| H | -0.1630020000 | 7.2337080000 | -2.3480910000 |
| H | -0.1490200000 | 6.7658740000 | -0.6276170000 |
| H | 1.3508210000 | 7.2497910000 | -1.4309430000 |

```
H      1.6778650000    3.5131950000   -1.0965820000
H      2.4250650000    5.0586360000   -0.6871790000
H      0.9052610000    4.5676160000    0.0888030000
H      2.4152890000    5.7345920000   -3.1826080000
H      1.6493860000    4.1921350000   -3.6157510000
H      0.8889320000    5.7247220000   -4.0869040000
C     -2.3785070000    2.9974540000   -2.5728480000
C     -2.8102180000    2.3573320000   -3.7363600000
C     -3.3026310000    3.2538310000   -1.5003980000
C     -4.1422510000    1.9735570000   -3.9061710000
H     -2.0902410000    2.1817670000   -4.5300160000
C     -4.6578250000    2.8645340000   -1.7191590000
C     -5.0656920000    2.2399220000   -2.8860700000
H     -4.4580030000    1.4942510000   -4.8269210000
H     -5.3851440000    3.0511690000   -0.9364890000
H     -6.1073240000    1.9587170000   -3.0101410000
C     -2.9094890000    3.8031930000   -0.2547120000
C     -3.8609160000    4.1272040000    0.8539800000
H     -4.2822430000    3.2211870000    1.3204960000
H     -3.3573440000    4.6870220000    1.6460510000
H     -4.7167470000    4.7223080000    0.5108030000
H     -1.4430290000    5.1916610000   -1.8715010000
C      1.9070710000    0.7222920000    1.5397440000
O      2.5623730000    1.4675620000    0.8001940000
O      0.9974160000   -0.1199190000    1.1028300000
C      2.1082380000    0.7180890000    3.0355560000
H      1.2031990000    1.1046060000    3.5176360000
H      2.2541350000   -0.3052740000    3.3946860000
H      2.9621110000    1.3397400000    3.3067880000
O      1.5847140000    0.9054870000   -1.5378410000
H      2.1143940000    1.2330520000   -0.7371370000
H      1.2558180000    1.6910920000   -2.0317870000
```

**$^{3}$13'**

```
Charge: 1
Multiplicity: 3
B3LYP-D3(BJ)/6-31G(d)-SDD SCF energy (au):              -1693.83571627
B3LYP-D3(BJ)/6-31G(d)-SDD enthalpy (au):                -1693.25546427
B3LYP-D3(BJ)/6-31G(d)-SDD free energy (au):             -1693.36866727
B3LYP-D3(BJ)/def2-TZVP/SMD SCF energy (au):             -2834.40136335
B3LYP-D3(BJ)/def2-TZVP/SMD enthalpy (au):               -2833.82111135
B3LYP-D3(BJ)/def2-TZVP/SMD free energy (au):            -2833.93431435
B3LYP-D3(BJ)/def2-TZVP/SMD free energy (quasi-harmonic) (au): -2833.92314408

Cartesian coordinates
ATOM    X               Y               Z
Fe    -0.0292070000   -0.0405310000   -0.6187560000
C     -3.7314380000   -2.3274390000    0.1179860000
N     -4.0988140000   -1.0205320000    0.3829950000
H     -5.0120170000   -0.7106210000    0.6849080000
C     -3.0270810000   -0.2217050000    0.1722200000
H     -3.0155460000    0.8499310000    0.2914770000
N     -1.9915700000   -0.9538200000   -0.2175830000
```

| | | | |
|---|---|---|---|
| C | -2.4160080000 | -2.2687840000 | -0.2556620000 |
| H | -1.7510950000 | -3.0655820000 | -0.5500490000 |
| H | -4.4204980000 | -3.1510260000 | 0.2188820000 |
| C | 1.4166810000 | -3.6526460000 | -2.0970060000 |
| N | 1.8440640000 | -3.6838930000 | -0.7797840000 |
| H | 2.3985550000 | -4.4136020000 | -0.3529150000 |
| C | 1.4245200000 | -2.5624690000 | -0.1607310000 |
| H | 1.6118780000 | -2.2992380000 | 0.8674840000 |
| N | 0.7393050000 | -1.8178400000 | -1.0220330000 |
| C | 0.7283260000 | -2.4808310000 | -2.2376830000 |
| H | 0.2446960000 | -2.0508880000 | -3.0988490000 |
| H | 1.6442650000 | -4.4426570000 | -2.7947740000 |
| O | -0.7454170000 | 0.0748050000 | -2.5660950000 |
| H | -0.5216040000 | 1.0185240000 | -2.8578040000 |
| H | -1.7137990000 | 0.0696220000 | -2.4724420000 |
| F | -0.8127430000 | 1.5172800000 | -0.2171910000 |
| C | -0.8819790000 | 3.3611570000 | -2.4785210000 |
| O | -0.0498200000 | 2.5611810000 | -2.9833440000 |
| N | -0.5498900000 | 4.5982820000 | -2.0968830000 |
| H | -1.5687220000 | 3.9672250000 | 0.0687300000 |
| C | 0.7830990000 | 5.2580310000 | -2.1737340000 |
| C | 0.5981310000 | 6.6245040000 | -1.4999060000 |
| C | 1.8411010000 | 4.4442160000 | -1.4121100000 |
| C | 1.1803650000 | 5.4316270000 | -3.6472290000 |
| H | -0.1817750000 | 7.2118420000 | -1.9995640000 |
| H | 0.3321450000 | 6.5111240000 | -0.4425850000 |
| H | 1.5290990000 | 7.1945840000 | -1.5545770000 |
| H | 2.1526530000 | 3.5657550000 | -1.9783720000 |
| H | 2.7296650000 | 5.0650320000 | -1.2602790000 |
| H | 1.4758470000 | 4.1233360000 | -0.4318520000 |
| H | 2.1574200000 | 5.9212450000 | -3.7124710000 |
| H | 1.2457630000 | 4.4607970000 | -4.1443270000 |
| H | 0.4497410000 | 6.0512310000 | -4.1776650000 |
| C | -2.3231020000 | 2.9745760000 | -2.3592040000 |
| C | -2.8569280000 | 2.2826550000 | -3.4477180000 |
| C | -3.1468260000 | 3.2787870000 | -1.2193180000 |
| C | -4.1958570000 | 1.8856560000 | -3.4761750000 |
| H | -2.2146430000 | 2.0794750000 | -4.2994920000 |
| C | -4.5160280000 | 2.8825360000 | -1.2975200000 |
| C | -5.0244100000 | 2.1998750000 | -2.3900450000 |
| H | -4.5920290000 | 1.3655760000 | -4.3421710000 |
| H | -5.1700980000 | 3.1079040000 | -0.4620820000 |
| H | -6.0723440000 | 1.9145680000 | -2.4068280000 |
| C | -2.6414610000 | 3.8691590000 | -0.0354910000 |
| C | -3.4788450000 | 4.2329590000 | 1.1489070000 |
| H | -3.7740280000 | 3.3444690000 | 1.7309860000 |
| H | -2.9234020000 | 4.8839370000 | 1.8292930000 |
| H | -4.4051900000 | 4.7487070000 | 0.8687210000 |
| H | -1.3168880000 | 5.1840710000 | -1.7925690000 |
| C | 1.3833770000 | 0.8624050000 | 1.6520760000 |
| O | 2.0043640000 | 1.7199330000 | 1.0181400000 |
| O | 0.7476140000 | -0.1543010000 | 1.1034190000 |
| C | 1.3085430000 | 0.8875130000 | 3.1609510000 |

H 0.2885920000 1.1582760000 3.4574030000
H 1.5144700000 -0.1046840000 3.5727550000
H 2.0114280000 1.6185480000 3.5621980000
O 1.6732330000 0.9462280000 -1.4688230000
H 1.9703530000 1.3664190000 -0.6051810000
H 1.3071490000 1.6769170000 -2.0120710000

**$^{5}$13**
Charge: 1
Multiplicity: 5
B3LYP-D3(BJ)/6-31G(d)-SDD SCF energy (au): -1690.98271601
B3LYP-D3(BJ)/6-31G(d)-SDD enthalpy (au): -1690.39465201
B3LYP-D3(BJ)/6-31G(d)-SDD free energy (au): -1690.51169001
B3LYP-D3(BJ)/def2-TZVP/SMD SCF energy (au): -2831.54132052
B3LYP-D3(BJ)/def2-TZVP/SMD enthalpy (au): -2830.95325652
B3LYP-D3(BJ)/def2-TZVP/SMD free energy (au): -2831.07029452
B3LYP-D3(BJ)/def2-TZVP/SMD free energy (quasi-harmonic) (au): -2831.05707651

Cartesian coordinates

| ATOM | X | Y | Z |
|---|---|---|---|
| Fe | 0.0509230000 | 0.2135410000 | -0.5651570000 |
| C | 1.3436160000 | -0.2875560000 | 3.5697400000 |
| N | 2.4075600000 | 0.3411550000 | 2.9468170000 |
| H | 3.2578120000 | 0.6531540000 | 3.3953330000 |
| C | 2.1109690000 | 0.4873750000 | 1.6343680000 |
| H | 2.7424030000 | 0.9462970000 | 0.8882020000 |
| N | 0.9118620000 | -0.0207830000 | 1.3888700000 |
| C | 0.4175440000 | -0.5043700000 | 2.5848300000 |
| H | -0.5586630000 | -0.9601260000 | 2.6430420000 |
| C | -3.6759560000 | -2.0072740000 | 0.5697930000 |
| N | -3.9963610000 | -0.6771720000 | 0.7766790000 |
| H | -4.8918630000 | -0.3233350000 | 1.0827660000 |
| C | -2.9084300000 | 0.0755680000 | 0.4850070000 |
| H | -2.8565640000 | 1.1515100000 | 0.5401290000 |
| N | -1.9062440000 | -0.7058370000 | 0.1027780000 |
| C | -2.3727850000 | -2.0068840000 | 0.1494250000 |
| H | -1.7433340000 | -2.8359150000 | -0.1322760000 |
| O | 1.6141640000 | 0.8643820000 | -1.3063450000 |
| H | 1.4305500000 | 1.6433490000 | -1.8633200000 |
| H | -4.3861250000 | -2.8025310000 | 0.7332660000 |
| H | 1.3441890000 | -0.5084690000 | 4.6254430000 |
| C | 1.1974520000 | -3.6940140000 | -2.1501570000 |
| N | 1.6170610000 | -3.8418170000 | -0.8405830000 |
| H | 2.1180290000 | -4.6339530000 | -0.4633840000 |
| C | 1.2674140000 | -2.7258730000 | -0.1557850000 |
| H | 1.4760150000 | -2.5678890000 | 0.8905840000 |
| N | 0.6400250000 | -1.8787240000 | -0.9570230000 |
| C | 0.5926770000 | -2.4678330000 | -2.2073160000 |
| H | 0.1367240000 | -1.9541930000 | -3.0378070000 |
| H | 1.3710870000 | -4.4515220000 | -2.8979270000 |
| O | -0.8962550000 | 0.1442170000 | -2.5114530000 |
| H | -0.6027800000 | 1.0417420000 | -2.8630240000 |
| H | -1.8548400000 | 0.2347530000 | -2.3817930000 |

| | | | |
|---|---|---|---|
| F | -0.8110930000 | 1.7961700000 | -0.1246390000 |
| C | -0.9991620000 | 3.3943160000 | -2.5586710000 |
| O | -0.1135050000 | 2.6123910000 | -2.9763820000 |
| N | -0.7346980000 | 4.6374780000 | -2.1301690000 |
| H | -1.9701800000 | 3.8997760000 | -0.0764640000 |
| C | 0.5970570000 | 5.2613520000 | -1.9201220000 |
| C | 0.3147090000 | 6.7163490000 | -1.5216790000 |
| C | 1.3282790000 | 4.5395880000 | -0.7766400000 |
| C | 1.4153100000 | 5.2254640000 | -3.2193900000 |
| H | -0.2196480000 | 7.2472350000 | -2.3176660000 |
| H | -0.2816320000 | 6.7674100000 | -0.6023330000 |
| H | 1.2557510000 | 7.2421320000 | -1.3388050000 |
| H | 1.5044250000 | 3.4888720000 | -1.0094140000 |
| H | 2.2951550000 | 5.0220300000 | -0.5971180000 |
| H | 0.7405550000 | 4.5835020000 | 0.1460260000 |
| H | 2.3680100000 | 5.7424480000 | -3.0646410000 |
| H | 1.6173320000 | 4.1996020000 | -3.5297840000 |
| H | 0.8764900000 | 5.7336120000 | -4.0259090000 |
| C | -2.4445370000 | 2.9957460000 | -2.6086770000 |
| C | -2.8519010000 | 2.3459350000 | -3.7750350000 |
| C | -3.3911990000 | 3.2542950000 | -1.5566650000 |
| C | -4.1789310000 | 1.9560910000 | -3.9688410000 |
| H | -2.1144070000 | 2.1661410000 | -4.5512310000 |
| C | -4.7408140000 | 2.8607910000 | -1.7996630000 |
| C | -5.1237360000 | 2.2271680000 | -2.9703100000 |
| H | -4.4740080000 | 1.4670200000 | -4.8913960000 |
| H | -5.4838560000 | 3.0488970000 | -1.0320860000 |
| H | -6.1618260000 | 1.9412710000 | -3.1129400000 |
| C | -3.0232860000 | 3.8093020000 | -0.3054260000 |
| C | -3.9980890000 | 4.1378350000 | 0.7815450000 |
| H | -4.4153040000 | 3.2331210000 | 1.2549430000 |
| H | -3.5148960000 | 4.7153360000 | 1.5740110000 |
| H | -4.8553680000 | 4.7170320000 | 0.4158800000 |
| H | -1.5369420000 | 5.1937160000 | -1.8664520000 |

**$^{3}$13**

Charge: 1
Multiplicity: 3
B3LYP-D3(BJ)/6-31G(d)-SDD SCF energy (au): -1690.96728770
B3LYP-D3(BJ)/6-31G(d)-SDD enthalpy (au): -1690.37816770
B3LYP-D3(BJ)/6-31G(d)-SDD free energy (au): -1690.49342170
B3LYP-D3(BJ)/def2-TZVP/SMD SCF energy (au): -2831.51638883
B3LYP-D3(BJ)/def2-TZVP/SMD enthalpy (au): -2830.92726883
B3LYP-D3(BJ)/def2-TZVP/SMD free energy (au): -2831.04252283
B3LYP-D3(BJ)/def2-TZVP/SMD free energy (quasi-harmonic) (au): -2831.02950413

Cartesian coordinates

| ATOM | X | Y | Z |
|---|---|---|---|
| Fe | -0.1226750000 | 0.1006100000 | -0.5524550000 |
| C | 1.2883620000 | -0.1837360000 | 3.6074020000 |
| N | 2.2064690000 | 0.6428840000 | 2.9840020000 |
| H | 2.9663030000 | 1.1326270000 | 3.4354290000 |
| C | 1.9020350000 | 0.6999790000 | 1.6644450000 |

| | | | |
|---|---|---|---|
| H | 2.4309510000 | 1.2738050000 | 0.9188370000 |
| N | 0.8411420000 | -0.0510010000 | 1.4128170000 |
| C | 0.4444060000 | -0.6047760000 | 2.6144590000 |
| H | -0.4125880000 | -1.2582580000 | 2.6756520000 |
| C | -3.4586050000 | -2.1023400000 | 0.9265000000 |
| N | -3.8662680000 | -0.7840320000 | 1.0193520000 |
| H | -4.7552120000 | -0.4617260000 | 1.3754340000 |
| C | -2.8729570000 | 0.0094470000 | 0.5571990000 |
| H | -2.8890090000 | 1.0872760000 | 0.5137100000 |
| N | -1.8487080000 | -0.7413110000 | 0.1706560000 |
| C | -2.1978630000 | -2.0606300000 | 0.3954910000 |
| H | -1.5282550000 | -2.8679580000 | 0.1482890000 |
| O | 1.3900650000 | 0.8023900000 | -1.2166780000 |
| H | 1.1154650000 | 1.5017410000 | -1.8433870000 |
| H | -4.0867170000 | -2.9235500000 | 1.2334500000 |
| H | 1.3226030000 | -0.3858750000 | 4.6664120000 |
| C | 1.0337480000 | -3.6654650000 | -2.2244030000 |
| N | 1.9129500000 | -3.5419440000 | -1.1640320000 |
| H | 2.6696040000 | -4.1738880000 | -0.9429340000 |
| C | 1.6090180000 | -2.4097260000 | -0.4876140000 |
| H | 2.1426060000 | -2.0486530000 | 0.3761930000 |
| N | 0.5752130000 | -1.8105830000 | -1.0571490000 |
| C | 0.2075300000 | -2.5775150000 | -2.1480170000 |
| H | -0.5931840000 | -2.2640840000 | -2.7969090000 |
| H | 1.0825410000 | -4.4884880000 | -2.9195250000 |
| O | -1.0927450000 | 0.0841530000 | -2.5585240000 |
| H | -0.7162600000 | 0.9386650000 | -2.9149500000 |
| H | -2.0287010000 | 0.2963940000 | -2.4115310000 |
| F | -0.8732400000 | 1.7195050000 | -0.1609790000 |
| C | -0.9814690000 | 3.3407200000 | -2.5730830000 |
| O | -0.1239730000 | 2.5287300000 | -2.9908460000 |
| N | -0.6732610000 | 4.5544730000 | -2.0917620000 |
| H | -2.0171060000 | 3.7903290000 | -0.1055050000 |
| C | 0.6761940000 | 5.1178280000 | -1.8301750000 |
| C | 0.4411900000 | 6.5779540000 | -1.4192980000 |
| C | 1.3414400000 | 4.3508300000 | -0.6758650000 |
| C | 1.5338500000 | 5.0661070000 | -3.1032020000 |
| H | -0.0451680000 | 7.1413710000 | -2.2236240000 |
| H | -0.1818620000 | 6.6412110000 | -0.5185330000 |
| H | 1.3966380000 | 7.0616920000 | -1.1988350000 |
| H | 1.4888800000 | 3.2989040000 | -0.9219870000 |
| H | 2.3170000000 | 4.7980660000 | -0.4551220000 |
| H | 0.7232130000 | 4.4002310000 | 0.2265020000 |
| H | 2.5007080000 | 5.5423870000 | -2.9101850000 |
| H | 1.7049280000 | 4.0372660000 | -3.4219200000 |
| H | 1.0423110000 | 5.6064050000 | -3.9190020000 |
| C | -2.4433130000 | 3.0225520000 | -2.6898250000 |
| C | -2.8406200000 | 2.4485700000 | -3.8986180000 |
| C | -3.4148390000 | 3.2927050000 | -1.6633810000 |
| C | -4.1803020000 | 2.1544040000 | -4.1614900000 |
| H | -2.0831360000 | 2.2529610000 | -4.6511840000 |
| C | -4.7752820000 | 2.9984860000 | -1.9766210000 |
| C | -5.1476090000 | 2.4439090000 | -3.1900650000 |

| | | | |
|---|---|---|---|
| H | -4.4663270000 | 1.7237130000 | -5.1154510000 |
| H | -5.5365190000 | 3.1991720000 | -1.2304480000 |
| H | -6.1947170000 | 2.2325130000 | -3.3862810000 |
| C | -3.0645090000 | 3.7677050000 | -0.3746870000 |
| C | -4.0621650000 | 4.1105150000 | 0.6874590000 |
| H | -4.5845940000 | 3.2196400000 | 1.0741480000 |
| H | -3.5727440000 | 4.5891110000 | 1.5396770000 |
| H | -4.8448750000 | 4.7885690000 | 0.3233320000 |
| H | -1.4580550000 | 5.1374270000 | -1.8331840000 |

**[5]14'**

Charge: 1

Multiplicity: 5

B3LYP-D3(BJ)/6-31G(d)-SDD SCF energy (au): -1693.88859309

B3LYP-D3(BJ)/6-31G(d)-SDD enthalpy (au): -1693.30724009

B3LYP-D3(BJ)/6-31G(d)-SDD free energy (au): -1693.42526509

B3LYP-D3(BJ)/def2-TZVP/SMD SCF energy (au): -2834.45284941

B3LYP-D3(BJ)/def2-TZVP/SMD enthalpy (au): -2833.87149641

B3LYP-D3(BJ)/def2-TZVP/SMD free energy (au): -2833.98952141

B3LYP-D3(BJ)/def2-TZVP/SMD free energy (quasi-harmonic) (au): -2833.97637932

Cartesian coordinates

| ATOM | X | Y | Z |
|---|---|---|---|
| Fe | 0.0051450000 | -0.0115060000 | -0.8166370000 |
| C | -2.2193880000 | -3.5365940000 | 0.4498370000 |
| N | -2.9276340000 | -2.5230320000 | 1.0692790000 |
| H | -3.6971170000 | -2.6451140000 | 1.7128150000 |
| C | -2.4001250000 | -1.3377960000 | 0.6795120000 |
| H | -2.7522850000 | -0.3745060000 | 1.0145290000 |
| N | -1.3932570000 | -1.5379060000 | -0.1575920000 |
| C | -1.2686400000 | -2.9077740000 | -0.3075740000 |
| H | -0.5037350000 | -3.3315550000 | -0.9400550000 |
| H | -2.4534650000 | -4.5780160000 | 0.6042540000 |
| C | 3.0089260000 | -2.9135960000 | -2.0683470000 |
| N | 3.3932070000 | -2.6285760000 | -0.7719480000 |
| H | 4.1793520000 | -3.0337410000 | -0.2831700000 |
| C | 2.5381390000 | -1.7039130000 | -0.2680970000 |
| H | 2.5872570000 | -1.2934390000 | 0.7291740000 |
| N | 1.6288280000 | -1.3847350000 | -1.1761780000 |
| C | 1.9092940000 | -2.1314620000 | -2.3059670000 |
| H | 1.3022190000 | -2.0432390000 | -3.1940800000 |
| H | 3.5413090000 | -3.6219720000 | -2.6832720000 |
| O | -0.8574530000 | -0.2712550000 | -2.7498470000 |
| H | -0.9159810000 | 0.5652480000 | -3.3066870000 |
| H | -1.7589820000 | -0.6186940000 | -2.6636850000 |
| F | -1.7389280000 | 1.4661110000 | -0.4674080000 |
| C | -1.8612520000 | 2.7657390000 | -4.2526910000 |
| O | -0.9455910000 | 1.9239410000 | -4.1900600000 |
| N | -1.7890740000 | 3.8644260000 | -5.0217880000 |
| H | -2.5613030000 | 4.5141590000 | -4.9543380000 |
| C | -0.6314490000 | 4.2736680000 | -5.8642090000 |
| C | -1.0429620000 | 5.5876730000 | -6.5388880000 |
| C | 0.5961150000 | 4.4959620000 | -4.9656560000 |

| | | | |
|---|---|---|---|
| C | -0.3567960000 | 3.1971350000 | -6.9246420000 |
| H | -1.9262110000 | 5.4483750000 | -7.1730270000 |
| H | -1.2601080000 | 6.3650600000 | -5.7964660000 |
| H | -0.2289750000 | 5.9504890000 | -7.1722370000 |
| H | 0.8607450000 | 3.5794900000 | -4.4337390000 |
| H | 1.4497990000 | 4.8069390000 | -5.5764700000 |
| H | 0.3976150000 | 5.2835940000 | -4.2306150000 |
| H | 0.4636630000 | 3.5223250000 | -7.5723370000 |
| H | -0.0786160000 | 2.2508510000 | -6.4573630000 |
| H | -1.2422350000 | 3.0365260000 | -7.5488250000 |
| C | -3.1410060000 | 2.6061630000 | -3.4783460000 |
| C | -4.3462620000 | 2.5168350000 | -4.1880200000 |
| C | -3.1432650000 | 2.5099060000 | -2.0748180000 |
| C | -5.5480230000 | 2.2986460000 | -3.5176600000 |
| H | -4.3320920000 | 2.5924590000 | -5.2714080000 |
| C | -4.3599710000 | 2.2987310000 | -1.4157680000 |
| C | -5.5528590000 | 2.1838050000 | -2.1263480000 |
| H | -6.4740840000 | 2.2165730000 | -4.0778910000 |
| H | -4.3689950000 | 2.2293550000 | -0.3322150000 |
| H | -6.4853210000 | 2.0157440000 | -1.5963840000 |
| C | -1.8829170000 | 2.6771170000 | -1.2675470000 |
| C | -1.8730880000 | 3.8376130000 | -0.2968560000 |
| H | -2.7239890000 | 3.7930000000 | 0.3897030000 |
| H | -0.9396030000 | 3.8282270000 | 0.2726090000 |
| H | -1.9351670000 | 4.7772680000 | -0.8562140000 |
| H | -0.9970880000 | 2.6888750000 | -1.8971520000 |
| C | 0.9857960000 | 1.6426650000 | 1.4443940000 |
| O | 1.2202660000 | 2.6189910000 | 0.6956540000 |
| O | 0.6450900000 | 0.4671040000 | 1.0315660000 |
| C | 1.0813820000 | 1.8135150000 | 2.9462210000 |
| H | 0.0745360000 | 1.7644430000 | 3.3770870000 |
| H | 1.6581180000 | 0.9926860000 | 3.3828590000 |
| H | 1.5350350000 | 2.7724770000 | 3.1997000000 |
| O | 1.0244640000 | 1.6974560000 | -1.6922530000 |
| H | 1.1999150000 | 2.1753300000 | -0.8042720000 |
| H | 1.8866510000 | 1.4522440000 | -2.0623800000 |

**$^{3}$14'**

Charge: 1
Multiplicity: 3
B3LYP-D3(BJ)/6-31G(d)-SDD SCF energy (au): -1693.86284553
B3LYP-D3(BJ)/6-31G(d)-SDD enthalpy (au): -1693.27937853
B3LYP-D3(BJ)/6-31G(d)-SDD free energy (au): -1693.39012353
B3LYP-D3(BJ)/def2-TZVP/SMD SCF energy (au): -2834.42976954
B3LYP-D3(BJ)/def2-TZVP/SMD enthalpy (au): -2833.84630254
B3LYP-D3(BJ)/def2-TZVP/SMD free energy (au): -2833.95704754
B3LYP-D3(BJ)/def2-TZVP/SMD free energy (quasi-harmonic) (au): -2833.94738301

Cartesian coordinates

| ATOM | X | Y | Z |
|---|---|---|---|
| Fe | 0.0454930000 | -0.3201330000 | -1.6373750000 |
| C | -2.8420940000 | -2.5568880000 | 0.3946550000 |
| N | -2.8621220000 | -1.3544810000 | 1.0757620000 |

| | | | |
|---|---|---|---|
| H | -3.4646070000 | -1.1228550000 | 1.8528940000 |
| C | -1.9213380000 | -0.5375120000 | 0.5445570000 |
| H | -1.7218530000 | 0.4763260000 | 0.8520050000 |
| N | -1.2985280000 | -1.1623840000 | -0.4477360000 |
| C | -1.8650190000 | -2.4235580000 | -0.5550060000 |
| H | -1.5193000000 | -3.1321640000 | -1.2923860000 |
| H | -3.4982410000 | -3.3748410000 | 0.6467870000 |
| C | 3.2953800000 | -3.0764710000 | -1.8443160000 |
| N | 2.6799770000 | -3.4964220000 | -0.6786170000 |
| H | 3.0087110000 | -4.2288470000 | -0.0650760000 |
| C | 1.5707620000 | -2.7438710000 | -0.4899540000 |
| H | 0.9013370000 | -2.8491570000 | 0.3496660000 |
| N | 1.4432390000 | -1.8660370000 | -1.4753690000 |
| C | 2.5165640000 | -2.0608230000 | -2.3269760000 |
| H | 2.6445760000 | -1.4421650000 | -3.2004330000 |
| H | 4.2055320000 | -3.5269970000 | -2.2071730000 |
| O | -1.1490930000 | -0.8051760000 | -3.2497240000 |
| H | -1.1481090000 | -0.0309370000 | -3.8801680000 |
| H | -2.0768500000 | -0.9317440000 | -2.9904470000 |
| F | -2.0449650000 | 2.7498050000 | 0.3554560000 |
| C | -1.6349890000 | 2.3867480000 | -3.9057510000 |
| O | -0.9198020000 | 1.5412960000 | -4.4973970000 |
| N | -1.4269460000 | 3.7045040000 | -3.9787550000 |
| H | -2.1506570000 | 4.2863920000 | -3.5791900000 |
| C | -0.3858720000 | 4.4231030000 | -4.7736270000 |
| C | -0.5524410000 | 5.9048210000 | -4.4126140000 |
| C | 1.0176800000 | 3.9453520000 | -4.3772530000 |
| C | -0.6496260000 | 4.1942150000 | -6.2690260000 |
| H | -1.5601560000 | 6.2646240000 | -4.6553140000 |
| H | -0.3609480000 | 6.0753780000 | -3.3471110000 |
| H | 0.1590440000 | 6.5069560000 | -4.9833910000 |
| H | 1.2080440000 | 2.9323800000 | -4.7362050000 |
| H | 1.7594320000 | 4.6047680000 | -4.8387710000 |
| H | 1.1579720000 | 3.9703630000 | -3.2932130000 |
| H | 0.0994430000 | 4.7286830000 | -6.8617670000 |
| H | -0.5876890000 | 3.1308750000 | -6.5131820000 |
| H | -1.6398570000 | 4.5656540000 | -6.5537860000 |
| C | -2.8052650000 | 1.9159860000 | -3.0888900000 |
| C | -3.6613160000 | 0.9969020000 | -3.7208480000 |
| C | -3.0229510000 | 2.2640920000 | -1.7432470000 |
| C | -4.7270130000 | 0.4245520000 | -3.0347450000 |
| H | -3.4787490000 | 0.7421980000 | -4.7606220000 |
| C | -4.0956190000 | 1.6685970000 | -1.0635180000 |
| C | -4.9392000000 | 0.7598370000 | -1.6940450000 |
| H | -5.3895060000 | -0.2706690000 | -3.5407520000 |
| H | -4.2495020000 | 1.9187740000 | -0.0197230000 |
| H | -5.7649940000 | 0.3171050000 | -1.1455290000 |
| C | -2.1494550000 | 3.2320920000 | -0.9699230000 |
| C | -2.7337300000 | 4.6348500000 | -0.8623580000 |
| H | -3.7284580000 | 4.5959330000 | -0.4085110000 |
| H | -2.0841080000 | 5.2598010000 | -0.2430840000 |
| H | -2.8317170000 | 5.1124880000 | -1.8423240000 |
| H | -1.1344200000 | 3.2571200000 | -1.3709980000 |

| | | | |
|---|---|---|---|
| C | 0.9691570000 | 2.1154460000 | -0.2452330000 |
| O | 1.4464400000 | 2.6177430000 | -1.2885120000 |
| O | 0.3219280000 | 0.9987040000 | -0.1999880000 |
| C | 1.1040820000 | 2.8439070000 | 1.0741330000 |
| H | 0.1082890000 | 3.1467340000 | 1.4118590000 |
| H | 1.5171520000 | 2.1691320000 | 1.8302010000 |
| H | 1.7436240000 | 3.7207900000 | 0.9661060000 |
| O | 1.2440510000 | 0.6596600000 | -2.9262520000 |
| H | 1.4513300000 | 1.4956910000 | -2.3778600000 |
| H | 0.7154450000 | 0.9902700000 | -3.6802460000 |

**[5]14**
Charge: 1
Multiplicity: 5
B3LYP-D3(BJ)/6-31G(d)-SDD SCF energy (au): -1691.01125007
B3LYP-D3(BJ)/6-31G(d)-SDD enthalpy (au): -1690.42029407
B3LYP-D3(BJ)/6-31G(d)-SDD free energy (au): -1690.53443707
B3LYP-D3(BJ)/def2-TZVP/SMD SCF energy (au): -2831.56803864
B3LYP-D3(BJ)/def2-TZVP/SMD enthalpy (au): -2830.97708264
B3LYP-D3(BJ)/def2-TZVP/SMD free energy (au): -2831.09122564
B3LYP-D3(BJ)/def2-TZVP/SMD free energy (quasi-harmonic) (au): -2831.08057757

Cartesian coordinates

| ATOM | X | Y | Z |
|---|---|---|---|
| Fe | -0.1607970000 | 0.2837470000 | -1.3770050000 |
| C | 0.7973150000 | 1.4659940000 | 2.8027270000 |
| N | 1.1949110000 | 2.5348920000 | 2.0210340000 |
| H | 1.5709500000 | 3.4094200000 | 2.3589400000 |
| C | 0.9870220000 | 2.2121910000 | 0.7193890000 |
| H | 1.1859340000 | 2.8465970000 | -0.1316020000 |
| N | 0.4787050000 | 0.9934650000 | 0.6276270000 |
| C | 0.3528390000 | 0.5169720000 | 1.9194050000 |
| H | -0.0474470000 | -0.4659770000 | 2.1192120000 |
| C | -3.5953420000 | -1.9362440000 | 0.1232020000 |
| N | -3.7308270000 | -0.7307070000 | 0.7878210000 |
| H | -4.4647250000 | -0.4925740000 | 1.4399190000 |
| C | -2.7161100000 | 0.0835690000 | 0.4089390000 |
| H | -2.5870440000 | 1.1017280000 | 0.7385090000 |
| N | -1.9375660000 | -0.5383440000 | -0.4633700000 |
| C | -2.4739830000 | -1.8016490000 | -0.6500520000 |
| H | -2.0058340000 | -2.5166570000 | -1.3096530000 |
| O | 0.3657200000 | 1.9398190000 | -2.1173980000 |
| H | 0.5832370000 | 1.8823660000 | -3.0588310000 |
| H | -4.2828020000 | -2.7544780000 | 0.2683250000 |
| H | 0.8693200000 | 1.4813440000 | 3.8789430000 |
| C | 2.3279150000 | -3.1932160000 | -2.0314180000 |
| N | 2.8803500000 | -2.6754650000 | -0.8735150000 |
| H | 3.6888190000 | -3.0382190000 | -0.3881340000 |
| C | 2.1607600000 | -1.5870230000 | -0.5103240000 |
| H | 2.3691470000 | -0.9791040000 | 0.3565430000 |
| N | 1.1764730000 | -1.3821910000 | -1.3735340000 |
| C | 1.2700830000 | -2.3784110000 | -2.3305160000 |
| H | 0.5741710000 | -2.4131140000 | -3.1536340000 |

| | | | |
|---|---|---|---|
| H | 2.7330810000 | -4.0650120000 | -2.5203150000 |
| O | -1.0713750000 | -0.6104330000 | -3.2694520000 |
| H | -0.9496570000 | 0.1020850000 | -3.9556490000 |
| H | -2.0285010000 | -0.5939410000 | -3.1051270000 |
| F | -2.0980730000 | 3.0829650000 | -0.1501190000 |
| C | -1.6552430000 | 2.4456680000 | -4.3368390000 |
| O | -0.9247790000 | 1.5745760000 | -4.8470130000 |
| N | -1.4695360000 | 3.7642760000 | -4.5217170000 |
| H | -2.2119290000 | 4.3690790000 | -4.1993480000 |
| C | -0.4048300000 | 4.4092440000 | -5.3352400000 |
| C | -0.5633360000 | 5.9173910000 | -5.1035600000 |
| C | 0.9799780000 | 3.9496260000 | -4.8550350000 |
| C | -0.6138820000 | 4.0645120000 | -6.8176460000 |
| H | -1.5573760000 | 6.2652150000 | -5.4113770000 |
| H | -0.4118210000 | 6.1723660000 | -4.0482850000 |
| H | 0.1757930000 | 6.4652640000 | -5.6941700000 |
| H | 1.1492420000 | 2.8996990000 | -5.1013670000 |
| H | 1.7513060000 | 4.5468620000 | -5.3517930000 |
| H | 1.0797130000 | 4.0783400000 | -3.7729480000 |
| H | 0.1615580000 | 4.5419410000 | -7.4256940000 |
| H | -0.5580950000 | 2.9840030000 | -6.9695330000 |
| H | -1.5895480000 | 4.4202610000 | -7.1660230000 |
| C | -2.8492450000 | 2.0585690000 | -3.5079180000 |
| C | -3.7002100000 | 1.0941470000 | -4.0719250000 |
| C | -3.1206740000 | 2.5690160000 | -2.2253270000 |
| C | -4.8282020000 | 0.6499560000 | -3.3886280000 |
| H | -3.4647080000 | 0.7051950000 | -5.0579490000 |
| C | -4.2672430000 | 2.1181030000 | -1.5568900000 |
| C | -5.1157210000 | 1.1728690000 | -2.1251270000 |
| H | -5.4838500000 | -0.0870790000 | -3.8418240000 |
| H | -4.4753360000 | 2.5052280000 | -0.5646950000 |
| H | -5.9981830000 | 0.8415990000 | -1.5858440000 |
| C | -2.1988860000 | 3.5289350000 | -1.5011280000 |
| C | -2.7101470000 | 4.9597630000 | -1.4378010000 |
| H | -3.7083140000 | 4.9974320000 | -0.9900690000 |
| H | -2.0309110000 | 5.5735120000 | -0.8386010000 |
| H | -2.7740270000 | 5.4037080000 | -2.4366540000 |
| H | -1.1877230000 | 3.4519030000 | -1.9015800000 |

**$^{3}$14**

Charge: 1
Multiplicity: 3
B3LYP-D3(BJ)/6-31G(d)-SDD SCF energy (au): -1690.98300497
B3LYP-D3(BJ)/6-31G(d)-SDD enthalpy (au): -1690.39092497
B3LYP-D3(BJ)/6-31G(d)-SDD free energy (au): -1690.50190697
B3LYP-D3(BJ)/def2-TZVP/SMD SCF energy (au): -2831.54336111
B3LYP-D3(BJ)/def2-TZVP/SMD enthalpy (au): -2830.95128111
B3LYP-D3(BJ)/def2-TZVP/SMD free energy (au): -2831.06226311
B3LYP-D3(BJ)/def2-TZVP/SMD free energy (quasi-harmonic) (au): -2831.05217414

Cartesian coordinates

| ATOM | X | Y | Z |
|---|---|---|---|
| Fe | -0.3246680000 | 0.3578660000 | -1.2683070000 |

| | | | |
|---|---|---|---|
| C | 0.7671980000 | 1.5300930000 | 2.6064100000 |
| N | 1.2239090000 | 2.5710460000 | 1.8191550000 |
| H | 1.6988450000 | 3.3994190000 | 2.1479550000 |
| C | 0.9274370000 | 2.2965690000 | 0.5259780000 |
| H | 1.1596750000 | 2.9050080000 | -0.3342450000 |
| N | 0.3004290000 | 1.1315350000 | 0.4476640000 |
| C | 0.1925850000 | 0.6408220000 | 1.7379640000 |
| H | -0.2824710000 | -0.3060780000 | 1.9407260000 |
| C | -3.3045630000 | -2.2594950000 | 0.2755910000 |
| N | -3.7155270000 | -1.0367230000 | 0.7753310000 |
| H | -4.5347960000 | -0.8737080000 | 1.3434400000 |
| C | -2.8346480000 | -0.0901350000 | 0.3699960000 |
| H | -2.9069840000 | 0.9626830000 | 0.5915510000 |
| N | -1.8823130000 | -0.6466420000 | -0.3663420000 |
| C | -2.1627790000 | -2.0010780000 | -0.4339370000 |
| H | -1.5143240000 | -2.6756760000 | -0.9711380000 |
| O | 0.4115320000 | 1.8521820000 | -2.1223180000 |
| H | 0.5374870000 | 1.7072080000 | -3.0711280000 |
| H | -3.8444320000 | -3.1721580000 | 0.4724190000 |
| H | 0.8900470000 | 1.5196060000 | 3.6778060000 |
| C | 2.2821850000 | -3.0939360000 | -2.0135910000 |
| N | 2.8484710000 | -2.5614160000 | -0.8702440000 |
| H | 3.6833780000 | -2.8958200000 | -0.4102100000 |
| C | 2.0974040000 | -1.4978790000 | -0.4856750000 |
| H | 2.3135090000 | -0.8862300000 | 0.3772020000 |
| N | 1.0827510000 | -1.3252210000 | -1.3161470000 |
| C | 1.1873610000 | -2.3141980000 | -2.2770010000 |
| H | 0.4766820000 | -2.3696820000 | -3.0863280000 |
| H | 2.7049330000 | -3.9481690000 | -2.5187020000 |
| O | -1.0745490000 | -0.4182160000 | -3.0466390000 |
| H | -0.9729480000 | 0.2436460000 | -3.7896090000 |
| H | -2.0355930000 | -0.4967850000 | -2.9322890000 |
| F | -2.2149460000 | 2.9663730000 | -0.1431940000 |
| C | -1.6651880000 | 2.4709920000 | -4.3319350000 |
| O | -0.9293900000 | 1.5836220000 | -4.8080490000 |
| N | -1.4497530000 | 3.7843740000 | -4.5072690000 |
| H | -2.1975550000 | 4.4000000000 | -4.2185220000 |
| C | -0.3428030000 | 4.4137340000 | -5.2772080000 |
| C | -0.4687300000 | 5.9212620000 | -5.0221990000 |
| C | 1.0151520000 | 3.9083070000 | -4.7673870000 |
| C | -0.5200140000 | 4.0996450000 | -6.7704780000 |
| H | -1.4460500000 | 6.3006140000 | -5.3462640000 |
| H | -0.3350160000 | 6.1537070000 | -3.9593900000 |
| H | 0.2976390000 | 6.4597450000 | -5.5860600000 |
| H | 1.1783720000 | 2.8682420000 | -5.0543310000 |
| H | 1.8123320000 | 4.5158050000 | -5.2078430000 |
| H | 1.0759340000 | 3.9802910000 | -3.6776020000 |
| H | 0.2871940000 | 4.5618570000 | -7.3481380000 |
| H | -0.4918080000 | 3.0202400000 | -6.9386050000 |
| H | -1.4737210000 | 4.4904550000 | -7.1415470000 |
| C | -2.8989160000 | 2.1022000000 | -3.5552030000 |
| C | -3.7652580000 | 1.1869190000 | -4.1737050000 |
| C | -3.1869770000 | 2.5805370000 | -2.2644250000 |

| | | | |
|---|---|---|---|
| C | -4.9331760000 | 0.7719490000 | -3.5402780000 |
| H | -3.5108490000 | 0.8136210000 | -5.1609960000 |
| C | -4.3724530000 | 2.1601040000 | -1.6457850000 |
| C | -5.2417850000 | 1.2714420000 | -2.2721710000 |
| H | -5.6022520000 | 0.0750280000 | -4.0352820000 |
| H | -4.5975490000 | 2.5257160000 | -0.6490560000 |
| H | -6.1565720000 | 0.9651620000 | -1.7733680000 |
| C | -2.2377940000 | 3.4632330000 | -1.4810100000 |
| C | -2.6629500000 | 4.9196820000 | -1.3869150000 |
| H | -3.6747440000 | 5.0050970000 | -0.9781680000 |
| H | -1.9729100000 | 5.4694610000 | -0.7398280000 |
| H | -2.6547460000 | 5.3971970000 | -2.3718400000 |
| H | -1.2174360000 | 3.3405170000 | -1.8499450000 |

**F□**

Charge: 0

Multiplicity: 2

B3LYP-D3(BJ)/6-31G(d)-SDD SCF energy (au): -99.7145779299

B3LYP-D3(BJ)/6-31G(d)-SDD enthalpy (au): -99.7122179299

B3LYP-D3(BJ)/6-31G(d)-SDD free energy (au): -99.7293909299

B3LYP-D3(BJ)/def2-TZVP/SMD SCF energy (au): -99.7699421456

B3LYP-D3(BJ)/def2-TZVP/SMD enthalpy (au): -99.7675821456

B3LYP-D3(BJ)/def2-TZVP/SMD free energy (au): -99.7847551456

B3LYP-D3(BJ)/def2-TZVP/SMD free energy (quasi-harmonic) (au): -99.7847552157

Cartesian coordinates

| ATOM | X | Y | Z |
|---|---|---|---|
| F | -0.0987430000 | 0.1885100000 | 0.0000000000 |

**$^{5}Im_2AcOWat_2$_FeF_remove-F**

Charge: 1

Multiplicity: 5

B3LYP-D3(BJ)/6-31G(d)-SDD SCF energy (au): -957.706375357

B3LYP-D3(BJ)/6-31G(d)-SDD enthalpy (au): -957.435398357

B3LYP-D3(BJ)/6-31G(d)-SDD free energy (au): -957.513740357

B3LYP-D3(BJ)/def2-TZVP/SMD SCF energy (au): -2097.97855970

B3LYP-D3(BJ)/def2-TZVP/SMD enthalpy (au): -2097.70758270

B3LYP-D3(BJ)/def2-TZVP/SMD free energy (au): -2097.78592470

B3LYP-D3(BJ)/def2-TZVP/SMD free energy (quasi-harmonic) (au): -2097.778447343

Cartesian coordinates

| ATOM | X | Y | Z |
|---|---|---|---|
| Fe | -0.2809980000 | -1.2035720000 | -0.4138570000 |
| C | -0.7585230000 | -1.4340070000 | 3.8928220000 |
| N | 0.2520610000 | -0.5067370000 | 3.7238310000 |
| H | 0.7527390000 | -0.0340860000 | 4.4642850000 |
| C | 0.4608910000 | -0.3357460000 | 2.3988680000 |
| H | 1.2003750000 | 0.3202500000 | 1.9652210000 |
| N | -0.3674220000 | -1.1097500000 | 1.7064310000 |
| C | -1.1332190000 | -1.8022700000 | 2.6295460000 |
| H | -1.8856980000 | -2.5121010000 | 2.3210440000 |
| H | -1.1048390000 | -1.7375940000 | 4.8680780000 |
| C | 0.7917530000 | -5.2893520000 | -1.3199320000 |

| | | | |
|---|---|---|---|
| N | 2.0163500000 | -4.6533700000 | -1.2773900000 |
| H | 2.9159930000 | -5.0879990000 | -1.4326010000 |
| C | 1.8106710000 | -3.3455610000 | -0.9969080000 |
| H | 2.5923570000 | -2.6065570000 | -0.9052130000 |
| N | 0.5116280000 | -3.1110240000 | -0.8579180000 |
| C | -0.1365130000 | -4.3182670000 | -1.0578350000 |
| H | -1.2111870000 | -4.4010470000 | -0.9961810000 |
| H | 0.6962950000 | -6.3435000000 | -1.5268630000 |
| O | -2.4448340000 | -1.1088610000 | -0.3920100000 |
| H | -2.5825700000 | -0.7951300000 | -1.3087550000 |
| H | -2.9132390000 | -0.5063540000 | 0.2090970000 |
| C | 1.6666340000 | 0.7127780000 | -1.5390390000 |
| O | 1.0238240000 | 0.8100530000 | -2.6008410000 |
| O | 1.3299570000 | -0.0607950000 | -0.5446560000 |
| C | 2.9195130000 | 1.5326430000 | -1.3314760000 |
| H | 2.7418750000 | 2.2698270000 | -0.5401410000 |
| H | 3.7395650000 | 0.8896670000 | -0.9968980000 |
| H | 3.1957020000 | 2.0506970000 | -2.2501560000 |
| O | -0.9064690000 | -0.8450350000 | -2.4849470000 |
| H | -0.1892750000 | -0.1270500000 | -2.6722630000 |
| H | -0.7357590000 | -1.5748630000 | -3.1019030000 |

**$^{6}$Im$_2$AcOWat$_2$_FeF**
Charge: 1
Multiplicity: 6
B3LYP-D3(BJ)/6-31G(d)-SDD SCF energy (au): -1057.56353877
B3LYP-D3(BJ)/6-31G(d)-SDD enthalpy (au): -1057.28793377
B3LYP-D3(BJ)/6-31G(d)-SDD free energy (au): -1057.36405277
B3LYP-D3(BJ)/def2-TZVP/SMD SCF energy (au): -2197.88044536
B3LYP-D3(BJ)/def2-TZVP/SMD enthalpy (au): -2197.60484036
B3LYP-D3(BJ)/def2-TZVP/SMD free energy (au): -2197.68095936
B3LYP-D3(BJ)/def2-TZVP/SMD free energy (quasi-harmonic) (au): -2197.67730759

Cartesian coordinates

| ATOM | X | Y | Z |
|---|---|---|---|
| Fe | -0.3190570000 | 0.2412120000 | -0.5632920000 |
| C | -3.6684990000 | -2.0299570000 | 0.8986390000 |
| N | -4.2245340000 | -0.8077070000 | 0.5650270000 |
| H | -5.1882350000 | -0.5353070000 | 0.7069260000 |
| C | -3.2643710000 | -0.0287510000 | 0.0281360000 |
| H | -3.4001070000 | 0.9877020000 | -0.3071010000 |
| N | -2.1147260000 | -0.6977390000 | -0.0058960000 |
| C | -2.3524440000 | -1.9474980000 | 0.5401050000 |
| H | -1.5679730000 | -2.6806270000 | 0.6344520000 |
| H | -4.2459530000 | -2.8214910000 | 1.3492190000 |
| C | 1.0998000000 | -3.7481230000 | -1.5065920000 |
| N | 2.1300920000 | -3.2700390000 | -0.7179830000 |
| H | 2.9766510000 | -3.7696320000 | -0.4792840000 |
| C | 1.8298300000 | -2.0132080000 | -0.3284840000 |
| H | 2.4682130000 | -1.3872690000 | 0.2782130000 |
| N | 0.6499040000 | -1.6588440000 | -0.8300780000 |
| C | 0.1845980000 | -2.7346190000 | -1.5716490000 |
| H | -0.7598170000 | -2.6908830000 | -2.0902090000 |

```
H      1.1147010000   -4.7366520000   -1.9375890000
O     -0.6940040000   -0.0928780000   -2.7405880000
H      0.1765710000   -0.0360940000   -3.1736250000
H     -1.1546600000    0.7390980000   -2.9559110000
F     -1.2058640000    1.7880880000   -0.9451810000
C      1.5094970000    0.7042900000    1.7251490000
O      2.5289490000    0.7746760000    1.0117640000
O      0.3170370000    0.4776340000    1.2424770000
C      1.5791500000    0.8690340000    3.2201570000
H      0.9792460000    1.7370360000    3.5139070000
H      1.1408830000   -0.0057780000    3.7111250000
H      2.6119970000    1.0068230000    3.5408130000
O      1.4835330000    1.1164630000   -1.3247840000
H      2.0797330000    1.0160990000   -0.5037240000
H      1.3144140000    2.0720510000   -1.4127240000
```

**$^{5}Im_2AcOWatN_3$_11**

```
Charge: 0
Multiplicity: 5
B3LYP-D3(BJ)/6-31G(d)-SDD SCF energy (au):              -1781.73725776
B3LYP-D3(BJ)/6-31G(d)-SDD enthalpy (au):                -1781.16923076
B3LYP-D3(BJ)/6-31G(d)-SDD free energy (au):             -1781.29071876
B3LYP-D3(BJ)/def2-TZVP/SMD SCF energy (au):             -2922.29646504
B3LYP-D3(BJ)/def2-TZVP/SMD enthalpy (au):               -2921.72843804
B3LYP-D3(BJ)/def2-TZVP/SMD free energy (au):            -2921.84992604
B3LYP-D3(BJ)/def2-TZVP/SMD free energy (quasi-harmonic) (au): -2921.83704328

Cartesian coordinates
ATOM    X               Y               Z
Fe     0.6756350000   -1.2910560000   -1.4525910000
C      1.7152430000    1.7022870000    1.5824320000
N      2.3579000000    2.2332060000    0.4752900000
H      2.9805660000    3.0272790000    0.4700380000
C      2.0784470000    1.4411480000   -0.5925320000
H      2.4478750000    1.6116080000   -1.5907900000
N      1.2835940000    0.4503790000   -0.2301840000
C      1.0509840000    0.5962920000    1.1225550000
H      0.4062900000   -0.0965540000    1.6396630000
H      1.7898050000    2.1543850000    2.5590230000
C      1.5790280000   -4.6032210000    1.2469260000
N      2.8026930000   -3.9561120000    1.2260140000
H      3.6482360000   -4.2628270000    1.6832880000
C      2.6805740000   -2.8449120000    0.4520070000
H      3.4926190000   -2.1635730000    0.2340930000
N      1.4476450000   -2.7526830000   -0.0162330000
C      0.7488970000   -3.8411510000    0.4689380000
H     -0.2863860000   -3.9973690000    0.2039560000
H      1.4238230000   -5.5199790000    1.7935840000
O     -0.2268420000   -2.8497240000   -2.6773420000
H     -0.4501540000   -2.3906840000   -3.5019590000
H     -1.0956250000   -2.9474530000   -2.1644450000
C     -4.5664240000    0.2305080000   -2.0786470000
C     -3.2862780000    0.2869340000   -2.6155320000
```

| | | | |
|---|---|---|---|
| C | -2.3160180000 | 1.1326600000 | -2.0579460000 |
| C | -2.5867920000 | 1.8700780000 | -0.8895060000 |
| C | -3.8853410000 | 1.7955930000 | -0.3676440000 |
| C | -4.8705090000 | 1.0078560000 | -0.9584130000 |
| H | -5.3145930000 | -0.4201980000 | -2.5197280000 |
| H | -3.0226120000 | -0.3158920000 | -3.4785650000 |
| H | -4.1148870000 | 2.3472550000 | 0.5407850000 |
| H | -5.8658360000 | 0.9724070000 | -0.5248320000 |
| C | -1.5301070000 | 2.6318620000 | -0.1148960000 |
| C | -1.7181360000 | 4.1543970000 | -0.1048340000 |
| H | -0.5375400000 | 2.3913150000 | -0.4930950000 |
| H | -1.5552750000 | 2.2614060000 | 0.9168520000 |
| H | -0.9969880000 | 4.6227360000 | 0.5742210000 |
| H | -1.5592790000 | 4.5760410000 | -1.1019490000 |
| H | -2.7240660000 | 4.4377570000 | 0.2251740000 |
| C | -0.9968860000 | 1.1567840000 | -2.7413740000 |
| O | -0.3501120000 | 0.1468710000 | -2.9871230000 |
| N | -0.5060710000 | 2.4427320000 | -3.0449400000 |
| C | -1.1226610000 | 3.2942370000 | -4.1289880000 |
| C | -2.5996690000 | 3.5760840000 | -3.8228530000 |
| C | -0.9911520000 | 2.5679340000 | -5.4768030000 |
| H | -2.9513350000 | 4.3059580000 | -4.5592110000 |
| H | -3.2232070000 | 2.6853530000 | -3.9064940000 |
| H | -2.7384320000 | 4.0008230000 | -2.8272230000 |
| H | 0.0512890000 | 2.3224240000 | -5.6886900000 |
| H | -1.5718610000 | 1.6392070000 | -5.4759490000 |
| H | -1.3718370000 | 3.2066010000 | -6.2802870000 |
| F | 0.8565790000 | 2.2865510000 | -3.4221590000 |
| C | -0.3457100000 | 4.6180860000 | -4.1316490000 |
| H | -0.4252920000 | 5.1113350000 | -3.1576480000 |
| H | 0.7101550000 | 4.4575330000 | -4.3555080000 |
| H | -0.7653190000 | 5.2834580000 | -4.8924920000 |
| C | -2.0920580000 | -1.9266290000 | -0.3074930000 |
| O | -2.3054120000 | -2.8740850000 | -1.1030390000 |
| O | -1.0701250000 | -1.1574730000 | -0.3190400000 |
| C | -3.0949430000 | -1.6732730000 | 0.8089660000 |
| H | -4.0548500000 | -2.1419780000 | 0.5829930000 |
| H | -3.2239530000 | -0.6003310000 | 0.9663290000 |
| H | -2.7025950000 | -2.1051770000 | 1.7382050000 |
| N | 2.4307990000 | -1.1357920000 | -2.5236190000 |
| N | 3.4623450000 | -0.8991260000 | -1.9543990000 |
| N | 4.4615730000 | -0.6578890000 | -1.4022410000 |

**$^{5}$Im$_{2}$AcOWatN$_{3}$_13**

Charge: 0
Multiplicity: 5
B3LYP-D3(BJ)/6-31G(d)-SDD SCF energy (au): -1781.80719873
B3LYP-D3(BJ)/6-31G(d)-SDD enthalpy (au): -1781.24039373
B3LYP-D3(BJ)/6-31G(d)-SDD free energy (au): -1781.35941773
B3LYP-D3(BJ)/def2-TZVP/SMD SCF energy (au): -2922.36776995
B3LYP-D3(BJ)/def2-TZVP/SMD enthalpy (au): -2921.80096495
B3LYP-D3(BJ)/def2-TZVP/SMD free energy (au): -2921.91998895
B3LYP-D3(BJ)/def2-TZVP/SMD free energy (quasi-harmonic) (au): -2921.90842522

Cartesian coordinates

| ATOM | X | Y | Z |
|---|---|---|---|
| H | 35.7177770000 | 28.1774830000 | 39.2753610000 |
| C | 35.2410310000 | 28.7404770000 | 38.4886380000 |
| C | 35.3300980000 | 28.6769800000 | 37.1242000000 |
| N | 34.3419480000 | 29.7594130000 | 38.7515440000 |
| C | 33.9219390000 | 30.2688630000 | 37.5677010000 |
| N | 34.5038000000 | 29.6320580000 | 36.5665950000 |
| H | 34.0333990000 | 30.0642150000 | 39.6626890000 |
| H | 35.9265880000 | 28.0267310000 | 36.5041140000 |
| H | 33.2221460000 | 31.0815910000 | 37.4615360000 |
| H | 37.1577050000 | 25.8546840000 | 33.0025250000 |
| C | 36.3474030000 | 26.5049160000 | 33.2924100000 |
| C | 36.3134060000 | 27.8141990000 | 33.6921820000 |
| N | 35.0295320000 | 26.0770990000 | 33.3071430000 |
| C | 34.2521740000 | 27.1149150000 | 33.7062700000 |
| N | 35.0056380000 | 28.1715400000 | 33.9492000000 |
| H | 34.6984380000 | 25.1580670000 | 33.0528890000 |
| H | 37.1023820000 | 28.5391650000 | 33.8129490000 |
| H | 33.1770160000 | 27.0936870000 | 33.8006120000 |
| Fe | 34.2191720000 | 30.1581100000 | 34.3990440000 |
| O | 33.6691210000 | 31.9942750000 | 35.2199970000 |
| H | 34.5693230000 | 32.3288940000 | 35.5494710000 |
| H | 33.3473600000 | 32.6185210000 | 34.5111980000 |
| F | 34.1264680000 | 30.5672830000 | 32.6184590000 |
| N | 33.8382080000 | 33.4331710000 | 31.0541420000 |
| O | 32.9634870000 | 33.5545430000 | 33.1512380000 |
| C | 35.3176790000 | 33.8952600000 | 32.9152830000 |
| C | 35.3589710000 | 34.8982330000 | 33.8821790000 |
| C | 36.5689170000 | 35.3697970000 | 34.3912050000 |
| C | 37.7677070000 | 34.8208310000 | 33.9211850000 |
| C | 37.7505580000 | 33.7995450000 | 32.9862790000 |
| C | 36.5322250000 | 33.2714970000 | 32.4673650000 |
| C | 33.9500950000 | 33.5768750000 | 32.3890320000 |
| C | 32.5939260000 | 33.1468170000 | 30.3044820000 |
| C | 31.9312630000 | 31.8598550000 | 30.8200140000 |
| C | 31.6364130000 | 34.3427380000 | 30.4357070000 |
| C | 33.0192500000 | 32.9664250000 | 28.8413760000 |
| C | 36.5307440000 | 32.1293600000 | 31.6316030000 |
| C | 37.7658630000 | 31.4024210000 | 31.2085340000 |
| H | 34.4179190000 | 35.3177420000 | 34.2226940000 |
| H | 36.5769570000 | 36.1540560000 | 35.1413810000 |
| H | 38.7184210000 | 35.1822750000 | 34.3040470000 |
| H | 38.6881860000 | 33.3568930000 | 32.6681720000 |
| H | 34.6971490000 | 33.4832600000 | 30.5238110000 |
| H | 35.5878540000 | 31.6210660000 | 31.4766690000 |
| H | 38.1581570000 | 30.7926770000 | 32.0391560000 |
| H | 37.5481050000 | 30.7152520000 | 30.3846940000 |
| H | 38.5744730000 | 32.0708080000 | 30.8881380000 |
| H | 31.0538220000 | 31.6340630000 | 30.2025520000 |
| H | 32.6290760000 | 31.0211470000 | 30.7809310000 |
| H | 31.6157440000 | 31.9750370000 | 31.8573580000 |

| | | | |
|---|---|---|---|
| H | 31.3743430000 | 34.5005440000 | 31.4839740000 |
| H | 32.1010250000 | 35.2559540000 | 30.0466830000 |
| H | 30.7185030000 | 34.1525930000 | 29.8681490000 |
| H | 33.7058880000 | 32.1180150000 | 28.7360750000 |
| H | 32.1417650000 | 32.7706780000 | 28.2180400000 |
| H | 33.5125080000 | 33.8678640000 | 28.4563830000 |
| N | 32.3683800000 | 29.3772920000 | 34.5473980000 |
| N | 31.7316100000 | 29.3115600000 | 35.5733290000 |
| N | 31.1022940000 | 29.2372160000 | 36.5412160000 |
| C | 36.7320270000 | 31.5440150000 | 35.3594000000 |
| O | 36.1045010000 | 32.4136080000 | 35.9975700000 |
| O | 36.2067910000 | 30.6675780000 | 34.5749280000 |
| C | 38.2406450000 | 31.4478640000 | 35.5185510000 |
| H | 38.7161070000 | 31.2728020000 | 34.5496440000 |
| H | 38.4732970000 | 30.5924790000 | 36.1651510000 |
| H | 38.6373100000 | 32.3576040000 | 35.9718570000 |

**$^5Im_2AcOWatN_3$_FeF_remove-F**
Charge: 0
Multiplicity: 5
B3LYP-D3(BJ)/6-31G(d)-SDD SCF energy (au): -1045.67205188
B3LYP-D3(BJ)/6-31G(d)-SDD enthalpy (au): -1045.41223888
B3LYP-D3(BJ)/6-31G(d)-SDD free energy (au): -1045.49315488
B3LYP-D3(BJ)/def2-TZVP/SMD SCF energy (au): -2185.93257811
B3LYP-D3(BJ)/def2-TZVP/SMD enthalpy (au): -2185.67276511
B3LYP-D3(BJ)/def2-TZVP/SMD free energy (au): -2185.75368111
B3LYP-D3(BJ)/def2-TZVP/SMD free energy (quasi-harmonic) (au): -2185.74648723

Cartesian coordinates

| ATOM | X | Y | Z |
|---|---|---|---|
| Fe | 0.4551210000 | -0.8536570000 | -0.8867460000 |
| C | 1.0781460000 | 1.6595090000 | 2.6632990000 |
| N | 2.1001530000 | 2.1481830000 | 1.8690370000 |
| H | 2.7222240000 | 2.9099870000 | 2.0966060000 |
| C | 2.0955950000 | 1.4577330000 | 0.6984240000 |
| H | 2.7748050000 | 1.6552300000 | -0.1201810000 |
| N | 1.1283840000 | 0.5520860000 | 0.7078420000 |
| C | 0.4845530000 | 0.6690420000 | 1.9274300000 |
| H | -0.3693840000 | 0.0487860000 | 2.1497860000 |
| H | 0.8719760000 | 2.0586410000 | 3.6439120000 |
| C | 2.6344130000 | -4.4803100000 | -0.0372150000 |
| N | 3.0900070000 | -3.6703290000 | 0.9900560000 |
| H | 3.7577060000 | -3.9307620000 | 1.7008770000 |
| C | 2.4861290000 | -2.4615550000 | 0.8713190000 |
| H | 2.6500060000 | -1.6291280000 | 1.5383050000 |
| N | 1.6735480000 | -2.4586860000 | -0.1715280000 |
| C | 1.7542640000 | -3.7118000000 | -0.7496960000 |
| H | 1.1678020000 | -3.9400550000 | -1.6259790000 |
| H | 2.9731200000 | -5.4971980000 | -0.1570440000 |
| O | -0.3167420000 | -2.2106700000 | -2.4898620000 |
| H | -0.1196110000 | -1.6249460000 | -3.2404270000 |
| H | -1.2936220000 | -2.0905940000 | -2.2714700000 |
| C | -2.4658650000 | -1.2846190000 | -0.3748670000 |

O     -2.7002200000   -1.7605480000   -1.5008070000
O     -1.3004080000   -0.9356370000    0.0671130000
C     -3.6078520000   -1.0715920000    0.6058110000
H     -3.6676990000   -0.0109580000    0.8734800000
H     -3.4095330000   -1.6285770000    1.5281560000
H     -4.5550840000   -1.3976780000    0.1733810000
N      1.3398670000    0.0974900000   -2.4294420000
N      2.2883090000    0.8410230000   -2.4062850000
N      3.1996490000    1.5605970000   -2.4046810000

**$^{5}$Im$_2$AcOWatN$_3$_FeF_remove-N$_3$**
Charge: 0
Multiplicity: 5
B3LYP-D3(BJ)/6-31G(d)-SDD SCF energy (au):               -981.331328730
B3LYP-D3(BJ)/6-31G(d)-SDD enthalpy (au):                 -981.085036730
B3LYP-D3(BJ)/6-31G(d)-SDD free energy (au):              -981.161505730
B3LYP-D3(BJ)/def2-TZVP/SMD SCF energy (au):              -2121.57381139
B3LYP-D3(BJ)/def2-TZVP/SMD enthalpy (au):                -2121.32751939
B3LYP-D3(BJ)/def2-TZVP/SMD free energy (au):             -2121.40398839
B3LYP-D3(BJ)/def2-TZVP/SMD free energy (quasi-harmonic) (au): -2121.397489660

Cartesian coordinates
ATOM    X               Y               Z
Fe     0.2661520000   -0.6255950000   -0.8586760000
C      1.1482660000    1.8995050000    2.6817740000
N      1.8371070000    2.5061720000    1.6452630000
H      2.3600120000    3.3675290000    1.7021380000
C      1.6547640000    1.7590240000    0.5237640000
H      2.0441030000    1.9861130000   -0.4574480000
N      0.8965630000    0.7115310000    0.7930920000
C      0.5681900000    0.7860750000    2.1308460000
H     -0.0801420000    0.0493700000    2.5804540000
H      1.1307430000    2.3124900000    3.6782620000
C      2.5348640000   -4.2719110000   -0.3003770000
N      2.9504350000   -3.5456940000    0.8037690000
H      3.6044350000   -3.8553980000    1.5073060000
C      2.3231340000   -2.3418860000    0.7734080000
H      2.4534540000   -1.5674940000    1.5142160000
N      1.5358670000   -2.2638950000   -0.2841220000
C      1.6555180000   -3.4599160000   -0.9649140000
H      1.0937400000   -3.6227930000   -1.8716220000
H      2.8971310000   -5.2682480000   -0.4988120000
O     -0.4681930000   -1.8563860000   -2.5843760000
H     -0.3967880000   -1.1801020000   -3.2774340000
H     -1.4336940000   -1.8665210000   -2.2825210000
F      1.0862720000    0.5352480000   -2.0543760000
C     -2.5598460000   -1.4343290000   -0.2474570000
O     -2.8006000000   -1.8135050000   -1.4124870000
O     -1.4323880000   -0.9798640000    0.1799430000
C     -3.6667280000   -1.4923630000    0.7956920000
H     -3.8511920000   -0.4873830000    1.1908300000
H     -3.3463770000   -2.1168660000    1.6369400000
H     -4.5853000000   -1.8948580000    0.3652310000

**$^{6}Im_2AcOWatN_3$_FeF**

Charge: 0
Multiplicity: 6
B3LYP-D3(BJ)/6-31G(d)-SDD SCF energy (au): -1145.52769527
B3LYP-D3(BJ)/6-31G(d)-SDD enthalpy (au): -1145.26437227
B3LYP-D3(BJ)/6-31G(d)-SDD free energy (au): -1145.34602827
B3LYP-D3(BJ)/def2-TZVP/SMD SCF energy (au): -2285.83596984
B3LYP-D3(BJ)/def2-TZVP/SMD enthalpy (au): -2285.57264684
B3LYP-D3(BJ)/def2-TZVP/SMD free energy (au): -2285.65430284
B3LYP-D3(BJ)/def2-TZVP/SMD free energy (quasi-harmonic) (au): -2285.64799357

Cartesian coordinates

| ATOM | X | Y | Z |
|---|---|---|---|
| Fe | 0.3240480000 | -0.3175630000 | -1.2057030000 |
| C | 0.9471070000 | 2.2135530000 | 2.2682710000 |
| N | 2.2490880000 | 2.1354900000 | 1.8022130000 |
| H | 3.0489690000 | 2.6221750000 | 2.1797230000 |
| C | 2.2689060000 | 1.3051440000 | 0.7316470000 |
| H | 3.1637320000 | 1.0586560000 | 0.1791870000 |
| N | 1.0494420000 | 0.8459650000 | 0.4972120000 |
| C | 0.2134450000 | 1.4037830000 | 1.4456210000 |
| H | -0.8373440000 | 1.1653350000 | 1.4477940000 |
| H | 0.6758870000 | 2.8257700000 | 3.1135600000 |
| C | 2.6110180000 | -3.9833720000 | -0.3540250000 |
| N | 2.0701840000 | -3.7032350000 | 0.8901530000 |
| H | 2.1873810000 | -4.2553250000 | 1.7269090000 |
| C | 1.3376080000 | -2.5667240000 | 0.7852450000 |
| H | 0.7740680000 | -2.1183590000 | 1.5880830000 |
| N | 1.3884870000 | -2.1092270000 | -0.4528310000 |
| C | 2.1767880000 | -2.9800110000 | -1.1764510000 |
| H | 2.3580670000 | -2.8187730000 | -2.2270510000 |
| H | 3.2362350000 | -4.8445490000 | -0.5277680000 |
| O | -0.4375060000 | -1.6972080000 | -2.6380730000 |
| H | -0.8440110000 | -1.0906080000 | -3.2814410000 |
| H | -1.2149970000 | -2.0775070000 | -2.0856970000 |
| F | -0.7683900000 | 0.9493080000 | -1.9133890000 |
| C | -2.0868230000 | -1.7052670000 | 0.0057120000 |
| O | -2.2911700000 | -2.4017000000 | -1.0110140000 |
| O | -1.0831580000 | -0.9196420000 | 0.1872490000 |
| C | -3.0893080000 | -1.7487710000 | 1.1472470000 |
| H | -3.6497970000 | -0.8067190000 | 1.1619460000 |
| H | -2.5707270000 | -1.8354460000 | 2.1072520000 |
| H | -3.7877290000 | -2.5771240000 | 1.0167230000 |
| N | 1.9767910000 | 0.0913920000 | -2.2654230000 |
| N | 3.1201730000 | -0.0933020000 | -1.9330190000 |
| N | 4.2274780000 | -0.2431430000 | -1.6195090000 |

**$^{5}Im_2AcOWatN_3$_TS-1**

Charge: 0
Multiplicity: 5
B3LYP-D3(BJ)/6-31G(d)-SDD SCF energy (au): -1781.71572522
B3LYP-D3(BJ)/6-31G(d)-SDD enthalpy (au): -1781.15005022

B3LYP-D3(BJ)/6-31G(d)-SDD free energy (au): -1781.27044122
B3LYP-D3(BJ)/def2-TZVP/SMD SCF energy (au): -2922.26972268
B3LYP-D3(BJ)/def2-TZVP/SMD enthalpy (au): -2921.70404768
B3LYP-D3(BJ)/def2-TZVP/SMD free energy (au): -2921.82443868
B3LYP-D3(BJ)/def2-TZVP/SMD free energy (quasi-harmonic) (au): -2921.81141646

Cartesian coordinates

| ATOM | X | Y | Z |
|---|---|---|---|
| Fe | 0.4281060000 | -0.4922090000 | -1.0939980000 |
| C | 0.7199030000 | 2.3160760000 | 2.3004230000 |
| N | 2.0313520000 | 2.3860150000 | 1.8620290000 |
| H | 2.7365460000 | 3.0308760000 | 2.1868510000 |
| C | 2.2064990000 | 1.4351810000 | 0.9056820000 |
| H | 3.1460660000 | 1.2552500000 | 0.4017380000 |
| N | 1.0825680000 | 0.7631830000 | 0.7162100000 |
| C | 0.1468290000 | 1.3029900000 | 1.5785440000 |
| H | -0.8589130000 | 0.9153140000 | 1.5875480000 |
| H | 0.3330490000 | 2.9772400000 | 3.0598770000 |
| C | 2.2985640000 | -4.2657680000 | -0.4736200000 |
| N | 2.7166430000 | -3.5818960000 | 0.6562360000 |
| H | 3.3731260000 | -3.9169060000 | 1.3454200000 |
| C | 2.1435680000 | -2.3542140000 | 0.6427140000 |
| H | 2.3013770000 | -1.5934610000 | 1.3901760000 |
| N | 1.3741080000 | -2.2282100000 | -0.4245990000 |
| C | 1.4623620000 | -3.4096630000 | -1.1355440000 |
| H | 0.9276100000 | -3.5314030000 | -2.0638810000 |
| H | 2.6278290000 | -5.2694090000 | -0.6904910000 |
| O | -0.4471110000 | -1.5862660000 | -2.7482940000 |
| H | -0.9298660000 | -0.8990360000 | -3.2515580000 |
| H | -1.1636820000 | -2.0576610000 | -2.2208140000 |
| C | -6.0150800000 | 2.5094870000 | -3.6650090000 |
| C | -4.7320980000 | 2.0013540000 | -3.8280830000 |
| C | -3.8340270000 | 1.9751780000 | -2.7496800000 |
| C | -4.2308840000 | 2.4066570000 | -1.4651240000 |
| C | -5.5362580000 | 2.8944790000 | -1.3251140000 |
| C | -6.4142440000 | 2.9644470000 | -2.4053140000 |
| H | -6.7010700000 | 2.5430210000 | -4.5059460000 |
| H | -4.4016800000 | 1.6237820000 | -4.7908890000 |
| H | -5.8707350000 | 3.2121760000 | -0.3404240000 |
| H | -7.4170820000 | 3.3567700000 | -2.2609990000 |
| C | -3.3474200000 | 2.2992620000 | -0.2374390000 |
| C | -2.7139910000 | 3.6382760000 | 0.1682770000 |
| H | -2.5608440000 | 1.5563960000 | -0.3850350000 |
| H | -3.9709640000 | 1.9348870000 | 0.5903710000 |
| H | -2.0879500000 | 3.5094000000 | 1.0580480000 |
| H | -2.0796420000 | 4.0095130000 | -0.6399620000 |
| H | -3.4790030000 | 4.3921630000 | 0.3885100000 |
| C | -2.4700860000 | 1.4394520000 | -3.0175240000 |
| O | -2.2782350000 | 0.3764290000 | -3.5920470000 |
| N | -1.4771000000 | 2.3190110000 | -2.5329240000 |
| C | -0.5773450000 | 2.9804140000 | -3.4928610000 |
| C | -1.4799010000 | 4.0085790000 | -4.2204400000 |
| C | 0.0798160000 | 2.0308550000 | -4.5072760000 |

| | | | |
|---|---|---|---|
| H | -0.8480180000 | 4.6263310000 | -4.8667260000 |
| H | -2.2300580000 | 3.5144980000 | -4.8454970000 |
| H | -1.9944290000 | 4.6584970000 | -3.5066370000 |
| H | 0.7080900000 | 1.3000000000 | -3.9933450000 |
| H | -0.6609180000 | 1.5017340000 | -5.1112230000 |
| H | 0.7123580000 | 2.6219720000 | -5.1790880000 |
| F | -0.3865000000 | 1.1791150000 | -1.7161720000 |
| C | 0.4944460000 | 3.7144850000 | -2.6753790000 |
| H | 0.0299220000 | 4.3645180000 | -1.9270480000 |
| H | 1.1352240000 | 2.9914920000 | -2.1671660000 |
| H | 1.1097580000 | 4.3282910000 | -3.3419080000 |
| C | -2.1471150000 | -1.6954060000 | -0.1196340000 |
| O | -2.2106160000 | -2.5264180000 | -1.0450830000 |
| O | -1.2396810000 | -0.7884780000 | 0.0198520000 |
| C | -3.2169430000 | -1.6893010000 | 0.9630050000 |
| H | -3.7491350000 | -0.7315180000 | 0.9388210000 |
| H | -2.7500110000 | -1.7775920000 | 1.9498250000 |
| H | -3.9257470000 | -2.5050340000 | 0.8110850000 |
| N | 2.1135510000 | 0.0305990000 | -2.1243560000 |
| N | 3.1783200000 | -0.1682810000 | -1.5979450000 |
| N | 4.2106020000 | -0.3355850000 | -1.0876140000 |

**$^5Im_2AcOWatN_3$_TS-3-F**

Charge: 0
Multiplicity: 5
B3LYP-D3(BJ)/6-31G(d)-SDD SCF energy (au): -1781.78835167
B3LYP-D3(BJ)/6-31G(d)-SDD enthalpy (au): -1781.22199867
B3LYP-D3(BJ)/6-31G(d)-SDD free energy (au): -1781.33894667
B3LYP-D3(BJ)/def2-TZVP/SMD SCF energy (au): -2922.35248229
B3LYP-D3(BJ)/def2-TZVP/SMD enthalpy (au): -2921.78612929
B3LYP-D3(BJ)/def2-TZVP/SMD free energy (au): -2921.90307729
B3LYP-D3(BJ)/def2-TZVP/SMD free energy (quasi-harmonic) (au): -2921.89227862

Cartesian coordinates

| ATOM | X | Y | Z |
|---|---|---|---|
| H | 34.4670740000 | 27.0762500000 | 39.1972890000 |
| C | 34.3514060000 | 27.7949540000 | 38.4013440000 |
| C | 35.1034860000 | 28.0738000000 | 37.2917910000 |
| N | 33.2910030000 | 28.6836950000 | 38.3586070000 |
| C | 33.4266630000 | 29.4503690000 | 37.2477790000 |
| N | 34.5173860000 | 29.1049810000 | 36.5852470000 |
| H | 32.5201100000 | 28.7267020000 | 39.0081280000 |
| H | 36.0198600000 | 27.6172140000 | 36.9517780000 |
| H | 32.7422900000 | 30.2258250000 | 36.9448330000 |
| H | 39.0250400000 | 26.3455830000 | 34.2889080000 |
| C | 38.0776920000 | 26.8578320000 | 34.2263600000 |
| C | 37.6600650000 | 28.0874090000 | 34.6653820000 |
| N | 36.9739040000 | 26.2910460000 | 33.6100020000 |
| C | 35.9440470000 | 27.1746660000 | 33.6930920000 |
| N | 36.3349740000 | 28.2626790000 | 34.3275930000 |
| H | 36.9376790000 | 25.3831640000 | 33.1707150000 |
| H | 38.1896680000 | 28.8658670000 | 35.1920910000 |
| H | 34.9502180000 | 27.0242870000 | 33.2976160000 |

| | | | |
|---|---|---|---|
| Fe | 35.0426130000 | 30.0260210000 | 34.6185950000 |
| O | 33.7911650000 | 31.5887530000 | 35.2665650000 |
| H | 34.4387640000 | 32.1117270000 | 35.8415190000 |
| H | 33.4676210000 | 32.1485620000 | 34.5182810000 |
| F | 35.7334360000 | 30.6313610000 | 32.9161350000 |
| N | 34.2307490000 | 33.1200090000 | 31.0549950000 |
| O | 33.0649040000 | 32.8766300000 | 32.9941080000 |
| C | 35.2519600000 | 33.7908260000 | 33.1613420000 |
| C | 34.9469440000 | 34.8179920000 | 34.0556770000 |
| C | 35.9196300000 | 35.3546470000 | 34.8955380000 |
| C | 37.2126880000 | 34.8233300000 | 34.8742890000 |
| C | 37.5195890000 | 33.7716680000 | 34.0255040000 |
| C | 36.5620950000 | 33.2302550000 | 33.1339340000 |
| C | 34.0976630000 | 33.2160170000 | 32.3948490000 |
| C | 33.3300910000 | 32.3623950000 | 30.1503730000 |
| C | 33.2509560000 | 30.8990470000 | 30.6146330000 |
| C | 31.9389940000 | 33.0144160000 | 30.1319850000 |
| C | 33.9680730000 | 32.4419790000 | 28.7576800000 |
| C | 36.9005750000 | 32.1021080000 | 32.3074690000 |
| C | 38.1950840000 | 31.3742720000 | 32.4525150000 |
| H | 33.9257440000 | 35.1840550000 | 34.0937530000 |
| H | 35.6663370000 | 36.1588920000 | 35.5787830000 |
| H | 37.9748330000 | 35.2171020000 | 35.5401330000 |
| H | 38.5185180000 | 33.3512990000 | 34.0418640000 |
| H | 35.0699510000 | 33.5119680000 | 30.6522160000 |
| H | 36.3895280000 | 31.9932390000 | 31.3629560000 |
| H | 38.3695470000 | 31.1229970000 | 33.5023800000 |
| H | 38.1679190000 | 30.4463550000 | 31.8773530000 |
| H | 39.0400660000 | 31.9789460000 | 32.0887580000 |
| H | 32.6230890000 | 30.3276850000 | 29.9214250000 |
| H | 34.2468890000 | 30.4480330000 | 30.6457890000 |
| H | 32.8296370000 | 30.8174790000 | 31.6174790000 |
| H | 31.4972270000 | 32.9902960000 | 31.1294430000 |
| H | 32.0049560000 | 34.0567980000 | 29.7998620000 |
| H | 31.2835270000 | 32.4723900000 | 29.4412790000 |
| H | 34.9629750000 | 31.9794680000 | 28.7520670000 |
| H | 33.3475220000 | 31.9076550000 | 28.0325010000 |
| H | 34.0612670000 | 33.4820720000 | 28.4210110000 |
| N | 33.4733780000 | 29.0232350000 | 33.7512370000 |
| N | 32.4615910000 | 28.7725150000 | 34.3632450000 |
| N | 31.4838860000 | 28.5165080000 | 34.9313940000 |
| C | 36.6868370000 | 31.7974990000 | 36.4581160000 |
| O | 35.7028340000 | 32.5022570000 | 36.7687140000 |
| O | 36.7010680000 | 30.8691050000 | 35.5705780000 |
| C | 38.0110870000 | 32.0265210000 | 37.1754010000 |
| H | 38.8157300000 | 32.1501680000 | 36.4430120000 |
| H | 38.2539070000 | 31.1449750000 | 37.7802380000 |
| H | 37.9569840000 | 32.9048020000 | 37.8214430000 |

**$^5$Im$_2$AcOWatN$_3$_TS-3-N$_3$**
Charge: 0
Multiplicity: 5
B3LYP-D3(BJ)/6-31G(d)-SDD SCF energy (au): -1781.78462992

B3LYP-D3(BJ)/6-31G(d)-SDD enthalpy (au): -1781.21879692
B3LYP-D3(BJ)/6-31G(d)-SDD free energy (au): -1781.33891292
B3LYP-D3(BJ)/def2-TZVP/SMD SCF energy (au): -2922.35361664
B3LYP-D3(BJ)/def2-TZVP/SMD enthalpy (au): -2921.78778364
B3LYP-D3(BJ)/def2-TZVP/SMD free energy (au): -2921.90789964
B3LYP-D3(BJ)/def2-TZVP/SMD free energy (quasi-harmonic) (au): -2921.89429772

Cartesian coordinates

| ATOM | X | Y | Z |
|---|---|---|---|
| H | 32.2000520000 | 26.7204960000 | 36.1669550000 |
| C | 32.7631780000 | 27.6235580000 | 35.9917100000 |
| C | 34.1052570000 | 27.8657290000 | 35.8660500000 |
| N | 32.1531370000 | 28.8556330000 | 35.8398180000 |
| C | 33.1226840000 | 29.7840430000 | 35.6318870000 |
| N | 34.3140620000 | 29.2116120000 | 35.6442890000 |
| H | 31.1609680000 | 29.0373880000 | 35.8649940000 |
| H | 34.9323740000 | 27.1761720000 | 35.9207990000 |
| H | 32.9527090000 | 30.8388310000 | 35.4866160000 |
| H | 38.9340250000 | 25.9722320000 | 36.6753400000 |
| C | 38.3869520000 | 26.6814230000 | 36.0742410000 |
| C | 37.7175840000 | 27.8353110000 | 36.3864020000 |
| N | 38.2574200000 | 26.5362040000 | 34.7022920000 |
| C | 37.5289100000 | 27.5853700000 | 34.2378730000 |
| N | 37.1869350000 | 28.3782820000 | 35.2343680000 |
| H | 38.6495180000 | 25.7939360000 | 34.1422210000 |
| H | 37.5808690000 | 28.3436190000 | 37.3277790000 |
| H | 37.2828580000 | 27.7465900000 | 33.1999230000 |
| Fe | 36.2018280000 | 30.3625620000 | 35.0657860000 |
| O | 34.9298460000 | 32.0009700000 | 35.3829000000 |
| H | 35.4441280000 | 32.5058600000 | 36.1029340000 |
| H | 34.5745480000 | 32.5859540000 | 34.6802830000 |
| F | 37.7501330000 | 31.0857880000 | 34.3933370000 |
| N | 33.4313060000 | 32.3506840000 | 31.1671620000 |
| O | 33.4171980000 | 33.0042950000 | 33.3498050000 |
| C | 35.4990470000 | 33.1133410000 | 32.2123060000 |
| C | 35.8814370000 | 34.3343610000 | 32.7716700000 |
| C | 37.2202110000 | 34.7140120000 | 32.8275200000 |
| C | 38.2004600000 | 33.8382090000 | 32.3505320000 |
| C | 37.8403710000 | 32.6057130000 | 31.8325840000 |
| C | 36.4868070000 | 32.2039320000 | 31.7278990000 |
| C | 34.0359060000 | 32.8077700000 | 32.2903980000 |
| C | 31.9940360000 | 32.0201080000 | 31.0262790000 |
| C | 31.5889210000 | 30.9643360000 | 32.0675500000 |
| C | 31.1522750000 | 33.2966960000 | 31.1879970000 |
| C | 31.8280600000 | 31.4434230000 | 29.6146460000 |
| C | 36.1593600000 | 30.8879690000 | 31.2629420000 |
| C | 37.2010330000 | 29.9166030000 | 30.8015710000 |
| H | 35.1097510000 | 34.9842640000 | 33.1717790000 |
| H | 37.4969660000 | 35.6716100000 | 33.2572190000 |
| H | 39.2501710000 | 34.1090320000 | 32.4120370000 |
| H | 38.6172240000 | 31.9213010000 | 31.5151700000 |
| H | 33.9755980000 | 32.3927630000 | 30.3165410000 |
| H | 35.1531980000 | 30.7118180000 | 30.9074330000 |

| | | | |
|---|---|---|---|
| H | 37.9854360000 | 29.8002790000 | 31.5556060000 |
| H | 36.7496840000 | 28.9380410000 | 30.6116530000 |
| H | 37.6762300000 | 30.2489380000 | 29.8659570000 |
| H | 30.5361920000 | 30.6971670000 | 31.9204890000 |
| H | 32.1938880000 | 30.0584970000 | 31.9656440000 |
| H | 31.7154680000 | 31.3566380000 | 33.0775810000 |
| H | 31.3106540000 | 33.7289570000 | 32.1785470000 |
| H | 31.4280290000 | 34.0404610000 | 30.4318650000 |
| H | 30.0876720000 | 33.0655190000 | 31.0702740000 |
| H | 32.4255020000 | 30.5328350000 | 29.4914080000 |
| H | 30.7795760000 | 31.1878880000 | 29.4352990000 |
| H | 32.1304210000 | 32.1702660000 | 28.8498990000 |
| N | 35.6146810000 | 29.8424150000 | 33.0837200000 |
| N | 34.7536450000 | 29.0556070000 | 32.7532730000 |
| N | 33.9545200000 | 28.3043720000 | 32.3824020000 |
| C | 36.8650570000 | 31.6697300000 | 37.6769580000 |
| O | 36.3924330000 | 32.7762650000 | 37.3332960000 |
| O | 36.6742530000 | 30.5510650000 | 37.0723270000 |
| C | 37.7526840000 | 31.5968530000 | 38.9111210000 |
| H | 38.7717090000 | 31.3384260000 | 38.6007220000 |
| H | 37.4021320000 | 30.8026340000 | 39.5786970000 |
| H | 37.7654220000 | 32.5518950000 | 39.4397320000 |

**$^{5}Im_{2}AcOWatOH\_11$**

Charge: 0
Multiplicity: 5
B3LYP-D3(BJ)/6-31G(d)-SDD SCF energy (au): -1693.35290801
B3LYP-D3(BJ)/6-31G(d)-SDD enthalpy (au): -1692.78523101
B3LYP-D3(BJ)/6-31G(d)-SDD free energy (au): -1692.90166501
B3LYP-D3(BJ)/def2-TZVP/SMD SCF energy (au): -2833.88358650
B3LYP-D3(BJ)/def2-TZVP/SMD enthalpy (au): -2833.31590950
B3LYP-D3(BJ)/def2-TZVP/SMD free energy (au): -2833.43234350
B3LYP-D3(BJ)/def2-TZVP/SMD free energy (quasi-harmonic) (au): -2833.41974149

Cartesian coordinates

| ATOM | X | Y | Z |
|---|---|---|---|
| Fe | 0.1870350000 | -0.4717890000 | -0.4695470000 |
| C | 0.7393360000 | 2.3855580000 | 2.8844300000 |
| N | 1.3476160000 | 3.0191740000 | 1.8130080000 |
| H | 1.7226240000 | 3.9560070000 | 1.8111260000 |
| C | 1.3208300000 | 2.1688510000 | 0.7506710000 |
| H | 1.6806920000 | 2.3861300000 | -0.2440970000 |
| N | 0.7365720000 | 1.0343030000 | 1.0906000000 |
| C | 0.3649520000 | 1.1534770000 | 2.4141800000 |
| H | -0.1676780000 | 0.3527130000 | 2.9032020000 |
| O | 0.6841640000 | 0.8638960000 | -1.7509460000 |
| H | 0.6341580000 | 0.5236660000 | -2.6562560000 |
| H | 0.6296160000 | 2.8631270000 | 3.8455950000 |
| C | 3.1158260000 | -3.6564710000 | -0.1689210000 |
| N | 3.5389300000 | -2.8284010000 | 0.8583540000 |
| H | 4.3230090000 | -2.9946830000 | 1.4715100000 |
| C | 2.7088100000 | -1.7535900000 | 0.8973280000 |
| H | 2.7953630000 | -0.9403260000 | 1.6020600000 |

| | | | |
|---|---|---|---|
| N | 1.7883970000 | -1.8530510000 | -0.0432940000 |
| C | 2.0265390000 | -3.0344580000 | -0.7180310000 |
| H | 1.3846850000 | -3.3259970000 | -1.5352040000 |
| H | 3.6185150000 | -4.5822390000 | -0.4005830000 |
| O | -0.5959120000 | -1.9297030000 | -1.9552590000 |
| H | -0.9970150000 | -1.3890950000 | -2.6671330000 |
| H | -1.3762780000 | -2.2430730000 | -1.4058240000 |
| C | -5.2175530000 | 3.0862390000 | -4.1180830000 |
| C | -4.0213930000 | 2.4285740000 | -4.4041680000 |
| C | -3.2946190000 | 1.8064960000 | -3.3852300000 |
| C | -3.7608050000 | 1.8042260000 | -2.0528180000 |
| C | -4.9521120000 | 2.4873050000 | -1.7887470000 |
| C | -5.6747230000 | 3.1207990000 | -2.8027090000 |
| H | -5.7819620000 | 3.5634790000 | -4.9136520000 |
| H | -3.6510300000 | 2.3823220000 | -5.4233270000 |
| H | -5.3269250000 | 2.5242880000 | -0.7715460000 |
| H | -6.6001680000 | 3.6362420000 | -2.5605710000 |
| C | -2.9846280000 | 1.0910760000 | -0.9636060000 |
| C | -3.2372090000 | 1.5825940000 | 0.4632920000 |
| H | -1.9139880000 | 1.1785240000 | -1.1761390000 |
| H | -3.1990490000 | 0.0171110000 | -1.0264410000 |
| H | -2.5512160000 | 1.0676430000 | 1.1403900000 |
| H | -3.0654150000 | 2.6624900000 | 0.5498940000 |
| H | -4.2602940000 | 1.3762800000 | 0.7987730000 |
| C | -2.0629950000 | 1.0319010000 | -3.7370520000 |
| O | -2.0158540000 | -0.1849900000 | -3.5953150000 |
| N | -1.0535140000 | 1.7511170000 | -4.3434320000 |
| C | -0.5336890000 | 3.1465010000 | -4.0578860000 |
| C | -0.8269320000 | 3.5637120000 | -2.6078640000 |
| C | -1.1778510000 | 4.1123400000 | -5.0627870000 |
| H | -0.3314120000 | 4.5231880000 | -2.4193070000 |
| H | -1.8940560000 | 3.6993920000 | -2.4247940000 |
| H | -0.4181110000 | 2.8060520000 | -1.9328210000 |
| H | -1.0008370000 | 3.7787970000 | -6.0905230000 |
| H | -2.2543030000 | 4.1980810000 | -4.9007800000 |
| H | -0.7370180000 | 5.1077370000 | -4.9408930000 |
| F | -0.0105780000 | 0.8667380000 | -4.6815170000 |
| C | 0.9913420000 | 3.1366250000 | -4.2566120000 |
| H | 1.4701260000 | 2.4855810000 | -3.5226090000 |
| H | 1.2717970000 | 2.8238000000 | -5.2648630000 |
| H | 1.3458910000 | 4.1608480000 | -4.1035160000 |
| C | -2.1883440000 | -1.8788880000 | 0.8093170000 |
| O | -2.5103110000 | -2.5044850000 | -0.2206940000 |
| O | -1.1918790000 | -1.0703460000 | 0.9201160000 |
| C | -3.0379100000 | -2.0276740000 | 2.0651080000 |
| H | -3.7770780000 | -2.8213390000 | 1.9419540000 |
| H | -3.5537290000 | -1.0810820000 | 2.2648110000 |
| H | -2.3991850000 | -2.2387070000 | 2.9290680000 |

**$^5$Im$_2$AcOWatOH_13**

Charge: 0

Multiplicity: 5

B3LYP-D3(BJ)/6-31G(d)-SDD SCF energy (au): -1693.41879127

B3LYP-D3(BJ)/6-31G(d)-SDD enthalpy (au): -1692.85339927
B3LYP-D3(BJ)/6-31G(d)-SDD free energy (au): -1692.96934927
B3LYP-D3(BJ)/def2-TZVP/SMD SCF energy (au): -2833.95216894
B3LYP-D3(BJ)/def2-TZVP/SMD enthalpy (au): -2833.38677694
B3LYP-D3(BJ)/def2-TZVP/SMD free energy (au): -2833.50272694
B3LYP-D3(BJ)/def2-TZVP/SMD free energy (quasi-harmonic) (au): -2833.49074067

Cartesian coordinates

| ATOM | X | Y | Z |
|---|---|---|---|
| H | 35.1091280000 | 29.9996210000 | 39.1938020000 |
| C | 34.8646290000 | 29.9573800000 | 38.1440420000 |
| C | 35.6374920000 | 29.8440270000 | 37.0191920000 |
| N | 33.5563860000 | 30.0315640000 | 37.6888600000 |
| C | 33.5761010000 | 29.9657950000 | 36.3329160000 |
| N | 34.8208350000 | 29.8447610000 | 35.9084280000 |
| H | 32.7334710000 | 30.1512060000 | 38.2606740000 |
| H | 36.7082040000 | 29.7830100000 | 36.9040910000 |
| H | 32.7204850000 | 30.0243170000 | 35.6765320000 |
| H | 36.5845240000 | 25.0391510000 | 32.3821530000 |
| C | 36.2284100000 | 25.8860480000 | 32.9471440000 |
| C | 36.2092330000 | 27.2291630000 | 32.6830350000 |
| N | 35.6508130000 | 25.7451520000 | 34.1983870000 |
| C | 35.3070490000 | 26.9826740000 | 34.6422270000 |
| N | 35.6349470000 | 27.8922920000 | 33.7468040000 |
| H | 35.5059950000 | 24.8779350000 | 34.6933540000 |
| H | 36.5491620000 | 27.7828740000 | 31.8237600000 |
| H | 34.8359450000 | 27.1804200000 | 35.5919460000 |
| Fe | 35.3007610000 | 30.0702200000 | 33.7553090000 |
| O | 35.2905800000 | 32.1704800000 | 34.1019390000 |
| H | 36.2693190000 | 32.3966440000 | 33.9316940000 |
| H | 34.7529530000 | 32.5732010000 | 33.4007570000 |
| F | 35.9354740000 | 30.0892120000 | 32.0113270000 |
| N | 34.6806350000 | 30.9696650000 | 29.1003800000 |
| O | 33.1380220000 | 30.4218060000 | 30.6813360000 |
| C | 34.2523730000 | 32.5264870000 | 30.9110710000 |
| C | 33.1272490000 | 33.1155280000 | 31.4892590000 |
| C | 33.1821470000 | 34.3821110000 | 32.0746920000 |
| C | 34.3981470000 | 35.0798340000 | 32.0793750000 |
| C | 35.5366150000 | 34.5036210000 | 31.5413280000 |
| C | 35.5259950000 | 33.1972200000 | 30.9664840000 |
| C | 33.9998320000 | 31.2011200000 | 30.2489900000 |
| C | 34.6789080000 | 29.7023930000 | 28.3422890000 |
| C | 35.1410110000 | 28.5479780000 | 29.2473200000 |
| C | 33.2726110000 | 29.4288020000 | 27.7845560000 |
| C | 35.6725120000 | 29.8946050000 | 27.1891680000 |
| C | 36.7419600000 | 32.5768020000 | 30.5856080000 |
| C | 38.0845740000 | 33.2248530000 | 30.6702960000 |
| H | 32.1989370000 | 32.5551540000 | 31.4682440000 |
| H | 32.2917960000 | 34.8189080000 | 32.5160000000 |
| H | 34.4580940000 | 36.0664590000 | 32.5305120000 |
| H | 36.4802990000 | 35.0320710000 | 31.6110980000 |
| H | 35.3599080000 | 31.6666370000 | 28.8288680000 |
| H | 36.7262510000 | 31.5049410000 | 30.4423090000 |

| | | | |
|---|---|---|---|
| H | 38.4443840000 | 33.1831430000 | 31.7104390000 |
| H | 38.8110080000 | 32.6885470000 | 30.0500960000 |
| H | 38.0794340000 | 34.2744480000 | 30.3535910000 |
| H | 35.1607480000 | 27.6124830000 | 28.6757340000 |
| H | 36.1434250000 | 28.7503630000 | 29.6354390000 |
| H | 34.4672450000 | 28.4353280000 | 30.0976620000 |
| H | 32.5567470000 | 29.3305590000 | 28.6022270000 |
| H | 32.9525800000 | 30.2504890000 | 27.1341730000 |
| H | 33.2751280000 | 28.5033100000 | 27.1971630000 |
| H | 36.6815490000 | 30.0965900000 | 27.5690720000 |
| H | 35.7178440000 | 28.9890060000 | 26.5765880000 |
| H | 35.3698660000 | 30.7269660000 | 26.5424210000 |
| C | 38.1284630000 | 31.1046800000 | 34.1405030000 |
| O | 37.8432120000 | 32.2811490000 | 33.8052720000 |
| O | 37.2863500000 | 30.2043040000 | 34.4873530000 |
| C | 39.5880420000 | 30.6757390000 | 34.1198970000 |
| H | 39.7629240000 | 30.0840670000 | 33.2130190000 |
| H | 39.8110310000 | 30.0343190000 | 34.9773810000 |
| H | 40.2508870000 | 31.5436520000 | 34.1091530000 |
| O | 33.4637580000 | 30.0271650000 | 33.4838140000 |
| H | 33.2488910000 | 30.0956730000 | 32.5283980000 |

**$^5Im_2AcOWatOH\_FeF\_remove\text{-}F$**

Charge: 0
Multiplicity: 5
B3LYP-D3(BJ)/6-31G(d)-SDD SCF energy (au): -957.283078025
B3LYP-D3(BJ)/6-31G(d)-SDD enthalpy (au): -957.025412025
B3LYP-D3(BJ)/6-31G(d)-SDD free energy (au): -957.101774025
B3LYP-D3(BJ)/def2-TZVP/SMD SCF energy (au): -2097.51413418
B3LYP-D3(BJ)/def2-TZVP/SMD enthalpy (au): -2097.25646818
B3LYP-D3(BJ)/def2-TZVP/SMD free energy (au): -2097.33283018
B3LYP-D3(BJ)/def2-TZVP/SMD free energy (quasi-harmonic) (au): -2097.327514155

Cartesian coordinates

| ATOM | X | Y | Z |
|---|---|---|---|
| Fe | 0.2859980000 | -0.8585110000 | 1.2009830000 |
| C | -2.8767280000 | -1.4591000000 | -1.7529960000 |
| N | -2.0110160000 | -0.6738630000 | -2.4968770000 |
| H | -2.1645910000 | -0.3473360000 | -3.4391270000 |
| C | -0.9293710000 | -0.3933670000 | -1.7200970000 |
| H | -0.0957160000 | 0.2225710000 | -2.0198490000 |
| N | -1.0591610000 | -0.9598120000 | -0.5362230000 |
| C | -2.2685460000 | -1.6254040000 | -0.5367570000 |
| H | -2.5966530000 | -2.1406740000 | 0.3532130000 |
| H | -3.8172800000 | -1.8073220000 | -2.1502360000 |
| C | 2.1987700000 | -4.6078250000 | 0.0518360000 |
| N | 3.2886200000 | -3.7582850000 | -0.0289250000 |
| H | 4.2277350000 | -4.0176360000 | -0.2923790000 |
| C | 2.8783480000 | -2.5089180000 | 0.3176850000 |
| H | 3.5120680000 | -1.6325080000 | 0.3577940000 |
| N | 1.5877310000 | -2.5206860000 | 0.6096130000 |
| C | 1.1510710000 | -3.8206280000 | 0.4501150000 |
| H | 0.1216150000 | -4.0813500000 | 0.6429950000 |

| | | | |
|---|---|---|---|
| H | 2.2734910000 | -5.6602180000 | -0.1726320000 |
| O | -1.0317520000 | -1.6143980000 | 2.3092580000 |
| H | -0.9383930000 | -1.3478270000 | 3.2355510000 |
| C | 2.4940860000 | 0.9445740000 | 0.1465570000 |
| O | 3.3416220000 | 0.5057950000 | 0.9640790000 |
| O | 1.2529010000 | 0.6323130000 | 0.1231480000 |
| C | 2.9506490000 | 1.9111950000 | -0.9381890000 |
| H | 2.2901910000 | 2.7837590000 | -0.9595050000 |
| H | 2.8752820000 | 1.4223630000 | -1.9172470000 |
| H | 3.9823750000 | 2.2260940000 | -0.7718810000 |
| O | 1.7571740000 | -0.4288900000 | 2.8814710000 |
| H | 2.0544030000 | -1.3236770000 | 3.1085990000 |
| H | 2.4682000000 | -0.0618150000 | 2.2762240000 |

**$^{5}$Im$_2$AcOWatOH_FeF_remove-OH**
Charge: 0
Multiplicity: 5
B3LYP-D3(BJ)/6-31G(d)-SDD SCF energy (au): -981.331484369
B3LYP-D3(BJ)/6-31G(d)-SDD enthalpy (au): -981.085064369
B3LYP-D3(BJ)/6-31G(d)-SDD free energy (au): -981.160461369
B3LYP-D3(BJ)/def2-TZVP/SMD SCF energy (au): -2121.57189667
B3LYP-D3(BJ)/def2-TZVP/SMD enthalpy (au): -2121.32547667
B3LYP-D3(BJ)/def2-TZVP/SMD free energy (au): -2121.40087367
B3LYP-D3(BJ)/def2-TZVP/SMD free energy (quasi-harmonic) (au): -2121.395374301

Cartesian coordinates

| ATOM | X | Y | Z |
|---|---|---|---|
| Fe | 0.4281050000 | -0.4134470000 | 0.9861720000 |
| C | -2.0488830000 | -0.6275320000 | -2.6623630000 |
| N | -2.9217210000 | -1.0031010000 | -1.6547990000 |
| H | -3.8983190000 | -1.2307290000 | -1.7676500000 |
| C | -2.2391190000 | -0.9858490000 | -0.4791260000 |
| H | -2.6517590000 | -1.2149540000 | 0.4917640000 |
| N | -0.9851840000 | -0.6264850000 | -0.6868020000 |
| C | -0.8504190000 | -0.3966370000 | -2.0395510000 |
| H | 0.0935470000 | -0.0681450000 | -2.4454300000 |
| H | -2.3588920000 | -0.5590930000 | -3.6933440000 |
| C | 1.6554640000 | -4.5877680000 | 0.8586660000 |
| N | 2.8752830000 | -3.9789280000 | 0.6203900000 |
| H | 3.7556790000 | -4.4483130000 | 0.4682800000 |
| C | 2.6875960000 | -2.6325580000 | 0.6349810000 |
| H | 3.4633290000 | -1.8917110000 | 0.4936590000 |
| N | 1.4143700000 | -2.3558050000 | 0.8648490000 |
| C | 0.7596110000 | -3.5628360000 | 1.0081380000 |
| H | -0.2983710000 | -3.5917470000 | 1.2190530000 |
| H | 1.5473430000 | -5.6601300000 | 0.9006770000 |
| F | -0.9533500000 | -0.8439780000 | 2.1441220000 |
| C | 2.8905310000 | 0.7058590000 | -0.3139180000 |
| O | 3.6280190000 | 0.3099410000 | 0.6232560000 |
| O | 1.6119260000 | 0.6219060000 | -0.3377410000 |
| C | 3.5304430000 | 1.3248300000 | -1.5478510000 |
| H | 3.0800940000 | 2.3032570000 | -1.7436770000 |
| H | 3.3258580000 | 0.6948110000 | -2.4214330000 |

| | | | |
|---|---|---|---|
| H | 4.6089490000 | 1.4299320000 | -1.4176090000 |
| O | 1.8263920000 | 0.2728090000 | 2.5722790000 |
| H | 1.9023980000 | -0.4815380000 | 3.1765400000 |
| H | 2.6427770000 | 0.2644030000 | 1.9911990000 |

**$^{6}$Im$_2$AcOWatOH_FeF**
Charge: 0
Multiplicity: 6
B3LYP-D3(BJ)/6-31G(d)-SDD SCF energy (au): -1057.14588096
B3LYP-D3(BJ)/6-31G(d)-SDD enthalpy (au): -1056.88392196
B3LYP-D3(BJ)/6-31G(d)-SDD free energy (au): -1056.96007596
B3LYP-D3(BJ)/def2-TZVP/SMD SCF energy (au): -2197.42509744
B3LYP-D3(BJ)/def2-TZVP/SMD enthalpy (au): -2197.16313844
B3LYP-D3(BJ)/def2-TZVP/SMD free energy (au): -2197.23929244
B3LYP-D3(BJ)/def2-TZVP/SMD free energy (quasi-harmonic) (au): -2197.23498448

Cartesian coordinates

| ATOM | X | Y | Z |
|---|---|---|---|
| Fe | 0.2658740000 | -0.5227400000 | 1.3283950000 |
| C | -1.9988520000 | -1.5957990000 | -2.3126840000 |
| N | -2.5461600000 | -0.3969390000 | -1.8881530000 |
| H | -3.2832140000 | 0.1176860000 | -2.3468030000 |
| C | -1.9164140000 | -0.0255840000 | -0.7456350000 |
| H | -2.1155500000 | 0.8690750000 | -0.1765310000 |
| N | -1.0017910000 | -0.9226150000 | -0.4223940000 |
| C | -1.0383300000 | -1.9060980000 | -1.3874870000 |
| H | -0.3798540000 | -2.7590020000 | -1.3415910000 |
| H | -2.3371350000 | -2.0983000000 | -3.2050580000 |
| C | 1.7889490000 | -4.5968040000 | 0.4167090000 |
| N | 2.8803940000 | -3.8638690000 | -0.0210910000 |
| H | 3.7331850000 | -4.2384070000 | -0.4102130000 |
| C | 2.6217740000 | -2.5484900000 | 0.1962120000 |
| H | 3.3184120000 | -1.7457570000 | 0.0041110000 |
| N | 1.4269180000 | -2.4063350000 | 0.7442890000 |
| C | 0.8969950000 | -3.6740700000 | 0.8947530000 |
| H | -0.0700840000 | -3.7995910000 | 1.3554060000 |
| H | 1.7584580000 | -5.6731970000 | 0.3539330000 |
| O | -0.5374880000 | -1.7277370000 | 2.5249580000 |
| H | -1.2523240000 | -1.2444110000 | 2.9709150000 |
| F | -0.7896350000 | 0.9376340000 | 1.6583040000 |
| C | 2.7242960000 | 0.7516280000 | -0.0418980000 |
| O | 3.5568660000 | 0.3447460000 | 0.8106110000 |
| O | 1.4797550000 | 0.4620650000 | -0.0504360000 |
| C | 3.2019300000 | 1.6473040000 | -1.1734110000 |
| H | 2.7062650000 | 2.6206950000 | -1.0915680000 |
| H | 2.9108330000 | 1.2173720000 | -2.1376140000 |
| H | 4.2841520000 | 1.7832340000 | -1.1321780000 |
| O | 1.8545950000 | -0.1592810000 | 2.7087020000 |
| H | 1.9362810000 | -1.0137370000 | 3.1617940000 |
| H | 2.6527840000 | -0.0383590000 | 2.0942570000 |

**$^{5}$Im$_2$AcOWatOH_TS-1**
Charge: 0

Multiplicity: 5
B3LYP-D3(BJ)/6-31G(d)-SDD SCF energy (au): -1693.32360708
B3LYP-D3(BJ)/6-31G(d)-SDD enthalpy (au): -1692.75945808
B3LYP-D3(BJ)/6-31G(d)-SDD free energy (au): -1692.87577008
B3LYP-D3(BJ)/def2-TZVP/SMD SCF energy (au): -2833.85173138
B3LYP-D3(BJ)/def2-TZVP/SMD enthalpy (au): -2833.28758238
B3LYP-D3(BJ)/def2-TZVP/SMD free energy (au): -2833.40389438
B3LYP-D3(BJ)/def2-TZVP/SMD free energy (quasi-harmonic) (au): -2833.39236730

Cartesian coordinates

| ATOM | X | Y | Z |
|---|---|---|---|
| Fe | 0.3588150000 | -0.6547090000 | 1.2343600000 |
| C | -2.1014670000 | -1.7021270000 | -2.3720780000 |
| N | -2.5279740000 | -0.4243930000 | -2.0536860000 |
| H | -3.2271950000 | 0.1106680000 | -2.5466120000 |
| C | -1.8417050000 | -0.0159930000 | -0.9520970000 |
| H | -1.9650980000 | 0.9433730000 | -0.4753060000 |
| N | -1.0060340000 | -0.9568270000 | -0.5561050000 |
| C | -1.1552250000 | -2.0121280000 | -1.4298710000 |
| H | -0.5773540000 | -2.9164890000 | -1.3124520000 |
| H | -2.5030030000 | -2.2493710000 | -3.2105770000 |
| C | 1.7958820000 | -4.6335480000 | 0.4540750000 |
| N | 2.8906560000 | -3.9308670000 | -0.0228350000 |
| H | 3.7312510000 | -4.3296150000 | -0.4136400000 |
| C | 2.6568280000 | -2.6048910000 | 0.1625390000 |
| H | 3.3540520000 | -1.8117010000 | -0.0653790000 |
| N | 1.4725540000 | -2.4312760000 | 0.7230920000 |
| C | 0.9237730000 | -3.6850610000 | 0.9176770000 |
| H | -0.0376840000 | -3.7782670000 | 1.3990360000 |
| H | 1.7492250000 | -5.7108230000 | 0.4252360000 |
| O | -0.5082430000 | -1.6588920000 | 2.5299650000 |
| H | -1.1320930000 | -1.0713790000 | 2.9937170000 |
| C | -5.2552050000 | 3.3534900000 | 5.1947570000 |
| C | -4.1252080000 | 2.6269100000 | 4.8370190000 |
| C | -3.8742250000 | 2.3144680000 | 3.4928140000 |
| C | -4.7935360000 | 2.6739740000 | 2.4882540000 |
| C | -5.9331190000 | 3.3918990000 | 2.8757400000 |
| C | -6.1590760000 | 3.7457920000 | 4.2040310000 |
| H | -5.4356320000 | 3.6048730000 | 6.2356090000 |
| H | -3.4159250000 | 2.2970200000 | 5.5899440000 |
| H | -6.6629400000 | 3.6633190000 | 2.1164330000 |
| H | -7.0492200000 | 4.3091480000 | 4.4695690000 |
| C | -4.6449830000 | 2.2634230000 | 1.0359860000 |
| C | -4.1473510000 | 3.3948040000 | 0.1251560000 |
| H | -3.9706150000 | 1.4071220000 | 0.9515970000 |
| H | -5.6274630000 | 1.9255900000 | 0.6813500000 |
| H | -4.1396230000 | 3.0744730000 | -0.9239860000 |
| H | -3.1305670000 | 3.6835920000 | 0.4019310000 |
| H | -4.7914790000 | 4.2781150000 | 0.2009740000 |
| C | -2.6137280000 | 1.5537050000 | 3.2004410000 |
| O | -2.3509730000 | 0.4987830000 | 3.7497240000 |
| N | -1.8485560000 | 2.1924050000 | 2.2081340000 |
| C | -0.8117140000 | 3.1829970000 | 2.5863520000 |

| | | | |
|---|---|---|---|
| C | -1.5570220000 | 4.4487350000 | 3.0722680000 |
| C | 0.1166110000 | 2.6594290000 | 3.6914940000 |
| H | -0.8147870000 | 5.2402340000 | 3.2189760000 |
| H | -2.0731970000 | 4.2828540000 | 4.0201620000 |
| H | -2.2891260000 | 4.7913710000 | 2.3358030000 |
| H | 0.6477050000 | 1.7605980000 | 3.3734210000 |
| H | -0.4461240000 | 2.4272420000 | 4.6021100000 |
| H | 0.8529860000 | 3.4327100000 | 3.9377280000 |
| F | -0.9205640000 | 0.8723260000 | 1.5888780000 |
| C | -0.0227150000 | 3.5254780000 | 1.3145400000 |
| H | -0.6961670000 | 3.8975880000 | 0.5343720000 |
| H | 0.5035970000 | 2.6498820000 | 0.9320880000 |
| H | 0.7056950000 | 4.3112210000 | 1.5428550000 |
| C | 2.6741750000 | 0.8052530000 | -0.0255560000 |
| O | 3.5297650000 | 0.2488000000 | 0.7020800000 |
| O | 1.4079400000 | 0.6041900000 | 0.0202940000 |
| C | 3.1259460000 | 1.8164550000 | -1.0706430000 |
| H | 2.6947470000 | 2.7959120000 | -0.8345190000 |
| H | 2.7479330000 | 1.5279980000 | -2.0571740000 |
| H | 4.2146050000 | 1.8933670000 | -1.0928180000 |
| O | 1.9273510000 | -0.0894660000 | 2.8215810000 |
| H | 1.8430110000 | -0.9692770000 | 3.2232020000 |
| H | 2.6876890000 | -0.1103040000 | 2.1806940000 |

**$^{5}$Im$_2$AcOWatOH_TS-3-F**
Charge: 0
Multiplicity: 5
B3LYP-D3(BJ)/6-31G(d)-SDD SCF energy (au): -1693.39728376
B3LYP-D3(BJ)/6-31G(d)-SDD enthalpy (au): -1692.83216776
B3LYP-D3(BJ)/6-31G(d)-SDD free energy (au): -1692.94522376
B3LYP-D3(BJ)/def2-TZVP/SMD SCF energy (au): -2833.93164307
B3LYP-D3(BJ)/def2-TZVP/SMD enthalpy (au): -2833.36652707
B3LYP-D3(BJ)/def2-TZVP/SMD free energy (au): -2833.47958307
B3LYP-D3(BJ)/def2-TZVP/SMD free energy (quasi-harmonic) (au): -2833.46954231

Cartesian coordinates

| ATOM | X | Y | Z |
|---|---|---|---|
| H | 33.5512180000 | 27.8926570000 | 39.6626750000 |
| C | 33.5856310000 | 28.2645230000 | 38.6505030000 |
| C | 34.5843660000 | 28.8382130000 | 37.9082490000 |
| N | 32.4791870000 | 28.2317320000 | 37.8161910000 |
| C | 32.8362920000 | 28.7777680000 | 36.6227130000 |
| N | 34.1020140000 | 29.1469230000 | 36.6554840000 |
| H | 31.5633840000 | 27.8827560000 | 38.0554860000 |
| H | 35.6064890000 | 29.0663700000 | 38.1666930000 |
| H | 32.1973430000 | 28.9070990000 | 35.7616250000 |
| H | 35.9290360000 | 25.4554440000 | 32.5874320000 |
| C | 36.0220490000 | 26.3298930000 | 33.2123730000 |
| C | 35.2426390000 | 27.4471990000 | 33.3630160000 |
| N | 37.0857500000 | 26.4991390000 | 34.0849860000 |
| C | 36.9177690000 | 27.6902240000 | 34.7232340000 |
| N | 35.8156540000 | 28.2758230000 | 34.3026550000 |
| H | 37.8484680000 | 25.8540160000 | 34.2255690000 |

| | | | |
|---|---|---|---|
| H | 34.3195300000 | 27.7379540000 | 32.8853000000 |
| H | 37.5827030000 | 28.1028270000 | 35.4664810000 |
| Fe | 35.0015370000 | 30.2201570000 | 34.9397070000 |
| O | 34.4341850000 | 32.1543550000 | 35.7004420000 |
| H | 35.3505260000 | 32.5148380000 | 35.8958790000 |
| H | 34.0571610000 | 32.6216350000 | 34.9276540000 |
| F | 36.1732160000 | 30.9898220000 | 33.5141040000 |
| N | 34.3953740000 | 32.9533030000 | 31.1144920000 |
| O | 33.2912510000 | 32.7256840000 | 33.1124760000 |
| C | 35.2364180000 | 34.0755210000 | 33.0915560000 |
| C | 34.7554870000 | 35.2373660000 | 33.6937990000 |
| C | 35.5984810000 | 36.0526990000 | 34.4488070000 |
| C | 36.9278070000 | 35.6667670000 | 34.6456480000 |
| C | 37.4055130000 | 34.4898820000 | 34.0885600000 |
| C | 36.5887240000 | 33.6696850000 | 33.2763220000 |
| C | 34.2256640000 | 33.1805910000 | 32.4357590000 |
| C | 33.6944540000 | 31.9531110000 | 30.2716000000 |
| C | 34.1009200000 | 30.5344310000 | 30.7037470000 |
| C | 32.1738860000 | 32.1481010000 | 30.3663650000 |
| C | 34.1602820000 | 32.2215740000 | 28.8336700000 |
| C | 37.0975740000 | 32.4427900000 | 32.7095140000 |
| C | 38.4887400000 | 31.9740320000 | 32.9837270000 |
| H | 33.7062770000 | 35.4917710000 | 33.5762820000 |
| H | 35.2164460000 | 36.9642860000 | 34.8980250000 |
| H | 37.5870730000 | 36.2759580000 | 35.2568440000 |
| H | 38.4266540000 | 34.1892570000 | 34.2859160000 |
| H | 35.1847110000 | 33.4217130000 | 30.6911740000 |
| H | 36.6733070000 | 32.1279850000 | 31.7664100000 |
| H | 38.6980810000 | 31.9963880000 | 34.0560940000 |
| H | 38.6125720000 | 30.9493200000 | 32.6269370000 |
| H | 39.2310560000 | 32.6088780000 | 32.4765200000 |
| H | 33.6079090000 | 29.7991190000 | 30.0562330000 |
| H | 35.1844120000 | 30.4005840000 | 30.6103600000 |
| H | 33.8283330000 | 30.3245510000 | 31.7416230000 |
| H | 31.8225130000 | 31.9730480000 | 31.3833260000 |
| H | 31.8979370000 | 33.1671580000 | 30.0730450000 |
| H | 31.6737400000 | 31.4460620000 | 29.6901920000 |
| H | 35.2472120000 | 32.0998980000 | 28.7408710000 |
| H | 33.6877400000 | 31.5105340000 | 28.1497340000 |
| H | 33.8922790000 | 33.2349530000 | 28.5123250000 |
| C | 37.3725350000 | 31.3530710000 | 36.4786210000 |
| O | 36.9817440000 | 32.5310380000 | 36.3270120000 |
| O | 36.6971840000 | 30.2943260000 | 36.2214700000 |
| C | 38.7889390000 | 31.1115980000 | 36.9960480000 |
| H | 39.4032460000 | 30.7017010000 | 36.1844120000 |
| H | 38.7767690000 | 30.3685830000 | 37.8002980000 |
| H | 39.2400290000 | 32.0400800000 | 37.3523380000 |
| O | 33.4004180000 | 29.9192800000 | 33.9515380000 |
| H | 33.0616650000 | 30.7735750000 | 33.6366260000 |

**$^5$Im$_2$AcOWatOH_TS-3-OH**
Charge: 0
Multiplicity: 5

B3LYP-D3(BJ)/6-31G(d)-SDD SCF energy (au): -1693.37592699
B3LYP-D3(BJ)/6-31G(d)-SDD enthalpy (au): -1692.81174199
B3LYP-D3(BJ)/6-31G(d)-SDD free energy (au): -1692.92628199
B3LYP-D3(BJ)/def2-TZVP/SMD SCF energy (au): -2833.93101328
B3LYP-D3(BJ)/def2-TZVP/SMD enthalpy (au): -2833.36682828
B3LYP-D3(BJ)/def2-TZVP/SMD free energy (au): -2833.48136828
B3LYP-D3(BJ)/def2-TZVP/SMD free energy (quasi-harmonic) (au): -2833.46975229

Cartesian coordinates

| ATOM | X | Y | Z |
|---|---|---|---|
| H | 31.9962890000 | 26.9758330000 | 36.7271170000 |
| C | 32.5868840000 | 27.8439290000 | 36.4800170000 |
| C | 33.7937810000 | 27.9895800000 | 35.8500070000 |
| N | 32.1823260000 | 29.1285150000 | 36.8007570000 |
| C | 33.1339130000 | 29.9942310000 | 36.3673830000 |
| N | 34.1185710000 | 29.3299210000 | 35.7864770000 |
| H | 31.3351850000 | 29.3839720000 | 37.2859710000 |
| H | 34.4608400000 | 27.2413390000 | 35.4543220000 |
| H | 33.1099270000 | 31.0676910000 | 36.4702820000 |
| H | 39.0131370000 | 26.0222830000 | 36.4121470000 |
| C | 38.3234500000 | 26.6645700000 | 35.8870990000 |
| C | 37.5205840000 | 27.6957060000 | 36.2994650000 |
| N | 38.1075870000 | 26.5509530000 | 34.5227030000 |
| C | 37.2031840000 | 27.5024500000 | 34.1638990000 |
| N | 36.8286140000 | 28.1975890000 | 35.2186290000 |
| H | 38.5694340000 | 25.9057630000 | 33.8993370000 |
| H | 37.4040150000 | 28.1466810000 | 37.2725100000 |
| H | 36.8717570000 | 27.6868360000 | 33.1537230000 |
| Fe | 35.9281710000 | 30.2778000000 | 35.0716060000 |
| O | 34.7571320000 | 32.0808380000 | 35.4126410000 |
| H | 35.4845790000 | 32.5993940000 | 35.8764930000 |
| H | 34.4072270000 | 32.5487010000 | 34.6250870000 |
| F | 37.5678390000 | 30.7764070000 | 34.3887710000 |
| N | 33.4893110000 | 32.4696990000 | 30.9920390000 |
| O | 33.2528820000 | 32.9016810000 | 33.2174480000 |
| C | 35.4163040000 | 33.2006020000 | 32.2796960000 |
| C | 35.7179290000 | 34.4192040000 | 32.8868980000 |
| C | 37.0393040000 | 34.7977300000 | 33.1195700000 |
| C | 38.0707660000 | 33.9169150000 | 32.7786070000 |
| C | 37.7836190000 | 32.6849360000 | 32.2138040000 |
| C | 36.4536350000 | 32.2857990000 | 31.9259390000 |
| C | 33.9690340000 | 32.8393330000 | 32.2055960000 |
| C | 32.1467600000 | 31.9111950000 | 30.7278990000 |
| C | 31.9353570000 | 30.6514850000 | 31.5863040000 |
| C | 31.0703580000 | 32.9647390000 | 31.0352900000 |
| C | 32.1163210000 | 31.5430890000 | 29.2394700000 |
| C | 36.1864040000 | 30.9781000000 | 31.4189860000 |
| C | 37.2667520000 | 29.9650780000 | 31.2347150000 |
| H | 34.9000210000 | 35.0660590000 | 33.1901620000 |
| H | 37.2596590000 | 35.7529110000 | 33.5857390000 |
| H | 39.1031200000 | 34.1834370000 | 32.9854090000 |
| H | 38.5938020000 | 31.9983070000 | 32.0054360000 |
| H | 34.1389700000 | 32.5106800000 | 30.2194630000 |

H 35.2160530000 30.7671100000 30.9960700000
H 37.8135220000 29.8539310000 32.1777940000
H 36.8429850000 28.9978660000 30.9490840000
H 37.9841900000 30.2607700000 30.4530180000
H 30.9486850000 30.2197370000 31.3847060000
H 32.6962980000 29.8981740000 31.3527200000
H 32.0023760000 30.9000220000 32.6473720000
H 31.1196680000 33.2574190000 32.0856030000
H 31.2201950000 33.8557300000 30.4158670000
H 30.0744540000 32.5595190000 30.8228280000
H 32.8813690000 30.7937730000 29.0018900000
H 31.1406420000 31.1241840000 28.9755410000
H 32.2855700000 32.4262340000 28.6118140000
C 37.0937320000 31.6823340000 37.2999610000
O 36.7681050000 32.8100680000 36.8816560000
O 36.5545880000 30.5602690000 36.9496260000
C 38.2249330000 31.5543890000 38.3087580000
H 39.0966940000 31.1321590000 37.7949410000
H 37.9469520000 30.8621370000 39.1099510000
H 38.4890580000 32.5287570000 38.7248690000
O 35.3292570000 30.0237300000 33.2801490000
H 34.4271450000 29.6553640000 33.2677480000

**$^{5}$Im$_2$WatOHN$_3$_11**
Charge: 0
Multiplicity: 5
B3LYP-D3(BJ)/6-31G(d)-SDD SCF energy (au): -1629.03798869
B3LYP-D3(BJ)/6-31G(d)-SDD enthalpy (au): -1628.51129569
B3LYP-D3(BJ)/6-31G(d)-SDD free energy (au): -1628.62336469
B3LYP-D3(BJ)/def2-TZVP/SMD SCF energy (au): -2769.53627779
B3LYP-D3(BJ)/def2-TZVP/SMD enthalpy (au): -2769.00958479
B3LYP-D3(BJ)/def2-TZVP/SMD free energy (au): -2769.12165379
B3LYP-D3(BJ)/def2-TZVP/SMD free energy (quasi-harmonic) (au): -2769.11101110

Cartesian coordinates
ATOM X Y Z
Fe 0.1193830000 -0.8928110000 -0.7224550000
C -1.6638260000 -0.2101600000 3.2900070000
N -1.2303740000 1.0410470000 2.8846010000
H -1.3745640000 1.9092220000 3.3781920000
C -0.5997170000 0.9013830000 1.6874670000
H -0.1882000000 1.7041610000 1.0942310000
N -0.6065640000 -0.3655630000 1.3145690000
C -1.2672380000 -1.0706580000 2.3000320000
H -1.4176780000 -2.1357880000 2.2117450000
H -2.2019230000 -0.3602360000 4.2127360000
C 3.6649010000 -2.0715110000 1.5521050000
N 4.0946040000 -2.2682090000 0.2515880000
H 5.0069980000 -2.5913140000 -0.0349750000
C 3.0751110000 -1.9539300000 -0.5869620000
H 3.1438450000 -2.0231580000 -1.6650360000
N 2.0166290000 -1.5650970000 0.1101920000
C 2.3728050000 -1.6338490000 1.4444700000

| | | | |
|---|---|---|---|
| H | 1.6699870000 | -1.3675020000 | 2.2178970000 |
| H | 4.3014220000 | -2.2584980000 | 2.4024990000 |
| O | -0.0168270000 | 0.9498360000 | -1.1749580000 |
| H | 0.3356350000 | 1.0998600000 | -2.0649400000 |
| C | -7.1893460000 | 1.5607970000 | -1.4726440000 |
| C | -6.2560240000 | 0.5554080000 | -1.7040570000 |
| C | -4.9158680000 | 0.7295440000 | -1.3297430000 |
| C | -4.4848500000 | 1.9207100000 | -0.7161960000 |
| C | -5.4426320000 | 2.9212400000 | -0.5044610000 |
| C | -6.7762410000 | 2.7508980000 | -0.8709260000 |
| H | -8.2276610000 | 1.4166230000 | -1.7556890000 |
| H | -6.5555490000 | -0.3786120000 | -2.1686370000 |
| H | -5.1288280000 | 3.8535950000 | -0.0418160000 |
| H | -7.4934890000 | 3.5455620000 | -0.6851690000 |
| C | -3.0387270000 | 2.1791450000 | -0.3556560000 |
| C | -2.3959130000 | 3.2736580000 | -1.2219230000 |
| H | -2.4374570000 | 1.2760350000 | -0.4512250000 |
| H | -2.9867700000 | 2.4751150000 | 0.7008900000 |
| H | -1.3298390000 | 3.3438370000 | -0.9908620000 |
| H | -2.4800640000 | 3.0228320000 | -2.2847940000 |
| H | -2.8697980000 | 4.2503920000 | -1.0698200000 |
| C | -3.9893860000 | -0.4295060000 | -1.4985430000 |
| O | -3.4675920000 | -0.9759440000 | -0.5302010000 |
| N | -3.9105470000 | -0.9468190000 | -2.7871950000 |
| C | -3.4652720000 | -0.2215540000 | -4.0329430000 |
| C | -4.3337980000 | 1.0211990000 | -4.2683380000 |
| C | -3.6779390000 | -1.2002930000 | -5.1976820000 |
| H | -4.0495140000 | 1.4305980000 | -5.2428790000 |
| H | -5.3987500000 | 0.7776740000 | -4.2952800000 |
| H | -4.1732510000 | 1.7966950000 | -3.5190860000 |
| H | -3.0566780000 | -2.0901390000 | -5.0875320000 |
| H | -4.7276000000 | -1.5061020000 | -5.2603240000 |
| H | -3.4039840000 | -0.7034560000 | -6.1333130000 |
| F | -3.1524680000 | -2.1421340000 | -2.7272800000 |
| C | -1.9875160000 | 0.1656180000 | -3.8964290000 |
| H | -1.8247170000 | 0.8289680000 | -3.0413800000 |
| H | -1.3506510000 | -0.7109460000 | -3.7529670000 |
| H | -1.6656180000 | 0.6849000000 | -4.8062530000 |
| O | -1.2566540000 | -2.5781640000 | -0.5447840000 |
| H | -1.0383850000 | -2.8911200000 | -1.4461180000 |
| H | -2.1445220000 | -2.1503500000 | -0.6108060000 |
| N | 2.3803370000 | -2.1026390000 | -3.8283300000 |
| N | 1.3848920000 | -2.0105920000 | -3.2294410000 |
| N | 0.3580120000 | -1.9217880000 | -2.6085160000 |

**$^{5}Im_2WatOHN_3$_13**
Charge: 0
Multiplicity: 5

| | |
|---|---|
| B3LYP-D3(BJ)/6-31G(d)-SDD SCF energy (au): | -1629.10744341 |
| B3LYP-D3(BJ)/6-31G(d)-SDD enthalpy (au): | -1628.58240141 |
| B3LYP-D3(BJ)/6-31G(d)-SDD free energy (au): | -1628.69489141 |
| B3LYP-D3(BJ)/def2-TZVP/SMD SCF energy (au): | -2769.61332194 |
| B3LYP-D3(BJ)/def2-TZVP/SMD enthalpy (au): | -2769.08827994 |

B3LYP-D3(BJ)/def2-TZVP/SMD free energy (au): -2769.20076994
B3LYP-D3(BJ)/def2-TZVP/SMD free energy (quasi-harmonic) (au): -2769.19001753

Cartesian coordinates

| ATOM | X | Y | Z |
|---|---|---|---|
| H | 32.4424500000 | 29.0907250000 | 38.1211620000 |
| C | 33.0201550000 | 29.1979380000 | 37.2168630000 |
| C | 34.2930280000 | 29.6449420000 | 36.9840180000 |
| N | 32.5080990000 | 28.8731390000 | 35.9713890000 |
| C | 33.4623250000 | 29.1303660000 | 35.0414070000 |
| N | 34.5506070000 | 29.5949930000 | 35.6299630000 |
| H | 31.5734920000 | 28.5466140000 | 35.7768000000 |
| H | 35.0326700000 | 30.0108350000 | 37.6777480000 |
| H | 33.3568150000 | 28.9772910000 | 33.9791290000 |
| H | 39.1430870000 | 26.8079350000 | 37.5028400000 |
| C | 38.5462240000 | 27.2695560000 | 36.7319700000 |
| C | 37.9957420000 | 28.5181370000 | 36.6137240000 |
| N | 38.1896360000 | 26.5968180000 | 35.5743660000 |
| C | 37.4520950000 | 27.4406080000 | 34.8068320000 |
| N | 37.3207750000 | 28.6041020000 | 35.4151340000 |
| H | 38.4430420000 | 25.6502500000 | 35.3337210000 |
| H | 38.0314360000 | 29.3638410000 | 37.2818750000 |
| H | 37.0273620000 | 27.2132550000 | 33.8407490000 |
| Fe | 36.2663040000 | 30.2772370000 | 34.4206630000 |
| O | 35.7572240000 | 29.0868780000 | 33.0456160000 |
| H | 36.0258150000 | 29.4786460000 | 32.1985310000 |
| F | 37.8687440000 | 31.0359420000 | 33.9089780000 |
| N | 33.4843700000 | 34.5014870000 | 36.0338270000 |
| O | 34.8953850000 | 33.0335750000 | 37.0425830000 |
| C | 35.8534800000 | 34.5894250000 | 35.5022390000 |
| C | 37.0369090000 | 34.6711150000 | 36.2371240000 |
| C | 38.1835070000 | 35.2708770000 | 35.7159820000 |
| C | 38.1455590000 | 35.8038810000 | 34.4214620000 |
| C | 36.9877380000 | 35.7221590000 | 33.6675220000 |
| C | 35.8027260000 | 35.0997370000 | 34.1587650000 |
| C | 34.7055430000 | 33.9605300000 | 36.2296080000 |
| C | 32.2253090000 | 34.0988690000 | 36.7008050000 |
| C | 31.9394340000 | 32.6095610000 | 36.4563610000 |
| C | 32.3265190000 | 34.3955140000 | 38.2062970000 |
| C | 31.1214330000 | 34.9502250000 | 36.0601380000 |
| C | 34.6829060000 | 34.9446910000 | 33.3062700000 |
| C | 34.6348310000 | 35.4262310000 | 31.8910830000 |
| H | 37.0423550000 | 34.2642100000 | 37.2430980000 |
| H | 39.0891160000 | 35.3275140000 | 36.3118870000 |
| H | 39.0296780000 | 36.2736650000 | 33.9995920000 |
| H | 36.9892390000 | 36.1093490000 | 32.6548370000 |
| H | 33.4434440000 | 35.3128640000 | 35.4327290000 |
| H | 33.8568760000 | 34.3270180000 | 33.6315500000 |
| H | 35.3035200000 | 34.8338380000 | 31.2471690000 |
| H | 33.6243710000 | 35.3269590000 | 31.4850790000 |
| H | 34.9419150000 | 36.4753430000 | 31.7880160000 |
| H | 30.9742960000 | 32.3497220000 | 36.9079320000 |
| H | 31.8986420000 | 32.3928770000 | 35.3850260000 |

H 32.7146440000 31.9881090000 36.9048060000
H 33.1463080000 33.8242930000 38.6475700000
H 32.5074120000 35.4622800000 38.3804350000
H 31.3927300000 34.1181720000 38.7083810000
H 31.0453610000 34.7439840000 34.9865310000
H 30.1560830000 34.7184070000 36.5198250000
H 31.3118260000 36.0217320000 36.2018140000
N 35.1184790000 31.8834780000 33.8878090000
N 33.9679310000 31.8020900000 33.5519390000
N 32.8508100000 31.7771830000 33.2273840000
O 36.8584620000 31.3443350000 36.3268810000
H 36.1594410000 32.0083830000 36.5774400000
H 37.5241200000 31.8285170000 35.8032180000

**$^{5}Im_2WatOHN_3$_FeF_remove-F**
Charge: 0
Multiplicity: 5
B3LYP-D3(BJ)/6-31G(d)-SDD SCF energy (au): -892.967172174
B3LYP-D3(BJ)/6-31G(d)-SDD enthalpy (au): -892.750210174
B3LYP-D3(BJ)/6-31G(d)-SDD free energy (au): -892.824324174
B3LYP-D3(BJ)/def2-TZVP/SMD SCF energy (au): -2033.17228999
B3LYP-D3(BJ)/def2-TZVP/SMD enthalpy (au): -2032.95532799
B3LYP-D3(BJ)/def2-TZVP/SMD free energy (au): -2033.02944199
B3LYP-D3(BJ)/def2-TZVP/SMD free energy (quasi-harmonic) (au): -2033.024075816

Cartesian coordinates
ATOM X Y Z
Fe 0.0865390000 -0.9413600000 1.4322710000
C -1.0592410000 -1.1035090000 -2.7777900000
N -2.2177710000 -1.4324260000 -2.0940420000
H -3.1090350000 -1.6624280000 -2.5074600000
C -1.9487350000 -1.3877460000 -0.7639540000
H -2.6513910000 -1.6034680000 0.0275640000
N -0.6852150000 -1.0498000000 -0.5689000000
C -0.1136340000 -0.8690470000 -1.8150480000
H 0.9284470000 -0.5923280000 -1.9074000000
H -1.0209520000 -1.0674710000 -3.8551530000
C 2.1820590000 -4.6865220000 1.8794830000
N 3.0831850000 -4.0079660000 1.0774870000
H 3.9642620000 -4.3643220000 0.7381720000
C 2.5836260000 -2.7705250000 0.8235610000
H 3.0648270000 -2.0212690000 0.2047590000
N 1.4116120000 -2.6295650000 1.4236840000
C 1.1475060000 -3.8141390000 2.0851110000
H 0.2341720000 -3.9332440000 2.6480410000
H 2.3603510000 -5.6946240000 2.2188520000
O -1.4345420000 -1.5589730000 2.3226920000
H -1.7132690000 -1.0747450000 3.1127900000
O 2.9220790000 -0.2263590000 -0.9169400000
H 2.4121240000 0.1543500000 -0.1562910000
H 3.5191540000 0.4866240000 -1.1882750000
N 1.3885230000 0.5753580000 1.3008880000
N 1.9113850000 1.2234870000 2.1878120000

N      2.4286430000    1.8658910000    2.9928020000

**⁵Im₂WatOHN₃_FeF_remove-N₃**
Charge: 0
Multiplicity: 5
B3LYP-D3(BJ)/6-31G(d)-SDD SCF energy (au):              -828.633665982
B3LYP-D3(BJ)/6-31G(d)-SDD enthalpy (au):                -828.429699982
B3LYP-D3(BJ)/6-31G(d)-SDD free energy (au):             -828.497182982
B3LYP-D3(BJ)/def2-TZVP/SMD SCF energy (au):             -1968.81692155
B3LYP-D3(BJ)/def2-TZVP/SMD enthalpy (au):               -1968.61295555
B3LYP-D3(BJ)/def2-TZVP/SMD free energy (au):            -1968.68043855
B3LYP-D3(BJ)/def2-TZVP/SMD free energy (quasi-harmonic) (au): -1968.677087568

Cartesian coordinates

| ATOM | X | Y | Z |
|---|---|---|---|
| Fe | -0.0458780000 | -0.9294730000 | 1.3099980000 |
| C | -2.3589010000 | -2.0961380000 | -2.1534760000 |
| N | -1.3710130000 | -1.3530350000 | -2.7771460000 |
| H | -1.2901750000 | -1.1837310000 | -3.7685530000 |
| C | -0.5251710000 | -0.8816210000 | -1.8225560000 |
| H | 0.3551780000 | -0.2771350000 | -2.0111860000 |
| N | -0.9265660000 | -1.2868430000 | -0.6288140000 |
| C | -2.0663090000 | -2.0442740000 | -0.8169740000 |
| H | -2.5711320000 | -2.4848500000 | 0.0295050000 |
| H | -3.1524590000 | -2.5743110000 | -2.7057690000 |
| C | 3.0227890000 | -3.9716010000 | 1.4785060000 |
| N | 3.6179460000 | -3.0958770000 | 0.5862940000 |
| H | 4.5195360000 | -3.2134130000 | 0.1488040000 |
| C | 2.7806840000 | -2.0404950000 | 0.4017720000 |
| H | 2.9717390000 | -1.1982530000 | -0.2540890000 |
| N | 1.6884830000 | -2.2026140000 | 1.1305150000 |
| C | 1.8228370000 | -3.3996860000 | 1.8065410000 |
| H | 1.0403770000 | -3.7405930000 | 2.4675920000 |
| H | 3.4987890000 | -4.8893430000 | 1.7860420000 |
| F | 0.7649800000 | 0.7379930000 | 0.9722920000 |
| O | -1.1460480000 | -1.9427020000 | 2.4262910000 |
| H | -1.5369190000 | -1.5060080000 | 3.1956740000 |
| O | 2.1701250000 | 0.5586280000 | -1.2066800000 |
| H | 2.5854000000 | 1.4178590000 | -1.3704690000 |
| H | 1.6500100000 | 0.6915720000 | -0.3653290000 |

**⁵Im₂WatOHN₃_FeF_remove-OH**
Charge: 0
Multiplicity: 5
B3LYP-D3(BJ)/6-31G(d)-SDD SCF energy (au):              -917.015831410
B3LYP-D3(BJ)/6-31G(d)-SDD enthalpy (au):                -916.809594410
B3LYP-D3(BJ)/6-31G(d)-SDD free energy (au):             -916.882425410
B3LYP-D3(BJ)/def2-TZVP/SMD SCF energy (au):             -2057.23153675
B3LYP-D3(BJ)/def2-TZVP/SMD enthalpy (au):               -2057.02529975
B3LYP-D3(BJ)/def2-TZVP/SMD free energy (au):            -2057.09813075
B3LYP-D3(BJ)/def2-TZVP/SMD free energy (quasi-harmonic) (au): -2057.092275340

Cartesian coordinates

| ATOM | X | Y | Z |
|---|---|---|---|
| Fe | 0.7042400000 | -0.1967990000 | 0.8762950000 |
| C | -2.6147420000 | -2.1907250000 | -1.1141540000 |
| N | -2.1082350000 | -1.3712630000 | -2.1071650000 |
| H | -2.4678520000 | -1.2815530000 | -3.0459810000 |
| C | -1.0477740000 | -0.6951900000 | -1.5967120000 |
| H | -0.4545370000 | 0.0265550000 | -2.1376860000 |
| N | -0.8527010000 | -1.0434480000 | -0.3360170000 |
| C | -1.8230660000 | -1.9757730000 | -0.0177740000 |
| H | -1.8833540000 | -2.4094870000 | 0.9700180000 |
| H | -3.4685690000 | -2.8292860000 | -1.2761740000 |
| C | 3.0130470000 | -3.8173580000 | 1.3140800000 |
| N | 3.8278350000 | -3.1076230000 | 0.4500760000 |
| H | 4.7042820000 | -3.4238130000 | 0.0622520000 |
| C | 3.2547110000 | -1.9012000000 | 0.2094390000 |
| H | 3.6668660000 | -1.1212500000 | -0.4288140000 |
| N | 2.1124900000 | -1.8098370000 | 0.8776430000 |
| C | 1.9488790000 | -2.9974410000 | 1.5719760000 |
| H | 1.0907320000 | -3.1715220000 | 2.2046910000 |
| H | 3.2615350000 | -4.8090250000 | 1.6574510000 |
| F | 1.2849180000 | 1.0383840000 | -0.3929250000 |
| O | 3.7501460000 | 0.6837180000 | -1.3227120000 |
| H | 4.2402680000 | 1.3134990000 | -0.7730470000 |
| H | 2.8084580000 | 0.8621000000 | -1.0856330000 |
| N | -0.0445340000 | -0.3473790000 | 2.6928470000 |
| N | -0.5709800000 | -1.3646960000 | 3.0796350000 |
| N | -1.0767390000 | -2.3320800000 | 3.4728450000 |

**$^6Im_2WatOHN_3$_FeF**
Charge: 0
Multiplicity: 6
B3LYP-D3(BJ)/6-31G(d)-SDD SCF energy (au): -992.832574410
B3LYP-D3(BJ)/6-31G(d)-SDD enthalpy (au): -992.611118410
B3LYP-D3(BJ)/6-31G(d)-SDD free energy (au): -992.685312410
B3LYP-D3(BJ)/def2-TZVP/SMD SCF energy (au): -2133.08149357
B3LYP-D3(BJ)/def2-TZVP/SMD enthalpy (au): -2132.86003757
B3LYP-D3(BJ)/def2-TZVP/SMD free energy (au): -2132.93423157
B3LYP-D3(BJ)/def2-TZVP/SMD free energy (quasi-harmonic) (au): -2132.929378160

Cartesian coordinates

| ATOM | X | Y | Z |
|---|---|---|---|
| Fe | 0.2441180000 | -0.3154020000 | 1.0676530000 |
| C | -1.5058930000 | -2.5584460000 | -2.3150320000 |
| N | -1.7954970000 | -1.2376550000 | -2.6134210000 |
| H | -2.3372290000 | -0.9092510000 | -3.3993020000 |
| C | -1.2325030000 | -0.4551930000 | -1.6605110000 |
| H | -1.2929320000 | 0.6205010000 | -1.6150620000 |
| N | -0.5970070000 | -1.2053860000 | -0.7760320000 |
| C | -0.7598270000 | -2.5180640000 | -1.1683680000 |
| H | -0.3369940000 | -3.3280370000 | -0.5952530000 |
| H | -1.8514420000 | -3.3753620000 | -2.9285780000 |
| C | 2.2770370000 | -4.1029110000 | 1.5827830000 |
| N | 3.1307050000 | -3.4879590000 | 0.6830600000 |

| | | | |
|---|---|---|---|
| H | 4.0054180000 | -3.8610890000 | 0.3442890000 |
| C | 2.6188250000 | -2.2725590000 | 0.3693310000 |
| H | 3.0607280000 | -1.5543290000 | -0.3050000000 |
| N | 1.4810430000 | -2.0896570000 | 1.0179080000 |
| C | 1.2530170000 | -3.2166960000 | 1.7827180000 |
| H | 0.3791820000 | -3.2801120000 | 2.4121060000 |
| H | 2.4818990000 | -5.0784880000 | 1.9939210000 |
| F | -0.1635710000 | 1.3024140000 | 0.2287750000 |
| O | -1.0604540000 | -0.9942100000 | 2.1827560000 |
| H | -1.0681180000 | -0.5627280000 | 3.0525100000 |
| O | 2.3323260000 | 0.6437480000 | -0.4047520000 |
| H | 2.5658820000 | 0.8553600000 | 0.5202400000 |
| H | 1.5088840000 | 1.1634920000 | -0.5165790000 |
| N | 1.5763590000 | 0.4700550000 | 2.3761720000 |
| N | 2.1803300000 | -0.2327010000 | 3.1621070000 |
| N | 2.7743910000 | -0.8777520000 | 3.9125620000 |

**$^5Im_2WatOHN_3$_TS-1**
Charge: 0
Multiplicity: 5
B3LYP-D3(BJ)/6-31G(d)-SDD SCF energy (au): -1629.01692323
B3LYP-D3(BJ)/6-31G(d)-SDD enthalpy (au): -1628.49323323
B3LYP-D3(BJ)/6-31G(d)-SDD free energy (au): -1628.60409323
B3LYP-D3(BJ)/def2-TZVP/SMD SCF energy (au): -2769.51613764
B3LYP-D3(BJ)/def2-TZVP/SMD enthalpy (au): -2768.99244764
B3LYP-D3(BJ)/def2-TZVP/SMD free energy (au): -2769.10330764
B3LYP-D3(BJ)/def2-TZVP/SMD free energy (quasi-harmonic) (au): -2769.09345541

Cartesian coordinates

| ATOM | X | Y | Z |
|---|---|---|---|
| Fe | -0.2740400000 | -0.3815460000 | 0.8826810000 |
| C | 0.8902540000 | -1.6643240000 | -3.2748880000 |
| N | -0.4362860000 | -1.2809170000 | -3.3672020000 |
| H | -1.0204500000 | -1.3522220000 | -4.1871800000 |
| C | -0.8255700000 | -0.8080750000 | -2.1532750000 |
| H | -1.8215960000 | -0.4794550000 | -1.8974470000 |
| N | 0.1784520000 | -0.8649340000 | -1.2976360000 |
| C | 1.2525840000 | -1.4006140000 | -1.9796510000 |
| H | 2.1969400000 | -1.5799380000 | -1.4867300000 |
| H | 1.4295400000 | -2.0779200000 | -4.1125270000 |
| C | 0.8402240000 | -4.5945780000 | 1.4039860000 |
| N | 1.6112720000 | -3.9775430000 | 2.3742550000 |
| H | 2.2112880000 | -4.4375230000 | 3.0427950000 |
| C | 1.4057370000 | -2.6368150000 | 2.2899940000 |
| H | 1.8667010000 | -1.9106180000 | 2.9448430000 |
| N | 0.5491010000 | -2.3711790000 | 1.3191180000 |
| C | 0.1869930000 | -3.5780290000 | 0.7586150000 |
| H | -0.5206010000 | -3.6151350000 | -0.0552070000 |
| H | 0.8301830000 | -5.6652430000 | 1.2737140000 |
| C | 0.7227760000 | 7.1795560000 | 1.8879460000 |
| C | 0.5627830000 | 5.8850920000 | 1.3980600000 |
| C | 0.6233210000 | 4.7843690000 | 2.2596150000 |
| C | 0.8825100000 | 4.9550700000 | 3.6352200000 |

```
C      1.0158230000    6.2667460000    4.1052470000
C      0.9361940000    7.3687220000    3.2526390000
H      0.6804940000    8.0280560000    1.2114210000
H      0.3930020000    5.7157880000    0.3388340000
H      1.2001540000    6.4212250000    5.1653680000
H      1.0545350000    8.3714690000    3.6538370000
C      0.9972870000    3.7914890000    4.5963300000
C     -0.3489430000    3.1235520000    4.9168040000
H      1.6734950000    3.0360400000    4.1882400000
H      1.4474560000    4.1589830000    5.5257200000
H     -0.2047640000    2.3336280000    5.6599960000
H     -0.7757690000    2.6493120000    4.0289810000
H     -1.0714750000    3.8491130000    5.3093930000
C      0.5449000000    3.4061260000    1.6647490000
O      1.5319890000    2.6687840000    1.7630210000
N     -0.5554910000    3.1433450000    0.9184020000
C     -2.0138460000    3.3832280000    0.9789850000
C     -2.3365710000    4.6560400000    1.7873750000
C     -2.4920620000    3.5606000000   -0.4706090000
H     -3.4268040000    4.7538160000    1.7957420000
H     -1.9142500000    5.5569210000    1.3420040000
H     -1.9922170000    4.5819470000    2.8206030000
H     -2.2608170000    2.6678300000   -1.0558550000
H     -2.0072350000    4.4252050000   -0.9366100000
H     -3.5756330000    3.7170210000   -0.4857820000
F     -0.3593090000    1.6844020000    0.2227640000
C     -2.7305860000    2.1918310000    1.6384260000
H     -2.3024910000    1.9679850000    2.6176020000
H     -2.6569910000    1.2915230000    1.0254190000
H     -3.7882880000    2.4507520000    1.7623060000
O     -2.0714570000   -0.8005780000    0.4284790000
H     -2.5634940000   -0.9392180000    1.2527900000
O      1.8855360000    0.2197730000    0.8285920000
H      1.7908330000    1.1265630000    1.2354320000
H      1.9964500000    0.4059500000   -0.1171120000
N      1.4259350000    0.1139470000    4.4163780000
N      0.5641000000    0.1622260000    3.6355480000
N     -0.3450270000    0.2193550000    2.8489850000
```

**$^{5}Im_2WatOHN_3$_TS-3-F**

```
Charge: 0
Multiplicity: 5
B3LYP-D3(BJ)/6-31G(d)-SDD SCF energy (au):              -1629.07821112
B3LYP-D3(BJ)/6-31G(d)-SDD enthalpy (au):                -1628.55356812
B3LYP-D3(BJ)/6-31G(d)-SDD free energy (au):             -1628.66590712
B3LYP-D3(BJ)/def2-TZVP/SMD SCF energy (au):             -2769.59243396
B3LYP-D3(BJ)/def2-TZVP/SMD enthalpy (au):               -2769.06779096
B3LYP-D3(BJ)/def2-TZVP/SMD free energy (au):            -2769.18012996
B3LYP-D3(BJ)/def2-TZVP/SMD free energy (quasi-harmonic) (au): -2769.16876084

Cartesian coordinates
ATOM    X               Y               Z
H     34.5516760000   26.9842370000   39.2252420000
```

| | | | |
|---|---|---|---|
| C | 34.3047120000 | 27.5905760000 | 38.3681030000 |
| C | 34.7204640000 | 27.5580830000 | 37.0642830000 |
| N | 33.3902090000 | 28.6285040000 | 38.4359190000 |
| C | 33.2857320000 | 29.1803250000 | 37.2007210000 |
| N | 34.0782770000 | 28.5509360000 | 36.3503290000 |
| H | 32.9017710000 | 28.9414290000 | 39.2613950000 |
| H | 35.4070160000 | 26.8770520000 | 36.5872620000 |
| H | 32.6703630000 | 30.0274330000 | 36.9422620000 |
| H | 39.1377580000 | 30.2797810000 | 36.7022730000 |
| C | 38.3607320000 | 29.9237710000 | 36.0442270000 |
| C | 37.0610940000 | 30.3143290000 | 35.8610350000 |
| N | 38.5689660000 | 28.8687170000 | 35.1674060000 |
| C | 37.4109190000 | 28.6478680000 | 34.4949200000 |
| N | 36.4920400000 | 29.5137040000 | 34.8875000000 |
| H | 39.4177280000 | 28.3312110000 | 35.0708520000 |
| H | 36.4640410000 | 31.0829320000 | 36.3433530000 |
| H | 37.2534760000 | 27.8695050000 | 33.7630160000 |
| Fe | 34.4389190000 | 29.3806530000 | 34.2626490000 |
| O | 33.3181210000 | 30.7671190000 | 34.8716070000 |
| H | 32.9983820000 | 31.3862870000 | 34.1875040000 |
| F | 34.6567560000 | 30.0659130000 | 32.4558550000 |
| N | 33.4631360000 | 33.1028500000 | 30.8315420000 |
| O | 32.5252450000 | 32.7007390000 | 32.8715190000 |
| C | 34.8585230000 | 33.1881810000 | 32.8143500000 |
| C | 34.9373010000 | 34.1743420000 | 33.7954930000 |
| C | 36.1266690000 | 34.3999630000 | 34.4928030000 |
| C | 37.2372810000 | 33.5872340000 | 34.2364760000 |
| C | 37.1606750000 | 32.5785080000 | 33.2872910000 |
| C | 35.9849510000 | 32.3611880000 | 32.5305010000 |
| C | 33.5123260000 | 32.9659040000 | 32.1792180000 |
| C | 32.2955490000 | 32.7870330000 | 29.9753580000 |
| C | 31.8644850000 | 31.3304340000 | 30.2131100000 |
| C | 31.1445640000 | 33.7569280000 | 30.2862190000 |
| C | 32.7665470000 | 32.9682680000 | 28.5271740000 |
| C | 35.8968160000 | 31.2848240000 | 31.5799120000 |
| C | 37.0251810000 | 30.3555180000 | 31.2834520000 |
| H | 34.0511340000 | 34.7614550000 | 34.0163850000 |
| H | 36.1820590000 | 35.1897890000 | 35.2361620000 |
| H | 38.1596930000 | 33.7354160000 | 34.7902030000 |
| H | 38.0234880000 | 31.9443600000 | 33.1191600000 |
| H | 34.3048640000 | 33.4219930000 | 30.3725150000 |
| H | 35.1576970000 | 31.3804850000 | 30.7982890000 |
| H | 37.5444680000 | 30.0424270000 | 32.1915210000 |
| H | 36.6424020000 | 29.4618860000 | 30.7845940000 |
| H | 37.7603890000 | 30.8312690000 | 30.6159630000 |
| H | 31.0386620000 | 31.0741200000 | 29.5404100000 |
| H | 32.6961460000 | 30.6439780000 | 30.0233860000 |
| H | 31.5369990000 | 31.1882130000 | 31.2440750000 |
| H | 30.8416650000 | 33.6592430000 | 31.3304880000 |
| H | 31.4528010000 | 34.7925480000 | 30.1035450000 |
| H | 30.2835580000 | 33.5351830000 | 29.6459270000 |
| H | 33.5872570000 | 32.2804980000 | 28.2896400000 |
| H | 31.9436010000 | 32.7595660000 | 27.8374690000 |

H 33.1052920000 33.9959360000 28.3455400000
N 34.5633160000 27.5086060000 33.5333090000
N 34.3921720000 26.4498480000 34.0914720000
N 34.2554510000 25.4109630000 34.5847760000
O 34.7320820000 32.2045770000 36.6781000000
H 35.0185510000 32.9527020000 36.1322270000
H 34.1632420000 31.6837510000 36.0469750000

**$^{5}Im_{2}WatOHN_{3}$_TS-3-$N_{3}$**

Charge: 0
Multiplicity: 5
B3LYP-D3(BJ)/6-31G(d)-SDD SCF energy (au): -1629.09315762
B3LYP-D3(BJ)/6-31G(d)-SDD enthalpy (au): -1628.56910662
B3LYP-D3(BJ)/6-31G(d)-SDD free energy (au): -1628.67823962
B3LYP-D3(BJ)/def2-TZVP/SMD SCF energy (au): -2769.59759527
B3LYP-D3(BJ)/def2-TZVP/SMD enthalpy (au): -2769.07354427
B3LYP-D3(BJ)/def2-TZVP/SMD free energy (au): -2769.18267727
B3LYP-D3(BJ)/def2-TZVP/SMD free energy (quasi-harmonic) (au): -2769.17356065

Cartesian coordinates

| ATOM | X | Y | Z |
|---|---|---|---|
| H | 32.2485150000 | 29.0967350000 | 38.5298990000 |
| C | 32.8013170000 | 29.1882470000 | 37.6082990000 |
| C | 33.9894600000 | 29.7980780000 | 37.3027230000 |
| N | 32.3505650000 | 28.6383080000 | 36.4201610000 |
| C | 33.2536930000 | 28.9344080000 | 35.4490090000 |
| N | 34.2559770000 | 29.6280910000 | 35.9599270000 |
| H | 31.4798820000 | 28.1474090000 | 36.2834560000 |
| H | 34.6551840000 | 30.3616450000 | 37.9367650000 |
| H | 33.1707840000 | 28.6689750000 | 34.4056640000 |
| H | 39.3626430000 | 26.8577750000 | 36.4035540000 |
| C | 38.6036030000 | 27.3915180000 | 35.8533660000 |
| C | 37.9746790000 | 28.5888400000 | 36.0711200000 |
| N | 38.0642730000 | 26.8987290000 | 34.6758670000 |
| C | 37.1468880000 | 27.7955670000 | 34.2266920000 |
| N | 37.0750290000 | 28.8219450000 | 35.0528560000 |
| H | 38.3145000000 | 26.0326870000 | 34.2229080000 |
| H | 38.0923570000 | 29.3068060000 | 36.8672690000 |
| H | 36.5524320000 | 27.7003030000 | 33.3304980000 |
| Fe | 35.7528110000 | 30.5254250000 | 34.6103930000 |
| O | 35.0065850000 | 29.5762020000 | 33.1367840000 |
| H | 35.2157630000 | 30.0589240000 | 32.3219280000 |
| F | 37.1737480000 | 31.6156840000 | 33.9625790000 |
| N | 33.7241630000 | 34.7434180000 | 36.7731860000 |
| O | 35.0237450000 | 33.1339850000 | 37.7353860000 |
| C | 36.0304420000 | 34.4816420000 | 36.0496130000 |
| C | 37.2193780000 | 34.7546620000 | 36.7332190000 |
| C | 38.3880480000 | 35.0738360000 | 36.0467340000 |
| C | 38.3772040000 | 35.0718860000 | 34.6472120000 |
| C | 37.2159180000 | 34.7598390000 | 33.9574110000 |
| C | 36.0026630000 | 34.4782770000 | 34.6262630000 |
| C | 34.8786220000 | 34.0514010000 | 36.9080900000 |
| C | 32.4935690000 | 34.5555670000 | 37.5768290000 |

| | | | |
|---|---|---|---|
| C | 31.9738600000 | 33.1178530000 | 37.4280040000 |
| C | 32.7904130000 | 34.8773670000 | 39.0512870000 |
| C | 31.4632900000 | 35.5417690000 | 37.0127270000 |
| C | 34.8222540000 | 34.1014610000 | 33.8787810000 |
| C | 34.8787380000 | 33.8524680000 | 32.4030630000 |
| H | 37.2172400000 | 34.7007870000 | 37.8177200000 |
| H | 39.2986010000 | 35.3000910000 | 36.5930510000 |
| H | 39.2853820000 | 35.2951940000 | 34.0946590000 |
| H | 37.2382890000 | 34.7238440000 | 32.8748020000 |
| H | 33.7646680000 | 35.5640510000 | 36.1845170000 |
| H | 33.8639270000 | 34.4263030000 | 34.2686590000 |
| H | 35.6984690000 | 33.1673890000 | 32.1668780000 |
| H | 33.9401940000 | 33.4089270000 | 32.0563920000 |
| H | 35.0307350000 | 34.7844310000 | 31.8366030000 |
| H | 31.0702340000 | 32.9948910000 | 38.0363700000 |
| H | 31.7209690000 | 32.8973190000 | 36.3874740000 |
| H | 32.7217610000 | 32.3978090000 | 37.7611490000 |
| H | 33.5519350000 | 34.1973990000 | 39.4393630000 |
| H | 33.1511860000 | 35.9067720000 | 39.1577070000 |
| H | 31.8805150000 | 34.7679390000 | 39.6518830000 |
| H | 31.2555750000 | 35.3285360000 | 35.9578340000 |
| H | 30.5233740000 | 35.4583180000 | 37.5663090000 |
| H | 31.8134500000 | 36.5781890000 | 37.1010660000 |
| N | 34.4478700000 | 32.1943680000 | 34.5982810000 |
| N | 33.2727800000 | 31.9731500000 | 34.3782190000 |
| N | 32.1487430000 | 31.8089510000 | 34.1691060000 |
| O | 36.7579500000 | 31.3497360000 | 36.5136680000 |
| H | 36.1984360000 | 32.0128040000 | 36.9848270000 |
| H | 37.2660570000 | 31.8471840000 | 35.8367670000 |

**$^5$Im$_2$WatOHN$_3$_TS-3-OH**

Charge: 0
Multiplicity: 5
B3LYP-D3(BJ)/6-31G(d)-SDD SCF energy (au): -1629.09196123
B3LYP-D3(BJ)/6-31G(d)-SDD enthalpy (au): -1628.56780423
B3LYP-D3(BJ)/6-31G(d)-SDD free energy (au): -1628.67675223
B3LYP-D3(BJ)/def2-TZVP/SMD SCF energy (au): -2769.59671465
B3LYP-D3(BJ)/def2-TZVP/SMD enthalpy (au): -2769.07255765
B3LYP-D3(BJ)/def2-TZVP/SMD free energy (au): -2769.18150565
B3LYP-D3(BJ)/def2-TZVP/SMD free energy (quasi-harmonic) (au): -2769.17243342

Cartesian coordinates

| ATOM | X | Y | Z |
|---|---|---|---|
| H | 32.9095350000 | 28.9017090000 | 39.1798000000 |
| C | 33.2167940000 | 29.0994710000 | 38.1650650000 |
| C | 34.4104880000 | 29.4980910000 | 37.6230630000 |
| N | 32.3501580000 | 28.9723350000 | 37.0945380000 |
| C | 33.0310610000 | 29.2909580000 | 35.9611030000 |
| N | 34.2792540000 | 29.6114900000 | 36.2544670000 |
| H | 31.3796590000 | 28.6994760000 | 37.1406190000 |
| H | 35.3466620000 | 29.7243670000 | 38.1075360000 |
| H | 32.5812520000 | 29.3070710000 | 34.9783670000 |
| H | 39.1100670000 | 26.5652860000 | 36.4185160000 |

| | | | |
|---|---|---|---|
| C | 38.3713220000 | 27.1497520000 | 35.8931210000 |
| C | 37.8087320000 | 28.3717360000 | 36.1486790000 |
| N | 37.7890800000 | 26.7139760000 | 34.7145190000 |
| C | 36.9125300000 | 27.6653630000 | 34.3004060000 |
| N | 36.9044020000 | 28.6738280000 | 35.1516200000 |
| H | 37.9905570000 | 25.8497140000 | 34.2342860000 |
| H | 37.9811490000 | 29.0642300000 | 36.9574320000 |
| H | 36.3116920000 | 27.6142200000 | 33.4050150000 |
| Fe | 35.7982790000 | 30.5469480000 | 34.8275600000 |
| O | 34.6225600000 | 32.0490560000 | 34.8498110000 |
| H | 33.7004660000 | 31.8201700000 | 34.6486110000 |
| F | 37.3575400000 | 31.3974800000 | 34.2431270000 |
| N | 33.7425540000 | 34.5880980000 | 36.7138580000 |
| O | 35.1215880000 | 33.1159690000 | 37.7633270000 |
| C | 36.0619170000 | 34.4183910000 | 35.9996270000 |
| C | 37.2578540000 | 34.7167880000 | 36.6585340000 |
| C | 38.4125460000 | 35.0438150000 | 35.9510500000 |
| C | 38.3795810000 | 35.0273100000 | 34.5516180000 |
| C | 37.2127060000 | 34.6925970000 | 33.8832190000 |
| C | 36.0117400000 | 34.3942170000 | 34.5722970000 |
| C | 34.9354390000 | 33.9724570000 | 36.8785990000 |
| C | 32.4755580000 | 34.2133890000 | 37.3783180000 |
| C | 32.1306760000 | 32.7478710000 | 37.0614260000 |
| C | 32.5977800000 | 34.4254330000 | 38.8963310000 |
| C | 31.3983080000 | 35.1413530000 | 36.8030260000 |
| C | 34.8486540000 | 33.9744440000 | 33.8429370000 |
| C | 34.8961290000 | 33.5725310000 | 32.4049160000 |
| H | 37.2722670000 | 34.6771510000 | 37.7437110000 |
| H | 39.3276880000 | 35.2904580000 | 36.4807770000 |
| H | 39.2755250000 | 35.2602330000 | 33.9828900000 |
| H | 37.2159710000 | 34.6504110000 | 32.7999010000 |
| H | 33.7080200000 | 35.3332880000 | 36.0331450000 |
| H | 33.8724370000 | 34.1655020000 | 34.2655120000 |
| H | 35.7465810000 | 32.9084610000 | 32.2190100000 |
| H | 33.9830310000 | 33.0360210000 | 32.1308960000 |
| H | 34.9876080000 | 34.4402410000 | 31.7311820000 |
| H | 31.2025560000 | 32.4686890000 | 37.5736100000 |
| H | 31.9817570000 | 32.6138920000 | 35.9851740000 |
| H | 32.9312900000 | 32.0830660000 | 37.3887560000 |
| H | 33.3831580000 | 33.7868570000 | 39.3039830000 |
| H | 32.8446870000 | 35.4695950000 | 39.1177130000 |
| H | 31.6480350000 | 34.1839090000 | 39.3872060000 |
| H | 31.3028380000 | 35.0105160000 | 35.7182240000 |
| H | 30.4278730000 | 34.9164730000 | 37.2555370000 |
| H | 31.6314500000 | 36.1928890000 | 37.0097920000 |
| N | 34.9073940000 | 29.6146830000 | 33.2004690000 |
| N | 33.7958160000 | 29.9901410000 | 32.9297330000 |
| N | 32.7128720000 | 30.3471240000 | 32.6939430000 |
| O | 36.7916040000 | 31.2220030000 | 36.7591780000 |
| H | 36.2268370000 | 31.9354930000 | 37.1610350000 |
| H | 37.4110070000 | 31.6792230000 | 36.1532460000 |

**$^{5}Im_2Wat_2N_3$_11**

Charge: 1
Multiplicity: 5
B3LYP-D3(BJ)/6-31G(d)-SDD SCF energy (au): -1629.47073885
B3LYP-D3(BJ)/6-31G(d)-SDD enthalpy (au): -1628.93062285
B3LYP-D3(BJ)/6-31G(d)-SDD free energy (au): -1629.04444785
B3LYP-D3(BJ)/def2-TZVP/SMD SCF energy (au): -2770.00736913
B3LYP-D3(BJ)/def2-TZVP/SMD enthalpy (au): -2769.46725313
B3LYP-D3(BJ)/def2-TZVP/SMD free energy (au): -2769.58107813
B3LYP-D3(BJ)/def2-TZVP/SMD free energy (quasi-harmonic) (au): -2769.56939928

Cartesian coordinates

| ATOM | X | Y | Z |
|---|---|---|---|
| Fe | 0.4164240000 | -0.5069930000 | -0.4839440000 |
| C | -0.4352850000 | 1.4290570000 | 3.3657690000 |
| N | 0.6445410000 | 2.1345630000 | 2.8699130000 |
| H | 1.0789520000 | 2.9378140000 | 3.3027840000 |
| C | 1.0186800000 | 1.5716840000 | 1.6939990000 |
| H | 1.8152700000 | 1.9296630000 | 1.0592760000 |
| N | 0.2342060000 | 0.5412810000 | 1.4136520000 |
| C | -0.6791830000 | 0.4416860000 | 2.4475470000 |
| H | -1.4537960000 | -0.3092370000 | 2.4392520000 |
| H | -0.9164100000 | 1.6902190000 | 4.2951410000 |
| C | 3.2052290000 | -3.6712660000 | 0.3192830000 |
| N | 2.8007430000 | -3.3537210000 | 1.6026290000 |
| H | 3.1599780000 | -3.7617960000 | 2.4548440000 |
| C | 1.8495140000 | -2.3946570000 | 1.5250750000 |
| H | 1.3669610000 | -1.9441610000 | 2.3791020000 |
| N | 1.6155500000 | -2.0869170000 | 0.2558150000 |
| C | 2.4595680000 | -2.8769100000 | -0.5079400000 |
| H | 2.4829160000 | -2.7900210000 | -1.5832790000 |
| H | 3.9691140000 | -4.4067120000 | 0.1224990000 |
| O | 0.3022350000 | -1.5457560000 | -2.4505490000 |
| H | -0.4024980000 | -1.0156840000 | -2.8968620000 |
| H | 0.0753120000 | -2.4792240000 | -2.5809000000 |
| C | -5.5685140000 | 2.5709640000 | -2.7543270000 |
| C | -4.4574390000 | 1.9180270000 | -3.2787190000 |
| C | -3.2513620000 | 1.9068240000 | -2.5670520000 |
| C | -3.1338250000 | 2.5303590000 | -1.3131790000 |
| C | -4.2675640000 | 3.1800490000 | -0.8079390000 |
| C | -5.4685680000 | 3.2053840000 | -1.5141950000 |
| H | -6.5031770000 | 2.5828210000 | -3.3056410000 |
| H | -4.5174310000 | 1.4160440000 | -4.2399580000 |
| H | -4.2058820000 | 3.6737210000 | 0.1577490000 |
| H | -6.3294810000 | 3.7172360000 | -1.0950150000 |
| C | -1.8380460000 | 2.5286630000 | -0.5284240000 |
| C | -1.3054610000 | 3.9341130000 | -0.2130010000 |
| H | -1.0581090000 | 1.9831770000 | -1.0678060000 |
| H | -1.9902660000 | 1.9848220000 | 0.4118850000 |
| H | -0.3606330000 | 3.8612000000 | 0.3351910000 |
| H | -1.1187290000 | 4.5027520000 | -1.1300540000 |
| H | -2.0086450000 | 4.5095810000 | 0.3974120000 |
| C | -2.1195620000 | 1.1019600000 | -3.1114980000 |
| O | -1.9119190000 | -0.0572760000 | -2.6834630000 |

| | | | |
|---|---|---|---|
| N | -1.3836680000 | 1.6013070000 | -4.1205460000 |
| C | -1.0304780000 | 2.9572550000 | -4.6641010000 |
| C | -2.1475900000 | 3.9732730000 | -4.4027660000 |
| C | -0.8456170000 | 2.7857530000 | -6.1801410000 |
| H | -1.8423930000 | 4.9036620000 | -4.8905910000 |
| H | -3.1005920000 | 3.6612030000 | -4.8340630000 |
| H | -2.2926680000 | 4.1805460000 | -3.3423510000 |
| H | -0.0484170000 | 2.0769740000 | -6.4093770000 |
| H | -1.7733120000 | 2.4416910000 | -6.6480680000 |
| H | -0.5789670000 | 3.7537000000 | -6.6135100000 |
| F | -0.3743640000 | 0.7019470000 | -4.4842920000 |
| C | 0.2745590000 | 3.4046480000 | -3.9891150000 |
| H | 0.1496310000 | 3.4660730000 | -2.9037870000 |
| H | 1.0930890000 | 2.7164610000 | -4.2069390000 |
| H | 0.5478890000 | 4.3958010000 | -4.3634690000 |
| O | -1.6911850000 | -1.0033380000 | -0.2381920000 |
| H | -1.9030700000 | -1.9504570000 | -0.2643090000 |
| H | -2.0565980000 | -0.6218530000 | -1.0843170000 |
| N | 2.6547410000 | 0.9853620000 | -3.2828160000 |
| N | 2.0206010000 | 0.9657710000 | -2.3224490000 |
| N | 1.3554580000 | 1.0058190000 | -1.3036040000 |

**$^{5}$Im$_2$Wat$_2$N$_3$_13**

Charge: 1
Multiplicity: 5
B3LYP-D3(BJ)/6-31G(d)-SDD SCF energy (au): -1629.53845012
B3LYP-D3(BJ)/6-31G(d)-SDD enthalpy (au): -1628.99961012
B3LYP-D3(BJ)/6-31G(d)-SDD free energy (au): -1629.11278012
B3LYP-D3(BJ)/def2-TZVP/SMD SCF energy (au): -2770.07343488
B3LYP-D3(BJ)/def2-TZVP/SMD enthalpy (au): -2769.53459488
B3LYP-D3(BJ)/def2-TZVP/SMD free energy (au): -2769.64776488
B3LYP-D3(BJ)/def2-TZVP/SMD free energy (quasi-harmonic) (au): -2769.63701376

Cartesian coordinates

| ATOM | X | Y | Z |
|---|---|---|---|
| H | 33.4033180000 | 26.6164660000 | 38.0126420000 |
| C | 33.5899820000 | 27.4377980000 | 37.3392870000 |
| C | 34.7455910000 | 28.0402430000 | 36.9266870000 |
| N | 32.5695630000 | 28.1147060000 | 36.6981990000 |
| C | 33.1103750000 | 29.0845780000 | 35.9289530000 |
| N | 34.4348910000 | 29.0665840000 | 36.0493780000 |
| H | 31.5821900000 | 27.9123920000 | 36.7736990000 |
| H | 35.7661990000 | 27.8156830000 | 37.1899110000 |
| H | 32.5377040000 | 29.7619180000 | 35.3109760000 |
| H | 39.9780710000 | 27.2139440000 | 36.4704520000 |
| C | 39.1554590000 | 27.6459140000 | 35.9229140000 |
| C | 38.2259880000 | 28.5909190000 | 36.2600600000 |
| N | 38.8618960000 | 27.2807700000 | 34.6221060000 |
| C | 37.7893720000 | 27.9911110000 | 34.2089480000 |
| N | 37.3782930000 | 28.7917740000 | 35.1836960000 |
| H | 39.3684370000 | 26.6050470000 | 34.0663660000 |
| H | 38.1011470000 | 29.1414790000 | 37.1787210000 |
| H | 37.3472250000 | 27.9259810000 | 33.2270740000 |

| | | | |
|---|---|---|---|
| Fe | 35.8517670000 | 30.2552010000 | 34.9147350000 |
| O | 34.4666400000 | 31.8707970000 | 35.2310040000 |
| H | 34.8795040000 | 32.5657880000 | 34.6838090000 |
| H | 33.5512140000 | 31.7385230000 | 34.8235110000 |
| F | 37.0018320000 | 31.5944780000 | 34.4365990000 |
| N | 31.2205380000 | 31.3983230000 | 31.9753870000 |
| O | 32.1556110000 | 31.4194870000 | 34.0545670000 |
| C | 33.3256090000 | 32.5433750000 | 32.3018430000 |
| C | 33.5723110000 | 33.7622860000 | 32.9367490000 |
| C | 34.6531440000 | 34.5636470000 | 32.5629060000 |
| C | 35.5086740000 | 34.1246880000 | 31.5415680000 |
| C | 35.2799130000 | 32.9175780000 | 30.9042670000 |
| C | 34.1840750000 | 32.0754010000 | 31.2537890000 |
| C | 32.1892810000 | 31.7304310000 | 32.8372830000 |
| C | 29.9619110000 | 30.6570630000 | 32.2637580000 |
| C | 30.2871920000 | 29.2484460000 | 32.7803260000 |
| C | 29.1278810000 | 31.4439120000 | 33.2862540000 |
| C | 29.2169920000 | 30.5635810000 | 30.9261820000 |
| C | 34.0073280000 | 30.8164290000 | 30.6314130000 |
| C | 34.9355700000 | 30.2530430000 | 29.6053400000 |
| H | 32.9002380000 | 34.0894430000 | 33.7257470000 |
| H | 34.8208520000 | 35.5173920000 | 33.0524460000 |
| H | 36.3575740000 | 34.7343050000 | 31.2478130000 |
| H | 35.9616550000 | 32.5900670000 | 30.1278010000 |
| H | 31.3330930000 | 31.7512280000 | 31.0335000000 |
| H | 33.1930370000 | 30.1839770000 | 30.9657720000 |
| H | 35.9308430000 | 30.0634790000 | 30.0352460000 |
| H | 34.5566220000 | 29.3053670000 | 29.2147240000 |
| H | 35.0828850000 | 30.9318990000 | 28.7547860000 |
| H | 29.3566710000 | 28.6888470000 | 32.9209620000 |
| H | 30.9159800000 | 28.7023850000 | 32.0716180000 |
| H | 30.8124050000 | 29.2920440000 | 33.7359560000 |
| H | 29.6693420000 | 31.5516210000 | 34.2286510000 |
| H | 28.8880120000 | 32.4414630000 | 32.9040010000 |
| H | 28.1879470000 | 30.9171400000 | 33.4802710000 |
| H | 29.8053590000 | 30.0127120000 | 30.1832330000 |
| H | 28.2699510000 | 30.0350790000 | 31.0640140000 |
| H | 28.9893930000 | 31.5596410000 | 30.5288000000 |
| N | 35.2356920000 | 29.5375280000 | 33.2229470000 |
| N | 34.3460770000 | 28.7367240000 | 33.0251240000 |
| N | 33.5019030000 | 27.9885760000 | 32.7911770000 |
| O | 36.4636160000 | 30.9706750000 | 36.9759860000 |
| H | 35.6539710000 | 31.4476320000 | 37.2327780000 |
| H | 37.0573750000 | 31.6572330000 | 36.6149180000 |

**$^{5}Im_2Wat_2N_3$_FeF_remove-F**
Charge: 1
Multiplicity: 5
B3LYP-D3(BJ)/6-31G(d)-SDD SCF energy (au): -893.385801270
B3LYP-D3(BJ)/6-31G(d)-SDD enthalpy (au): -893.154772270
B3LYP-D3(BJ)/6-31G(d)-SDD free energy (au): -893.229899270
B3LYP-D3(BJ)/def2-TZVP/SMD SCF energy (au): -2033.63568239
B3LYP-D3(BJ)/def2-TZVP/SMD enthalpy (au): -2033.40465339

B3LYP-D3(BJ)/def2-TZVP/SMD free energy (au):            -2033.47978039
B3LYP-D3(BJ)/def2-TZVP/SMD free energy (quasi-harmonic) (au): -2033.472277120

Cartesian coordinates

| ATOM | X | Y | Z |
|---|---|---|---|
| Fe | 0.1271510000 | -1.2015290000 | 0.2586320000 |
| C | -1.3820120000 | -2.2231640000 | -3.6200950000 |
| N | -2.4984590000 | -1.8281660000 | -2.9069170000 |
| H | -3.4470200000 | -1.8008350000 | -3.2573070000 |
| C | -2.1155460000 | -1.4843960000 | -1.6600660000 |
| H | -2.7786460000 | -1.1319070000 | -0.8833900000 |
| N | -0.7981470000 | -1.6367850000 | -1.5365300000 |
| C | -0.3297930000 | -2.0993810000 | -2.7562070000 |
| H | 0.7163700000 | -2.3084000000 | -2.9187580000 |
| H | -1.4392410000 | -2.5466100000 | -4.6472820000 |
| C | 2.4894410000 | -4.4346460000 | 1.7166770000 |
| N | 3.4515630000 | -3.4934660000 | 1.3995610000 |
| H | 4.4475690000 | -3.5928320000 | 1.5445460000 |
| C | 2.8401500000 | -2.4109360000 | 0.8764960000 |
| H | 3.3484570000 | -1.5180460000 | 0.5466930000 |
| N | 1.5245360000 | -2.6094680000 | 0.8387310000 |
| C | 1.2949810000 | -3.8731540000 | 1.3630720000 |
| H | 0.2976120000 | -4.2755230000 | 1.4528420000 |
| H | 2.7384670000 | -5.3879760000 | 2.1549450000 |
| O | 2.0076080000 | 1.0143520000 | 2.6111760000 |
| H | 1.1654790000 | 1.0660950000 | 3.1132710000 |
| H | 2.5257750000 | 1.7890870000 | 2.8794500000 |
| O | 1.5376780000 | 0.3707140000 | 0.1117980000 |
| H | 1.7628220000 | 0.7111320000 | 1.0364000000 |
| H | 1.2411010000 | 1.1374550000 | -0.4045760000 |
| N | -1.2222630000 | -0.4487890000 | 1.4286090000 |
| N | -0.9946790000 | 0.1878100000 | 2.4322080000 |
| N | -0.7902300000 | 0.8124560000 | 3.3830540000 |

**$^5Im_2Wat_2N_3$_FeF_remove-$N_3$**
Charge: 1
Multiplicity: 5
B3LYP-D3(BJ)/6-31G(d)-SDD SCF energy (au):              -829.055172156
B3LYP-D3(BJ)/6-31G(d)-SDD enthalpy (au):                -828.836857156
B3LYP-D3(BJ)/6-31G(d)-SDD free energy (au):             -828.905716156
B3LYP-D3(BJ)/def2-TZVP/SMD SCF energy (au):             -1969.28221291
B3LYP-D3(BJ)/def2-TZVP/SMD enthalpy (au):               -1969.06389791
B3LYP-D3(BJ)/def2-TZVP/SMD free energy (au):            -1969.13275691
B3LYP-D3(BJ)/def2-TZVP/SMD free energy (quasi-harmonic) (au): -1969.126740754

Cartesian coordinates

| ATOM | X | Y | Z |
|---|---|---|---|
| Fe | 0.6350380000 | -0.9463530000 | 0.4958530000 |
| C | -1.5548860000 | -2.1070060000 | -3.0232960000 |
| N | -2.3472680000 | -1.1324680000 | -2.4486240000 |
| H | -3.2341410000 | -0.8020770000 | -2.8055300000 |
| C | -1.7421950000 | -0.6916470000 | -1.3245100000 |
| H | -2.1358120000 | 0.0731180000 | -0.6712620000 |

| | | | |
|---|---|---|---|
| N | -0.5919080000 | -1.3367530000 | -1.1498880000 |
| C | -0.4648750000 | -2.2249470000 | -2.2053940000 |
| H | 0.3891870000 | -2.8786750000 | -2.2953600000 |
| H | -1.8322020000 | -2.6120100000 | -3.9349520000 |
| C | 2.3624700000 | -4.6685870000 | 1.6620500000 |
| N | 3.4664670000 | -3.9149430000 | 1.3085500000 |
| H | 4.4307540000 | -4.2161030000 | 1.3613910000 |
| C | 3.0476310000 | -2.7035850000 | 0.8896120000 |
| H | 3.6917520000 | -1.9047240000 | 0.5550820000 |
| N | 1.7188680000 | -2.6400550000 | 0.9564790000 |
| C | 1.2803990000 | -3.8646120000 | 1.4391400000 |
| H | 0.2323930000 | -4.0723120000 | 1.5919970000 |
| H | 2.4478930000 | -5.6780830000 | 2.0316870000 |
| F | -0.2457570000 | 0.3855130000 | 1.4029430000 |
| O | 1.6918800000 | 2.1715570000 | 1.3661230000 |
| H | 0.8312480000 | 1.7311240000 | 1.5633790000 |
| H | 2.0951670000 | 2.3905140000 | 2.2200910000 |
| O | 2.3526800000 | 0.1688460000 | -0.1216050000 |
| H | 2.2916290000 | 1.0039670000 | 0.4579660000 |
| H | 2.3160460000 | 0.4863410000 | -1.0388620000 |

**$^{6}$Im$_{2}$Wat$_{2}$N$_{3}$_FeF**
Charge: 1
Multiplicity: 6
B3LYP-D3(BJ)/6-31G(d)-SDD SCF energy (au): -993.242139814
B3LYP-D3(BJ)/6-31G(d)-SDD enthalpy (au): -993.007251814
B3LYP-D3(BJ)/6-31G(d)-SDD free energy (au): -993.083283814
B3LYP-D3(BJ)/def2-TZVP/SMD SCF energy (au): -2133.53069231
B3LYP-D3(BJ)/def2-TZVP/SMD enthalpy (au): -2133.29580431
B3LYP-D3(BJ)/def2-TZVP/SMD free energy (au): -2133.37183631
B3LYP-D3(BJ)/def2-TZVP/SMD free energy (quasi-harmonic) (au): -2133.366455496

Cartesian coordinates

| ATOM | X | Y | Z |
|---|---|---|---|
| Fe | 0.3704350000 | -0.4675560000 | 0.7344300000 |
| C | -1.3784400000 | -2.5370990000 | -2.6119120000 |
| N | -1.9516250000 | -1.2804510000 | -2.6794160000 |
| H | -2.6610550000 | -0.9864250000 | -3.3377040000 |
| C | -1.4186450000 | -0.5099220000 | -1.7082070000 |
| H | -1.6855640000 | 0.5164180000 | -1.5078480000 |
| N | -0.5233810000 | -1.2145080000 | -1.0220090000 |
| C | -0.4910550000 | -2.4830760000 | -1.5730950000 |
| H | 0.1527880000 | -3.2523630000 | -1.1778380000 |
| H | -1.6491070000 | -3.3303930000 | -3.2905100000 |
| C | 2.3441880000 | -4.1860760000 | 1.6156170000 |
| N | 3.2309110000 | -3.6545610000 | 0.6959740000 |
| H | 4.0751070000 | -4.0995590000 | 0.3605130000 |
| C | 2.7888230000 | -2.4341750000 | 0.3256080000 |
| H | 3.2775740000 | -1.7914470000 | -0.3891440000 |
| N | 1.6565550000 | -2.1539620000 | 0.9652650000 |
| C | 1.3668720000 | -3.2436870000 | 1.7741940000 |
| H | 0.4824190000 | -3.2520660000 | 2.3912610000 |
| H | 2.4890330000 | -5.1582010000 | 2.0597030000 |

```
F     -0.2875800000    1.2092980000    0.3985200000
O      1.6517000000    0.6473280000    2.2164980000
H      1.0790760000    1.4154340000    2.3992830000
H      1.7348000000    0.1641760000    3.0567380000
O      2.1567220000    0.3053270000   -0.4978200000
H      2.5914790000    0.8514210000    0.1825640000
H      1.7304280000    0.9421660000   -1.0976070000
N     -0.8413930000   -1.1317530000    2.0594440000
N     -1.9735720000   -0.7274640000    2.2806730000
N     -3.0428760000   -0.3878450000    2.5148350000
```

**$^5Im_2Wat_2N_3$_TS-1**

```
Charge: 1
Multiplicity: 5
B3LYP-D3(BJ)/6-31G(d)-SDD SCF energy (au):              -1629.44760838
B3LYP-D3(BJ)/6-31G(d)-SDD enthalpy (au):                -1628.91108338
B3LYP-D3(BJ)/6-31G(d)-SDD free energy (au):             -1629.02372938
B3LYP-D3(BJ)/def2-TZVP/SMD SCF energy (au):             -2769.99195039
B3LYP-D3(BJ)/def2-TZVP/SMD enthalpy (au):               -2769.45542539
B3LYP-D3(BJ)/def2-TZVP/SMD free energy (au):            -2769.56807139
B3LYP-D3(BJ)/def2-TZVP/SMD free energy (quasi-harmonic) (au): -2769.55753301

Cartesian coordinates
ATOM    X               Y               Z
Fe     0.3498840000   -0.4676800000    0.8439300000
C     -1.1143950000   -2.5015960000   -2.7524730000
N     -2.0023940000   -1.4506870000   -2.6205460000
H     -2.8485910000   -1.3183440000   -3.1573710000
C     -1.5595480000   -0.6425840000   -1.6287070000
H     -2.0702730000    0.2468310000   -1.2957990000
N     -0.4310420000   -1.1175240000   -1.1189850000
C     -0.1454630000   -2.2809140000   -1.8114850000
H      0.7225260000   -2.8750950000   -1.5732050000
H     -1.2543800000   -3.2844840000   -3.4809910000
C      2.2383680000   -4.2611350000    1.4910660000
N      3.2449410000   -3.6047170000    0.8071650000
H      4.1491450000   -3.9888470000    0.5694660000
C      2.8181250000   -2.3500580000    0.5215540000
H      3.3960990000   -1.6089480000   -0.0095890000
N      1.5885980000   -2.1774760000    0.9875480000
C      1.2124620000   -3.3607480000    1.5966150000
H      0.2413300000   -3.4546730000    2.0579470000
H      2.3446870000   -5.2788440000    1.8319590000
C      1.5268730000    6.4272680000    3.3463200000
C      1.3987260000    5.4103800000    2.4129760000
C      0.6007510000    4.2837720000    2.6951950000
C     -0.0448190000    4.1353970000    3.9442600000
C      0.1210320000    5.1749260000    4.8709420000
C      0.8747120000    6.3083530000    4.5798570000
H      2.1323700000    7.3001200000    3.1248870000
H      1.9029280000    5.4712370000    1.4536370000
H     -0.3457020000    5.0791090000    5.8473860000
H      0.9705680000    7.0958140000    5.3213460000
```

| | | | |
|---|---|---|---|
| C | -0.8263310000 | 2.9071180000 | 4.3648280000 |
| C | -2.3403210000 | 3.1503830000 | 4.4486610000 |
| H | -0.6318640000 | 2.0679090000 | 3.6952070000 |
| H | -0.4619310000 | 2.6084080000 | 5.3559030000 |
| H | -2.8501780000 | 2.2444720000 | 4.7862690000 |
| H | -2.7460000000 | 3.4100670000 | 3.4670260000 |
| H | -2.5764610000 | 3.9651590000 | 5.1417320000 |
| C | 0.5089240000 | 3.2715530000 | 1.6116990000 |
| O | 1.5217940000 | 2.7903040000 | 1.0780030000 |
| N | -0.7916590000 | 2.9252380000 | 1.3475410000 |
| C | -1.6701900000 | 3.6052290000 | 0.3857010000 |
| C | -1.8998520000 | 5.0453260000 | 0.9203420000 |
| C | -1.0228740000 | 3.6725810000 | -1.0077820000 |
| H | -2.6467740000 | 5.5162500000 | 0.2741680000 |
| H | -0.9848320000 | 5.6397520000 | 0.8896740000 |
| H | -2.2782860000 | 5.0343470000 | 1.9440230000 |
| H | -0.8583130000 | 2.6672040000 | -1.4018890000 |
| H | -0.0628980000 | 4.1976220000 | -0.9754520000 |
| H | -1.6845320000 | 4.2132500000 | -1.6912050000 |
| F | -0.6235490000 | 1.2295260000 | 0.6014670000 |
| C | -3.0093370000 | 2.8592150000 | 0.3609960000 |
| H | -3.4365710000 | 2.7914860000 | 1.3643000000 |
| H | -2.8792130000 | 1.8415730000 | -0.0087830000 |
| H | -3.7111880000 | 3.3893620000 | -0.2896970000 |
| O | 1.5048800000 | 0.4505990000 | 2.5274850000 |
| H | 2.1830220000 | -0.1369790000 | 2.8955500000 |
| H | 1.9401610000 | 1.2867470000 | 2.2736650000 |
| O | 1.8678130000 | 0.5394240000 | -0.3691160000 |
| H | 1.7415890000 | 1.4668540000 | -0.0348960000 |
| H | 1.4490180000 | 0.5085020000 | -1.2457920000 |
| N | -1.0586240000 | -1.1925670000 | 1.9932920000 |
| N | -1.9580000000 | -0.4932430000 | 2.4184590000 |
| N | -2.8292160000 | 0.1312770000 | 2.8402840000 |

**$^{5}Im_2Wat_2N_3$_TS-3-F**
Charge: 1
Multiplicity: 5
B3LYP-D3(BJ)/6-31G(d)-SDD SCF energy (au): -1629.52101210
B3LYP-D3(BJ)/6-31G(d)-SDD enthalpy (au): -1628.98378110
B3LYP-D3(BJ)/6-31G(d)-SDD free energy (au): -1629.09740210
B3LYP-D3(BJ)/def2-TZVP/SMD SCF energy (au): -2770.06182226
B3LYP-D3(BJ)/def2-TZVP/SMD enthalpy (au): -2769.52459126
B3LYP-D3(BJ)/def2-TZVP/SMD free energy (au): -2769.63821226
B3LYP-D3(BJ)/def2-TZVP/SMD free energy (quasi-harmonic) (au): -2769.62710116

Cartesian coordinates

| ATOM | X | Y | Z |
|---|---|---|---|
| H | 32.9101970000 | 26.2664420000 | 37.5700320000 |
| C | 33.2027260000 | 27.2466180000 | 37.2285820000 |
| C | 34.3730980000 | 27.7272590000 | 36.7093170000 |
| N | 32.3405470000 | 28.3250610000 | 37.2568730000 |
| C | 32.9872390000 | 29.4026490000 | 36.7583980000 |
| N | 34.2300160000 | 29.0753270000 | 36.4233400000 |

| | | | |
|---|---|---|---|
| H | 31.3739300000 | 28.3051220000 | 37.5508500000 |
| H | 35.2972960000 | 27.2079060000 | 36.5143630000 |
| H | 32.5536080000 | 30.3830880000 | 36.6445490000 |
| H | 39.4991130000 | 26.6891040000 | 35.8183770000 |
| C | 38.6655590000 | 27.2580570000 | 35.4378150000 |
| C | 37.9301690000 | 28.2826850000 | 35.9706120000 |
| N | 38.1214410000 | 27.0281990000 | 34.1889230000 |
| C | 37.0970060000 | 27.8946810000 | 34.0020540000 |
| N | 36.9561940000 | 28.6679780000 | 35.0690250000 |
| H | 38.4305980000 | 26.3303160000 | 33.5263080000 |
| H | 38.0343380000 | 28.7588580000 | 36.9332690000 |
| H | 36.4785870000 | 27.9493200000 | 33.1188300000 |
| Fe | 35.4641220000 | 30.2746120000 | 35.0709410000 |
| O | 34.0433790000 | 31.8422560000 | 35.4936090000 |
| H | 34.5545320000 | 32.6351420000 | 35.7233450000 |
| H | 33.4940540000 | 32.0800700000 | 34.6806880000 |
| F | 36.6856740000 | 31.4335340000 | 34.2334240000 |
| N | 33.3489030000 | 32.6558230000 | 31.1207820000 |
| O | 32.7082690000 | 32.3527470000 | 33.2932210000 |
| C | 34.5735760000 | 33.7884240000 | 32.8542340000 |
| C | 34.1827320000 | 34.9647410000 | 33.4901510000 |
| C | 35.1272860000 | 35.9361550000 | 33.8318610000 |
| C | 36.4806860000 | 35.7230210000 | 33.5383060000 |
| C | 36.8880790000 | 34.5396500000 | 32.9432260000 |
| C | 35.9548850000 | 33.5252010000 | 32.6030950000 |
| C | 33.4734080000 | 32.8491490000 | 32.4439370000 |
| C | 32.2632770000 | 31.9134910000 | 30.4207960000 |
| C | 32.2175190000 | 30.4609810000 | 30.9098800000 |
| C | 30.9228260000 | 32.6202060000 | 30.6751140000 |
| C | 32.6224470000 | 31.9606140000 | 28.9305820000 |
| C | 36.3706690000 | 32.2399760000 | 32.1627460000 |
| C | 37.7408400000 | 31.8705890000 | 31.7238510000 |
| H | 33.1286950000 | 35.1303060000 | 33.6918540000 |
| H | 34.8088790000 | 36.8572860000 | 34.3093980000 |
| H | 37.2159150000 | 36.4807780000 | 33.7907030000 |
| H | 37.9431690000 | 34.3716620000 | 32.7613400000 |
| H | 33.9676100000 | 33.2019450000 | 30.5354540000 |
| H | 35.5983260000 | 31.4956860000 | 31.9992610000 |
| H | 38.5234570000 | 32.4169050000 | 32.2547980000 |
| H | 37.9030280000 | 30.8004730000 | 31.8746500000 |
| H | 37.8608340000 | 32.0753760000 | 30.6481910000 |
| H | 31.4969670000 | 29.8982320000 | 30.3078330000 |
| H | 33.1990240000 | 29.9871500000 | 30.8091310000 |
| H | 31.9103260000 | 30.4055110000 | 31.9537280000 |
| H | 30.6907260000 | 32.6300720000 | 31.7427100000 |
| H | 30.9499130000 | 33.6520930000 | 30.3084640000 |
| H | 30.1196130000 | 32.0917810000 | 30.1515630000 |
| H | 33.5808340000 | 31.4639320000 | 28.7396660000 |
| H | 31.8540250000 | 31.4460510000 | 28.3475610000 |
| H | 32.6809890000 | 32.9935940000 | 28.5657560000 |
| N | 34.5727240000 | 29.5656480000 | 33.4279350000 |
| N | 33.5346450000 | 28.9383910000 | 33.5169780000 |
| N | 32.5451640000 | 28.3496280000 | 33.5774740000 |

O 36.3829260000 31.1829720000 36.9174660000
H 36.3843330000 30.6497800000 37.7279090000
H 37.3026280000 31.4119240000 36.7040980000

**$^5Im_2Wat_2N_3$_TS-3-$N_3$**

Charge: 1
Multiplicity: 5
B3LYP-D3(BJ)/6-31G(d)-SDD SCF energy (au): -1629.53760394
B3LYP-D3(BJ)/6-31G(d)-SDD enthalpy (au): -1628.99956994
B3LYP-D3(BJ)/6-31G(d)-SDD free energy (au): -1629.11023894
B3LYP-D3(BJ)/def2-TZVP/SMD SCF energy (au): -2770.07238075
B3LYP-D3(BJ)/def2-TZVP/SMD enthalpy (au): -2769.53434675
B3LYP-D3(BJ)/def2-TZVP/SMD free energy (au): -2769.64501575
B3LYP-D3(BJ)/def2-TZVP/SMD free energy (quasi-harmonic) (au): -2769.63561081

Cartesian coordinates

| ATOM | X | Y | Z |
|---|---|---|---|
| H | 33.0382110000 | 27.2832910000 | 38.5628570000 |
| C | 33.2678790000 | 27.9747910000 | 37.7679290000 |
| C | 34.4516150000 | 28.3717490000 | 37.2104670000 |
| N | 32.2866150000 | 28.6737400000 | 37.0904630000 |
| C | 32.8789960000 | 29.4556000000 | 36.1603410000 |
| N | 34.1972910000 | 29.2962420000 | 36.2115440000 |
| H | 31.2909150000 | 28.6032050000 | 37.2477630000 |
| H | 35.4572100000 | 28.0647860000 | 37.4474780000 |
| H | 32.3450750000 | 30.1036350000 | 35.4800590000 |
| H | 39.4683580000 | 26.7287410000 | 36.5189160000 |
| C | 38.6727130000 | 27.2039300000 | 35.9673070000 |
| C | 37.9091000000 | 28.3092890000 | 36.2261400000 |
| N | 38.2325550000 | 26.7248460000 | 34.7474730000 |
| C | 37.2374140000 | 27.5285070000 | 34.3074000000 |
| N | 37.0157050000 | 28.4963410000 | 35.1852560000 |
| H | 38.5957900000 | 25.9168180000 | 34.2609710000 |
| H | 37.9334700000 | 28.9847360000 | 37.0665580000 |
| H | 36.7172460000 | 27.4042010000 | 33.3705570000 |
| Fe | 35.6657880000 | 30.1354670000 | 34.8491350000 |
| O | 34.5290380000 | 31.9900700000 | 35.1023300000 |
| H | 35.0043470000 | 32.5524550000 | 34.4637040000 |
| H | 33.5944110000 | 31.9323740000 | 34.7479040000 |
| F | 36.9261390000 | 31.2453560000 | 34.0841530000 |
| N | 31.2539280000 | 31.5547360000 | 31.8700270000 |
| O | 32.1332800000 | 31.6378320000 | 33.9758550000 |
| C | 33.4511360000 | 32.5164460000 | 32.1956710000 |
| C | 33.7097690000 | 33.8257260000 | 32.5959930000 |
| C | 34.8997900000 | 34.4621650000 | 32.2293010000 |
| C | 35.8512920000 | 33.7650280000 | 31.4732650000 |
| C | 35.6121460000 | 32.4580940000 | 31.0803670000 |
| C | 34.4048760000 | 31.7884610000 | 31.4177890000 |
| C | 32.2193810000 | 31.8583560000 | 32.7467770000 |
| C | 29.9320740000 | 30.9310250000 | 32.1624410000 |
| C | 30.1411720000 | 29.5331680000 | 32.7611220000 |
| C | 29.1400920000 | 31.8340250000 | 33.1199430000 |
| C | 29.2119210000 | 30.8228120000 | 30.8128480000 |

| | | | |
|---|---|---|---|
| C | 34.1868940000 | 30.4356060000 | 31.0525600000 |
| C | 35.1539530000 | 29.6105900000 | 30.2735900000 |
| H | 32.9771740000 | 34.3498290000 | 33.2033060000 |
| H | 35.0828290000 | 35.4875220000 | 32.5337640000 |
| H | 36.7841830000 | 34.2468130000 | 31.1986480000 |
| H | 36.3663390000 | 31.9270950000 | 30.5116800000 |
| H | 31.4206170000 | 31.8414980000 | 30.9137890000 |
| H | 33.2110500000 | 30.0070590000 | 31.2502280000 |
| H | 36.1607140000 | 29.6625570000 | 30.7028010000 |
| H | 34.8423300000 | 28.5634360000 | 30.2558780000 |
| H | 35.2244840000 | 29.9531450000 | 29.2299250000 |
| H | 29.1694560000 | 29.0571680000 | 32.9273090000 |
| H | 30.7221780000 | 28.8968070000 | 32.0867880000 |
| H | 30.6675570000 | 29.5859190000 | 33.7155330000 |
| H | 29.6650130000 | 31.9487310000 | 34.0706840000 |
| H | 28.9890720000 | 32.8255420000 | 32.6806470000 |
| H | 28.1570310000 | 31.3931230000 | 33.3137240000 |
| H | 29.7721980000 | 30.1907710000 | 30.1139730000 |
| H | 28.2252800000 | 30.3735370000 | 30.9535690000 |
| H | 29.0680000000 | 31.8104850000 | 30.3593200000 |
| N | 34.8238530000 | 29.2685720000 | 33.2792450000 |
| N | 33.9088230000 | 28.4690200000 | 33.3149330000 |
| N | 33.0385650000 | 27.7134280000 | 33.3023810000 |
| O | 36.5948390000 | 31.0205610000 | 36.7327360000 |
| H | 35.8893250000 | 31.6535420000 | 36.9572970000 |
| H | 37.2214110000 | 31.5293510000 | 36.1813550000 |

**$^{5}$Im$_2$Wat$_2$OH_11**
Charge: 1
Multiplicity: 5
B3LYP-D3(BJ)/6-31G(d)-SDD SCF energy (au): -1541.09343838
B3LYP-D3(BJ)/6-31G(d)-SDD enthalpy (au): -1540.55497738
B3LYP-D3(BJ)/6-31G(d)-SDD free energy (au): -1540.66585538
B3LYP-D3(BJ)/def2-TZVP/SMD SCF energy (au): -2681.59541738
B3LYP-D3(BJ)/def2-TZVP/SMD enthalpy (au): -2681.05695638
B3LYP-D3(BJ)/def2-TZVP/SMD free energy (au): -2681.16783438
B3LYP-D3(BJ)/def2-TZVP/SMD free energy (quasi-harmonic) (au): -2681.15607500

Cartesian coordinates

| ATOM | X | Y | Z |
|---|---|---|---|
| Fe | -0.0276010000 | -0.7357270000 | 1.5877250000 |
| C | 1.0020500000 | 1.0737990000 | -2.1606470000 |
| N | 2.1666950000 | 1.2810940000 | -1.4404230000 |
| H | 2.9937290000 | 1.7585570000 | -1.7707840000 |
| C | 1.9919480000 | 0.7872680000 | -0.1940480000 |
| H | 2.7273910000 | 0.8276050000 | 0.5945770000 |
| N | 0.7771480000 | 0.2623970000 | -0.0833250000 |
| C | 0.1459640000 | 0.4367020000 | -1.3031450000 |
| H | -0.8637840000 | 0.0876210000 | -1.4519390000 |
| H | 0.9000660000 | 1.3834300000 | -3.1887440000 |
| C | 1.1201100000 | -4.7745200000 | 2.3808580000 |
| N | 2.0079680000 | -4.5369200000 | 1.3490740000 |
| H | 2.6763570000 | -5.1974760000 | 0.9775120000 |

| | | | |
|---|---|---|---|
| C | 1.8277040000 | -3.2629670000 | 0.9198320000 |
| H | 2.3918190000 | -2.8118140000 | 0.1174400000 |
| N | 0.8721200000 | -2.6759010000 | 1.6232780000 |
| C | 0.4191600000 | -3.6098700000 | 2.5421910000 |
| H | -0.3684630000 | -3.3543670000 | 3.2356360000 |
| H | 1.0725810000 | -5.7240920000 | 2.8900580000 |
| O | -1.8610820000 | -1.4907700000 | 0.3306320000 |
| H | -2.1876610000 | -1.3748580000 | 1.2617800000 |
| H | -1.8102660000 | -2.4522470000 | 0.2062250000 |
| C | 1.8990850000 | 4.9478810000 | -0.4302250000 |
| C | 0.9327690000 | 4.0418390000 | -0.0042340000 |
| C | 0.5998090000 | 3.9335740000 | 1.3560610000 |
| C | 1.2533570000 | 4.7336940000 | 2.3219930000 |
| C | 2.2079090000 | 5.6473700000 | 1.8625650000 |
| C | 2.5331530000 | 5.7598530000 | 0.5099280000 |
| H | 2.1475510000 | 5.0267850000 | -1.4845200000 |
| H | 0.4304950000 | 3.3968460000 | -0.7162580000 |
| H | 2.7083270000 | 6.2845820000 | 2.5860930000 |
| H | 3.2831330000 | 6.4791930000 | 0.1948230000 |
| C | 0.9241910000 | 4.6861090000 | 3.7991740000 |
| C | -0.4560990000 | 5.2777270000 | 4.1279470000 |
| H | 0.9684700000 | 3.6560410000 | 4.1610320000 |
| H | 1.6923720000 | 5.2493750000 | 4.3391870000 |
| H | -0.6222700000 | 5.2846490000 | 5.2096390000 |
| H | -1.2592930000 | 4.6870270000 | 3.6758750000 |
| H | -0.5419830000 | 6.3066260000 | 3.7625830000 |
| C | -0.3740290000 | 2.8816560000 | 1.7438140000 |
| O | -0.1902030000 | 2.1007870000 | 2.6829570000 |
| N | -1.4273510000 | 2.7010620000 | 0.8532340000 |
| C | -2.5714200000 | 3.6384130000 | 0.5635080000 |
| C | -2.0647600000 | 5.0734150000 | 0.3794540000 |
| C | -3.2170360000 | 3.1416600000 | -0.7363700000 |
| H | -2.9373410000 | 5.6951790000 | 0.1585390000 |
| H | -1.3646370000 | 5.1636160000 | -0.4527340000 |
| H | -1.5903130000 | 5.4693440000 | 1.2797120000 |
| H | -3.5868390000 | 2.1196690000 | -0.6286320000 |
| H | -2.4933260000 | 3.1684920000 | -1.5573420000 |
| H | -4.0602720000 | 3.7880410000 | -0.9966180000 |
| F | -2.0058840000 | 1.4333990000 | 1.1091180000 |
| C | -3.5633770000 | 3.5842750000 | 1.7376680000 |
| H | -3.0939040000 | 3.9282540000 | 2.6646810000 |
| H | -3.9433610000 | 2.5731450000 | 1.8922920000 |
| H | -4.4109800000 | 4.2417360000 | 1.5228710000 |
| O | -1.3834510000 | -0.9707970000 | 2.8751450000 |
| H | -1.6813300000 | -0.1603030000 | 3.3143060000 |
| O | 1.6359650000 | 0.1795900000 | 2.7738800000 |
| H | 1.6718350000 | -0.3131970000 | 3.6097490000 |
| H | 1.1688090000 | 1.0270740000 | 2.9697790000 |

**$^{5}$Im$_2$Wat$_2$OH_13**
Charge: 1
Multiplicity: 5
B3LYP-D3(BJ)/6-31G(d)-SDD SCF energy (au): -1541.15440206

B3LYP-D3(BJ)/6-31G(d)-SDD enthalpy (au): -1540.61714106
B3LYP-D3(BJ)/6-31G(d)-SDD free energy (au): -1540.72822706
B3LYP-D3(BJ)/def2-TZVP/SMD SCF energy (au): -2681.66201061
B3LYP-D3(BJ)/def2-TZVP/SMD enthalpy (au): -2681.12474961
B3LYP-D3(BJ)/def2-TZVP/SMD free energy (au): -2681.23583561
B3LYP-D3(BJ)/def2-TZVP/SMD free energy (quasi-harmonic) (au): -2681.22380355

Cartesian coordinates

| ATOM | X | Y | Z |
|---|---|---|---|
| Fe | 0.1182290000 | 0.1861450000 | -0.5069700000 |
| C | 1.7884360000 | -0.2564820000 | 3.4773530000 |
| N | 2.8209040000 | 0.2866870000 | 2.7331900000 |
| H | 3.7357340000 | 0.5378870000 | 3.0820330000 |
| C | 2.3974190000 | 0.4417550000 | 1.4589430000 |
| H | 2.9778090000 | 0.8506800000 | 0.6457750000 |
| N | 1.1439160000 | 0.0203380000 | 1.3518340000 |
| C | 0.7478860000 | -0.4165640000 | 2.6034770000 |
| H | -0.2474190000 | -0.7969680000 | 2.7707430000 |
| O | 1.5472480000 | 0.9700790000 | -1.3352430000 |
| H | 1.2814850000 | 1.7079730000 | -1.9206050000 |
| H | 1.8865710000 | -0.4649340000 | 4.5309670000 |
| C | 1.2641540000 | -3.6393790000 | -2.1180680000 |
| N | 1.1892240000 | -3.9542440000 | -0.7725900000 |
| H | 1.4034810000 | -4.8517890000 | -0.3604950000 |
| C | 0.7825080000 | -2.8541510000 | -0.0993890000 |
| H | 0.6472150000 | -2.8075700000 | 0.9697840000 |
| N | 0.5906090000 | -1.8536290000 | -0.9475220000 |
| C | 0.8898340000 | -2.3275520000 | -2.2116850000 |
| H | 0.8069980000 | -1.6906410000 | -3.0768520000 |
| H | 1.5698080000 | -4.3590510000 | -2.8606900000 |
| O | -0.9908150000 | 0.1190630000 | -2.3846290000 |
| H | -0.6695800000 | 0.9975120000 | -2.7625950000 |
| H | -1.9435010000 | 0.2735820000 | -2.2681450000 |
| F | -1.0665860000 | 1.5236600000 | 0.0355630000 |
| C | -1.0378680000 | 3.3634790000 | -2.5483260000 |
| O | -0.1403850000 | 2.5587900000 | -2.9059150000 |
| N | -0.7772240000 | 4.5997120000 | -2.1037800000 |
| H | -2.1202770000 | 3.8210740000 | -0.1181250000 |
| C | 0.5458910000 | 5.2166870000 | -1.8229640000 |
| C | 0.2499970000 | 6.6780350000 | -1.4583820000 |
| C | 1.2033030000 | 4.5050580000 | -0.6291000000 |
| C | 1.4378600000 | 5.1610190000 | -3.0718430000 |
| H | -0.2344280000 | 7.2020700000 | -2.2898820000 |
| H | -0.3975090000 | 6.7434310000 | -0.5754530000 |
| H | 1.1823340000 | 7.2005510000 | -1.2281300000 |
| H | 1.4052870000 | 3.4543890000 | -0.8426330000 |
| H | 2.1528310000 | 4.9949590000 | -0.3881610000 |
| H | 0.5562760000 | 4.5534240000 | 0.2531100000 |
| H | 2.3839060000 | 5.6721740000 | -2.8661830000 |
| H | 1.6510520000 | 4.1315640000 | -3.3626770000 |
| H | 0.9516250000 | 5.6656840000 | -3.9130300000 |
| C | -2.4794710000 | 2.9899110000 | -2.6891920000 |
| C | -2.8311130000 | 2.3646910000 | -3.8878010000 |

| | | | |
|---|---|---|---|
| C | -3.4781110000 | 3.2629640000 | -1.6899860000 |
| C | -4.1563110000 | 2.0276370000 | -4.1694650000 |
| H | -2.0527000000 | 2.1699030000 | -4.6192840000 |
| C | -4.8233230000 | 2.9254780000 | -2.0248210000 |
| C | -5.1521450000 | 2.3248360000 | -3.2286150000 |
| H | -4.4105880000 | 1.5597910000 | -5.1148210000 |
| H | -5.6051480000 | 3.1214350000 | -1.2995390000 |
| H | -6.1886140000 | 2.0790560000 | -3.4401320000 |
| C | -3.1623850000 | 3.7735660000 | -0.4070130000 |
| C | -4.1806450000 | 4.1098620000 | 0.6352630000 |
| H | -4.6242890000 | 3.2050080000 | 1.0805110000 |
| H | -3.7266650000 | 4.6778890000 | 1.4517780000 |
| H | -5.0126300000 | 4.7014980000 | 0.2337400000 |
| H | -1.5879770000 | 5.1654280000 | -1.8884840000 |
| O | -1.6745080000 | -0.8360300000 | 0.6974670000 |
| H | -2.3107310000 | -1.3295560000 | 0.1562810000 |
| H | -1.9799980000 | 0.0978290000 | 0.7173460000 |

**$^5Im_2Wat_2OH$_FeF_remove-F**
Charge: 1
Multiplicity: 5
B3LYP-D3(BJ)/6-31G(d)-SDD SCF energy (au): -805.005489630
B3LYP-D3(BJ)/6-31G(d)-SDD enthalpy (au): -804.776436630
B3LYP-D3(BJ)/6-31G(d)-SDD free energy (au): -804.846817630
B3LYP-D3(BJ)/def2-TZVP/SMD SCF energy (au): -1945.22035199
B3LYP-D3(BJ)/def2-TZVP/SMD enthalpy (au): -1944.99129899
B3LYP-D3(BJ)/def2-TZVP/SMD free energy (au): -1945.06167999
B3LYP-D3(BJ)/def2-TZVP/SMD free energy (quasi-harmonic) (au): -1945.057093360

Cartesian coordinates

| ATOM | X | Y | Z |
|---|---|---|---|
| Fe | 0.4671800000 | -0.7918340000 | 0.8674320000 |
| C | -1.7316720000 | -0.4701070000 | -2.7944250000 |
| N | -2.4699210000 | -1.4030480000 | -2.0897160000 |
| H | -3.3011800000 | -1.8742700000 | -2.4210650000 |
| C | -1.8970580000 | -1.5792020000 | -0.8798070000 |
| H | -2.2540620000 | -2.2632690000 | -0.1251680000 |
| N | -0.8237360000 | -0.8014350000 | -0.7705960000 |
| C | -0.7084230000 | -0.1056610000 | -1.9640230000 |
| H | 0.1002740000 | 0.5883270000 | -2.1286490000 |
| H | -1.9985880000 | -0.1598430000 | -3.7922020000 |
| C | 2.5698830000 | -4.4744370000 | -0.0781160000 |
| N | 2.8841840000 | -4.2418630000 | 1.2466490000 |
| H | 3.4438820000 | -4.8417230000 | 1.8375950000 |
| C | 2.3086500000 | -3.0777250000 | 1.6267230000 |
| H | 2.3812660000 | -2.6418690000 | 2.6123080000 |
| N | 1.6411970000 | -2.5494100000 | 0.6075550000 |
| C | 1.7957600000 | -3.4129640000 | -0.4629540000 |
| H | 1.3405580000 | -3.2104220000 | -1.4206210000 |
| H | 2.9168980000 | -5.3473970000 | -0.6078920000 |
| O | -1.1551080000 | -1.8106760000 | 2.2266160000 |
| H | -0.5301980000 | -1.3828450000 | 2.8650180000 |
| H | -1.9910510000 | -1.3328750000 | 2.3552580000 |

| | | | |
|---|---|---|---|
| O | 0.9654550000 | -0.2901780000 | 2.6036920000 |
| H | 1.0060540000 | 0.6083850000 | 2.9575210000 |
| O | 2.0252390000 | 0.3235330000 | -0.3227430000 |
| H | 2.2849100000 | 1.1572400000 | 0.1041030000 |
| H | 2.8070220000 | -0.2515760000 | -0.2598520000 |

**$^{5}$Im$_2$Wat$_2$OH_FeF_remove-OH**

Charge: 1
Multiplicity: 5
B3LYP-D3(BJ)/6-31G(d)-SDD SCF energy (au): -829.053220256
B3LYP-D3(BJ)/6-31G(d)-SDD enthalpy (au): -828.835091256
B3LYP-D3(BJ)/6-31G(d)-SDD free energy (au): -828.903144256
B3LYP-D3(BJ)/def2-TZVP/SMD SCF energy (au): -1969.27415149
B3LYP-D3(BJ)/def2-TZVP/SMD enthalpy (au): -1969.05602249
B3LYP-D3(BJ)/def2-TZVP/SMD free energy (au): -1969.12407549
B3LYP-D3(BJ)/def2-TZVP/SMD free energy (quasi-harmonic) (au): -1969.119686234

Cartesian coordinates

| ATOM | X | Y | Z |
|---|---|---|---|
| Fe | 0.2660210000 | -0.9982670000 | 0.7269780000 |
| C | -1.3815070000 | -0.6476530000 | -3.2118810000 |
| N | -2.5041050000 | -0.8048300000 | -2.4209680000 |
| H | -3.4622560000 | -0.8028860000 | -2.7448810000 |
| C | -2.1106720000 | -0.9569030000 | -1.1385190000 |
| H | -2.7755780000 | -1.1058050000 | -0.3013020000 |
| N | -0.7836520000 | -0.9027570000 | -1.0644460000 |
| C | -0.3177600000 | -0.7104910000 | -2.3555090000 |
| H | 0.7373410000 | -0.6415750000 | -2.5721760000 |
| H | -1.4434620000 | -0.5112580000 | -4.2798860000 |
| C | 2.2890340000 | -4.7739340000 | 0.6988370000 |
| N | 3.3299130000 | -3.8676900000 | 0.7748350000 |
| H | 4.3129870000 | -4.0979410000 | 0.8342630000 |
| C | 2.8167910000 | -2.6190410000 | 0.7625420000 |
| H | 3.3988140000 | -1.7109140000 | 0.8043240000 |
| N | 1.4901880000 | -2.6773740000 | 0.6842180000 |
| C | 1.1498620000 | -4.0202620000 | 0.6429350000 |
| H | 0.1222880000 | -4.3417930000 | 0.5677390000 |
| H | 2.4524460000 | -5.8399080000 | 0.6903260000 |
| O | -1.5865710000 | -1.6905140000 | 1.8654270000 |
| H | -1.5087690000 | -2.4604670000 | 2.4523640000 |
| H | -1.3832410000 | -0.9112990000 | 2.4294330000 |
| F | 0.1281490000 | 0.2204940000 | 2.1368880000 |
| O | 2.0401090000 | 0.4009220000 | 0.4093930000 |
| H | 1.9727570000 | 1.0857050000 | -0.2761110000 |
| H | 1.7038300000 | 0.8216780000 | 1.2318280000 |

**$^{6}$Im$_2$Wat$_2$OH_FeF**

Charge: 1
Multiplicity: 6
B3LYP-D3(BJ)/6-31G(d)-SDD SCF energy (au): -904.865739905
B3LYP-D3(BJ)/6-31G(d)-SDD enthalpy (au): -904.632340905
B3LYP-D3(BJ)/6-31G(d)-SDD free energy (au): -904.702620905
B3LYP-D3(BJ)/def2-TZVP/SMD SCF energy (au): -2045.12474063

B3LYP-D3(BJ)/def2-TZVP/SMD enthalpy (au): -2044.89134163
B3LYP-D3(BJ)/def2-TZVP/SMD free energy (au): -2044.96162163
B3LYP-D3(BJ)/def2-TZVP/SMD free energy (quasi-harmonic) (au): -2044.958718725

Cartesian coordinates

| ATOM | X | Y | Z |
|---|---|---|---|
| Fe | 0.3081710000 | -0.7274540000 | 1.1870350000 |
| C | -1.1073190000 | -1.1579100000 | -2.9452490000 |
| N | -2.2190910000 | -0.6842230000 | -2.2726260000 |
| H | -3.1193570000 | -0.4782630000 | -2.6847390000 |
| C | -1.8986700000 | -0.5309040000 | -0.9691410000 |
| H | -2.5639550000 | -0.1501360000 | -0.2101020000 |
| N | -0.6359240000 | -0.8907490000 | -0.7689870000 |
| C | -0.1288350000 | -1.2801890000 | -1.9973180000 |
| H | 0.8921460000 | -1.6107470000 | -2.1040270000 |
| H | -1.1161720000 | -1.3542180000 | -4.0057400000 |
| C | 2.1299840000 | -4.5070430000 | -0.0193260000 |
| N | 3.1107850000 | -3.9016680000 | 0.7459550000 |
| H | 4.0189320000 | -4.2904120000 | 0.9625810000 |
| C | 2.6514560000 | -2.7031200000 | 1.1648190000 |
| H | 3.1864150000 | -2.0177640000 | 1.8039740000 |
| N | 1.4225410000 | -2.5052060000 | 0.6988320000 |
| C | 1.0835860000 | -3.6270400000 | -0.0393740000 |
| H | 0.1220400000 | -3.7148220000 | -0.5193840000 |
| H | 2.2642240000 | -5.4813030000 | -0.4620000000 |
| O | -1.0280210000 | -2.1700660000 | 2.1681130000 |
| H | -0.5344880000 | -2.6997310000 | 2.8178530000 |
| H | -1.6167720000 | -1.5807040000 | 2.6746160000 |
| F | -1.0301030000 | 0.4309660000 | 1.6497040000 |
| O | 1.5256550000 | -0.4188460000 | 2.5606370000 |
| H | 1.1994660000 | 0.1959170000 | 3.2387460000 |
| O | 1.5834960000 | 0.6521930000 | 0.0504890000 |
| H | 1.0680520000 | 1.3727790000 | -0.3514260000 |
| H | 2.1098340000 | 1.0429740000 | 0.7721180000 |

**$^{5}$Im$_2$Wat$_2$OH_TS-1**
Charge: 1
Multiplicity: 5
B3LYP-D3(BJ)/6-31G(d)-SDD SCF energy (au): -1541.07398927
B3LYP-D3(BJ)/6-31G(d)-SDD enthalpy (au): -1540.53838627
B3LYP-D3(BJ)/6-31G(d)-SDD free energy (au): -1540.64632327
B3LYP-D3(BJ)/def2-TZVP/SMD SCF energy (au): -2681.57376980
B3LYP-D3(BJ)/def2-TZVP/SMD enthalpy (au): -2681.03816680
B3LYP-D3(BJ)/def2-TZVP/SMD free energy (au): -2681.14610380
B3LYP-D3(BJ)/def2-TZVP/SMD free energy (quasi-harmonic) (au): -2681.13651553

Cartesian coordinates

| ATOM | X | Y | Z |
|---|---|---|---|
| Fe | 0.4933960000 | -0.5962390000 | 1.0027650000 |
| C | -1.3928300000 | -0.9556550000 | -2.8430270000 |
| N | -2.2707080000 | -1.6431120000 | -2.0247920000 |
| H | -3.1084350000 | -2.1234870000 | -2.3237110000 |
| C | -1.8178380000 | -1.5707930000 | -0.7514840000 |

| | | | |
|---|---|---|---|
| H | -2.2862050000 | -2.0335940000 | 0.1036110000 |
| N | -0.6945840000 | -0.8667450000 | -0.7161660000 |
| C | -0.4136560000 | -0.4778460000 | -2.0141230000 |
| H | 0.4654720000 | 0.1046700000 | -2.2400870000 |
| H | -1.5443380000 | -0.8718620000 | -3.9075520000 |
| C | 2.2808400000 | -4.3246870000 | -0.3225990000 |
| N | 2.7228130000 | -4.1973080000 | 0.9806620000 |
| H | 3.3152700000 | -4.8507830000 | 1.4737580000 |
| C | 2.2224790000 | -3.0479910000 | 1.4928130000 |
| H | 2.3917290000 | -2.6743340000 | 2.4912650000 |
| N | 1.4801400000 | -2.4346330000 | 0.5811170000 |
| C | 1.5077900000 | -3.2196720000 | -0.5569040000 |
| H | 0.9799580000 | -2.9322740000 | -1.4525970000 |
| H | 2.5514830000 | -5.1639390000 | -0.9435720000 |
| O | -1.3463940000 | -1.5073070000 | 2.2501760000 |
| H | -0.5889520000 | -1.4710890000 | 2.8812180000 |
| H | -1.8075940000 | -0.6670150000 | 2.4134560000 |
| C | 1.3917770000 | 6.8812220000 | 2.7649300000 |
| C | 1.3550160000 | 5.6573290000 | 2.1136660000 |
| C | 0.5808890000 | 4.5993470000 | 2.6288180000 |
| C | -0.1280130000 | 4.7400090000 | 3.8430860000 |
| C | -0.0571030000 | 5.9835280000 | 4.4850820000 |
| C | 0.6717310000 | 7.0444980000 | 3.9537760000 |
| H | 1.9791060000 | 7.6988500000 | 2.3599050000 |
| H | 1.9120170000 | 5.5000230000 | 1.1956220000 |
| H | -0.5740430000 | 6.1113430000 | 5.4323330000 |
| H | 0.6959510000 | 7.9959340000 | 4.4764510000 |
| C | -0.8769020000 | 3.6145580000 | 4.5290950000 |
| C | -2.4041200000 | 3.7516750000 | 4.4463380000 |
| H | -0.5789450000 | 2.6463170000 | 4.1195060000 |
| H | -0.5782290000 | 3.6151980000 | 5.5846490000 |
| H | -2.8916880000 | 2.9731220000 | 5.0422200000 |
| H | -2.7446300000 | 3.6526110000 | 3.4117990000 |
| H | -2.7383580000 | 4.7238930000 | 4.8238120000 |
| C | 0.5693070000 | 3.3575480000 | 1.8159530000 |
| O | 1.5952390000 | 2.8445720000 | 1.3747930000 |
| N | -0.7349360000 | 2.8855950000 | 1.6244270000 |
| C | -1.4892680000 | 3.2669120000 | 0.4125530000 |
| C | -1.9002220000 | 4.7509680000 | 0.6124250000 |
| C | -0.6616420000 | 3.1244580000 | -0.8723900000 |
| H | -2.5604460000 | 5.0229730000 | -0.2167560000 |
| H | -1.0355130000 | 5.4168310000 | 0.6015240000 |
| H | -2.4399980000 | 4.8936690000 | 1.5507320000 |
| H | -0.3491410000 | 2.0893980000 | -1.0155440000 |
| H | 0.2272570000 | 3.7619870000 | -0.8559400000 |
| H | -1.2747270000 | 3.4215950000 | -1.7289440000 |
| F | -0.4522790000 | 1.1132090000 | 1.3985180000 |
| C | -2.7541660000 | 2.3994850000 | 0.3578250000 |
| H | -3.2972990000 | 2.4498320000 | 1.3060930000 |
| H | -2.4973050000 | 1.3588940000 | 0.1519840000 |
| H | -3.4104350000 | 2.7642420000 | -0.4381340000 |
| O | 1.1802440000 | -0.6934890000 | 2.7523150000 |
| H | 1.2301700000 | 0.1782260000 | 3.1743420000 |

| | | | |
|---|---|---|---|
| O | 1.9045890000 | 0.6175650000 | -0.1183770000 |
| H | 1.8157390000 | 1.4848610000 | 0.3601560000 |
| H | 2.8068370000 | 0.3013900000 | 0.0499800000 |

**$^{5}$Im$_2$Wat$_2$OH_TS-3-F**
Charge: 1
Multiplicity: 5
B3LYP-D3(BJ)/6-31G(d)-SDD SCF energy (au): -1541.14373953
B3LYP-D3(BJ)/6-31G(d)-SDD enthalpy (au): -1540.60683953
B3LYP-D3(BJ)/6-31G(d)-SDD free energy (au): -1540.71279553
B3LYP-D3(BJ)/def2-TZVP/SMD SCF energy (au): -2681.64814221
B3LYP-D3(BJ)/def2-TZVP/SMD enthalpy (au): -2681.11124221
B3LYP-D3(BJ)/def2-TZVP/SMD free energy (au): -2681.21719821
B3LYP-D3(BJ)/def2-TZVP/SMD free energy (quasi-harmonic) (au): -2681.20954768

Cartesian coordinates

| ATOM | X | Y | Z |
|---|---|---|---|
| Fe | -0.6000410000 | 0.9701060000 | -0.9216350000 |
| C | 1.0928410000 | 1.1939180000 | 3.1117970000 |
| N | 1.4709980000 | 2.3828250000 | 2.5156760000 |
| H | 1.9990280000 | 3.1244660000 | 2.9538830000 |
| C | 1.0046210000 | 2.3893830000 | 1.2432180000 |
| H | 1.1387430000 | 3.1850190000 | 0.5260020000 |
| N | 0.3483340000 | 1.2651930000 | 0.9966100000 |
| C | 0.3934280000 | 0.5094280000 | 2.1540060000 |
| H | -0.0824110000 | -0.4573140000 | 2.2070120000 |
| O | 0.1714870000 | 2.5291320000 | -1.6839430000 |
| H | 0.9668490000 | 2.3260530000 | -2.1971290000 |
| H | 1.3461240000 | 0.9562050000 | 4.1329690000 |
| C | 2.3341240000 | -1.4602660000 | -3.0328570000 |
| N | 2.6401360000 | -1.7703530000 | -1.7204460000 |
| H | 3.3839230000 | -2.3813810000 | -1.4129460000 |
| C | 1.7660000000 | -1.1185070000 | -0.9159260000 |
| H | 1.7752420000 | -1.1707830000 | 0.1620390000 |
| N | 0.9176810000 | -0.4103760000 | -1.6444530000 |
| C | 1.2613820000 | -0.6124570000 | -2.9708820000 |
| H | 0.7144880000 | -0.1346310000 | -3.7704340000 |
| H | 2.8925020000 | -1.8629030000 | -3.8631070000 |
| O | -1.9133620000 | -0.0982290000 | -2.4162440000 |
| H | -1.6288030000 | 0.2094670000 | -3.3224770000 |
| H | -2.7072620000 | 0.4328260000 | -2.2263290000 |
| F | -2.3058680000 | 1.8013660000 | -0.5546140000 |
| C | -1.7498940000 | 2.3145990000 | -4.4569530000 |
| O | -1.1069360000 | 1.2520430000 | -4.5679770000 |
| N | -1.2947570000 | 3.5145580000 | -4.8480530000 |
| H | -1.9509380000 | 4.2811400000 | -4.7771100000 |
| C | -0.0197540000 | 3.8009940000 | -5.5612820000 |
| C | -0.0059370000 | 5.3160160000 | -5.7996610000 |
| C | 1.1786340000 | 3.3980020000 | -4.6917660000 |
| C | -0.0069920000 | 3.0473840000 | -6.9003210000 |
| H | -0.8616170000 | 5.6302860000 | -6.4093520000 |
| H | -0.0268720000 | 5.8649640000 | -4.8510890000 |
| H | 0.9045730000 | 5.6003770000 | -6.3338740000 |

| | | | |
|---|---|---|---|
| H | 1.1880480000 | 2.3193600000 | -4.5226510000 |
| H | 2.1060470000 | 3.6780610000 | -5.2018030000 |
| H | 1.1377390000 | 3.9080760000 | -3.7257480000 |
| H | 0.9158010000 | 3.2733570000 | -7.4442770000 |
| H | -0.0590870000 | 1.9685140000 | -6.7374110000 |
| H | -0.8544570000 | 3.3509480000 | -7.5240800000 |
| C | -3.1737940000 | 2.2964570000 | -3.9577560000 |
| C | -4.0838330000 | 1.6054000000 | -4.7568930000 |
| C | -3.6108080000 | 2.9330690000 | -2.7545670000 |
| C | -5.4404130000 | 1.5688050000 | -4.4272780000 |
| H | -3.7262630000 | 1.1065420000 | -5.6523370000 |
| C | -4.9969610000 | 2.8845730000 | -2.4522620000 |
| C | -5.8945860000 | 2.2233440000 | -3.2755040000 |
| H | -6.1382960000 | 1.0387540000 | -5.0674090000 |
| H | -5.3551250000 | 3.3549840000 | -1.5440910000 |
| H | -6.9487810000 | 2.2014680000 | -3.0179590000 |
| C | -2.6691260000 | 3.5014770000 | -1.8449410000 |
| C | -3.0184680000 | 4.3799120000 | -0.6999860000 |
| H | -3.9173100000 | 4.0581320000 | -0.1687550000 |
| H | -2.1875310000 | 4.4136160000 | 0.0068350000 |
| H | -3.1949560000 | 5.4044120000 | -1.0641960000 |
| H | -1.6160950000 | 3.4618640000 | -2.1148400000 |
| O | -1.4873050000 | -0.8957330000 | 0.1541100000 |
| H | -1.7616270000 | -1.3238940000 | -0.6781020000 |
| H | -2.3013120000 | -0.5059140000 | 0.5173060000 |

**$^{5}Im_2Wat_2OH$_TS-3-OH**
Charge: 1
Multiplicity: 5
B3LYP-D3(BJ)/6-31G(d)-SDD SCF energy (au): -1541.14678503
B3LYP-D3(BJ)/6-31G(d)-SDD enthalpy (au): -1540.61114203
B3LYP-D3(BJ)/6-31G(d)-SDD free energy (au): -1540.71889003
B3LYP-D3(BJ)/def2-TZVP/SMD SCF energy (au): -2681.65427957
B3LYP-D3(BJ)/def2-TZVP/SMD enthalpy (au): -2681.11863657
B3LYP-D3(BJ)/def2-TZVP/SMD free energy (au): -2681.22638457
B3LYP-D3(BJ)/def2-TZVP/SMD free energy (quasi-harmonic) (au): -2681.21638854

Cartesian coordinates

| ATOM | X | Y | Z |
|---|---|---|---|
| Fe | 0.2679990000 | 0.3922120000 | -1.4093650000 |
| C | 1.2491170000 | 1.6557280000 | 2.6518910000 |
| N | 0.4751920000 | 2.6847350000 | 2.1448330000 |
| H | 0.2625590000 | 3.5535310000 | 2.6150610000 |
| C | 0.0421180000 | 2.3312940000 | 0.9134970000 |
| H | -0.5713150000 | 2.9362550000 | 0.2642120000 |
| N | 0.5020380000 | 1.1237640000 | 0.6061960000 |
| C | 1.2588980000 | 0.6931720000 | 1.6802060000 |
| H | 1.7445720000 | -0.2695390000 | 1.6761900000 |
| O | 1.8836330000 | 1.0435140000 | -2.0940990000 |
| H | 2.2015700000 | 0.4913740000 | -2.8287570000 |
| H | 1.7023590000 | 1.7037120000 | 3.6293650000 |
| C | 1.5444830000 | -3.6744750000 | -0.4467490000 |
| N | 2.7344250000 | -2.9705780000 | -0.4375810000 |

| | | | |
|---|---|---|---|
| H | 3.6541130000 | -3.3624590000 | -0.2877600000 |
| C | 2.4615230000 | -1.6687250000 | -0.6891710000 |
| H | 3.2002540000 | -0.8843180000 | -0.7523470000 |
| N | 1.1551350000 | -1.5034030000 | -0.8492130000 |
| C | 0.5709280000 | -2.7477130000 | -0.7042490000 |
| H | -0.4938370000 | -2.8803990000 | -0.8113260000 |
| H | 1.5045490000 | -4.7390730000 | -0.2787690000 |
| O | -0.2297600000 | -0.4366630000 | -3.2936230000 |
| H | -0.2914350000 | 0.3409770000 | -3.9339710000 |
| H | -1.1116510000 | -0.8428320000 | -3.2529220000 |
| F | -1.1737990000 | 1.6404140000 | -1.5125720000 |
| C | -0.7380050000 | 2.7084150000 | -4.9260910000 |
| O | -0.3553790000 | 1.5222470000 | -5.0179130000 |
| N | -1.4143940000 | 3.3148090000 | -5.9193210000 |
| H | -1.5881740000 | 4.3042660000 | -5.8047860000 |
| C | -1.8495340000 | 2.7016970000 | -7.2022630000 |
| C | -2.6175330000 | 3.7966770000 | -7.9531060000 |
| C | -0.6188860000 | 2.2642150000 | -8.0119300000 |
| C | -2.7757430000 | 1.5097600000 | -6.9169820000 |
| H | -3.4931010000 | 4.1288760000 | -7.3831210000 |
| H | -1.9772250000 | 4.6641520000 | -8.1531940000 |
| H | -2.9683230000 | 3.4126390000 | -8.9148240000 |
| H | -0.0488090000 | 1.5075880000 | -7.4693650000 |
| H | -0.9370810000 | 1.8424880000 | -8.9709270000 |
| H | 0.0334500000 | 3.1204300000 | -8.2142340000 |
| H | -3.1376390000 | 1.0905980000 | -7.8612670000 |
| H | -2.2450260000 | 0.7276170000 | -6.3710500000 |
| H | -3.6426800000 | 1.8278840000 | -6.3279970000 |
| C | -0.4885700000 | 3.5638790000 | -3.7222540000 |
| C | -1.5667710000 | 4.2595080000 | -3.1786590000 |
| C | 0.8113620000 | 3.6742840000 | -3.1326500000 |
| C | -1.4141540000 | 5.0630530000 | -2.0472010000 |
| H | -2.5500540000 | 4.1447890000 | -3.6251090000 |
| C | 0.9388000000 | 4.5185830000 | -1.9978780000 |
| C | -0.1481590000 | 5.1948430000 | -1.4652240000 |
| H | -2.2700780000 | 5.5865810000 | -1.6332430000 |
| H | 1.9150700000 | 4.6355500000 | -1.5415070000 |
| H | -0.0116520000 | 5.8381360000 | -0.5999740000 |
| C | 1.9423370000 | 2.9992790000 | -3.6610390000 |
| C | 3.3500950000 | 3.2673370000 | -3.2441770000 |
| H | 3.4712340000 | 3.2208070000 | -2.1574570000 |
| H | 4.0322950000 | 2.5376540000 | -3.6879810000 |
| H | 3.6760670000 | 4.2657280000 | -3.5738030000 |
| H | 1.7841360000 | 2.3700430000 | -4.5285490000 |
| O | -1.7748320000 | -0.6953150000 | -0.8744470000 |
| H | -2.0235430000 | 0.2506570000 | -1.0390850000 |
| H | -1.8543900000 | -0.8176600000 | 0.0852610000 |

**$^{5}$Im$_2$Wat$_3$_11**
Charge: 2
Multiplicity: 5
B3LYP-D3(BJ)/6-31G(d)-SDD SCF energy (au): -1541.41077592
B3LYP-D3(BJ)/6-31G(d)-SDD enthalpy (au): -1540.85962092

B3LYP-D3(BJ)/6-31G(d)-SDD free energy (au): -1540.97374692
B3LYP-D3(BJ)/def2-TZVP/SMD SCF energy (au): -2682.02920429
B3LYP-D3(BJ)/def2-TZVP/SMD enthalpy (au): -2681.47804929
B3LYP-D3(BJ)/def2-TZVP/SMD free energy (au): -2681.59217529
B3LYP-D3(BJ)/def2-TZVP/SMD free energy (quasi-harmonic) (au): -2681.58042037

Cartesian coordinates

| ATOM | X | Y | Z |
|---|---|---|---|
| Fe | 0.3443720000 | -1.0430780000 | -0.7976230000 |
| C | 1.6863430000 | 2.0969600000 | 1.8199650000 |
| N | 1.8557060000 | 2.6945560000 | 0.5861100000 |
| H | 2.2354250000 | 3.6190730000 | 0.4252850000 |
| C | 1.4561610000 | 1.8300630000 | -0.3714510000 |
| H | 1.4829170000 | 2.0378410000 | -1.4291740000 |
| N | 1.0340250000 | 0.6991590000 | 0.1877560000 |
| C | 1.1763540000 | 0.8548810000 | 1.5586010000 |
| H | 0.9223980000 | 0.0643400000 | 2.2480830000 |
| O | 1.8277010000 | -1.0023850000 | -2.3845030000 |
| H | 1.9254300000 | -1.8699430000 | -2.8145640000 |
| H | 1.4711340000 | -0.4030230000 | -3.0654890000 |
| H | 1.9449000000 | 2.5939700000 | 2.7419720000 |
| C | 1.8911490000 | -4.0944350000 | 1.8280070000 |
| N | 3.0497900000 | -3.4774340000 | 1.4002480000 |
| H | 3.9912180000 | -3.7382940000 | 1.6673190000 |
| C | 2.7196020000 | -2.4733330000 | 0.5603560000 |
| H | 3.4347950000 | -1.8222400000 | 0.0801170000 |
| N | 1.3972230000 | -2.4103370000 | 0.4230990000 |
| C | 0.8699900000 | -3.4218820000 | 1.2141590000 |
| H | -0.1944710000 | -3.5872880000 | 1.2891450000 |
| H | 1.9009080000 | -4.9305960000 | 2.5097890000 |
| O | -0.4227930000 | -2.7252600000 | -2.0057780000 |
| H | -1.2323880000 | -2.4608860000 | -2.4763360000 |
| H | -0.6303920000 | -3.5500760000 | -1.5349770000 |
| C | -5.1843820000 | 1.3509630000 | -3.0936400000 |
| C | -3.8344900000 | 1.1102460000 | -3.3340670000 |
| C | -2.8807530000 | 1.4411680000 | -2.3622840000 |
| C | -3.2569930000 | 1.9843460000 | -1.1176830000 |
| C | -4.6213330000 | 2.2232140000 | -0.9076470000 |
| C | -5.5746650000 | 1.9192320000 | -1.8789020000 |
| H | -5.9238660000 | 1.0985810000 | -3.8462890000 |
| H | -3.5176560000 | 0.6723190000 | -4.2767550000 |
| H | -4.9383300000 | 2.6544340000 | 0.0380320000 |
| H | -6.6238130000 | 2.1171830000 | -1.6842290000 |
| C | -2.2547230000 | 2.3419870000 | -0.0365620000 |
| C | -1.6786420000 | 3.7624640000 | -0.1622900000 |
| H | -1.4231780000 | 1.6296710000 | -0.0283110000 |
| H | -2.7524410000 | 2.2531080000 | 0.9359260000 |
| H | -1.0423800000 | 3.9917720000 | 0.6985950000 |
| H | -1.0728400000 | 3.8648860000 | -1.0682340000 |
| H | -2.4765190000 | 4.5100370000 | -0.2020300000 |
| C | -1.4585450000 | 1.0737640000 | -2.6134410000 |
| O | -1.0066880000 | -0.0047250000 | -2.1617230000 |
| N | -0.6761730000 | 1.9323390000 | -3.2928740000 |

| | | | |
|---|---|---|---|
| C | -0.8232450000 | 3.0952500000 | -4.2534880000 |
| C | -2.1317180000 | 3.8562240000 | -4.0021210000 |
| C | -0.7887580000 | 2.5227750000 | -5.6767180000 |
| H | -2.0886110000 | 4.7558790000 | -4.6226000000 |
| H | -3.0137690000 | 3.2901080000 | -4.2957900000 |
| H | -2.2397150000 | 4.1704970000 | -2.9629680000 |
| H | 0.1401910000 | 1.9806180000 | -5.8695030000 |
| H | -1.6369730000 | 1.8527430000 | -5.8484280000 |
| H | -0.8562020000 | 3.3450950000 | -6.3944610000 |
| F | 0.6262480000 | 1.3880390000 | -3.4636380000 |
| C | 0.3660150000 | 4.0326270000 | -3.9882490000 |
| H | 0.3504550000 | 4.4015400000 | -2.9572990000 |
| H | 1.3240350000 | 3.5485530000 | -4.1857160000 |
| H | 0.2830620000 | 4.8949750000 | -4.6542980000 |
| O | -1.5742760000 | -1.1741980000 | 0.2962770000 |
| H | -1.6995980000 | -1.0198670000 | 1.2470440000 |
| H | -2.1930600000 | -0.5817730000 | -0.1685250000 |

**$^5Im_2Wat_3$_13**
Charge: 2
Multiplicity: 5
B3LYP-D3(BJ)/6-31G(d)-SDD SCF energy (au): -1541.47036790
B3LYP-D3(BJ)/6-31G(d)-SDD enthalpy (au): -1540.91940290
B3LYP-D3(BJ)/6-31G(d)-SDD free energy (au): -1541.03258790
B3LYP-D3(BJ)/def2-TZVP/SMD SCF energy (au): -2682.07872557
B3LYP-D3(BJ)/def2-TZVP/SMD enthalpy (au): -2681.52776057
B3LYP-D3(BJ)/def2-TZVP/SMD free energy (au): -2681.64094557
B3LYP-D3(BJ)/def2-TZVP/SMD free energy (quasi-harmonic) (au): -2681.62973867

Cartesian coordinates

| ATOM | X | Y | Z |
|---|---|---|---|
| Fe | -0.0195520000 | 0.6318740000 | -1.1762060000 |
| C | 2.2482340000 | 1.9190960000 | 2.3241040000 |
| N | 2.6668940000 | 2.7777370000 | 1.3271160000 |
| H | 3.2950190000 | 3.5618000000 | 1.4475260000 |
| C | 2.0900260000 | 2.3992290000 | 0.1641840000 |
| H | 2.2343790000 | 2.8987180000 | -0.7813640000 |
| N | 1.3199570000 | 1.3359830000 | 0.3610600000 |
| C | 1.4094600000 | 1.0290960000 | 1.7097630000 |
| H | 0.8738200000 | 0.1963870000 | 2.1401220000 |
| O | 1.4311440000 | 1.0755260000 | -2.7637560000 |
| H | 1.9423830000 | 0.2890460000 | -3.0172110000 |
| H | 0.8960510000 | 1.3286460000 | -3.5542750000 |
| H | 2.5754190000 | 2.0192640000 | 3.3471770000 |
| C | 1.0370210000 | -3.5850850000 | -1.2558200000 |
| N | 2.0252440000 | -3.0833380000 | -0.4307810000 |
| H | 2.7656460000 | -3.6214160000 | 0.0012220000 |
| C | 1.8394910000 | -1.7522720000 | -0.2979830000 |
| H | 2.4685140000 | -1.1074850000 | 0.2957060000 |
| N | 0.7745560000 | -1.3689260000 | -0.9990610000 |
| C | 0.2662180000 | -2.5101690000 | -1.6042100000 |
| H | -0.5982180000 | -2.4576380000 | -2.2456550000 |
| H | 0.9772130000 | -4.6313600000 | -1.5116360000 |

```
O     -1.3426690000   -0.2005640000   -2.7518200000
H     -1.2409620000    0.4162310000   -3.5169680000
H     -2.2363570000   -0.0617710000   -2.3974570000
F     -0.8962680000    2.2587910000   -1.3743490000
C     -1.0456220000    2.6144120000   -5.0749270000
O     -0.4760150000    1.5926920000   -4.6303770000
N     -0.4430560000    3.6037430000   -5.7387070000
H     -1.0249570000    4.4033870000   -5.9591110000
C      1.0123550000    3.7699970000   -6.0557550000
C      1.0871420000    4.9700820000   -7.0090930000
C      1.7796580000    4.0745160000   -4.7603590000
C      1.5517430000    2.5138790000   -6.7524750000
H      0.5272680000    4.7811430000   -7.9318470000
H      0.6980550000    5.8806090000   -6.5379270000
H      2.1277200000    5.1604540000   -7.2832890000
H      1.7419700000    3.2383250000   -4.0586230000
H      2.8302110000    4.2695660000   -4.9970580000
H      1.3711780000    4.9623430000   -4.2671680000
H      2.5990180000    2.6785300000   -7.0216560000
H      1.4956990000    1.6337920000   -6.1097040000
H      0.9990710000    2.3084450000   -7.6746160000
C     -2.5200980000    2.8063830000   -4.8399730000
C     -2.9190560000    2.9724210000   -3.5236420000
C     -3.4900120000    2.9059650000   -5.9043240000
C     -4.2546990000    3.3027850000   -3.2262270000
H     -2.1881150000    2.8757140000   -2.7203920000
C     -4.8292770000    3.2898130000   -5.5574750000
C     -5.2045340000    3.4796450000   -4.2417000000
H     -4.5427570000    3.4471430000   -2.1892190000
H     -5.5625340000    3.3850750000   -6.3526600000
H     -6.2249560000    3.7546600000   -3.9976880000
C     -3.2747790000    2.5483200000   -7.2372310000
C     -2.1278310000    1.8591240000   -7.8614420000
H     -1.4202620000    1.4208360000   -7.1599690000
H     -2.5097510000    1.0768050000   -8.5311920000
H     -1.5910340000    2.5604070000   -8.5210860000
H     -4.1020400000    2.7618300000   -7.9134300000
O     -1.7390910000    0.0780910000    0.1520710000
H     -2.0054290000    0.8905100000    0.6163060000
H     -1.6517360000   -0.6163330000    0.8255330000
```

**$^{5}$Im$_2$Wat$_3$_FeF_remove-F**

Charge: 2
Multiplicity: 5
B3LYP-D3(BJ)/6-31G(d)-SDD SCF energy (au): -805.286859892
B3LYP-D3(BJ)/6-31G(d)-SDD enthalpy (au): -805.045067892
B3LYP-D3(BJ)/6-31G(d)-SDD free energy (au): -805.119312892
B3LYP-D3(BJ)/def2-TZVP/SMD SCF energy (au): -1945.64142437
B3LYP-D3(BJ)/def2-TZVP/SMD enthalpy (au): -1945.39963237
B3LYP-D3(BJ)/def2-TZVP/SMD free energy (au): -1945.47387737
B3LYP-D3(BJ)/def2-TZVP/SMD free energy (quasi-harmonic) (au): -1945.467435478

Cartesian coordinates

| ATOM | X | Y | Z |
|---|---|---|---|
| Fe | 0.7890410000 | -0.5303620000 | -0.6914240000 |
| C | -0.0337380000 | 2.1328810000 | 2.4983000000 |
| N | 1.1741890000 | 2.6208260000 | 2.0395540000 |
| H | 1.6696650000 | 3.4207400000 | 2.4174870000 |
| C | 1.5863560000 | 1.8671670000 | 1.0062240000 |
| H | 2.5068580000 | 2.0249620000 | 0.4668960000 |
| N | 0.6923180000 | 0.9015230000 | 0.7668220000 |
| C | -0.3262920000 | 1.0629130000 | 1.7023670000 |
| H | -1.1776650000 | 0.4029140000 | 1.7354490000 |
| O | 2.7571850000 | 0.0539580000 | -1.4272730000 |
| H | 3.5023550000 | -0.5540640000 | -1.2811570000 |
| H | 3.0645940000 | 0.7429000000 | -2.0410570000 |
| H | -0.5607100000 | 2.5803120000 | 3.3268940000 |
| C | 2.4379750000 | -4.4711980000 | -0.3617190000 |
| N | 2.3740280000 | -4.1029360000 | 0.9652640000 |
| H | 2.6709670000 | -4.6706660000 | 1.7508400000 |
| C | 1.8530420000 | -2.8617240000 | 1.0486860000 |
| H | 1.7003280000 | -2.3313270000 | 1.9771200000 |
| N | 1.5758000000 | -2.4041320000 | -0.1738540000 |
| C | 1.9404380000 | -3.4081570000 | -1.0645950000 |
| H | 1.8094740000 | -3.2999890000 | -2.1306940000 |
| H | 2.8178460000 | -5.4304180000 | -0.6783040000 |
| O | 0.1593960000 | -0.5601900000 | -2.6941730000 |
| H | 0.6416420000 | -0.3183990000 | -3.5029110000 |
| H | -0.7079710000 | -0.9159140000 | -2.9534550000 |
| O | -1.2427170000 | -1.3149900000 | -0.4002930000 |
| H | -1.3753450000 | -2.1993810000 | -0.0182210000 |
| H | -2.1057850000 | -0.8662090000 | -0.3945520000 |

**$^{6}Im_2Wat_3$_FeF**

Charge: 2
Multiplicity: 6
B3LYP-D3(BJ)/6-31G(d)-SDD SCF energy (au): -905.123838114
B3LYP-D3(BJ)/6-31G(d)-SDD enthalpy (au): -904.877612114
B3LYP-D3(BJ)/6-31G(d)-SDD free energy (au): -904.948797114
B3LYP-D3(BJ)/def2-TZVP/SMD SCF energy (au): -2045.52474341
B3LYP-D3(BJ)/def2-TZVP/SMD enthalpy (au): -2045.27851741
B3LYP-D3(BJ)/def2-TZVP/SMD free energy (au): -2045.34970241
B3LYP-D3(BJ)/def2-TZVP/SMD free energy (quasi-harmonic) (au): -2045.346362296

Cartesian coordinates

| ATOM | X | Y | Z |
|---|---|---|---|
| Fe | 0.6212690000 | -0.3050430000 | -0.9737230000 |
| C | 0.3150530000 | 1.6464850000 | 2.7941680000 |
| N | 1.4663810000 | 2.1992330000 | 2.2662810000 |
| H | 2.0171810000 | 2.9368360000 | 2.6924840000 |
| C | 1.7308850000 | 1.6192830000 | 1.0850200000 |
| H | 2.5690990000 | 1.8668170000 | 0.4533230000 |
| N | 0.7947100000 | 0.6996480000 | 0.8213240000 |
| C | -0.1009960000 | 0.7139710000 | 1.8874440000 |
| H | -0.9707820000 | 0.0779960000 | 1.9090040000 |
| O | 2.5754190000 | 0.3046490000 | -1.5753870000 |

```
H      3.3070820000   -0.3281620000   -1.6897690000
H      2.5789180000    0.9104620000   -2.3409550000
H     -0.0975670000    1.9636130000    3.7397070000
C      2.5249380000   -4.0569090000   -0.1325750000
N      2.1454470000   -3.6290350000    1.1248620000
H      2.3265810000   -4.1194190000    1.9944320000
C      1.5013490000   -2.4553710000    1.0132020000
H      1.1117030000   -1.8909050000    1.8465650000
N      1.4362340000   -2.0983540000   -0.2756990000
C      2.0807730000   -3.0994720000   -0.9996670000
H      2.1855420000   -3.0536680000   -2.0722980000
H      3.0634160000   -4.9788280000   -0.2919570000
O      0.3896710000   -1.3907470000   -2.8796850000
H     -0.0058440000   -0.7983860000   -3.5464270000
H      0.0711480000   -2.2913210000   -3.0637610000
F      0.0124190000    1.0385550000   -1.9801310000
O     -1.3748210000   -0.8384740000   -0.4283010000
H     -1.7399400000   -1.7397420000   -0.4771870000
H     -2.0466250000   -0.2262760000   -0.7863420000
```

**$^{5}Im_2Wat_3$_TS-1**

```
Charge: 2
Multiplicity: 5
B3LYP-D3(BJ)/6-31G(d)-SDD SCF energy (au):              -1541.39191887
B3LYP-D3(BJ)/6-31G(d)-SDD enthalpy (au):                -1540.84375687
B3LYP-D3(BJ)/6-31G(d)-SDD free energy (au):             -1540.95204187
B3LYP-D3(BJ)/def2-TZVP/SMD SCF energy (au):             -2682.00860728
B3LYP-D3(BJ)/def2-TZVP/SMD enthalpy (au):               -2681.46044528
B3LYP-D3(BJ)/def2-TZVP/SMD free energy (au):            -2681.56873028
B3LYP-D3(BJ)/def2-TZVP/SMD free energy (quasi-harmonic) (au): -2681.55967541

Cartesian coordinates
ATOM    X               Y               Z
Fe     0.6111460000   -0.2947400000   -0.9807000000
C      0.1966610000    1.8541370000    2.7074650000
N      1.4817470000    2.1895660000    2.3253410000
H      2.0981070000    2.8263860000    2.8152270000
C      1.7816440000    1.5291410000    1.1886920000
H      2.7187440000    1.6096580000    0.6601810000
N      0.7474830000    0.7754060000    0.8200010000
C     -0.2530450000    0.9744170000    1.7619450000
H     -1.2082410000    0.4822980000    1.6762650000
O      2.6527180000    0.2794420000   -1.5466510000
H      3.3031760000   -0.4306830000   -1.6799230000
H      2.6483510000    0.8189640000   -2.3560110000
H     -0.2652490000    2.2606130000    3.5935950000
C      2.5943410000   -4.0788710000   -0.1991910000
N      2.2243330000   -3.6845270000    1.0694740000
H      2.4232000000   -4.1873840000    1.9259780000
C      1.5559020000   -2.5138310000    0.9806740000
H      1.1657730000   -1.9710440000    1.8286350000
N      1.4735830000   -2.1317910000   -0.2916090000
C      2.1233680000   -3.1054760000   -1.0383960000
```

| | | | |
|---|---|---|---|
| H | 2.2065320000 | -3.0399080000 | -2.1130860000 |
| H | 3.1415150000 | -4.9902430000 | -0.3845940000 |
| O | 0.2421540000 | -1.2137760000 | -2.9131810000 |
| H | -0.5655230000 | -0.6990750000 | -3.1753960000 |
| H | 0.0136700000 | -2.1541640000 | -2.9994920000 |
| C | -5.9345040000 | 2.1038370000 | -3.0677890000 |
| C | -4.6753010000 | 1.5491520000 | -3.2111180000 |
| C | -3.6406580000 | 1.9007710000 | -2.3124660000 |
| C | -3.8729230000 | 2.7895290000 | -1.2282290000 |
| C | -5.1624950000 | 3.3193000000 | -1.1090740000 |
| C | -6.1722190000 | 3.0032200000 | -2.0161790000 |
| H | -6.7301150000 | 1.8431200000 | -3.7574760000 |
| H | -4.4631600000 | 0.8510910000 | -4.0144910000 |
| H | -5.3819080000 | 3.9820880000 | -0.2770410000 |
| H | -7.1570530000 | 3.4441490000 | -1.8971230000 |
| C | -2.8510580000 | 3.1093260000 | -0.1570500000 |
| C | -2.2920340000 | 4.5372420000 | -0.2466600000 |
| H | -2.0237690000 | 2.3958380000 | -0.1820300000 |
| H | -3.3462690000 | 2.9842590000 | 0.8141220000 |
| H | -1.6192130000 | 4.7350470000 | 0.5938350000 |
| H | -1.7289260000 | 4.6747520000 | -1.1740480000 |
| H | -3.0917850000 | 5.2838610000 | -0.2198080000 |
| C | -2.3218000000 | 1.3144220000 | -2.5990590000 |
| O | -2.1251240000 | 0.1013640000 | -2.7969490000 |
| N | -1.3596410000 | 2.2954610000 | -2.6114530000 |
| C | -0.8763670000 | 2.9428310000 | -3.8303850000 |
| C | -2.0424150000 | 3.8896160000 | -4.2732960000 |
| C | -0.5759800000 | 1.9372320000 | -4.9526740000 |
| H | -1.6643950000 | 4.4733890000 | -5.1174520000 |
| H | -2.9171370000 | 3.3226880000 | -4.5954160000 |
| H | -2.3288250000 | 4.5707350000 | -3.4702420000 |
| H | 0.2134450000 | 1.2457810000 | -4.6484610000 |
| H | -1.4640310000 | 1.3623580000 | -5.2315150000 |
| H | -0.2359280000 | 2.4820300000 | -5.8377180000 |
| F | 0.1992730000 | 1.2718280000 | -2.0658410000 |
| C | 0.3440730000 | 3.8010480000 | -3.4742520000 |
| H | 0.1138680000 | 4.4841060000 | -2.6524740000 |
| H | 1.1834800000 | 3.1701550000 | -3.1784960000 |
| H | 0.6377410000 | 4.3910290000 | -4.3464900000 |
| O | -1.3981750000 | -0.9308400000 | -0.4246580000 |
| H | -1.5186190000 | -1.8643490000 | -0.1835850000 |
| H | -2.0020200000 | -0.7382880000 | -1.1787820000 |

**$^{5}$Im$_2$Wat$_3$_TS-3-F**
Charge: 2
Multiplicity: 5
B3LYP-D3(BJ)/6-31G(d)-SDD SCF energy (au): -1541.47459734
B3LYP-D3(BJ)/6-31G(d)-SDD enthalpy (au): -1540.92533834
B3LYP-D3(BJ)/6-31G(d)-SDD free energy (au): -1541.03668634
B3LYP-D3(BJ)/def2-TZVP/SMD SCF energy (au): -2682.08321435
B3LYP-D3(BJ)/def2-TZVP/SMD enthalpy (au): -2681.53395535
B3LYP-D3(BJ)/def2-TZVP/SMD free energy (au): -2681.64530335
B3LYP-D3(BJ)/def2-TZVP/SMD free energy (quasi-harmonic) (au): -2681.63477801

Cartesian coordinates

| ATOM | X | Y | Z |
|---|---|---|---|
| Fe | -0.0850670000 | 0.5774010000 | -1.0041760000 |
| C | 1.9834590000 | 1.3350250000 | 2.7798070000 |
| N | 2.6212190000 | 2.1927260000 | 1.9057990000 |
| H | 3.3503640000 | 2.8502020000 | 2.1497610000 |
| C | 2.1013570000 | 2.0076100000 | 0.6693610000 |
| H | 2.4144520000 | 2.5462980000 | -0.2120780000 |
| N | 1.1607380000 | 1.0721570000 | 0.7024790000 |
| C | 1.0778240000 | 0.6467290000 | 2.0178230000 |
| H | 0.3778890000 | -0.1159660000 | 2.3237290000 |
| O | 1.3863720000 | 1.5501050000 | -2.4103280000 |
| H | 1.9607430000 | 0.8423890000 | -2.7473610000 |
| H | 0.8465670000 | 1.8464850000 | -3.1770410000 |
| H | 2.2240100000 | 1.2984840000 | 3.8308200000 |
| C | 1.5156060000 | -3.3448110000 | -2.0182230000 |
| N | 2.2872610000 | -3.0185610000 | -0.9193740000 |
| H | 3.0062050000 | -3.5992890000 | -0.5073410000 |
| C | 1.9182780000 | -1.7932800000 | -0.4849510000 |
| H | 2.3553140000 | -1.2936550000 | 0.3655220000 |
| N | 0.9439820000 | -1.3114970000 | -1.2518930000 |
| C | 0.6841670000 | -2.2763250000 | -2.2151100000 |
| H | -0.0767730000 | -2.1187680000 | -2.9621370000 |
| H | 1.6268740000 | -4.2816840000 | -2.5412700000 |
| O | -1.2660900000 | -0.0201360000 | -2.8185010000 |
| H | -1.2735090000 | 0.7918650000 | -3.3714050000 |
| H | -2.1511170000 | -0.1067530000 | -2.4275770000 |
| F | -1.2139670000 | 2.0994260000 | -0.8919070000 |
| C | -1.0668030000 | 3.1375700000 | -4.9264020000 |
| O | -0.5425510000 | 2.2403770000 | -4.2360300000 |
| N | -0.4346520000 | 3.8936550000 | -5.8241560000 |
| H | -0.9884070000 | 4.6072980000 | -6.2836920000 |
| C | 1.0097660000 | 3.8248840000 | -6.2234490000 |
| C | 1.1673760000 | 4.8252270000 | -7.3752440000 |
| C | 1.8857540000 | 4.2439890000 | -5.0336070000 |
| C | 1.3434750000 | 2.4053570000 | -6.7025820000 |
| H | 0.5365130000 | 4.5522450000 | -8.2284020000 |
| H | 0.9164770000 | 5.8440540000 | -7.0575170000 |
| H | 2.2050210000 | 4.8321090000 | -7.7180040000 |
| H | 1.7859640000 | 3.5571010000 | -4.1896310000 |
| H | 2.9361540000 | 4.2489740000 | -5.3396240000 |
| H | 1.6301440000 | 5.2549280000 | -4.6987590000 |
| H | 2.3830320000 | 2.3755830000 | -7.0418190000 |
| H | 1.2190370000 | 1.6702690000 | -5.9046490000 |
| H | 0.7049080000 | 2.1187680000 | -7.5439870000 |
| C | -2.5480130000 | 3.3725700000 | -4.7830240000 |
| C | -3.3673840000 | 3.0937280000 | -5.8635840000 |
| C | -3.1444400000 | 3.7034200000 | -3.5172580000 |
| C | -4.7692530000 | 3.1142200000 | -5.7306920000 |
| H | -2.9263840000 | 2.8237910000 | -6.8180400000 |
| C | -4.5664280000 | 3.6700740000 | -3.4058690000 |
| C | -5.3703240000 | 3.3902620000 | -4.5000890000 |

| | | | |
|---|---|---|---|
| H | -5.3865430000 | 2.8949780000 | -6.5967660000 |
| H | -5.0190830000 | 3.9110790000 | -2.4486400000 |
| H | -6.4504280000 | 3.3898790000 | -4.4032820000 |
| C | -2.4526500000 | 4.1861740000 | -2.3920270000 |
| C | -1.0612680000 | 4.5648730000 | -2.2247400000 |
| H | -0.4847340000 | 4.7106740000 | -3.1351620000 |
| H | -0.9723490000 | 5.4231930000 | -1.5495970000 |
| H | -0.6757460000 | 3.6855750000 | -1.6485590000 |
| H | -3.0629210000 | 4.3009440000 | -1.4974240000 |
| O | -1.8267950000 | -0.4630420000 | 0.0293330000 |
| H | -2.2386980000 | 0.2057750000 | 0.6021330000 |
| H | -1.7130790000 | -1.2631300000 | 0.5674340000 |

**$^{5}Im_3OHN_3$_11**

Charge: 0
Multiplicity: 5
B3LYP-D3(BJ)/6-31G(d)-SDD SCF energy (au): -1778.85087037
B3LYP-D3(BJ)/6-31G(d)-SDD enthalpy (au): -1778.27440437
B3LYP-D3(BJ)/6-31G(d)-SDD free energy (au): -1778.39363737
B3LYP-D3(BJ)/def2-TZVP/SMD SCF energy (au): -2919.40274584
B3LYP-D3(BJ)/def2-TZVP/SMD enthalpy (au): -2918.82627984
B3LYP-D3(BJ)/def2-TZVP/SMD free energy (au): -2918.94551284
B3LYP-D3(BJ)/def2-TZVP/SMD free energy (quasi-harmonic) (au): -2918.93436047

Cartesian coordinates

| ATOM | X | Y | Z |
|---|---|---|---|
| Fe | 1.1474490000 | 0.1577120000 | 0.7495890000 |
| C | -0.1085660000 | 1.5685710000 | -3.2527010000 |
| N | 0.9856850000 | 0.7827700000 | -3.5667310000 |
| H | 1.2845420000 | 0.5206550000 | -4.4939510000 |
| C | 1.5352070000 | 0.3377980000 | -2.4067340000 |
| H | 2.3699180000 | -0.3476150000 | -2.3641740000 |
| N | 0.8572600000 | 0.8078320000 | -1.3722130000 |
| C | -0.1709130000 | 1.5743570000 | -1.8858370000 |
| H | -0.8865550000 | 2.0463890000 | -1.2348240000 |
| C | 0.1314440000 | 4.4673660000 | 1.6427450000 |
| N | -0.0850440000 | 3.7490180000 | 2.8065330000 |
| H | -0.4297940000 | 4.1193630000 | 3.6795920000 |
| C | 0.2457390000 | 2.4506250000 | 2.5704100000 |
| H | 0.2110930000 | 1.6405420000 | 3.2857000000 |
| N | 0.6595500000 | 2.3039820000 | 1.3239050000 |
| C | 0.5951620000 | 3.5505670000 | 0.7348610000 |
| H | 0.8838540000 | 3.6966720000 | -0.2950980000 |
| H | -0.0596920000 | 5.5271010000 | 1.5765790000 |
| H | -0.7343430000 | 2.0122780000 | -4.0097840000 |
| C | 5.2863820000 | 1.5868290000 | 1.1116650000 |
| N | 5.5306030000 | 0.4059250000 | 0.4324810000 |
| H | 6.4361900000 | 0.0169770000 | 0.2157700000 |
| C | 4.3325770000 | -0.1574060000 | 0.1194670000 |
| H | 4.2275820000 | -1.0871400000 | -0.4263430000 |
| N | 3.3413870000 | 0.6027240000 | 0.5563800000 |
| C | 3.9225500000 | 1.6905200000 | 1.1778350000 |
| H | 3.3118790000 | 2.4528340000 | 1.6351420000 |

| | | | |
|---|---|---|---|
| H | 6.0836460000 | 2.2154890000 | 1.4760820000 |
| O | 1.1327950000 | -0.2242360000 | 2.6326290000 |
| H | 1.4776950000 | -1.1216550000 | 2.7548060000 |
| C | -1.3455630000 | -1.6952980000 | -3.5505220000 |
| C | -1.2060360000 | -1.7176190000 | -2.1646390000 |
| C | -2.1221720000 | -1.0332290000 | -1.3629160000 |
| C | -3.2019080000 | -0.3328750000 | -1.9249500000 |
| C | -3.3229360000 | -0.3265170000 | -3.3193370000 |
| C | -2.4050850000 | -0.9947760000 | -4.1294220000 |
| H | -0.6311620000 | -2.2285500000 | -4.1714320000 |
| H | -0.3848000000 | -2.2404650000 | -1.6872950000 |
| H | -4.1568000000 | 0.2059060000 | -3.7715260000 |
| H | -2.5265360000 | -0.9797590000 | -5.2096140000 |
| C | -4.1983310000 | 0.4178270000 | -1.0675300000 |
| C | -3.7253360000 | 1.8327080000 | -0.6902380000 |
| H | -4.4010550000 | -0.1486020000 | -0.1510410000 |
| H | -5.1497770000 | 0.4852020000 | -1.6086640000 |
| H | -3.5048200000 | 2.4201850000 | -1.5892910000 |
| H | -4.4987820000 | 2.3607260000 | -0.1206330000 |
| H | -2.8203640000 | 1.7837550000 | -0.0786940000 |
| C | -1.8869560000 | -0.9864230000 | 0.1213880000 |
| O | -1.2774540000 | -0.0581470000 | 0.6577780000 |
| N | -2.4825640000 | -1.9719980000 | 0.8566090000 |
| C | -2.1779090000 | -2.5020720000 | 2.2129060000 |
| C | -1.8671510000 | -1.3298670000 | 3.1571080000 |
| C | -0.9887900000 | -3.4684270000 | 2.1134570000 |
| H | -2.6489910000 | -0.5652830000 | 3.0992590000 |
| H | -1.8618980000 | -1.7334050000 | 4.1768220000 |
| H | -0.8935040000 | -0.8756450000 | 2.9526090000 |
| H | -1.2376180000 | -4.3203970000 | 1.4730750000 |
| H | -0.1136460000 | -2.9644490000 | 1.6946770000 |
| H | -0.7435630000 | -3.8478080000 | 3.1115930000 |
| F | -2.9568110000 | -3.0177430000 | 0.0362120000 |
| C | -3.4456780000 | -3.2253380000 | 2.6965920000 |
| H | -4.2915090000 | -2.5312410000 | 2.7460660000 |
| H | -3.7126090000 | -4.0568150000 | 2.0409530000 |
| H | -3.2639920000 | -3.6209210000 | 3.7005580000 |
| N | 2.9213140000 | -2.5341540000 | -1.5694130000 |
| N | 2.1136950000 | -2.2399600000 | -0.7694420000 |
| N | 1.2815920000 | -1.9216580000 | 0.0290150000 |

**$^{5}Im_3OHN_3$_13**

Charge: 0
Multiplicity: 5
B3LYP-D3(BJ)/6-31G(d)-SDD SCF energy (au): -1778.90731059
B3LYP-D3(BJ)/6-31G(d)-SDD enthalpy (au): -1778.33179559
B3LYP-D3(BJ)/6-31G(d)-SDD free energy (au): -1778.45313159
B3LYP-D3(BJ)/def2-TZVP/SMD SCF energy (au): -2919.47186337
B3LYP-D3(BJ)/def2-TZVP/SMD enthalpy (au): -2918.89634837
B3LYP-D3(BJ)/def2-TZVP/SMD free energy (au): -2919.01768437
B3LYP-D3(BJ)/def2-TZVP/SMD free energy (quasi-harmonic) (au): -2919.00419978

Cartesian coordinates

| ATOM | X | Y | Z |
|---|---|---|---|
| H | 33.1174460000 | 26.1231140000 | 36.2656960000 |
| C | 33.4014470000 | 27.1623930000 | 36.2130210000 |
| C | 34.6195600000 | 27.7876710000 | 36.2117300000 |
| N | 32.4672340000 | 28.1785010000 | 36.1226940000 |
| C | 33.1322760000 | 29.3595470000 | 36.0545710000 |
| N | 34.4380080000 | 29.1542230000 | 36.1166870000 |
| H | 31.4710440000 | 28.0663950000 | 36.0073710000 |
| H | 35.6085800000 | 27.3613640000 | 36.2616330000 |
| H | 32.6605470000 | 30.3203600000 | 35.9301120000 |
| H | 39.9575000000 | 27.2098870000 | 36.5324760000 |
| C | 39.1809160000 | 27.8001780000 | 36.0713890000 |
| C | 38.1436480000 | 28.5341240000 | 36.5870260000 |
| N | 39.0783350000 | 27.9402220000 | 34.6979920000 |
| C | 38.0049860000 | 28.7354730000 | 34.4368640000 |
| N | 37.4207640000 | 29.1039830000 | 35.5595690000 |
| H | 39.6990710000 | 27.5412550000 | 34.0096630000 |
| H | 37.8658590000 | 28.6939470000 | 37.6178990000 |
| H | 37.6779390000 | 29.0363620000 | 33.4527470000 |
| H | 36.3322630000 | 30.7889810000 | 40.9214290000 |
| C | 36.4577250000 | 30.9462610000 | 39.8615970000 |
| C | 35.8050530000 | 30.4406370000 | 38.7675730000 |
| N | 37.4182250000 | 31.8053320000 | 39.3522140000 |
| C | 37.3185150000 | 31.7934380000 | 37.9971720000 |
| N | 36.3547120000 | 30.9738200000 | 37.6230060000 |
| H | 38.0634710000 | 32.3638180000 | 39.8906810000 |
| H | 34.9839660000 | 29.7432910000 | 38.7127090000 |
| H | 37.9254130000 | 32.3619940000 | 37.3093690000 |
| Fe | 35.8255490000 | 30.8730370000 | 35.4616480000 |
| O | 34.4209040000 | 32.0422770000 | 35.7209470000 |
| H | 33.7061420000 | 31.9850330000 | 35.0557240000 |
| F | 37.3279620000 | 31.9187630000 | 35.1835520000 |
| N | 31.2702130000 | 31.2822540000 | 31.9702920000 |
| O | 32.1680220000 | 31.6925920000 | 34.0169040000 |
| C | 33.4745220000 | 32.2947210000 | 32.1049370000 |
| C | 34.0545930000 | 33.3802920000 | 32.7593370000 |
| C | 35.1479720000 | 34.0571350000 | 32.2215200000 |
| C | 35.6713830000 | 33.6451330000 | 30.9903600000 |
| C | 35.1260870000 | 32.5584370000 | 30.3294150000 |
| C | 34.0329990000 | 31.8169770000 | 30.8670690000 |
| C | 32.2710520000 | 31.7105760000 | 32.7785800000 |
| C | 30.0159580000 | 30.6400200000 | 32.4075490000 |
| C | 30.3214170000 | 29.4042100000 | 33.2717990000 |
| C | 29.1613030000 | 31.6504980000 | 33.1908570000 |
| C | 29.2886710000 | 30.2092590000 | 31.1271060000 |
| C | 33.6030640000 | 30.6300640000 | 30.2238210000 |
| C | 34.1694970000 | 30.1274910000 | 28.9325300000 |
| H | 33.6433940000 | 33.6917090000 | 33.7118350000 |
| H | 35.5876990000 | 34.8898440000 | 32.7604040000 |
| H | 36.5211490000 | 34.1664870000 | 30.5578670000 |
| H | 35.5715230000 | 32.2271730000 | 29.3976920000 |
| H | 31.4216670000 | 31.3731570000 | 30.9756040000 |
| H | 32.9633590000 | 29.9509770000 | 30.7723310000 |

| | | | |
|---|---|---|---|
| H | 35.1923670000 | 29.7382540000 | 29.0613410000 |
| H | 33.5623070000 | 29.3073310000 | 28.5371920000 |
| H | 34.2240830000 | 30.9083190000 | 28.1630400000 |
| H | 29.3867590000 | 28.8819380000 | 33.5093570000 |
| H | 30.9967680000 | 28.7216290000 | 32.7493040000 |
| H | 30.8100590000 | 29.7110730000 | 34.1970870000 |
| H | 29.7085100000 | 32.0018380000 | 34.0681310000 |
| H | 28.9153960000 | 32.5139610000 | 32.5630060000 |
| H | 28.2250570000 | 31.1833230000 | 33.5179980000 |
| H | 29.8856500000 | 29.4833990000 | 30.5624980000 |
| H | 28.3313990000 | 29.7414660000 | 31.3764490000 |
| H | 29.0841100000 | 31.0718460000 | 30.4810980000 |
| N | 35.5278690000 | 30.1720220000 | 33.5507220000 |
| N | 34.6187840000 | 29.4555590000 | 33.2246920000 |
| N | 33.7517150000 | 28.7656080000 | 32.8732890000 |

**$^{5}Im_3OHN_3$_FeF_remove-F**

Charge: 0
Multiplicity: 5
B3LYP-D3(BJ)/6-31G(d)-SDD SCF energy (au): -1042.77025016
B3LYP-D3(BJ)/6-31G(d)-SDD enthalpy (au): -1042.50234616
B3LYP-D3(BJ)/6-31G(d)-SDD free energy (au): -1042.58180416
B3LYP-D3(BJ)/def2-TZVP/SMD SCF energy (au): -2183.03131501
B3LYP-D3(BJ)/def2-TZVP/SMD enthalpy (au): -2182.76341101
B3LYP-D3(BJ)/def2-TZVP/SMD free energy (au): -2182.84286901
B3LYP-D3(BJ)/def2-TZVP/SMD free energy (quasi-harmonic) (au): -2182.83679385

Cartesian coordinates

| ATOM | X | Y | Z |
|---|---|---|---|
| Fe | 0.4786150000 | -0.2373030000 | -1.3442140000 |
| C | 0.1728390000 | 0.7685970000 | 2.9912350000 |
| N | 0.5388300000 | 1.9887100000 | 2.4524980000 |
| H | 0.8617440000 | 2.7948560000 | 2.9670210000 |
| C | 0.5261870000 | 1.8705670000 | 1.1006660000 |
| H | 0.8229560000 | 2.6527790000 | 0.4188160000 |
| N | 0.1493990000 | 0.6559760000 | 0.7438820000 |
| C | -0.0661390000 | -0.0452880000 | 1.9154750000 |
| H | -0.3532260000 | -1.0854230000 | 1.8932900000 |
| C | -3.4891350000 | -2.1023460000 | -1.7825100000 |
| N | -3.7250680000 | -1.5583310000 | -0.5310200000 |
| H | -4.5857040000 | -1.6173040000 | -0.0076280000 |
| C | -2.5957640000 | -0.9014710000 | -0.1457790000 |
| H | -2.5025570000 | -0.3720170000 | 0.7901300000 |
| N | -1.6617140000 | -1.0013460000 | -1.0723080000 |
| C | -2.2061370000 | -1.7457970000 | -2.1045220000 |
| H | -1.6172140000 | -1.9151700000 | -2.9937830000 |
| H | -4.2413210000 | -2.6667570000 | -2.3111990000 |
| H | 0.1260280000 | 0.6019680000 | 4.0559700000 |
| C | 1.6976610000 | -4.1112060000 | 0.2999920000 |
| N | 2.7200060000 | -3.2946860000 | 0.7512230000 |
| H | 3.5348770000 | -3.5887280000 | 1.2690400000 |
| C | 2.4561820000 | -2.0221060000 | 0.3521120000 |
| H | 3.0891770000 | -1.1710250000 | 0.5743200000 |

N 1.3181480000 -1.9865670000 -0.3227650000
C 0.8350780000 -3.2791860000 -0.3636310000
H -0.0866060000 -3.5080640000 -0.8766550000
H 1.6879970000 -5.1741890000 0.4829540000
O 0.2545450000 -0.7069910000 -3.1564140000
H 1.0541150000 -0.4509870000 -3.6411850000
N 3.6174610000 1.0932350000 0.4978500000
N 2.9100440000 1.0672150000 -0.4335620000
N 2.1632950000 1.0508320000 -1.3735880000

**$^5Im_3OHN_3$_FeF_remove-$N_3$**

Charge: 0
Multiplicity: 5
B3LYP-D3(BJ)/6-31G(d)-SDD SCF energy (au): -978.437199641
B3LYP-D3(BJ)/6-31G(d)-SDD enthalpy (au): -978.182543641
B3LYP-D3(BJ)/6-31G(d)-SDD free energy (au): -978.258219641
B3LYP-D3(BJ)/def2-TZVP/SMD SCF energy (au): -2118.66981038
B3LYP-D3(BJ)/def2-TZVP/SMD enthalpy (au): -2118.41515438
B3LYP-D3(BJ)/def2-TZVP/SMD free energy (au): -2118.49083038
B3LYP-D3(BJ)/def2-TZVP/SMD free energy (quasi-harmonic) (au): -2118.485611739

Cartesian coordinates

| ATOM | X | Y | Z |
|---|---|---|---|
| Fe | -0.0906920000 | -0.8289540000 | -1.7987910000 |
| C | 0.6097590000 | 1.4010360000 | 3.5879920000 |
| N | 0.7788380000 | 1.7706090000 | 2.2659210000 |
| H | 0.5501720000 | 2.6632270000 | 1.8528490000 |
| C | 1.3218490000 | 0.7106000000 | 1.5987470000 |
| H | 1.4546750000 | 0.7263180000 | 0.5227980000 |
| N | 1.5108990000 | -0.3117650000 | 2.4149210000 |
| C | 1.0734370000 | 0.1085930000 | 3.6543030000 |
| H | 1.1283080000 | -0.5368270000 | 4.5205860000 |
| C | -1.7658080000 | -2.2133680000 | 1.9627750000 |
| N | -1.7971650000 | -0.8361660000 | 2.0962590000 |
| H | -1.8318320000 | -0.3233950000 | 2.9655390000 |
| C | -1.4924420000 | -0.2956520000 | 0.8892060000 |
| H | -1.3878130000 | 0.7605760000 | 0.7049670000 |
| N | -1.2930820000 | -1.2440780000 | -0.0086080000 |
| C | -1.4638930000 | -2.4477250000 | 0.6476770000 |
| H | -1.3495320000 | -3.3847650000 | 0.1244760000 |
| H | -1.9681850000 | -2.8741740000 | 2.7908350000 |
| H | 0.2280710000 | 2.0817530000 | 4.3337770000 |
| C | 2.7261190000 | -4.0638230000 | -1.3466190000 |
| N | 2.8200530000 | -3.5475290000 | -0.0648840000 |
| H | 3.4088540000 | -3.8947610000 | 0.6772780000 |
| C | 1.9917660000 | -2.4728300000 | 0.0254330000 |
| H | 1.8629900000 | -1.8853430000 | 0.9289000000 |
| N | 1.3822540000 | -2.2795510000 | -1.1337410000 |
| C | 1.8269110000 | -3.2632740000 | -1.9979440000 |
| H | 1.4407270000 | -3.3062780000 | -3.0052280000 |
| H | 3.2918610000 | -4.9267910000 | -1.6608020000 |
| O | -0.5760270000 | -1.7617500000 | -3.3407650000 |
| H | -1.0783410000 | -1.2859890000 | -4.0166630000 |

F      0.4449270000    0.8567310000   -1.1933130000

**$^{5}Im_3OHN_3$_FeF_remove-OH**
Charge: 0
Multiplicity: 5
B3LYP-D3(BJ)/6-31G(d)-SDD SCF energy (au):              -1066.82043566
B3LYP-D3(BJ)/6-31G(d)-SDD enthalpy (au):                -1066.56367666
B3LYP-D3(BJ)/6-31G(d)-SDD free energy (au):             -1066.64240966
B3LYP-D3(BJ)/def2-TZVP/SMD SCF energy (au):             -2207.09170294
B3LYP-D3(BJ)/def2-TZVP/SMD enthalpy (au):               -2206.83494394
B3LYP-D3(BJ)/def2-TZVP/SMD free energy (au):            -2206.91367694
B3LYP-D3(BJ)/def2-TZVP/SMD free energy (quasi-harmonic) (au): -2206.90733428

Cartesian coordinates

| ATOM | X | Y | Z |
|---|---|---|---|
| Fe | 0.5163210000 | -0.3038960000 | -0.6386170000 |
| C | 1.3631560000 | 1.3021980000 | 3.3673890000 |
| N | 2.0365580000 | 2.1588470000 | 2.5133660000 |
| H | 2.6135920000 | 2.9416540000 | 2.7836200000 |
| C | 1.8268390000 | 1.7398550000 | 1.2419580000 |
| H | 2.2319750000 | 2.2158810000 | 0.3633800000 |
| N | 1.0520530000 | 0.6667940000 | 1.2347500000 |
| C | 0.7546490000 | 0.3841480000 | 2.5546310000 |
| H | 0.1218550000 | -0.4495970000 | 2.8159610000 |
| C | -2.4404630000 | -2.6089680000 | 1.9915270000 |
| N | -3.1799040000 | -1.7081690000 | 1.2458710000 |
| H | -4.1739640000 | -1.5489320000 | 1.3186780000 |
| C | -2.3340100000 | -1.0730410000 | 0.3899770000 |
| H | -2.6043050000 | -0.3070430000 | -0.3220610000 |
| N | -1.0997170000 | -1.5175270000 | 0.5483520000 |
| C | -1.1513370000 | -2.4750270000 | 1.5424570000 |
| H | -0.2635720000 | -3.0061400000 | 1.8534520000 |
| H | -2.8893890000 | -3.2403010000 | 2.7425130000 |
| H | 1.3808960000 | 1.4277090000 | 4.4385560000 |
| C | 1.9674220000 | -4.4263160000 | -0.9825670000 |
| N | 3.1598490000 | -3.7407430000 | -0.8316140000 |
| H | 4.0871020000 | -4.1384510000 | -0.8653380000 |
| C | 2.8789150000 | -2.4258200000 | -0.6518160000 |
| H | 3.6364890000 | -1.6615380000 | -0.5240030000 |
| N | 1.5665570000 | -2.2362420000 | -0.6741500000 |
| C | 0.9883250000 | -3.4756970000 | -0.8798010000 |
| H | -0.0830290000 | -3.5833650000 | -0.9402580000 |
| H | 1.9326050000 | -5.4912230000 | -1.1495790000 |
| F | -0.9395940000 | 0.4791500000 | -1.4556530000 |
| N | 4.3834640000 | 0.4864590000 | -0.6463030000 |
| N | 3.3106530000 | 0.5425320000 | -1.1164500000 |
| N | 2.2111490000 | 0.5916690000 | -1.5900200000 |

**$^{6}Im_3OHN_3$_FeF**
Charge: 0
Multiplicity: 6
B3LYP-D3(BJ)/6-31G(d)-SDD SCF energy (au):              -1142.64169990
B3LYP-D3(BJ)/6-31G(d)-SDD enthalpy (au):                -1142.36998490

B3LYP-D3(BJ)/6-31G(d)-SDD free energy (au): -1142.45093790
B3LYP-D3(BJ)/def2-TZVP/SMD SCF energy (au): -2282.94860223
B3LYP-D3(BJ)/def2-TZVP/SMD enthalpy (au): -2282.67688723
B3LYP-D3(BJ)/def2-TZVP/SMD free energy (au): -2282.75784023
B3LYP-D3(BJ)/def2-TZVP/SMD free energy (quasi-harmonic) (au): -2282.75221733

Cartesian coordinates

| ATOM | X | Y | Z |
|---|---|---|---|
| Fe | 0.1792580000 | -0.3788940000 | -0.9852090000 |
| C | 1.6232030000 | 0.9610590000 | 3.1197600000 |
| N | 2.0334410000 | 1.9670510000 | 2.2617710000 |
| H | 2.4898600000 | 2.8277700000 | 2.5255480000 |
| C | 1.7188820000 | 1.5919190000 | 0.9942740000 |
| H | 1.9075820000 | 2.1855090000 | 0.1127350000 |
| N | 1.1365870000 | 0.4084640000 | 0.9965210000 |
| C | 1.0645090000 | 0.0036840000 | 2.3131720000 |
| H | 0.6099590000 | -0.9383450000 | 2.5804080000 |
| C | -2.5219350000 | -2.6651680000 | 1.7049750000 |
| N | -3.1356020000 | -1.4359720000 | 1.5294390000 |
| H | -4.0354630000 | -1.1617380000 | 1.8948610000 |
| C | -2.3307090000 | -0.6687490000 | 0.7508680000 |
| H | -2.5324880000 | 0.3402950000 | 0.4256690000 |
| N | -1.2404420000 | -1.3406980000 | 0.4283660000 |
| C | -1.3435360000 | -2.5860820000 | 1.0104960000 |
| H | -0.5736390000 | -3.3293000000 | 0.8742060000 |
| H | -2.9733290000 | -3.4553420000 | 2.2841360000 |
| H | 1.7615790000 | 1.0251370000 | 4.1877280000 |
| C | 2.4232540000 | -4.1704550000 | -0.8860690000 |
| N | 3.1886200000 | -3.4757000000 | 0.0369000000 |
| H | 4.0751420000 | -3.7698550000 | 0.4190170000 |
| C | 2.5988650000 | -2.2713120000 | 0.2495250000 |
| H | 3.0005740000 | -1.5107230000 | 0.8979250000 |
| N | 1.4966700000 | -2.1675660000 | -0.4718150000 |
| C | 1.3760040000 | -3.3434710000 | -1.1910570000 |
| H | 0.5539200000 | -3.4602140000 | -1.8799810000 |
| H | 2.6946030000 | -5.1566460000 | -1.2285030000 |
| O | -0.5006790000 | -1.3752290000 | -2.4142520000 |
| H | -0.0523150000 | -1.0931040000 | -3.2282630000 |
| F | -0.8870450000 | 1.1283880000 | -0.8457680000 |
| N | 4.0347710000 | 0.0634010000 | -1.1757250000 |
| N | 2.9448670000 | 0.2357140000 | -1.5380080000 |
| N | 1.8159500000 | 0.4394620000 | -1.9069840000 |

**$^{5}Im_3OHN_3$_TS-1**
Charge: 0
Multiplicity: 5
B3LYP-D3(BJ)/6-31G(d)-SDD SCF energy (au): -1778.82815305
B3LYP-D3(BJ)/6-31G(d)-SDD enthalpy (au): -1778.25406805
B3LYP-D3(BJ)/6-31G(d)-SDD free energy (au): -1778.37267705
B3LYP-D3(BJ)/def2-TZVP/SMD SCF energy (au): -2919.37921648
B3LYP-D3(BJ)/def2-TZVP/SMD enthalpy (au): -2918.80513148
B3LYP-D3(BJ)/def2-TZVP/SMD free energy (au): -2918.92374048
B3LYP-D3(BJ)/def2-TZVP/SMD free energy (quasi-harmonic) (au): -2918.91257943

Cartesian coordinates

| ATOM | X | Y | Z |
|---|---|---|---|
| Fe | 0.4401060000 | -0.1494650000 | -1.3728300000 |
| C | -0.0360300000 | 0.5930410000 | 3.0832060000 |
| N | 0.4952530000 | 1.8101770000 | 2.6917790000 |
| H | 0.8631240000 | 2.5265380000 | 3.3003400000 |
| C | 0.5405220000 | 1.8288750000 | 1.3344680000 |
| H | 0.9361520000 | 2.6523580000 | 0.7629380000 |
| N | 0.0533670000 | 0.7069250000 | 0.8344790000 |
| C | -0.3033590000 | -0.0756620000 | 1.9180060000 |
| H | -0.7028570000 | -1.0672970000 | 1.7735550000 |
| C | -3.4740300000 | -2.0852920000 | -1.8269860000 |
| N | -3.8220260000 | -1.2452900000 | -0.7818640000 |
| H | -4.7509590000 | -1.1039970000 | -0.4140830000 |
| C | -2.6889130000 | -0.6219390000 | -0.3557880000 |
| H | -2.6668310000 | 0.1093380000 | 0.4373770000 |
| N | -1.6492370000 | -1.0221010000 | -1.0614020000 |
| C | -2.1224170000 | -1.9302380000 | -1.9904280000 |
| H | -1.4391240000 | -2.3299250000 | -2.7246500000 |
| H | -4.2052500000 | -2.6864470000 | -2.3442940000 |
| H | -0.1655240000 | 0.3301130000 | 4.1214030000 |
| C | 1.9283370000 | -3.8718200000 | 0.4208060000 |
| N | 2.8005860000 | -2.9629320000 | 0.9953960000 |
| H | 3.6106770000 | -3.1838420000 | 1.5549460000 |
| C | 2.4061070000 | -1.7127450000 | 0.6335520000 |
| H | 2.9238480000 | -0.8077400000 | 0.9227610000 |
| N | 1.3309710000 | -1.7792390000 | -0.1322520000 |
| C | 1.0240350000 | -3.1161520000 | -0.2770490000 |
| H | 0.1909440000 | -3.4273880000 | -0.8869050000 |
| H | 2.0379800000 | -4.9364780000 | 0.5547560000 |
| O | 0.5256070000 | -1.2372110000 | -2.8970360000 |
| H | 0.9444710000 | -0.8048270000 | -3.6551500000 |
| C | 0.8591010000 | -0.0511500000 | -6.1798300000 |
| C | 0.6767020000 | 1.1209080000 | -5.4509490000 |
| C | -0.5346380000 | 1.3838420000 | -4.7964130000 |
| C | -1.6070960000 | 0.4696810000 | -4.8893470000 |
| C | -1.4065380000 | -0.6876570000 | -5.6491500000 |
| C | -0.1940850000 | -0.9600650000 | -6.2796960000 |
| H | 1.8105440000 | -0.2498500000 | -6.6639900000 |
| H | 1.4747520000 | 1.8508380000 | -5.3681740000 |
| H | -2.2251060000 | -1.3976290000 | -5.7368480000 |
| H | -0.0720330000 | -1.8807150000 | -6.8435110000 |
| C | -2.9697290000 | 0.7055270000 | -4.2731050000 |
| C | -3.7560600000 | 1.8262220000 | -4.9709630000 |
| H | -2.8623060000 | 0.9333250000 | -3.2122520000 |
| H | -3.5388260000 | -0.2286040000 | -4.3398830000 |
| H | -4.7458150000 | 1.9472430000 | -4.5145600000 |
| H | -3.2330310000 | 2.7840400000 | -4.8931250000 |
| H | -3.8960610000 | 1.6044660000 | -6.0350370000 |
| C | -0.5852790000 | 2.7013580000 | -4.0929250000 |
| O | -0.0931000000 | 3.7117070000 | -4.5913320000 |
| N | -1.2925160000 | 2.7686050000 | -2.8987520000 |

```
C     -1.1555260000    3.8819990000   -1.9481650000
C     -1.8440620000    5.0965680000   -2.6128320000
C      0.3088960000    4.1943680000   -1.6069800000
H     -1.8584990000    5.9232540000   -1.8933230000
H     -1.3065130000    5.4084860000   -3.5075790000
H     -2.8776400000    4.8534300000   -2.8806400000
H      0.8419450000    3.2781730000   -1.3369580000
H      0.8193450000    4.6327830000   -2.4664750000
H      0.3544650000    4.9055960000   -0.7734820000
F     -0.9091900000    1.3996200000   -2.0826340000
C     -1.9596730000    3.5053020000   -0.6926770000
H     -2.9824740000    3.2267310000   -0.9680530000
H     -1.5042450000    2.6679670000   -0.1648480000
H     -2.0033280000    4.3692110000   -0.0206030000
N      3.5475360000    1.4139860000    0.4219970000
N      2.8170820000    1.2782790000   -0.4775760000
N      2.0490820000    1.1585200000   -1.3934840000
```

**$^{5}Im_{3}OHN_{3}$_TS-3-F**

```
Charge: 0
Multiplicity: 5
B3LYP-D3(BJ)/6-31G(d)-SDD SCF energy (au):              -1778.89653373
B3LYP-D3(BJ)/6-31G(d)-SDD enthalpy (au):                -1778.32085673
B3LYP-D3(BJ)/6-31G(d)-SDD free energy (au):             -1778.43637273
B3LYP-D3(BJ)/def2-TZVP/SMD SCF energy (au):             -2919.44846472
B3LYP-D3(BJ)/def2-TZVP/SMD enthalpy (au):               -2918.87278772
B3LYP-D3(BJ)/def2-TZVP/SMD free energy (au):            -2918.98830372
B3LYP-D3(BJ)/def2-TZVP/SMD free energy (quasi-harmonic) (au): -2918.97828499

Cartesian coordinates
ATOM    X                Y                Z
H     32.7208220000   25.9665630000   37.0550260000
C     33.0719900000   26.6787660000   36.3248840000
C     34.1100480000   27.5720190000   36.3298660000
N     32.4510630000   26.8432700000   35.0982070000
C     33.1266990000   27.7945100000   34.4061840000
N     34.1279440000   28.2611800000   35.1336820000
H     31.6995360000   26.2783770000   34.7317490000
H     34.8421810000   27.7711020000   37.0975330000
H     32.9072550000   28.1326220000   33.4045790000
H     38.3878370000   29.6747510000   38.7884860000
C     38.0164030000   29.3872660000   37.8172950000
C     37.0538850000   29.9203920000   37.0006560000
N     38.5144520000   28.2913580000   37.1322840000
C     37.8515680000   28.1972170000   35.9491430000
N     36.9630490000   29.1689480000   35.8473640000
H     39.2487580000   27.6747730000   37.4469480000
H     36.4227030000   30.7834860000   37.1419450000
H     38.0126130000   27.4424240000   35.1933850000
H     32.2080820000   33.2700690000   35.5477760000
C     33.1361460000   32.7592910000   35.3440620000
C     33.5814520000   31.4974140000   35.6476030000
N     34.1708500000   33.3635810000   34.6535630000
```

| | | | |
|---|---|---|---|
| C | 35.1745470000 | 32.4604770000 | 34.5440850000 |
| N | 34.8544720000 | 31.3200060000 | 35.1384490000 |
| H | 34.1211900000 | 34.1868160000 | 34.0585120000 |
| H | 33.0791870000 | 30.6993430000 | 36.1727990000 |
| H | 36.0887710000 | 32.6348500000 | 34.0012800000 |
| Fe | 35.6740330000 | 29.4227420000 | 34.0439370000 |
| O | 34.3765540000 | 29.7764590000 | 32.6377560000 |
| H | 34.1810350000 | 30.7238650000 | 32.6130040000 |
| F | 37.0236410000 | 30.6888730000 | 33.3090850000 |
| N | 34.3522970000 | 32.4687050000 | 30.8035240000 |
| O | 34.3481000000 | 34.3862680000 | 32.0322070000 |
| C | 36.4816940000 | 33.5085800000 | 31.4161200000 |
| C | 37.0023030000 | 34.8090380000 | 31.4523580000 |
| C | 38.3660650000 | 35.0613930000 | 31.3504300000 |
| C | 39.2502760000 | 33.9846130000 | 31.2187040000 |
| C | 38.7634610000 | 32.6883890000 | 31.2137640000 |
| C | 37.3800120000 | 32.3913210000 | 31.3203590000 |
| C | 34.9753960000 | 33.4800460000 | 31.4561730000 |
| C | 32.8948950000 | 32.3407160000 | 30.5519640000 |
| C | 32.4396560000 | 33.4821150000 | 29.6282870000 |
| C | 32.7152200000 | 30.9736640000 | 29.8766640000 |
| C | 32.1006670000 | 32.3682430000 | 31.8674090000 |
| C | 36.9777660000 | 31.0064420000 | 31.4293280000 |
| C | 37.8449390000 | 29.8608570000 | 31.0145460000 |
| H | 36.2965620000 | 35.6261590000 | 31.5501970000 |
| H | 38.7347660000 | 36.0825690000 | 31.3670710000 |
| H | 40.3192150000 | 34.1592450000 | 31.1335000000 |
| H | 39.4658670000 | 31.8665880000 | 31.1476270000 |
| H | 34.9374940000 | 31.8734310000 | 30.2346130000 |
| H | 35.9234800000 | 30.7619870000 | 31.4450180000 |
| H | 38.8634450000 | 29.9363180000 | 31.4041500000 |
| H | 37.4102270000 | 28.9296870000 | 31.3854780000 |
| H | 37.9053470000 | 29.8135460000 | 29.9165570000 |
| H | 31.3699500000 | 33.3890580000 | 29.4098660000 |
| H | 32.6173590000 | 34.4494170000 | 30.1053540000 |
| H | 32.9868200000 | 33.4523590000 | 28.6792650000 |
| H | 33.2543440000 | 30.9328180000 | 28.9210140000 |
| H | 33.0887800000 | 30.1788480000 | 30.5317150000 |
| H | 31.6556640000 | 30.7969880000 | 29.6670400000 |
| H | 32.2339470000 | 33.3195620000 | 32.3815990000 |
| H | 31.0366440000 | 32.2343470000 | 31.6423860000 |
| H | 32.4090050000 | 31.5556680000 | 32.5310410000 |
| N | 36.5048950000 | 27.6561830000 | 33.3388990000 |
| N | 35.7527700000 | 26.7528370000 | 33.0543360000 |
| N | 35.0461950000 | 25.8735460000 | 32.7861810000 |

**$^{5}Im_3OHN_3$_TS-3-$N_3$**

Charge: 0

Multiplicity: 5

B3LYP-D3(BJ)/6-31G(d)-SDD SCF energy (au): -1778.89107324

B3LYP-D3(BJ)/6-31G(d)-SDD enthalpy (au): -1778.31647324

B3LYP-D3(BJ)/6-31G(d)-SDD free energy (au): -1778.43443124

B3LYP-D3(BJ)/def2-TZVP/SMD SCF energy (au): -2919.45410297

B3LYP-D3(BJ)/def2-TZVP/SMD enthalpy (au): -2918.87950297
B3LYP-D3(BJ)/def2-TZVP/SMD free energy (au): -2918.99746097
B3LYP-D3(BJ)/def2-TZVP/SMD free energy (quasi-harmonic) (au): -2918.98560073

Cartesian coordinates

| ATOM | X | Y | Z |
|---|---|---|---|
| H | 31.2872020000 | 27.7496970000 | 37.0589890000 |
| C | 32.0662240000 | 28.4702530000 | 36.8648260000 |
| C | 33.3193780000 | 28.3544570000 | 36.3222270000 |
| N | 31.9169080000 | 29.8168030000 | 37.1523270000 |
| C | 33.0542830000 | 30.4571710000 | 36.7706010000 |
| N | 33.9263220000 | 29.5965120000 | 36.2725250000 |
| H | 31.0851960000 | 30.2634440000 | 37.5083580000 |
| H | 33.8140620000 | 27.4731900000 | 35.9447770000 |
| H | 33.2183630000 | 31.5238610000 | 36.8017480000 |
| H | 37.5410310000 | 25.5787100000 | 36.2178320000 |
| C | 37.2708790000 | 26.4945620000 | 35.7157560000 |
| C | 36.5402920000 | 27.5865460000 | 36.1075410000 |
| N | 37.6683190000 | 26.7685980000 | 34.4181780000 |
| C | 37.1737150000 | 27.9897700000 | 34.0785000000 |
| N | 36.4844540000 | 28.5016420000 | 35.0786790000 |
| H | 38.2140000000 | 26.1644360000 | 33.8221680000 |
| H | 36.0566740000 | 27.7849470000 | 37.0514620000 |
| H | 37.3445170000 | 28.4835130000 | 33.1351780000 |
| H | 38.1401750000 | 31.7216750000 | 39.5874320000 |
| C | 37.8890920000 | 31.5300100000 | 38.5559220000 |
| C | 36.7433340000 | 31.0873280000 | 37.9479070000 |
| N | 38.7966580000 | 31.7280050000 | 37.5278220000 |
| C | 38.1817350000 | 31.4081100000 | 36.3574130000 |
| N | 36.9453300000 | 31.0152630000 | 36.5888680000 |
| H | 39.7390540000 | 32.0746750000 | 37.6248500000 |
| H | 35.7901060000 | 30.8239560000 | 38.3802480000 |
| H | 38.6178740000 | 31.4703440000 | 35.3717720000 |
| Fe | 35.5984500000 | 30.6490670000 | 34.8735340000 |
| O | 34.7084890000 | 32.2504930000 | 35.2961610000 |
| H | 33.8545550000 | 32.3445050000 | 34.8401450000 |
| F | 37.1060380000 | 30.8718000000 | 33.7179920000 |
| N | 31.5029940000 | 31.1298980000 | 32.0471140000 |
| O | 31.9049100000 | 32.1006570000 | 34.0737540000 |
| C | 33.3579700000 | 32.6952810000 | 32.2891900000 |
| C | 33.3977390000 | 34.0592810000 | 32.5831150000 |
| C | 34.4806400000 | 34.8485390000 | 32.2052180000 |
| C | 35.5721600000 | 34.2494920000 | 31.5710380000 |
| C | 35.5632470000 | 32.8877540000 | 31.3048310000 |
| C | 34.4513470000 | 32.0747120000 | 31.6194110000 |
| C | 32.2024090000 | 31.9419200000 | 32.8833750000 |
| C | 30.3415580000 | 30.2972350000 | 32.4252850000 |
| C | 30.7388110000 | 29.3171050000 | 33.5408040000 |
| C | 29.1825480000 | 31.1997940000 | 32.8822250000 |
| C | 29.9424740000 | 29.5175960000 | 31.1663570000 |
| C | 34.4999630000 | 30.6522720000 | 31.3648000000 |
| C | 35.7472610000 | 29.9887490000 | 30.8637520000 |
| H | 32.5722840000 | 34.4905890000 | 33.1404340000 |

H     34.4877710000   35.9103490000   32.4323430000
H     36.4398780000   34.8439020000   31.2987460000
H     36.4317890000   32.4339860000   30.8429110000
H     31.7236190000   31.1954450000   31.0636690000
H     33.5684140000   30.1536420000   31.1261250000
H     36.5943800000   30.2745610000   31.4942060000
H     35.6316930000   28.8998490000   30.8930660000
H     35.9657960000   30.2648420000   29.8204510000
H     29.8631000000   28.7295700000   33.8412710000
H     31.5133810000   28.6250980000   33.1992580000
H     31.1161160000   29.8576480000   34.4090440000
H     29.4852360000   31.7827750000   33.7545380000
H     28.8935950000   31.8910050000   32.0822660000
H     28.3088200000   30.5921060000   33.1445170000
H     30.7651340000   28.8778420000   30.8261200000
H     29.0815790000   28.8764620000   31.3789370000
H     29.6626440000   30.1959450000   30.3501830000
N     34.4138760000   29.9011240000   33.2973090000
N     34.1631940000   28.7221150000   33.1710680000
N     33.9112620000   27.6050440000   33.0028660000

**$^{5}Im_3OHN_3$_TS-3-OH**
Charge: 0
Multiplicity: 5
B3LYP-D3(BJ)/6-31G(d)-SDD SCF energy (au):               -1778.89402979
B3LYP-D3(BJ)/6-31G(d)-SDD enthalpy (au):                 -1778.31851379
B3LYP-D3(BJ)/6-31G(d)-SDD free energy (au):              -1778.43335679
B3LYP-D3(BJ)/def2-TZVP/SMD SCF energy (au):              -2919.45706218
B3LYP-D3(BJ)/def2-TZVP/SMD enthalpy (au):                -2918.88154618
B3LYP-D3(BJ)/def2-TZVP/SMD free energy (au):             -2918.99638918
B3LYP-D3(BJ)/def2-TZVP/SMD free energy (quasi-harmonic) (au): -2918.98622939

Cartesian coordinates
ATOM    X               Y               Z
H     -0.3883420000   -3.1885880000   -3.6500190000
C     -0.2069060000   -2.3467770000   -3.0008040000
C      0.6856820000   -2.1446960000   -1.9812760000
N     -0.9289360000   -1.1721610000   -3.0978730000
C     -0.4854410000   -0.3213350000   -2.1422090000
N      0.5016660000   -0.8789610000   -1.4606230000
H     -1.7715430000   -1.0334920000   -3.6366850000
H      1.4346320000   -2.8088410000   -1.5803040000
H     -0.8722270000    0.6683070000   -1.9575090000
H      5.4072800000    1.3189480000   -2.7297680000
C      4.7341970000    1.1359120000   -1.9072180000
C      3.5672450000    0.4269840000   -1.8127500000
N      4.9581240000    1.6532570000   -0.6426620000
C      3.9343790000    1.2748900000    0.1591480000
N      3.0897780000    0.5174350000   -0.5205610000
H      5.6845550000    2.3064330000   -0.3894640000
H      3.0289700000   -0.1247320000   -2.5676700000
H      3.7976080000    1.5460820000    1.1945260000
H      4.5468630000   -4.2570970000    1.3493850000

| | | | |
|---|---|---|---|
| C | 3.7082870000 | -3.5806910000 | 1.2941920000 |
| C | 3.4203850000 | -2.5269960000 | 0.4655700000 |
| N | 2.6368560000 | -3.6631970000 | 2.1659970000 |
| C | 1.7574650000 | -2.6769810000 | 1.8411920000 |
| N | 2.2031560000 | -1.9804920000 | 0.8153860000 |
| H | 2.5375230000 | -4.3192990000 | 2.9266050000 |
| H | 4.0019330000 | -2.1151640000 | -0.3453920000 |
| H | 0.8339200000 | -2.4855270000 | 2.3626490000 |
| Fe | 1.2322700000 | 0.0743740000 | 0.3048650000 |
| O | -0.3909060000 | -0.3929070000 | 1.1710690000 |
| H | -0.2182740000 | -0.0179280000 | 2.0577660000 |
| F | 1.8443200000 | 0.6531580000 | 1.9887740000 |
| N | -2.4987240000 | 1.6184410000 | 0.8600610000 |
| O | -4.5100030000 | 1.9988390000 | -0.1745070000 |
| C | -3.4923920000 | -0.1125170000 | -0.5464820000 |
| C | -4.1148130000 | -0.2110480000 | -1.7934980000 |
| C | -4.1730130000 | -1.4167130000 | -2.4929400000 |
| C | -3.5913010000 | -2.5604620000 | -1.9313470000 |
| C | -2.9815290000 | -2.4895670000 | -0.6885490000 |
| C | -2.9338350000 | -1.2827270000 | 0.0589120000 |
| C | -3.5454000000 | 1.2683900000 | 0.0747870000 |
| C | -2.3415140000 | 2.9430780000 | 1.5008890000 |
| C | -3.5609780000 | 3.2440980000 | 2.3871830000 |
| C | -1.0825960000 | 2.8563920000 | 2.3747640000 |
| C | -2.1696750000 | 4.0259160000 | 0.4223280000 |
| C | -2.4142610000 | -1.2945360000 | 1.3948910000 |
| C | -1.9786580000 | -2.5494720000 | 2.0802320000 |
| H | -4.5765090000 | 0.6873090000 | -2.1891000000 |
| H | -4.6808420000 | -1.4685930000 | -3.4525630000 |
| H | -3.6246510000 | -3.5094490000 | -2.4604310000 |
| H | -2.5569430000 | -3.3901300000 | -0.2598630000 |
| H | -1.6589960000 | 1.0376980000 | 0.8137310000 |
| H | -2.6703460000 | -0.4533560000 | 2.0248290000 |
| H | -1.5224300000 | -2.3175400000 | 3.0472420000 |
| H | -2.8314370000 | -3.2199520000 | 2.2706440000 |
| H | -1.2498470000 | -3.1054090000 | 1.4816140000 |
| H | -3.4129210000 | 4.2008190000 | 2.9009290000 |
| H | -4.4728080000 | 3.2960580000 | 1.7910840000 |
| H | -3.6817710000 | 2.4631920000 | 3.1475320000 |
| H | -0.9082040000 | 3.8177950000 | 2.8687850000 |
| H | -1.2043190000 | 2.0953860000 | 3.1564790000 |
| H | -0.2004350000 | 2.6068520000 | 1.7800780000 |
| H | -2.0127580000 | 5.0053200000 | 0.8890930000 |
| H | -1.3037640000 | 3.7911720000 | -0.2025690000 |
| H | -3.0628400000 | 4.0748960000 | -0.2055600000 |
| N | 0.5510440000 | 1.8948300000 | -0.5011820000 |
| N | 1.3491680000 | 2.7973950000 | -0.5882340000 |
| N | 2.1148550000 | 3.6647160000 | -0.6767520000 |

**$^{5}Im_{3}Wat_{2}$_FeF_remove-F**
Charge: 2
Multiplicity: 5
B3LYP-D3(BJ)/6-31G(d)-SDD SCF energy (au): -955.134744862

B3LYP-D3(BJ)/6-31G(d)-SDD enthalpy (au): -954.841582862
B3LYP-D3(BJ)/6-31G(d)-SDD free energy (au): -954.921788862
B3LYP-D3(BJ)/def2-TZVP/SMD SCF energy (au): -2095.53168490
B3LYP-D3(BJ)/def2-TZVP/SMD enthalpy (au): -2095.23852290
B3LYP-D3(BJ)/def2-TZVP/SMD free energy (au): -2095.31872890
B3LYP-D3(BJ)/def2-TZVP/SMD free energy (quasi-harmonic) (au): -2095.310899038

Cartesian coordinates

| ATOM | X | Y | Z |
|---|---|---|---|
| Fe | 0.1290200000 | -0.2784260000 | -0.6121820000 |
| C | 1.0039420000 | 1.6378360000 | 3.0852060000 |
| N | 2.0058800000 | 2.1637540000 | 2.2935170000 |
| H | 2.7107960000 | 2.8267370000 | 2.5946750000 |
| C | 1.8856260000 | 1.6597000000 | 1.0506370000 |
| H | 2.5483160000 | 1.9018680000 | 0.2350590000 |
| N | 0.8441220000 | 0.8281140000 | 0.9955520000 |
| C | 0.2863690000 | 0.8081440000 | 2.2700160000 |
| H | -0.5787680000 | 0.2059870000 | 2.4981490000 |
| C | -3.6470800000 | -1.8027650000 | 0.7800250000 |
| N | -3.8706050000 | -0.4452620000 | 0.8972140000 |
| H | -4.7225720000 | -0.0135020000 | 1.2357620000 |
| C | -2.7682110000 | 0.2122820000 | 0.4826840000 |
| H | -2.6740420000 | 1.2883820000 | 0.4762470000 |
| N | -1.8342380000 | -0.6609520000 | 0.1008190000 |
| C | -2.3783170000 | -1.9260770000 | 0.2843040000 |
| H | -1.8203570000 | -2.8191290000 | 0.0490930000 |
| O | 1.8697030000 | 0.1616840000 | -1.9464040000 |
| H | 2.5376480000 | -0.5384620000 | -2.0380240000 |
| H | 2.2494930000 | 0.9712640000 | -2.3268220000 |
| H | -4.3914530000 | -2.5348910000 | 1.0523800000 |
| H | 0.8963850000 | 1.8974600000 | 4.1269830000 |
| C | 1.4541840000 | -4.3636370000 | -0.7621400000 |
| N | 1.8810510000 | -3.9876270000 | 0.4940010000 |
| H | 2.3530360000 | -4.5862780000 | 1.1613230000 |
| C | 1.5713020000 | -2.6875560000 | 0.6862870000 |
| H | 1.7906580000 | -2.1388450000 | 1.5899930000 |
| N | 0.9621600000 | -2.2017290000 | -0.3938960000 |
| C | 0.8833610000 | -3.2457120000 | -1.3077140000 |
| H | 0.4249510000 | -3.1226640000 | -2.2775460000 |
| H | 1.5927000000 | -5.3629880000 | -1.1449390000 |
| O | -0.8444530000 | 0.1645010000 | -2.4456590000 |
| H | -0.4015110000 | 0.2139660000 | -3.3095160000 |
| H | -1.7861840000 | -0.0295830000 | -2.5889860000 |

**$^{6}$Im$_3$Wat$_2$_FeF**
Charge: 2
Multiplicity: 6
B3LYP-D3(BJ)/6-31G(d)-SDD SCF energy (au): -1054.98309009
B3LYP-D3(BJ)/6-31G(d)-SDD enthalpy (au): -1054.68541309
B3LYP-D3(BJ)/6-31G(d)-SDD free energy (au): -1054.76279009
B3LYP-D3(BJ)/def2-TZVP/SMD SCF energy (au): -2195.42360882
B3LYP-D3(BJ)/def2-TZVP/SMD enthalpy (au): -2195.12593182
B3LYP-D3(BJ)/def2-TZVP/SMD free energy (au): -2195.20330882

B3LYP-D3(BJ)/def2-TZVP/SMD free energy (quasi-harmonic) (au): -2195.19898573

Cartesian coordinates

| ATOM | X | Y | Z |
|---|---|---|---|
| Fe | 0.0964840000 | -0.0998000000 | -0.7458960000 |
| C | 1.2103150000 | 0.9276020000 | 3.2585740000 |
| N | 1.4917950000 | 2.0837160000 | 2.5543800000 |
| H | 1.8152070000 | 2.9569540000 | 2.9554150000 |
| C | 1.2541210000 | 1.8686300000 | 1.2486300000 |
| H | 1.3550300000 | 2.6132690000 | 0.4740400000 |
| N | 0.8414700000 | 0.6101820000 | 1.0714020000 |
| C | 0.8017620000 | 0.0157980000 | 2.3267560000 |
| H | 0.4656830000 | -0.9994640000 | 2.4656430000 |
| C | -3.6404820000 | -1.2953720000 | 0.9285800000 |
| N | -3.8413150000 | 0.0372660000 | 0.6173760000 |
| H | -4.6932940000 | 0.5611010000 | 0.7837580000 |
| C | -2.7264430000 | 0.5278420000 | 0.0497420000 |
| H | -2.5965450000 | 1.5434950000 | -0.2929720000 |
| N | -1.8051720000 | -0.4394580000 | -0.0217430000 |
| C | -2.3668630000 | -1.5842540000 | 0.5276370000 |
| H | -1.8237460000 | -2.5139880000 | 0.5845900000 |
| O | 1.8772600000 | 0.1242620000 | -2.0382480000 |
| H | 2.7267780000 | -0.3399820000 | -1.9507540000 |
| H | 2.0588020000 | 1.0388710000 | -2.3198080000 |
| H | -4.4044270000 | -1.9008070000 | 1.3914690000 |
| H | 1.3188670000 | 0.8640190000 | 4.3303400000 |
| C | 1.1694930000 | -4.2624490000 | -0.5687450000 |
| N | 2.1127880000 | -3.7181120000 | 0.2807390000 |
| H | 2.8370840000 | -4.2340980000 | 0.7675100000 |
| C | 1.9145170000 | -2.3886570000 | 0.3567560000 |
| H | 2.5085790000 | -1.7194990000 | 0.9600170000 |
| N | 0.8762790000 | -2.0427860000 | -0.4104070000 |
| C | 0.4057960000 | -3.2132300000 | -0.9977820000 |
| H | -0.4223800000 | -3.2033510000 | -1.6874660000 |
| H | 1.1312080000 | -5.3183670000 | -0.7883580000 |
| O | -0.6471930000 | -0.9212450000 | -2.6374660000 |
| H | 0.0109180000 | -0.7626070000 | -3.3392530000 |
| H | -1.4706940000 | -0.4924740000 | -2.9325990000 |
| F | -0.2212120000 | 1.5735220000 | -1.3398050000 |

**$^{5}Im_{3}WatN_{3}$_11**

Charge: 1

Multiplicity: 5

B3LYP-D3(BJ)/6-31G(d)-SDD SCF energy (au): -1779.29954894

B3LYP-D3(BJ)/6-31G(d)-SDD enthalpy (au): -1778.70854694

B3LYP-D3(BJ)/6-31G(d)-SDD free energy (au): -1778.82893794

B3LYP-D3(BJ)/def2-TZVP/SMD SCF energy (au): -2919.88077577

B3LYP-D3(BJ)/def2-TZVP/SMD enthalpy (au): -2919.28977377

B3LYP-D3(BJ)/def2-TZVP/SMD free energy (au): -2919.41016477

B3LYP-D3(BJ)/def2-TZVP/SMD free energy (quasi-harmonic) (au): -2919.39838683

Cartesian coordinates

| ATOM | X | Y | Z |
|---|---|---|---|

| | | | |
|---|---|---|---|
| Fe | 0.0460890000 | -1.1547180000 | -0.8650850000 |
| C | 1.6100680000 | 2.1758070000 | 1.5047020000 |
| N | 2.0626770000 | 2.4800640000 | 0.2341950000 |
| H | 2.7129120000 | 3.2154240000 | -0.0052420000 |
| C | 1.5369100000 | 1.5800600000 | -0.6319580000 |
| H | 1.7287090000 | 1.5684650000 | -1.6925870000 |
| N | 0.7668350000 | 0.7173350000 | 0.0156160000 |
| C | 0.8051920000 | 1.0780500000 | 1.3512820000 |
| H | 0.2476970000 | 0.5292630000 | 2.0945230000 |
| C | -2.9869900000 | -1.6872550000 | 2.2410430000 |
| N | -3.7787890000 | -1.3352490000 | 1.1635930000 |
| H | -4.7861650000 | -1.2565890000 | 1.1656430000 |
| C | -2.9751450000 | -1.1300150000 | 0.0928760000 |
| H | -3.3216950000 | -0.8284560000 | -0.8818580000 |
| N | -1.7082470000 | -1.3283580000 | 0.4282980000 |
| C | -1.7029170000 | -1.6802060000 | 1.7673230000 |
| H | -0.7825420000 | -1.9196800000 | 2.2752080000 |
| H | -3.4010080000 | -1.9102060000 | 3.2116620000 |
| H | 1.8955720000 | 2.7521080000 | 2.3706500000 |
| C | 2.0582560000 | -4.2759750000 | 1.3745220000 |
| N | 2.9292840000 | -3.2187860000 | 1.5651860000 |
| H | 3.8553150000 | -3.2755630000 | 1.9652410000 |
| C | 2.3538970000 | -2.0984350000 | 1.0657760000 |
| H | 2.8329900000 | -1.1325990000 | 1.0474920000 |
| N | 1.1572350000 | -2.3808810000 | 0.5687500000 |
| C | 0.9625860000 | -3.7398490000 | 0.7544600000 |
| H | 0.0582300000 | -4.2222750000 | 0.4166760000 |
| H | 2.2962760000 | -5.2801460000 | 1.6879080000 |
| O | -0.5648790000 | -2.9660690000 | -2.0926380000 |
| H | 0.3239520000 | -2.9294970000 | -2.5066140000 |
| H | -1.1533170000 | -2.5698000000 | -2.7572880000 |
| C | -5.3873280000 | 1.8088860000 | -3.1552480000 |
| C | -4.0840960000 | 1.3719190000 | -3.3807690000 |
| C | -3.0977710000 | 1.5642510000 | -2.4059780000 |
| C | -3.4050410000 | 2.1502990000 | -1.1637590000 |
| C | -4.7206520000 | 2.5888890000 | -0.9660710000 |
| C | -5.7004100000 | 2.4324040000 | -1.9460130000 |
| H | -6.1486720000 | 1.6646460000 | -3.9150730000 |
| H | -3.8258340000 | 0.8860530000 | -4.3176960000 |
| H | -4.9777460000 | 3.0560490000 | -0.0188400000 |
| H | -6.7105610000 | 2.7849180000 | -1.7610220000 |
| C | -2.3793820000 | 2.3038810000 | -0.0603420000 |
| C | -1.5146070000 | 3.5696220000 | -0.1662610000 |
| H | -1.7226880000 | 1.4313310000 | -0.0375730000 |
| H | -2.9098110000 | 2.3166450000 | 0.8989700000 |
| H | -0.8608220000 | 3.6491020000 | 0.7078540000 |
| H | -0.8780760000 | 3.5348610000 | -1.0547610000 |
| H | -2.1314990000 | 4.4730090000 | -0.2190210000 |
| C | -1.7357490000 | 0.9973300000 | -2.6554030000 |
| O | -1.4607450000 | -0.1465540000 | -2.2572250000 |
| N | -0.8139410000 | 1.7830630000 | -3.2672500000 |
| C | -0.8441150000 | 2.9328110000 | -4.2351930000 |
| C | -1.9806020000 | 3.9139580000 | -3.9170450000 |

| | | | |
|---|---|---|---|
| C | -0.9965240000 | 2.3589890000 | -5.6511850000 |
| H | -1.8280590000 | 4.7915540000 | -4.5523190000 |
| H | -2.9664850000 | 3.5087960000 | -4.1381220000 |
| H | -1.9601030000 | 4.2432540000 | -2.8767780000 |
| H | -0.2025970000 | 1.6409630000 | -5.8705970000 |
| H | -1.9650150000 | 1.8614390000 | -5.7651720000 |
| H | -0.9418570000 | 3.1695210000 | -6.3840580000 |
| F | 0.3622810000 | 1.0532410000 | -3.4938080000 |
| C | 0.4994540000 | 3.6654450000 | -4.0735960000 |
| H | 0.6226640000 | 4.0211810000 | -3.0451610000 |
| H | 1.3445860000 | 3.0261410000 | -4.3330720000 |
| H | 0.5103160000 | 4.5331430000 | -4.7384330000 |
| N | 3.7715500000 | -0.7022220000 | -1.5002780000 |
| N | 2.7094040000 | -0.9985550000 | -1.8554670000 |
| N | 1.5921920000 | -1.2845650000 | -2.2189700000 |

**$^{5}Im_{3}WatN_{3}$_13**

Charge: 1
Multiplicity: 5
B3LYP-D3(BJ)/6-31G(d)-SDD SCF energy (au): -1779.36491787
B3LYP-D3(BJ)/6-31G(d)-SDD enthalpy (au): -1778.77521587
B3LYP-D3(BJ)/6-31G(d)-SDD free energy (au): -1778.89570387
B3LYP-D3(BJ)/def2-TZVP/SMD SCF energy (au): -2919.95371781
B3LYP-D3(BJ)/def2-TZVP/SMD enthalpy (au): -2919.36401581
B3LYP-D3(BJ)/def2-TZVP/SMD free energy (au): -2919.48450381
B3LYP-D3(BJ)/def2-TZVP/SMD free energy (quasi-harmonic) (au): -2919.47132394

Cartesian coordinates

| ATOM | X | Y | Z |
|---|---|---|---|
| H | 33.1276900000 | 26.2897170000 | 37.3348880000 |
| C | 33.3807290000 | 27.2461090000 | 36.9056760000 |
| C | 34.5820460000 | 27.8430770000 | 36.6370280000 |
| N | 32.4198240000 | 28.1511260000 | 36.4988330000 |
| C | 33.0411730000 | 29.2451220000 | 36.0014270000 |
| N | 34.3574740000 | 29.0909330000 | 36.0785400000 |
| H | 31.4195510000 | 28.0100980000 | 36.5242260000 |
| H | 35.5810740000 | 27.4707830000 | 36.7969860000 |
| H | 32.5227880000 | 30.0892090000 | 35.5716530000 |
| H | 39.8346410000 | 26.9499830000 | 36.4325760000 |
| C | 39.0639100000 | 27.5121750000 | 35.9294520000 |
| C | 38.1370380000 | 28.4105220000 | 36.3836850000 |
| N | 38.8501890000 | 27.4155610000 | 34.5668930000 |
| C | 37.8241290000 | 28.2324060000 | 34.2361340000 |
| N | 37.3684440000 | 28.8463600000 | 35.3189270000 |
| H | 39.3755590000 | 26.8413660000 | 33.9220210000 |
| H | 37.9666850000 | 28.7739890000 | 37.3844070000 |
| H | 37.4418400000 | 28.3749520000 | 33.2373420000 |
| H | 36.4614910000 | 31.2095160000 | 40.5411610000 |
| C | 36.5581150000 | 31.2166640000 | 39.4669830000 |
| C | 35.9039430000 | 30.5293670000 | 38.4802370000 |
| N | 37.4663310000 | 32.0293010000 | 38.8127350000 |
| C | 37.3439150000 | 31.8209880000 | 37.4811580000 |
| N | 36.4056080000 | 30.9127030000 | 37.2508120000 |

| | | | |
|---|---|---|---|
| H | 38.1027080000 | 32.6815410000 | 39.2496960000 |
| H | 35.1142800000 | 29.7999400000 | 38.5615930000 |
| H | 37.9162140000 | 32.3194620000 | 36.7145320000 |
| Fe | 35.8777290000 | 30.4170610000 | 35.1854270000 |
| O | 34.4720730000 | 32.0148840000 | 35.3363580000 |
| H | 34.9522030000 | 32.6998460000 | 34.8363670000 |
| H | 33.6013080000 | 31.8940090000 | 34.8463880000 |
| F | 37.0734060000 | 31.6928840000 | 34.6502500000 |
| N | 31.3040420000 | 31.3068090000 | 31.9964170000 |
| O | 32.1893290000 | 31.5809800000 | 34.0775460000 |
| C | 33.3840360000 | 32.5210940000 | 32.2365680000 |
| C | 33.6670380000 | 33.7492890000 | 32.8379840000 |
| C | 34.7098640000 | 34.5562750000 | 32.3811070000 |
| C | 35.4912170000 | 34.1168950000 | 31.3026000000 |
| C | 35.2293940000 | 32.9006620000 | 30.6968720000 |
| C | 34.1746570000 | 32.0474020000 | 31.1367610000 |
| C | 32.2529560000 | 31.7485140000 | 32.8351670000 |
| C | 30.0409100000 | 30.6062940000 | 32.3483190000 |
| C | 30.3466310000 | 29.2915490000 | 33.0806650000 |
| C | 29.1692240000 | 31.5296620000 | 33.2137140000 |
| C | 29.3440280000 | 30.3079070000 | 31.0146690000 |
| C | 33.9807190000 | 30.7771160000 | 30.5438240000 |
| C | 34.8240970000 | 30.2340380000 | 29.4355960000 |
| H | 33.0425930000 | 34.0828660000 | 33.6622300000 |
| H | 34.9023230000 | 35.5167430000 | 32.8480880000 |
| H | 36.3078650000 | 34.7331030000 | 30.9388190000 |
| H | 35.8563980000 | 32.5718950000 | 29.8759800000 |
| H | 31.4260880000 | 31.5580090000 | 31.0238770000 |
| H | 33.2409800000 | 30.1115760000 | 30.9723940000 |
| H | 35.8571610000 | 30.0542420000 | 29.7713400000 |
| H | 34.4258260000 | 29.2827250000 | 29.0738260000 |
| H | 34.8868900000 | 30.9201190000 | 28.5806370000 |
| H | 29.4125080000 | 28.7442430000 | 33.2459860000 |
| H | 31.0247020000 | 28.6627110000 | 32.4979710000 |
| H | 30.8128260000 | 29.4864890000 | 34.0476220000 |
| H | 29.6831780000 | 31.7857880000 | 34.1430300000 |
| H | 28.9341580000 | 32.4547320000 | 32.6771190000 |
| H | 28.2273530000 | 31.0286030000 | 33.4604890000 |
| H | 29.9584650000 | 29.6531410000 | 30.3861160000 |
| H | 28.3908010000 | 29.8041620000 | 31.1964100000 |
| H | 29.1335900000 | 31.2314340000 | 30.4623200000 |
| N | 35.3071900000 | 29.7654520000 | 33.4090700000 |
| N | 34.4424400000 | 28.9544320000 | 33.1667140000 |
| N | 33.6180570000 | 28.1934950000 | 32.8939020000 |

**$^{5}Im_3WatN_3$_FeF_remove-F**
Charge: 1
Multiplicity: 5

| | |
|---|---|
| B3LYP-D3(BJ)/6-31G(d)-SDD SCF energy (au): | -1043.21464517 |
| B3LYP-D3(BJ)/6-31G(d)-SDD enthalpy (au): | -1042.93275317 |
| B3LYP-D3(BJ)/6-31G(d)-SDD free energy (au): | -1043.01460517 |
| B3LYP-D3(BJ)/def2-TZVP/SMD SCF energy (au): | -2183.51101043 |
| B3LYP-D3(BJ)/def2-TZVP/SMD enthalpy (au): | -2183.22911843 |

B3LYP-D3(BJ)/def2-TZVP/SMD free energy (au): -2183.31097043
B3LYP-D3(BJ)/def2-TZVP/SMD free energy (quasi-harmonic) (au): -2183.30341626

Cartesian coordinates

| ATOM | X | Y | Z |
|---|---|---|---|
| Fe | 0.3578050000 | -0.7767130000 | -0.5548850000 |
| C | 2.1106280000 | 0.9118400000 | 3.0684070000 |
| N | 2.0077290000 | 2.0259530000 | 2.2578220000 |
| H | 2.3768470000 | 2.9452750000 | 2.4602430000 |
| C | 1.3721970000 | 1.6674210000 | 1.1184430000 |
| H | 1.1777530000 | 2.3394440000 | 0.2963420000 |
| N | 1.0553000000 | 0.3791280000 | 1.1563230000 |
| C | 1.5156510000 | -0.1044660000 | 2.3704860000 |
| H | 1.3923280000 | -1.1414820000 | 2.6434750000 |
| C | -3.4255840000 | -2.3238330000 | 0.7826200000 |
| N | -3.1887310000 | -1.3356610000 | 1.7196040000 |
| H | -3.7930280000 | -1.0965000000 | 2.4943000000 |
| C | -2.0157670000 | -0.7343190000 | 1.4212210000 |
| H | -1.5790400000 | 0.0709930000 | 1.9911600000 |
| N | -1.4867430000 | -1.2856760000 | 0.3349130000 |
| C | -2.3594970000 | -2.2832130000 | -0.0727770000 |
| H | -2.1503670000 | -2.8880300000 | -0.9412870000 |
| H | -4.3062930000 | -2.9456760000 | 0.8120270000 |
| H | 2.5848880000 | 0.9466360000 | 4.0365530000 |
| C | 2.7274450000 | -4.3783200000 | -0.2426410000 |
| N | 3.6697090000 | -3.3735180000 | -0.1332090000 |
| H | 4.6711100000 | -3.5015100000 | -0.0796750000 |
| C | 3.0285690000 | -2.1820220000 | -0.1341560000 |
| H | 3.5267790000 | -1.2243270000 | -0.0865110000 |
| N | 1.7180780000 | -2.3732970000 | -0.2411510000 |
| C | 1.5185910000 | -3.7408420000 | -0.3122580000 |
| H | 0.5293580000 | -4.1599830000 | -0.4117180000 |
| H | 2.9981450000 | -5.4220640000 | -0.2614040000 |
| O | -0.4236710000 | -1.7886900000 | -2.4758550000 |
| H | 0.3507990000 | -1.4411450000 | -2.9611020000 |
| H | -1.1704340000 | -1.2838230000 | -2.8404400000 |
| N | 3.4565680000 | 1.1528430000 | -1.4469140000 |
| N | 2.3943950000 | 0.7626210000 | -1.6744640000 |
| N | 1.2715220000 | 0.3604960000 | -1.9111140000 |

**$^5Im_3WatN_3$_FeF_remove-$N_3$**
Charge: 1
Multiplicity: 5
B3LYP-D3(BJ)/6-31G(d)-SDD SCF energy (au): -978.881546340
B3LYP-D3(BJ)/6-31G(d)-SDD enthalpy (au): -978.613679340
B3LYP-D3(BJ)/6-31G(d)-SDD free energy (au): -978.687327340
B3LYP-D3(BJ)/def2-TZVP/SMD SCF energy (au): -2119.15587470
B3LYP-D3(BJ)/def2-TZVP/SMD enthalpy (au): -2118.88800770
B3LYP-D3(BJ)/def2-TZVP/SMD free energy (au): -2118.96165570
B3LYP-D3(BJ)/def2-TZVP/SMD free energy (quasi-harmonic) (au): -2118.956150360

Cartesian coordinates

ATOM X Y Z

| | | | |
|---|---|---|---|
| Fe | 0.2353100000 | -0.5913820000 | -0.4129880000 |
| C | 2.0425790000 | 1.1506760000 | 3.1944440000 |
| N | 1.9141050000 | 2.2172810000 | 2.3264200000 |
| H | 2.1626880000 | 3.1769550000 | 2.5238030000 |
| C | 1.3868370000 | 1.7620290000 | 1.1641320000 |
| H | 1.1545110000 | 2.3729960000 | 0.3048940000 |
| N | 1.1738040000 | 0.4555110000 | 1.2424890000 |
| C | 1.5758760000 | 0.0620750000 | 2.5062710000 |
| H | 1.4867330000 | -0.9631360000 | 2.8333140000 |
| C | -2.9961530000 | -2.7628090000 | 1.4671040000 |
| N | -3.4688760000 | -1.4657470000 | 1.4434050000 |
| H | -4.3796360000 | -1.1598920000 | 1.7574110000 |
| C | -2.5082050000 | -0.6708700000 | 0.9152660000 |
| H | -2.6103120000 | 0.3939460000 | 0.7679360000 |
| N | -1.4431180000 | -1.3944090000 | 0.5996150000 |
| C | -1.7352920000 | -2.7030320000 | 0.9376240000 |
| H | -1.0248750000 | -3.4990590000 | 0.7753530000 |
| H | -3.5843350000 | -3.5832810000 | 1.8463630000 |
| H | 2.4366680000 | 1.2636610000 | 4.1921090000 |
| C | 2.4713140000 | -4.2244680000 | -0.9268760000 |
| N | 3.3566430000 | -3.4780380000 | -0.1725430000 |
| H | 4.2770970000 | -3.7729720000 | 0.1246110000 |
| C | 2.7900390000 | -2.2783250000 | 0.0901300000 |
| H | 3.2588750000 | -1.4945110000 | 0.6652860000 |
| N | 1.5838840000 | -2.2163050000 | -0.4600040000 |
| C | 1.3726210000 | -3.4278560000 | -1.0995800000 |
| H | 0.4544110000 | -3.6267670000 | -1.6309290000 |
| H | 2.7011010000 | -5.2235070000 | -1.2621430000 |
| O | -0.6320700000 | -1.2364800000 | -2.4825160000 |
| H | 0.0374100000 | -1.3857240000 | -3.1701810000 |
| H | -0.7976260000 | -0.2677980000 | -2.5101750000 |
| F | -0.1744980000 | 1.0127420000 | -1.2610050000 |

**$^{6}$Im$_3$WatN$_3$_FeF**
Charge: 1
Multiplicity: 6
B3LYP-D3(BJ)/6-31G(d)-SDD SCF energy (au): -1143.07370529
B3LYP-D3(BJ)/6-31G(d)-SDD enthalpy (au): -1142.78776629
B3LYP-D3(BJ)/6-31G(d)-SDD free energy (au): -1142.86892629
B3LYP-D3(BJ)/def2-TZVP/SMD SCF energy (au): -2283.41282535
B3LYP-D3(BJ)/def2-TZVP/SMD enthalpy (au): -2283.12688635
B3LYP-D3(BJ)/def2-TZVP/SMD free energy (au): -2283.20804635
B3LYP-D3(BJ)/def2-TZVP/SMD free energy (quasi-harmonic) (au): -2283.20235306

Cartesian coordinates

| ATOM | X | Y | Z |
|---|---|---|---|
| Fe | 0.3290050000 | -0.3119630000 | -0.7178980000 |
| C | 1.5482200000 | 1.0664080000 | 3.2063450000 |
| N | 2.0335510000 | 2.0443290000 | 2.3571740000 |
| H | 2.5032910000 | 2.8951220000 | 2.6372630000 |
| C | 1.7659700000 | 1.6805340000 | 1.0847230000 |
| H | 2.0188970000 | 2.2588920000 | 0.2090400000 |
| N | 1.1330440000 | 0.5128880000 | 1.0761460000 |

| | | | |
|---|---|---|---|
| C | 0.9857780000 | 0.1194250000 | 2.3951820000 |
| H | 0.4847070000 | -0.8001340000 | 2.6539250000 |
| C | -2.4246610000 | -2.7064160000 | 1.7400900000 |
| N | -3.1192080000 | -1.5386110000 | 1.4832760000 |
| H | -4.0480330000 | -1.3145910000 | 1.8143310000 |
| C | -2.3540470000 | -0.7480300000 | 0.6986650000 |
| H | -2.6323710000 | 0.2293600000 | 0.3367920000 |
| N | -1.1986590000 | -1.3510380000 | 0.4389190000 |
| C | -1.2311090000 | -2.5755440000 | 1.0839390000 |
| H | -0.4087940000 | -3.2690580000 | 1.0163040000 |
| H | -2.8366380000 | -3.5012240000 | 2.3416770000 |
| H | 1.6445590000 | 1.1359960000 | 4.2783220000 |
| C | 2.3109660000 | -4.1931670000 | -0.9103930000 |
| N | 3.1207160000 | -3.5248830000 | -0.0116710000 |
| H | 3.9897160000 | -3.8685900000 | 0.3742860000 |
| C | 2.5887080000 | -2.3033260000 | 0.2167120000 |
| H | 3.0259070000 | -1.5640810000 | 0.8686460000 |
| N | 1.4702700000 | -2.1577850000 | -0.4834060000 |
| C | 1.2864320000 | -3.3326370000 | -1.1958200000 |
| H | 0.4408650000 | -3.4580180000 | -1.8531920000 |
| H | 2.5340880000 | -5.1906530000 | -1.2545370000 |
| O | -0.4569890000 | -1.2242110000 | -2.5994920000 |
| H | 0.1967650000 | -0.8713600000 | -3.2320510000 |
| H | -1.2401490000 | -0.6531480000 | -2.7088450000 |
| F | -0.9432870000 | 0.9570200000 | -1.0554140000 |
| N | 4.0781960000 | -0.1889670000 | -1.6234650000 |
| N | 2.9667150000 | 0.0979140000 | -1.7158390000 |
| N | 1.7983470000 | 0.4196880000 | -1.8119130000 |

**$^{5}$Im$_3$WatN$_3$_TS-1**
Charge: 1
Multiplicity: 5
B3LYP-D3(BJ)/6-31G(d)-SDD SCF energy (au): -1779.27826697
B3LYP-D3(BJ)/6-31G(d)-SDD enthalpy (au): -1778.69037497
B3LYP-D3(BJ)/6-31G(d)-SDD free energy (au): -1778.80857397
B3LYP-D3(BJ)/def2-TZVP/SMD SCF energy (au): -2919.86033980
B3LYP-D3(BJ)/def2-TZVP/SMD enthalpy (au): -2919.27244780
B3LYP-D3(BJ)/def2-TZVP/SMD free energy (au): -2919.39064680
B3LYP-D3(BJ)/def2-TZVP/SMD free energy (quasi-harmonic) (au): -2919.37915083

Cartesian coordinates

| ATOM | X | Y | Z |
|---|---|---|---|
| Fe | 0.3118910000 | -0.4076850000 | -0.8764230000 |
| C | 1.5588250000 | 0.9756170000 | 3.1414590000 |
| N | 2.2915340000 | 1.7847110000 | 2.2945640000 |
| H | 2.9617040000 | 2.4875750000 | 2.5741310000 |
| C | 1.9635900000 | 1.4700160000 | 1.0172120000 |
| H | 2.3854100000 | 1.9418210000 | 0.1436830000 |
| N | 1.0588660000 | 0.5036020000 | 1.0016280000 |
| C | 0.7952220000 | 0.1863080000 | 2.3220170000 |
| H | 0.0795160000 | -0.5778840000 | 2.5825600000 |
| C | -2.5999300000 | -2.5009630000 | 1.7116860000 |
| N | -3.2296950000 | -1.2891290000 | 1.4976400000 |

| | | | |
|---|---|---|---|
| H | -4.1283110000 | -1.0136700000 | 1.8692990000 |
| C | -2.4460700000 | -0.5392990000 | 0.6855070000 |
| H | -2.6884990000 | 0.4557090000 | 0.3512760000 |
| N | -1.3460670000 | -1.2056370000 | 0.3703400000 |
| C | -1.4287790000 | -2.4333720000 | 1.0043970000 |
| H | -0.6412070000 | -3.1638070000 | 0.9044270000 |
| H | -3.0332880000 | -3.2742660000 | 2.3261920000 |
| H | 1.6440290000 | 1.0390780000 | 4.2149170000 |
| C | 2.9120410000 | -3.8192870000 | -1.1696270000 |
| N | 3.2298190000 | -3.4022370000 | 0.1079850000 |
| H | 4.0307220000 | -3.7023380000 | 0.6459400000 |
| C | 2.3475250000 | -2.4431030000 | 0.4738120000 |
| H | 2.3802930000 | -1.9182980000 | 1.4159840000 |
| N | 1.4626940000 | -2.2401920000 | -0.4932190000 |
| C | 1.8120070000 | -3.0911660000 | -1.5302650000 |
| H | 1.2708240000 | -3.0864060000 | -2.4637460000 |
| H | 3.4881060000 | -4.5674220000 | -1.6904820000 |
| O | -0.7526810000 | -1.4058690000 | -2.4731460000 |
| H | -1.5285260000 | -0.7933370000 | -2.6102030000 |
| H | -1.1486020000 | -2.2194720000 | -2.1198660000 |
| C | -5.8153100000 | 2.6216730000 | -4.5465880000 |
| C | -4.7138020000 | 1.9023240000 | -4.1056380000 |
| C | -4.0896050000 | 2.2314830000 | -2.8877490000 |
| C | -4.6028200000 | 3.2550180000 | -2.0612330000 |
| C | -5.7263950000 | 3.9514760000 | -2.5257790000 |
| C | -6.3160590000 | 3.6593760000 | -3.7532430000 |
| H | -6.2847300000 | 2.3764990000 | -5.4937420000 |
| H | -4.3088440000 | 1.0886580000 | -4.6987590000 |
| H | -6.1536440000 | 4.7300890000 | -1.8993930000 |
| H | -7.1797110000 | 4.2280110000 | -4.0845600000 |
| C | -4.0743960000 | 3.5789400000 | -0.6769430000 |
| C | -3.3062670000 | 4.9060210000 | -0.5971130000 |
| H | -3.4298640000 | 2.7749920000 | -0.3121580000 |
| H | -4.9361500000 | 3.6281030000 | 0.0008350000 |
| H | -3.0559510000 | 5.1415130000 | 0.4425350000 |
| H | -2.3743360000 | 4.8465710000 | -1.1651200000 |
| H | -3.8999700000 | 5.7345130000 | -0.9976620000 |
| C | -2.8831970000 | 1.4353810000 | -2.5449690000 |
| O | -2.8795880000 | 0.2042270000 | -2.5846280000 |
| N | -1.8150370000 | 2.2461540000 | -2.1691560000 |
| C | -0.8529470000 | 2.7596430000 | -3.1754100000 |
| C | -1.6187270000 | 3.8476480000 | -3.9764550000 |
| C | -0.3589940000 | 1.6679250000 | -4.1355000000 |
| H | -0.8921820000 | 4.3267480000 | -4.6393620000 |
| H | -2.4155170000 | 3.4214900000 | -4.5881500000 |
| H | -2.0471670000 | 4.6075580000 | -3.3204390000 |
| H | 0.2286480000 | 0.9108060000 | -3.6146520000 |
| H | -1.1921010000 | 1.1905050000 | -4.6621580000 |
| H | 0.2906690000 | 2.1277460000 | -4.8868820000 |
| F | -0.8909120000 | 1.2269600000 | -1.1043460000 |
| C | 0.3098040000 | 3.4187020000 | -2.4226940000 |
| H | -0.0655540000 | 4.1158390000 | -1.6674280000 |
| H | 0.9353880000 | 2.6621980000 | -1.9477420000 |

| | | | |
|---|---|---|---|
| H | 0.9244350000 | 3.9747780000 | -3.1370740000 |
| N | 4.0705440000 | -0.3383140000 | -1.4850870000 |
| N | 2.9835150000 | -0.0030440000 | -1.6814800000 |
| N | 1.8460170000 | 0.3693010000 | -1.8807360000 |

**$^5Im_3WatN_3$_TS-3-F**

Charge: 1
Multiplicity: 5
B3LYP-D3(BJ)/6-31G(d)-SDD SCF energy (au): -1779.35280166
B3LYP-D3(BJ)/6-31G(d)-SDD enthalpy (au): -1778.76423366
B3LYP-D3(BJ)/6-31G(d)-SDD free energy (au): -1778.88297966
B3LYP-D3(BJ)/def2-TZVP/SMD SCF energy (au): -2919.94294440
B3LYP-D3(BJ)/def2-TZVP/SMD enthalpy (au): -2919.35437640
B3LYP-D3(BJ)/def2-TZVP/SMD free energy (au): -2919.47312240
B3LYP-D3(BJ)/def2-TZVP/SMD free energy (quasi-harmonic) (au): -2919.46087774

Cartesian coordinates

| ATOM | X | Y | Z |
|---|---|---|---|
| H | 32.9776240000 | 26.2852680000 | 37.5967610000 |
| C | 33.2479890000 | 27.2595430000 | 37.2216580000 |
| C | 34.4338620000 | 27.7777000000 | 36.7770440000 |
| N | 32.3314080000 | 28.2865970000 | 37.1100520000 |
| C | 32.9654410000 | 29.3691470000 | 36.6024160000 |
| N | 34.2468800000 | 29.0958560000 | 36.3973700000 |
| H | 31.3425730000 | 28.2247540000 | 37.3075640000 |
| H | 35.3996210000 | 27.3044660000 | 36.7007940000 |
| H | 32.4903240000 | 30.3146570000 | 36.3961960000 |
| H | 39.4340380000 | 26.6222620000 | 35.9365790000 |
| C | 38.6337480000 | 27.2185080000 | 35.5273680000 |
| C | 37.9473610000 | 28.2984390000 | 36.0152520000 |
| N | 38.0909760000 | 26.9682740000 | 34.2803470000 |
| C | 37.1124140000 | 27.8773510000 | 34.0514380000 |
| N | 37.0032030000 | 28.6950990000 | 35.0870570000 |
| H | 38.3712600000 | 26.2322670000 | 33.6472380000 |
| H | 38.0549430000 | 28.8103730000 | 36.9586160000 |
| H | 36.5036600000 | 27.9291680000 | 33.1615400000 |
| H | 37.4165060000 | 31.4194760000 | 40.1198340000 |
| C | 37.2479390000 | 31.4021420000 | 39.0545900000 |
| C | 36.3042760000 | 30.7809800000 | 38.2804740000 |
| N | 38.0524390000 | 32.0986170000 | 38.1696630000 |
| C | 37.5871730000 | 31.8858030000 | 36.9146040000 |
| N | 36.5282100000 | 31.0901590000 | 36.9522160000 |
| H | 38.8474190000 | 32.6716200000 | 38.4162450000 |
| H | 35.4870470000 | 30.1396180000 | 38.5701590000 |
| H | 38.0167930000 | 32.2925770000 | 36.0124810000 |
| Fe | 35.5628480000 | 30.3356070000 | 35.1212270000 |
| O | 34.1051960000 | 31.9031970000 | 35.5088250000 |
| H | 34.5952050000 | 32.7199020000 | 35.6947550000 |
| H | 33.5355950000 | 32.1011530000 | 34.7013180000 |
| F | 36.7446350000 | 31.4915230000 | 34.1780550000 |
| N | 33.3388470000 | 32.6632610000 | 31.1432390000 |
| O | 32.7115790000 | 32.3645500000 | 33.3209020000 |
| C | 34.5749330000 | 33.7993640000 | 32.8638450000 |

| | | | |
|---|---|---|---|
| C | 34.1894900000 | 34.9835440000 | 33.4900340000 |
| C | 35.1370550000 | 35.9568540000 | 33.8161590000 |
| C | 36.4873560000 | 35.7383460000 | 33.5130610000 |
| C | 36.8883470000 | 34.5477620000 | 32.9264230000 |
| C | 35.9521000000 | 33.5316600000 | 32.6056880000 |
| C | 33.4711100000 | 32.8581610000 | 32.4657100000 |
| C | 32.2548120000 | 31.9126460000 | 30.4508460000 |
| C | 32.2303960000 | 30.4581010000 | 30.9347330000 |
| C | 30.9091060000 | 32.6035370000 | 30.7204390000 |
| C | 32.5985890000 | 31.9697840000 | 28.9573280000 |
| C | 36.3654800000 | 32.2340770000 | 32.1844990000 |
| C | 37.7209940000 | 31.8783100000 | 31.6891030000 |
| H | 33.1367220000 | 35.1522940000 | 33.6960490000 |
| H | 34.8233410000 | 36.8837160000 | 34.2857350000 |
| H | 37.2255440000 | 36.4989280000 | 33.7488010000 |
| H | 37.9415320000 | 34.3771150000 | 32.7346370000 |
| H | 33.9581000000 | 33.2039760000 | 30.5537300000 |
| H | 35.5921380000 | 31.4865630000 | 32.0426890000 |
| H | 38.5191380000 | 32.4236790000 | 32.1981550000 |
| H | 37.8953170000 | 30.8079560000 | 31.8218080000 |
| H | 37.7999900000 | 32.0987970000 | 30.6130580000 |
| H | 31.5054500000 | 29.8900750000 | 30.3428870000 |
| H | 33.2145010000 | 29.9940850000 | 30.8159370000 |
| H | 31.9422610000 | 30.3948690000 | 31.9834680000 |
| H | 30.6870390000 | 32.6062640000 | 31.7901340000 |
| H | 30.9218200000 | 33.6373260000 | 30.3582280000 |
| H | 30.1070030000 | 32.0686120000 | 30.2017260000 |
| H | 33.5609150000 | 31.4851990000 | 28.7551940000 |
| H | 31.8308380000 | 31.4479810000 | 28.3798300000 |
| H | 32.6413530000 | 33.0048210000 | 28.5960990000 |
| N | 34.6577910000 | 29.6504580000 | 33.4484250000 |
| N | 33.6473940000 | 28.9800170000 | 33.5055120000 |
| N | 32.6811900000 | 28.3491280000 | 33.5387610000 |

**$^{5}$Im$_3$WatN$_3$_TS-3-N$_3$**
Charge: 1
Multiplicity: 5
B3LYP-D3(BJ)/6-31G(d)-SDD SCF energy (au): -1779.36187779
B3LYP-D3(BJ)/6-31G(d)-SDD enthalpy (au): -1778.77304579
B3LYP-D3(BJ)/6-31G(d)-SDD free energy (au): -1778.89114779
B3LYP-D3(BJ)/def2-TZVP/SMD SCF energy (au): -2919.95008180
B3LYP-D3(BJ)/def2-TZVP/SMD enthalpy (au): -2919.36124980
B3LYP-D3(BJ)/def2-TZVP/SMD free energy (au): -2919.47935180
B3LYP-D3(BJ)/def2-TZVP/SMD free energy (quasi-harmonic) (au): -2919.46759701

Cartesian coordinates

| ATOM | X | Y | Z |
|---|---|---|---|
| H | 32.8400230000 | 27.0610100000 | 38.2516600000 |
| C | 33.1280790000 | 27.8623910000 | 37.5900080000 |
| C | 34.3399720000 | 28.2459540000 | 37.0832270000 |
| N | 32.2119260000 | 28.7552200000 | 37.0690950000 |
| C | 32.8713880000 | 29.6337930000 | 36.2775980000 |
| N | 34.1682670000 | 29.3540090000 | 36.2714400000 |

| | | | |
|---|---|---|---|
| H | 31.2136450000 | 28.7409830000 | 37.2236780000 |
| H | 35.3126600000 | 27.8056660000 | 37.2329260000 |
| H | 32.3874170000 | 30.4171910000 | 35.7137220000 |
| H | 39.2839600000 | 26.4261650000 | 36.3124050000 |
| C | 38.5385970000 | 27.0217870000 | 35.8094230000 |
| C | 37.8209880000 | 28.1206280000 | 36.1992190000 |
| N | 38.1248420000 | 26.7494450000 | 34.5189950000 |
| C | 37.1880820000 | 27.6644320000 | 34.1715800000 |
| N | 36.9800180000 | 28.5066490000 | 35.1712760000 |
| H | 38.4647940000 | 26.0009270000 | 33.9314840000 |
| H | 37.8485090000 | 28.6605010000 | 37.1325470000 |
| H | 36.6965440000 | 27.7076720000 | 33.2123600000 |
| H | 37.3982120000 | 31.8465080000 | 39.9101660000 |
| C | 37.2928070000 | 31.6518710000 | 38.8545360000 |
| C | 36.3522770000 | 30.9648460000 | 38.1338340000 |
| N | 38.1959450000 | 32.1297540000 | 37.9223150000 |
| C | 37.7894760000 | 31.7305860000 | 36.6925120000 |
| N | 36.6764670000 | 31.0195040000 | 36.7923440000 |
| H | 39.0114460000 | 32.6926070000 | 38.1195730000 |
| H | 35.4702640000 | 30.4470580000 | 38.4762270000 |
| H | 38.2886900000 | 31.9578010000 | 35.7632200000 |
| Fe | 35.7011430000 | 30.2849110000 | 34.9691120000 |
| O | 34.5165260000 | 32.0946210000 | 35.1377510000 |
| H | 35.0287550000 | 32.6905260000 | 34.5636460000 |
| H | 33.6125930000 | 32.0387640000 | 34.7224330000 |
| F | 36.9352580000 | 31.2336210000 | 33.9747570000 |
| N | 31.2866370000 | 31.5123130000 | 31.8652840000 |
| O | 32.1131480000 | 31.7387700000 | 33.9820470000 |
| C | 33.4552410000 | 32.5446350000 | 32.1910900000 |
| C | 33.6669830000 | 33.8823370000 | 32.5181810000 |
| C | 34.8539830000 | 34.5253870000 | 32.1555100000 |
| C | 35.8512260000 | 33.8056540000 | 31.4855870000 |
| C | 35.6602140000 | 32.4689810000 | 31.1729870000 |
| C | 34.4554210000 | 31.7946740000 | 31.5011290000 |
| C | 32.2240720000 | 31.8893600000 | 32.7468410000 |
| C | 29.9729610000 | 30.8797380000 | 32.1697770000 |
| C | 30.1968430000 | 29.5369370000 | 32.8789830000 |
| C | 29.1354080000 | 31.8324180000 | 33.0365150000 |
| C | 29.2912740000 | 30.6521590000 | 30.8151710000 |
| C | 34.2797190000 | 30.4131820000 | 31.2136990000 |
| C | 35.2880630000 | 29.5779020000 | 30.4992860000 |
| H | 32.9000510000 | 34.4223970000 | 33.0660630000 |
| H | 35.0004000000 | 35.5726670000 | 32.3995270000 |
| H | 36.7832950000 | 34.2923500000 | 31.2162440000 |
| H | 36.4518760000 | 31.9207540000 | 30.6767230000 |
| H | 31.4643540000 | 31.7557960000 | 30.8992750000 |
| H | 33.2899320000 | 29.9921970000 | 31.3460730000 |
| H | 36.2837050000 | 29.6907000000 | 30.9409180000 |
| H | 35.0099480000 | 28.5216400000 | 30.5378480000 |
| H | 35.3619540000 | 29.8623330000 | 29.4383640000 |
| H | 29.2310410000 | 29.0549220000 | 33.0624970000 |
| H | 30.8045540000 | 28.8630920000 | 32.2678530000 |
| H | 30.7039330000 | 29.6755220000 | 33.8350940000 |

H 29.6342180000 32.0308450000 33.9876300000
H 28.9735340000 32.7842980000 32.5199170000
H 28.1577690000 31.3838370000 33.2405370000
H 29.8850110000 29.9839820000 30.1805380000
H 28.3118790000 30.1899120000 30.9649760000
H 29.1364640000 31.5989240000 30.2842120000
N 34.7754610000 29.4051090000 33.3851790000
N 33.9028070000 28.5643010000 33.4703230000
N 33.0731200000 27.7634030000 33.5117990000

**$^{5}Im_3$WatOH_11**
Charge: 1
Multiplicity: 5
B3LYP-D3(BJ)/6-31G(d)-SDD SCF energy (au): -1690.91838365
B3LYP-D3(BJ)/6-31G(d)-SDD enthalpy (au): -1690.32872265
B3LYP-D3(BJ)/6-31G(d)-SDD free energy (au): -1690.44564165
B3LYP-D3(BJ)/def2-TZVP/SMD SCF energy (au): -2831.47273599
B3LYP-D3(BJ)/def2-TZVP/SMD enthalpy (au): -2830.88307499
B3LYP-D3(BJ)/def2-TZVP/SMD free energy (au): -2830.99999399
B3LYP-D3(BJ)/def2-TZVP/SMD free energy (quasi-harmonic) (au): -2830.98779634

Cartesian coordinates
ATOM X Y Z
Fe 0.0196270000 -0.4964380000 -0.5244470000
C -0.0367070000 1.7868010000 3.3159830000
N 0.8731290000 2.4292590000 2.4969210000
H 1.3731470000 3.2793390000 2.7164550000
C 0.9588260000 1.7381750000 1.3311120000
H 1.5514440000 2.0212950000 0.4738340000
N 0.1561870000 0.6846030000 1.3649920000
C -0.4729800000 0.7052500000 2.5959240000
H -1.1979230000 -0.0464370000 2.8695150000
C -3.2512380000 -2.5073610000 1.5168650000
N -3.9329520000 -1.8809930000 0.4907890000
H -4.9327120000 -1.8989800000 0.3438960000
C -3.0340930000 -1.2477420000 -0.3013790000
H -3.2725010000 -0.6925290000 -1.1990430000
N -1.8071620000 -1.4343390000 0.1701380000
C -1.9295980000 -2.2211620000 1.3027240000
H -1.0623890000 -2.5301470000 1.8662810000
O 0.6232780000 1.0121640000 -1.4756850000
H 0.8799260000 0.7856830000 -2.3817960000
H -3.7548630000 -3.0869770000 2.2742840000
H -0.2814770000 2.1505860000 4.3015180000
C 3.0866300000 -3.4917850000 0.0372520000
N 3.0319790000 -2.9411470000 1.3047030000
H 3.6259150000 -3.1879100000 2.0840580000
C 2.0581050000 -1.9991660000 1.3159950000
H 1.8042870000 -1.3949320000 2.1736870000
N 1.4825570000 -1.9233440000 0.1245900000
C 2.1175910000 -2.8489130000 -0.6845310000
H 1.8359020000 -2.9606040000 -1.7199790000
H 3.7944830000 -4.2625620000 -0.2236520000

| | | | |
|---|---|---|---|
| O | -0.2539310000 | -1.7209550000 | -2.4326460000 |
| H | -0.8968220000 | -1.0990400000 | -2.8630760000 |
| H | -0.7897020000 | -2.4986020000 | -2.2080490000 |
| C | -5.5189670000 | 3.0020550000 | -3.5032460000 |
| C | -4.3750300000 | 2.2835230000 | -3.8397450000 |
| C | -3.3708140000 | 2.0746840000 | -2.8868490000 |
| C | -3.4893790000 | 2.5673980000 | -1.5757710000 |
| C | -4.6520660000 | 3.2836630000 | -1.2620430000 |
| C | -5.6538060000 | 3.5022100000 | -2.2068290000 |
| H | -6.2958950000 | 3.1685370000 | -4.2426760000 |
| H | -4.2534020000 | 1.8821230000 | -4.8419340000 |
| H | -4.7738570000 | 3.6841360000 | -0.2602640000 |
| H | -6.5407050000 | 4.0648140000 | -1.9307740000 |
| C | -2.3999010000 | 2.3481460000 | -0.5427610000 |
| C | -2.0201580000 | 3.6089830000 | 0.2453010000 |
| H | -1.4960420000 | 1.9555720000 | -1.0194600000 |
| H | -2.7255450000 | 1.5750440000 | 0.1673700000 |
| H | -1.2153530000 | 3.3780060000 | 0.9486980000 |
| H | -1.6693650000 | 4.4020220000 | -0.4243880000 |
| H | -2.8599230000 | 4.0092150000 | 0.8221670000 |
| C | -2.2283490000 | 1.1871270000 | -3.2664080000 |
| O | -2.2543320000 | -0.0146880000 | -2.9476440000 |
| N | -1.2346580000 | 1.7073450000 | -4.0138650000 |
| C | -0.7392320000 | 3.0371480000 | -4.4835770000 |
| C | -1.5451680000 | 4.1863890000 | -3.8668710000 |
| C | -0.8600340000 | 3.0546030000 | -6.0141060000 |
| H | -1.0842980000 | 5.1153470000 | -4.2156980000 |
| H | -2.5894240000 | 4.1898460000 | -4.1778550000 |
| H | -1.4985320000 | 4.1736950000 | -2.7760530000 |
| H | -0.3242890000 | 2.2103720000 | -6.4557880000 |
| H | -1.9098000000 | 3.0043470000 | -6.3205360000 |
| H | -0.4287080000 | 3.9801340000 | -6.4073240000 |
| F | -0.3098040000 | 0.7189040000 | -4.3820370000 |
| C | 0.7277580000 | 3.1643800000 | -4.0329200000 |
| H | 0.8108850000 | 3.0057350000 | -2.9542680000 |
| H | 1.3676820000 | 2.4424430000 | -4.5444500000 |
| H | 1.0843310000 | 4.1682100000 | -4.2809280000 |

**$^{5}Im_3$WatOH_FeF_remove-F**
Charge: 1
Multiplicity: 5
B3LYP-D3(BJ)/6-31G(d)-SDD SCF energy (au): -954.840129310
B3LYP-D3(BJ)/6-31G(d)-SDD enthalpy (au): -954.559932310
B3LYP-D3(BJ)/6-31G(d)-SDD free energy (au): -954.635646310
B3LYP-D3(BJ)/def2-TZVP/SMD SCF energy (au): -2095.10486805
B3LYP-D3(BJ)/def2-TZVP/SMD enthalpy (au): -2094.82467105
B3LYP-D3(BJ)/def2-TZVP/SMD free energy (au): -2094.90038505
B3LYP-D3(BJ)/def2-TZVP/SMD free energy (quasi-harmonic) (au): -2094.894789740

Cartesian coordinates

| ATOM | X | Y | Z |
|---|---|---|---|
| Fe | 0.0511950000 | -0.2493030000 | -0.5206620000 |
| C | 1.6383310000 | 0.7355820000 | 3.4676790000 |

| | | | |
|---|---|---|---|
| N | 2.1222220000 | 1.7381070000 | 2.6508510000 |
| H | 2.6967610000 | 2.5168640000 | 2.9432080000 |
| C | 1.6908900000 | 1.5048880000 | 1.3869820000 |
| H | 1.9167570000 | 2.1228270000 | 0.5305590000 |
| N | 0.9528220000 | 0.4039310000 | 1.3523700000 |
| C | 0.9134540000 | -0.0871370000 | 2.6459580000 |
| H | 0.3736440000 | -0.9899170000 | 2.8905610000 |
| C | -3.9056140000 | -0.3519560000 | 1.1870620000 |
| N | -4.1354600000 | -0.6791940000 | -0.1367670000 |
| H | -5.0390540000 | -0.8457450000 | -0.5582460000 |
| C | -2.9474060000 | -0.7242240000 | -0.7806040000 |
| H | -2.8142940000 | -0.9380290000 | -1.8306810000 |
| N | -1.9627870000 | -0.4439880000 | 0.0666650000 |
| C | -2.5501040000 | -0.2079410000 | 1.2982100000 |
| H | -1.9583750000 | 0.0569150000 | 2.1605860000 |
| O | 0.4658120000 | 1.2712540000 | -1.5806080000 |
| H | -0.0277320000 | 2.0875760000 | -1.4121280000 |
| H | -4.7064050000 | -0.2527990000 | 1.9026110000 |
| H | 1.8461740000 | 0.7035450000 | 4.5256780000 |
| C | 1.3861590000 | -4.3260810000 | -0.8737860000 |
| N | 2.3336930000 | -3.8092330000 | -0.0105260000 |
| H | 3.1223830000 | -4.3116670000 | 0.3736250000 |
| C | 2.0343510000 | -2.5110040000 | 0.2245420000 |
| H | 2.6065480000 | -1.8609120000 | 0.8680850000 |
| N | 0.9428650000 | -2.1672650000 | -0.4469840000 |
| C | 0.5280260000 | -3.2940320000 | -1.1372140000 |
| H | -0.3514230000 | -3.2739220000 | -1.7615290000 |
| H | 1.4113030000 | -5.3508350000 | -1.2090500000 |
| O | -0.6729020000 | -0.7277610000 | -2.8054580000 |
| H | -0.3088880000 | 0.2001390000 | -2.7355250000 |
| H | -0.0276830000 | -1.2046270000 | -3.3513410000 |

**$^5$Im$_3$WatOH_FeF_remove-OH**

Charge: 1
Multiplicity: 5
B3LYP-D3(BJ)/6-31G(d)-SDD SCF energy (au): -978.885373045
B3LYP-D3(BJ)/6-31G(d)-SDD enthalpy (au): -978.616364045
B3LYP-D3(BJ)/6-31G(d)-SDD free energy (au): -978.691752045
B3LYP-D3(BJ)/def2-TZVP/SMD SCF energy (au): -2119.15781167
B3LYP-D3(BJ)/def2-TZVP/SMD enthalpy (au): -2118.88880267
B3LYP-D3(BJ)/def2-TZVP/SMD free energy (au): -2118.96419067
B3LYP-D3(BJ)/def2-TZVP/SMD free energy (quasi-harmonic) (au): -2118.958198625

Cartesian coordinates

| ATOM | X | Y | Z |
|---|---|---|---|
| Fe | -0.0869750000 | -0.2235080000 | -0.5234440000 |
| C | 1.4761480000 | 0.7775120000 | 3.4645860000 |
| N | 1.6527930000 | 1.9255050000 | 2.7179240000 |
| H | 2.0377230000 | 2.7983150000 | 3.0525260000 |
| C | 1.2213220000 | 1.6869710000 | 1.4558770000 |
| H | 1.2363540000 | 2.4008310000 | 0.6459280000 |
| N | 0.7765920000 | 0.4416400000 | 1.3554320000 |
| C | 0.9307860000 | -0.1376120000 | 2.6029350000 |

| | | | |
|---|---|---|---|
| H | 0.6424660000 | -1.1614550000 | 2.7902640000 |
| C | -3.9462160000 | -0.5917510000 | 1.3302410000 |
| N | -4.2537120000 | -0.6948020000 | -0.0141150000 |
| H | -5.1818010000 | -0.7823030000 | -0.4052530000 |
| C | -3.1045080000 | -0.6379690000 | -0.7230240000 |
| H | -3.0362890000 | -0.6816010000 | -1.7994510000 |
| N | -2.0708990000 | -0.5061220000 | 0.1016540000 |
| C | -2.5850950000 | -0.4743870000 | 1.3869870000 |
| H | -1.9422570000 | -0.3569850000 | 2.2453040000 |
| H | -4.7047210000 | -0.6078670000 | 2.0966860000 |
| H | 1.7464120000 | 0.7204190000 | 4.5072480000 |
| C | 1.3330640000 | -4.2628440000 | -0.9572650000 |
| N | 2.3494160000 | -3.6865800000 | -0.2193680000 |
| H | 3.2001970000 | -4.1455620000 | 0.0770200000 |
| C | 2.0192920000 | -2.3988480000 | 0.0310700000 |
| H | 2.6318430000 | -1.7100720000 | 0.5925100000 |
| N | 0.8410610000 | -2.1183100000 | -0.5116650000 |
| C | 0.4021330000 | -3.2758050000 | -1.1322990000 |
| H | -0.5462440000 | -3.3120720000 | -1.6461510000 |
| H | 1.3663860000 | -5.2922700000 | -1.2772010000 |
| O | -0.8254950000 | -0.4412240000 | -2.7626970000 |
| H | -0.2200790000 | -0.9409750000 | -3.3336450000 |
| H | -0.4404300000 | 0.4668750000 | -2.7199700000 |
| F | 0.4172390000 | 1.3433150000 | -1.3921640000 |

**$^{6}$Im$_3$WatOH_FeF**

Charge: 1
Multiplicity: 6
B3LYP-D3(BJ)/6-31G(d)-SDD SCF energy (au): -1054.69822175
B3LYP-D3(BJ)/6-31G(d)-SDD enthalpy (au): -1054.41368875
B3LYP-D3(BJ)/6-31G(d)-SDD free energy (au): -1054.49054475
B3LYP-D3(BJ)/def2-TZVP/SMD SCF energy (au): -2195.00741246
B3LYP-D3(BJ)/def2-TZVP/SMD enthalpy (au): -2194.72287946
B3LYP-D3(BJ)/def2-TZVP/SMD free energy (au): -2194.79973546
B3LYP-D3(BJ)/def2-TZVP/SMD free energy (quasi-harmonic) (au): -2194.79487071

Cartesian coordinates

| ATOM | X | Y | Z |
|---|---|---|---|
| Fe | 0.1804000000 | 0.0181660000 | -0.7926600000 |
| C | 0.7325220000 | 1.4418660000 | 3.2800900000 |
| N | 1.9339540000 | 1.7394060000 | 2.6628430000 |
| H | 2.7109570000 | 2.2317900000 | 3.0822440000 |
| C | 1.8876320000 | 1.2815630000 | 1.3928180000 |
| H | 2.6783910000 | 1.3742370000 | 0.6649610000 |
| N | 0.7150090000 | 0.7000140000 | 1.1660070000 |
| C | -0.0168340000 | 0.7958330000 | 2.3361200000 |
| H | -1.0148930000 | 0.3964650000 | 2.4100640000 |
| C | -3.4783380000 | -1.4858480000 | 1.2072720000 |
| N | -3.9041840000 | -0.3764190000 | 0.5004190000 |
| H | -4.8453190000 | -0.0074630000 | 0.4837180000 |
| C | -2.8479030000 | 0.1322250000 | -0.1743610000 |
| H | -2.8727620000 | 1.0132340000 | -0.7963180000 |
| N | -1.7648110000 | -0.5976590000 | 0.0602930000 |

| | | | |
|---|---|---|---|
| C | -2.1449250000 | -1.6103630000 | 0.9240400000 |
| H | -1.4384600000 | -2.3456820000 | 1.2752890000 |
| O | 1.8929380000 | 0.3715800000 | -1.4568400000 |
| H | 1.9195430000 | 1.2400790000 | -1.8932880000 |
| H | -4.1451960000 | -2.0667010000 | 1.8247780000 |
| H | 0.5288930000 | 1.7126900000 | 4.3039760000 |
| C | 1.0204660000 | -4.2686340000 | -0.6399520000 |
| N | 2.2133150000 | -3.6995220000 | -0.2310370000 |
| H | 3.0695800000 | -4.1973010000 | -0.0286550000 |
| C | 2.0482900000 | -2.3588360000 | -0.1718280000 |
| H | 2.8253240000 | -1.6551680000 | 0.0812850000 |
| N | 0.8075600000 | -2.0406190000 | -0.5125770000 |
| C | 0.1552190000 | -3.2234390000 | -0.8142120000 |
| H | -0.8716830000 | -3.2274710000 | -1.1431470000 |
| H | 0.9052990000 | -5.3331010000 | -0.7693570000 |
| O | -0.5115570000 | -0.8093480000 | -2.7678910000 |
| H | 0.3309110000 | -0.9548430000 | -3.2336980000 |
| H | -0.8445330000 | 0.0480460000 | -3.0941860000 |
| F | -0.7058170000 | 1.4933430000 | -1.4498580000 |

**$^{5}$Im$_3$WatOH_TS-1**
Charge: 1
Multiplicity: 5
B3LYP-D3(BJ)/6-31G(d)-SDD SCF energy (au): -1690.89186837
B3LYP-D3(BJ)/6-31G(d)-SDD enthalpy (au): -1690.30575237
B3LYP-D3(BJ)/6-31G(d)-SDD free energy (au): -1690.42189937
B3LYP-D3(BJ)/def2-TZVP/SMD SCF energy (au): -2831.44546097
B3LYP-D3(BJ)/def2-TZVP/SMD enthalpy (au): -2830.85934497
B3LYP-D3(BJ)/def2-TZVP/SMD free energy (au): -2830.97549197
B3LYP-D3(BJ)/def2-TZVP/SMD free energy (quasi-harmonic) (au): -2830.96342160

Cartesian coordinates

| ATOM | X | Y | Z |
|---|---|---|---|
| Fe | 0.3748020000 | -0.6192920000 | -0.9983780000 |
| C | 2.1921940000 | 1.8331830000 | 2.2649330000 |
| N | 2.9841950000 | 2.0577460000 | 1.1545780000 |
| H | 3.7647720000 | 2.6970100000 | 1.1014590000 |
| C | 2.5263580000 | 1.2793970000 | 0.1403610000 |
| H | 2.9331010000 | 1.2403510000 | -0.8595980000 |
| N | 1.4837840000 | 0.5705090000 | 0.5457190000 |
| C | 1.2637330000 | 0.9063940000 | 1.8668940000 |
| H | 0.4600280000 | 0.4603770000 | 2.4331250000 |
| C | -2.3305160000 | -1.8775510000 | 2.2031530000 |
| N | -2.8010650000 | -0.6165850000 | 1.8854710000 |
| H | -3.5737610000 | -0.1418860000 | 2.3315420000 |
| C | -2.0619930000 | -0.1290630000 | 0.8602520000 |
| H | -2.1953690000 | 0.8389800000 | 0.4053320000 |
| N | -1.1387900000 | -1.0135760000 | 0.5081930000 |
| C | -1.2940390000 | -2.1087000000 | 1.3386540000 |
| H | -0.6536530000 | -2.9719630000 | 1.2496370000 |
| O | 1.6152840000 | -0.0893160000 | -2.2852460000 |
| H | 1.7372800000 | -0.7884660000 | -2.9469880000 |
| H | -2.7668980000 | -2.4742820000 | 2.9885220000 |

| | | | |
|---|---|---|---|
| H | 2.3598310000 | 2.3384450000 | 3.2031690000 |
| C | 2.2418230000 | -4.4789340000 | -0.6958030000 |
| N | 2.8493270000 | -3.8062680000 | 0.3487900000 |
| H | 3.5914030000 | -4.1655200000 | 0.9331680000 |
| C | 2.2975010000 | -2.5717520000 | 0.4380970000 |
| H | 2.5879790000 | -1.8208950000 | 1.1562370000 |
| N | 1.3646240000 | -2.4244910000 | -0.4897450000 |
| C | 1.3209990000 | -3.6054310000 | -1.2086780000 |
| H | 0.6466620000 | -3.7214420000 | -2.0429180000 |
| H | 2.5189380000 | -5.4844670000 | -0.9703790000 |
| O | -0.9336400000 | -1.7531280000 | -2.4214500000 |
| H | -1.4344840000 | -0.9884480000 | -2.8055120000 |
| H | -1.6087540000 | -2.2486850000 | -1.9305930000 |
| C | -5.3670740000 | 2.9109510000 | -5.1200120000 |
| C | -4.2757970000 | 2.1923390000 | -4.6489030000 |
| C | -3.9805400000 | 2.1705960000 | -3.2754150000 |
| C | -4.8146030000 | 2.8206170000 | -2.3444410000 |
| C | -5.9164400000 | 3.5259830000 | -2.8461270000 |
| C | -6.1838400000 | 3.5908490000 | -4.2114550000 |
| H | -5.5856320000 | 2.9362240000 | -6.1827490000 |
| H | -3.6326550000 | 1.6490320000 | -5.3340210000 |
| H | -6.5846460000 | 4.0197180000 | -2.1452030000 |
| H | -7.0419110000 | 4.1533560000 | -4.5667780000 |
| C | -4.6314540000 | 2.7269770000 | -0.8422530000 |
| C | -4.0219920000 | 3.9831990000 | -0.2036750000 |
| H | -4.0172180000 | 1.8558540000 | -0.5941940000 |
| H | -5.6187210000 | 2.5467560000 | -0.3987950000 |
| H | -4.0229540000 | 3.9022810000 | 0.8892080000 |
| H | -2.9896600000 | 4.1213640000 | -0.5345350000 |
| H | -4.5920350000 | 4.8784670000 | -0.4735010000 |
| C | -2.7564720000 | 1.4081840000 | -2.8794600000 |
| O | -2.6076950000 | 0.2302070000 | -3.1858390000 |
| N | -1.8890170000 | 2.1849760000 | -2.1181510000 |
| C | -0.9056660000 | 3.1043720000 | -2.7470440000 |
| C | -1.6991850000 | 4.2918110000 | -3.3506850000 |
| C | -0.0954100000 | 2.3976500000 | -3.8433240000 |
| H | -0.9667040000 | 5.0338380000 | -3.6827780000 |
| H | -2.2974700000 | 3.9898970000 | -4.2109940000 |
| H | -2.3563670000 | 4.7587460000 | -2.6138380000 |
| H | 0.4868440000 | 1.5707480000 | -3.4272100000 |
| H | -0.7519850000 | 2.0162450000 | -4.6333560000 |
| H | 0.5904140000 | 3.1189390000 | -4.3002540000 |
| F | -0.7902360000 | 0.9632170000 | -1.3761280000 |
| C | 0.0053100000 | 3.6375850000 | -1.6324060000 |
| H | -0.5869460000 | 4.0919540000 | -0.8315920000 |
| H | 0.6104390000 | 2.8348610000 | -1.2118110000 |
| H | 0.6685740000 | 4.4027010000 | -2.0478290000 |

**$N_3$□**
Charge: 0
Multiplicity: 2
B3LYP-D3(BJ)/6-31G(d)-SDD SCF energy (au): -164.135645456
B3LYP-D3(BJ)/6-31G(d)-SDD enthalpy (au): -164.123282456

B3LYP-D3(BJ)/6-31G(d)-SDD free energy (au): -164.145918456
B3LYP-D3(BJ)/def2-TZVP/SMD SCF energy (au): -164.204699031
B3LYP-D3(BJ)/def2-TZVP/SMD enthalpy (au): -164.192336031
B3LYP-D3(BJ)/def2-TZVP/SMD free energy (au): -164.214972031
B3LYP-D3(BJ)/def2-TZVP/SMD free energy (quasi-harmonic) (au): -164.214972031

Cartesian coordinates

| ATOM | X | Y | Z |
|---|---|---|---|
| N | 4.0850900000 | -0.1948240000 | -1.6230990000 |
| N | 2.9482470000 | 0.1105640000 | -1.7169730000 |
| N | 1.8099210000 | 0.4128950000 | -1.8111450000 |

**N-centered radical**
Charge: 0
Multiplicity: 2
B3LYP-D3(BJ)/6-31G(d)-SDD SCF energy (au): -636.215951469
B3LYP-D3(BJ)/6-31G(d)-SDD enthalpy (au): -635.915028469
B3LYP-D3(BJ)/6-31G(d)-SDD free energy (au): -635.974910469
B3LYP-D3(BJ)/def2-TZVP/SMD SCF energy (au): -636.478583618
B3LYP-D3(BJ)/def2-TZVP/SMD enthalpy (au): -636.177660618
B3LYP-D3(BJ)/def2-TZVP/SMD free energy (au): -636.237542618
B3LYP-D3(BJ)/def2-TZVP/SMD free energy (quasi-harmonic) (au): -636.235282149

Cartesian coordinates

| ATOM | X | Y | Z |
|---|---|---|---|
| C | -3.4987470000 | -3.4231720000 | -5.9770130000 |
| C | -2.7455640000 | -2.2869970000 | -5.7113140000 |
| C | -3.2356070000 | -1.2673920000 | -4.8785970000 |
| C | -4.5273010000 | -1.3755720000 | -4.3183830000 |
| C | -5.2720290000 | -2.5272760000 | -4.6074520000 |
| C | -4.7723250000 | -3.5438690000 | -5.4173450000 |
| H | -3.1003650000 | -4.2057460000 | -6.6158920000 |
| H | -1.7571300000 | -2.1579640000 | -6.1381000000 |
| H | -6.2709650000 | -2.6181760000 | -4.1875660000 |
| H | -5.3789150000 | -4.4226330000 | -5.6184690000 |
| C | -5.1574150000 | -0.3388280000 | -3.4133790000 |
| C | -4.8709110000 | -0.6010130000 | -1.9261290000 |
| H | -6.2418050000 | -0.3484730000 | -3.5804520000 |
| H | -4.8138550000 | 0.6651130000 | -3.6794000000 |
| H | -5.3532730000 | 0.1581480000 | -1.2999200000 |
| H | -5.2513670000 | -1.5838850000 | -1.6267410000 |
| H | -3.7952010000 | -0.5710750000 | -1.7361790000 |
| C | -2.3158450000 | -0.1045980000 | -4.6881210000 |
| O | -1.2980900000 | 0.0376500000 | -5.3802270000 |
| N | -2.5216870000 | 0.7360210000 | -3.6197140000 |
| C | -2.3252810000 | 2.1794290000 | -3.7112960000 |
| C | -2.7598510000 | 2.7692710000 | -5.0634520000 |
| C | -0.8161750000 | 2.4391230000 | -3.4654520000 |
| H | -2.6080370000 | 3.8539410000 | -5.0662180000 |
| H | -2.1823780000 | 2.3353340000 | -5.8824420000 |
| H | -3.8241540000 | 2.5770550000 | -5.2430490000 |
| H | -0.5015130000 | 1.9901610000 | -2.5186490000 |
| H | -0.2257250000 | 2.0079560000 | -4.2748460000 |

| | | | |
|---|---|---|---|
| H | -0.6455130000 | 3.5207910000 | -3.4164470000 |
| C | -3.1350730000 | 2.8060080000 | -2.5616460000 |
| H | -4.2095770000 | 2.6539510000 | -2.7078120000 |
| H | -2.8521140000 | 2.3580280000 | -1.6046380000 |
| H | -2.9462440000 | 3.8838070000 | -2.5176360000 |

**OH□**
Charge: 0
Multiplicity: 2
B3LYP-D3(BJ)/6-31G(d)-SDD SCF energy (au): -75.7217884823
B3LYP-D3(BJ)/6-31G(d)-SDD enthalpy (au): -75.7102224823
B3LYP-D3(BJ)/6-31G(d)-SDD free energy (au): -75.7304694823
B3LYP-D3(BJ)/def2-TZVP/SMD SCF energy (au): -75.7734433827
B3LYP-D3(BJ)/def2-TZVP/SMD enthalpy (au): -75.7618773827
B3LYP-D3(BJ)/def2-TZVP/SMD free energy (au): -75.7821243827
B3LYP-D3(BJ)/def2-TZVP/SMD free energy (quasi-harmonic) (au): -75.7821239004

Cartesian coordinates

| ATOM | X | Y | Z |
|---|---|---|---|
| O | -0.4595260000 | -1.2255800000 | -2.5970370000 |
| H | 0.1993020000 | -0.8699910000 | -3.2345060000 |

**$^{5}$TS-1'**
Charge: 1
Multiplicity: 5
B3LYP-D3(BJ)/6-31G(d)-SDD SCF energy (au): -1693.77688517
B3LYP-D3(BJ)/6-31G(d)-SDD enthalpy (au): -1693.19952117
B3LYP-D3(BJ)/6-31G(d)-SDD free energy (au): -1693.31489017
B3LYP-D3(BJ)/def2-TZVP/SMD SCF energy (au): -2834.33807910
B3LYP-D3(BJ)/def2-TZVP/SMD enthalpy (au): -2833.76071510
B3LYP-D3(BJ)/def2-TZVP/SMD free energy (au): -2833.87608410
B3LYP-D3(BJ)/def2-TZVP/SMD free energy (quasi-harmonic) (au): -2833.86476393

Cartesian coordinates

| ATOM | X | Y | Z |
|---|---|---|---|
| Fe | 0.0380170000 | -0.6593910000 | -0.8120440000 |
| C | -0.8973410000 | -1.7926800000 | 3.3080600000 |
| N | -1.4780580000 | -0.5417560000 | 3.2073140000 |
| H | -1.9369190000 | -0.0388540000 | 3.9544800000 |
| C | -1.3224110000 | -0.0996510000 | 1.9385680000 |
| H | -1.6756800000 | 0.8499310000 | 1.5700770000 |
| N | -0.6694020000 | -1.0032160000 | 1.2184470000 |
| C | -0.3971780000 | -2.0649130000 | 2.0642200000 |
| H | 0.1321080000 | -2.9367710000 | 1.7146140000 |
| H | -0.8966960000 | -2.3541580000 | 4.2288830000 |
| C | 1.9448700000 | -4.5886280000 | -1.0212690000 |
| N | 2.9345220000 | -3.8207140000 | -0.4377740000 |
| H | 3.8668540000 | -4.1328410000 | -0.2027430000 |
| C | 2.4519200000 | -2.5680910000 | -0.2573410000 |
| H | 3.0090130000 | -1.7546150000 | 0.1801840000 |
| N | 1.2014910000 | -2.4965450000 | -0.6902910000 |
| C | 0.8716670000 | -3.7515480000 | -1.1728490000 |
| H | -0.1033700000 | -3.9472950000 | -1.5915530000 |

| | | | |
|---|---|---|---|
| H | 2.0938290000 | -5.6276010000 | -1.2695950000 |
| O | -1.6133130000 | -1.8123750000 | -1.5531760000 |
| H | -2.0931680000 | -1.0938230000 | -2.0689000000 |
| H | -2.1882650000 | -2.0123400000 | -0.7957600000 |
| C | -3.7360990000 | 3.4139350000 | -5.7544710000 |
| C | -3.0110970000 | 2.5910940000 | -4.8983520000 |
| C | -3.5194250000 | 2.2841570000 | -3.6291820000 |
| C | -4.7799320000 | 2.7474070000 | -3.2106190000 |
| C | -5.4866040000 | 3.5752360000 | -4.0917640000 |
| C | -4.9709140000 | 3.9204360000 | -5.3396480000 |
| H | -3.3459350000 | 3.6522810000 | -6.7387600000 |
| H | -2.0571790000 | 2.1785850000 | -5.2113390000 |
| H | -6.4657610000 | 3.9414880000 | -3.7943910000 |
| H | -5.5417070000 | 4.5671880000 | -5.9989410000 |
| C | -5.4004920000 | 2.3502750000 | -1.8866960000 |
| C | -5.2219110000 | 3.3858610000 | -0.7653060000 |
| H | -4.9978180000 | 1.3852300000 | -1.5606590000 |
| H | -6.4720260000 | 2.1911090000 | -2.0545540000 |
| H | -5.8089760000 | 3.1013480000 | 0.1143680000 |
| H | -4.1728610000 | 3.4557490000 | -0.4644210000 |
| H | -5.5544770000 | 4.3786700000 | -1.0867810000 |
| C | -2.7184410000 | 1.3844210000 | -2.7290260000 |
| O | -2.7985420000 | 0.1608360000 | -2.8306820000 |
| N | -2.0762620000 | 2.0839530000 | -1.7469240000 |
| C | -1.1115890000 | 3.2043430000 | -1.8221020000 |
| C | -1.8931160000 | 4.4942330000 | -2.1899500000 |
| C | -0.0016390000 | 2.9472590000 | -2.8466410000 |
| H | -1.1878510000 | 5.3275230000 | -2.1122290000 |
| H | -2.2826860000 | 4.4644090000 | -3.2061740000 |
| H | -2.7172270000 | 4.6717670000 | -1.4972620000 |
| H | 0.5512810000 | 2.0369640000 | -2.6203280000 |
| H | -0.4086780000 | 2.8657730000 | -3.8577540000 |
| H | 0.7034120000 | 3.7845470000 | -2.8361110000 |
| F | -1.2892960000 | 0.8328620000 | -0.7558350000 |
| C | -0.5219490000 | 3.4068160000 | -0.4125310000 |
| H | -1.3201890000 | 3.4881080000 | 0.3317990000 |
| H | 0.1433480000 | 2.5879110000 | -0.1380420000 |
| H | 0.0478610000 | 4.3404720000 | -0.4069190000 |
| C | 2.5737640000 | 0.8880660000 | -1.0463880000 |
| O | 2.6953170000 | 0.6437160000 | -2.2626320000 |
| O | 1.5919900000 | 0.4647110000 | -0.3076640000 |
| C | 3.5936610000 | 1.7505030000 | -0.3370940000 |
| H | 3.1332620000 | 2.7161720000 | -0.0974270000 |
| H | 3.8889390000 | 1.2927310000 | 0.6117740000 |
| H | 4.4656680000 | 1.9167680000 | -0.9706240000 |
| O | 0.4742390000 | -0.4788880000 | -2.8945180000 |
| H | 0.5640780000 | -1.3450710000 | -3.3237120000 |
| H | 1.4119750000 | -0.0812360000 | -2.8135270000 |

**$^{3}$TS-1'**

Charge: 1

Multiplicity: 3

B3LYP-D3(BJ)/6-31G(d)-SDD SCF energy (au): -1693.75025590

B3LYP-D3(BJ)/6-31G(d)-SDD enthalpy (au): -1693.17163990
B3LYP-D3(BJ)/6-31G(d)-SDD free energy (au): -1693.28401490
B3LYP-D3(BJ)/def2-TZVP/SMD SCF energy (au): -2834.31268992
B3LYP-D3(BJ)/def2-TZVP/SMD enthalpy (au): -2833.73407392
B3LYP-D3(BJ)/def2-TZVP/SMD free energy (au): -2833.84644892
B3LYP-D3(BJ)/def2-TZVP/SMD free energy (quasi-harmonic) (au): -2833.83569902

Cartesian coordinates

| ATOM | X | Y | Z |
|---|---|---|---|
| Fe | 0.3716990000 | -0.6737230000 | -0.9888760000 |
| C | -2.2649440000 | -1.9603890000 | 2.0349940000 |
| N | -2.5001390000 | -0.6003050000 | 1.9699500000 |
| H | -3.1494840000 | -0.0824170000 | 2.5455210000 |
| C | -1.7085720000 | -0.0711640000 | 1.0084620000 |
| H | -1.6790460000 | 0.9700040000 | 0.7310530000 |
| N | -0.9760330000 | -1.0291580000 | 0.4560760000 |
| C | -1.3113190000 | -2.2143710000 | 1.0865150000 |
| H | -0.8372110000 | -3.1455920000 | 0.8199100000 |
| H | -2.7803520000 | -2.6022940000 | 2.7315690000 |
| C | 2.2833480000 | -4.5840100000 | -0.7194940000 |
| N | 2.7933360000 | -3.9194420000 | 0.3794380000 |
| H | 3.4745530000 | -4.2859390000 | 1.0302530000 |
| C | 2.2405100000 | -2.6828730000 | 0.4259730000 |
| H | 2.4551880000 | -1.9345710000 | 1.1732740000 |
| N | 1.4000830000 | -2.5256370000 | -0.5858200000 |
| C | 1.4176680000 | -3.7029940000 | -1.3122210000 |
| H | 0.8146570000 | -3.8212410000 | -2.1996410000 |
| H | 2.5783040000 | -5.5904160000 | -0.9717230000 |
| O | -0.8060030000 | -1.6298120000 | -2.3830200000 |
| H | -1.2738620000 | -0.8743970000 | -2.8449110000 |
| H | -1.5207360000 | -2.1122680000 | -1.9347860000 |
| C | -5.3703060000 | 2.8998160000 | -5.0052250000 |
| C | -4.2403630000 | 2.1955800000 | -4.6107050000 |
| C | -3.8774810000 | 2.1427320000 | -3.2539320000 |
| C | -4.6803600000 | 2.7415350000 | -2.2615980000 |
| C | -5.8222400000 | 3.4320900000 | -2.6878460000 |
| C | -6.1570130000 | 3.5308260000 | -4.0363710000 |
| H | -5.6417850000 | 2.9524560000 | -6.0545810000 |
| H | -3.6174610000 | 1.6912000000 | -5.3426890000 |
| H | -6.4683480000 | 3.8862610000 | -1.9411090000 |
| H | -7.0447060000 | 4.0816990000 | -4.3320490000 |
| C | -4.4250710000 | 2.5996010000 | -0.7732000000 |
| C | -3.8446650000 | 3.8582880000 | -0.1124530000 |
| H | -3.7625920000 | 1.7494510000 | -0.5828800000 |
| H | -5.3838400000 | 2.3593160000 | -0.2967270000 |
| H | -3.7975010000 | 3.7378190000 | 0.9757940000 |
| H | -2.8336630000 | 4.0549140000 | -0.4781020000 |
| H | -4.4626520000 | 4.7372090000 | -0.3239130000 |
| C | -2.6193880000 | 1.4124100000 | -2.9333520000 |
| O | -2.4327330000 | 0.2523380000 | -3.2924490000 |
| N | -1.7489910000 | 2.1934030000 | -2.1751980000 |
| C | -0.8092530000 | 3.1450420000 | -2.8288730000 |
| C | -1.6539100000 | 4.3475410000 | -3.3217800000 |

| | | | |
|---|---|---|---|
| C | -0.0763720000 | 2.4947520000 | -4.0096670000 |
| H | -0.9541920000 | 5.0991590000 | -3.6993870000 |
| H | -2.3323950000 | 4.0726270000 | -4.1299290000 |
| H | -2.2350640000 | 4.7914330000 | -2.5107660000 |
| H | 0.4754980000 | 1.6089710000 | -3.6895180000 |
| H | -0.7757510000 | 2.2083200000 | -4.8026610000 |
| H | 0.6334270000 | 3.2129310000 | -4.4319570000 |
| F | -0.6935920000 | 1.0003840000 | -1.4568230000 |
| C | 0.1738370000 | 3.6402950000 | -1.7591480000 |
| H | -0.3634270000 | 3.9647860000 | -0.8624470000 |
| H | 0.8902190000 | 2.8653860000 | -1.4887400000 |
| H | 0.7306640000 | 4.4946460000 | -2.1553370000 |
| C | 2.3711800000 | 1.1409520000 | 0.1079440000 |
| O | 2.8392590000 | 1.3866230000 | -1.0259120000 |
| O | 1.4352610000 | 0.2819210000 | 0.3564540000 |
| C | 2.8886930000 | 1.9014080000 | 1.3089570000 |
| H | 2.1762680000 | 2.6996400000 | 1.5495240000 |
| H | 2.9569690000 | 1.2478720000 | 2.1823950000 |
| H | 3.8569350000 | 2.3523950000 | 1.0871740000 |
| O | 1.5839200000 | -0.1990700000 | -2.5166850000 |
| H | 2.1310590000 | -0.9588900000 | -2.7741300000 |
| H | 2.2008910000 | 0.4867820000 | -2.0410930000 |

**[5]TS-1**

Charge: 2
Multiplicity: 5
B3LYP-D3(BJ)/6-31G(d)-SDD SCF energy (au): -1691.22827201
B3LYP-D3(BJ)/6-31G(d)-SDD enthalpy (au): -1690.62899401
B3LYP-D3(BJ)/6-31G(d)-SDD free energy (au): -1690.74560901
B3LYP-D3(BJ)/def2-TZVP/SMD SCF energy (au): -2831.89376700
B3LYP-D3(BJ)/def2-TZVP/SMD enthalpy (au): -2831.29448900
B3LYP-D3(BJ)/def2-TZVP/SMD free energy (au): -2831.41110400
B3LYP-D3(BJ)/def2-TZVP/SMD free energy (quasi-harmonic) (au): -2831.39943799

Cartesian coordinates

| ATOM | X | Y | Z |
|---|---|---|---|
| Fe | 0.3653060000 | -0.3631130000 | -0.7281460000 |
| C | 1.1320730000 | 1.3526830000 | 3.1568150000 |
| N | 2.1041450000 | 1.9540980000 | 2.3807180000 |
| H | 2.7876600000 | 2.6271670000 | 2.7045120000 |
| C | 1.9892980000 | 1.4996740000 | 1.1150750000 |
| H | 2.6260290000 | 1.7940040000 | 0.2957600000 |
| N | 0.9850150000 | 0.6308650000 | 1.0370620000 |
| C | 0.4401240000 | 0.5316980000 | 2.3091650000 |
| H | -0.4007900000 | -0.1120230000 | 2.5125430000 |
| C | -3.0231650000 | -2.4910060000 | 0.8748660000 |
| N | -3.5335500000 | -1.2087300000 | 0.8186960000 |
| H | -4.4811940000 | -0.9392520000 | 1.0506640000 |
| C | -2.5616820000 | -0.3766030000 | 0.3868880000 |
| H | -2.6930180000 | 0.6849500000 | 0.2492350000 |
| N | -1.4431730000 | -1.0596910000 | 0.1656450000 |
| C | -1.7212900000 | -2.3852520000 | 0.4680380000 |
| H | -0.9693450000 | -3.1531810000 | 0.3750130000 |

| | | | |
|---|---|---|---|
| O | 2.2607130000 | 0.2999700000 | -1.7173040000 |
| H | 2.8424330000 | -0.4309390000 | -1.9865930000 |
| H | 2.0432350000 | 0.7978810000 | -2.5231880000 |
| H | -3.6165110000 | -3.3340670000 | 1.1927330000 |
| H | 1.0245140000 | 1.5614710000 | 4.2097650000 |
| C | 2.7447980000 | -4.0196890000 | -0.5899790000 |
| N | 2.7415810000 | -3.6448910000 | 0.7373210000 |
| H | 3.2125870000 | -4.1314180000 | 1.4900490000 |
| C | 2.0112610000 | -2.5135250000 | 0.8641360000 |
| H | 1.8515350000 | -1.9961480000 | 1.7979480000 |
| N | 1.5379250000 | -2.1410320000 | -0.3206610000 |
| C | 1.9935660000 | -3.0769110000 | -1.2394720000 |
| H | 1.7503010000 | -3.0065990000 | -2.2889870000 |
| H | 3.2651740000 | -4.8971690000 | -0.9413320000 |
| O | -0.2273290000 | -1.2678110000 | -2.5808560000 |
| H | -1.0007880000 | -0.6800630000 | -2.8313250000 |
| H | -0.6014870000 | -2.1607050000 | -2.4945610000 |
| C | -6.0296960000 | 2.2351680000 | -3.7045310000 |
| C | -4.7731640000 | 1.6619940000 | -3.6094490000 |
| C | -3.8790040000 | 2.0619960000 | -2.5901110000 |
| C | -4.2593030000 | 3.0295230000 | -1.6241360000 |
| C | -5.5441980000 | 3.5747760000 | -1.7417510000 |
| C | -6.4113990000 | 3.2049090000 | -2.7671610000 |
| H | -6.7106720000 | 1.9341520000 | -4.4935210000 |
| H | -4.4532430000 | 0.9065980000 | -4.3194650000 |
| H | -5.8729070000 | 4.3005010000 | -1.0032580000 |
| H | -7.3938220000 | 3.6624120000 | -2.8309160000 |
| C | -3.4169930000 | 3.4520850000 | -0.4356320000 |
| C | -2.8818050000 | 4.8885050000 | -0.5439510000 |
| H | -2.5704520000 | 2.7760080000 | -0.2932890000 |
| H | -4.0477180000 | 3.3802860000 | 0.4597480000 |
| H | -2.3611710000 | 5.1726320000 | 0.3757810000 |
| H | -2.1775470000 | 4.9781560000 | -1.3757230000 |
| H | -3.6922740000 | 5.6053880000 | -0.7071750000 |
| C | -2.5534780000 | 1.4107210000 | -2.6372300000 |
| O | -2.4019360000 | 0.2036040000 | -2.8364740000 |
| N | -1.5324470000 | 2.3414400000 | -2.4486050000 |
| C | -0.8222550000 | 2.9216660000 | -3.5962530000 |
| C | -1.8530990000 | 3.8722010000 | -4.2840130000 |
| C | -0.3563950000 | 1.8670290000 | -4.6129210000 |
| H | -1.3111640000 | 4.4119810000 | -5.0659860000 |
| H | -2.6710180000 | 3.3143340000 | -4.7428360000 |
| H | -2.2653400000 | 4.5954220000 | -3.5788080000 |
| H | 0.3402500000 | 1.1609010000 | -4.1540760000 |
| H | -1.1942210000 | 1.3000700000 | -5.0274900000 |
| H | 0.1534390000 | 2.3695090000 | -5.4395860000 |
| F | -0.1863890000 | 1.2863110000 | -1.6808400000 |
| C | 0.3430280000 | 3.7629420000 | -3.0597360000 |
| H | -0.0117110000 | 4.4915410000 | -2.3259610000 |
| H | 1.0923060000 | 3.1307150000 | -2.5792700000 |
| H | 0.8151920000 | 4.3016910000 | -3.8857640000 |

**$^{3}$TS-1**

Charge: 2
Multiplicity: 3
B3LYP-D3(BJ)/6-31G(d)-SDD SCF energy (au): -1691.20095744
B3LYP-D3(BJ)/6-31G(d)-SDD enthalpy (au): -1690.60052244
B3LYP-D3(BJ)/6-31G(d)-SDD free energy (au): -1690.71334744
B3LYP-D3(BJ)/def2-TZVP/SMD SCF energy (au): -2831.86896007
B3LYP-D3(BJ)/def2-TZVP/SMD enthalpy (au): -2831.26852507
B3LYP-D3(BJ)/def2-TZVP/SMD free energy (au): -2831.38135007
B3LYP-D3(BJ)/def2-TZVP/SMD free energy (quasi-harmonic) (au): -2831.37042963

Cartesian coordinates

| ATOM | X | Y | Z |
|---|---|---|---|
| Fe | 0.3726390000 | -0.4260220000 | -0.6901880000 |
| C | 1.5961460000 | 1.1984960000 | 2.9627970000 |
| N | 2.2619490000 | 1.9986320000 | 2.0548450000 |
| H | 2.8641640000 | 2.7795540000 | 2.2840380000 |
| C | 1.9742760000 | 1.5729270000 | 0.8075100000 |
| H | 2.3579860000 | 2.0154900000 | -0.0970900000 |
| N | 1.1509920000 | 0.5291150000 | 0.8698020000 |
| C | 0.9027200000 | 0.2879330000 | 2.2139600000 |
| H | 0.2485780000 | -0.5069080000 | 2.5343680000 |
| C | -2.6166150000 | -2.3720990000 | 1.4875150000 |
| N | -3.1466090000 | -1.1085550000 | 1.3241090000 |
| H | -4.0520460000 | -0.8002760000 | 1.6549950000 |
| C | -2.2657040000 | -0.3528100000 | 0.6332340000 |
| H | -2.4180260000 | 0.6781180000 | 0.3595210000 |
| N | -1.1873500000 | -1.0760840000 | 0.3482840000 |
| C | -1.3919580000 | -2.3423020000 | 0.8766280000 |
| H | -0.6542620000 | -3.1219830000 | 0.7761980000 |
| O | 1.9532040000 | 0.2606320000 | -1.8496260000 |
| H | 2.5826350000 | -0.4793620000 | -1.9260300000 |
| H | 1.6451830000 | 0.4489180000 | -2.7543970000 |
| H | -3.1439430000 | -3.1573430000 | 2.0062660000 |
| H | 1.6716670000 | 1.3486700000 | 4.0284110000 |
| C | 2.5490950000 | -4.2350050000 | -0.7439630000 |
| N | 2.9354450000 | -3.7241250000 | 0.4774690000 |
| H | 3.5591060000 | -4.1745460000 | 1.1359250000 |
| C | 2.3522090000 | -2.5145680000 | 0.6424440000 |
| H | 2.4848510000 | -1.8932780000 | 1.5149890000 |
| N | 1.5999730000 | -2.2212730000 | -0.4141020000 |
| C | 1.7181050000 | -3.2928910000 | -1.2890610000 |
| H | 1.2188310000 | -3.3036010000 | -2.2459050000 |
| H | 2.8936810000 | -5.1914900000 | -1.1055130000 |
| O | -0.4509460000 | -1.2530330000 | -2.3616640000 |
| H | -1.2217230000 | -0.6338340000 | -2.5760750000 |
| H | -0.8743530000 | -2.1067760000 | -2.1675560000 |
| C | -5.8274920000 | 2.4521340000 | -4.4184280000 |
| C | -4.6626010000 | 1.7978800000 | -4.0520700000 |
| C | -4.0041010000 | 2.1366200000 | -2.8495280000 |
| C | -4.5419580000 | 3.1081350000 | -1.9699030000 |
| C | -5.7283230000 | 3.7383850000 | -2.3648060000 |
| C | -6.3544550000 | 3.4365050000 | -3.5721440000 |
| H | -6.3278270000 | 2.1997780000 | -5.3473070000 |

```
H     -4.2352270000    1.0290200000   -4.6873800000
H     -6.1784000000    4.4701820000   -1.6999280000
H     -7.2676550000    3.9546970000   -3.8477610000
C     -3.9816600000    3.4292470000   -0.5978620000
C     -3.3342600000    4.8178460000   -0.4922760000
H     -3.2519850000    2.6737120000   -0.2932390000
H     -4.8131350000    3.3738000000    0.1157100000
H     -3.0675640000    5.0374110000    0.5462850000
H     -2.4233540000    4.8690210000   -1.0952120000
H     -4.0151790000    5.6024960000   -0.8364600000
C     -2.7293350000    1.4319280000   -2.6108810000
O     -2.5926700000    0.2141580000   -2.7463130000
N     -1.7164010000    2.3236950000   -2.2422100000
C     -0.8540700000    2.9337290000   -3.2767860000
C     -1.7133450000    4.0300540000   -3.9710380000
C     -0.3858840000    1.9198900000   -4.3311820000
H     -1.0549270000    4.5606760000   -4.6652250000
H     -2.5423040000    3.6003240000   -4.5344810000
H     -2.1080570000    4.7453290000   -3.2483110000
H      0.1366760000    1.0768790000   -3.8694000000
H     -1.2224940000    1.5120590000   -4.9056610000
H      0.2960930000    2.4099010000   -5.0314930000
F     -0.5894300000    1.2236090000   -1.3409820000
C      0.3237470000    3.6134250000   -2.5651370000
H     -0.0379040000    4.2731870000   -1.7718340000
H      0.9937550000    2.8741390000   -2.1225830000
H      0.8871710000    4.2116370000   -3.2864640000
```

**[5]TS-2'**

```
Charge: 1
Multiplicity: 5
B3LYP-D3(BJ)/6-31G(d)-SDD SCF energy (au):              -1693.81375146
B3LYP-D3(BJ)/6-31G(d)-SDD enthalpy (au):                -1693.24029146
B3LYP-D3(BJ)/6-31G(d)-SDD free energy (au):             -1693.35562146
B3LYP-D3(BJ)/def2-TZVP/SMD SCF energy (au):             -2834.37665990
B3LYP-D3(BJ)/def2-TZVP/SMD enthalpy (au):               -2833.80319990
B3LYP-D3(BJ)/def2-TZVP/SMD free energy (au):            -2833.91852990
B3LYP-D3(BJ)/def2-TZVP/SMD free energy (quasi-harmonic) (au): -2833.90629244

Cartesian coordinates
ATOM    X               Y               Z
Fe    -0.1453580000   -0.1141520000   -0.1035820000
C     -2.8354440000   -3.1324240000    1.5263070000
N     -3.2083120000   -1.9679860000    2.1744760000
H     -3.9113530000   -1.8897220000    2.8966770000
C     -2.4674570000   -0.9492730000    1.6872330000
H     -2.5322950000    0.0798600000    2.0040740000
N     -1.6360870000   -1.4048080000    0.7566440000
C     -1.8540370000   -2.7664730000    0.6472450000
H     -1.2970250000   -3.3691420000   -0.0517440000
H     -3.2927420000   -4.0843190000    1.7450320000
C      1.2785020000   -3.4689130000   -2.4663060000
N      2.1406740000   -3.4727480000   -1.3840520000
```

| | | | |
|---|---|---|---|
| H | 2.9299630000 | -4.0910380000 | -1.2559370000 |
| C | 1.7665270000 | -2.4894400000 | -0.5353420000 |
| H | 2.2586000000 | -2.2494280000 | 0.3938250000 |
| N | 0.7037440000 | -1.8606910000 | -1.0185670000 |
| C | 0.3886680000 | -2.4578730000 | -2.2259800000 |
| H | -0.4359280000 | -2.1003300000 | -2.8210980000 |
| H | 1.3819420000 | -4.1618960000 | -3.2861340000 |
| O | -1.4189720000 | 0.0004600000 | -1.8856450000 |
| H | -1.2152550000 | 0.9025220000 | -2.2765740000 |
| H | -2.3124690000 | 0.0902890000 | -1.5132670000 |
| F | -1.1127310000 | 1.2995900000 | 0.5478490000 |
| C | -1.3261250000 | 3.4714080000 | -2.5637790000 |
| O | -0.6261730000 | 2.4188970000 | -2.6092500000 |
| N | -0.8199000000 | 4.6749330000 | -2.2776560000 |
| H | -3.1667890000 | 5.4968250000 | -0.7424570000 |
| C | 0.3669090000 | 5.0081840000 | -1.4705490000 |
| C | 0.3908570000 | 6.5397710000 | -1.3387840000 |
| C | 0.2827470000 | 4.3558020000 | -0.0815930000 |
| C | 1.6261240000 | 4.5627030000 | -2.2470460000 |
| H | 0.4227420000 | 7.0167290000 | -2.3231980000 |
| H | -0.4935110000 | 6.9017580000 | -0.8026250000 |
| H | 1.2748410000 | 6.8477270000 | -0.7731460000 |
| H | 0.1025780000 | 3.2824530000 | -0.1397810000 |
| H | 1.2138090000 | 4.5164500000 | 0.4706240000 |
| H | -0.5374420000 | 4.7973450000 | 0.4954390000 |
| H | 2.5117850000 | 4.9161710000 | -1.7092250000 |
| H | 1.6897000000 | 3.4790920000 | -2.3429630000 |
| H | 1.6295890000 | 4.9975140000 | -3.2505920000 |
| C | -2.7728670000 | 3.4215450000 | -2.9321830000 |
| C | -3.2483290000 | 2.3147830000 | -3.6470970000 |
| C | -3.6475430000 | 4.4776190000 | -2.5977040000 |
| C | -4.5847060000 | 2.2456000000 | -4.0297010000 |
| H | -2.5555510000 | 1.5322370000 | -3.9361620000 |
| C | -4.9849840000 | 4.4002430000 | -3.0022500000 |
| C | -5.4534710000 | 3.2898380000 | -3.7031790000 |
| H | -4.9439340000 | 1.3929310000 | -4.5973890000 |
| H | -5.6664880000 | 5.2080820000 | -2.7557980000 |
| H | -6.4963520000 | 3.2438740000 | -4.0020910000 |
| C | -3.1187000000 | 5.6479670000 | -1.8278000000 |
| C | -3.5471950000 | 7.0347110000 | -2.2593000000 |
| H | -4.6120530000 | 7.1983490000 | -2.0437820000 |
| H | -2.9871790000 | 7.8033310000 | -1.7180050000 |
| H | -3.3970350000 | 7.1802590000 | -3.3332210000 |
| H | -1.9109500000 | 5.5064410000 | -2.0423210000 |
| C | 1.9660950000 | 0.7274170000 | 1.7309870000 |
| O | 2.3418140000 | 1.6578180000 | 1.0067380000 |
| O | 1.1664110000 | -0.2348430000 | 1.3254450000 |
| C | 2.4071460000 | 0.6191640000 | 3.1704370000 |
| H | 1.5411290000 | 0.7852500000 | 3.8211830000 |
| H | 2.7799750000 | -0.3884970000 | 3.3775480000 |
| H | 3.1758910000 | 1.3608010000 | 3.3900080000 |
| O | 1.1714370000 | 1.0345420000 | -1.2223640000 |
| H | 1.7260300000 | 1.4397040000 | -0.4786620000 |

H      0.6900830000    1.7513320000   -1.7041810000

**[3]TS-2'**

Charge: 1
Multiplicity: 3
B3LYP-D3(BJ)/6-31G(d)-SDD SCF energy (au): -1693.78985435
B3LYP-D3(BJ)/6-31G(d)-SDD enthalpy (au): -1693.21719735
B3LYP-D3(BJ)/6-31G(d)-SDD free energy (au): -1693.33048435
B3LYP-D3(BJ)/def2-TZVP/SMD SCF energy (au): -2834.35668777
B3LYP-D3(BJ)/def2-TZVP/SMD enthalpy (au): -2833.78403077
B3LYP-D3(BJ)/def2-TZVP/SMD free energy (au): -2833.89731777
B3LYP-D3(BJ)/def2-TZVP/SMD free energy (quasi-harmonic) (au): -2833.88591142

Cartesian coordinates

| ATOM | X | Y | Z |
|---|---|---|---|
| Fe | -0.2685590000 | -0.1846270000 | -0.0116610000 |
| C | -2.8721200000 | -3.0376350000 | 1.5242460000 |
| N | -2.2511550000 | -2.5377310000 | 2.6558320000 |
| H | -2.4433260000 | -2.8142820000 | 3.6090020000 |
| C | -1.3439020000 | -1.6115250000 | 2.2816460000 |
| H | -0.6922140000 | -1.0554050000 | 2.9361680000 |
| N | -1.3606580000 | -1.4983540000 | 0.9603220000 |
| C | -2.3060930000 | -2.3790610000 | 0.4685190000 |
| H | -2.4911780000 | -2.4743920000 | -0.5894720000 |
| H | -3.6365110000 | -3.7965540000 | 1.5738180000 |
| C | 1.2383330000 | -3.2315800000 | -2.4941040000 |
| N | 2.1049530000 | -3.2630100000 | -1.4160780000 |
| H | 2.9168090000 | -3.8581340000 | -1.3244580000 |
| C | 1.7002090000 | -2.3487070000 | -0.5084670000 |
| H | 2.1815310000 | -2.1320500000 | 0.4320930000 |
| N | 0.6114530000 | -1.7403520000 | -0.9587680000 |
| C | 0.3082750000 | -2.2740160000 | -2.1972140000 |
| H | -0.5374550000 | -1.9191220000 | -2.7632810000 |
| H | 1.3632520000 | -3.8727960000 | -3.3520230000 |
| O | -1.8059950000 | -0.0941700000 | -1.7113370000 |
| H | -1.5446920000 | 0.7740400000 | -2.1249390000 |
| H | -2.6006840000 | 0.1351400000 | -1.2002520000 |
| F | -1.2070850000 | 1.1445570000 | 0.6851780000 |
| C | -1.4600000000 | 3.4106140000 | -2.5411630000 |
| O | -0.8567790000 | 2.3022450000 | -2.4506460000 |
| N | -0.8787510000 | 4.5827600000 | -2.2687840000 |
| H | -3.3000300000 | 5.7472560000 | -1.0969550000 |
| C | 0.2501250000 | 4.8700830000 | -1.3669660000 |
| C | 0.3584270000 | 6.4012900000 | -1.2734040000 |
| C | 0.0067740000 | 4.2722220000 | 0.0280670000 |
| C | 1.5407990000 | 4.3290960000 | -2.0215100000 |
| H | 0.4968520000 | 6.8450990000 | -2.2641860000 |
| H | -0.5405120000 | 6.8317710000 | -0.8183960000 |
| H | 1.2136270000 | 6.6722660000 | -0.6477970000 |
| H | -0.2349550000 | 3.2101440000 | -0.0092690000 |
| H | 0.8977370000 | 4.3931800000 | 0.6517660000 |
| H | -0.8276250000 | 4.7863270000 | 0.5182320000 |
| H | 2.3953070000 | 4.6065320000 | -1.3960380000 |

| | | | |
|---|---|---|---|
| H | 1.5287320000 | 3.2446950000 | -2.1261870000 |
| H | 1.6751270000 | 4.7675700000 | -3.0145120000 |
| C | -2.8591710000 | 3.4582540000 | -3.0620340000 |
| C | -3.3657460000 | 2.3383990000 | -3.7334940000 |
| C | -3.6560400000 | 4.6141660000 | -2.9133260000 |
| C | -4.6535570000 | 2.3563010000 | -4.2602350000 |
| H | -2.7327070000 | 1.4687490000 | -3.8675250000 |
| C | -4.9431450000 | 4.6224810000 | -3.4622420000 |
| C | -5.4425030000 | 3.5008490000 | -4.1225870000 |
| H | -5.0356620000 | 1.4896270000 | -4.7904710000 |
| H | -5.5635630000 | 5.5068690000 | -3.3586370000 |
| H | -6.4471870000 | 3.5223940000 | -4.5339350000 |
| C | -3.1048030000 | 5.7947860000 | -2.1753290000 |
| C | -3.3332160000 | 7.1709600000 | -2.7646330000 |
| H | -4.3940690000 | 7.4499550000 | -2.7030250000 |
| H | -2.7676780000 | 7.9302630000 | -2.2159550000 |
| H | -3.0391840000 | 7.2051070000 | -3.8179550000 |
| H | -1.9030340000 | 5.5257910000 | -2.2213910000 |
| C | 1.9304260000 | 0.7091960000 | 1.8101710000 |
| O | 2.1829670000 | 1.6354560000 | 0.9985970000 |
| O | 1.1449110000 | -0.2823960000 | 1.5663810000 |
| C | 2.5612980000 | 0.7527980000 | 3.1852020000 |
| H | 1.8217750000 | 1.1370120000 | 3.8983820000 |
| H | 2.8456600000 | -0.2514940000 | 3.5106310000 |
| H | 3.4269130000 | 1.4168410000 | 3.1916070000 |
| O | 0.8952150000 | 0.9651770000 | -0.9959180000 |
| H | 1.4884370000 | 1.3870170000 | -0.2120370000 |
| H | 0.3691850000 | 1.6483860000 | -1.4895520000 |

**$^{5}$TS-2**

Charge: 1
Multiplicity: 5
B3LYP-D3(BJ)/6-31G(d)-SDD SCF energy (au): -1690.94148217
B3LYP-D3(BJ)/6-31G(d)-SDD enthalpy (au): -1690.36062717
B3LYP-D3(BJ)/6-31G(d)-SDD free energy (au): -1690.46996717
B3LYP-D3(BJ)/def2-TZVP/SMD SCF energy (au): -2831.49602977
B3LYP-D3(BJ)/def2-TZVP/SMD enthalpy (au): -2830.91517477
B3LYP-D3(BJ)/def2-TZVP/SMD free energy (au): -2831.02451477
B3LYP-D3(BJ)/def2-TZVP/SMD free energy (quasi-harmonic) (au): -2831.01585260

Cartesian coordinates

| ATOM | X | Y | Z |
|---|---|---|---|
| Fe | 0.1277060000 | 0.3699270000 | -0.4839610000 |
| C | 0.8407880000 | -0.0826320000 | 3.7897910000 |
| N | 1.9977410000 | 0.5069900000 | 3.3116660000 |
| H | 2.7888880000 | 0.7993020000 | 3.8686870000 |
| C | 1.8852260000 | 0.6474620000 | 1.9703570000 |
| H | 2.6267920000 | 1.0789120000 | 1.3146630000 |
| N | 0.7145390000 | 0.1730400000 | 1.5690490000 |
| C | 0.0490570000 | -0.2828020000 | 2.6907770000 |
| H | -0.9402620000 | -0.7071580000 | 2.6196120000 |
| C | -3.6885050000 | -1.9344750000 | 0.0873620000 |
| N | -4.0696380000 | -0.6063660000 | 0.1540450000 |

| | | | |
|---|---|---|---|
| H | -5.0149550000 | -0.2548920000 | 0.2134200000 |
| C | -2.9612140000 | 0.1634350000 | 0.0596200000 |
| H | -2.9549710000 | 1.2406740000 | 0.0559210000 |
| N | -1.8839130000 | -0.6033680000 | -0.0616910000 |
| C | -2.3256860000 | -1.9153310000 | -0.0473840000 |
| H | -1.6359620000 | -2.7387010000 | -0.1439820000 |
| O | 1.7637740000 | 1.0371640000 | -1.0260470000 |
| H | 1.6826430000 | 1.8316080000 | -1.5797950000 |
| H | -4.4011240000 | -2.7424490000 | 0.1406800000 |
| H | 0.6921040000 | -0.2909340000 | 4.8375590000 |
| C | 1.6985300000 | -3.4629500000 | -1.8436900000 |
| N | 1.7830550000 | -3.6571510000 | -0.4768220000 |
| H | 2.1882800000 | -4.4594150000 | -0.0152730000 |
| C | 1.2542500000 | -2.5717920000 | 0.1384800000 |
| H | 1.1969170000 | -2.4502990000 | 1.2089580000 |
| N | 0.8292770000 | -1.7006900000 | -0.7642170000 |
| C | 1.1041720000 | -2.2413190000 | -2.0066960000 |
| H | 0.8573440000 | -1.7016710000 | -2.9063320000 |
| H | 2.0658380000 | -4.1896510000 | -2.5508590000 |
| O | -0.5896980000 | 0.2168430000 | -2.5099910000 |
| H | -0.5273210000 | 1.1649380000 | -2.8525990000 |
| H | -1.5424230000 | 0.0466100000 | -2.4257000000 |
| F | -0.8295020000 | 1.9343620000 | -0.1915360000 |
| C | -1.3184450000 | 3.4750290000 | -2.5492890000 |
| O | -0.4429070000 | 2.7697370000 | -3.1042780000 |
| N | -1.1215860000 | 4.7516760000 | -2.1808490000 |
| H | -3.7746220000 | 5.5785120000 | -0.6623480000 |
| C | 0.1518740000 | 5.4254880000 | -1.8839710000 |
| C | -0.2040530000 | 6.8552070000 | -1.4466330000 |
| C | 0.9266670000 | 4.7093580000 | -0.7678830000 |
| C | 0.9764130000 | 5.4804960000 | -3.1883880000 |
| H | -0.7892970000 | 7.3629410000 | -2.2192560000 |
| H | -0.7850800000 | 6.8496180000 | -0.5170440000 |
| H | 0.7100600000 | 7.4289710000 | -1.2682470000 |
| H | 1.1939910000 | 3.6919610000 | -1.0543940000 |
| H | 1.8474140000 | 5.2605910000 | -0.5481750000 |
| H | 0.3308380000 | 4.6508330000 | 0.1472420000 |
| H | 1.8880010000 | 6.0597280000 | -3.0053920000 |
| H | 1.2454570000 | 4.4774470000 | -3.5211130000 |
| H | 0.4054020000 | 5.9721390000 | -3.9815270000 |
| C | -2.7189330000 | 2.9639690000 | -2.3534930000 |
| C | -3.1718360000 | 1.9007080000 | -3.1435260000 |
| C | -3.5775470000 | 3.5454080000 | -1.4020400000 |
| C | -4.4537840000 | 1.3833160000 | -2.9685750000 |
| H | -2.5263420000 | 1.5147790000 | -3.9259310000 |
| C | -4.8649750000 | 3.0196340000 | -1.2344390000 |
| C | -5.2990180000 | 1.9388120000 | -2.0035890000 |
| H | -4.8024980000 | 0.5709710000 | -3.5989890000 |
| H | -5.5335340000 | 3.4739160000 | -0.5078090000 |
| H | -6.3082370000 | 1.5557930000 | -1.8772050000 |
| C | -3.0999220000 | 4.7182360000 | -0.6002930000 |
| C | -2.6126080000 | 4.4225370000 | 0.8068530000 |
| H | -1.8673990000 | 3.6231310000 | 0.7947120000 |

| | | | |
|---|---|---|---|
| H | -2.1844350000 | 5.3163230000 | 1.2722290000 |
| H | -3.4565860000 | 4.0985960000 | 1.4333260000 |
| H | -2.1165200000 | 5.0495520000 | -1.2953840000 |

**[3]TS-2**
Charge: 1
Multiplicity: 3
B3LYP-D3(BJ)/6-31G(d)-SDD SCF energy (au): -1690.92552005
B3LYP-D3(BJ)/6-31G(d)-SDD enthalpy (au): -1690.34233105
B3LYP-D3(BJ)/6-31G(d)-SDD free energy (au): -1690.45304105
B3LYP-D3(BJ)/def2-TZVP/SMD SCF energy (au): -2831.47118872
B3LYP-D3(BJ)/def2-TZVP/SMD enthalpy (au): -2830.88799972
B3LYP-D3(BJ)/def2-TZVP/SMD free energy (au): -2830.99870972
B3LYP-D3(BJ)/def2-TZVP/SMD free energy (quasi-harmonic) (au): -2830.98821467

Cartesian coordinates

| ATOM | X | Y | Z |
|---|---|---|---|
| Fe | -0.1724940000 | 0.2513850000 | -0.5693510000 |
| C | 0.8357680000 | 0.3247600000 | 3.7134860000 |
| N | 1.8244350000 | 1.0810420000 | 3.1095960000 |
| H | 2.5528730000 | 1.5944100000 | 3.5860660000 |
| C | 1.6404060000 | 1.0320400000 | 1.7677130000 |
| H | 2.2471720000 | 1.5335320000 | 1.0291030000 |
| N | 0.5887510000 | 0.2798020000 | 1.4838730000 |
| C | 0.0731290000 | -0.1645690000 | 2.6861780000 |
| H | -0.8041880000 | -0.7925590000 | 2.7229780000 |
| C | -3.5317270000 | -2.1591400000 | 0.4694090000 |
| N | -4.0566400000 | -0.8845300000 | 0.3561610000 |
| H | -5.0267110000 | -0.6264020000 | 0.4718940000 |
| C | -3.0558260000 | -0.0281520000 | 0.0535150000 |
| H | -3.1622230000 | 1.0347680000 | -0.0926860000 |
| N | -1.9110310000 | -0.6976010000 | -0.0346300000 |
| C | -2.1921210000 | -2.0275590000 | 0.2216760000 |
| H | -1.4201170000 | -2.7789500000 | 0.1939840000 |
| O | 1.3255050000 | 1.0561590000 | -1.1478460000 |
| H | 1.0393740000 | 1.7120630000 | -1.8140730000 |
| H | -4.1421640000 | -3.0162640000 | 0.7056560000 |
| H | 0.7694380000 | 0.2139950000 | 4.7844050000 |
| C | 1.6731130000 | -3.3494820000 | -1.9387840000 |
| N | 2.0522710000 | -3.3498610000 | -0.6090920000 |
| H | 2.6725120000 | -4.0146850000 | -0.1684760000 |
| C | 1.4772540000 | -2.2852030000 | -0.0014240000 |
| H | 1.6118720000 | -2.0267820000 | 1.0363380000 |
| N | 0.7431050000 | -1.6125620000 | -0.8756180000 |
| C | 0.8592150000 | -2.2606490000 | -2.0918000000 |
| H | 0.3611350000 | -1.8771370000 | -2.9666450000 |
| H | 2.0138680000 | -4.0998820000 | -2.6343070000 |
| O | -0.8891340000 | 0.0843670000 | -2.6580360000 |
| H | -0.7064870000 | 1.0142940000 | -2.9818190000 |
| H | -1.8568280000 | 0.0436780000 | -2.5929480000 |
| F | -1.0860910000 | 1.8139000000 | -0.3061220000 |
| C | -1.2507920000 | 3.4174590000 | -2.5554870000 |
| O | -0.4068650000 | 2.6651060000 | -3.1031250000 |

| | | | |
|---|---|---|---|
| N | -0.9771630000 | 4.6512400000 | -2.1065820000 |
| H | -3.6304310000 | 5.6004240000 | -0.7036330000 |
| C | 0.3139520000 | 5.2371360000 | -1.7150170000 |
| C | 0.0257280000 | 6.6982500000 | -1.3340310000 |
| C | 0.9341590000 | 4.4934080000 | -0.5217070000 |
| C | 1.2521570000 | 5.2076390000 | -2.9410240000 |
| H | -0.4435830000 | 7.2315540000 | -2.1661530000 |
| H | -0.6410850000 | 6.7539270000 | -0.4655960000 |
| H | 0.9587090000 | 7.2073230000 | -1.0747430000 |
| H | 1.1348270000 | 3.4473210000 | -0.7536450000 |
| H | 1.8769840000 | 4.9774220000 | -0.2428940000 |
| H | 0.2644910000 | 4.5231190000 | 0.3431450000 |
| H | 2.1753410000 | 5.7401610000 | -2.6880700000 |
| H | 1.4926020000 | 4.1840880000 | -3.2287580000 |
| H | 0.7842660000 | 5.7092920000 | -3.7932700000 |
| C | -2.6928740000 | 3.0050790000 | -2.4476730000 |
| C | -3.1880810000 | 2.0193940000 | -3.3093510000 |
| C | -3.5380200000 | 3.5748160000 | -1.4779600000 |
| C | -4.4934280000 | 1.5495050000 | -3.1745390000 |
| H | -2.5523800000 | 1.6497650000 | -4.1073600000 |
| C | -4.8462740000 | 3.0928970000 | -1.3461300000 |
| C | -5.3189540000 | 2.0758060000 | -2.1769010000 |
| H | -4.8734140000 | 0.7968160000 | -3.8588010000 |
| H | -5.5013220000 | 3.5321310000 | -0.5982330000 |
| H | -6.3410420000 | 1.7227070000 | -2.0719520000 |
| C | -3.0267960000 | 4.6905240000 | -0.6163270000 |
| C | -2.6602250000 | 4.3357090000 | 0.8126400000 |
| H | -1.9744210000 | 3.4854660000 | 0.8303640000 |
| H | -2.2050280000 | 5.1882560000 | 1.3271470000 |
| H | -3.5662440000 | 4.0605420000 | 1.3723630000 |
| H | -1.9842750000 | 4.9700180000 | -1.2429970000 |

**$^5$TS-3'**

Charge: 1
Multiplicity: 5
B3LYP-D3(BJ)/6-31G(d)-SDD SCF energy (au): -1693.84972847
B3LYP-D3(BJ)/6-31G(d)-SDD enthalpy (au): -1693.27195547
B3LYP-D3(BJ)/6-31G(d)-SDD free energy (au): -1693.38574947
B3LYP-D3(BJ)/def2-TZVP/SMD SCF energy (au): -2834.41477160
B3LYP-D3(BJ)/def2-TZVP/SMD enthalpy (au): -2833.83699860
B3LYP-D3(BJ)/def2-TZVP/SMD free energy (au): -2833.95079260
B3LYP-D3(BJ)/def2-TZVP/SMD free energy (quasi-harmonic) (au): -2833.94062613

Cartesian coordinates

| ATOM | X | Y | Z |
|---|---|---|---|
| Fe | -0.6607560000 | 0.5403190000 | -1.0337070000 |
| C | -2.3643250000 | -2.9390010000 | 0.9697140000 |
| N | -3.1491910000 | -1.8894820000 | 1.4142350000 |
| H | -3.9020410000 | -1.9546110000 | 2.0851480000 |
| C | -2.7318490000 | -0.7566440000 | 0.8036870000 |
| H | -3.1550740000 | 0.2244540000 | 0.9526050000 |
| N | -1.7195970000 | -1.0281780000 | -0.0081920000 |
| C | -1.4788480000 | -2.3859650000 | 0.0855650000 |

| | | | |
|---|---|---|---|
| H | -0.6976920000 | -2.8570210000 | -0.4899310000 |
| H | -2.5090440000 | -3.9507110000 | 1.3140760000 |
| C | 2.1840740000 | -2.5138040000 | -2.3702270000 |
| N | 2.8050010000 | -1.9934630000 | -1.2490080000 |
| H | 3.6874680000 | -2.2996640000 | -0.8630090000 |
| C | 2.0298540000 | -0.9997660000 | -0.7542050000 |
| H | 2.2498680000 | -0.4189670000 | 0.1279370000 |
| N | 0.9447130000 | -0.8569820000 | -1.5029450000 |
| C | 1.0266820000 | -1.7970840000 | -2.5159300000 |
| H | 0.2472820000 | -1.8806050000 | -3.2565000000 |
| H | 2.6142170000 | -3.3195160000 | -2.9439240000 |
| O | -1.5716050000 | -0.1954140000 | -2.8520550000 |
| H | -1.4080450000 | 0.4032000000 | -3.6471620000 |
| H | -2.5290100000 | -0.1503990000 | -2.6930290000 |
| F | -2.0936460000 | 1.6868480000 | -0.7027840000 |
| C | -1.7635950000 | 2.5541820000 | -4.5586550000 |
| O | -1.1445390000 | 1.5045270000 | -4.8242080000 |
| N | -1.3368320000 | 3.7771490000 | -4.9160660000 |
| H | -1.9759610000 | 4.5407450000 | -4.7371430000 |
| C | -0.1350030000 | 4.0986800000 | -5.7324250000 |
| C | -0.0812320000 | 5.6291330000 | -5.8146150000 |
| C | 1.1309520000 | 3.5657530000 | -5.0460690000 |
| C | -0.2945980000 | 3.4889080000 | -7.1339490000 |
| H | -0.9831240000 | 6.0330940000 | -6.2902620000 |
| H | 0.0223800000 | 6.0752160000 | -4.8187810000 |
| H | 0.7778200000 | 5.9398940000 | -6.4152300000 |
| H | 1.1110900000 | 2.4752470000 | -5.0006060000 |
| H | 2.0108900000 | 3.8759210000 | -5.6184930000 |
| H | 1.2168120000 | 3.9670230000 | -4.0316820000 |
| H | 0.5752660000 | 3.7385440000 | -7.7501310000 |
| H | -0.3771370000 | 2.4012140000 | -7.0736580000 |
| H | -1.1884270000 | 3.8838860000 | -7.6283300000 |
| C | -3.1020410000 | 2.5032400000 | -3.8763460000 |
| C | -4.0795450000 | 1.7324350000 | -4.5023060000 |
| C | -3.3941280000 | 3.1878390000 | -2.6532850000 |
| C | -5.3782080000 | 1.6708500000 | -3.9891080000 |
| H | -3.8247700000 | 1.1987420000 | -5.4129090000 |
| C | -4.7285060000 | 3.1207730000 | -2.1687620000 |
| C | -5.7001350000 | 2.3850520000 | -2.8269060000 |
| H | -6.1351020000 | 1.0838100000 | -4.4994040000 |
| H | -4.9808030000 | 3.6365050000 | -1.2495890000 |
| H | -6.7113210000 | 2.3519360000 | -2.4331220000 |
| C | -2.3553670000 | 3.7794610000 | -1.8945890000 |
| C | -2.4965360000 | 4.5338840000 | -0.6247460000 |
| H | -3.3507890000 | 4.2120800000 | -0.0249240000 |
| H | -1.5825370000 | 4.4195810000 | -0.0355430000 |
| H | -2.6181350000 | 5.6094230000 | -0.8320160000 |
| H | -1.3570010000 | 3.7301340000 | -2.3106610000 |
| C | 0.7103900000 | 2.3450400000 | 0.8431770000 |
| O | 0.7933900000 | 3.2577040000 | -0.0098300000 |
| O | 0.4238040000 | 1.1138160000 | 0.5640150000 |
| C | 0.9402340000 | 2.6560990000 | 2.3045100000 |
| H | -0.0247740000 | 2.6333760000 | 2.8244270000 |

| | | | |
|---|---|---|---|
| H | 1.5720740000 | 1.8893290000 | 2.7615910000 |
| H | 1.3890260000 | 3.6429700000 | 2.4247210000 |
| O | 0.2722780000 | 2.0684080000 | -2.1965970000 |
| H | 0.5687110000 | 2.6501370000 | -1.3966680000 |
| H | 1.0682660000 | 1.7838820000 | -2.6720480000 |

**$^3$TS-3'**
Charge: 1
Multiplicity: 3
B3LYP-D3(BJ)/6-31G(d)-SDD SCF energy (au): -1693.81918759
B3LYP-D3(BJ)/6-31G(d)-SDD enthalpy (au): -1693.24074959
B3LYP-D3(BJ)/6-31G(d)-SDD free energy (au): -1693.35265159
B3LYP-D3(BJ)/def2-TZVP/SMD SCF energy (au): -2834.38717027
B3LYP-D3(BJ)/def2-TZVP/SMD enthalpy (au): -2833.80873227
B3LYP-D3(BJ)/def2-TZVP/SMD free energy (au): -2833.92063427
B3LYP-D3(BJ)/def2-TZVP/SMD free energy (quasi-harmonic) (au): -2833.91087168

Cartesian coordinates

| ATOM | X | Y | Z |
|---|---|---|---|
| Fe | -0.5337290000 | 0.4741070000 | -1.1181160000 |
| C | -2.4177410000 | -2.8962510000 | 0.9101520000 |
| N | -3.0939730000 | -1.7991090000 | 1.4129170000 |
| H | -3.8090920000 | -1.8189930000 | 2.1263970000 |
| C | -2.6258700000 | -0.6903480000 | 0.7914700000 |
| H | -2.9651990000 | 0.3158220000 | 0.9804360000 |
| N | -1.6865410000 | -1.0215030000 | -0.0807800000 |
| C | -1.5449110000 | -2.3944240000 | -0.0169310000 |
| H | -0.8354990000 | -2.9145230000 | -0.6414650000 |
| H | -2.6195050000 | -3.9003290000 | 1.2482730000 |
| C | 2.1582460000 | -2.5043060000 | -2.2338980000 |
| N | 2.7310380000 | -2.0071560000 | -1.0768900000 |
| H | 3.5867720000 | -2.3338640000 | -0.6499500000 |
| C | 1.9551110000 | -1.0048180000 | -0.6067580000 |
| H | 2.1351700000 | -0.4319100000 | 0.2886950000 |
| N | 0.9114520000 | -0.8378240000 | -1.4108430000 |
| C | 1.0227040000 | -1.7675420000 | -2.4324150000 |
| H | 0.2762970000 | -1.8287500000 | -3.2075760000 |
| H | 2.6008810000 | -3.3123090000 | -2.7944210000 |
| O | -1.4985220000 | -0.2189390000 | -2.8952590000 |
| H | -1.3557070000 | 0.3936350000 | -3.6779270000 |
| H | -2.4390790000 | -0.1226340000 | -2.6716700000 |
| F | -1.9129950000 | 1.6338180000 | -0.8149180000 |
| C | -1.7689780000 | 2.5553600000 | -4.5818390000 |
| O | -1.1524440000 | 1.5171310000 | -4.8830720000 |
| N | -1.3565460000 | 3.7929150000 | -4.9076240000 |
| H | -2.0022260000 | 4.5446290000 | -4.7034210000 |
| C | -0.1778170000 | 4.1460010000 | -5.7455950000 |
| C | -0.1366990000 | 5.6784400000 | -5.7865940000 |
| C | 1.1081260000 | 3.6029960000 | -5.1068800000 |
| C | -0.3704810000 | 3.5738090000 | -7.1587580000 |
| H | -1.0527170000 | 6.0891670000 | -6.2284970000 |
| H | -0.0108540000 | 6.0974730000 | -4.7816440000 |
| H | 0.7050520000 | 6.0118150000 | -6.3994660000 |

| | | | |
|---|---|---|---|
| H | 1.0933140000 | 2.5120180000 | -5.0853700000 |
| H | 1.9703650000 | 3.9321520000 | -5.6955640000 |
| H | 1.2188500000 | 3.9781320000 | -4.0852050000 |
| H | 0.4808940000 | 3.8470250000 | -7.7905140000 |
| H | -0.4431390000 | 2.4841720000 | -7.1265320000 |
| H | -1.2798790000 | 3.9750680000 | -7.6187010000 |
| C | -3.0989510000 | 2.4816260000 | -3.8798250000 |
| C | -4.1114720000 | 1.7940600000 | -4.5459410000 |
| C | -3.3482290000 | 3.0710700000 | -2.6035320000 |
| C | -5.3996760000 | 1.7273430000 | -4.0082490000 |
| H | -3.8920380000 | 1.3285670000 | -5.5018620000 |
| C | -4.6683000000 | 2.9996700000 | -2.0900080000 |
| C | -5.6765050000 | 2.3483550000 | -2.7836810000 |
| H | -6.1840300000 | 1.2049540000 | -4.5466610000 |
| H | -4.8839200000 | 3.4431450000 | -1.1247450000 |
| H | -6.6792150000 | 2.3086650000 | -2.3696510000 |
| C | -2.2720630000 | 3.5782480000 | -1.8227660000 |
| C | -2.3944590000 | 4.3307950000 | -0.5500700000 |
| H | -3.1983640000 | 3.9578290000 | 0.0895220000 |
| H | -1.4469650000 | 4.2845030000 | -0.0096240000 |
| H | -2.6059760000 | 5.3907350000 | -0.7642950000 |
| H | -1.2884650000 | 3.5569740000 | -2.2742460000 |
| C | 0.6778640000 | 2.3608230000 | 0.7634850000 |
| O | 0.7884410000 | 3.2493620000 | -0.1158310000 |
| O | 0.4261730000 | 1.1209180000 | 0.5218310000 |
| C | 0.8313800000 | 2.7369550000 | 2.2218620000 |
| H | -0.1681270000 | 2.8395130000 | 2.6619280000 |
| H | 1.3511440000 | 1.9480790000 | 2.7716150000 |
| H | 1.3580730000 | 3.6871970000 | 2.3231130000 |
| O | 0.4178110000 | 1.9923840000 | -2.2957140000 |
| H | 0.6574970000 | 2.5885040000 | -1.4889300000 |
| H | 1.2459080000 | 1.7327440000 | -2.7278370000 |

**$^5$TS-3-OH**
Charge: 1
Multiplicity: 5
B3LYP-D3(BJ)/6-31G(d)-SDD SCF energy (au): -1690.97145734
B3LYP-D3(BJ)/6-31G(d)-SDD enthalpy (au): -1690.38479934
B3LYP-D3(BJ)/6-31G(d)-SDD free energy (au): -1690.50114534
B3LYP-D3(BJ)/def2-TZVP/SMD SCF energy (au): -2831.53192102
B3LYP-D3(BJ)/def2-TZVP/SMD enthalpy (au): -2830.94526302
B3LYP-D3(BJ)/def2-TZVP/SMD free energy (au): -2831.06160902
B3LYP-D3(BJ)/def2-TZVP/SMD free energy (quasi-harmonic) (au): -2831.04842568

Cartesian coordinates

| ATOM | X | Y | Z |
|---|---|---|---|
| Fe | -0.0357060000 | 0.6259070000 | -1.1433790000 |
| C | 0.5411030000 | 1.3783600000 | 3.1572830000 |
| N | 1.2998640000 | 2.3454820000 | 2.5223460000 |
| H | 1.7769210000 | 3.1146000000 | 2.9714520000 |
| C | 1.2789830000 | 2.0955700000 | 1.1903010000 |
| H | 1.7671790000 | 2.6910080000 | 0.4339880000 |
| N | 0.5493630000 | 1.0197300000 | 0.9424870000 |

| | | | |
|---|---|---|---|
| C | 0.0771110000 | 0.5641600000 | 2.1586080000 |
| H | -0.5571730000 | -0.3068200000 | 2.2219170000 |
| C | -3.1241020000 | -2.0355100000 | 0.7206580000 |
| N | -3.6864550000 | -0.7714910000 | 0.7208130000 |
| H | -4.5847660000 | -0.5211470000 | 1.1098850000 |
| C | -2.8303700000 | 0.0754890000 | 0.0984230000 |
| H | -2.9839800000 | 1.1337720000 | -0.0467090000 |
| N | -1.7488520000 | -0.5805880000 | -0.2986980000 |
| C | -1.9191250000 | -1.8983100000 | 0.0833370000 |
| H | -1.1740820000 | -2.6463520000 | -0.1383760000 |
| O | 1.5613480000 | 1.4262830000 | -1.7890870000 |
| H | 2.2276090000 | 0.7473320000 | -1.9867290000 |
| H | -3.6240750000 | -2.8877060000 | 1.1535850000 |
| H | 0.4076170000 | 1.3637820000 | 4.2275480000 |
| C | 1.8185300000 | -3.4038890000 | -1.4553040000 |
| N | 2.3028630000 | -2.9939380000 | -0.2265660000 |
| H | 2.9156580000 | -3.5243320000 | 0.3769800000 |
| C | 1.8280430000 | -1.7473560000 | 0.0207570000 |
| H | 2.0415360000 | -1.1819920000 | 0.9144080000 |
| N | 1.0646150000 | -1.3421060000 | -0.9817490000 |
| C | 1.0508250000 | -2.3654590000 | -1.9126090000 |
| H | 0.4916960000 | -2.2628780000 | -2.8289300000 |
| H | 2.0635430000 | -4.3658350000 | -1.8773390000 |
| O | -0.7132030000 | -0.1505230000 | -3.0225950000 |
| H | -0.5486320000 | 0.5316640000 | -3.7454160000 |
| H | -1.6781030000 | -0.1985820000 | -2.9261700000 |
| F | -1.2231940000 | 2.0370700000 | -1.1297490000 |
| C | -0.8282290000 | 2.7218870000 | -5.0237580000 |
| O | -0.5126330000 | 1.5121000000 | -5.0205710000 |
| N | -1.4596150000 | 3.2835570000 | -6.0742500000 |
| H | -1.5712870000 | 4.2877890000 | -6.0448590000 |
| C | -1.9079830000 | 2.5942690000 | -7.3111640000 |
| C | -2.6087760000 | 3.6628960000 | -8.1595280000 |
| C | -0.6917780000 | 2.0314930000 | -8.0637060000 |
| C | -2.8983080000 | 1.4762350000 | -6.9505560000 |
| H | -3.4725100000 | 4.0840260000 | -7.6318030000 |
| H | -1.9217120000 | 4.4780590000 | -8.4165170000 |
| H | -2.9669140000 | 3.2218330000 | -9.0938030000 |
| H | -0.1685900000 | 1.2937450000 | -7.4523390000 |
| H | -1.0186770000 | 1.5509240000 | -8.9917740000 |
| H | 0.0065180000 | 2.8349220000 | -8.3215720000 |
| H | -3.2682360000 | 1.0017560000 | -7.8652610000 |
| H | -2.4152570000 | 0.7153260000 | -6.3347820000 |
| H | -3.7558640000 | 1.8840370000 | -6.4043530000 |
| C | -0.5406700000 | 3.6550260000 | -3.8915630000 |
| C | -1.5660210000 | 4.4966620000 | -3.4560870000 |
| C | 0.7542000000 | 3.7281370000 | -3.2887490000 |
| C | -1.3604360000 | 5.4168970000 | -2.4291470000 |
| H | -2.5490510000 | 4.4106610000 | -3.9103530000 |
| C | 0.9408200000 | 4.7045070000 | -2.2770220000 |
| C | -0.0915820000 | 5.5234890000 | -1.8497900000 |
| H | -2.1767070000 | 6.0469690000 | -2.0908210000 |
| H | 1.9194540000 | 4.8011740000 | -1.8205980000 |

| | | | |
|---|---|---|---|
| H | 0.0861900000 | 6.2505960000 | -1.0626020000 |
| C | 1.8243640000 | 2.8724490000 | -3.6728880000 |
| C | 3.2554520000 | 3.1206170000 | -3.3256850000 |
| H | 3.3948500000 | 3.2901870000 | -2.2532740000 |
| H | 3.8786440000 | 2.2698850000 | -3.6158470000 |
| H | 3.6429390000 | 4.0059060000 | -3.8524940000 |
| H | 1.6145610000 | 2.1196340000 | -4.4214260000 |

**$^{5}$TS-3**
Charge: 1
Multiplicity: 5
B3LYP-D3(BJ)/6-31G(d)-SDD SCF energy (au): -1690.96849496
B3LYP-D3(BJ)/6-31G(d)-SDD enthalpy (au): -1690.38134196
B3LYP-D3(BJ)/6-31G(d)-SDD free energy (au): -1690.49561096
B3LYP-D3(BJ)/def2-TZVP/SMD SCF energy (au): -2831.52617049
B3LYP-D3(BJ)/def2-TZVP/SMD enthalpy (au): -2830.93901749
B3LYP-D3(BJ)/def2-TZVP/SMD free energy (au): -2831.05328649
B3LYP-D3(BJ)/def2-TZVP/SMD free energy (quasi-harmonic) (au): -2831.04225653

Cartesian coordinates

| ATOM | X | Y | Z |
|---|---|---|---|
| Fe | -0.6871620000 | 0.7391400000 | -1.0785410000 |
| C | 0.8973180000 | 1.5048730000 | 2.9791100000 |
| N | 1.2791790000 | 2.6092450000 | 2.2380460000 |
| H | 1.7574920000 | 3.4252480000 | 2.5929410000 |
| C | 0.8915340000 | 2.4136550000 | 0.9526030000 |
| H | 1.0450470000 | 3.0961090000 | 0.1296750000 |
| N | 0.2840250000 | 1.2428730000 | 0.8368110000 |
| C | 0.2776070000 | 0.6656500000 | 2.0913700000 |
| H | -0.1662620000 | -0.3035880000 | 2.2589250000 |
| C | -2.4215880000 | -2.6918650000 | 1.1645100000 |
| N | -3.2898600000 | -1.6502240000 | 1.4390220000 |
| H | -4.0990820000 | -1.6955610000 | 2.0421880000 |
| C | -2.8746900000 | -0.5593180000 | 0.7482530000 |
| H | -3.3499010000 | 0.4088250000 | 0.7679530000 |
| N | -1.7877940000 | -0.8483360000 | 0.0494420000 |
| C | -1.4943380000 | -2.1753280000 | 0.2984930000 |
| H | -0.6490210000 | -2.6567910000 | -0.1676380000 |
| O | 0.2228690000 | 2.1804140000 | -1.9370180000 |
| H | 0.8421680000 | 1.8776140000 | -2.6167020000 |
| H | -2.5475690000 | -3.6734240000 | 1.5936640000 |
| H | 1.0972880000 | 1.4249910000 | 4.0361240000 |
| C | 2.0970960000 | -2.4442510000 | -2.3480190000 |
| N | 2.8394520000 | -1.8020580000 | -1.3741810000 |
| H | 3.7839560000 | -2.0322390000 | -1.0992640000 |
| C | 2.0833600000 | -0.8001280000 | -0.8590280000 |
| H | 2.4096160000 | -0.1331670000 | -0.0765730000 |
| N | 0.9000150000 | -0.7689290000 | -1.4502220000 |
| C | 0.8945760000 | -1.7895880000 | -2.3845820000 |
| H | 0.0284360000 | -1.9626710000 | -3.0030560000 |
| H | 2.4857630000 | -3.2793070000 | -2.9092830000 |
| O | -1.6568980000 | -0.2082690000 | -2.8003740000 |
| H | -1.4532810000 | 0.3270060000 | -3.6243320000 |

| | | | |
|---|---|---|---|
| H | -2.6144420000 | -0.1207460000 | -2.6677850000 |
| F | -2.2136900000 | 1.8499900000 | -0.6640580000 |
| C | -1.7660400000 | 2.4651650000 | -4.5444870000 |
| O | -1.1146270000 | 1.4383210000 | -4.8156900000 |
| N | -1.3459480000 | 3.7112970000 | -4.8231700000 |
| H | -2.0107370000 | 4.4530710000 | -4.6480880000 |
| C | -0.1053790000 | 4.0918700000 | -5.5484930000 |
| C | -0.0669090000 | 5.6251070000 | -5.5346370000 |
| C | 1.1284370000 | 3.5308980000 | -4.8276670000 |
| C | -0.1818540000 | 3.5677880000 | -6.9911480000 |
| H | -0.9505410000 | 6.0497680000 | -6.0268410000 |
| H | -0.0162970000 | 6.0083620000 | -4.5089380000 |
| H | 0.8155640000 | 5.9808400000 | -6.0733390000 |
| H | 1.1403530000 | 2.4405400000 | -4.8813080000 |
| H | 2.0336400000 | 3.9139880000 | -5.3102860000 |
| H | 1.1298140000 | 3.8317280000 | -3.7764890000 |
| H | 0.7209700000 | 3.8532650000 | -7.5410930000 |
| H | -0.2624120000 | 2.4781310000 | -6.9982430000 |
| H | -1.0481210000 | 3.9892350000 | -7.5124550000 |
| C | -3.1477700000 | 2.3655220000 | -3.9585660000 |
| C | -4.0743930000 | 1.6340500000 | -4.7023570000 |
| C | -3.5346180000 | 2.9871490000 | -2.7336060000 |
| C | -5.4081930000 | 1.5513140000 | -4.2982300000 |
| H | -3.7470110000 | 1.1488680000 | -5.6168170000 |
| C | -4.9007240000 | 2.9052740000 | -2.3616740000 |
| C | -5.8194800000 | 2.2081340000 | -3.1309500000 |
| H | -6.1228000000 | 0.9938330000 | -4.8953020000 |
| H | -5.2257990000 | 3.3710580000 | -1.4383560000 |
| H | -6.8583480000 | 2.1599700000 | -2.8188430000 |
| C | -2.5489360000 | 3.5375090000 | -1.8572450000 |
| C | -2.8402200000 | 4.4296490000 | -0.7030080000 |
| H | -3.7459310000 | 4.1482140000 | -0.1601480000 |
| H | -2.0006460000 | 4.4162900000 | -0.0042640000 |
| H | -2.9708780000 | 5.4655670000 | -1.0524030000 |
| H | -1.5119210000 | 3.4794100000 | -2.1745900000 |

**$^{3}$TS-3**

Charge: 1
Multiplicity: 3
B3LYP-D3(BJ)/6-31G(d)-SDD SCF energy (au): -1690.94345980
B3LYP-D3(BJ)/6-31G(d)-SDD enthalpy (au): -1690.35548580
B3LYP-D3(BJ)/6-31G(d)-SDD free energy (au): -1690.46779380
B3LYP-D3(BJ)/def2-TZVP/SMD SCF energy (au): -2831.50402869
B3LYP-D3(BJ)/def2-TZVP/SMD enthalpy (au): -2830.91605469
B3LYP-D3(BJ)/def2-TZVP/SMD free energy (au): -2831.02836269
B3LYP-D3(BJ)/def2-TZVP/SMD free energy (quasi-harmonic) (au): -2831.01784689

Cartesian coordinates

| ATOM | X | Y | Z |
|---|---|---|---|
| Fe | -0.7006130000 | 0.5611590000 | -1.1149670000 |
| C | 0.6711990000 | 1.8952590000 | 2.9877220000 |
| N | 1.0095660000 | 2.9323860000 | 2.1380810000 |
| H | 1.3967950000 | 3.8232560000 | 2.4153430000 |

| | | | |
|---|---|---|---|
| C | 0.7333040000 | 2.5414490000 | 0.8646490000 |
| H | 0.8861100000 | 3.1392100000 | -0.0216260000 |
| N | 0.2393910000 | 1.3156050000 | 0.8578860000 |
| C | 0.1935580000 | 0.9008560000 | 2.1729050000 |
| H | -0.1778030000 | -0.0779500000 | 2.4384570000 |
| C | -2.2718530000 | -2.7480630000 | 1.0973560000 |
| N | -3.1584950000 | -1.7238770000 | 1.3786440000 |
| H | -3.9682170000 | -1.7876060000 | 1.9793250000 |
| C | -2.7657500000 | -0.6213110000 | 0.6980470000 |
| H | -3.2598030000 | 0.3364550000 | 0.7201090000 |
| N | -1.6722170000 | -0.8874150000 | -0.0021920000 |
| C | -1.3531880000 | -2.2113310000 | 0.2368610000 |
| H | -0.5018950000 | -2.6770100000 | -0.2322410000 |
| O | 0.1272690000 | 1.9237430000 | -2.0305390000 |
| H | 0.6623640000 | 1.5749670000 | -2.7593600000 |
| H | -2.3794610000 | -3.7349400000 | 1.5187860000 |
| H | 0.8009120000 | 1.9626270000 | 4.0566450000 |
| C | 2.0209250000 | -2.3373700000 | -2.4162590000 |
| N | 2.7703850000 | -1.6381320000 | -1.4875080000 |
| H | 3.7343310000 | -1.8185080000 | -1.2452260000 |
| C | 1.9944170000 | -0.6603340000 | -0.9607040000 |
| H | 2.3096500000 | 0.0432390000 | -0.2072110000 |
| N | 0.7865750000 | -0.7047820000 | -1.5047460000 |
| C | 0.7866420000 | -1.7455550000 | -2.4193070000 |
| H | -0.0978690000 | -1.9680870000 | -2.9942730000 |
| H | 2.4262880000 | -3.1650170000 | -2.9763470000 |
| O | -1.8433960000 | -0.3844910000 | -2.9302470000 |
| H | -1.6084720000 | 0.1690760000 | -3.7211280000 |
| H | -2.7627010000 | -0.1437390000 | -2.7362080000 |
| F | -2.1656310000 | 1.6820240000 | -0.7382780000 |
| C | -1.7668370000 | 2.4246490000 | -4.5361810000 |
| O | -1.1804380000 | 1.3846090000 | -4.8814650000 |
| N | -1.2805950000 | 3.6640170000 | -4.7414270000 |
| H | -1.9049510000 | 4.4281540000 | -4.5200360000 |
| C | -0.0386830000 | 4.0184400000 | -5.4761750000 |
| C | 0.0886490000 | 5.5434170000 | -5.3728940000 |
| C | 1.1782520000 | 3.3471820000 | -4.8236120000 |
| C | -0.1769080000 | 3.5878420000 | -6.9450830000 |
| H | -0.7798020000 | 6.0459260000 | -5.8163750000 |
| H | 0.1833580000 | 5.8607450000 | -4.3279100000 |
| H | 0.9779100000 | 5.8812980000 | -5.9119920000 |
| H | 1.1251090000 | 2.2629000000 | -4.9398480000 |
| H | 2.0925460000 | 3.7044970000 | -5.3088100000 |
| H | 1.2248620000 | 3.5876810000 | -3.7577730000 |
| H | 0.7280800000 | 3.8548520000 | -7.5007750000 |
| H | -0.3207320000 | 2.5069490000 | -7.0142160000 |
| H | -1.0290270000 | 4.0881500000 | -7.4177200000 |
| C | -3.1448280000 | 2.3688890000 | -3.9319930000 |
| C | -4.1233680000 | 1.7381840000 | -4.7015280000 |
| C | -3.4777110000 | 2.9304100000 | -2.6648510000 |
| C | -5.4517160000 | 1.6988170000 | -4.2745910000 |
| H | -3.8382320000 | 1.2940480000 | -5.6501780000 |
| C | -4.8354260000 | 2.8881380000 | -2.2640310000 |

| | | | |
|---|---|---|---|
| C | -5.8055100000 | 2.2924520000 | -3.0567460000 |
| H | -6.2056940000 | 1.2196520000 | -4.8909570000 |
| H | -5.1175930000 | 3.3075450000 | -1.3048960000 |
| H | -6.8389380000 | 2.2752780000 | -2.7241250000 |
| C | -2.4463520000 | 3.3878860000 | -1.7784080000 |
| C | -2.6822270000 | 4.2368120000 | -0.5781090000 |
| H | -3.5726290000 | 3.9463150000 | -0.0148380000 |
| H | -1.8193920000 | 4.1814750000 | 0.0897150000 |
| H | -2.8092000000 | 5.2869190000 | -0.8823910000 |
| H | -1.4257740000 | 3.3348840000 | -2.1427980000 |

**benzyl radical**
Charge: 0
Multiplicity: 2
B3LYP-D3(BJ)/6-31G(d)-SDD SCF energy (au): -636.239341674
B3LYP-D3(BJ)/6-31G(d)-SDD enthalpy (au): -635.938361674
B3LYP-D3(BJ)/6-31G(d)-SDD free energy (au): -635.998826674
B3LYP-D3(BJ)/def2-TZVP/SMD SCF energy (au): -636.509902367
B3LYP-D3(BJ)/def2-TZVP/SMD enthalpy (au): -636.208922367
B3LYP-D3(BJ)/def2-TZVP/SMD free energy (au): -636.269387367
B3LYP-D3(BJ)/def2-TZVP/SMD free energy (quasi-harmonic) (au): -636.267443693

Cartesian coordinates

| ATOM | X | Y | Z |
|---|---|---|---|
| C | -0.0183400000 | 0.2415300000 | -3.6542560000 |
| O | -0.6218130000 | 0.0465660000 | -2.5976030000 |
| N | -0.2011640000 | -0.5294420000 | -4.7684740000 |
| H | 0.2116950000 | -0.1935130000 | -5.6265490000 |
| C | -1.1344330000 | -1.6696330000 | -4.8758600000 |
| C | -0.9812620000 | -2.2192990000 | -6.2998290000 |
| C | -0.7503950000 | -2.7514810000 | -3.8535510000 |
| C | -2.5792960000 | -1.1942350000 | -4.6449780000 |
| H | -1.2433410000 | -1.4601750000 | -7.0475780000 |
| H | 0.0471560000 | -2.5514860000 | -6.4847350000 |
| H | -1.6461070000 | -3.0759130000 | -6.4459790000 |
| H | -0.8210470000 | -2.3573010000 | -2.8384790000 |
| H | -1.4224660000 | -3.6116710000 | -3.9489590000 |
| H | 0.2758630000 | -3.0932590000 | -4.0273560000 |
| H | -3.2747410000 | -2.0356880000 | -4.7411690000 |
| H | -2.6786600000 | -0.7655400000 | -3.6460250000 |
| H | -2.8552090000 | -0.4332560000 | -5.3837770000 |
| C | 0.9982210000 | 1.3363440000 | -3.8279580000 |
| C | 2.1761030000 | 1.0563240000 | -4.5212120000 |
| C | 0.7863920000 | 2.6467810000 | -3.2735270000 |
| C | 3.1690430000 | 2.0225150000 | -4.7042510000 |
| H | 2.3355030000 | 0.0503170000 | -4.8992710000 |
| C | 1.8118080000 | 3.6126580000 | -3.4954290000 |
| C | 2.9715810000 | 3.3091970000 | -4.1889610000 |
| H | 4.0830690000 | 1.7724470000 | -5.2340220000 |
| H | 1.6773970000 | 4.6139130000 | -3.1004220000 |
| H | 3.7325110000 | 4.0727320000 | -4.3258870000 |
| C | -0.3852870000 | 2.9917540000 | -2.5611530000 |
| C | -0.6479340000 | 4.3567830000 | -2.0066660000 |

| | | | |
|---|---|---|---|
| H | -0.6140680000 | 5.1357650000 | -2.7825860000 |
| H | -1.6329400000 | 4.4018390000 | -1.5343500000 |
| H | 0.0951410000 | 4.6437410000 | -1.2470110000 |
| H | -1.1141880000 | 2.2137100000 | -2.3790770000 |

**benzyl radical_Arg**
Charge: 1
Multiplicity: 2
B3LYP-D3(BJ)/6-31G(d)-SDD SCF energy (au): -881.348122470
B3LYP-D3(BJ)/6-31G(d)-SDD enthalpy (au): -880.919631470
B3LYP-D3(BJ)/6-31G(d)-SDD free energy (au): -881.001373470
B3LYP-D3(BJ)/def2-TZVP/SMD SCF energy (au): -881.780057450
B3LYP-D3(BJ)/def2-TZVP/SMD enthalpy (au): -881.351566450
B3LYP-D3(BJ)/def2-TZVP/SMD free energy (au): -881.433308450
B3LYP-D3(BJ)/def2-TZVP/SMD free energy (quasi-harmonic) (au): -881.425923980

Cartesian coordinates

| ATOM | X | Y | Z |
|---|---|---|---|
| C | -5.6633590000 | -0.8097060000 | -4.3164370000 |
| C | -4.4326890000 | -0.1684470000 | -4.3448790000 |
| C | -3.3117700000 | -0.7761480000 | -4.9482540000 |
| C | -3.4261260000 | -2.0633970000 | -5.5287470000 |
| C | -4.6755190000 | -2.6909000000 | -5.4701180000 |
| C | -5.7818260000 | -2.0816310000 | -4.8806130000 |
| H | -6.5212270000 | -0.3245650000 | -3.8626110000 |
| H | -4.3261850000 | 0.8251160000 | -3.9226350000 |
| H | -4.7774410000 | -3.6849680000 | -5.8971220000 |
| H | -6.7355940000 | -2.6002130000 | -4.8614840000 |
| C | -2.2935020000 | -2.7922130000 | -6.2166810000 |
| C | -2.2968440000 | -2.5334710000 | -7.7339450000 |
| H | -2.4135070000 | -3.8671870000 | -6.0421300000 |
| H | -1.3297770000 | -2.5024020000 | -5.7937130000 |
| H | -1.4433190000 | -3.0253920000 | -8.2153690000 |
| H | -3.2101950000 | -2.9230330000 | -8.1944660000 |
| H | -2.2634630000 | -1.4600280000 | -7.9504790000 |
| C | -2.0297670000 | -0.0486250000 | -4.8720460000 |
| O | -1.0714250000 | -0.1967610000 | -5.6620930000 |
| N | -1.9353170000 | 0.9053850000 | -3.8904120000 |
| C | -1.2639790000 | 0.6453170000 | -2.6120680000 |
| C | -1.6258680000 | -0.7468800000 | -2.0613360000 |
| C | -1.6945860000 | 1.7500120000 | -1.6392870000 |
| H | -1.1447950000 | -0.8903130000 | -1.0892060000 |
| H | -2.7068170000 | -0.8457460000 | -1.9269770000 |
| H | -1.2864160000 | -1.5431100000 | -2.7322700000 |
| H | -1.4601080000 | 2.7375360000 | -2.0473250000 |
| H | -2.7721220000 | 1.7012610000 | -1.4542990000 |
| H | -1.1767830000 | 1.6332850000 | -0.6821370000 |
| C | 0.2603200000 | 0.7293550000 | -2.8754400000 |
| H | 0.5868250000 | -0.0520110000 | -3.5666710000 |
| H | 0.5278570000 | 1.7101390000 | -3.2818270000 |
| H | 0.7867170000 | 0.5970110000 | -1.9249590000 |
| H | 2.8362310000 | 3.0654020000 | -8.1724410000 |
| C | 3.2275850000 | 2.0452470000 | -8.2658140000 |

N 2.1752240000 1.0871530000 -8.5922790000
C 1.2076680000 0.7158030000 -7.7402360000
N 1.1308180000 1.2475480000 -6.5178770000
N 0.3055060000 -0.2069190000 -8.1013060000
H 3.9635040000 2.0314390000 -9.0698660000
H 3.7305900000 1.7537180000 -7.3388740000
H 2.0972160000 0.7746050000 -9.5495020000
H 1.6320080000 2.0914200000 -6.2871750000
H 0.3778830000 0.9165360000 -5.9031750000
H 0.4653520000 -0.8068450000 -8.8966860000
H -0.4116110000 -0.4732010000 -7.4260360000

**benzyl radical_water**
Charge: 0
Multiplicity: 2
B3LYP-D3(BJ)/6-31G(d)-SDD SCF energy (au): -712.642754030
B3LYP-D3(BJ)/6-31G(d)-SDD enthalpy (au): -712.313877030
B3LYP-D3(BJ)/6-31G(d)-SDD free energy (au): -712.381467030
B3LYP-D3(BJ)/def2-TZVP/SMD SCF energy (au): -712.958136430
B3LYP-D3(BJ)/def2-TZVP/SMD enthalpy (au): -712.629259430
B3LYP-D3(BJ)/def2-TZVP/SMD free energy (au): -712.696849430
B3LYP-D3(BJ)/def2-TZVP/SMD free energy (quasi-harmonic) (au): -712.693335400

Cartesian coordinates
ATOM X Y Z
N 33.8843920000 34.8698210000 34.2929100000
O 35.1005740000 36.1612670000 32.8754590000
C 35.8116800000 33.8830540000 33.1964570000
C 35.2549500000 32.6081780000 33.4060460000
C 36.0147940000 31.4579170000 33.2293950000
C 37.3540220000 31.5770200000 32.8548450000
C 37.9126040000 32.8384130000 32.6572020000
C 37.1670840000 34.0138710000 32.8123600000
C 34.9473790000 35.0697350000 33.4367240000
C 33.9216970000 35.3146410000 35.6896000000
C 35.0657180000 36.2934310000 35.9978020000
C 32.5602360000 35.9664260000 35.9914770000
C 34.0867930000 34.0177750000 36.5190120000
C 37.8482590000 35.3386410000 32.5488890000
C 37.6922580000 35.7895160000 31.0867740000
H 34.2106430000 32.5379170000 33.6897460000
H 35.5679220000 30.4800660000 33.3806260000
H 37.9645570000 30.6888770000 32.7176400000
H 38.9587480000 32.9226290000 32.3736880000
H 37.4465710000 36.1153620000 33.2034760000
H 38.9141960000 35.2281810000 32.7831030000
H 38.2297030000 36.7290360000 30.9161510000
H 38.0934790000 35.0363360000 30.3991820000
H 36.6367430000 35.9489750000 30.8545120000
H 35.0403010000 36.5603460000 37.0595070000
H 36.0428610000 35.8447660000 35.7840700000
H 34.9490390000 37.2058080000 35.4091650000
H 32.4509940000 36.8838550000 35.4070400000

H     31.7450780000   35.2799870000   35.7430370000
H     32.5001120000   36.2142690000   37.0574390000
H     35.0421760000   33.5303320000   36.3013300000
H     34.0573140000   34.2720290000   37.5839690000
H     33.2769470000   33.3146550000   36.3033990000
O     33.4831690000   38.3233320000   33.7487720000
H     33.9601660000   37.5713820000   33.3413700000
H     34.1080390000   39.0595490000   33.6762130000

**water**
Charge: 0
Multiplicity: 1
B3LYP-D3(BJ)/6-31G(d)-SDD SCF energy (au): -76.4075979388
B3LYP-D3(BJ)/6-31G(d)-SDD enthalpy (au): -76.3826819388
B3LYP-D3(BJ)/6-31G(d)-SDD free energy (au): -76.4047809388
B3LYP-D3(BJ)/def2-TZVP/SMD SCF energy (au): -76.4696566367
B3LYP-D3(BJ)/def2-TZVP/SMD enthalpy (au): -76.4447406367
B3LYP-D3(BJ)/def2-TZVP/SMD free energy (au): -76.4668396367
B3LYP-D3(BJ)/def2-TZVP/SMD free energy (quasi-harmonic) (au): -76.4668396979

Cartesian coordinates
ATOM    X               Y               Z
O     -0.3884090000   -0.2449700000    0.0000000000
H      0.5789960000   -0.1974570000    0.0000000000
H     -0.6665480000    0.6828060000    0.0000000000

**$His_2AspWat_2$_FeF_theozyme**
Charge: 2
Multiplicity: 5
B3LYP-D3(BJ)/6-31G(d)-SDD SCF energy (au): -3260.75679370
B3LYP-D3(BJ)/6-31G(d)-SDD enthalpy (au): -3259.48349470
B3LYP-D3(BJ)/6-31G(d)-SDD free energy (au): -3259.68033270
B3LYP-D3(BJ)/def2-TZVP/SMD SCF energy (au): -4402.21860609
B3LYP-D3(BJ)/def2-TZVP/SMD enthalpy (au): -4400.94530709
B3LYP-D3(BJ)/def2-TZVP/SMD free energy (au): -4401.14214509
B3LYP-D3(BJ)/def2-TZVP/SMD free energy (quasi-harmonic) (au): -4401.11949639

Cartesian coordinates
ATOM    X               Y               Z
H     42.5038290000   47.0446510000   28.5015760000
C     43.4097510000   46.4756090000   28.4865960000
H     43.4937590000   45.9793960000   27.5146630000
C     44.5125520000   47.5439690000   28.6682810000
H     45.5153780000   47.1039310000   28.6482410000
H     44.4681730000   48.2433070000   27.8227030000
C     44.2771730000   48.3108750000   29.9870720000
H     43.2127580000   48.2452540000   30.2413430000
H     44.8190650000   47.8397210000   30.8171090000
C     44.6367980000   49.8021590000   29.9383540000
H     45.7173310000   49.9619200000   29.9367560000
H     44.2668820000   50.2585040000   29.0115860000
N     44.0991390000   50.5429770000   31.0895400000
H     44.7397990000   50.8577090000   31.8040370000

| | | | |
|---|---|---|---|
| C | 42.7966790000 | 50.7988960000 | 31.2943320000 |
| N | 41.8903610000 | 50.5790820000 | 30.3374100000 |
| H | 40.9025070000 | 50.7178520000 | 30.5823600000 |
| H | 42.1633620000 | 50.4678390000 | 29.3731180000 |
| N | 42.3835440000 | 51.3025300000 | 32.4699820000 |
| H | 42.8832280000 | 51.0260170000 | 33.3071460000 |
| H | 41.3671930000 | 51.4025400000 | 32.5685710000 |
| H | 43.5181060000 | 45.7284070000 | 29.2446310000 |
| H | 45.3297990000 | 43.5663220000 | 33.0145030000 |
| C | 45.7855520000 | 43.8763940000 | 33.9322950000 |
| H | 46.8683800000 | 43.9190670000 | 33.7730810000 |
| C | 45.2568460000 | 45.3079000000 | 34.2582520000 |
| H | 45.7951300000 | 45.7278610000 | 35.1113620000 |
| H | 45.4484120000 | 45.9586740000 | 33.3940780000 |
| C | 43.8029580000 | 45.3070980000 | 34.6111580000 |
| N | 42.8128840000 | 44.8159050000 | 33.7676890000 |
| C | 41.6155920000 | 44.9236530000 | 34.3851350000 |
| H | 40.6758380000 | 44.5869170000 | 33.9785270000 |
| N | 41.7823940000 | 45.4637900000 | 35.5852850000 |
| H | 42.9631160000 | 44.3836600000 | 32.8672010000 |
| C | 43.1359880000 | 45.7069640000 | 35.7409550000 |
| H | 43.5408130000 | 46.1269020000 | 36.6468150000 |
| C | 45.4329520000 | 42.9857360000 | 35.1195990000 |
| O | 44.4716150000 | 42.2142480000 | 35.1154890000 |
| N | 46.2069390000 | 43.2279150000 | 36.2061310000 |
| H | 46.9799770000 | 43.8770580000 | 36.1211260000 |
| C | 45.8521890000 | 42.9774280000 | 37.5861110000 |
| H | 45.1576370000 | 42.1351800000 | 37.6359040000 |
| C | 47.1638900000 | 42.6567900000 | 38.3667570000 |
| H | 46.9265780000 | 42.6349240000 | 39.4411780000 |
| C | 47.7720240000 | 41.3280450000 | 37.9493450000 |
| H | 48.7098110000 | 41.1701190000 | 38.4877020000 |
| H | 47.0913300000 | 40.5017840000 | 38.1784360000 |
| H | 47.9825950000 | 41.3233260000 | 36.8763270000 |
| O | 48.1206330000 | 43.6716630000 | 38.0865810000 |
| H | 47.8133010000 | 44.5139420000 | 38.4615870000 |
| C | 45.2489260000 | 44.2640110000 | 38.1964770000 |
| O | 45.5430570000 | 45.3679180000 | 37.7344770000 |
| N | 44.4717130000 | 44.1101440000 | 39.3020370000 |
| H | 44.3711190000 | 43.1830550000 | 39.6971190000 |
| C | 44.0650740000 | 45.2523440000 | 40.1274560000 |
| H | 44.0968570000 | 46.1290670000 | 39.4806700000 |
| C | 42.6970750000 | 45.0881860000 | 40.7775310000 |
| H | 42.6080530000 | 44.1377400000 | 41.3209850000 |
| H | 42.5936980000 | 45.8561690000 | 41.5527550000 |
| C | 41.4711030000 | 45.2098360000 | 39.8923470000 |
| O | 41.6486140000 | 45.2199470000 | 38.6180170000 |
| O | 40.3544130000 | 45.2838100000 | 40.4615150000 |
| H | 44.7755570000 | 45.4080890000 | 40.9409840000 |
| H | 36.1387690000 | 40.7413690000 | 41.5246690000 |
| C | 37.1406330000 | 40.3189580000 | 41.4362380000 |
| H | 37.0060690000 | 39.2318320000 | 41.4754870000 |
| C | 37.7543780000 | 40.6575730000 | 40.0625020000 |

| | | | |
|---|---|---|---|
| H | 38.6631060000 | 40.0592080000 | 39.9084240000 |
| H | 37.0585600000 | 40.3867610000 | 39.2647300000 |
| C | 38.1499990000 | 42.1017650000 | 39.8306860000 |
| O | 37.8146550000 | 42.7154190000 | 38.7859580000 |
| N | 38.9126050000 | 42.6869990000 | 40.7555370000 |
| H | 39.2836730000 | 43.6196570000 | 40.6079760000 |
| H | 39.2251680000 | 42.2398070000 | 41.6075050000 |
| H | 37.7806810000 | 40.5614190000 | 42.2840840000 |
| H | 42.7026240000 | 40.1710440000 | 33.7692100000 |
| C | 41.7036340000 | 40.3127830000 | 33.3572800000 |
| H | 41.5048180000 | 39.4419230000 | 32.7232670000 |
| C | 40.6429290000 | 40.3579750000 | 34.4755690000 |
| H | 39.6411900000 | 40.4264140000 | 34.0387110000 |
| H | 40.6702300000 | 39.4080720000 | 35.0267010000 |
| C | 40.7932440000 | 41.4886480000 | 35.4453280000 |
| N | 41.9747370000 | 41.7558540000 | 36.1245610000 |
| H | 42.9098890000 | 41.4365600000 | 35.8750100000 |
| C | 41.7749360000 | 42.8026240000 | 36.9454250000 |
| H | 42.5464860000 | 43.2664930000 | 37.5387510000 |
| N | 40.5182760000 | 43.2360760000 | 36.8527670000 |
| C | 39.8964270000 | 42.4152910000 | 35.9180600000 |
| H | 38.8549960000 | 42.5315850000 | 35.6623170000 |
| H | 41.6373040000 | 41.1894040000 | 32.7130610000 |
| Fe | 40.3114430000 | 45.3845840000 | 37.1392470000 |
| O | 38.7018410000 | 45.1706390000 | 38.4116430000 |
| H | 38.3781660000 | 44.2240110000 | 38.5453840000 |
| H | 39.1263420000 | 45.4188810000 | 39.2798420000 |
| O | 38.8681680000 | 45.3127660000 | 35.5752920000 |
| H | 38.0752000000 | 44.7600790000 | 35.8483230000 |
| H | 38.5615310000 | 46.2297980000 | 35.4060560000 |
| C | 40.5392630000 | 48.7974970000 | 34.3590820000 |
| C | 41.1586230000 | 48.1495590000 | 33.2917920000 |
| C | 42.5480390000 | 48.0451980000 | 33.2165310000 |
| C | 43.3351400000 | 48.6251530000 | 34.2220690000 |
| C | 42.7419940000 | 49.3125460000 | 35.2691700000 |
| C | 41.3227540000 | 49.4347420000 | 35.3784800000 |
| H | 40.5344800000 | 47.7077340000 | 32.5204380000 |
| H | 43.0097580000 | 47.5075950000 | 32.3977780000 |
| H | 44.4175500000 | 48.5324090000 | 34.1861970000 |
| H | 43.3647090000 | 49.7593920000 | 36.0365560000 |
| C | 40.7425190000 | 50.1889380000 | 36.4238740000 |
| H | 39.0454010000 | 48.0864240000 | 36.2431510000 |
| H | 39.6669140000 | 50.3209810000 | 36.4261080000 |
| C | 41.5159950000 | 50.8100070000 | 37.5405680000 |
| H | 42.2595670000 | 51.5357960000 | 37.1805690000 |
| H | 42.0662470000 | 50.0517370000 | 38.1160260000 |
| H | 40.8527770000 | 51.3352540000 | 38.2311060000 |
| C | 39.0371670000 | 48.7984500000 | 34.3523650000 |
| O | 38.4154890000 | 49.1826930000 | 33.3452460000 |
| N | 38.4261820000 | 48.2974220000 | 35.4546120000 |
| C | 36.9710520000 | 48.4505800000 | 35.7702000000 |
| C | 36.5847820000 | 49.9373310000 | 35.7365480000 |
| C | 36.1334780000 | 47.6498390000 | 34.7619570000 |

| | | | |
|---|---|---|---|
| C | 36.7808320000 | 47.8899270000 | 37.1855680000 |
| H | 37.1711650000 | 50.5059000000 | 36.4670680000 |
| H | 36.7438650000 | 50.3625620000 | 34.7435380000 |
| H | 35.5272720000 | 50.0510910000 | 35.9940480000 |
| H | 35.0706280000 | 47.7483690000 | 35.0037730000 |
| H | 36.2955880000 | 48.0085690000 | 33.7441980000 |
| H | 36.3870410000 | 46.5832070000 | 34.8009300000 |
| H | 35.7321990000 | 47.9908920000 | 37.4798280000 |
| H | 37.0548590000 | 46.8324630000 | 37.2491630000 |
| H | 37.3904820000 | 48.4355320000 | 37.9141710000 |
| F | 40.1901060000 | 47.2136770000 | 37.2906480000 |
| O | 36.9490280000 | 43.6063540000 | 36.3120050000 |
| H | 37.1090850000 | 43.1359880000 | 37.1602340000 |
| H | 36.0037840000 | 43.8197300000 | 36.2957480000 |
| O | 40.6766660000 | 43.5603320000 | 42.7641170000 |
| H | 40.6519180000 | 43.7558270000 | 43.7129760000 |
| H | 40.4776980000 | 44.4041350000 | 42.3177260000 |
| O | 39.6441080000 | 51.0622760000 | 31.8821780000 |
| H | 38.9753150000 | 51.7567890000 | 31.7786750000 |
| H | 39.1875180000 | 50.3107780000 | 32.3543050000 |

**$His_2AspWat_2$_Fe_theozyme**
Charge: 2
Multiplicity: 5
B3LYP-D3(BJ)/6-31G(d)-SDD SCF energy (au): -3260.68853372
B3LYP-D3(BJ)/6-31G(d)-SDD enthalpy (au): -3259.41609572
B3LYP-D3(BJ)/6-31G(d)-SDD free energy (au): -3259.61069672
B3LYP-D3(BJ)/def2-TZVP/SMD SCF energy (au): -4402.15419091
B3LYP-D3(BJ)/def2-TZVP/SMD enthalpy (au): -4400.88175291
B3LYP-D3(BJ)/def2-TZVP/SMD free energy (au): -4401.07635391
B3LYP-D3(BJ)/def2-TZVP/SMD free energy (quasi-harmonic) (au): -4401.05406419

Cartesian coordinates

| ATOM | X | Y | Z |
|---|---|---|---|
| H | 42.5036580000 | 47.0445370000 | 28.5014600000 |
| C | 43.4099960000 | 46.4760150000 | 28.4870150000 |
| H | 43.4454140000 | 45.9517690000 | 27.5255990000 |
| C | 44.5598170000 | 47.5010810000 | 28.5632380000 |
| H | 45.5366920000 | 47.0060850000 | 28.5009590000 |
| H | 44.4957770000 | 48.1669730000 | 27.6922100000 |
| C | 44.4825130000 | 48.3403050000 | 29.8479020000 |
| H | 43.4426740000 | 48.6470600000 | 30.0157530000 |
| H | 44.7724300000 | 47.7313960000 | 30.7142950000 |
| C | 45.3645930000 | 49.5934430000 | 29.8041490000 |
| H | 46.4047180000 | 49.3257320000 | 29.6030410000 |
| H | 45.0604140000 | 50.2542890000 | 28.9824620000 |
| N | 45.3764890000 | 50.3489030000 | 31.0677770000 |
| H | 46.2408290000 | 50.3738640000 | 31.5902320000 |
| C | 44.3647020000 | 51.0720290000 | 31.5707300000 |
| N | 43.2520230000 | 51.3025090000 | 30.8661560000 |
| H | 42.4904080000 | 51.7978060000 | 31.3365560000 |
| H | 43.2353180000 | 51.1944260000 | 29.8632920000 |
| N | 44.4629670000 | 51.5913550000 | 32.8051390000 |

| | | | |
|---|---|---|---|
| H | 45.0123090000 | 51.0932680000 | 33.4925460000 |
| H | 43.6318780000 | 52.0874830000 | 33.1496280000 |
| H | 43.5178380000 | 45.7281210000 | 29.2446000000 |
| H | 45.3308130000 | 43.5670830000 | 33.0134060000 |
| C | 45.7849840000 | 43.8750270000 | 33.9320090000 |
| H | 46.8685770000 | 43.9109920000 | 33.7699320000 |
| C | 45.2783230000 | 45.3084310000 | 34.2609800000 |
| H | 45.8177600000 | 45.7131370000 | 35.1211810000 |
| H | 45.5042690000 | 45.9518510000 | 33.4012410000 |
| C | 43.8204550000 | 45.3659210000 | 34.5928380000 |
| N | 42.8117650000 | 45.0566340000 | 33.6883630000 |
| C | 41.6137060000 | 45.2161570000 | 34.3049520000 |
| H | 40.6603690000 | 45.0197140000 | 33.8380180000 |
| N | 41.7937340000 | 45.6157660000 | 35.5558120000 |
| H | 42.9424420000 | 44.7178040000 | 32.7457140000 |
| C | 43.1614620000 | 45.7098090000 | 35.7472220000 |
| H | 43.5797810000 | 46.0040100000 | 36.6951180000 |
| C | 45.4463510000 | 42.9765450000 | 35.1152710000 |
| O | 44.5219880000 | 42.1642420000 | 35.1004160000 |
| N | 46.2009290000 | 43.2496240000 | 36.2093370000 |
| H | 46.9540990000 | 43.9212830000 | 36.1285080000 |
| C | 45.8498930000 | 42.9843170000 | 37.5871480000 |
| H | 45.1451370000 | 42.1501340000 | 37.6261880000 |
| C | 47.1577210000 | 42.6385160000 | 38.3631110000 |
| H | 46.9176140000 | 42.5987400000 | 39.4365210000 |
| C | 47.7544720000 | 41.3125800000 | 37.9217810000 |
| H | 48.6909050000 | 41.1369100000 | 38.4570900000 |
| H | 47.0668090000 | 40.4881910000 | 38.1360630000 |
| H | 47.9646770000 | 41.3258770000 | 36.8487420000 |
| O | 48.1238340000 | 43.6503830000 | 38.1038450000 |
| H | 47.8091110000 | 44.4916950000 | 38.4751040000 |
| C | 45.2678270000 | 44.2715970000 | 38.2135390000 |
| O | 45.6077650000 | 45.3775080000 | 37.7810260000 |
| N | 44.4634120000 | 44.1177470000 | 39.2952610000 |
| H | 44.3087360000 | 43.1847550000 | 39.6575290000 |
| C | 44.0640040000 | 45.2539410000 | 40.1299920000 |
| H | 44.1008310000 | 46.1337730000 | 39.4881470000 |
| C | 42.6872830000 | 45.1046110000 | 40.7663250000 |
| H | 42.5620030000 | 44.1358300000 | 41.2669130000 |
| H | 42.6097860000 | 45.8352190000 | 41.5802710000 |
| C | 41.4709030000 | 45.3206790000 | 39.8763330000 |
| O | 41.6550840000 | 45.5055380000 | 38.6249080000 |
| O | 40.3451380000 | 45.3129170000 | 40.4473220000 |
| H | 44.7764580000 | 45.4078360000 | 40.9404650000 |
| H | 36.1386980000 | 40.7413310000 | 41.5253400000 |
| C | 37.1400150000 | 40.3200700000 | 41.4359780000 |
| H | 37.0002510000 | 39.2332230000 | 41.4832750000 |
| C | 37.7727030000 | 40.6417910000 | 40.0692120000 |
| H | 38.6725550000 | 40.0256750000 | 39.9329450000 |
| H | 37.0826940000 | 40.3805480000 | 39.2636250000 |
| C | 38.1924630000 | 42.0794820000 | 39.8283110000 |
| O | 37.8646150000 | 42.6726390000 | 38.7781440000 |
| N | 38.9632940000 | 42.6567250000 | 40.7620650000 |

| | | | |
|---|---|---|---|
| H | 39.3541130000 | 43.5796230000 | 40.6132840000 |
| H | 39.2473570000 | 42.2084530000 | 41.6218380000 |
| H | 37.7810690000 | 40.5604590000 | 42.2841310000 |
| H | 42.7023980000 | 40.1715690000 | 33.7695640000 |
| C | 41.7039960000 | 40.3109990000 | 33.3550050000 |
| H | 41.5182300000 | 39.4431700000 | 32.7122880000 |
| C | 40.6347830000 | 40.3346850000 | 34.4655150000 |
| H | 39.6353060000 | 40.4118050000 | 34.0254210000 |
| H | 40.6590340000 | 39.3753230000 | 35.0002650000 |
| C | 40.7974440000 | 41.4512990000 | 35.4478410000 |
| N | 42.0148400000 | 41.7416950000 | 36.0428070000 |
| H | 42.9355000000 | 41.4162300000 | 35.7557310000 |
| C | 41.8540660000 | 42.7884880000 | 36.8736000000 |
| H | 42.6550450000 | 43.2665860000 | 37.4148220000 |
| N | 40.5871100000 | 43.1895550000 | 36.8723510000 |
| C | 39.9143870000 | 42.3568090000 | 35.9887330000 |
| H | 38.8495150000 | 42.4471210000 | 35.8347390000 |
| H | 41.6373820000 | 41.1898020000 | 32.7136170000 |
| Fe | 40.2827760000 | 45.2481020000 | 37.1241640000 |
| O | 38.6023580000 | 45.1418450000 | 38.4624060000 |
| H | 38.2988970000 | 44.1670260000 | 38.5628270000 |
| H | 39.0910990000 | 45.3476400000 | 39.3080880000 |
| O | 39.1199130000 | 46.4776060000 | 35.8729680000 |
| H | 38.2001970000 | 46.6552450000 | 36.2444050000 |
| H | 39.5869330000 | 47.3303370000 | 35.8453110000 |
| C | 41.9331000000 | 49.7042660000 | 35.0330150000 |
| C | 41.7613140000 | 49.2595280000 | 33.7142550000 |
| C | 42.7894850000 | 48.5981420000 | 33.0538450000 |
| C | 44.0171620000 | 48.4397350000 | 33.6970170000 |
| C | 44.1907440000 | 48.8905070000 | 35.0036540000 |
| C | 43.1496430000 | 49.5043780000 | 35.7154350000 |
| H | 40.8137860000 | 49.4371850000 | 33.2141350000 |
| H | 42.6407440000 | 48.2305310000 | 32.0457280000 |
| H | 44.8435080000 | 47.9656640000 | 33.1783500000 |
| H | 45.1439650000 | 48.7383870000 | 35.5037720000 |
| C | 43.3885110000 | 49.9120050000 | 37.1539020000 |
| H | 42.5870780000 | 50.5651100000 | 37.5074890000 |
| H | 44.3135090000 | 50.5000300000 | 37.2021010000 |
| C | 43.5190340000 | 48.7016650000 | 38.0982420000 |
| H | 44.3572660000 | 48.0571160000 | 37.8122870000 |
| H | 42.6102980000 | 48.0943930000 | 38.0944640000 |
| H | 43.6982190000 | 49.0475600000 | 39.1217010000 |
| C | 40.7886720000 | 50.4923240000 | 35.5872370000 |
| O | 40.3857500000 | 51.5203900000 | 35.0326470000 |
| N | 40.2029850000 | 50.0577730000 | 36.7483270000 |
| C | 38.8407690000 | 50.4159140000 | 37.2613390000 |
| C | 38.8886480000 | 51.9104390000 | 37.6211880000 |
| C | 37.7801730000 | 50.1216230000 | 36.1915200000 |
| C | 38.5896700000 | 49.5897910000 | 38.5315470000 |
| H | 39.6698130000 | 52.1034980000 | 38.3625200000 |
| H | 39.0659290000 | 52.5314980000 | 36.7435340000 |
| H | 37.9251170000 | 52.1897010000 | 38.0576440000 |
| H | 36.7904020000 | 50.3833410000 | 36.5783730000 |

| | | | |
|---|---|---|---|
| H | 37.9560120000 | 50.7108560000 | 35.2885520000 |
| H | 37.7670460000 | 49.0586310000 | 35.9305800000 |
| H | 37.6896360000 | 49.9790330000 | 39.0160770000 |
| H | 38.4259010000 | 48.5333870000 | 38.3063160000 |
| H | 39.4206130000 | 49.6780890000 | 39.2370140000 |
| F | 40.4817590000 | 48.6706330000 | 36.9356440000 |
| O | 36.8451960000 | 46.6448780000 | 37.1273710000 |
| H | 37.2424190000 | 46.1278030000 | 37.8686010000 |
| H | 36.3298030000 | 47.3690540000 | 37.5100370000 |
| O | 40.6806900000 | 43.6709350000 | 42.7885140000 |
| H | 40.5735330000 | 43.9400010000 | 43.7129580000 |
| H | 40.4689860000 | 44.4623580000 | 42.2576660000 |
| O | 41.8682620000 | 52.6100150000 | 32.9837380000 |
| H | 41.6555060000 | 53.5566070000 | 33.0044480000 |
| H | 41.3004580000 | 52.1938230000 | 33.6802040000 |

**$His_2$AspWatOH_FeF_theozyme**

Charge: 1
Multiplicity: 5
B3LYP-D3(BJ)/6-31G(d)-SDD SCF energy (au): -3260.39205093
B3LYP-D3(BJ)/6-31G(d)-SDD enthalpy (au): -3259.13184093
B3LYP-D3(BJ)/6-31G(d)-SDD free energy (au): -3259.33195893
B3LYP-D3(BJ)/def2-TZVP/SMD SCF energy (au): -4401.75413527
B3LYP-D3(BJ)/def2-TZVP/SMD enthalpy (au): -4400.49392527
B3LYP-D3(BJ)/def2-TZVP/SMD free energy (au): -4400.69404327
B3LYP-D3(BJ)/def2-TZVP/SMD free energy (quasi-harmonic) (au): -4400.66830534

Cartesian coordinates

| ATOM | X | Y | Z |
|---|---|---|---|
| H | 42.5038250000 | 47.0446350000 | 28.5015430000 |
| C | 43.4097330000 | 46.4756580000 | 28.4866620000 |
| H | 43.4782840000 | 45.9734340000 | 27.5158730000 |
| C | 44.5359900000 | 47.5207590000 | 28.6438600000 |
| H | 45.5274190000 | 47.0602990000 | 28.5541360000 |
| H | 44.4604370000 | 48.2541580000 | 27.8291100000 |
| C | 44.4031840000 | 48.2384010000 | 29.9965050000 |
| H | 43.3415490000 | 48.4301100000 | 30.1986280000 |
| H | 44.7518660000 | 47.5786750000 | 30.8004780000 |
| C | 45.1536070000 | 49.5733150000 | 30.0867960000 |
| H | 46.2363770000 | 49.4256610000 | 30.0659980000 |
| H | 44.9253110000 | 50.2053330000 | 29.2196020000 |
| N | 44.8540590000 | 50.3239250000 | 31.3126630000 |
| H | 45.4877190000 | 50.2288630000 | 32.0932870000 |
| C | 43.6735580000 | 50.9120060000 | 31.5810770000 |
| N | 42.7593190000 | 51.0915730000 | 30.6252840000 |
| H | 41.8283210000 | 51.4205730000 | 30.9200650000 |
| H | 42.9610860000 | 50.9028570000 | 29.6570250000 |
| N | 43.4149640000 | 51.3770480000 | 32.8144570000 |
| H | 43.8251040000 | 50.8822530000 | 33.6007530000 |
| H | 42.4481180000 | 51.6964890000 | 32.9539320000 |
| H | 43.5181230000 | 45.7283700000 | 29.2446050000 |
| H | 45.3298900000 | 43.5663100000 | 33.0144610000 |
| C | 45.7854340000 | 43.8763900000 | 33.9323340000 |

| | | | |
|---|---|---|---|
| H | 46.8701340000 | 43.9110900000 | 33.7773710000 |
| C | 45.2630990000 | 45.2989370000 | 34.2819540000 |
| H | 45.8062780000 | 45.7064570000 | 35.1390680000 |
| H | 45.4640160000 | 45.9641820000 | 33.4304640000 |
| C | 43.8110900000 | 45.3097600000 | 34.6349840000 |
| N | 42.8062980000 | 44.9236710000 | 33.7536590000 |
| C | 41.6084890000 | 45.0553140000 | 34.3781800000 |
| H | 40.6435480000 | 44.8134200000 | 33.9598200000 |
| N | 41.7947890000 | 45.5009130000 | 35.6084720000 |
| H | 42.9400980000 | 44.5574630000 | 32.8227920000 |
| C | 43.1530580000 | 45.6636270000 | 35.7861890000 |
| H | 43.5614800000 | 46.0233660000 | 36.7166070000 |
| C | 45.4310110000 | 42.9423280000 | 35.0871890000 |
| O | 44.5537290000 | 42.0814900000 | 35.0130230000 |
| N | 46.1287880000 | 43.2233580000 | 36.2169270000 |
| H | 46.8047200000 | 43.9752130000 | 36.1861910000 |
| C | 45.7349960000 | 42.8984850000 | 37.5715560000 |
| H | 44.9357080000 | 42.1544490000 | 37.5392280000 |
| C | 46.9640180000 | 42.3438430000 | 38.3486390000 |
| H | 46.6656700000 | 42.2232090000 | 39.4027090000 |
| C | 47.4361040000 | 41.0098950000 | 37.7948960000 |
| H | 48.3278900000 | 40.6827260000 | 38.3360710000 |
| H | 46.6578570000 | 40.2474820000 | 37.9041360000 |
| H | 47.6879220000 | 41.1079900000 | 36.7352730000 |
| O | 48.0458260000 | 43.2578220000 | 38.2357380000 |
| H | 47.7573450000 | 44.1271850000 | 38.5629550000 |
| C | 45.2989910000 | 44.2060530000 | 38.2657660000 |
| O | 45.8059480000 | 45.2783170000 | 37.9164730000 |
| N | 44.4507090000 | 44.0989020000 | 39.3109430000 |
| H | 44.0796150000 | 43.1907310000 | 39.5578900000 |
| C | 44.0650720000 | 45.2522590000 | 40.1274620000 |
| H | 44.0968190000 | 46.1280810000 | 39.4776870000 |
| C | 42.6960710000 | 45.0936390000 | 40.7719280000 |
| H | 42.6218640000 | 44.1551740000 | 41.3375650000 |
| H | 42.5873190000 | 45.8771160000 | 41.5308240000 |
| C | 41.4680590000 | 45.1917310000 | 39.8707390000 |
| O | 41.6243960000 | 45.2555900000 | 38.6200980000 |
| O | 40.3546700000 | 45.2009360000 | 40.4861720000 |
| H | 44.7755670000 | 45.4081540000 | 40.9409840000 |
| H | 36.1387560000 | 40.7413420000 | 41.5246710000 |
| C | 37.1406500000 | 40.3190270000 | 41.4362150000 |
| H | 37.0013490000 | 39.2316140000 | 41.4837970000 |
| C | 37.7603000000 | 40.6443650000 | 40.0642460000 |
| H | 38.6743720000 | 40.0508800000 | 39.9262510000 |
| H | 37.0701020000 | 40.3591120000 | 39.2665910000 |
| C | 38.1372020000 | 42.0941800000 | 39.8204870000 |
| O | 37.7361290000 | 42.7107570000 | 38.8094860000 |
| N | 38.9469520000 | 42.6720600000 | 40.7122840000 |
| H | 39.3195080000 | 43.6030910000 | 40.5404170000 |
| H | 39.2950250000 | 42.2232280000 | 41.5489670000 |
| H | 37.7806830000 | 40.5613950000 | 42.2840980000 |
| H | 42.7025920000 | 40.1709240000 | 33.7692310000 |
| C | 41.7038120000 | 40.3129750000 | 33.3573220000 |

| | | | |
|---|---|---|---|
| H | 41.4986720000 | 39.4397450000 | 32.7272260000 |
| C | 40.6568720000 | 40.3677440000 | 34.4876460000 |
| H | 39.6500650000 | 40.4339440000 | 34.0613180000 |
| H | 40.6908140000 | 39.4170010000 | 35.0389750000 |
| C | 40.8104460000 | 41.5036750000 | 35.4517910000 |
| N | 42.0018850000 | 41.8070140000 | 36.1017430000 |
| H | 42.9383110000 | 41.5207400000 | 35.8198970000 |
| C | 41.7724930000 | 42.8457470000 | 36.9377420000 |
| H | 42.5367570000 | 43.3471390000 | 37.5084960000 |
| N | 40.5038310000 | 43.2285500000 | 36.8832110000 |
| C | 39.8945070000 | 42.3919630000 | 35.9593230000 |
| H | 38.8478740000 | 42.4859910000 | 35.7173820000 |
| H | 41.6371760000 | 41.1893510000 | 32.7129960000 |
| Fe | 40.1224680000 | 45.4757470000 | 37.0128900000 |
| O | 38.6447600000 | 45.2559610000 | 38.4742720000 |
| H | 38.3097450000 | 44.3208960000 | 38.5391050000 |
| H | 39.1534640000 | 45.4003870000 | 39.3204340000 |
| O | 38.9344050000 | 45.3304270000 | 35.6094690000 |
| H | 38.2065310000 | 44.6902440000 | 35.7773760000 |
| C | 41.3257420000 | 49.0342180000 | 34.5013200000 |
| C | 42.0228000000 | 48.4160730000 | 33.4654140000 |
| C | 43.4018660000 | 48.2165600000 | 33.5362680000 |
| C | 44.1092510000 | 48.6879390000 | 34.6525470000 |
| C | 43.4396860000 | 49.3296100000 | 35.6824040000 |
| C | 42.0230360000 | 49.5170500000 | 35.6584910000 |
| H | 41.4622320000 | 48.0745820000 | 32.6004710000 |
| H | 43.9148520000 | 47.6902050000 | 32.7413340000 |
| H | 45.1815460000 | 48.5230800000 | 34.7306890000 |
| H | 43.9980070000 | 49.6752470000 | 36.5453460000 |
| C | 41.3614500000 | 50.1659600000 | 36.7252720000 |
| H | 39.5006180000 | 48.1574040000 | 36.0104200000 |
| H | 40.2894130000 | 50.2970430000 | 36.6538370000 |
| C | 42.0291160000 | 50.5944580000 | 37.9904270000 |
| H | 42.9247040000 | 51.2057810000 | 37.8134890000 |
| H | 42.3482920000 | 49.7203650000 | 38.5782260000 |
| H | 41.3459950000 | 51.1762520000 | 38.6139320000 |
| C | 39.8406950000 | 49.1803080000 | 34.3001570000 |
| O | 39.4204570000 | 49.7103470000 | 33.2392610000 |
| N | 39.0513660000 | 48.6936770000 | 35.2530110000 |
| C | 37.5655160000 | 48.6906420000 | 35.2245350000 |
| C | 37.0444570000 | 50.1290810000 | 35.0813150000 |
| C | 37.0814790000 | 47.8026330000 | 34.0670490000 |
| C | 37.1107140000 | 48.1006810000 | 36.5659090000 |
| H | 37.4036950000 | 50.7526000000 | 35.9083030000 |
| H | 37.3662910000 | 50.5721630000 | 34.1372420000 |
| H | 35.9497820000 | 50.1253290000 | 35.1110280000 |
| H | 35.9866710000 | 47.7525010000 | 34.0703270000 |
| H | 37.4071070000 | 48.2088960000 | 33.1060860000 |
| H | 37.4859860000 | 46.7939170000 | 34.1909790000 |
| H | 36.0166140000 | 48.0979860000 | 36.6088870000 |
| H | 37.4733490000 | 47.0772430000 | 36.6827620000 |
| H | 37.4845750000 | 48.6952980000 | 37.4067670000 |
| F | 40.2736530000 | 47.3209060000 | 37.2713850000 |

| | | | |
|---|---|---|---|
| O | 36.8794060000 | 43.4540010000 | 36.2725970000 |
| H | 37.0783350000 | 43.0243520000 | 37.1303500000 |
| H | 36.1029700000 | 44.0066710000 | 36.4483790000 |
| O | 40.6747390000 | 43.5666720000 | 42.7711670000 |
| H | 40.4974450000 | 43.9129990000 | 43.6579800000 |
| H | 40.5122800000 | 44.3197380000 | 42.1634630000 |
| O | 40.7138030000 | 51.7535550000 | 32.2880190000 |
| H | 40.0854710000 | 52.4866460000 | 32.3666710000 |
| H | 40.2257620000 | 50.9291170000 | 32.6194380000 |

**$His_2$AspWatOH_Fe_theozyme**

Charge: 1

Multiplicity: 5

B3LYP-D3(BJ)/6-31G(d)-SDD SCF energy (au): -3260.30710059

B3LYP-D3(BJ)/6-31G(d)-SDD enthalpy (au): -3259.04719559

B3LYP-D3(BJ)/6-31G(d)-SDD free energy (au): -3259.23969659

B3LYP-D3(BJ)/def2-TZVP/SMD SCF energy (au): -4401.67722663

B3LYP-D3(BJ)/def2-TZVP/SMD enthalpy (au): -4400.41732163

B3LYP-D3(BJ)/def2-TZVP/SMD free energy (au): -4400.60982263

B3LYP-D3(BJ)/def2-TZVP/SMD free energy (quasi-harmonic) (au): -4400.58935104

Cartesian coordinates

| ATOM | X | Y | Z |
|---|---|---|---|
| H | 42.5036320000 | 47.0445310000 | 28.5015590000 |
| C | 43.4100850000 | 46.4759930000 | 28.4868640000 |
| H | 43.4693630000 | 45.9692820000 | 27.5177580000 |
| C | 44.5457800000 | 47.5129010000 | 28.6212690000 |
| H | 45.5301960000 | 47.0436100000 | 28.5036890000 |
| H | 44.4548160000 | 48.2455640000 | 27.8074490000 |
| C | 44.4779750000 | 48.2418490000 | 29.9751770000 |
| H | 43.4289350000 | 48.4377710000 | 30.2332080000 |
| H | 44.8747430000 | 47.5970410000 | 30.7695980000 |
| C | 45.2473040000 | 49.5687140000 | 29.9790720000 |
| H | 46.3011840000 | 49.4016880000 | 29.7424630000 |
| H | 44.8667920000 | 50.2365010000 | 29.1951030000 |
| N | 45.2225140000 | 50.2729970000 | 31.2686940000 |
| H | 46.0799230000 | 50.3171930000 | 31.8002360000 |
| C | 44.1717960000 | 50.9245390000 | 31.7927520000 |
| N | 43.0327330000 | 51.0725050000 | 31.1135420000 |
| H | 42.2273430000 | 51.4588160000 | 31.6202840000 |
| H | 42.9630140000 | 50.8413650000 | 30.1354540000 |
| N | 44.2684340000 | 51.4746520000 | 33.0173010000 |
| H | 44.8348690000 | 50.9823430000 | 33.6994080000 |
| H | 43.3856590000 | 51.8556860000 | 33.3819250000 |
| H | 43.5177820000 | 45.7281480000 | 29.2446500000 |
| H | 45.3308450000 | 43.5670550000 | 33.0133770000 |
| C | 45.7849360000 | 43.8750640000 | 33.9320460000 |
| H | 46.8713170000 | 43.8528880000 | 33.7746160000 |
| C | 45.3500350000 | 45.3272910000 | 34.2446200000 |
| H | 45.9364180000 | 45.7376850000 | 35.0719100000 |
| H | 45.5733040000 | 45.9499590000 | 33.3668250000 |
| C | 43.9122710000 | 45.4369890000 | 34.6231760000 |
| N | 42.8656860000 | 45.2031970000 | 33.7390230000 |

| | | | |
|---|---|---|---|
| C | 41.6929710000 | 45.3984650000 | 34.4048110000 |
| H | 40.7008950000 | 45.2886970000 | 33.9913900000 |
| N | 41.9354090000 | 45.7365890000 | 35.6577580000 |
| H | 42.9526030000 | 44.8875860000 | 32.7842310000 |
| C | 43.3041000000 | 45.7623510000 | 35.8105900000 |
| H | 43.7643800000 | 46.0222860000 | 36.7508150000 |
| C | 45.4203630000 | 42.9475780000 | 35.0859350000 |
| O | 44.6050060000 | 42.0306380000 | 34.9872950000 |
| N | 46.0675830000 | 43.2701690000 | 36.2326170000 |
| H | 46.6958430000 | 44.0627090000 | 36.2271580000 |
| C | 45.6898720000 | 42.8826420000 | 37.5750770000 |
| H | 44.8575310000 | 42.1773220000 | 37.5191540000 |
| C | 46.9000410000 | 42.2308320000 | 38.3048600000 |
| H | 46.6022060000 | 42.0648710000 | 39.3536360000 |
| C | 47.2973270000 | 40.9079320000 | 37.6711220000 |
| H | 48.1787500000 | 40.5078660000 | 38.1794220000 |
| H | 46.4837620000 | 40.1794320000 | 37.7491680000 |
| H | 47.5409710000 | 41.0535270000 | 36.6149230000 |
| O | 48.0248180000 | 43.0929080000 | 38.2323640000 |
| H | 47.7655720000 | 43.9712870000 | 38.5610280000 |
| C | 45.3263390000 | 44.1735770000 | 38.3284020000 |
| O | 45.9350490000 | 45.2192000000 | 38.0620990000 |
| N | 44.4150340000 | 44.0909130000 | 39.3174680000 |
| H | 43.9430260000 | 43.2136360000 | 39.4932140000 |
| C | 44.0640170000 | 45.2538890000 | 40.1299790000 |
| H | 44.1199450000 | 46.1263380000 | 39.4784380000 |
| C | 42.6820890000 | 45.1442360000 | 40.7640730000 |
| H | 42.5542390000 | 44.1940280000 | 41.2981730000 |
| H | 42.6150810000 | 45.9032240000 | 41.5528900000 |
| C | 41.4548930000 | 45.3501180000 | 39.8706180000 |
| O | 41.6180080000 | 45.5908660000 | 38.6390190000 |
| O | 40.3381680000 | 45.2805980000 | 40.4700940000 |
| H | 44.7764550000 | 45.4078750000 | 40.9404710000 |
| H | 36.1386980000 | 40.7413340000 | 41.5253420000 |
| C | 37.1400160000 | 40.3200660000 | 41.4359730000 |
| H | 36.9920850000 | 39.2336870000 | 41.5006050000 |
| C | 37.7815560000 | 40.6080350000 | 40.0685310000 |
| H | 38.6803650000 | 39.9854830000 | 39.9567980000 |
| H | 37.0965800000 | 40.3259480000 | 39.2655360000 |
| C | 38.2033610000 | 42.0397130000 | 39.7842480000 |
| O | 37.8741030000 | 42.5968670000 | 38.7229580000 |
| N | 38.9811430000 | 42.6414070000 | 40.7019050000 |
| H | 39.3707310000 | 43.5598780000 | 40.5176120000 |
| H | 39.2613460000 | 42.2253490000 | 41.5785720000 |
| H | 37.7810680000 | 40.5604610000 | 42.2841320000 |
| H | 42.7023990000 | 40.1715690000 | 33.7695640000 |
| C | 41.7039930000 | 40.3110030000 | 33.3550100000 |
| H | 41.4997820000 | 39.4368090000 | 32.7261590000 |
| C | 40.6490280000 | 40.3790540000 | 34.4803190000 |
| H | 39.6449330000 | 40.4421370000 | 34.0475070000 |
| H | 40.6774120000 | 39.4368210000 | 35.0463440000 |
| C | 40.8152900000 | 41.5327120000 | 35.4219840000 |
| N | 42.0349240000 | 41.8525220000 | 36.0009870000 |

| | | | |
|---|---|---|---|
| H | 42.9561100000 | 41.5351300000 | 35.7049080000 |
| C | 41.8511160000 | 42.9201690000 | 36.8119000000 |
| H | 42.6472690000 | 43.4343790000 | 37.3252830000 |
| N | 40.5833780000 | 43.3021900000 | 36.8110080000 |
| C | 39.9243150000 | 42.4386680000 | 35.9525990000 |
| H | 38.8606810000 | 42.5249640000 | 35.7887680000 |
| H | 41.6373830000 | 41.1898020000 | 32.7136140000 |
| Fe | 40.1780920000 | 45.4524670000 | 37.0166470000 |
| O | 38.5538480000 | 45.1688390000 | 38.4780470000 |
| H | 38.3158840000 | 44.1932390000 | 38.5163520000 |
| H | 39.0446640000 | 45.3492370000 | 39.3205140000 |
| O | 39.1123280000 | 46.1285220000 | 35.6151020000 |
| H | 38.1675410000 | 45.9385250000 | 35.7791700000 |
| C | 42.2220410000 | 49.2576720000 | 35.4384900000 |
| C | 42.3482880000 | 48.7401830000 | 34.1399810000 |
| C | 43.5915030000 | 48.3781360000 | 33.6421260000 |
| C | 44.7323550000 | 48.5836090000 | 34.4239840000 |
| C | 44.6073510000 | 49.0990930000 | 35.7101350000 |
| C | 43.3527390000 | 49.4213720000 | 36.2578220000 |
| H | 41.4535740000 | 48.6113400000 | 33.5399830000 |
| H | 43.6709240000 | 47.9419200000 | 32.6538330000 |
| H | 45.7142670000 | 48.3137600000 | 34.0435520000 |
| H | 45.4931560000 | 49.2342090000 | 36.3253460000 |
| C | 43.2990530000 | 49.9049600000 | 37.6907290000 |
| H | 42.3210370000 | 50.3357060000 | 37.9181100000 |
| H | 44.0345300000 | 50.7095020000 | 37.8183300000 |
| C | 43.6117280000 | 48.7747830000 | 38.6895650000 |
| H | 44.6063090000 | 48.3520730000 | 38.5078930000 |
| H | 42.8799680000 | 47.9683280000 | 38.6053030000 |
| H | 43.5908190000 | 49.1598660000 | 39.7148380000 |
| C | 40.8545800000 | 49.7613010000 | 35.7902340000 |
| O | 40.3022230000 | 50.6209340000 | 35.0759960000 |
| N | 40.2334650000 | 49.3154710000 | 36.8986180000 |
| C | 38.8093040000 | 49.5417680000 | 37.3146360000 |
| C | 38.6798270000 | 51.0347370000 | 37.6562160000 |
| C | 37.8768920000 | 49.1284880000 | 36.1670210000 |
| C | 38.5376220000 | 48.6946520000 | 38.5632610000 |
| H | 39.3408690000 | 51.3030250000 | 38.4870720000 |
| H | 38.9154970000 | 51.6598860000 | 36.7934540000 |
| H | 37.6477680000 | 51.2377430000 | 37.9588100000 |
| H | 36.8390240000 | 49.1924860000 | 36.5151550000 |
| H | 37.9872790000 | 49.7984080000 | 35.3120200000 |
| H | 38.1124970000 | 48.1056550000 | 35.8548730000 |
| H | 37.5443130000 | 48.9630650000 | 38.9357170000 |
| H | 38.5424410000 | 47.6278570000 | 38.3375430000 |
| H | 39.2631900000 | 48.8938850000 | 39.3569990000 |
| F | 40.8306270000 | 48.1763140000 | 37.4458150000 |
| O | 36.4678940000 | 45.9610060000 | 36.8003140000 |
| H | 37.0523080000 | 45.7527020000 | 37.5583400000 |
| H | 36.3562110000 | 46.9215850000 | 36.8441800000 |
| O | 40.6310260000 | 43.7006730000 | 42.7945960000 |
| H | 40.3909040000 | 44.0810000000 | 43.6520750000 |
| H | 40.4649870000 | 44.4142680000 | 42.1417640000 |

```
O     41.5576090000   52.0734010000   33.2388760000
H     41.1357940000   52.9407000000   33.3372820000
H     41.0926200000   51.4733300000   33.8921120000
```

**$His_3Wat_2$_FeF_theozyme**

Charge: 3
Multiplicity: 5

```
B3LYP-D3(BJ)/6-31G(d)-SDD SCF energy (au):              -2898.45553725
B3LYP-D3(BJ)/6-31G(d)-SDD enthalpy (au):                -2897.25744725
B3LYP-D3(BJ)/6-31G(d)-SDD free energy (au):             -2897.43442025
B3LYP-D3(BJ)/def2-TZVP/SMD SCF energy (au):             -4039.91226972
B3LYP-D3(BJ)/def2-TZVP/SMD enthalpy (au):               -4038.71417972
B3LYP-D3(BJ)/def2-TZVP/SMD free energy (au):            -4038.89115272
B3LYP-D3(BJ)/def2-TZVP/SMD free energy (quasi-harmonic) (au): -4038.87283447
```

Cartesian coordinates

```
ATOM    X               Y               Z
H     50.4463900000   18.1352380000   40.5491810000
C     51.0759990000   18.9330260000   40.8839960000
H     50.6793990000   19.4297890000   41.7447000000
C     51.2268060000   19.9765890000   39.7422130000
C     51.3938080000   21.3859220000   40.2225600000
C     50.7252910000   22.5383650000   39.8978060000
N     52.3265160000   21.7784080000   41.1741030000
C     52.2148750000   23.1082850000   41.3826300000
N     51.2454170000   23.6075940000   40.6225390000
H     52.0449570000   18.5043540000   41.1613940000
H     52.0632360000   19.7050500000   39.0853190000
H     50.3335870000   19.9718220000   39.1098730000
H     52.9875030000   21.1649000000   41.6329450000
H     49.9453510000   22.6712910000   39.1643040000
H     52.8441750000   23.6665490000   42.0579120000
H     46.2257260000   28.1487280000   45.7837450000
C     46.7069880000   27.8000540000   44.8939800000
H     46.0294720000   27.2078360000   44.3150500000
C     48.0774520000   27.1375970000   45.0362480000
C     48.7832830000   27.1098890000   43.7082210000
C     49.7704090000   26.2690460000   43.1705780000
N     48.5586180000   28.0414110000   42.7306680000
C     49.4045690000   27.8026920000   41.6883330000
N     50.1298670000   26.6976190000   41.9245370000
H     46.8269930000   28.7333850000   44.3333340000
H     48.6932250000   27.6954030000   45.7549750000
H     48.0056230000   26.1168100000   45.4250050000
H     47.9491800000   28.8715240000   42.7990610000
H     50.1980900000   25.3802400000   43.6127960000
H     49.3264820000   28.2953520000   40.7352770000
H     43.0995510000   29.3205140000   42.0312280000
C     43.5559960000   29.7580370000   41.1680110000
H     42.8881030000   30.5217770000   40.7643040000
C     44.9349550000   30.3581750000   41.5682110000
H     45.6776160000   29.5548370000   41.5244240000
H     45.2328650000   31.0914430000   40.8144740000
```

| | | | |
|---|---|---|---|
| C | 45.0353460000 | 30.9708710000 | 42.9660780000 |
| H | 44.3826570000 | 31.8435430000 | 43.0748090000 |
| H | 44.7124670000 | 30.2395840000 | 43.7207120000 |
| C | 46.4775970000 | 31.3372850000 | 43.2961980000 |
| O | 47.4475760000 | 30.5946590000 | 43.0363490000 |
| N | 46.6627300000 | 32.5220360000 | 43.9040750000 |
| H | 45.8890050000 | 33.1205260000 | 44.1578440000 |
| H | 47.5887910000 | 32.7823470000 | 44.2166110000 |
| C | 43.9419700000 | 28.7781530000 | 40.0441780000 |
| O | 44.3516380000 | 29.2020820000 | 38.9683910000 |
| N | 44.0007740000 | 27.4422170000 | 40.3729160000 |
| C | 44.2630090000 | 26.4159910000 | 39.3470010000 |
| H | 43.3155570000 | 26.1085120000 | 38.9562240000 |
| C | 45.0449730000 | 25.2032480000 | 39.9015770000 |
| C | 46.5031050000 | 25.2235640000 | 39.5799410000 |
| C | 47.6309190000 | 25.4513010000 | 40.3236610000 |
| N | 46.9749050000 | 24.9755200000 | 38.2978240000 |
| C | 48.3235540000 | 25.0454580000 | 38.2978200000 |
| N | 48.7599060000 | 25.3394960000 | 39.5184930000 |
| H | 43.4312070000 | 27.1454000000 | 41.1567250000 |
| H | 44.8030100000 | 26.9216520000 | 38.5449490000 |
| H | 44.9235230000 | 25.1424400000 | 40.9875400000 |
| H | 44.6057350000 | 24.2830280000 | 39.4975810000 |
| H | 46.4097120000 | 24.7187380000 | 37.4983530000 |
| H | 47.7032290000 | 25.6526480000 | 41.3803070000 |
| H | 48.9276470000 | 24.8007980000 | 37.4370970000 |
| H | 48.4060690000 | 38.5789460000 | 37.9572130000 |
| C | 49.2070030000 | 38.8929080000 | 37.3210340000 |
| H | 48.8580890000 | 39.8190460000 | 36.8497580000 |
| C | 50.5176970000 | 39.1785140000 | 38.0627950000 |
| H | 51.2460890000 | 39.5807670000 | 37.3507670000 |
| H | 50.3626040000 | 39.9656870000 | 38.8123350000 |
| C | 51.1449330000 | 37.9539560000 | 38.7421530000 |
| H | 51.2389170000 | 37.1302650000 | 38.0195850000 |
| H | 52.1625050000 | 38.1928470000 | 39.0753930000 |
| C | 50.3371460000 | 37.4893350000 | 39.9569150000 |
| H | 50.2557880000 | 38.2948180000 | 40.6919180000 |
| H | 49.3119660000 | 37.2115690000 | 39.6949760000 |
| N | 50.9543800000 | 36.3483390000 | 40.6547600000 |
| H | 51.7253180000 | 36.5654870000 | 41.2735070000 |
| C | 50.7842720000 | 35.0629880000 | 40.3385900000 |
| N | 49.9446760000 | 34.6850670000 | 39.3643200000 |
| H | 49.6167650000 | 35.3469260000 | 38.6771290000 |
| H | 49.8170880000 | 33.6919250000 | 39.1911390000 |
| N | 51.4373980000 | 34.1046360000 | 41.0175320000 |
| H | 51.9119910000 | 34.3230750000 | 41.8804840000 |
| H | 51.2566730000 | 33.1309840000 | 40.7652470000 |
| H | 49.3592450000 | 38.1981040000 | 36.5217270000 |
| Fe | 50.7998110000 | 25.5813340000 | 40.0577790000 |
| O | 53.2174970000 | 26.2620060000 | 40.5173470000 |
| H | 52.7351630000 | 26.9697590000 | 40.0233840000 |
| N | 49.9162300000 | 29.7303970000 | 38.9139520000 |
| C | 51.9202780000 | 30.1171590000 | 40.1810520000 |

| | | | |
|---|---|---|---|
| C | 53.0630550000 | 30.0633930000 | 39.3934460000 |
| C | 54.2810850000 | 29.6560430000 | 39.9474200000 |
| C | 54.3678610000 | 29.2851950000 | 41.3008300000 |
| C | 53.2368040000 | 29.3259840000 | 42.0954640000 |
| C | 51.9880520000 | 29.7569980000 | 41.5636050000 |
| C | 50.6264640000 | 30.6286430000 | 39.5904110000 |
| C | 48.5785990000 | 29.9392980000 | 38.2755270000 |
| C | 48.0821820000 | 28.5464030000 | 37.8660070000 |
| C | 47.5990010000 | 30.5466400000 | 39.2887590000 |
| C | 48.7408570000 | 30.8413010000 | 37.0454140000 |
| C | 50.8356470000 | 29.8484270000 | 42.3784950000 |
| C | 50.8021090000 | 29.6045040000 | 43.8377180000 |
| O | 50.3174730000 | 31.8208490000 | 39.8023440000 |
| F | 51.2402090000 | 27.3623250000 | 39.3184040000 |
| H | 53.0089770000 | 30.3393210000 | 38.3453710000 |
| H | 55.1703620000 | 29.6293070000 | 39.3250100000 |
| H | 55.3217350000 | 28.9806350000 | 41.7188850000 |
| H | 53.3042710000 | 29.0511580000 | 43.1428430000 |
| H | 49.9591370000 | 30.3266480000 | 41.9507230000 |
| H | 50.3411570000 | 28.7960290000 | 38.8496390000 |
| H | 50.8544380000 | 28.5224600000 | 44.0567130000 |
| H | 51.6625810000 | 30.0579500000 | 44.3431460000 |
| H | 49.8769070000 | 29.9901780000 | 44.2676250000 |
| H | 48.7724900000 | 28.0659150000 | 37.1636250000 |
| H | 47.9619360000 | 27.8950970000 | 38.7382090000 |
| H | 47.1052610000 | 28.6384780000 | 37.3843510000 |
| H | 47.8526610000 | 31.5798740000 | 39.5331090000 |
| H | 46.5849540000 | 30.5084660000 | 38.8812420000 |
| H | 47.5956270000 | 29.9707170000 | 40.2197790000 |
| H | 49.1455280000 | 31.8177640000 | 37.3275300000 |
| H | 49.4124000000 | 30.3863920000 | 36.3104270000 |
| H | 47.7682750000 | 30.9984930000 | 36.5695620000 |
| O | 51.6037600000 | 24.9209500000 | 38.0390990000 |
| H | 52.5307660000 | 24.5837190000 | 38.2343950000 |
| H | 51.7128180000 | 25.7501680000 | 37.5445670000 |
| O | 53.9865800000 | 24.1764040000 | 38.8816230000 |
| H | 54.0786240000 | 24.8957000000 | 39.5380270000 |
| H | 54.8015060000 | 24.1568730000 | 38.3558670000 |
| O | 49.8684580000 | 23.0756720000 | 36.7451260000 |
| H | 50.6937410000 | 23.5187430000 | 37.0138630000 |
| H | 50.0935180000 | 22.5190770000 | 35.9842810000 |
| H | 53.3593820000 | 26.6189650000 | 41.4065430000 |

**$His_3Wat_2$_Fe_theozyme**
Charge: 3
Multiplicity: 5
B3LYP-D3(BJ)/6-31G(d)-SDD SCF energy (au): -2898.39635455
B3LYP-D3(BJ)/6-31G(d)-SDD enthalpy (au): -2897.19775655
B3LYP-D3(BJ)/6-31G(d)-SDD free energy (au): -2897.38135555
B3LYP-D3(BJ)/def2-TZVP/SMD SCF energy (au): -4039.85390262
B3LYP-D3(BJ)/def2-TZVP/SMD enthalpy (au): -4038.65530462
B3LYP-D3(BJ)/def2-TZVP/SMD free energy (au): -4038.83890362
B3LYP-D3(BJ)/def2-TZVP/SMD free energy (quasi-harmonic) (au): -4038.81484307

Cartesian coordinates

| ATOM | X | Y | Z |
|---|---|---|---|
| H | 50.4463830000 | 18.1352940000 | 40.5491870000 |
| C | 51.0760190000 | 18.9328850000 | 40.8839940000 |
| H | 50.6794050000 | 19.4298360000 | 41.7446950000 |
| C | 51.1468560000 | 19.9004970000 | 39.6716580000 |
| C | 51.5196870000 | 21.3088880000 | 39.9899240000 |
| C | 50.7500390000 | 22.4445440000 | 40.0672880000 |
| N | 52.8003480000 | 21.7331930000 | 40.3115630000 |
| C | 52.7824880000 | 23.0605590000 | 40.5651060000 |
| N | 51.5436950000 | 23.5274390000 | 40.4311700000 |
| H | 52.0596990000 | 18.5285220000 | 41.1399980000 |
| H | 51.8260650000 | 19.4951440000 | 38.9114120000 |
| H | 50.1589970000 | 19.9444170000 | 39.2009690000 |
| H | 53.6206190000 | 21.1396560000 | 40.3495780000 |
| H | 49.6856250000 | 22.5533060000 | 39.9208130000 |
| H | 53.6627840000 | 23.6481460000 | 40.7737830000 |
| H | 46.2257400000 | 28.1487440000 | 45.7837720000 |
| C | 46.7069460000 | 27.8000210000 | 44.8939860000 |
| H | 46.0294650000 | 27.2078320000 | 44.3150560000 |
| C | 48.0552520000 | 27.1151870000 | 45.1427630000 |
| C | 48.8412130000 | 27.0212590000 | 43.8711560000 |
| C | 49.5991030000 | 26.0391080000 | 43.2834810000 |
| N | 48.8915790000 | 28.0816640000 | 42.9765550000 |
| C | 49.6312050000 | 27.7414650000 | 41.9104000000 |
| N | 50.0875660000 | 26.5018740000 | 42.0627540000 |
| H | 46.8535470000 | 28.7246650000 | 44.3171920000 |
| H | 48.6287960000 | 27.6865600000 | 45.8842500000 |
| H | 47.9239290000 | 26.1120480000 | 45.5588380000 |
| H | 48.4193520000 | 28.9710830000 | 43.0737760000 |
| H | 49.8144900000 | 25.0432780000 | 43.6421800000 |
| H | 49.7845070000 | 28.3743750000 | 41.0531020000 |
| H | 43.0996680000 | 29.3204490000 | 42.0312650000 |
| C | 43.5558780000 | 29.7581050000 | 41.1679350000 |
| H | 42.7437900000 | 30.0450530000 | 40.4856890000 |
| C | 44.4028340000 | 30.9713570000 | 41.5502280000 |
| H | 45.1867900000 | 30.6605020000 | 42.2495120000 |
| H | 44.9242530000 | 31.3449900000 | 40.6660790000 |
| C | 43.5902320000 | 32.1029040000 | 42.1734480000 |
| H | 42.7950210000 | 32.4290470000 | 41.4890240000 |
| H | 43.0839460000 | 31.7694830000 | 43.0893870000 |
| C | 44.4713020000 | 33.3094870000 | 42.5054420000 |
| O | 45.6753110000 | 33.3436930000 | 42.2549070000 |
| N | 43.8257650000 | 34.3451220000 | 43.0967130000 |
| H | 42.8349960000 | 34.3315300000 | 43.2898950000 |
| H | 44.3485080000 | 35.1775270000 | 43.3319740000 |
| C | 44.4153550000 | 28.6917190000 | 40.4337890000 |
| O | 45.5050600000 | 28.9528010000 | 39.9226290000 |
| N | 43.8825450000 | 27.4295730000 | 40.3585890000 |
| C | 44.2630890000 | 26.4160490000 | 39.3468240000 |
| H | 43.3154910000 | 26.1084650000 | 38.9563840000 |
| C | 45.0994750000 | 25.2256200000 | 39.8757860000 |

| | | | |
|---|---|---|---|
| C | 46.5706140000 | 25.3148250000 | 39.6032180000 |
| C | 47.6717700000 | 25.3420000000 | 40.4195100000 |
| N | 47.0938720000 | 25.3107690000 | 38.3158050000 |
| C | 48.4442240000 | 25.3370280000 | 38.3826440000 |
| N | 48.8347120000 | 25.3508720000 | 39.6529440000 |
| H | 42.9562640000 | 27.3135100000 | 40.7510780000 |
| H | 44.7981500000 | 26.9437780000 | 38.5528710000 |
| H | 44.9485810000 | 25.1260940000 | 40.9539580000 |
| H | 44.7170850000 | 24.3009820000 | 39.4239450000 |
| H | 46.5547860000 | 25.2882600000 | 37.4598510000 |
| H | 47.7058600000 | 25.3563430000 | 41.4977990000 |
| H | 49.0961190000 | 25.3088250000 | 37.5227220000 |
| H | 48.4060880000 | 38.5788630000 | 37.9572150000 |
| C | 49.2070090000 | 38.8930950000 | 37.3209780000 |
| H | 48.8387400000 | 39.8101190000 | 36.8455320000 |
| C | 50.5217700000 | 39.2098510000 | 38.0453890000 |
| H | 51.1639920000 | 39.7847560000 | 37.3699570000 |
| H | 50.3325900000 | 39.8611620000 | 38.9083530000 |
| C | 51.3072030000 | 37.9702960000 | 38.4887830000 |
| H | 51.4139990000 | 37.2848030000 | 37.6359610000 |
| H | 52.3241180000 | 38.2519040000 | 38.7877550000 |
| C | 50.6446990000 | 37.2312720000 | 39.6550460000 |
| H | 50.7048880000 | 37.8181150000 | 40.5751460000 |
| H | 49.5797890000 | 37.0498070000 | 39.4810940000 |
| N | 51.2852140000 | 35.9347600000 | 39.9324940000 |
| H | 52.1644920000 | 35.9663270000 | 40.4332010000 |
| C | 51.0255340000 | 34.8102310000 | 39.2586250000 |
| N | 50.0274080000 | 34.7456920000 | 38.3674910000 |
| H | 49.6083850000 | 35.5818670000 | 37.9893040000 |
| H | 49.8128530000 | 33.8523820000 | 37.9368550000 |
| N | 51.7527280000 | 33.7020880000 | 39.4775770000 |
| H | 52.3302740000 | 33.6224360000 | 40.3013010000 |
| H | 51.4580580000 | 32.8470920000 | 39.0062920000 |
| H | 49.3592250000 | 38.1980050000 | 36.5217850000 |
| Fe | 50.8358130000 | 25.4791760000 | 40.4354460000 |
| O | 52.8978780000 | 26.0967430000 | 41.7226970000 |
| H | 52.5713930000 | 26.9943620000 | 41.9199050000 |
| N | 48.9888840000 | 29.8556680000 | 38.1480110000 |
| C | 50.7206570000 | 30.3475220000 | 39.8028810000 |
| C | 51.7540040000 | 29.4265620000 | 39.5997490000 |
| C | 52.6597820000 | 29.1559130000 | 40.6268860000 |
| C | 52.5072240000 | 29.7863190000 | 41.8655070000 |
| C | 51.4536870000 | 30.6824770000 | 42.0683820000 |
| C | 50.5494410000 | 30.9954110000 | 41.0438790000 |
| C | 49.8830870000 | 30.7535790000 | 38.6262510000 |
| C | 47.8423600000 | 30.0913180000 | 37.2056820000 |
| C | 47.1357910000 | 28.7452500000 | 36.9969450000 |
| C | 46.8862450000 | 31.1213650000 | 37.8220820000 |
| C | 48.4434930000 | 30.5737300000 | 35.8757940000 |
| C | 49.3959430000 | 31.9439720000 | 41.2930260000 |
| C | 48.1439760000 | 31.2376000000 | 41.8439720000 |
| O | 50.0745040000 | 31.8451240000 | 38.0678850000 |
| F | 48.7137270000 | 28.8436190000 | 39.0934270000 |

| | | | |
|---|---|---|---|
| H | 51.8581060000 | 28.9368670000 | 38.6372160000 |
| H | 53.4939000000 | 28.4866700000 | 40.4367970000 |
| H | 53.2200520000 | 29.6072070000 | 42.6660680000 |
| H | 51.3406940000 | 31.1668330000 | 43.0348730000 |
| H | 49.7088120000 | 32.7179520000 | 42.0021720000 |
| H | 49.1285110000 | 32.4611330000 | 40.3679560000 |
| H | 47.8335230000 | 30.4056360000 | 41.1998720000 |
| H | 48.3467020000 | 30.8646990000 | 42.8582530000 |
| H | 47.3066170000 | 31.9377780000 | 41.9237820000 |
| H | 47.8393850000 | 27.9857950000 | 36.6391880000 |
| H | 46.6562140000 | 28.4099590000 | 37.9174900000 |
| H | 46.3676450000 | 28.8870900000 | 36.2313290000 |
| H | 47.3991320000 | 32.0694500000 | 38.0078070000 |
| H | 46.0669230000 | 31.3145680000 | 37.1224750000 |
| H | 46.4579790000 | 30.7391370000 | 38.7520090000 |
| H | 48.9434700000 | 31.5363910000 | 35.9780010000 |
| H | 49.1588850000 | 29.8443680000 | 35.4833380000 |
| H | 47.6302670000 | 30.6794210000 | 35.1518800000 |
| O | 51.8674280000 | 26.1424870000 | 38.7826480000 |
| H | 52.8553690000 | 26.1452030000 | 38.9297490000 |
| H | 51.6962850000 | 25.4844720000 | 38.0571460000 |
| O | 54.4258360000 | 25.9141980000 | 39.4854440000 |
| H | 54.3486470000 | 26.0918830000 | 40.4410670000 |
| H | 55.2562660000 | 26.3062270000 | 39.1724910000 |
| O | 51.1461530000 | 24.1318340000 | 37.1385940000 |
| H | 51.5508500000 | 23.9020590000 | 36.2860230000 |
| H | 51.1367630000 | 23.3134120000 | 37.6649160000 |
| H | 52.8764310000 | 25.6308160000 | 42.5761030000 |

**His$_3$WatOH_FeF_theozyme**
Charge: 2
Multiplicity: 5
B3LYP-D3(BJ)/6-31G(d)-SDD SCF energy (au): -2898.20021760
B3LYP-D3(BJ)/6-31G(d)-SDD enthalpy (au): -2897.01463660
B3LYP-D3(BJ)/6-31G(d)-SDD free energy (au): -2897.19253760
B3LYP-D3(BJ)/def2-TZVP/SMD SCF energy (au): -4039.45699808
B3LYP-D3(BJ)/def2-TZVP/SMD enthalpy (au): -4038.27141708
B3LYP-D3(BJ)/def2-TZVP/SMD free energy (au): -4038.44931808
B3LYP-D3(BJ)/def2-TZVP/SMD free energy (quasi-harmonic) (au): -4038.42895048

Cartesian coordinates

| ATOM | X | Y | Z |
|---|---|---|---|
| H | 50.4463900000 | 18.1352380000 | 40.5491810000 |
| C | 51.0759990000 | 18.9330260000 | 40.8839960000 |
| H | 50.6793990000 | 19.4297890000 | 41.7447000000 |
| C | 51.2173580000 | 19.9841330000 | 39.7461770000 |
| C | 51.3113050000 | 21.3978930000 | 40.2320760000 |
| C | 50.6127030000 | 22.5253390000 | 39.8872000000 |
| N | 52.2073070000 | 21.8232950000 | 41.2082960000 |
| C | 52.0506630000 | 23.1491050000 | 41.4062350000 |
| N | 51.0828310000 | 23.6091300000 | 40.6184640000 |
| H | 52.0473820000 | 18.5084080000 | 41.1587450000 |
| H | 52.0836850000 | 19.7486280000 | 39.1142950000 |

| | | | |
|---|---|---|---|
| H | 50.3450370000 | 19.9424660000 | 39.0866760000 |
| H | 52.8850920000 | 21.2384990000 | 41.6784090000 |
| H | 49.8500460000 | 22.6354990000 | 39.1333600000 |
| H | 52.6482800000 | 23.7510210000 | 42.0719080000 |
| H | 46.2257260000 | 28.1487280000 | 45.7837450000 |
| C | 46.7069880000 | 27.8000540000 | 44.8939800000 |
| H | 46.0294720000 | 27.2078360000 | 44.3150500000 |
| C | 48.0971960000 | 27.1728470000 | 45.0320360000 |
| C | 48.7729580000 | 27.1125830000 | 43.6893910000 |
| C | 49.5607600000 | 26.1707020000 | 43.0682680000 |
| N | 48.6629150000 | 28.1478570000 | 42.7667580000 |
| C | 49.3510490000 | 27.8188380000 | 41.6584150000 |
| N | 49.9105930000 | 26.6235630000 | 41.8043520000 |
| H | 46.8099210000 | 28.7305830000 | 44.3217610000 |
| H | 48.7037440000 | 27.7745040000 | 45.7220510000 |
| H | 48.0532580000 | 26.1643150000 | 45.4547100000 |
| H | 48.1542430000 | 29.0280610000 | 42.8876680000 |
| H | 49.8895350000 | 25.2086270000 | 43.4311100000 |
| H | 49.4112400000 | 28.4340890000 | 40.7794010000 |
| H | 43.0995510000 | 29.3205140000 | 42.0312280000 |
| C | 43.5559960000 | 29.7580370000 | 41.1680110000 |
| H | 42.8859430000 | 30.5232500000 | 40.7692940000 |
| C | 44.9220370000 | 30.3615220000 | 41.5804850000 |
| H | 45.6590570000 | 29.5531370000 | 41.6245640000 |
| H | 45.2595060000 | 31.0445690000 | 40.7974380000 |
| C | 44.9481950000 | 31.0677430000 | 42.9357500000 |
| H | 44.2939710000 | 31.9469240000 | 42.9458460000 |
| H | 44.5797770000 | 30.3906760000 | 43.7197830000 |
| C | 46.3723620000 | 31.4536100000 | 43.3210380000 |
| O | 47.3518010000 | 30.7292010000 | 43.0908720000 |
| N | 46.5142050000 | 32.6418130000 | 43.9467600000 |
| H | 45.7218300000 | 33.2207290000 | 44.1840560000 |
| H | 47.4239560000 | 32.9041510000 | 44.3020860000 |
| C | 43.9008500000 | 28.7760010000 | 40.0334240000 |
| O | 44.1942690000 | 29.1925700000 | 38.9178730000 |
| N | 44.0291990000 | 27.4472400000 | 40.3718300000 |
| C | 44.2630090000 | 26.4159910000 | 39.3470010000 |
| H | 43.3155570000 | 26.1085120000 | 38.9562240000 |
| C | 45.0362300000 | 25.1919190000 | 39.8912230000 |
| C | 46.4958890000 | 25.1809360000 | 39.5722880000 |
| C | 47.6196740000 | 25.3376330000 | 40.3387940000 |
| N | 46.9757240000 | 24.9601480000 | 38.2873460000 |
| C | 48.3285090000 | 24.9859230000 | 38.3104260000 |
| N | 48.7538750000 | 25.2146110000 | 39.5461870000 |
| H | 43.5647080000 | 27.1424320000 | 41.2183480000 |
| H | 44.8035250000 | 26.9115700000 | 38.5386030000 |
| H | 44.9185770000 | 25.1258630000 | 40.9778000000 |
| H | 44.5774660000 | 24.2815900000 | 39.4850890000 |
| H | 46.4153100000 | 24.7676240000 | 37.4679900000 |
| H | 47.6851440000 | 25.5136500000 | 41.4001540000 |
| H | 48.9392730000 | 24.7690150000 | 37.4466420000 |
| H | 48.4060690000 | 38.5789460000 | 37.9572130000 |
| C | 49.2070030000 | 38.8929080000 | 37.3210340000 |

| | | | |
|---|---|---|---|
| H | 48.8723190000 | 39.8278260000 | 36.8572450000 |
| C | 50.5192500000 | 39.1431750000 | 38.0761160000 |
| H | 51.2545950000 | 39.5585890000 | 37.3784570000 |
| H | 50.3692310000 | 39.9092570000 | 38.8486000000 |
| C | 51.1291740000 | 37.8895460000 | 38.7194570000 |
| H | 51.2026080000 | 37.0849430000 | 37.9736230000 |
| H | 52.1550800000 | 38.1000380000 | 39.0464920000 |
| C | 50.3295960000 | 37.4011600000 | 39.9320980000 |
| H | 50.2871650000 | 38.1825240000 | 40.6964830000 |
| H | 49.2916410000 | 37.1688030000 | 39.6760700000 |
| N | 50.9120890000 | 36.2145410000 | 40.5729330000 |
| H | 51.7062690000 | 36.3691400000 | 41.1798400000 |
| C | 50.6836890000 | 34.9464530000 | 40.2004870000 |
| N | 49.8271900000 | 34.6587060000 | 39.2097260000 |
| H | 49.5757900000 | 35.3582130000 | 38.5285430000 |
| H | 49.6694810000 | 33.6750300000 | 38.9918410000 |
| N | 51.2850570000 | 33.9328300000 | 40.8335450000 |
| H | 51.7362500000 | 34.0731570000 | 41.7247530000 |
| H | 51.0498400000 | 32.9798560000 | 40.5228470000 |
| H | 49.3592450000 | 38.1981040000 | 36.5217270000 |
| Fe | 50.8367090000 | 25.6701770000 | 40.1435230000 |
| O | 52.5804900000 | 25.9489910000 | 40.8358240000 |
| H | 52.7629990000 | 26.8978730000 | 40.9529690000 |
| N | 49.6702060000 | 29.7416700000 | 38.7467500000 |
| C | 51.6329070000 | 30.1567980000 | 40.0565950000 |
| C | 52.6446580000 | 29.5957930000 | 39.2867670000 |
| C | 53.8061620000 | 29.0922030000 | 39.8824530000 |
| C | 53.9374690000 | 29.1350900000 | 41.2778080000 |
| C | 52.9290530000 | 29.6772130000 | 42.0619950000 |
| C | 51.7423170000 | 30.2171760000 | 41.4873820000 |
| C | 50.3914290000 | 30.6673520000 | 39.3864890000 |
| C | 48.3329470000 | 29.9233290000 | 38.1208580000 |
| C | 47.8178130000 | 28.5113830000 | 37.8043800000 |
| C | 47.3728280000 | 30.5925810000 | 39.1140080000 |
| C | 48.4714160000 | 30.7502340000 | 36.8358170000 |
| C | 50.6999060000 | 30.7269610000 | 42.3006280000 |
| C | 50.7052650000 | 30.6896280000 | 43.7959610000 |
| O | 50.0678750000 | 31.8751610000 | 39.4829650000 |
| F | 50.7238740000 | 27.2220380000 | 39.0634460000 |
| H | 52.5267040000 | 29.5502760000 | 38.2085770000 |
| H | 54.5986360000 | 28.6785510000 | 39.2671800000 |
| H | 54.8354800000 | 28.7483350000 | 41.7507720000 |
| H | 53.0463310000 | 29.6962300000 | 43.1398910000 |
| H | 49.8153220000 | 31.1310560000 | 41.8206350000 |
| H | 50.0525990000 | 28.7908490000 | 38.7563000000 |
| H | 50.9598370000 | 29.6922340000 | 44.1798510000 |
| H | 51.4381320000 | 31.3846870000 | 44.2336240000 |
| H | 49.7158970000 | 30.9521810000 | 44.1774640000 |
| H | 48.4897770000 | 27.9872620000 | 37.1159630000 |
| H | 47.7256560000 | 27.9139690000 | 38.7180280000 |
| H | 46.8274380000 | 28.5803910000 | 37.3461910000 |
| H | 47.6374240000 | 31.6363180000 | 39.2918270000 |
| H | 46.3499520000 | 30.5343730000 | 38.7318550000 |

H     47.3997440000   30.0691080000   40.0748140000
H     48.8901470000   31.7364320000   37.0552510000
H     49.1274520000   30.2478340000   36.1174170000
H     47.4903170000   30.8869010000   36.3698600000
O     51.6126800000   24.9517760000   38.1516190000
H     52.5697820000   24.7054630000   38.3799760000
H     51.6361000000   25.8471240000   37.7673120000
O     53.9584960000   24.4407810000   39.1581430000
H     53.6845110000   25.0481450000   39.9003620000
H     54.7768010000   24.8012990000   38.7835760000
O     50.0151320000   23.1417540000   36.6870860000
H     50.7709310000   23.6189460000   37.0798090000
H     50.3618100000   22.6958860000   35.9001190000

**$His_3$WatOH_Fe_theozyme**

Charge: 2
Multiplicity: 5
B3LYP-D3(BJ)/6-31G(d)-SDD SCF energy (au):               -2898.12129405
B3LYP-D3(BJ)/6-31G(d)-SDD enthalpy (au):                 -2896.93719805
B3LYP-D3(BJ)/6-31G(d)-SDD free energy (au):              -2897.11655105
B3LYP-D3(BJ)/def2-TZVP/SMD SCF energy (au):              -4039.37969258
B3LYP-D3(BJ)/def2-TZVP/SMD enthalpy (au):                -4038.19559658
B3LYP-D3(BJ)/def2-TZVP/SMD free energy (au):             -4038.37494958
B3LYP-D3(BJ)/def2-TZVP/SMD free energy (quasi-harmonic) (au): -4038.35223353

Cartesian coordinates

| ATOM | X | Y | Z |
|---|---|---|---|
| H | 50.4463980000 | 18.1352480000 | 40.5491820000 |
| C | 51.0759960000 | 18.9330110000 | 40.8839900000 |
| H | 50.6794020000 | 19.4297940000 | 41.7447040000 |
| C | 51.1394800000 | 19.9463450000 | 39.7088270000 |
| C | 51.4659100000 | 21.3642670000 | 40.0669470000 |
| C | 50.6541790000 | 22.4691000000 | 40.2035930000 |
| N | 52.7437000000 | 21.8478860000 | 40.3103420000 |
| C | 52.6729370000 | 23.1802530000 | 40.5634670000 |
| N | 51.4146130000 | 23.5897050000 | 40.5019670000 |
| H | 52.0634470000 | 18.5358710000 | 41.1366140000 |
| H | 51.8370720000 | 19.5867470000 | 38.9412840000 |
| H | 50.1545770000 | 19.9738640000 | 39.2286780000 |
| H | 53.5928480000 | 21.2984460000 | 40.3043970000 |
| H | 49.5795710000 | 22.5286220000 | 40.1113950000 |
| H | 53.5322620000 | 23.8198510000 | 40.7017470000 |
| H | 46.2257140000 | 28.1487440000 | 45.7837420000 |
| C | 46.7070450000 | 27.7999780000 | 44.8940520000 |
| H | 46.0294340000 | 27.2078880000 | 44.3149890000 |
| C | 47.9600900000 | 26.9755720000 | 45.2785370000 |
| C | 48.9336300000 | 26.7934890000 | 44.1569860000 |
| C | 49.1519670000 | 25.7935480000 | 43.2387430000 |
| N | 49.8453700000 | 27.7825510000 | 43.8123330000 |
| C | 50.5546890000 | 27.3698580000 | 42.7363810000 |
| N | 50.1517690000 | 26.1687020000 | 42.3500010000 |
| H | 46.9849530000 | 28.6771070000 | 44.2996990000 |
| H | 48.4744040000 | 27.4652270000 | 46.1155330000 |

| | | | |
|---|---|---|---|
| H | 47.6545730000 | 25.9915280000 | 45.6486220000 |
| H | 49.9703170000 | 28.6741180000 | 44.2731070000 |
| H | 48.6643480000 | 24.8324710000 | 43.1667140000 |
| H | 51.3166510000 | 27.9422910000 | 42.2392300000 |
| H | 43.0995770000 | 29.3205290000 | 42.0312600000 |
| C | 43.5559650000 | 29.7579950000 | 41.1679510000 |
| H | 42.7356390000 | 30.0598120000 | 40.5002590000 |
| C | 44.3955470000 | 30.9720560000 | 41.5597900000 |
| H | 45.1824990000 | 30.6570700000 | 42.2499840000 |
| H | 44.9077370000 | 31.3602180000 | 40.6760620000 |
| C | 43.5639770000 | 32.0778020000 | 42.2053080000 |
| H | 42.7490970000 | 32.3895120000 | 41.5374330000 |
| H | 43.0801890000 | 31.7161670000 | 43.1238060000 |
| C | 44.4082680000 | 33.3024990000 | 42.5584570000 |
| O | 45.6318060000 | 33.3264130000 | 42.4553190000 |
| N | 43.6989280000 | 34.3767860000 | 42.9983950000 |
| H | 42.6960430000 | 34.3584890000 | 43.1083070000 |
| H | 44.2000980000 | 35.2042890000 | 43.2894240000 |
| C | 44.3984680000 | 28.7193850000 | 40.3838990000 |
| O | 45.4292660000 | 29.0257530000 | 39.7910080000 |
| N | 43.8999350000 | 27.4412750000 | 40.3510540000 |
| C | 44.2630290000 | 26.4159850000 | 39.3469960000 |
| H | 43.3155310000 | 26.1085250000 | 38.9562350000 |
| C | 45.0286330000 | 25.2019510000 | 39.9295330000 |
| C | 46.4946700000 | 25.1879390000 | 39.6455810000 |
| C | 47.5887600000 | 25.6029220000 | 40.3581510000 |
| N | 47.0182840000 | 24.6713900000 | 38.4666680000 |
| C | 48.3676590000 | 24.7749860000 | 38.5014990000 |
| N | 48.7470710000 | 25.3335550000 | 39.6450510000 |
| H | 43.0100970000 | 27.2975610000 | 40.8114590000 |
| H | 44.8367160000 | 26.9149440000 | 38.5637390000 |
| H | 44.8772600000 | 25.1782290000 | 41.0124680000 |
| H | 44.5840270000 | 24.2786260000 | 39.5372330000 |
| H | 46.4833000000 | 24.2648110000 | 37.7110750000 |
| H | 47.6235980000 | 26.0766520000 | 41.3254300000 |
| H | 49.0178810000 | 24.4153720000 | 37.7187250000 |
| H | 48.4060500000 | 38.5789320000 | 37.9572110000 |
| C | 49.2070490000 | 38.8929260000 | 37.3210370000 |
| H | 48.8644770000 | 39.8227310000 | 36.8518660000 |
| C | 50.5103860000 | 39.1712160000 | 38.0807030000 |
| H | 51.2397350000 | 39.6044040000 | 37.3873410000 |
| H | 50.3352270000 | 39.9346370000 | 38.8505360000 |
| C | 51.1553390000 | 37.9416330000 | 38.7342900000 |
| H | 51.3161220000 | 37.1585180000 | 37.9793460000 |
| H | 52.1492920000 | 38.2074510000 | 39.1153430000 |
| C | 50.3309410000 | 37.3821020000 | 39.8999240000 |
| H | 50.1480560000 | 38.1623830000 | 40.6440440000 |
| H | 49.3462380000 | 37.0267890000 | 39.5812940000 |
| N | 51.0045760000 | 36.2818410000 | 40.6049100000 |
| H | 51.7031430000 | 36.5411660000 | 41.2890090000 |
| C | 51.0158330000 | 35.0045750000 | 40.2022480000 |
| N | 50.3227630000 | 34.6104850000 | 39.1261790000 |
| H | 50.0024450000 | 35.2779940000 | 38.4421270000 |

| | | | |
|---|---|---|---|
| H | 50.3136500000 | 33.6191240000 | 38.8959700000 |
| N | 51.7148020000 | 34.0858560000 | 40.8832050000 |
| H | 51.9931170000 | 34.2517740000 | 41.8392020000 |
| H | 51.6751450000 | 33.1211780000 | 40.5449620000 |
| H | 49.3592180000 | 38.1980880000 | 36.5217260000 |
| Fe | 50.7680370000 | 25.6520110000 | 40.2696990000 |
| O | 51.9737540000 | 27.1785270000 | 40.2240820000 |
| H | 51.7315900000 | 27.7138870000 | 39.4515440000 |
| N | 49.8223320000 | 29.8431580000 | 39.5790070000 |
| C | 51.1273750000 | 30.6980780000 | 41.4777280000 |
| C | 52.3738150000 | 30.0792990000 | 41.6351090000 |
| C | 52.9752350000 | 30.0332370000 | 42.8913520000 |
| C | 52.3317170000 | 30.6135750000 | 43.9871160000 |
| C | 51.0970910000 | 31.2425970000 | 43.8183100000 |
| C | 50.4741940000 | 31.3069130000 | 42.5626580000 |
| C | 50.5746250000 | 30.8427040000 | 40.0914490000 |
| C | 48.9426340000 | 29.8593140000 | 38.3628350000 |
| C | 48.3380740000 | 28.4586980000 | 38.1998340000 |
| C | 47.8423370000 | 30.9119180000 | 38.5547970000 |
| C | 49.8425250000 | 30.1644470000 | 37.1541630000 |
| C | 49.1205730000 | 31.9631260000 | 42.4234790000 |
| C | 47.9731340000 | 31.0302490000 | 42.8396070000 |
| O | 50.8699110000 | 31.8389750000 | 39.4035880000 |
| F | 49.3609830000 | 28.9630200000 | 40.5682470000 |
| H | 52.8474900000 | 29.5964330000 | 40.7881990000 |
| H | 53.9363870000 | 29.5444120000 | 43.0131110000 |
| H | 52.7960670000 | 30.5866820000 | 44.9684870000 |
| H | 50.6064020000 | 31.7085540000 | 44.6698270000 |
| H | 49.0874420000 | 32.8645800000 | 43.0462780000 |
| H | 48.9563910000 | 32.2976050000 | 41.3932650000 |
| H | 47.9428460000 | 30.1408770000 | 42.2025610000 |
| H | 48.1037910000 | 30.7043510000 | 43.8785480000 |
| H | 47.0277780000 | 31.5700830000 | 42.7594310000 |
| H | 49.1166540000 | 27.6913100000 | 38.1769120000 |
| H | 47.6174580000 | 28.2331830000 | 38.9858300000 |
| H | 47.8074590000 | 28.4395450000 | 37.2430420000 |
| H | 48.2709270000 | 31.9074810000 | 38.7048600000 |
| H | 47.2130360000 | 30.9429570000 | 37.6599640000 |
| H | 47.1990830000 | 30.6515090000 | 39.3985330000 |
| H | 50.2982210000 | 31.1519390000 | 37.2174890000 |
| H | 50.6358960000 | 29.4150270000 | 37.0655510000 |
| H | 49.2281680000 | 30.1178390000 | 36.2501340000 |
| O | 51.4477690000 | 25.4333170000 | 38.0881110000 |
| H | 52.4212790000 | 25.4476130000 | 38.2703600000 |
| H | 51.2651510000 | 24.5665280000 | 37.6616570000 |
| O | 53.8935780000 | 25.6707860000 | 39.2594550000 |
| H | 53.3319470000 | 26.3701310000 | 39.7188240000 |
| H | 54.6619470000 | 26.1302670000 | 38.8886260000 |
| O | 50.6763820000 | 22.9559720000 | 37.1300440000 |
| H | 51.1487710000 | 22.5275880000 | 36.3990540000 |
| H | 50.8374580000 | 22.4097070000 | 37.9215180000 |

**12-A_theozyme**

Charge: 1
Multiplicity: 5
B3LYP-D3(BJ)/6-31G(d)-SDD SCF energy (au): -3132.02019251
B3LYP-D3(BJ)/6-31G(d)-SDD enthalpy (au): -3130.75165351
B3LYP-D3(BJ)/6-31G(d)-SDD free energy (au): -3130.96752751
B3LYP-D3(BJ)/def2-TZVP/SMD SCF energy (au): -4273.31794966
B3LYP-D3(BJ)/def2-TZVP/SMD enthalpy (au): -4272.04941066
B3LYP-D3(BJ)/def2-TZVP/SMD free energy (au): -4272.26528466
B3LYP-D3(BJ)/def2-TZVP/SMD free energy (quasi-harmonic) (au): -4272.23042515

Cartesian coordinates

| ATOM | X | Y | Z |
|---|---|---|---|
| H | 36.6176320000 | 24.2867520000 | 40.6866460000 |
| C | 36.2409030000 | 24.5417870000 | 39.6919670000 |
| H | 37.0603690000 | 24.9555310000 | 39.0977930000 |
| C | 35.1530830000 | 25.6387090000 | 39.8376860000 |
| C | 34.8540810000 | 26.4613860000 | 38.6137860000 |
| C | 35.3921710000 | 27.6429320000 | 38.1552420000 |
| N | 33.8399240000 | 26.1777390000 | 37.7072800000 |
| C | 33.7941660000 | 27.1644850000 | 36.7739600000 |
| N | 34.7291110000 | 28.0713380000 | 37.0162090000 |
| H | 35.8669220000 | 23.6420700000 | 39.1953140000 |
| H | 34.2222680000 | 25.1972630000 | 40.2158660000 |
| H | 35.4821650000 | 26.3415140000 | 40.6115750000 |
| H | 33.2148930000 | 25.3837820000 | 37.7420980000 |
| H | 36.1900720000 | 28.2303640000 | 38.5841860000 |
| H | 33.0560830000 | 27.2068670000 | 35.9870260000 |
| H | 29.7025460000 | 22.7110720000 | 37.5034010000 |
| C | 29.2400330000 | 23.2259100000 | 36.6470680000 |
| H | 28.9461840000 | 24.2328180000 | 36.9439750000 |
| C | 28.0795660000 | 22.5380320000 | 35.9473760000 |
| C | 27.6692710000 | 23.3702340000 | 34.7205710000 |
| O | 28.2803640000 | 24.4010160000 | 34.3660840000 |
| O | 26.6569620000 | 22.9448210000 | 34.0751320000 |
| H | 30.0531650000 | 23.3689370000 | 35.9271930000 |
| H | 28.3431640000 | 21.5320730000 | 35.6003510000 |
| H | 27.2024990000 | 22.4242520000 | 36.5942300000 |
| H | 25.6136270000 | 30.4503390000 | 42.5541610000 |
| C | 26.4050450000 | 30.7305890000 | 43.2551910000 |
| H | 26.1176250000 | 30.4443670000 | 44.2716060000 |
| C | 26.9041320000 | 32.1650410000 | 43.0765190000 |
| C | 27.8470050000 | 32.3214570000 | 41.8692950000 |
| O | 27.9549520000 | 31.3815500000 | 41.0286270000 |
| O | 28.4960920000 | 33.4057700000 | 41.7731560000 |
| H | 27.2333400000 | 30.0559060000 | 43.0022690000 |
| H | 26.0806360000 | 32.8793530000 | 42.9504880000 |
| H | 27.4593320000 | 32.5055660000 | 43.9586160000 |
| H | 41.2540500000 | 32.9343430000 | 33.7554340000 |
| C | 40.4492010000 | 32.4364000000 | 34.3021300000 |
| H | 40.7812270000 | 32.2475160000 | 35.3275560000 |
| C | 39.9620170000 | 31.1484880000 | 33.6123080000 |
| C | 38.7859560000 | 30.4867170000 | 34.2859730000 |
| C | 37.4755220000 | 30.8604020000 | 34.4965930000 |

| | | | |
|---|---|---|---|
| N | 38.8697810000 | 29.2118940000 | 34.8325250000 |
| C | 37.6522790000 | 28.8666880000 | 35.3323780000 |
| N | 36.7867700000 | 29.8474480000 | 35.1453620000 |
| H | 39.6217060000 | 33.1531550000 | 34.3690500000 |
| H | 39.6904210000 | 31.3802960000 | 32.5751230000 |
| H | 40.7855460000 | 30.4259470000 | 33.5507910000 |
| H | 39.6954370000 | 28.6293300000 | 34.8297580000 |
| H | 36.9642120000 | 31.7794130000 | 34.2527110000 |
| H | 37.4428710000 | 27.9296290000 | 35.8269200000 |
| H | 39.3343920000 | 34.2947870000 | 39.1395590000 |
| C | 38.6576600000 | 33.8739940000 | 39.8917060000 |
| H | 39.0647710000 | 34.1441680000 | 40.8703130000 |
| C | 38.5867600000 | 32.3248090000 | 39.8050950000 |
| C | 37.3324930000 | 31.7739200000 | 39.2127610000 |
| C | 37.0484590000 | 31.2501680000 | 37.9792780000 |
| N | 36.1337290000 | 31.6885440000 | 39.9115760000 |
| C | 35.1938640000 | 31.1292350000 | 39.1007840000 |
| N | 35.7244200000 | 30.8532450000 | 37.9214470000 |
| H | 37.6736890000 | 34.3395800000 | 39.7832610000 |
| H | 39.4294350000 | 31.9458920000 | 39.2183440000 |
| H | 38.7132380000 | 31.9006190000 | 40.8097050000 |
| H | 35.9837200000 | 31.9824330000 | 40.8666910000 |
| H | 37.7022380000 | 31.1489460000 | 37.1281330000 |
| H | 34.1582360000 | 30.9880800000 | 39.3858940000 |
| O | 29.8417420000 | 32.4804310000 | 39.4953330000 |
| H | 29.7707330000 | 33.1859880000 | 40.1733750000 |
| H | 29.1153730000 | 31.8971810000 | 39.8727050000 |
| H | 33.8251030000 | 43.3258380000 | 32.3960350000 |
| C | 32.7408300000 | 43.4797130000 | 32.3962030000 |
| H | 32.3584030000 | 43.2660970000 | 33.4029550000 |
| C | 32.0150470000 | 42.6263230000 | 31.3488040000 |
| C | 31.9474840000 | 41.1179360000 | 31.6422350000 |
| C | 33.3109740000 | 40.4075670000 | 31.6293880000 |
| N | 33.1930940000 | 38.9586980000 | 31.4043770000 |
| C | 33.1780260000 | 37.9754880000 | 32.3186150000 |
| N | 33.2796350000 | 38.2179420000 | 33.6308290000 |
| N | 33.0812700000 | 36.7016960000 | 31.9168260000 |
| H | 32.5734640000 | 44.5434240000 | 32.2003970000 |
| H | 30.9830580000 | 42.9840070000 | 31.2555510000 |
| H | 32.4765980000 | 42.7756510000 | 30.3629830000 |
| H | 31.4465670000 | 40.9397420000 | 32.6036010000 |
| H | 31.3168830000 | 40.6464510000 | 30.8771920000 |
| H | 33.9303710000 | 40.7962660000 | 30.8139750000 |
| H | 33.8801040000 | 40.5695120000 | 32.5488260000 |
| H | 32.9295560000 | 38.6851160000 | 30.4668160000 |
| H | 33.0598340000 | 39.1269650000 | 34.0076720000 |
| H | 33.2956760000 | 37.4049480000 | 34.2559090000 |
| H | 33.2561910000 | 36.4412640000 | 30.9581460000 |
| H | 33.0247250000 | 35.9660040000 | 32.6274630000 |
| Fe | 34.7380400000 | 30.1732650000 | 36.0735690000 |
| N | 32.7670480000 | 35.4079770000 | 36.6089630000 |
| O | 33.0876830000 | 35.4882290000 | 34.3904610000 |
| C | 31.7453520000 | 33.6649590000 | 35.2617480000 |

| | | | |
|---|---|---|---|
| C | 30.7934540000 | 33.4706420000 | 36.2832090000 |
| C | 29.9387930000 | 32.3755790000 | 36.2742080000 |
| C | 30.0333020000 | 31.4573040000 | 35.2261790000 |
| C | 30.9948240000 | 31.6257860000 | 34.2273440000 |
| C | 31.8748310000 | 32.7170850000 | 34.2191360000 |
| C | 32.6034610000 | 34.8683720000 | 35.3634290000 |
| C | 33.8989180000 | 35.1037930000 | 37.4857760000 |
| C | 35.1792210000 | 34.7742050000 | 36.6995560000 |
| C | 34.0924140000 | 36.3164780000 | 38.4071110000 |
| C | 33.4713640000 | 33.8696830000 | 38.3204740000 |
| C | 32.8998640000 | 32.7930590000 | 33.1121350000 |
| C | 32.2803730000 | 33.3044900000 | 31.8008120000 |
| F | 34.6615310000 | 31.9327640000 | 35.3872580000 |
| H | 30.7158430000 | 34.1936440000 | 37.0863690000 |
| H | 29.2374030000 | 32.2483840000 | 37.0904980000 |
| H | 29.3657300000 | 30.5994130000 | 35.2022620000 |
| H | 31.0907550000 | 30.8857590000 | 33.4367820000 |
| H | 33.7395730000 | 33.4159350000 | 33.4157040000 |
| H | 33.3025470000 | 31.7862910000 | 32.9639590000 |
| H | 33.0448640000 | 33.4006690000 | 31.0198500000 |
| H | 31.5132420000 | 32.6165090000 | 31.4295380000 |
| H | 31.7928130000 | 34.2761040000 | 31.9414240000 |
| H | 36.0024000000 | 34.6267430000 | 37.4078070000 |
| H | 35.0590460000 | 33.8508680000 | 36.1240670000 |
| H | 35.4477000000 | 35.5974910000 | 36.0287330000 |
| H | 34.3898240000 | 37.1983460000 | 37.8295700000 |
| H | 33.1675500000 | 36.5488960000 | 38.9433960000 |
| H | 34.8794410000 | 36.1070890000 | 39.1391010000 |
| H | 33.3700760000 | 32.9880280000 | 37.6867200000 |
| H | 34.2580920000 | 33.6768100000 | 39.0554300000 |
| H | 32.5321710000 | 34.0471720000 | 38.8506910000 |
| O | 33.8605220000 | 29.2691750000 | 34.2921440000 |
| H | 34.4946780000 | 28.8130570000 | 33.7209420000 |
| H | 32.9982390000 | 28.7708040000 | 34.2572110000 |
| O | 32.6593030000 | 30.2938220000 | 36.8233310000 |
| H | 32.2599600000 | 30.9625190000 | 36.2441460000 |
| H | 32.4136790000 | 30.5636890000 | 37.7532980000 |
| O | 31.4542590000 | 28.1476200000 | 34.1567820000 |
| H | 31.0433370000 | 27.3002650000 | 34.4817710000 |
| H | 30.8568370000 | 28.8479440000 | 34.4558600000 |
| O | 31.9985190000 | 30.9705330000 | 39.3516510000 |
| H | 31.2230950000 | 31.6217470000 | 39.4280790000 |
| H | 31.6751660000 | 30.1751720000 | 39.8029570000 |
| O | 30.2874290000 | 26.0233710000 | 35.3283050000 |
| H | 29.6607700000 | 25.3920860000 | 34.8856740000 |
| H | 29.7400330000 | 26.4106440000 | 36.0283930000 |

**TS-2-A_theozyme**
Charge: 1
Multiplicity: 5
B3LYP-D3(BJ)/6-31G(d)-SDD SCF energy (au): -3131.99502982
B3LYP-D3(BJ)/6-31G(d)-SDD enthalpy (au): -3130.73011182
B3LYP-D3(BJ)/6-31G(d)-SDD free energy (au): -3130.94853882

B3LYP-D3(BJ)/def2-TZVP/SMD SCF energy (au): -4273.31274133
B3LYP-D3(BJ)/def2-TZVP/SMD enthalpy (au): -4272.04782333
B3LYP-D3(BJ)/def2-TZVP/SMD free energy (au): -4272.26625033
B3LYP-D3(BJ)/def2-TZVP/SMD free energy (quasi-harmonic) (au): -4272.22883951

Cartesian coordinates

| ATOM | X | Y | Z |
|---|---|---|---|
| H | 36.6175660000 | 24.2867220000 | 40.6866880000 |
| C | 36.2408890000 | 24.5419910000 | 39.6917670000 |
| H | 37.0604330000 | 24.9554790000 | 39.0978540000 |
| C | 35.1618170000 | 25.6479650000 | 39.8214130000 |
| C | 34.9235550000 | 26.4526410000 | 38.5766730000 |
| C | 35.3973080000 | 27.6800280000 | 38.1807320000 |
| N | 34.1041120000 | 26.0486450000 | 37.5306340000 |
| C | 34.0931930000 | 27.0087790000 | 36.5749880000 |
| N | 34.8774740000 | 28.0183670000 | 36.9385130000 |
| H | 35.8649530000 | 23.6418980000 | 39.1973060000 |
| H | 34.2148760000 | 25.2211050000 | 40.1755380000 |
| H | 35.4798490000 | 26.3553580000 | 40.5949670000 |
| H | 33.5565750000 | 25.1988830000 | 37.5021050000 |
| H | 36.0579710000 | 28.3504790000 | 38.7078850000 |
| H | 33.4838930000 | 26.9526910000 | 35.6818400000 |
| H | 29.7028830000 | 22.7111790000 | 37.5032770000 |
| C | 29.2395430000 | 23.2255600000 | 36.6474040000 |
| H | 28.9461860000 | 24.2329230000 | 36.9437490000 |
| C | 28.0802600000 | 22.5419620000 | 35.9425570000 |
| C | 27.6724570000 | 23.3619780000 | 34.6964840000 |
| O | 28.2684840000 | 24.4190310000 | 34.3754130000 |
| O | 26.7079170000 | 22.8844650000 | 34.0302810000 |
| H | 30.0518550000 | 23.3682230000 | 35.9232390000 |
| H | 28.3348570000 | 21.5296190000 | 35.6082060000 |
| H | 27.1988250000 | 22.4426500000 | 36.5861830000 |
| H | 25.6135410000 | 30.4504430000 | 42.5542670000 |
| C | 26.4055960000 | 30.7302760000 | 43.2550480000 |
| H | 26.1172760000 | 30.4445490000 | 44.2715920000 |
| C | 26.8898180000 | 32.1702510000 | 43.0832190000 |
| C | 27.8035480000 | 32.3287930000 | 41.8630500000 |
| O | 27.8648280000 | 31.4159200000 | 40.9907890000 |
| O | 28.4957350000 | 33.3873150000 | 41.7638270000 |
| H | 27.2373510000 | 30.0596670000 | 43.0037280000 |
| H | 26.0582310000 | 32.8780670000 | 42.9740300000 |
| H | 27.4584090000 | 32.5096030000 | 43.9570440000 |
| H | 41.2540250000 | 32.9342630000 | 33.7553020000 |
| C | 40.4491780000 | 32.4364960000 | 34.3023150000 |
| H | 40.7812680000 | 32.2475690000 | 35.3275380000 |
| C | 39.9875970000 | 31.1384020000 | 33.6147690000 |
| C | 38.8004100000 | 30.4711940000 | 34.2642080000 |
| C | 37.5902800000 | 30.9277440000 | 34.7341700000 |
| N | 38.7406290000 | 29.0933340000 | 34.4457240000 |
| C | 37.5438590000 | 28.7697960000 | 34.9985520000 |
| N | 36.8249810000 | 29.8628140000 | 35.1901770000 |
| H | 39.6173360000 | 33.1486940000 | 34.3593480000 |
| H | 39.7426910000 | 31.3543490000 | 32.5662660000 |

| | | | |
|---|---|---|---|
| H | 40.8192110000 | 30.4227970000 | 33.5868290000 |
| H | 39.4648590000 | 28.4383440000 | 34.1848000000 |
| H | 37.2054210000 | 31.9330350000 | 34.7813040000 |
| H | 37.2364290000 | 27.7656520000 | 35.2478240000 |
| H | 39.3343460000 | 34.2947510000 | 39.1394480000 |
| C | 38.6576290000 | 33.8740690000 | 39.8920130000 |
| H | 39.0648550000 | 34.1441840000 | 40.8702140000 |
| C | 38.5331830000 | 32.3316130000 | 39.8152130000 |
| C | 37.2884850000 | 31.8339960000 | 39.1575540000 |
| C | 37.0779710000 | 31.0434370000 | 38.0610550000 |
| N | 36.0143740000 | 32.1105460000 | 39.6382530000 |
| C | 35.0982660000 | 31.5060020000 | 38.8431540000 |
| N | 35.7179770000 | 30.8454020000 | 37.8739620000 |
| H | 37.6874100000 | 34.3699390000 | 39.7861320000 |
| H | 39.3889970000 | 31.9029820000 | 39.2848560000 |
| H | 38.5770610000 | 31.9192090000 | 40.8323130000 |
| H | 35.7973060000 | 32.6746260000 | 40.4483100000 |
| H | 37.8065820000 | 30.6196590000 | 37.3907220000 |
| H | 34.0277450000 | 31.5515270000 | 39.0089010000 |
| O | 29.8097870000 | 32.4063410000 | 39.4389750000 |
| H | 29.8023200000 | 33.1243310000 | 40.1031510000 |
| H | 29.0560150000 | 31.8685600000 | 39.8176050000 |
| H | 33.8251850000 | 43.3260790000 | 32.3959540000 |
| C | 32.7408580000 | 43.4791760000 | 32.3963020000 |
| H | 32.3583450000 | 43.2663120000 | 33.4029670000 |
| C | 32.0162570000 | 42.5945600000 | 31.3569180000 |
| C | 31.3803800000 | 41.3201010000 | 31.9371750000 |
| C | 32.3737500000 | 40.2725470000 | 32.4519250000 |
| N | 31.6324710000 | 39.1883420000 | 33.1094630000 |
| C | 32.0325670000 | 37.9159700000 | 33.2749930000 |
| N | 33.2488670000 | 37.5080410000 | 32.8915190000 |
| N | 31.2120080000 | 37.0128920000 | 33.8241900000 |
| H | 32.5726240000 | 44.5407890000 | 32.1944700000 |
| H | 31.2022800000 | 43.1663400000 | 30.8982550000 |
| H | 32.6935130000 | 42.3300000000 | 30.5348990000 |
| H | 30.7208470000 | 41.6131990000 | 32.7679380000 |
| H | 30.7443310000 | 40.8442530000 | 31.1811000000 |
| H | 32.9365720000 | 39.8475850000 | 31.6136290000 |
| H | 33.0887890000 | 40.7225180000 | 33.1532510000 |
| H | 30.7634220000 | 39.4489620000 | 33.5560280000 |
| H | 33.9930260000 | 38.1724970000 | 32.7478180000 |
| H | 33.4667740000 | 36.5124240000 | 33.0463560000 |
| H | 30.2172050000 | 37.1791680000 | 33.8604060000 |
| H | 31.5834850000 | 36.0465240000 | 33.9124680000 |
| Fe | 34.8100700000 | 30.0636820000 | 36.0848860000 |
| N | 33.5200490000 | 32.9171400000 | 32.7265850000 |
| O | 32.8888630000 | 34.9121040000 | 33.7033280000 |
| C | 31.6880400000 | 32.9082770000 | 34.2512220000 |
| C | 31.1903650000 | 33.4765170000 | 35.4275750000 |
| C | 30.3317470000 | 32.7658080000 | 36.2672230000 |
| C | 29.9726680000 | 31.4583530000 | 35.9287880000 |
| C | 30.4613780000 | 30.8762720000 | 34.7559280000 |
| C | 31.3070850000 | 31.5901760000 | 33.8880630000 |

| | | | |
|---|---|---|---|
| C | 32.7567660000 | 33.6593820000 | 33.5013160000 |
| C | 34.7318310000 | 33.3319600000 | 32.0196640000 |
| C | 35.7129670000 | 34.0139710000 | 32.9937570000 |
| C | 35.3790610000 | 32.0393980000 | 31.4919600000 |
| C | 34.3748830000 | 34.2545800000 | 30.8414050000 |
| C | 31.8452490000 | 30.9752670000 | 32.6483520000 |
| C | 31.2148640000 | 31.4222420000 | 31.3334410000 |
| F | 34.6910820000 | 31.7124280000 | 35.2993700000 |
| H | 31.5335540000 | 34.4670450000 | 35.7081020000 |
| H | 29.9778850000 | 33.1864400000 | 37.2024500000 |
| H | 29.3228850000 | 30.8921260000 | 36.5899600000 |
| H | 30.1948150000 | 29.8545710000 | 34.4972440000 |
| H | 32.9894370000 | 31.7182080000 | 32.6215010000 |
| H | 31.8891580000 | 29.8840160000 | 32.7126130000 |
| H | 31.7672950000 | 31.0189440000 | 30.4766150000 |
| H | 30.1751170000 | 31.0744170000 | 31.2524870000 |
| H | 31.2090970000 | 32.5134970000 | 31.2470770000 |
| H | 36.6928670000 | 34.1367960000 | 32.5166680000 |
| H | 35.3499170000 | 34.9943800000 | 33.3078860000 |
| H | 35.8140700000 | 33.3941210000 | 33.8881020000 |
| H | 35.6596320000 | 31.3864890000 | 32.3259780000 |
| H | 34.6865050000 | 31.4957130000 | 30.8396610000 |
| H | 36.2803330000 | 32.2704640000 | 30.9143530000 |
| H | 33.8612860000 | 35.1534280000 | 31.1933950000 |
| H | 35.2789010000 | 34.5586820000 | 30.3010510000 |
| H | 33.7123130000 | 33.7382830000 | 30.1387490000 |
| O | 33.9486550000 | 29.2042950000 | 34.3119310000 |
| H | 33.7356880000 | 29.9649330000 | 33.7390670000 |
| H | 33.1277200000 | 28.6410650000 | 34.3444250000 |
| O | 32.8116840000 | 30.1276450000 | 36.8328110000 |
| H | 32.2316960000 | 30.5709020000 | 36.1849700000 |
| H | 32.5034080000 | 30.4590760000 | 37.7307990000 |
| O | 31.9312730000 | 27.4309220000 | 34.2050960000 |
| H | 31.1100500000 | 27.0850640000 | 34.6730700000 |
| H | 31.9216350000 | 27.0032140000 | 33.3356580000 |
| O | 32.0267500000 | 31.0152120000 | 39.2161710000 |
| H | 31.1929110000 | 31.5938520000 | 39.2836290000 |
| H | 31.8447050000 | 30.2635620000 | 39.8012410000 |
| O | 29.7918160000 | 26.3798420000 | 35.3936070000 |
| H | 29.3429070000 | 25.5647490000 | 35.0252820000 |
| H | 29.0555210000 | 26.9987420000 | 35.5117070000 |

**TS-6_theozyme**
Charge: 1
Multiplicity: 5
B3LYP-D3(BJ)/6-31G(d)-SDD SCF energy (au): -3132.00372344
B3LYP-D3(BJ)/6-31G(d)-SDD enthalpy (au): -3130.73977144
B3LYP-D3(BJ)/6-31G(d)-SDD free energy (au): -3130.95690644
B3LYP-D3(BJ)/def2-TZVP/SMD SCF energy (au): -4273.31285587
B3LYP-D3(BJ)/def2-TZVP/SMD enthalpy (au): -4272.04890387
B3LYP-D3(BJ)/def2-TZVP/SMD free energy (au): -4272.26603887
B3LYP-D3(BJ)/def2-TZVP/SMD free energy (quasi-harmonic) (au): -4272.23053643

Cartesian coordinates

| ATOM | X | Y | Z |
|---|---|---|---|
| H | 36.6176230000 | 24.2867560000 | 40.6866550000 |
| C | 36.2409240000 | 24.5417860000 | 39.6919380000 |
| H | 37.0603580000 | 24.9555310000 | 39.0978090000 |
| C | 35.1335020000 | 25.6187170000 | 39.8176060000 |
| C | 34.8526960000 | 26.4160480000 | 38.5747350000 |
| C | 35.3870720000 | 27.6014140000 | 38.1250200000 |
| N | 33.9059580000 | 26.0843540000 | 37.6132770000 |
| C | 33.8889580000 | 27.0437500000 | 36.6576860000 |
| N | 34.7855570000 | 27.9853720000 | 36.9356750000 |
| H | 35.8814830000 | 23.6349430000 | 39.1966870000 |
| H | 34.2040790000 | 25.1635980000 | 40.1819030000 |
| H | 35.4362830000 | 26.3365590000 | 40.5884010000 |
| H | 33.2783240000 | 25.2899400000 | 37.6180490000 |
| H | 36.1446870000 | 28.2157210000 | 38.5878090000 |
| H | 33.1998410000 | 27.0223540000 | 35.8269910000 |
| H | 29.7025730000 | 22.7110700000 | 37.5033840000 |
| C | 29.2400130000 | 23.2259190000 | 36.6470920000 |
| H | 28.9461660000 | 24.2328070000 | 36.9439730000 |
| C | 28.2219370000 | 22.6546690000 | 35.6906950000 |
| C | 28.0881040000 | 23.5773370000 | 34.4465520000 |
| O | 28.8810100000 | 24.5967950000 | 34.2487680000 |
| O | 27.2019670000 | 23.2940740000 | 33.6400950000 |
| H | 30.1362700000 | 23.4331930000 | 36.0458820000 |
| H | 28.5095830000 | 21.6655300000 | 35.3159320000 |
| H | 27.2198950000 | 22.5477680000 | 36.1178560000 |
| H | 25.6136260000 | 30.4503430000 | 42.5541740000 |
| C | 26.4050520000 | 30.7305820000 | 43.2551680000 |
| H | 26.1176190000 | 30.4443690000 | 44.2716140000 |
| C | 26.8985150000 | 32.1675620000 | 43.0781490000 |
| C | 27.8238410000 | 32.3194320000 | 41.8650260000 |
| O | 27.8879910000 | 31.4102260000 | 40.9892550000 |
| O | 28.5253410000 | 33.3728350000 | 41.7719190000 |
| H | 27.2342110000 | 30.0558660000 | 43.0047670000 |
| H | 26.0714180000 | 32.8789460000 | 42.9583630000 |
| H | 27.4614960000 | 32.5090420000 | 43.9547530000 |
| H | 41.2540120000 | 32.9343590000 | 33.7554210000 |
| C | 40.4492780000 | 32.4363820000 | 34.3021410000 |
| H | 40.7811990000 | 32.2475180000 | 35.3275580000 |
| C | 39.9500440000 | 31.1484890000 | 33.6151170000 |
| C | 38.7763930000 | 30.4744160000 | 34.2882690000 |
| C | 37.4529190000 | 30.8164080000 | 34.4687790000 |
| N | 38.8791080000 | 29.2065910000 | 34.8521670000 |
| C | 37.6636890000 | 28.8337140000 | 35.3308110000 |
| N | 36.7763470000 | 29.7913460000 | 35.1156370000 |
| H | 39.6246430000 | 33.1565420000 | 34.3712540000 |
| H | 39.6692920000 | 31.3836220000 | 32.5814030000 |
| H | 40.7720040000 | 30.4252430000 | 33.5427930000 |
| H | 39.7175560000 | 28.6423860000 | 34.8694820000 |
| H | 36.9238550000 | 31.7183630000 | 34.2029470000 |
| H | 37.4665930000 | 27.8969660000 | 35.8305950000 |
| H | 39.3343860000 | 34.2947850000 | 39.1395590000 |

| | | | |
|---|---|---|---|
| C | 38.6576720000 | 33.8739960000 | 39.8917060000 |
| H | 39.0647670000 | 34.1441690000 | 40.8703150000 |
| C | 38.5555640000 | 32.3272700000 | 39.7821870000 |
| C | 37.3095650000 | 31.7912660000 | 39.1481470000 |
| C | 37.0586010000 | 31.2252730000 | 37.9244490000 |
| N | 36.0902280000 | 31.7274430000 | 39.8136370000 |
| C | 35.1731370000 | 31.1390090000 | 39.0012340000 |
| N | 35.7347570000 | 30.8216540000 | 37.8445900000 |
| H | 37.6804680000 | 34.3557210000 | 39.7863970000 |
| H | 39.4095460000 | 31.9380220000 | 39.2190970000 |
| H | 38.6418900000 | 31.8891640000 | 40.7851610000 |
| H | 35.9142260000 | 32.0469670000 | 40.7561540000 |
| H | 37.7355030000 | 31.0962410000 | 37.0956600000 |
| H | 34.1366360000 | 30.9891940000 | 39.2775320000 |
| O | 29.8604480000 | 32.4048480000 | 39.4456930000 |
| H | 29.8423280000 | 33.1124760000 | 40.1212520000 |
| H | 29.0936450000 | 31.8734020000 | 39.8045830000 |
| H | 33.8251060000 | 43.3258300000 | 32.3960420000 |
| C | 32.7408190000 | 43.4797310000 | 32.3961920000 |
| H | 32.3584110000 | 43.2660890000 | 33.4029580000 |
| C | 32.0082650000 | 42.6361890000 | 31.3446280000 |
| C | 31.8793280000 | 41.1339720000 | 31.6504070000 |
| C | 33.2151060000 | 40.3727080000 | 31.6838600000 |
| N | 33.0573530000 | 38.9299870000 | 31.4408210000 |
| C | 32.9093680000 | 37.9513560000 | 32.3483960000 |
| N | 32.8993420000 | 38.1953630000 | 33.6644230000 |
| N | 32.7916130000 | 36.6818670000 | 31.9387370000 |
| H | 32.5773480000 | 44.5449000000 | 32.2044440000 |
| H | 30.9910030000 | 43.0271860000 | 31.2266430000 |
| H | 32.4933690000 | 42.7586310000 | 30.3664830000 |
| H | 31.3439670000 | 40.9857580000 | 32.5984180000 |
| H | 31.2535640000 | 40.6779860000 | 30.8723560000 |
| H | 33.8792350000 | 40.7429320000 | 30.8959020000 |
| H | 33.7539790000 | 40.5067640000 | 32.6260730000 |
| H | 32.8654090000 | 38.6682000000 | 30.4827940000 |
| H | 32.6840220000 | 39.1156560000 | 34.0159660000 |
| H | 32.8132600000 | 37.3857400000 | 34.2885330000 |
| H | 33.0406210000 | 36.4106410000 | 30.9996160000 |
| H | 32.6453440000 | 35.9503470000 | 32.6400950000 |
| Fe | 34.7281940000 | 30.0541860000 | 36.0302360000 |
| N | 32.2557040000 | 35.2366090000 | 36.6234900000 |
| O | 32.5563640000 | 35.4791830000 | 34.4101840000 |
| C | 31.5192680000 | 33.4280490000 | 35.1883900000 |
| C | 30.5740740000 | 33.0463530000 | 36.1636210000 |
| C | 29.9514760000 | 31.8050610000 | 36.1070530000 |
| C | 30.2887320000 | 30.9167480000 | 35.0796740000 |
| C | 31.2456680000 | 31.2836290000 | 34.1313130000 |
| C | 31.8759390000 | 32.5335750000 | 34.1516470000 |
| C | 32.1632540000 | 34.7503290000 | 35.3475460000 |
| C | 33.4357060000 | 35.0127580000 | 37.4654030000 |
| C | 34.7096600000 | 34.7270870000 | 36.6537750000 |
| C | 33.5941640000 | 36.2612000000 | 38.3470750000 |
| C | 33.0936470000 | 33.7875590000 | 38.3478990000 |

| | | | |
|---|---|---|---|
| C | 32.9017700000 | 32.8283240000 | 33.0830030000 |
| C | 32.2365430000 | 33.2742930000 | 31.7700580000 |
| F | 34.6951190000 | 31.7553150000 | 35.2843310000 |
| H | 30.3229730000 | 33.7266050000 | 36.9688400000 |
| H | 29.2315340000 | 31.5378990000 | 36.8725120000 |
| H | 29.8586490000 | 29.9211760000 | 35.0295930000 |
| H | 31.5418320000 | 30.5633300000 | 33.3765210000 |
| H | 33.6102700000 | 33.5821240000 | 33.4275140000 |
| H | 33.4785860000 | 31.9145090000 | 32.9140540000 |
| H | 32.9911860000 | 33.5179060000 | 31.0112710000 |
| H | 31.6009170000 | 32.4815340000 | 31.3618740000 |
| H | 31.5930440000 | 34.1478580000 | 31.9257250000 |
| H | 35.5605270000 | 34.6412680000 | 37.3389500000 |
| H | 34.6314070000 | 33.7845860000 | 36.1021090000 |
| H | 34.9183540000 | 35.5424240000 | 35.9527200000 |
| H | 33.8314370000 | 37.1389770000 | 37.7364340000 |
| H | 32.6731050000 | 36.4631430000 | 38.9013910000 |
| H | 34.4103930000 | 36.1116800000 | 39.0616270000 |
| H | 33.0107220000 | 32.8835250000 | 37.7430300000 |
| H | 33.9073440000 | 33.6507780000 | 39.0670050000 |
| H | 32.1592720000 | 33.9381280000 | 38.8933930000 |
| O | 33.7418310000 | 29.1035980000 | 34.3701980000 |
| H | 34.3553110000 | 28.6038360000 | 33.8116980000 |
| H | 32.8574190000 | 28.5731980000 | 34.4022670000 |
| O | 32.7802080000 | 30.1667180000 | 36.8930920000 |
| H | 32.1323390000 | 30.5084980000 | 36.2528890000 |
| H | 32.4658930000 | 30.4442420000 | 37.8026270000 |
| O | 31.5103500000 | 27.7989930000 | 34.3368150000 |
| H | 31.1251040000 | 26.9002580000 | 34.8574940000 |
| H | 31.2750680000 | 27.6392270000 | 33.4096990000 |
| O | 31.9860650000 | 30.8377730000 | 39.3496180000 |
| H | 31.2097450000 | 31.4881370000 | 39.4023910000 |
| H | 31.6581570000 | 30.0365590000 | 39.7871390000 |
| O | 30.6690410000 | 25.7680100000 | 35.4991500000 |
| H | 29.7047490000 | 25.0334130000 | 34.8928510000 |
| H | 30.1831120000 | 26.1321390000 | 36.2566010000 |

#### XIV. $^{1}H$ and $^{13}C$ NMR Spectra of Compounds

**[1]H NMR Spectrum of 1 (400 MHz, CDCl₃)**

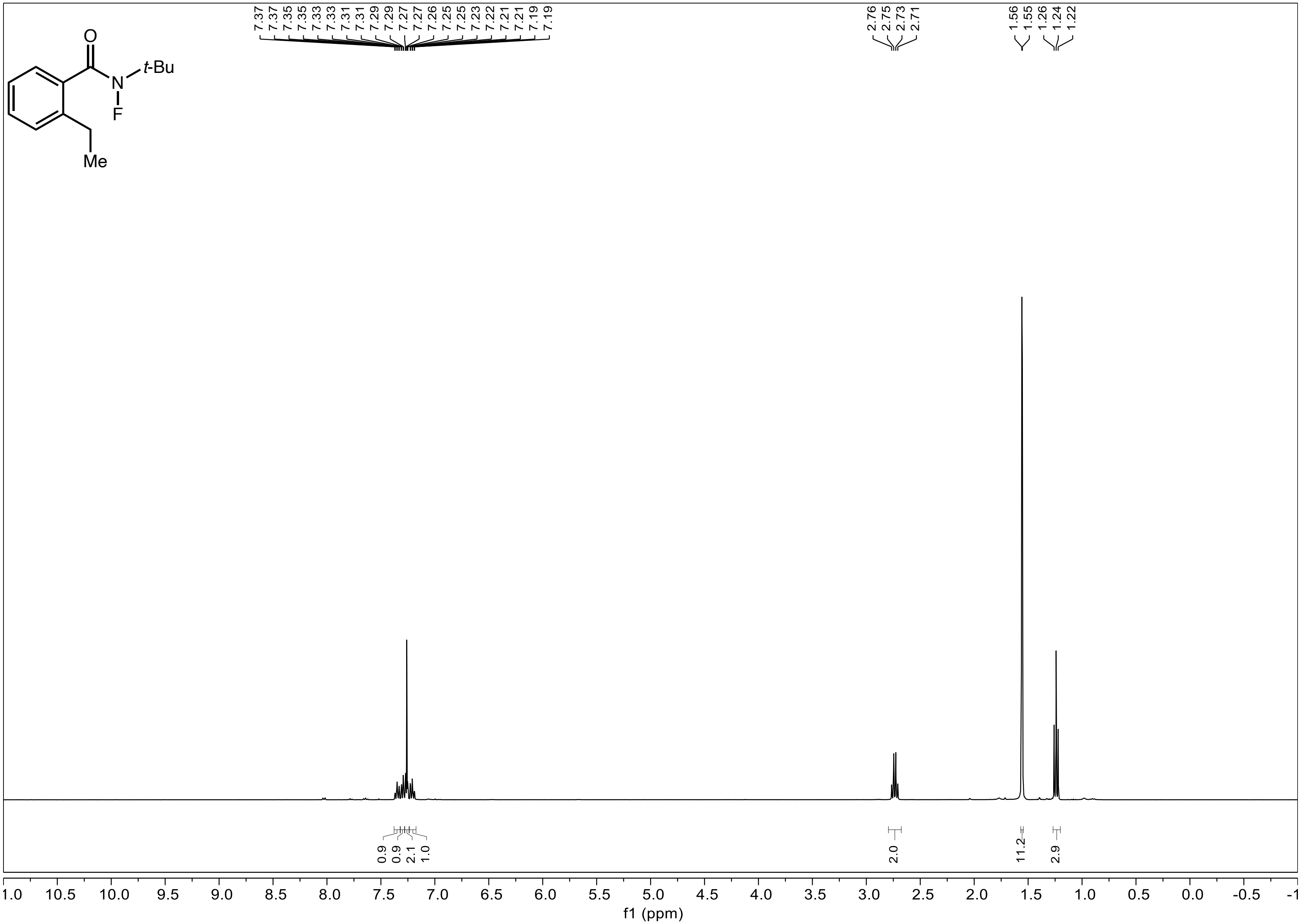

**[13]C NMR Spectrum of 1 (101 MHz, CDCl$_3$)**

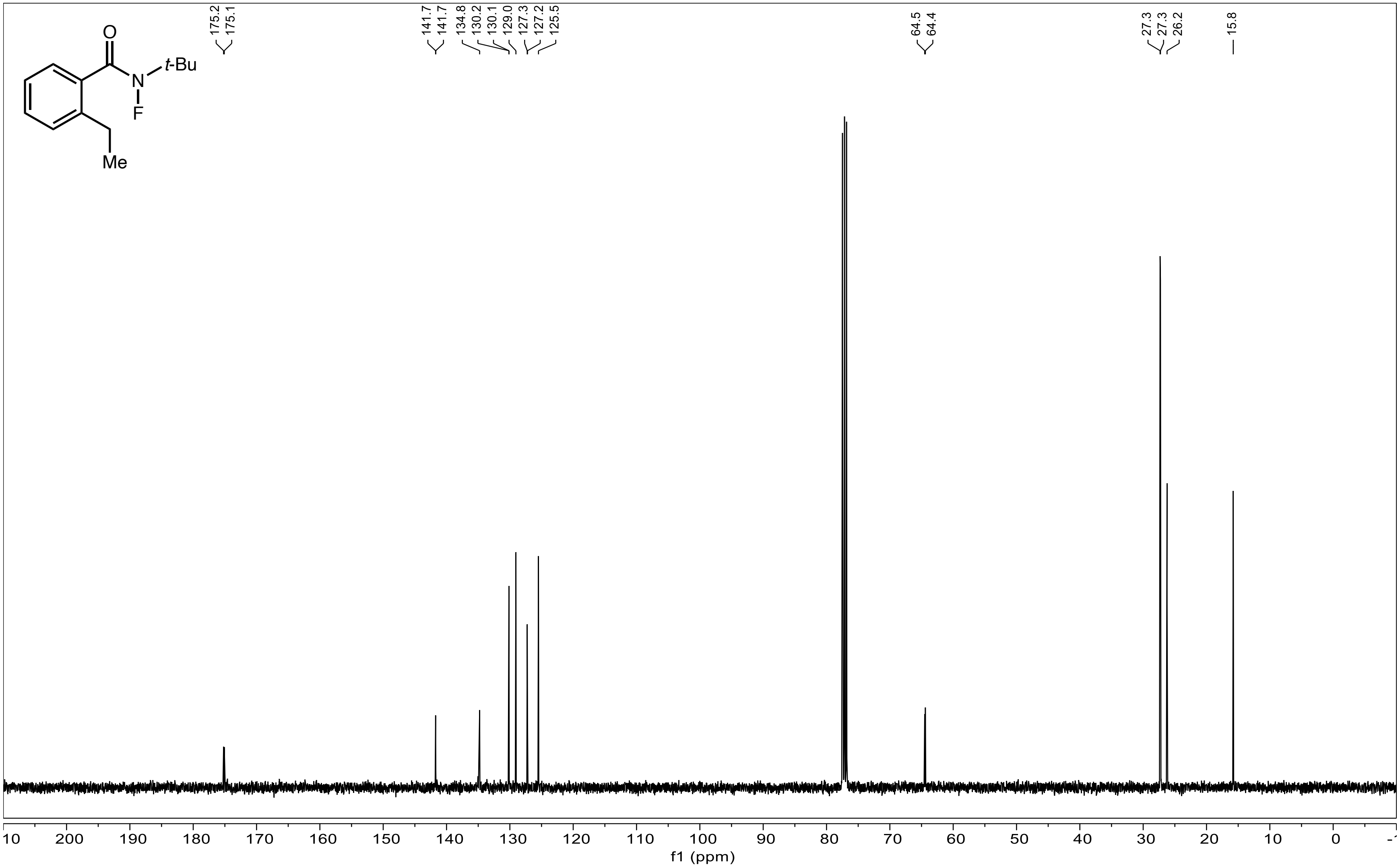

#### $^{19}$F NMR Spectrum of 1 (376 MHz, $CDCl_3$)

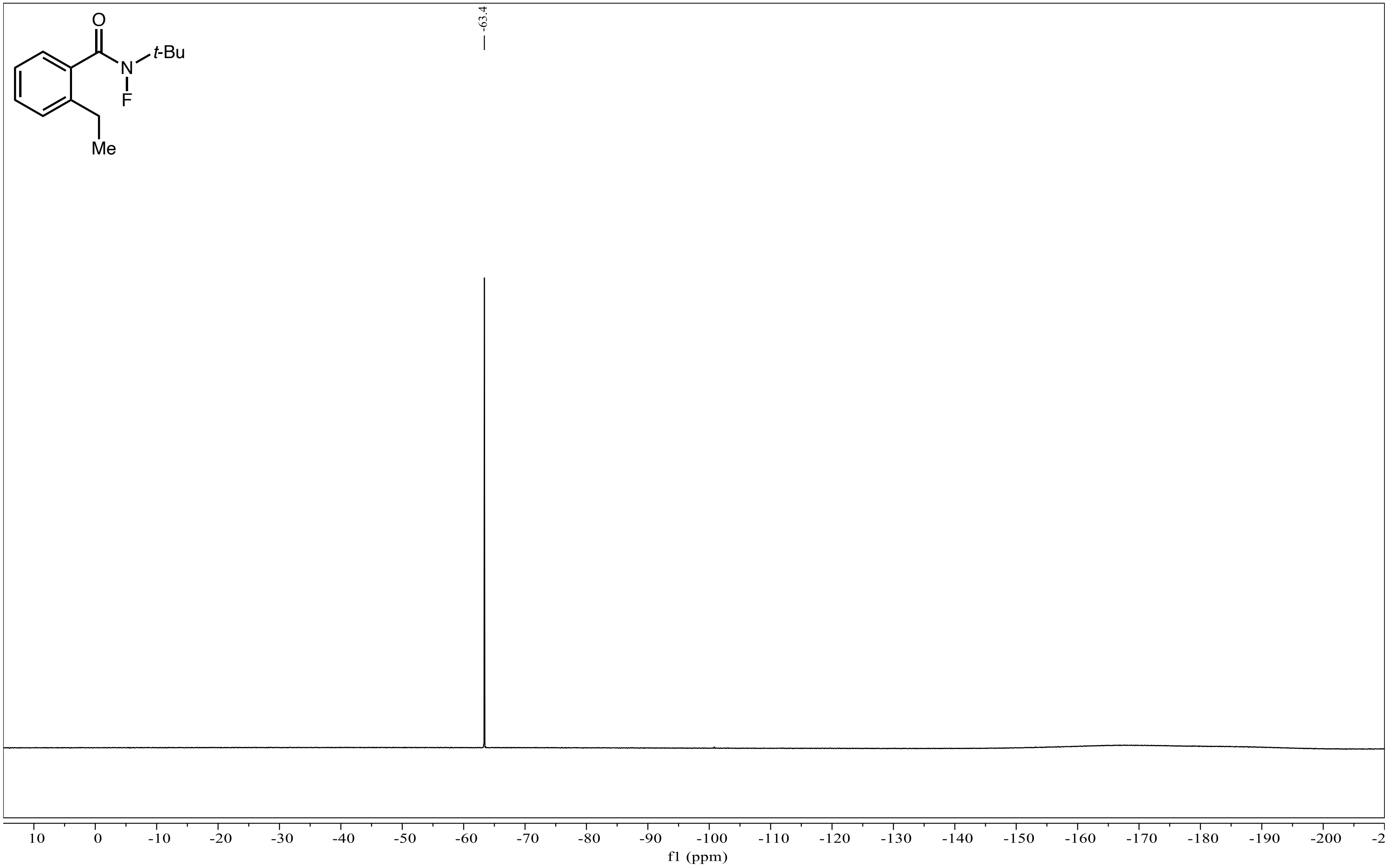

**[1]H NMR Spectrum of 2 (400 MHz, CDCl₃)**

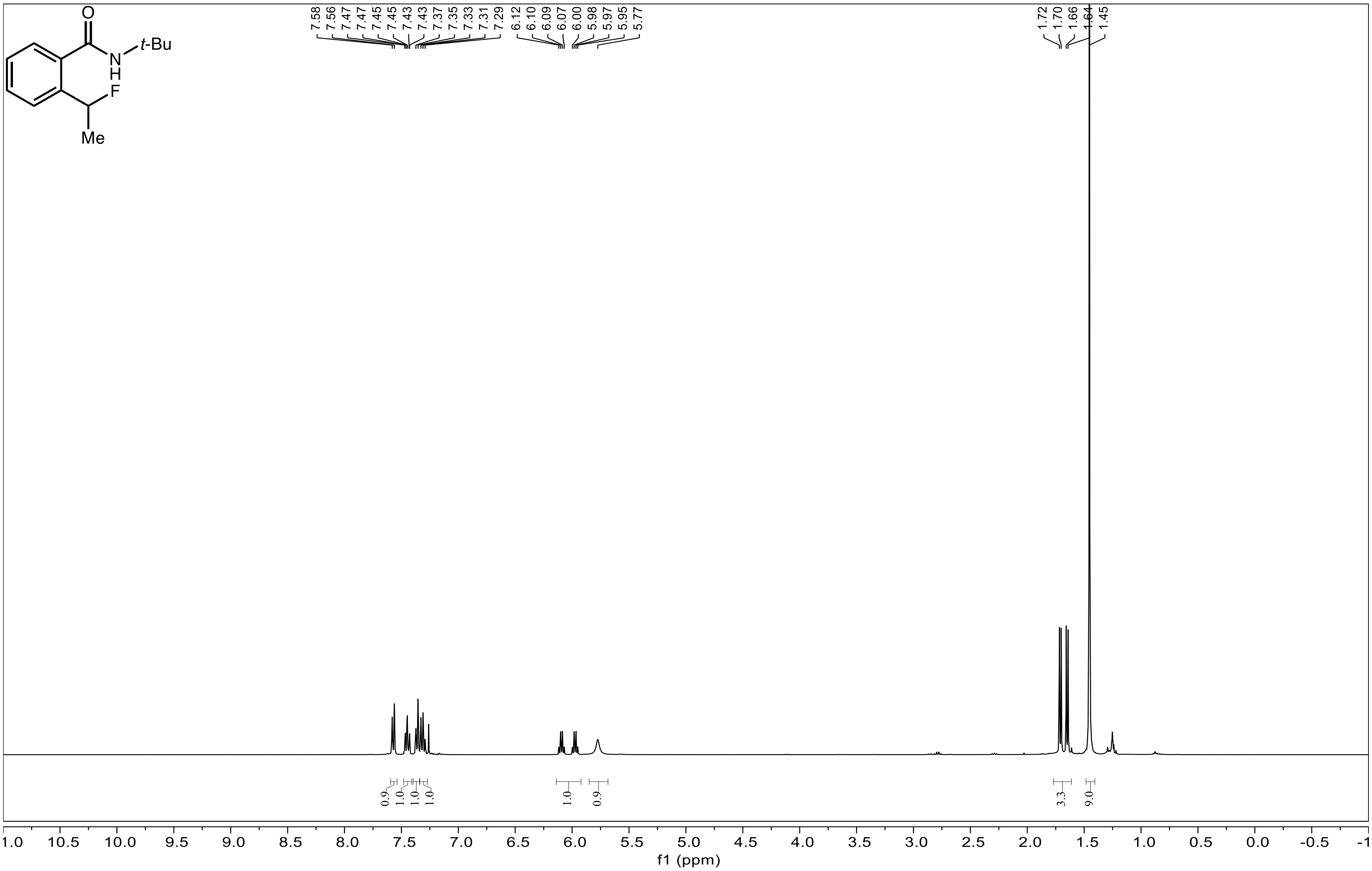

**[13]C NMR Spectrum of 2 (101 MHz, CDCl$_3$)**

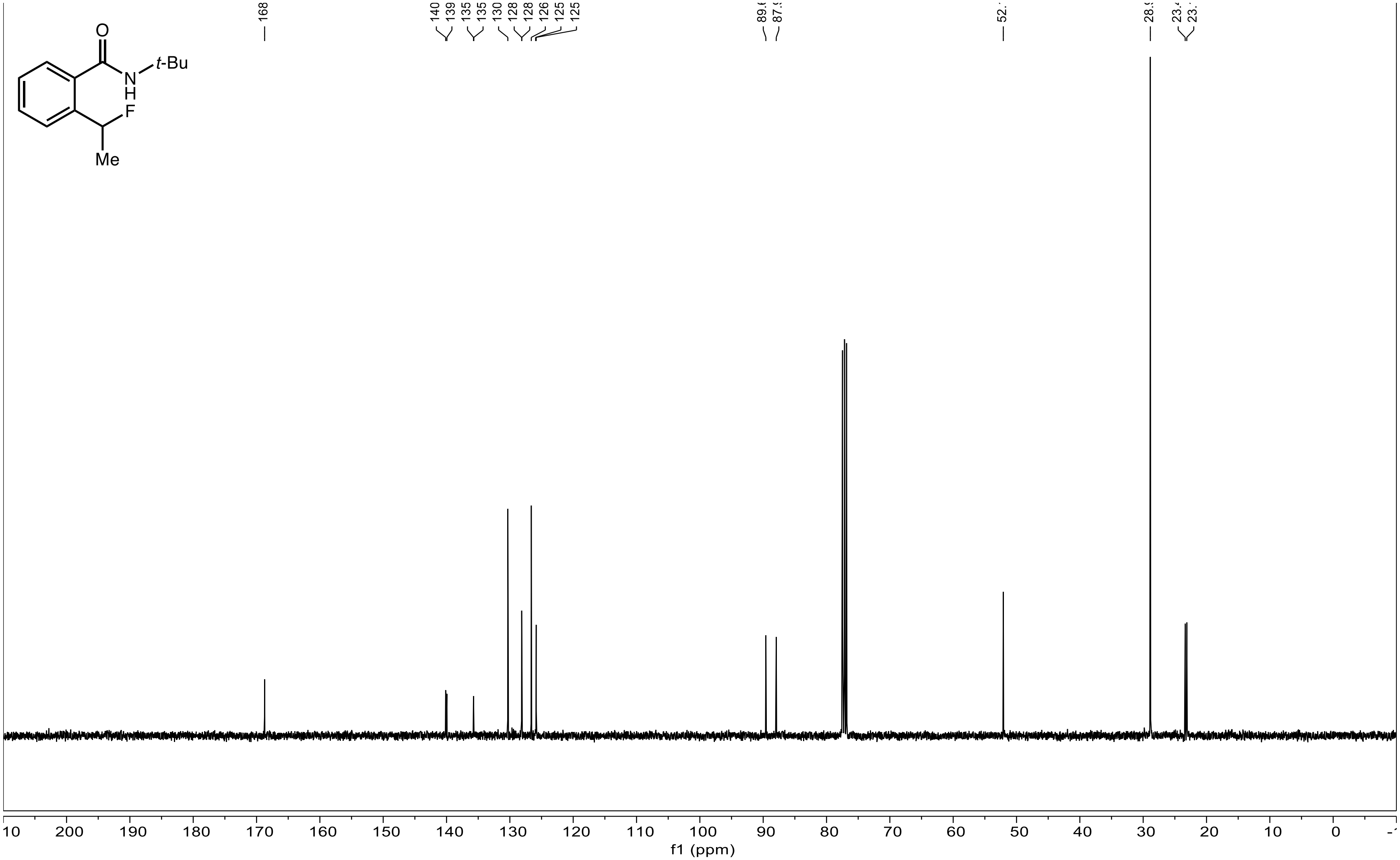

**[19]F NMR Spectrum of 2 (376 MHz, $CDCl_3$)**

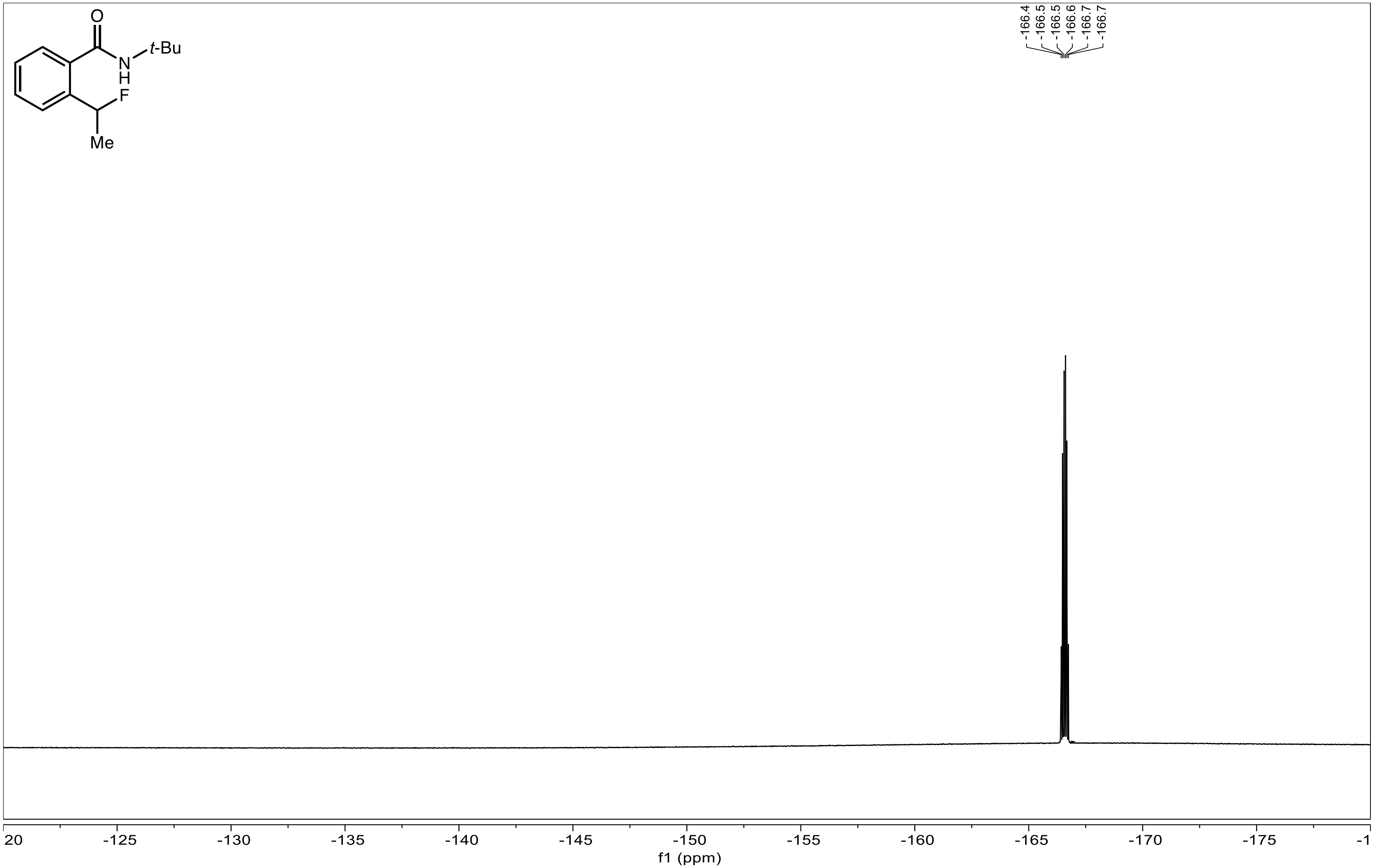

**[1]H NMR Spectrum of 3 (400 MHz, CDCl$_3$)**

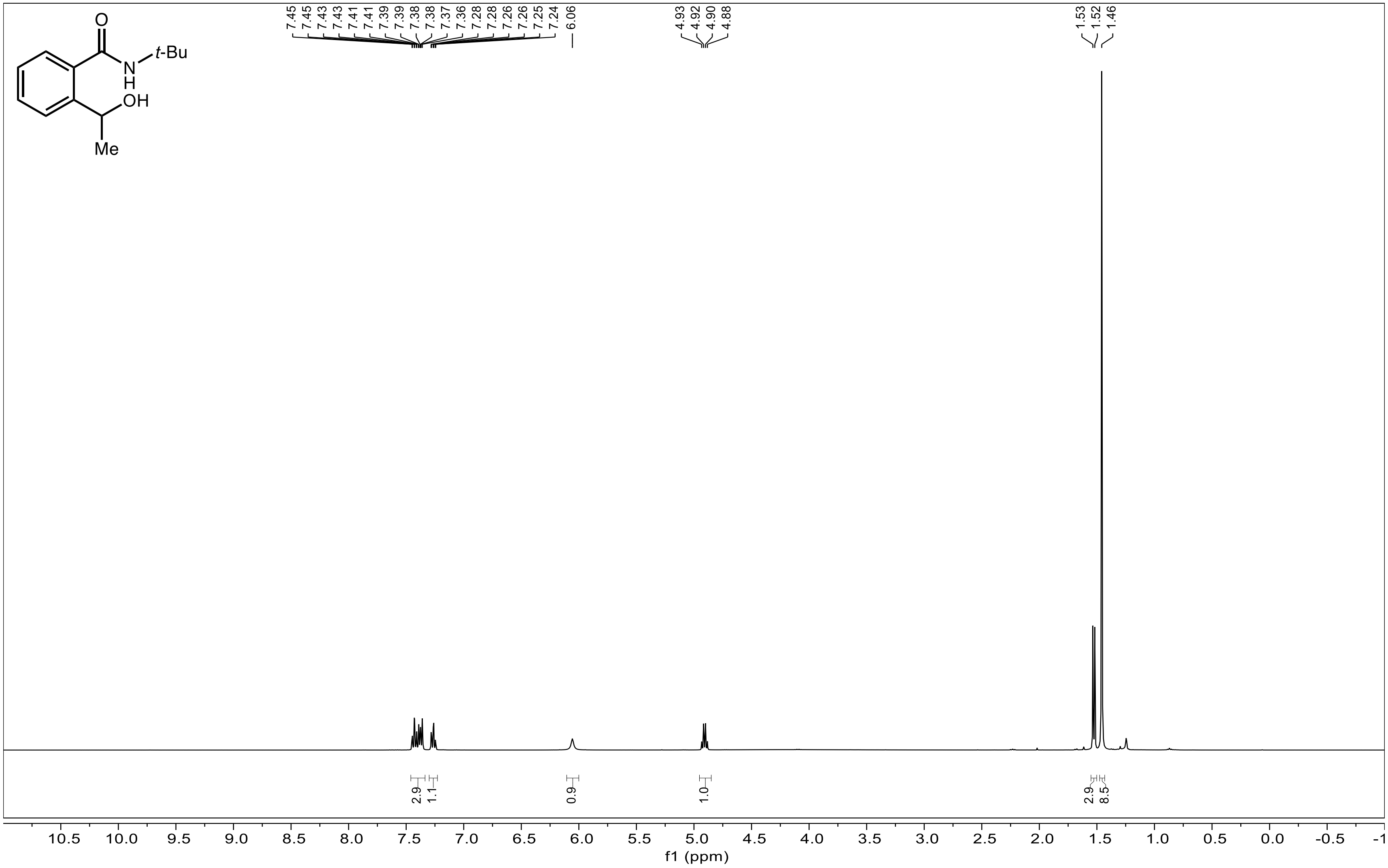

**[13]C NMR Spectrum of 3 (101 MHz, CDCl$_3$)**

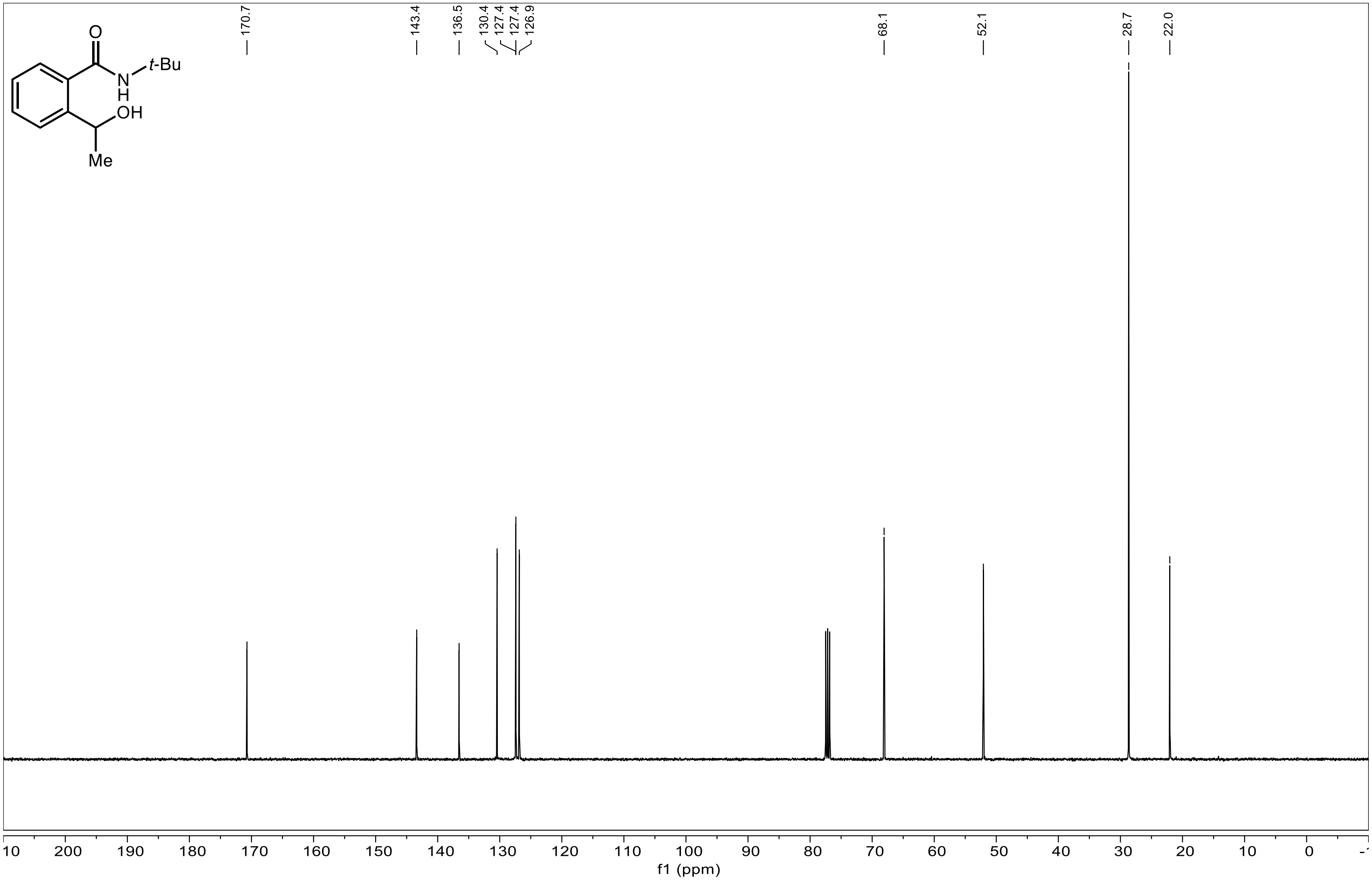

### $^{1}H$ NMR Spectrum of 4 (400 MHz, $CDCl_3$)

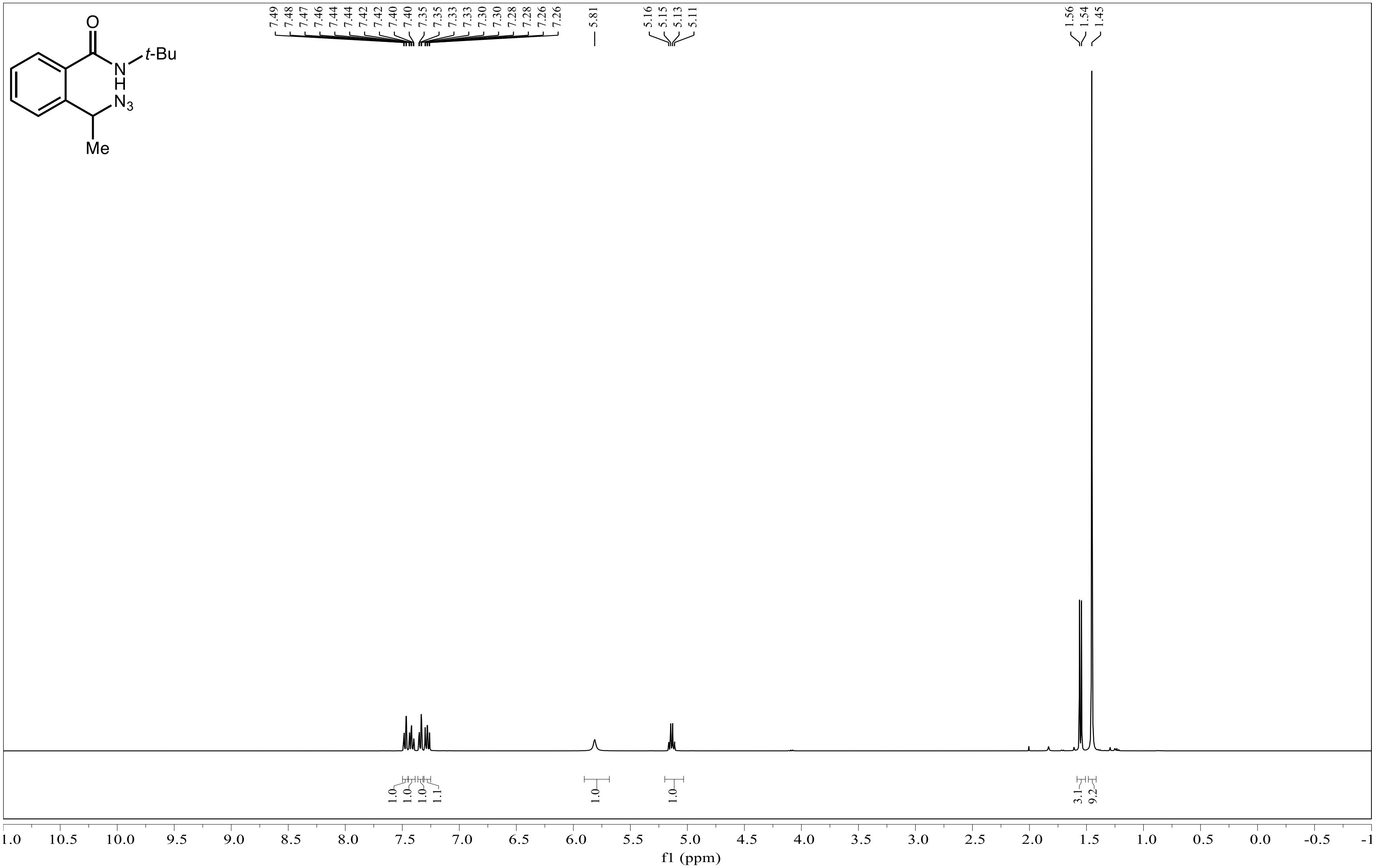

**[13]C NMR Spectrum of 4 (101 MHz, $CDCl_3$)**

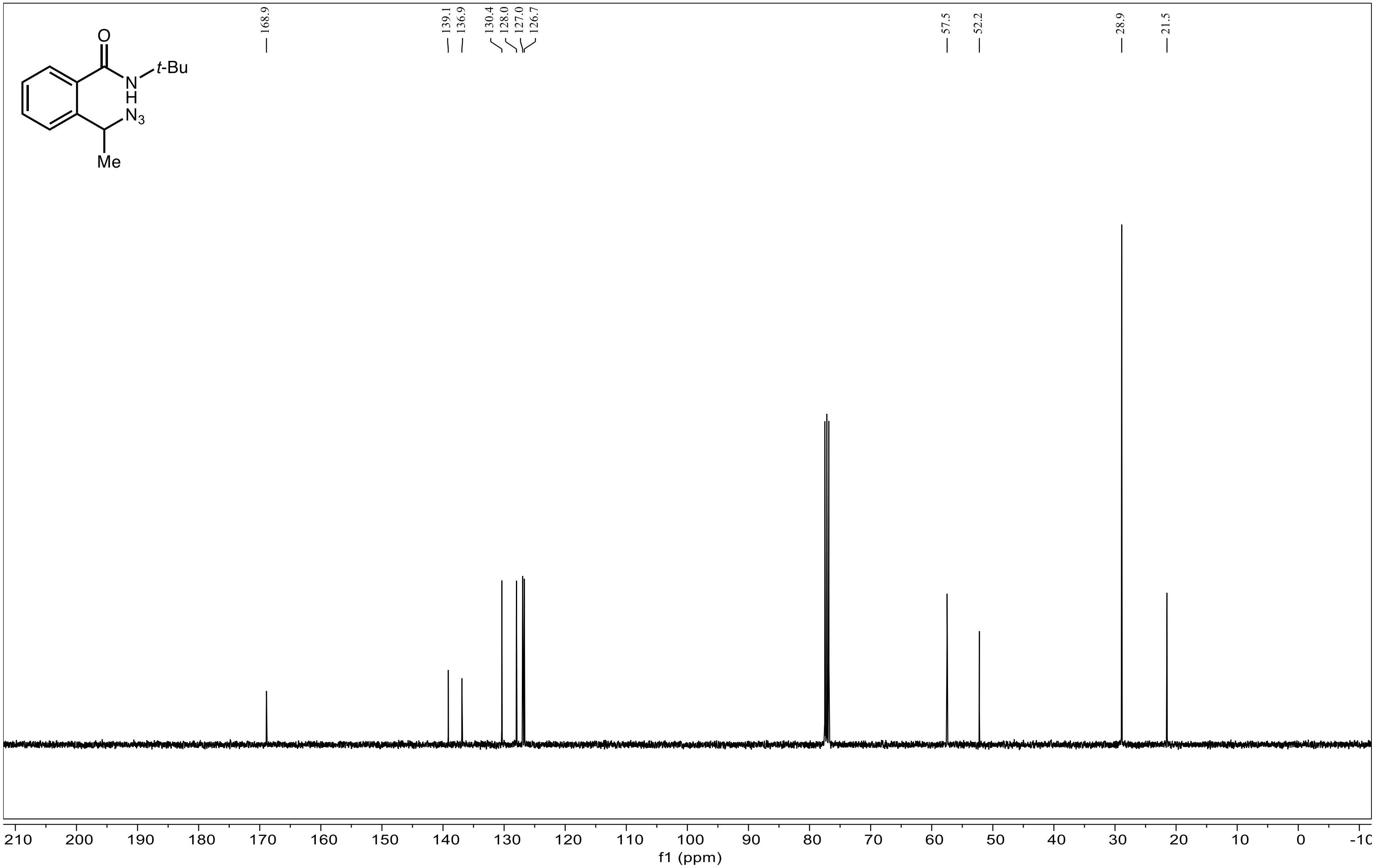

**[1]H NMR Spectrum of ethyl 2-(1-azidoethyl)benzoate (400 MHz, $CDCl_3$)**

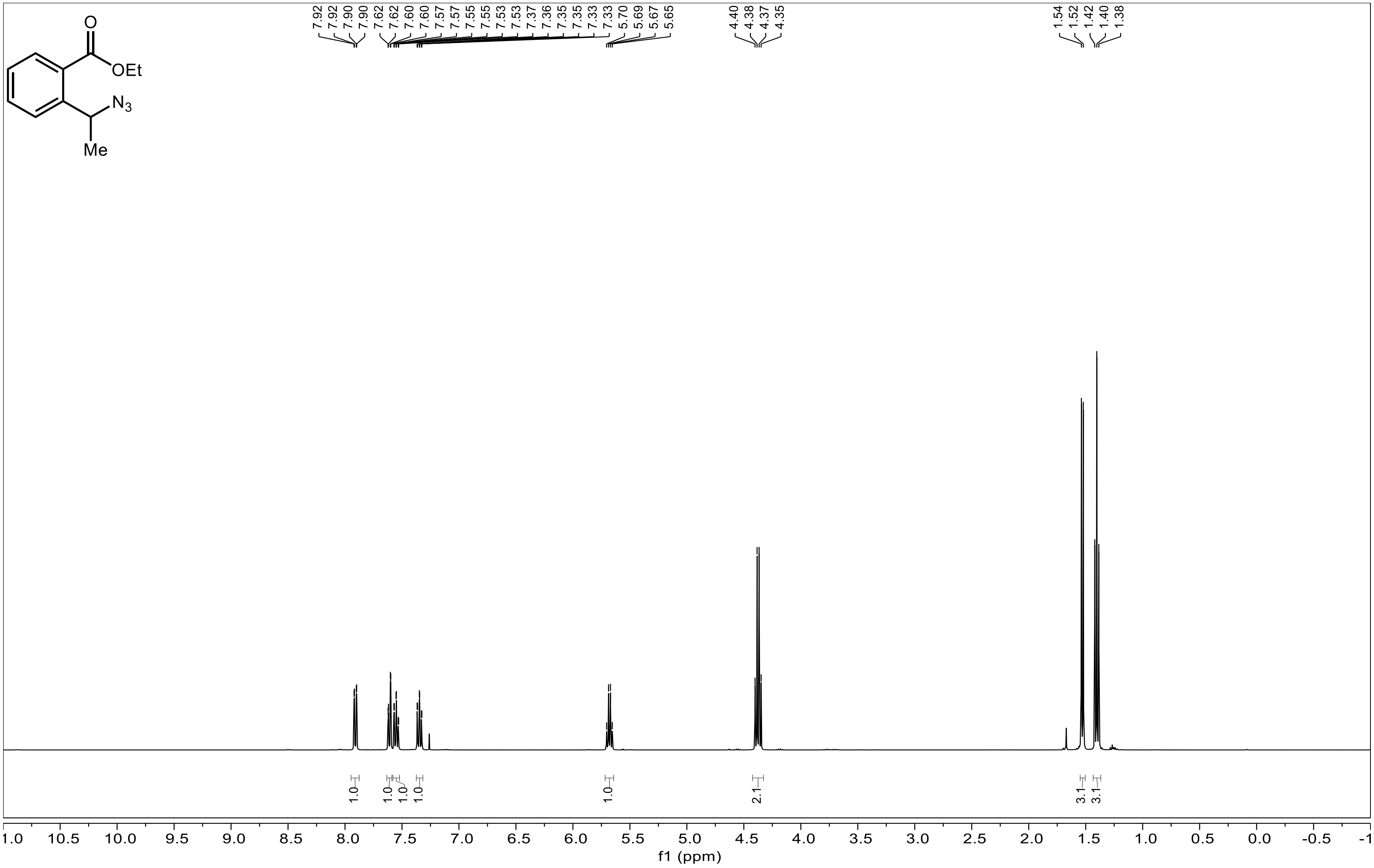

**[13]C NMR Spectrum of ethyl 2-(1-azidoethyl)benzoate (101 MHz, $CDCl_3$)**

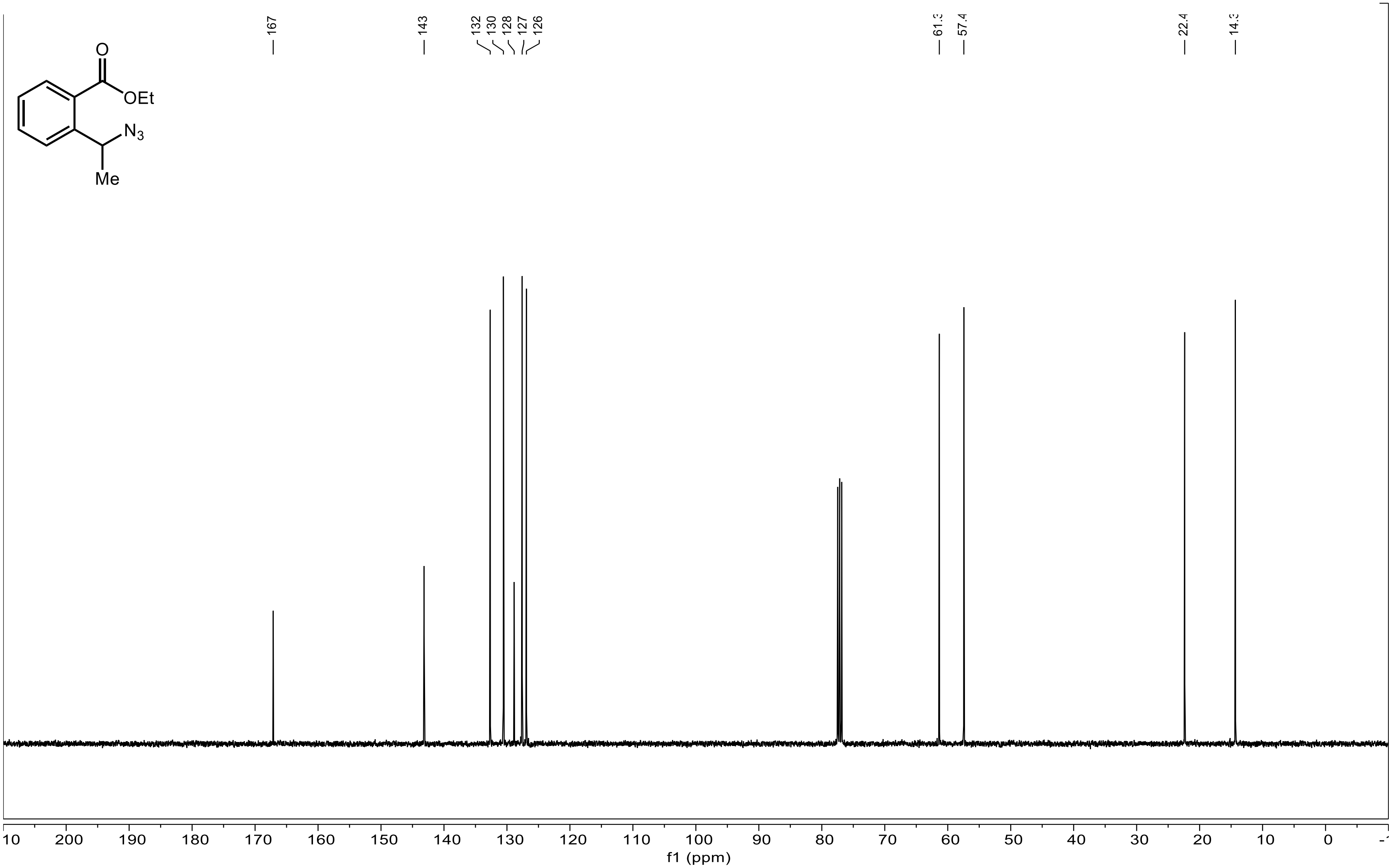
